# Incorporating prior information into signal-detection analyses across biologically informed gene-sets

**DOI:** 10.1101/525840

**Authors:** Mengqi Zhang, Sahar Gelfman, Janice McCarthy, David B. Goldstein, Andrew S. Allen

## Abstract

Signal detection analyses are used to assess whether there is any evidence of signal within a large collection of hypotheses. For example, we may wish to assess whether there is any evidence of association with disease among a set of biologically related genes. Such an analysis typically treats all genes within the sets similarly, even though there is substantial information concerning the likely importance of each gene within each set. For example, deleterious variants within genes that show evidence of purifying selection are more likely to substantially affect the phenotype than genes that are not under purifying selection, at least for traits that are themselves subject to purifying selection. Here we improve such analyses by incorporating prior information into a higher-criticism-based signal detection analysis. We show that when this prior information is predictive of whether a gene is associated with disease, our approach can lead to a significant increase in power. We illustrate our approach with a gene-set analysis of amyotrophic lateral sclerosis (ALS), which implicates a number of gene-sets containing *SOD1* and *NEK1* as well as showing enrichment of small p-values for gene-sets containing known ALS genes.

## INTRODUCTION

High-throughput sequencing (HTS) studies, both whole exome studies (WES) and, more recently, whole genome studies (WGS), have emerged as the primary approach to identifying genetic variation that may be associated with disease. Unlike genome-wide association studies which depend on linkage disequilibrium between tag SNPs and pathogenic variation, WES and WGS studies are able to assay pathogenic variation directly, and as a result, are able to directly interrogate the role of rare variation in disease. When the disease phenotype impacts the fitness of an individual, variants with a large effect on the phenotype will tend to be rare, as they will tend to be pruned out of the population before reaching appreciable frequency by purifying selection. This has been seen empirically (Park et al., 2011), and rare-variant focused disease mapping studies have successfully implicated a number of newly discovered disease genes, including studies in predominantly later onset diseases such as amyotrophic lateral sclerosis (ALS, Poppe et al, 2014; Cirulli et al. 2015) and idiopathic pulmonary fibrosis (IPF, Palmer et al, 2018). As each individual variant is rare, single-variant analyses are likely underpowered for detecting disease association. Therefore, rare-variant analyses usually integrate the effects of many variants across a gene or other genetic region. In these analyses, variants are often filtered by surrogates of the variant’s likely deleteriousness including the variant’s frequency in the general population as well as annotations of the variant’s likely functional impact on the ultimate protein product. Further, when such an analysis is restricted to rare variation, a gene that demonstrates an excess of deleterious variants in cases over controls provides strong evidence for the direct involvement of that gene in disease etiology as any indirect effects such as linkage disequilibrium are minimized.

A primary goal of such studies is *signal identification*, i.e., identifying individual genes that show significant differences in the burden of qualifying variants between cases and controls. For example, in the whole exome ALS sequencing study (Gelfman et al. 2018), the exomes of 18536 genes were sequenced across 3093 ALS cases and 8186 neurologically normal controls and tested for differences in the burden of rare variation, identifying *SOD1* [MIM: 147450], *NEK1*[MIM: 604588], and *TARDBP*[MIM:605078] as being involved in ALS (Gelfman et al. 2018). Though these findings are undoubtedly an advance for ALS genetics, there is likely signal in this dataset that is below the identification threshold and, thus, remains latent. Our goal here is to enhance our understanding of disease etiology by uncovering this latent signal.

An alternate to *signal identification* is *signal detection*, i.e., the detection of whether any genes within a set of genes participating in a biologic process show significant differences in the burden of qualifying variants between cases and controls. Signal detection does not attempt to identify which genes in the set are non-null, just that there exists some subset of the genes within the gene-set that show signal. As a result, the signal detection problem can be addressed by a goodness of fit test that assesses whether the distribution of *p*-values, for the individual gene-level tests within the gene-set, follow a uniform distribution (i.e., all tests in the gene-set are null). Rejecting such a test implies that at least some of the genes in the gene-set have differences in the burden of qualifying variants between cases and controls. If the gene sets are chosen carefully, signal detection within the set can provide mechanistic insight into disease etiology, for example by emphasizing the importance of specific pathways or otherwise related gene sets. Higher Criticism (HC) is one such approach. Donoho and Jin (Donoho & Jin, 2004) showed that the HC statistic obtains the optimal detection limit for “rare-weak” alternatives where a small proportion of hypotheses weakly deviate from the null.

A limitation of such an approach is that it treats all genes within the focal gene-set equally, even though there is substantial external prior information about the relative importance of a gene within a gene-set. Genic intolerance (Petrovski et al, 2013), network centrality (Barabási & Albert, 1999; White & Smyth, 2003) and gene expression in disease relevant tissues are examples of sources of prior information that can be used to quantify the relative importance of genes within gene-sets. We discuss these sources of prior information in detail in the materials and methods section below.

Here we develop a novel HC statistic that incorporates prior information concerning the relative importance of genes within a gene-set into the analysis. We develop an asymptotic theory for the null distribution of our statistics and describe permutation procedures that can be used when the asymptotic approximation is likely to be poor. We conduct extensive simulation studies and show that when the prior information is correlated with those genes that are involved in disease our approach leads to a substantial increase in power. We illustrate our approach with a gene-set analysis of a whole exome sequencing study of amyotrophic lateral sclerosis.

## MATERIALS AND METHODS

### Overview of our approach

We develop a self-contained gene-set test based on a higher criticism statistic that explicitly weights the contribution of each gene in the set by prior information reflecting a genes likelihood of being involved in disease pathology. The higher criticism statistic our approach is based on can be thought of as a goodness of fit test. Specifically, consider a set of *n* statistics, *X*_1_, *X*_2_, …, *X*_*n*_ and empirical distribution function 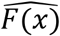. In our application, these statistics represent tests of gene/phenotype association for each of the *n* genes in a gene-set selected on the basis of biological relationships. The higher criticism test assesses whether the observed distribution, 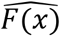, is consistent with *F*_0_(*x*), the distribution of the test statistics under the global null. In most cases, the statistics we are working with are the *p*-values associated with each gene in the set. Therefore, as these *p*-values will be uniform under the null, we have that *F*_0_ (*x*) = *x* and we could assess whether the observed distribution of *p*-values is consistent with the global null using the Kolomorgorov-Smirnov (KS) test statistic,

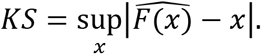

However, Donoho and Jin (2004) showed that the power to detect small shifts in the distribution function due to a small number of weak signals can be improved by scaling the KS test by 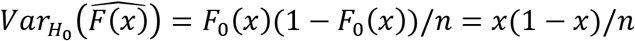 and restricting the domain over which the supremum is taken to the tail. These modifications of the KS test give rise to the higher-criticism (HC) test,

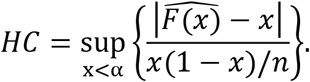

#### Incorporating prior information into higher-criticism statistics

To incorporate prior information into the HC framework, we assume that for the *i*^*th*^ gene (*i*^*th*^ Hypothesis being tested) there is affiliated a weight, *w*_*i*_ ≥ 0, such that *w*_*i*_ quantifies the relative importance of a gene within the gene-set. Let *w* = (*w*_1_, *w*_2_, …, *w*_*n*_). Let *x*_*i*_ be *i*^*th*^ gene’s *p*-value and let 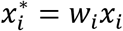. Define

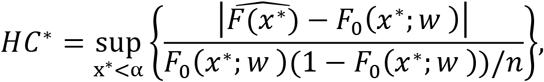

where 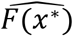 is the empirical distribution function of the *x*\*s and *F*_0_ (*x*\*) is cumulative distribution function of *x*\* under the global null hypothesis. We can show that 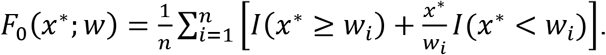.

It is not difficult to see that *HC*\* is of the same form as the unweighted HC statistic studied by (Jaeschke, 1979) which was shown to converge in distribution to the Gumbel distribution as *n* goes to infinity. However, as noted by Barrett and Lin (2014), this convergence is extremely slow and unlikely to yield a good approximation in most cases. As a result, we use permutation to approximate the null distribution of *HC*\*. When testing across a large number of gene-sets, we use the algorithm proposed by Ge, Dudoit, & Speed (2003) to account for testing multiple, potentially correlated, hypotheses.

### Choosing the weights

The weights summarize prior information concerning the relative importance of a gene within a gene-set. We consider three main sources of this information: 1) Genic intolerance; 2) Network centrality; and 3) Gene expression in disease relevant tissues.

#### Genic Intolerance

Genic Intolerance quantifies the amount of purifying selection affecting a gene relative to a genome-wide average. Specifically, using standing human variation in large, publicly available, databases such as gnomAD (Lek et al, 2016), the number of common functional variants within a gene is regressed against the total number of variants within that gene. The Residual Variance Intolerance Score (RVIS) is the gene-level residual from that regression (Petrovski, Wang, Heinzen, Allen, & Goldstein, 2013). Thus, if a gene has less functional variation than expected given the total amount of variation within the gene, it will have a negative RVIS score. If it has more functional variation than expected, it will tend to have a positive score. RVIS has been shown to be strongly predictive of Mendelian disease genes, especially those that lead to early-onset severe disease phenotypes (Petrovski, Wang, Heinzen, Allen, & Goldstein, 2013).

Here, we calculate a gene’s intolerance-based weight, *w*_*ri*_, as the gene’s intolerance percentile among all 18536 scored genes, scaled to be between 0 and 2. By rescaling, we ensure that those genes that have intolerance scores that are less than the mean, and hence are more likely to be important in disease etiology, are given more importance in the overall gene-set, by decreasing their p-values.

#### Connectivity of genes within biologic gene sets

Interactions between genes in a genetic gene set can be represented by a network. In such a representation, nodes denote genes and the edges connecting them represent gene-gene interactions. It is quite common in biologic networks for a few genes to have a much larger number of connections than other genes. These highly connected genes are referred to as “hub” genes, and it is reasonable to hypothesize that deleterious mutations within such genes might be more disruptive of the biologic process represented by the network than mutations falling within less connected, more distal, genes.

The connectivity of a node is captured in the graph theory concept of “centrality” (White & Smyth, 2003; Borgatti, 2005), which can be quantified in a number of ways. Here, we investigate 3 measures of network centrality: degree centrality (White & Smyth, 2003; Borgatti, 2005), Eigenvector centrality (Freeman, 1979; Stephenson and Zelen, 1989; Wasserman and Faust, 1994; White & Smyth, 2003) and PageRank centrality (Page et al, 1998, White & Smyth, 2003). Degree centrality is simply the number of edges from a given node, i.e., the number of genes that interact with a given gene. Eigenvector centrality of a node is a measure of the importance of the nodes it is connected to, i.e. a node is important if it is connected to other important nodes. (Freeman, 1979; Stephenson and Zelen, 1989; Wasserman and Faust, 1994; White & Smyth, 2003). PageRank centrality is defined by the pageRank algorithm used in web searches by Google (Page et al, 1998, White & Smyth, 2003). pageRank assumes that a web page (node) is more important if it receives more links (directed connections) from other high pageRank scored web pages (Page et al, 1998, White & Smyth, 2003). Unlike degree centrality and eigenvector centrality, pageRank considers the directionality of the connection.

For a given centrality measure, Let *c*_*i*_ be the centrality for the *i*^*th*^ gene. In order to generate weights that result in smaller p-values for more highly connected genes, we take *w*_*ci*_ = *λ* (*ac*_*i*_ + *bc̅*)^−1^, where *c̅* is the mean centrality across the gene set, *a* and *b* are user-defined constants (here we take *a* = 0.95 and *b* = 0.05), and *λ* is a scaling factor so that the mean of the weights is one.

#### Gene expression in disease-related tissues

Genes that are important in disease etiology are more likely to be expressed in disease-related tissues during the developmental period leading to the disease. Therefore, for the *i*^*th*^ gene we define a weight *w*_*ei*_ = *λ* (*ae*_*i*_ + *be̅*)^−1^, where *e*_*i*_ is the expression level (appropriately normalized) in a disease-related tissue, *e̅* is the mean expression across all genes in the gene set, *a* and *b* are user-defined constants (here we take *a* = 0.95 and *b* = 0.05), and *λ* is a scaling factor so that the mean of the weights is one.

### Simulation study

We conduct a simple simulation study to evaluate the utility of our approach. For each scenario, we simulate 1e+4 datasets. For each simulated dataset we generate *n* independent statistics *X*_*i*_, *i* = 1, …, *n*, associated with *n* hypotheses. Let *π* be the proportion of the *X*_*i′s*_ that are generated under the alternative. We assume *X*_*i*_ ∼ *N*(*μ*, 1) under the alternative and *X*_*i*_ ∼ *N*(0,1) under the null. Thus, marginally, *X*_*i*_ ∼ *πN*(*μ*, 1) + (1 − *π*)*N*(0,1). Note that *π* characterizes the sparsity of the alternatives among all the hypotheses tested while *μ* controls the location shift from null to alternative. Thus, in our simulations, we evaluate the power of our approach as *π* and *μ* vary and choose configurations that explore the detection boundary outlined by Donoho & Jin, 2014. Each *X*_*i*_ is converted to a p-value via *p*_*i*_ = Φ(−|*X*_*i*_|). Weights are generated from a truncated exponential distribution and then scaled to have mean one. We consider three different scenarios: 1) weights are randomly assigned to genes; 2) weights are negatively correlated with disease-associated genes so that their p-values in the *HC*\* statistic are decreased, increasing their influence on the statistic; and 3) weights are positively correlated with disease-associated genes so that these genes will have less influence on the *HC*\* statistic while the influence of genes that are not disease-associated will be increased. We generate a large number of simulated datasets under the global null (i.e., *π* = 0) and use these to calculate a rejection threshold for each scenario. Specifically, we take the top 5^th^ percentile of *HC* and the *HC*\* statistics (corresponding to the weighting scenarios) calculated using the global null simulated datasets and use them as rejection thresholds for the corresponding statistics under the various alternative hypotheses. Please see supplementary materials for complete details.

### Amyotrophic lateral sclerosis whole exome study

We illustrate our approach through an analysis of data from a whole exome sequencing study comprised of 3093 ALS patients and 8186 controls of European ancestry (Gelfman et al, 2018). The sample information is available online (alsdb.org). The 18536 genes were sequenced and captured with the standard approach of Gelfman et al, 2018. Gene-level qualifying variant collapsing analyses were conducted according to two definitions of qualifying variants (table 1). Specifically, for a given qualifying variant definition and a given gene, we create an indicator variable of whether a given subject has a qualifying variant in that gene. We then test for association between the presence of a qualifying variant in the gene and case-control status using the Cochran–Mantel–Haenszel test, where strata are defined by the stratification score (Epstein et al, 2007). This results in two p-values per gene (corresponding to two definitions of qualifying variant); we take the minimum of the two to get a single gene-level p-value.

**Table 1.** Different Genetic Models and qualifying criteria

(MAF=Minor Allele Frequency, LoF=Loss-of-function variants, Dom=dominant)
| Model | type | LOO MAF | ExAC MAF | Qualifying Variant Effect Criteria |
| --- | --- | --- | --- | --- |
| **Damaging rare**<br>(coding) | Dom | 0.05% | 0.01% | LoF, inframe indels and PolyPhen-2 (HumDiv) “probably” |
| **LoF** | Dom | 0.1% | 0.1% | LoF |

We analyzed 4436 candidate gene sets extracted from the hallmark, GO biological process collections (c5.bp, version 6.1, Subramanian et al., 2005, Liberzon et al., 2015). Gene sets containing less than 10 genes were not analyzed.

We consider three different sources of prior information in developing gene-level weights: genetic networks from bioGRID (*BioGRID Version 3.4.147*, Stark et al., 2006, Chatraryamontri et al., 2017), genic intolerance (Petrovski, Wang, Heinzen, Allen, & Goldstein, 2013), and gene expression levels in disease-relevant tissue (brain) from GTex (GTEx Consortium, 2015). We used three different metrics for summarizing a genes importance within a genetic network: degree centrality (White & Smyth, 2003), eigenvector centrality (Freeman, 1979; Stephenson and Zelen, 1989; Wasserman and Faust, 1994; White & Smyth, 2003), and pageRank centrality (Page et al, 1998, White & Smyth, 2003).

For each gene-set, we conducted both weighted (denoted by *w*), using the weighting schemes highlighted above, and unweighted (denoted by *u*) analyses. Since we are interested in whether gene set analyses can help uncover specific gene sets that disease genes participate in, we also removed two genes that were found to be significantly associated with ALS: *SOD1* (raw *p*-value: 4.1e-15, Bonferroni adjusted *p*-value: 7.6e-11) and *NEK1* (raw *p*-value: 6.74e-10, Bonferroni adjusted *p*-value: 1.25e-05) from the gene set analyses. These analyses are denoted by *wr* and *ur* for weighted and unweighted analyses, respectively.

We obtained empirical null distributions of our statistics by randomly permuting case/control status (each distribution was generated using 2e+6 permutations). Because genes may participate in multiple gene sets, leading to correlation between tests, we used the step-down minP algorithm (Box 4: Ge, Dudoit, & Speed, 2003) to obtain multiplicity adjusted p-values across all the gene sets analyzed.

## RESULTS

### Simulation study

As expected, when the effect size, *μ*, is fixed, power increases with an increasing proportion of non-null hypotheses, *π* (figure 1). Similarly, when *π* is fixed, power increases with *μ*. More interestingly, we can see that weighting can substantially increase the power of the HC analysis if the values of weights are negatively correlated with non-null hypotheses, i.e. the weights tend to make p-values affiliated with non-null hypotheses smaller, so that those hypotheses have more influence on the final HC statistics (orange dashed lines). Further, the power is very similar to an unweighted analysis (solid black lines) when the weights are uncorrelated, i.e., are random noise (blue lines). It is only when the weights are positively correlated with the non-null hypotheses, so that null hypothesis are given more influence on the HC statistic, that we see a substantial negative effect on power when using weighting (red dash lines). However, in real applications one would expect that most weighting schemes would be somewhat informative of which genes would be disease-related. Thus, these results suggest that there is little downside to weighting individual hypotheses in HC analyses.

**Figure 1.**
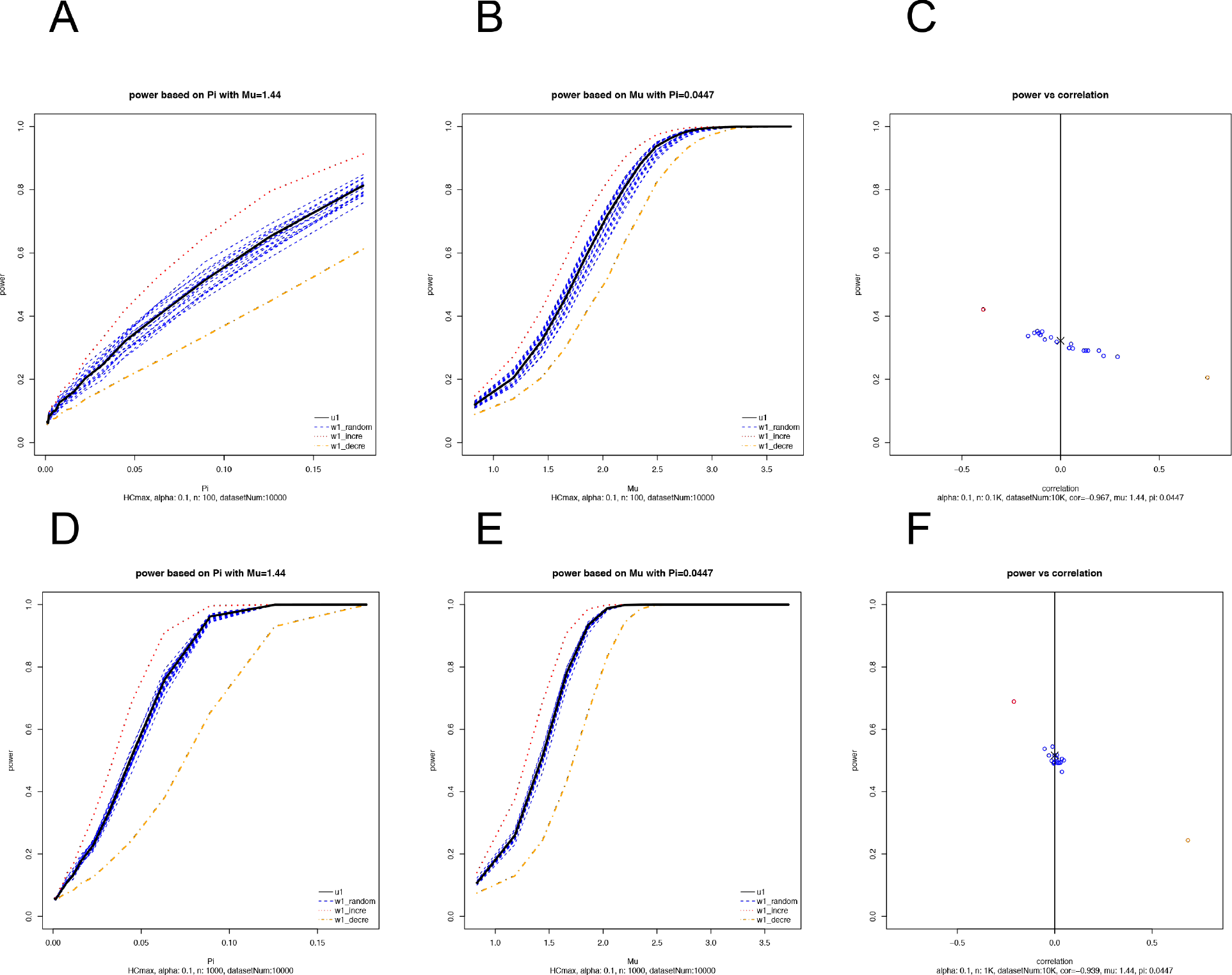
Selected Theoretical Simulation Results. Figure 1 shows the selected simulation results of the power of weighted HC (mu=1.44, pi=0.447). The black lines (points) represent HC with no weights, the blue dashed lines(points) represent HC with uncorrelated weights. The red dashed lines (points) represent HC with weights that negatively correlated with non-null hypotheses (favors the alternative). The orange dashed lines (points) represent HC with weights that negatively correlated with non-null hypotheses (favors the null). 1a) gene set size:100, Fixed mu. 1b) gene set size:100, Fixed pi. 1c) gene set size:100. power vs. correlations between weights and non-null hypothesis. 1d) gene set size:1000, Fixed mu. 1e) gene set size:1000, Fixed pi. 1f) gene set size:1000. power vs. correlations between weights and non-null hypothesis. See support information for more results.

### ALS data analysis

We found that marginally associated genes, had a strong effect on all HC analyses (weighted or not). For example, all gene-sets containing *SOD1* (260) and *NEK1* (21) are significantly associated with ALS after multiplicity adjustment, regardless of the HC statistic used (table 2). GSEA (Subramanian et al., 2005) fails to detect any significant gene sets. To investigate whether there is residual signal in gene sets after the marginally significant genes are removed, we conducted gene set analyses that excluded *SOD1* and *NEK1* from inclusion in any gene set. This analysis did not detect any significant gene sets after multiplicity adjustment, regardless of the method used.

**Table 2.** The number of significant gene sets by method

| Gene sets with | Total | Significance in HC (u) or wHC(GI, Exp, Deg, Eigen or PR) | Significance in GSEA(u or w) |
| --- | --- | --- | --- |
| **SOD1** | 260 | 260 | 0 |
| **NEK1** | 21 | 21 | 0 |

We then investigated whether there were enrichment differences between methods for gene sets involving 51 known ALS disease genes highlighted in Cirulli et al. 2015 (Table S1). The results of these analyses are presented in table 3 and one can see that weighting based on pageRank centralities performs well. Since many of these gene sets are likely devoid of any signal, we repeated this analysis while further restricting the gene sets considered to those where there was at least one gene-set analysis approach yielding a marginally significant result (p<=0.05) (table 4). Once again, we find that pageRank centrality does well and that HC outperforms GSEA.

**Table 3.** Method Comparison on ALS-related gene sets (Total gene sets: 899)

| Method 1 | Method 2 | Numbers of *p*-values<br>Method 1<=Method 2 | Numbers of *p*-values<br>Method 1>=Method 2 | *p*-values for<br>Method 1 better than Method 2 |
| --- | --- | --- | --- | --- |
| wHC(GI) | HC(u) | 468 | 432 | 1.22e-1 |
| wHC(Exp) | HC(u) | 397 | 503 | 1.00e-0 |
| wHC(Deg) | HC(u) | 490 | 409 | 3.80e-3 |
| wHC(Eigen) | HC(u) | 440 | 460 | 7.58e-1 |
| wHC(PR) | HC(u) | 501 | 398 | 3.31e-4 |
| HC(u) | GSEA(u) | 451 | 448 | 4.73e-1 |
| HC(u) | GSEA(w) | 436 | 463 | 8.25e-1 |

**Table 4.** Method Comparison on core ALS-related gene sets (Total gene sets: 186)

| Method 1 | Method 2 | Numbers of *p*-values<br>Method 1<=Method 2 | Numbers of *p*-values<br>Method 1>=Method 2 | *p*-values for<br>Method 1 better than Method 2 |
| --- | --- | --- | --- | --- |
| wHC(GI) | HC(u) | 89 | 98 | 7.68e-1 |
| wHC(Exp) | HC(u) | 76 | 111 | 9.96e-1 |
| wHC(Deg) | HC(u) | 116 | 70 | 4.59e-4 |
| wHC(Eigen) | HC(u) | 89 | 98 | 7.68e-1 |
| wHC(PR) | HC(u) | 123 | 64 | 9.57e-6 |
| HC(u) | GSEA(u) | 161 | 25 | 7.83e-26 |
| HC(u) | GSEA(w) | 158 | 28 | 1.67e-23 |

## DISCUSSION

We have presented a new gene-set based analysis that incorporates prior information into the analysis using a higher criticism approach. In both simulation studies and real data analyses, we showed that such an approach can lead to higher power. However, the choice of weights is important and consideration should be made for what information is most likely to be predictive of truly associated disease genes. For example, in our p-value enrichment analyses of known ALS genes, we found little enrichment when we used genic intolerance measures as our weights. As genic intolerance is indicative of purifying selection, this choice of weights may be less informative in a late-onset disorder such as ALS. Results would likely be different for earlier-onset disorders such as autism spectrum disorder, epilepsy, or schizophrenia. Further applications across a spectrum of diseases are needed before general recommendations can be made with respect to weighting schemes.

The higher criticism analysis can be thought of as a goodness of fit test that focuses on extreme deviations (by taking a max) from expectation under a global null that none of the genes within the gene set are associated with the disease. Though this approach has been shown to be optimal in detecting sparse signals within a large collection of hypotheses, it may be less sensitive to detecting signal that is more diffuse. In such a case, there may be an advantage in integrating over the tail of the distribution of deviations rather than taking a max. We are currently investigating this approach and plan to highlight it in a future manuscript.

## DESCRIPTION OF SUPPLEMEMTAL DATA

Supplemental Data include two tables and 4 series of figures. Table S1 lists the ALS-related genes according to Cirulli et al, 2015. Table S2 describes the simulation procedure. The text describes profiling details in the simulation study. The 4 series of figures provides the complete results of the simulation under various condition settings.

## Supporting information

wHC_support_text

supplementary_figure_series_of_correlation_alpha=0.1_n=100

supplementary_figure_series_of_correlation_alpha=0.1_n=1000

supplementary_figure_series_of_power_curve_alpha=0.1_n=100

supplementary_figure_series_of_power_curve_alpha=0.1_n=1000

## ACKNOWLEDGEMENTS

We thank Chong Jin from University of North Carolina at Chapel Hill and Dr. Zhiguo Li from Duke University, for suggestions on the asymptotic distribution of Higher Criticism based on the *p*-values and weighted *p*-values under the null hypothesis.

## WEB RESOURCES

An R package implementing the approach is available at https://github.com/mqzhanglab/wHC

## REFERENCES

Adzhubei, I. A., Schmidt, S., Peshkin, L., Ramensky, V. E., Gerasimova, A., Bork, P., … Sunyaev, S. R. (2010). A method and server for predicting damaging missense mutations. Nature Methods, 7(4), 248–249. https://doi.org/10.1038/nmeth0410-248

Al-Chalabi, A., Fang, F., Hanby, M. F., Leigh, P. N., Shaw, C. E., Ye, W., & Rijsdijk, F. (2010). An estimate of amyotrophic lateral sclerosis heritability using twin data. Journal of Neurology, Neurosurgery & Psychiatry, 81(12), 1324–1326. doi.org/10.1136/jnnp.2010.207464

Andersen, P. M., Borasio, G. D., Dengler, R., Hardiman, O., Kollewe, K., Leigh, P. N., … Tomik, B. (2007). Good practice in the management of amyotrophic lateral sclerosis: Clinical guidelines. An evidence‐based review with good practice points. EALSC Working Group. Amyotrophic Lateral Sclerosis, 8(4), 195–213. https://doi.org/10.1080/17482960701262376

Bapat, R. B., & Beg, M. I. (1989). Order Statistics for Nonidentically Distributed Variables and Permanents. Sankhyā: The Indian Journal of Statistics, Series A (1961-2002), 51(1), 79–93.

Barabási, A.-L., & Albert, R. (1999). Emergence of Scaling in Random Networks. Science, 286(5439), 509–512. https://doi.org/10.1126/science.286.5439.509

Barnett, I. J., & Lin, X. (2014). Analytic P-value calculation for the higher criticism test in finite d problems. Biometrika, 101(4), 964–970. https://doi.org/10.1093/biomet/asu033

Barnett, I., Mukherjee, R., & Lin, X. (2017). The Generalized Higher Criticism for Testing SNP-Set Effects in Genetic Association Studies. Journal of the American Statistical Association, 112(517), 64–76. https://doi.org/10.1080/01621459.2016.1192039

Benjamini, Y., & Hochberg, Y. (1995). Controlling the False Discovery Rate: A Practical and Powerful Approach to Multiple Testing. Journal of the Royal Statistical Society. Series B (Methodological), 57(1), 289–300.

Borgatti, S. P. (2005). Centrality and network flow. Social Networks, 27(1), 55–71. https://doi.10.1016/j.socnet.2004.11.008

Boyle, E. A., Li, Y. I., & Pritchard, J. K. (2017). An Expanded View of Complex Traits: From Polygenic to Omnigenic. Cell, 169(7), 1177–1186. https://doi.org/10.1016/j.cell.2017.05.038

Brenner, D., Müller, K., Wieland, T., Weydt, P., Böhm, S., Lulé, D., … Weishaupt, J. H. (2016). NEK1 mutations in familial amyotrophic lateral sclerosis. Brain, 139(5), e28–e28. https://doi.org/10.1093/brain/aww033

Chatr-aryamontri, A., Oughtred, R., Boucher, L., Rust, J., Chang, C., Kolas, N. K., … Tyers, M. (2017). The BioGRID interaction database: 2017 update. Nucleic Acids Research, 45(Database issue), D369–D379. https://doi.org/10.1093/nar/gkw1102

Cirulli, E. T., Lasseigne, B. N., Petrovski, S., Sapp, P. C., Dion, P. A., Leblond, C. S., … Goldstein, D. B. (2015). Exome sequencing in amyotrophic lateral sclerosis identifies risk genes and pathways. Science, 347(6229), 1436–1441. https://doi.org/10.1126/science.aaa3650

Donoho, D., & Jin, J. (2004). Higher Criticism for Detecting Sparse Heterogeneous Mixtures. The Annals of Statistics, 32(3), 962–994. Download. (n.d.). Retrieved December 27, 2018, from http://alsdb.org/download.jsp

Epstein, M. P., Allen, A. S., & Satten, G. A. (2007). A Simple and Improved Correction for Population Stratification in Case-Control Studies. The American Journal of Human Genetics, 80(5), 921–930. https://doi.org/10.1086/516842

Freeman, L. C., Roeder, D., & Mulholland, R. R. (1979). Centrality in social networks: ii. experimental results. Social Networks, 2(2), 119–141. https://doi.org/10.1016/0378-8733(79)90002-9

Ge, Y., Dudoit, S., & Speed, T. P. (2003). Resampling-based multiple testing for microarray data analysis. Test, 12(1), 1–77. https://doi.org/10.1007/BF02595811

Gelfman, S., Dugger, S. A., Moreno, C. A. M., Ren, Z., Wolock, C. J., Shneider, N. A., … Goldstein, D. B. (2018). Regional collapsing of rare variation implicates specific genic regions in ALS. BioRxiv, 375774. https://doi.org/10.1101/375774

Genovese, C. R., Roeder, K., & Wasserman, L. (2006). False Discovery Control with p-Value Weighting. Biometrika, 93(3), 509–524.

Goh, K.-I., Cusick, M. E., Valle, D., Childs, B., Vidal, M., & Barabási, A.-L. (2007). The human disease network. Proceedings of the National Academy of Sciences, 104(21), 8685–8690.

Hall, P., & Jin, J. (2010). Innovated higher criticism for detecting sparse signals in correlated noise. The Annals of Statistics, 38(3), 1686–1732. https://doi.org/10.1214/09-AOS764

Kabashi, E., Valdmanis, P. N., Dion, P., Spiegelman, D., McConkey, B. J., Velde, C. V., … Rouleau, G. A. (2008). TARDBP mutations in individuals with sporadic and familial amyotrophic lateral sclerosis. Nature Genetics, 40(5), 572–574. https://doi.org/10.1038/ng.132

Kircher, M., Witten, D. M., Jain, P., O’Roak, B. J., Cooper, G. M., & Shendure, J. (2014). A general framework for estimating the relative pathogenicity of human genetic variants. Nature Genetics, 46(3), 310–315. https://doi.org/10.1038/ng.2892

Kosorok, M. R. (2008). Introduction to empirical processes and semiparametric inference. New York: Springer.

Lek, M., Karczewski, K. J., Minikel, E. V., Samocha, K. E., Banks, E., Fennell, T., … Exome Aggregation Consortium. (2016). Analysis of protein-coding genetic variation in 60,706 humans. Nature, 536(7616), 285–291. https://doi.org/10.1038/nature19057

Liberzon, A., Birger, C., Thorvaldsdóttir, H., Ghandi, M., Mesirov, J. P., & Tamayo, P. (2015). The Molecular Signatures Database Hallmark Gene Set Collection. Cell Systems, 1(6), 417–425. https://doi.org/10.1016/j.cels.2015.12.004

Liberzon, A., Subramanian, A., Pinchback, R., Thorvaldsdóttir, H., Tamayo, P., & Mesirov, J. P. (2011). Molecular signatures database (MSigDB) 3.0. Bioinformatics, 27(12), 1739–1740. https://doi.org/10.1093/bioinformatics/btr260

Lowe, D. G. (1999). Object recognition from local scale-invariant features. In Proceedings of the Seventh IEEE International Conference on Computer Vision (Vol. 2, pp. 1150–1157 vol.2). https://doi.org/10.1109/ICCV.1999.790410

Mann, H. B., & Wald, A. (1943). On Stochastic Limit and Order Relationships. The Annals of Mathematical Statistics, 14(3), 217–226.

Mooney, M. A., Nigg, J. T., McWeeney, S. K., & Wilmot, B. (2014). Functional and genomic context in pathway analysis of GWAS data. Trends in Genetics, 30(9), 390–400. https://doi.org/10.1016/j.tig.2014.07.004

OMIM - Online Mendelian Inheritance in Man. (n.d.). Retrieved December 17, 2018, from https://www.omim.org/

Page, L., Brin, S., Motwani, R., & Winograd, T. (1999). The PageRank citation ranking: Bringing order to the web. Stanford InfoLab. Retrieved from http://ilpubs.stanford.edu:8090/422

Palmer, S.M., Snyder, L., Todd, J. L., Soule, B., Christian, R., Anstrom, K., Luo, Y., Gagnon, R., and Rosen, G. (2018). Randomized, Double-Blind, Placebo-Controlled, Phase 2 Trial of BMS-986020, a Lysophosphatidic Acid Receptor Antagonist for the Treatment of Idiopathic Pulmonary Fibrosis. Chest 154, 1061–1069.

Park, J.-H., Gail, M. H., Weinberg, C. R., Carroll, R. J., Chung, C. C., Wang, Z., Chanock, S. J., Fraumeni, J. F., and Chatterjee, N. (2011). Distribution of allele frequencies and effect sizes and their interrelationships for common genetic susceptibility variants. Proceedings of the National Academy of Sciences 108, 18026–18031.

Petrovski, S., Todd, J. L., Durheim, M. T., Wang, Q., Chien, J. W., Kelly, F. L., … Goldstein, D. B. (2017). An Exome Sequencing Study to Assess the Role of Rare Genetic Variation in Pulmonary Fibrosis. American Journal of Respiratory and Critical Care Medicine, 196(1), 82–93. https://doi.org/10.1164/rccm.201610-2088OC

Petrovski, S., Wang, Q., Heinzen, E.L., Allen, A.S., & Goldstein, D.B. (2013). Genic Intolerance to Functional Variation and the Interpretation of Personal Genomes. PLOS Genetics, 9(8), e1003709. https://doi.org/10.1371/journal.pgen.1003709

Poppe, L., Rué, L., Robberecht, W., & Van Den Bosch, L. (2014). Translating biological findings into new treatment strategies for amyotrophic lateral sclerosis (ALS). Experimental Neurology, 262, 138–151. https://doi.org/10.1016/j.expneurol.2014.07.001

Rao, J.S., & Sethuraman, J. (1975). Weak Convergence of Empirical Distribution Functions of Random Variables Subject to Perturbations and Scale Factors. The Annals of Statistics, 3(2), 299–313.

Renton, A. E., Chiò, A., & Traynor, B. J. (2014). State of play in amyotrophic lateral sclerosis genetics. Nature Neuroscience, 17(1), 17–23. https://doi.org/10.1038/nn.3584

Roeder, K., & Wasserman, L. (2009). Genome-Wide Significance Levels and Weighted Hypothesis Testing. Statistical Science: A Review Journal of the Institute of Mathematical Statistics, 24(4), 398–413. https://doi.org/10.1214/09-STS289

Rosen, D. R., Siddique, T., Patterson, D., Figlewicz, D. A., Sapp, P., Hentati, A., … Brown Jr, R. H. (1993). Mutations in Cu/Zn superoxide dismutase gene are associated with familial amyotrophic lateral sclerosis. Nature, 362(6415), 59–62. https://doi.org/10.1038/362059a0

Stark, C., Breitkreutz, B.-J., Reguly, T., Boucher, L., Breitkreutz, A., & Tyers, M. (2006). BioGRID: a general repository for interaction datasets. Nucleic Acids Research, 34(Database issue), D535–D539. https://doi.org/10.1093/nar/gkj109

Stephenson, K., & Zelen, M. (1989). Rethinking centrality: Methods and examples. Social Networks, 11(1), 1–37. https://doi.org/10.1016/0378-8733(89)90016-6

Storey, J.D. (2003). The Positive False Discovery Rate: A Bayesian Interpretation and the q-Value. The Annals of Statistics, 31(6), 2013–2035.

Subramanian, A., Tamayo, P., Mootha, V. K., Mukherjee, S., Ebert, B.L., Gillette, M. A., … Mesirov, J. P. (2005). Gene set enrichment analysis: A knowledge-based approach for interpreting genome-wide expression profiles. Proceedings of the National Academy of Sciences, 102(43), 15545–15550. https://doi.org/10.1073/pnas.0506580102

Wang, K., Li, M., & Bucan, M. (2007). Pathway-Based Approaches for Analysis of Genomewide Association Studies. The American Journal of Human Genetics, 81(6), 1278–1283. https://doi.org/10.1086/522374

Wasserman, S., & Faust, K. (1994). Social Network Analysis: Methods and Applications. Cambridge University Press.

White, S., & Smyth, P. (2003). Algorithms for Estimating Relative Importance in Networks. In Proceedings of the Ninth ACM SIGKDD International Conference on Knowledge Discovery and Data Mining (pp. 266–275). New York, NY, USA: ACM. https://doi.org/10.1145/956750.956782

