## Supplementary material for "Incorporating prior information into signal-detection analyses across biologically informed gene-sets": wHC_support_text

**Supporting Information**

Table S1: list of ALS related genes (according to Cirulli et al, 2015)

| | | | | |
|---|---|---|---|---|
| SOD1 | VAPB | hnRNPA1 | DCTN1 | DPP6 |
| NEK1 | TBK1 | DAO | PRPH | FGGY |
| C9ORF72 | FIG4 | TAF15 | ATXN2 | FLJ10986 |
| TARDBP | ERBB4 | EWSR1 | GRN | ITPR2 |
| FUS | SPG11 | SQSTM1 | PON1-3 | NIPA1 |
| UBQLN2 | ALS2 | MATR3 | ELP3 | HFE |
| OPTN | SIGMAR1 | TUBB4 | UNC13A | KIFAP3 |
| PFN1 | ERLIN2 | SSI8L1 | SMN1 | APEX1 |
| VCP | ANG | CHMP2B | TREM2 | PLCD1 |
| SETX | hnRNPA2B1 | NEFH | 17q11.2 | ARHGEF28 |
| VEGF | | | | |

**Simulation**

Data Generation and Parameters

We generate $X_i, i = 1,2, \dots n$ corresponding to $n$ independent hypotheses. Under the null, the $X_i$s are assumed to follow a standard normal, i.e.,

$$X_i \sim N(0,1), i = 1,2, \dots n.$$

Under the alternative hypothesis, we assume that most of the $X_i$s still come from a standard normal except for a small proportion, $\pi$, that experience a mean shift, i.e.,

$$X_i \sim (1-\pi)N(0,1) + \pi\, N(\mu, 1), \qquad i = 1,2, \dots n$$

Thus, $\pi$ and $\mu$ jointly control the departure of the alternative from the null. In our simulations, we take multiple combination of $\pi$ and $\mu$ to explore the detection boundary obtained by Donoho & Jin (2004). Specifically, we take $\pi = n^{-\gamma}$, where $\gamma$ is a scalar that varies from 0.25 to 1. Similarly, we take $\mu = \sqrt{2rlog(n)}$ where $r$ is a scalar that varies from 0.05 and 1. *P*-values are calculated via the upper-tail of the standard normal distribution, i.e.,

$$p_i = \Phi(-|X_i|)$$

Weights, $w_i, i = 1, \dots, n,$ are generated from a truncated exponential distribution and scaled to have mean 1. Specifically, we generate $w_i^* \sim \exp{(1)}$, threshold so that $w_i^* > 1$, and then rescale so that $w_i = w_i^* / mean(w_i^*)$.

Given the generated weights and p-values generated under a specific alternative hypothesis, we investigate three scenarios. First, the weights are taken as generated so that there is no relationship between the weights and those genes that are under the alternate hypothesis. We denote these weights by $w_{random}$. Second, we take the largest of the generated weights and assign them to those genes that are under the alternate hypothesis. We denote these weights by $w_{inc}$. Finally, we take the smallest of the generated weights and assign them to those genes that are under the alternate hypothesis. We denote these weights by $w_{dec}$. The general structure of our simulation algorithm is described in Table S2.

**Table S2: Simulation Procedure**

Distribution under null hypothesis $H_0$:

1. Generate *X* from null (for 1e+4 datasets, each with n=100 or n=1000)
2. Calculate *p*-values
3. Calculate non-weighted HC *p*-values for each dataset
4. Calculate weighted HC *p*-values for each dataset,
5. Obtain $\alpha_u$, the top 5th quantile of the non-weighted HC statistic
6. Obtain $\alpha_w$, the top 5th quantile of the weighted HC statistic

Distribution under alternative hypothesis $H_1$:

For each scenario ($\pi$ and $\mu$), iterate through 7-10

7. Generate *X* from alternative. (for 1e+4 datasets, each with n=100 or n=1000)
8. Repeat step 2-4 above
9. Find the proportion of non-weighted HC statistic which are greater than $\alpha_u$.
10. Find the proportion of weighted HC statistic which are greater than $\alpha_w$.

The proportions computed in steps 9 and 10 are empirical estimates of the power of the test.

References

Cirulli, E.T., Lasseigne, B.N., Petrovski, S., Sapp, P.C., Dion, P.A., Leblond, C.S., Couthouis, J., Lu, Y.-F., Wang, Q., Krueger, B.J., et al. (2015). Exome sequencing in amyotrophic lateral sclerosis identifies risk genes and pathways. Science 347, 1436–1441.

Donoho, D., and Jin, J. (2004). Higher Criticism for Detecting Sparse Heterogeneous Mixtures. The Annals of Statistics 32, 962–994.
