## supplementary_figure_series_of_correlation_alpha=0.1_n=100 for "Incorporating prior information into signal-detection analyses across biologically informed gene-sets"

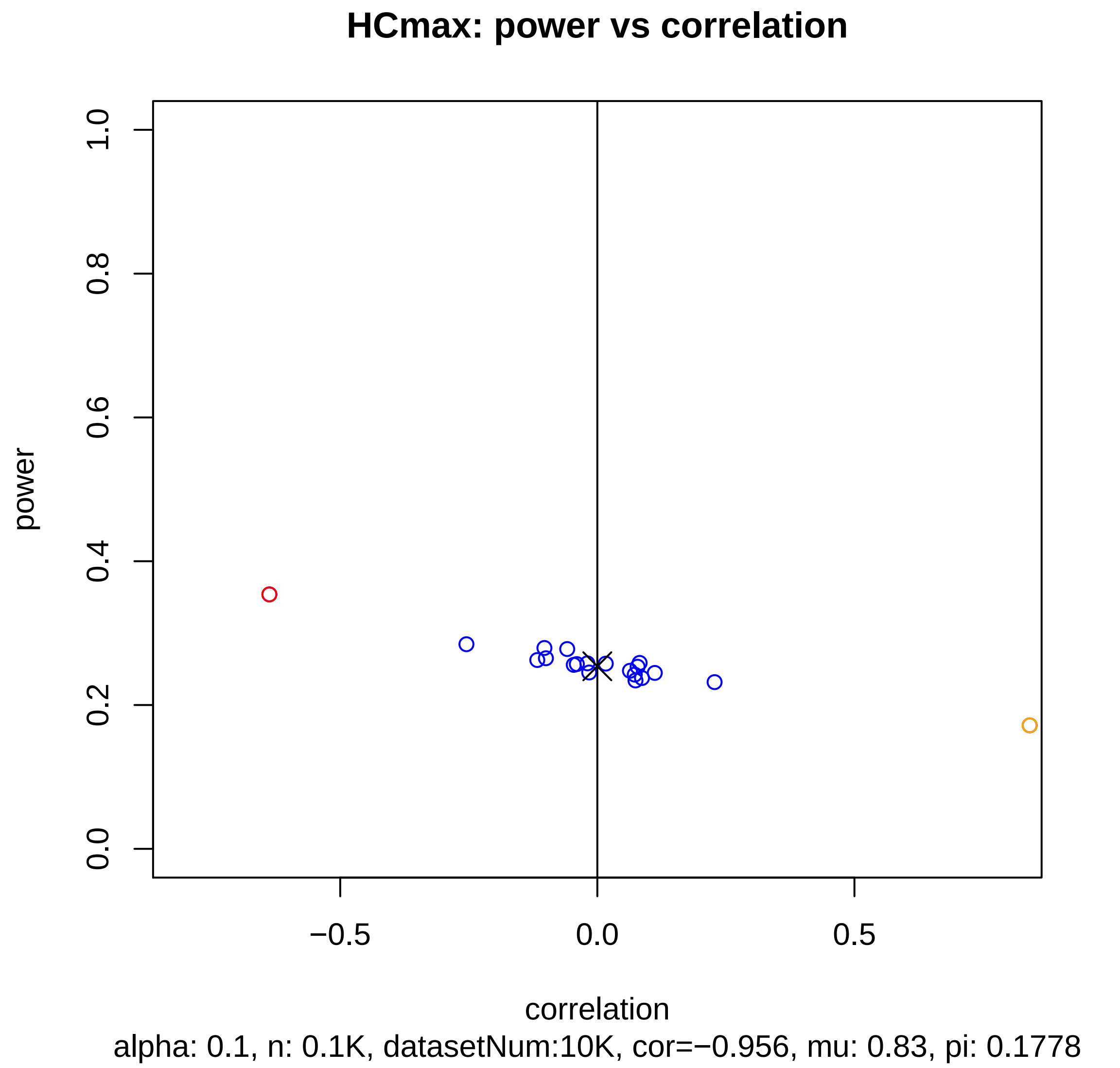
HCmax: power vs correlation
power
1.0
0.8
0.6
0.4
0.2
0.0
−0.5
0.0
0.5
correlation
alpha: 0.1, n: 0.1K, datasetNum:10K, cor=−0.956, mu: 0.83, pi: 0.1778

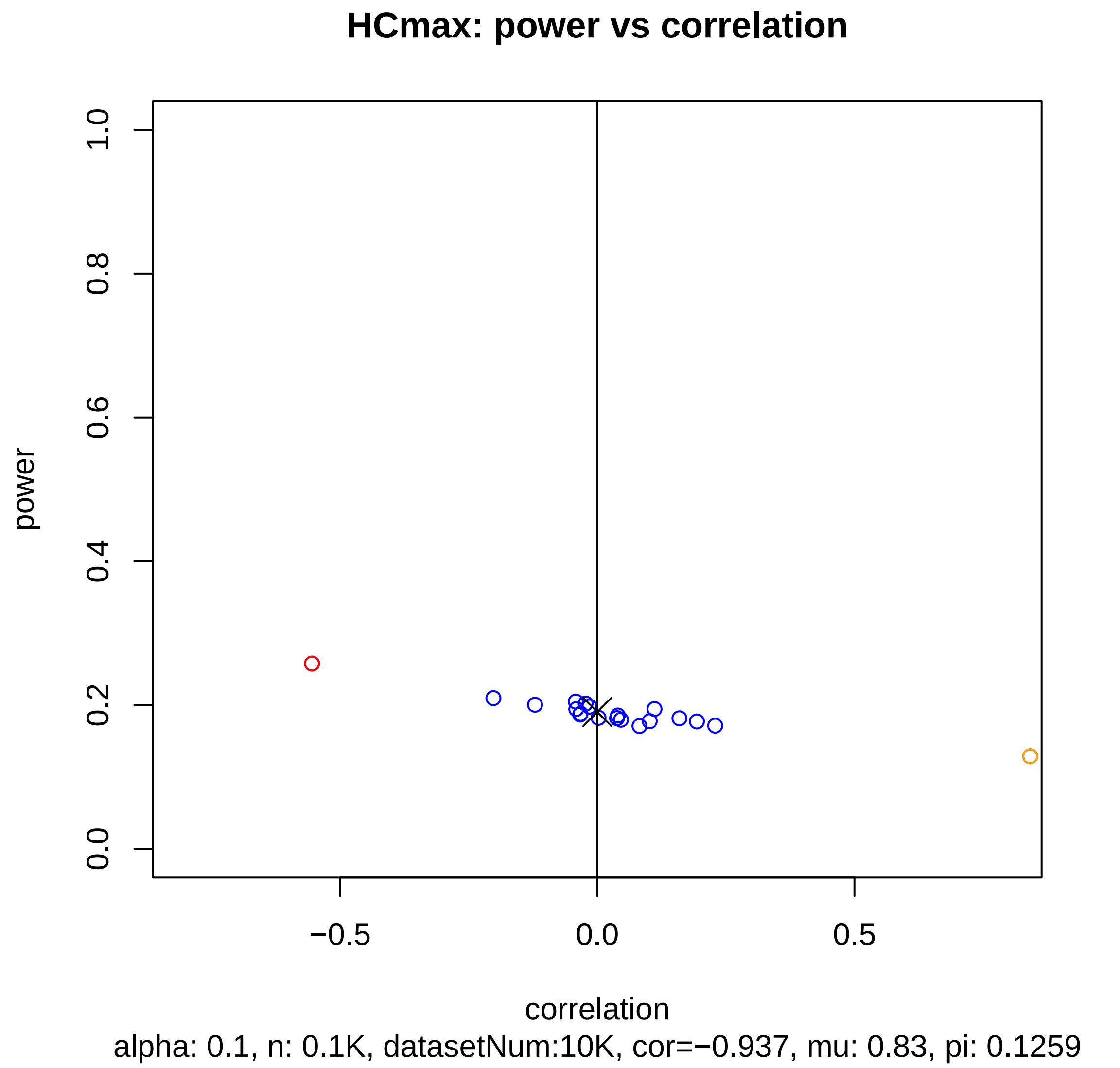
HCmax: power vs correlation
power
1.0
0.8
0.6
0.4
0.2
0.0
−0.5
0.0
0.5
correlation
alpha: 0.1, n: 0.1K, datasetNum:10K, cor=−0.937, mu: 0.83, pi: 0.1259

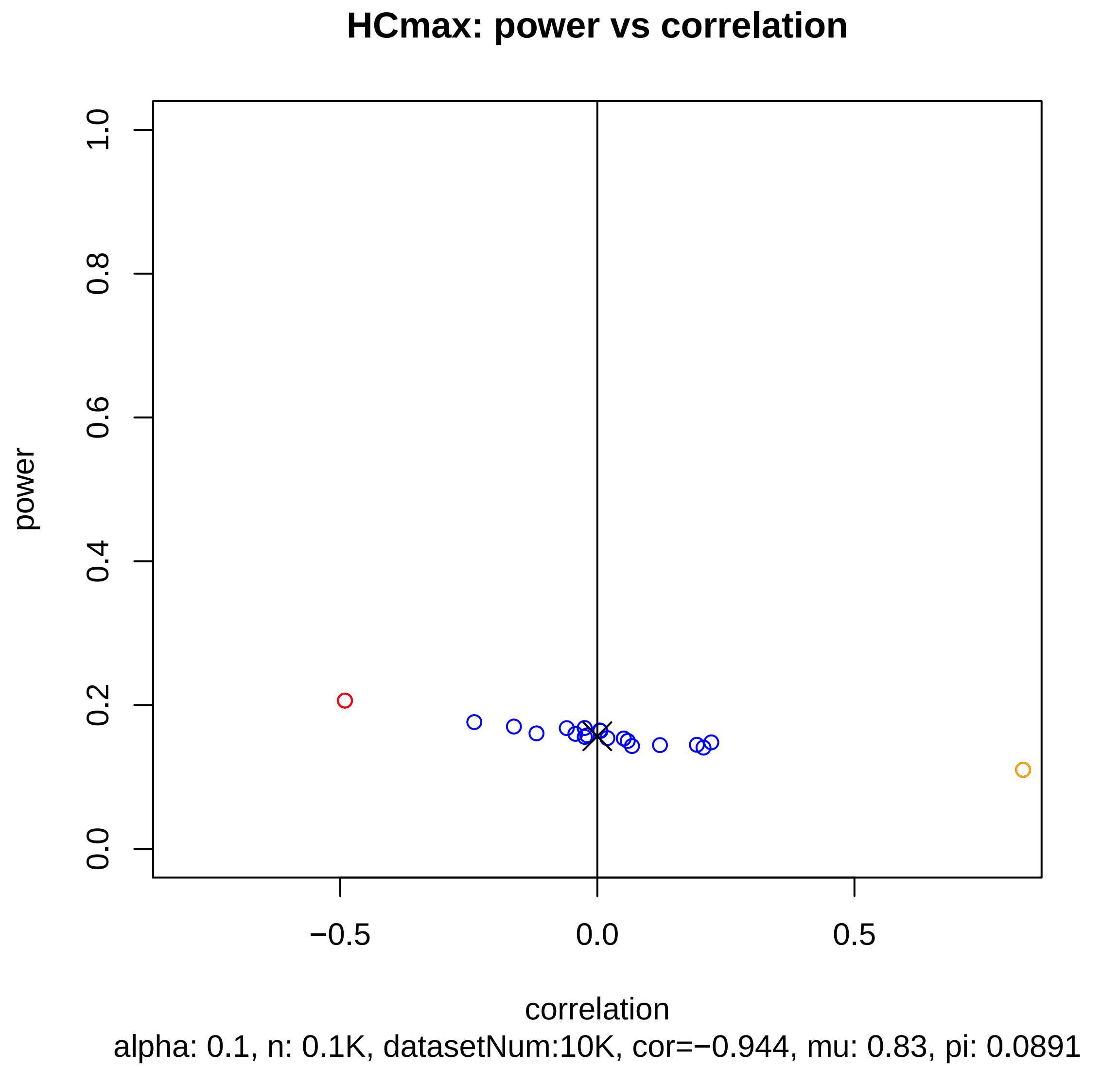

HCmax: power vs correlation
power
1.0
0.8
0.6
0.4
0.2
0.0
−0.5
0.0
0.5
correlation
alpha: 0.1, n: 0.1K, datasetNum:10K, cor=−0.944, mu: 0.83, pi: 0.0891

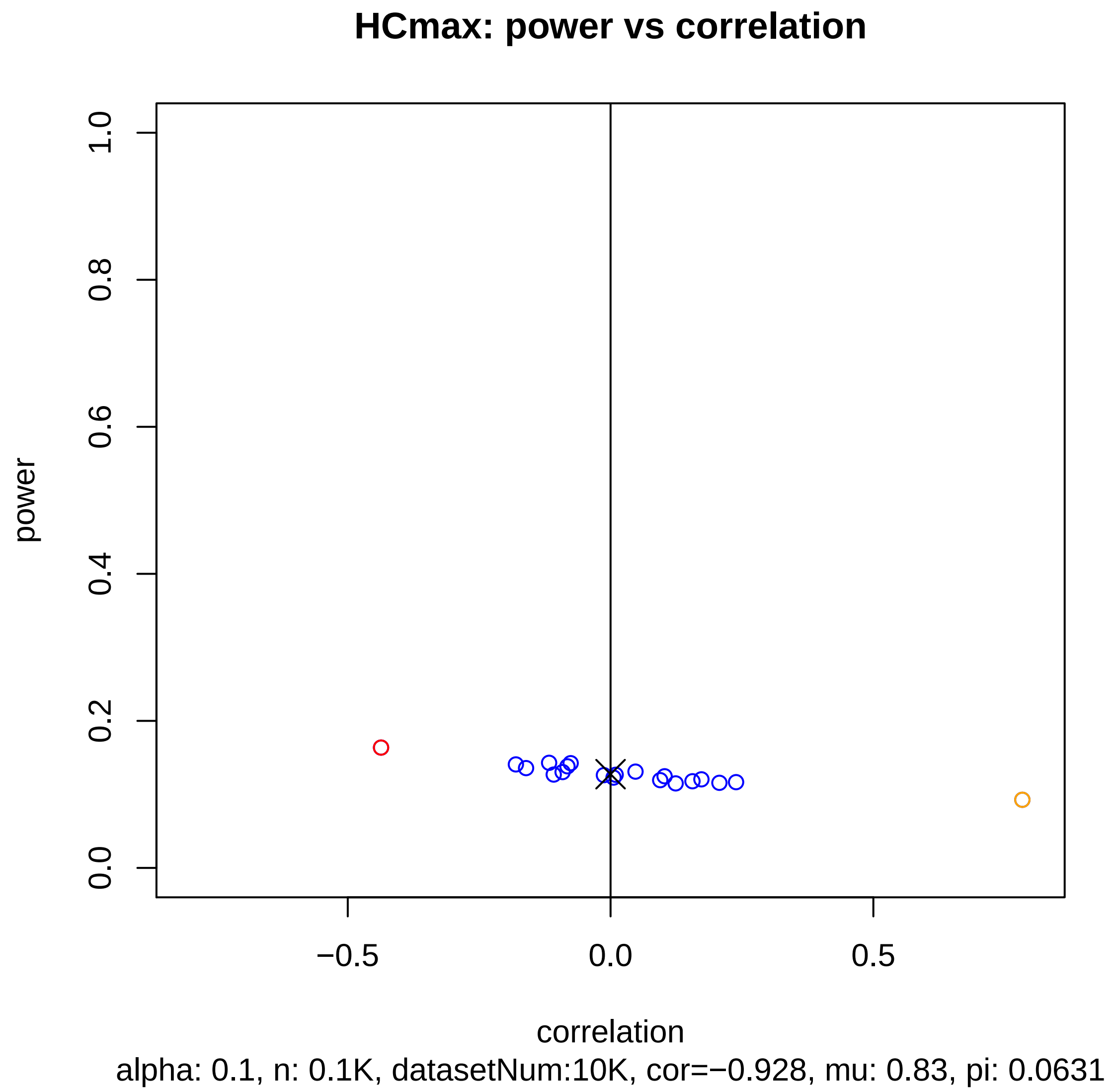

HCmax: power vs correlation
power
1.0
0.8
0.6
0.4
0.2
0.0
−0.5
0.0
0.5
correlation
alpha: 0.1, n: 0.1K, datasetNum:10K, cor=−0.928, mu: 0.83, pi: 0.0631

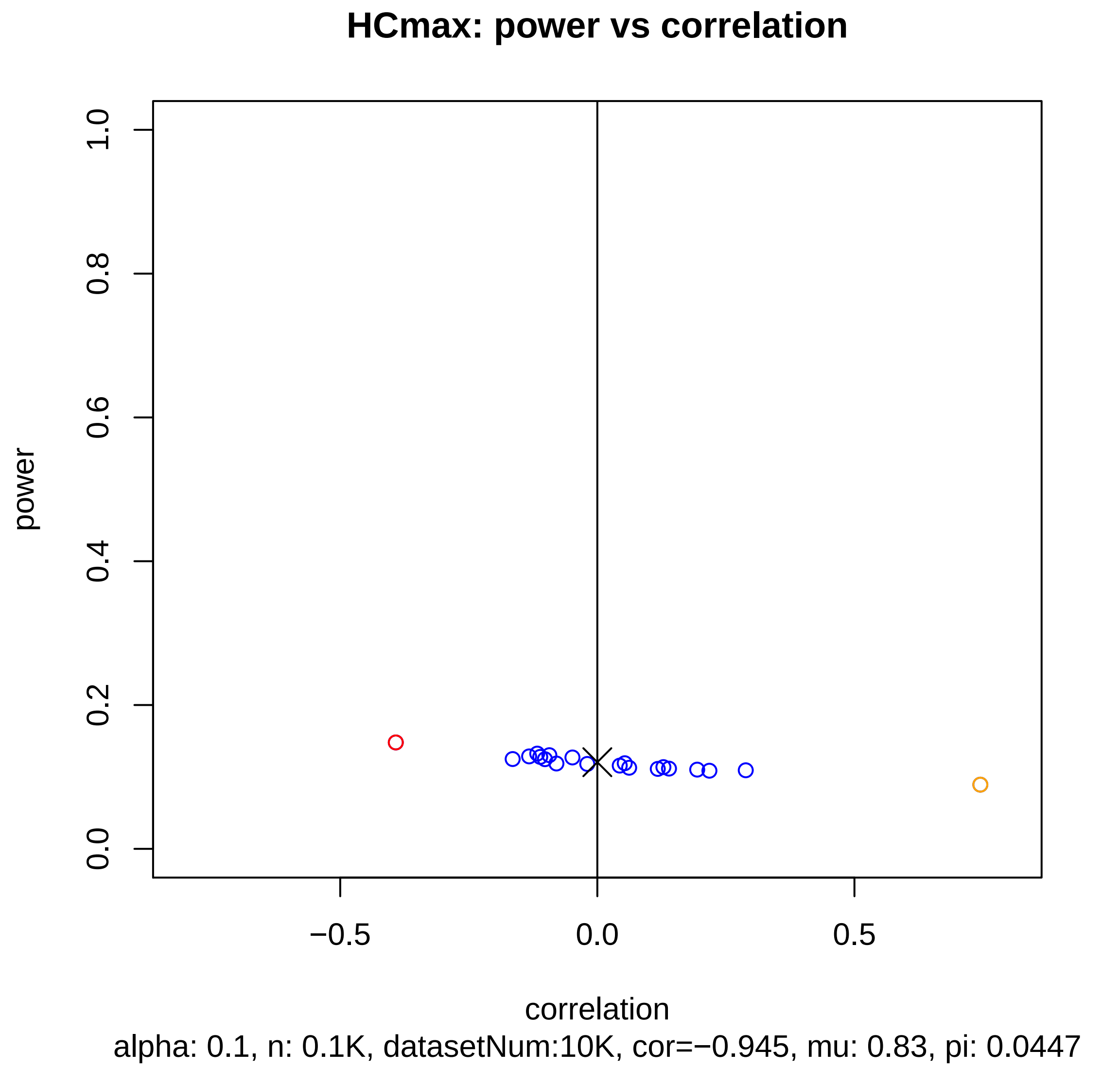
HCmax: power vs correlation
power
1.0
0.8
0.6
0.4
0.2
0.0
−0.5
0.0
0.5
correlation
alpha: 0.1, n: 0.1K, datasetNum:10K, cor=−0.945, mu: 0.83, pi: 0.0447

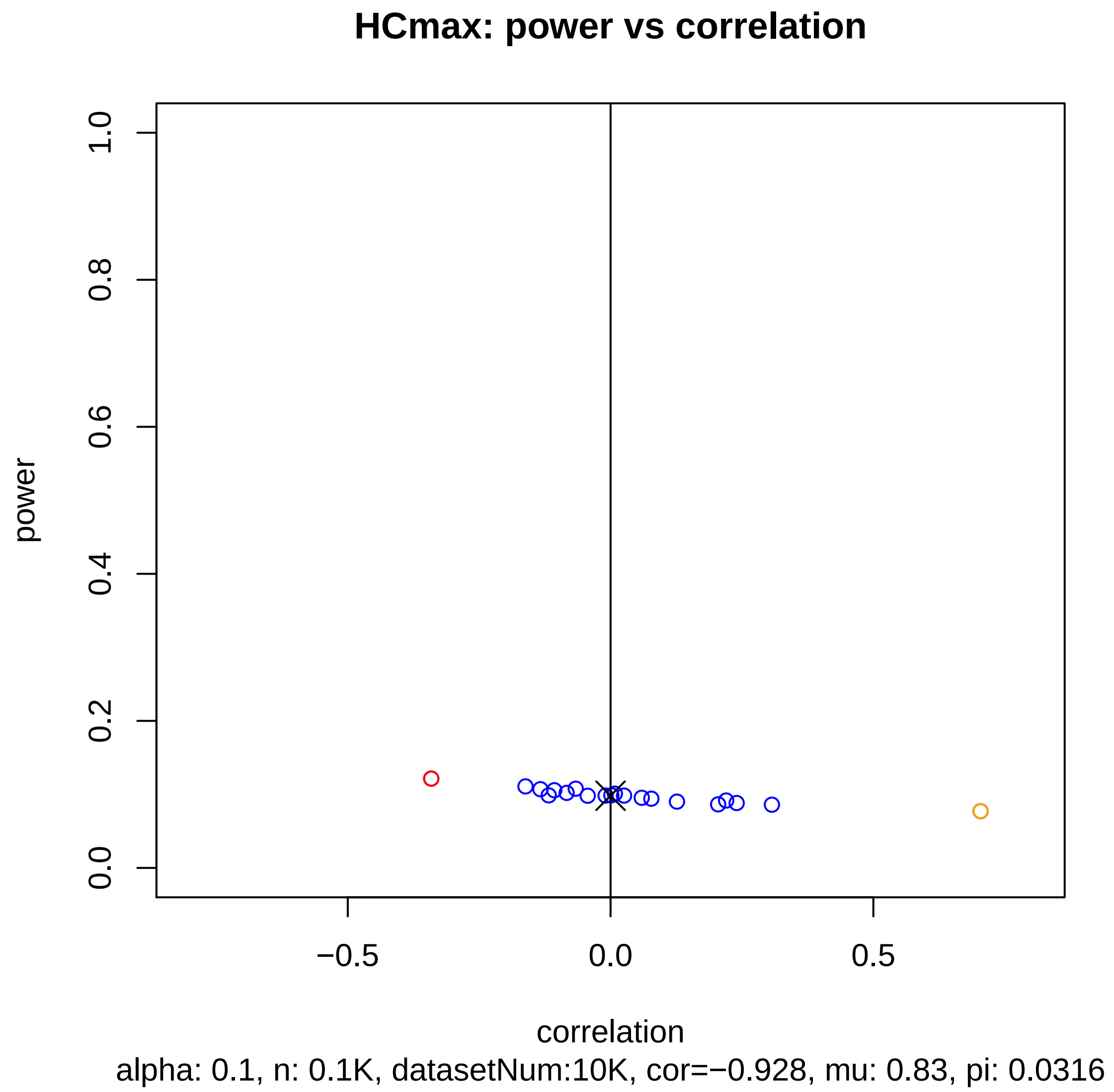
HCmax: power vs correlation
power
1.0
0.8
0.6
0.4
0.2
0.0
−0.5
0.0
0.5
correlation
alpha: 0.1, n: 0.1K, datasetNum:10K, cor=−0.928, mu: 0.83, pi: 0.0316

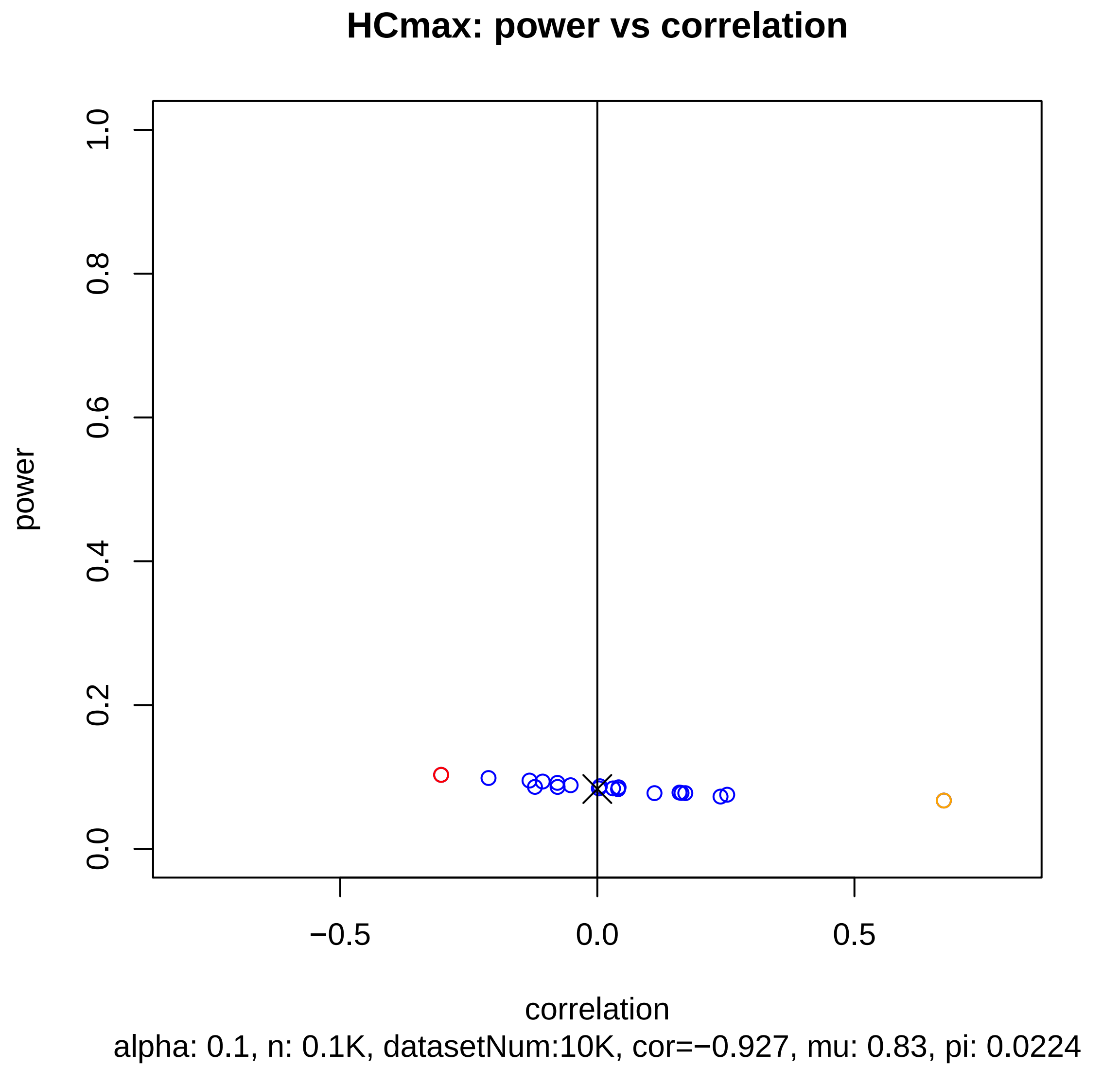
HCmax: power vs correlation
power
1.0
0.8
0.6
0.4
0.2
0.0
−0.5
0.0
0.5
correlation
alpha: 0.1, n: 0.1K, datasetNum:10K, cor=−0.927, mu: 0.83, pi: 0.0224

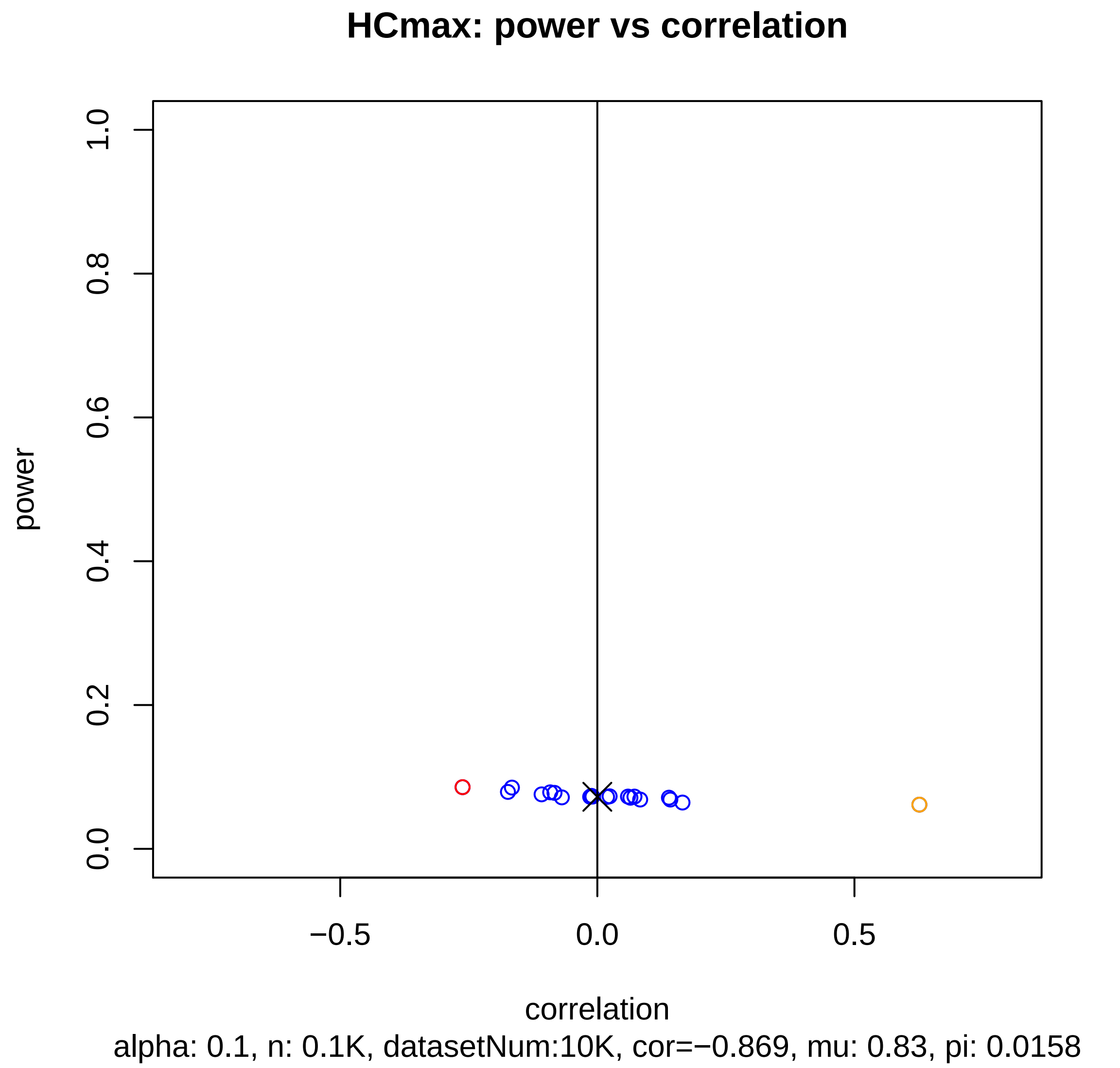
HCmax: power vs correlation
power
0.0
0.2
0.4
0.6
0.8
1.0
−0.5
0.0
0.5
correlation
alpha: 0.1, n: 0.1K, datasetNum:10K, cor=−0.869, mu: 0.83, pi: 0.0158

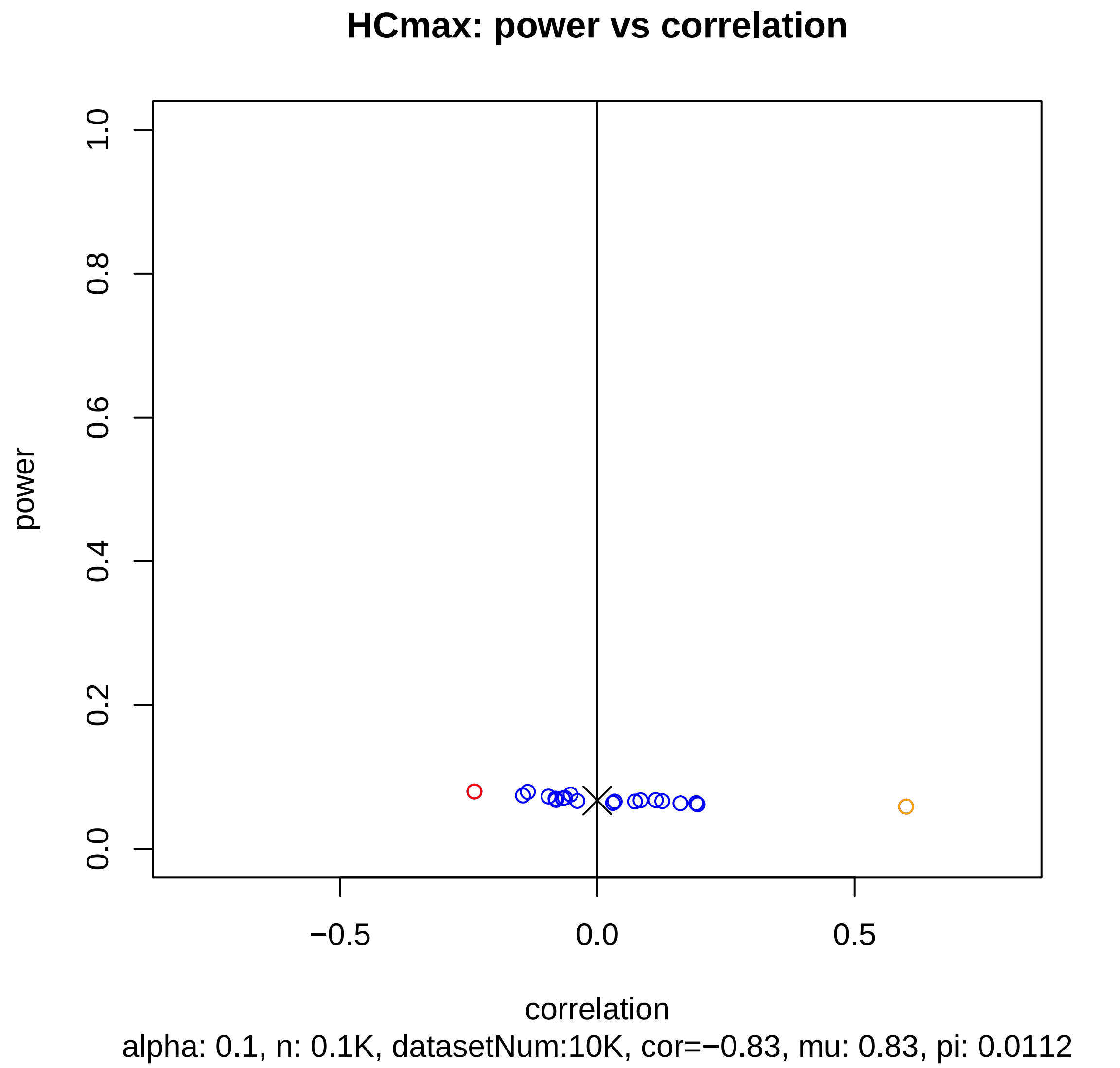
HCmax: power vs correlation
power
1.0
0.8
0.6
0.4
0.2
0.0
−0.5
0.0
0.5
correlation
alpha: 0.1, n: 0.1K, datasetNum:10K, cor=−0.83, mu: 0.83, pi: 0.0112

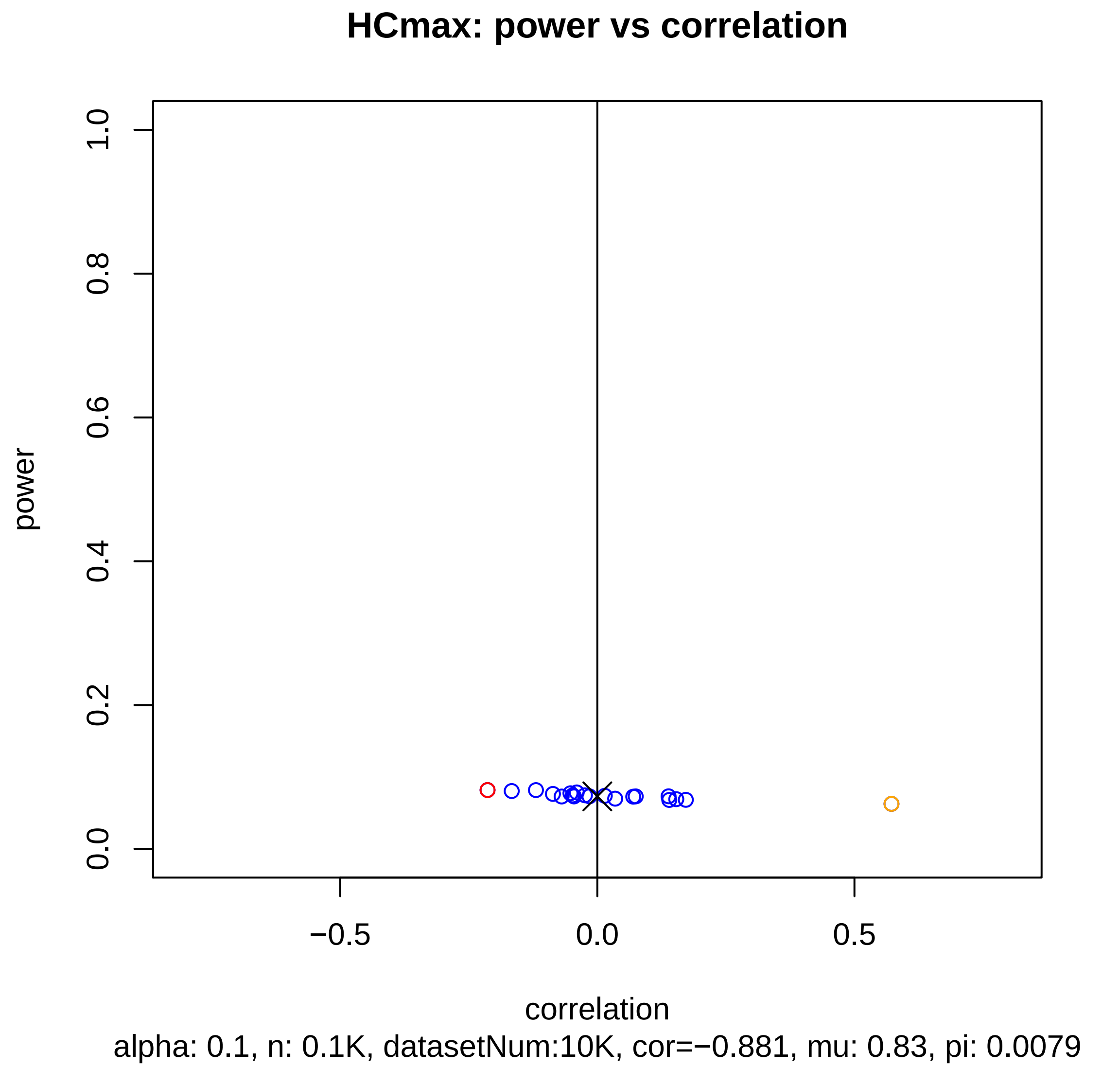

HCmax: power vs correlation
power
1.0
0.8
0.6
0.4
0.2
0.0
−0.5
0.0
0.5
correlation
alpha: 0.1, n: 0.1K, datasetNum:10K, cor=−0.881, mu: 0.83, pi: 0.0079

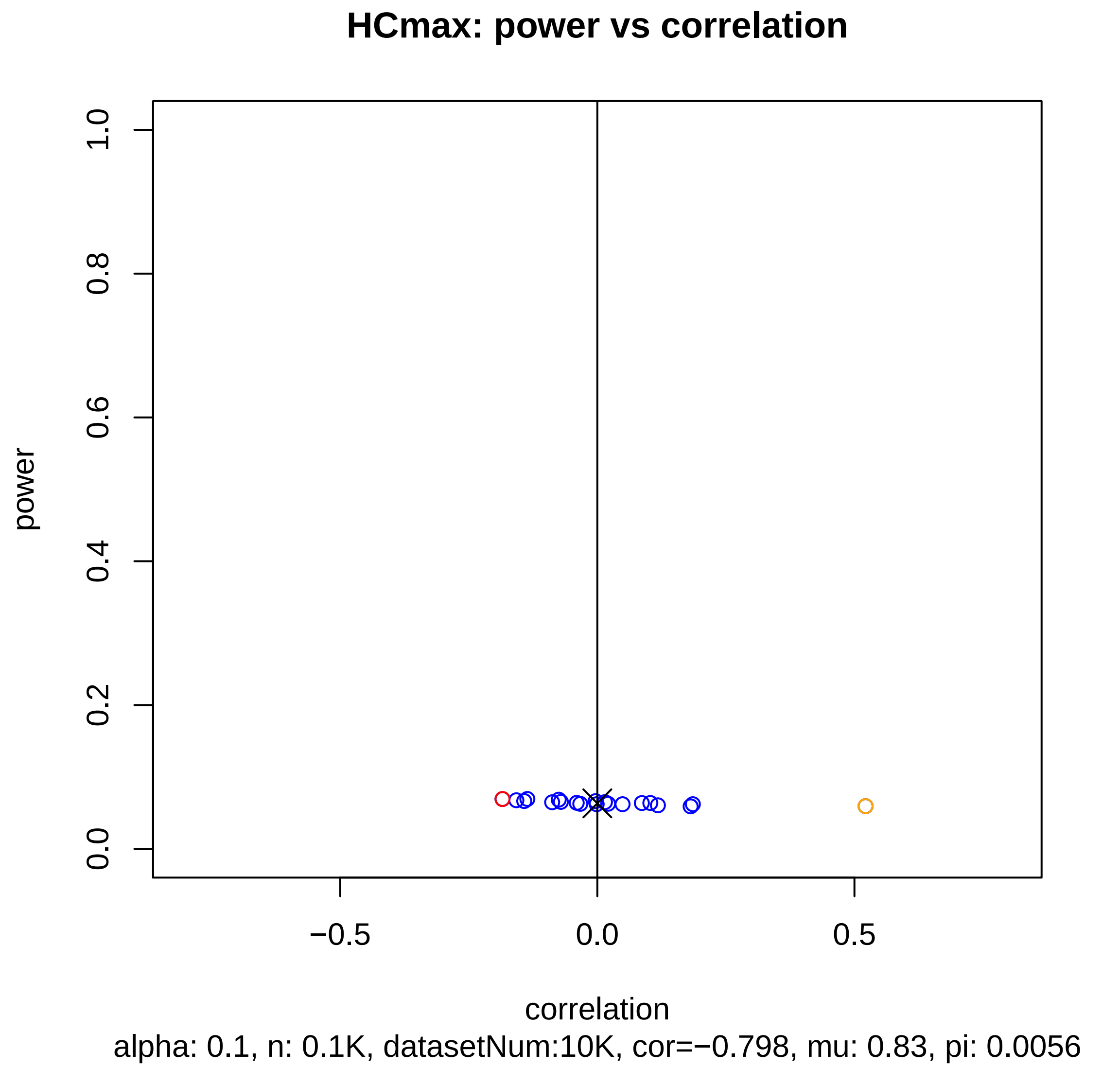
HCmax: power vs correlation
power
1.0
0.8
0.6
0.4
0.2
0.0
−0.5
0.0
0.5
correlation
alpha: 0.1, n: 0.1K, datasetNum:10K, cor=−0.798, mu: 0.83, pi: 0.0056

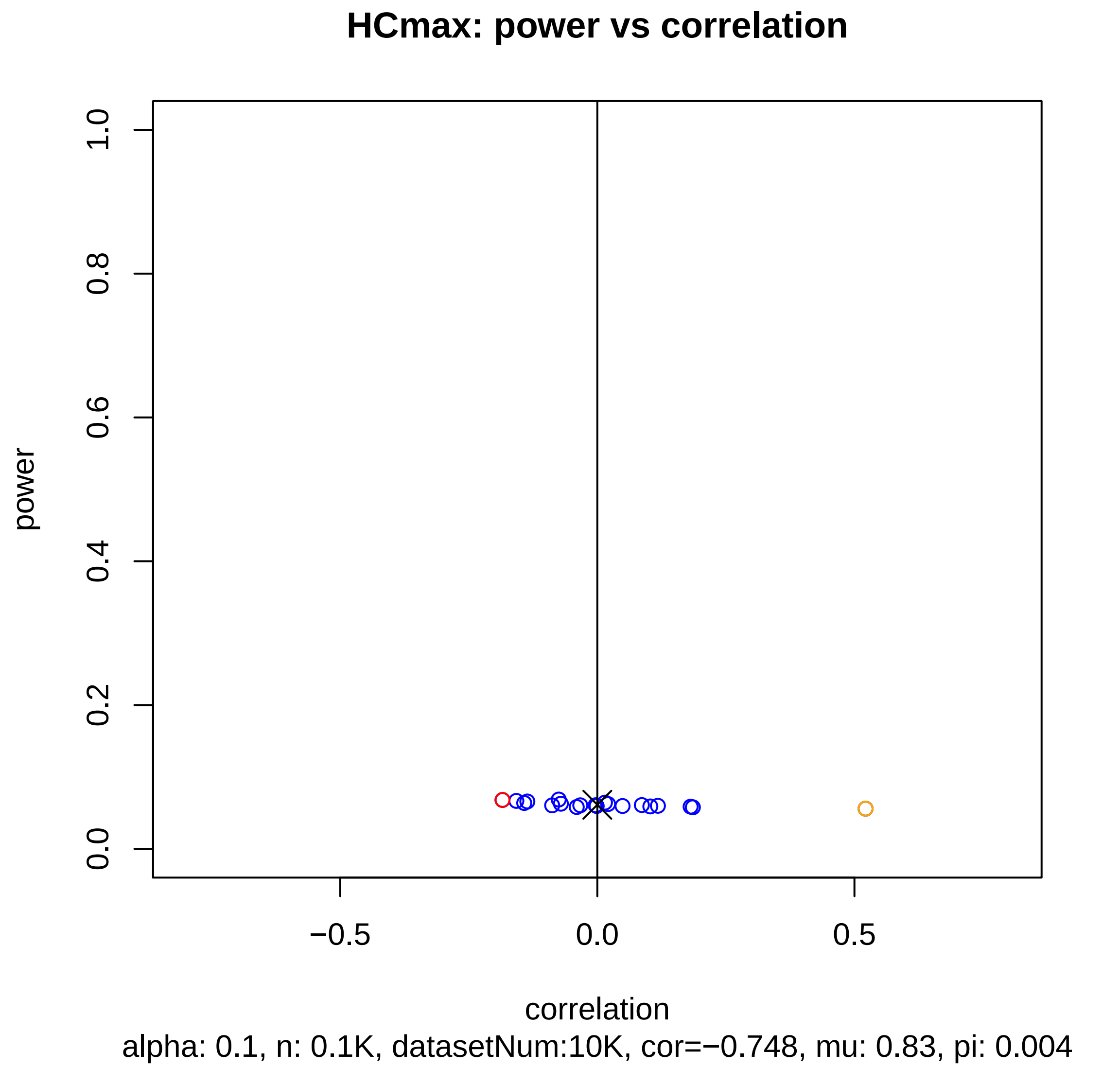

HCmax: power vs correlation
power
1.0
0.8
0.6
0.4
0.2
0.0
−0.5
0.0
0.5
correlation
alpha: 0.1, n: 0.1K, datasetNum:10K, cor=−0.748, mu: 0.83, pi: 0.004

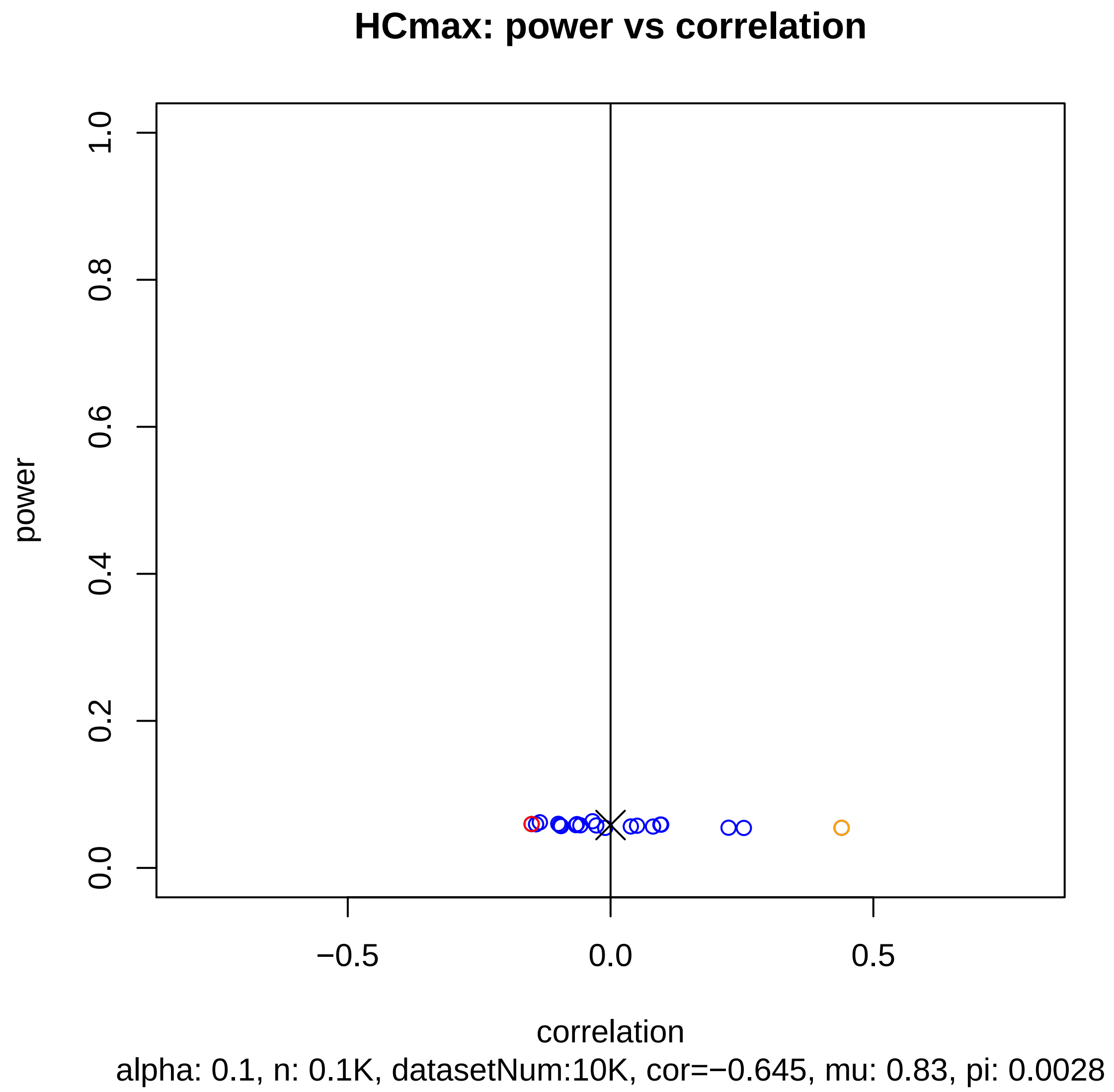

HCmax: power vs correlation
power
correlation
0.0
0.2
0.4
0.6
0.8
1.0
−0.5
0.0
0.5
alpha: 0.1, n: 0.1K, datasetNum:10K, cor=−0.645, mu: 0.83, pi: 0.0028

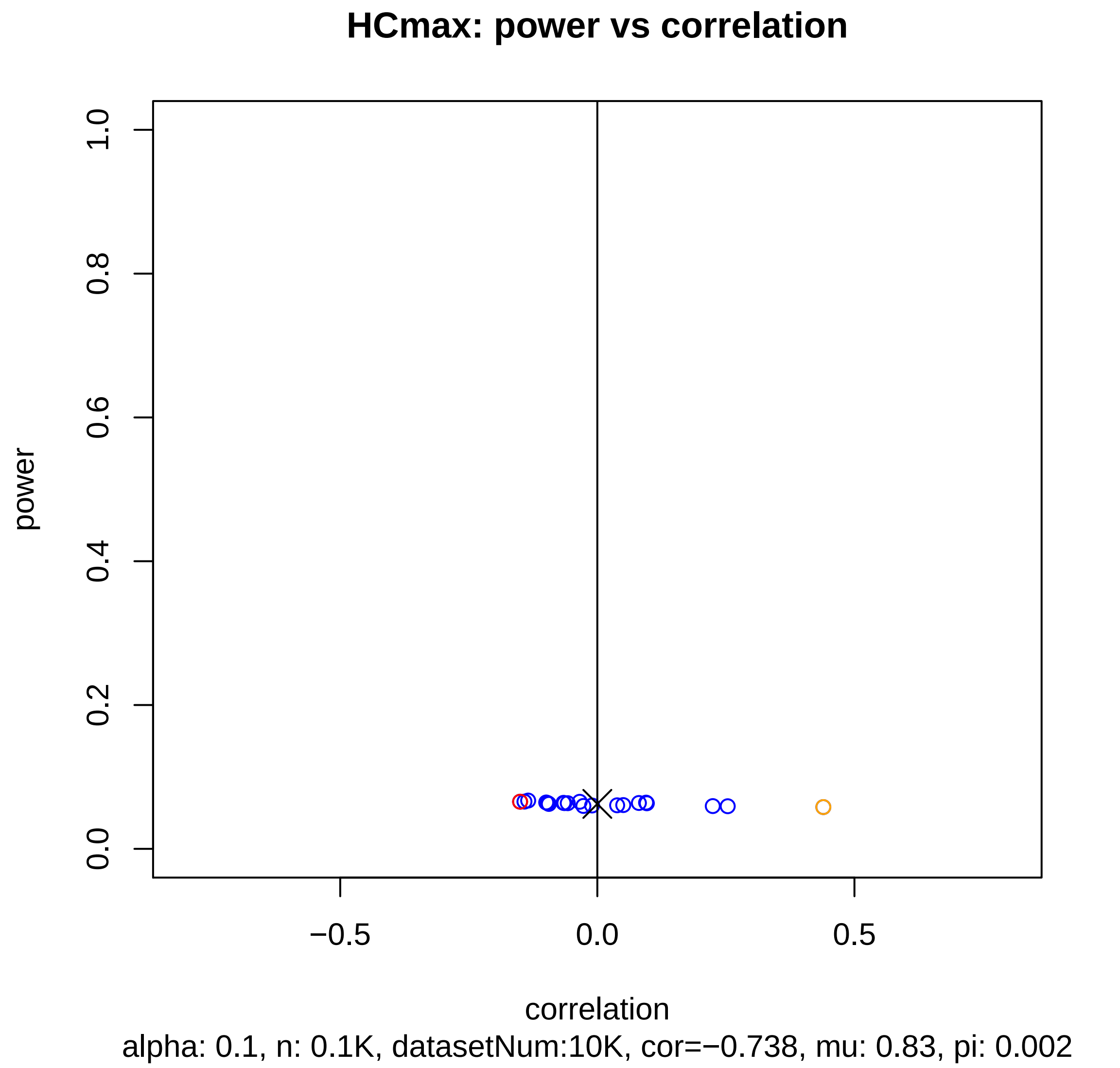
HCmax: power vs correlation
power
1.0
0.8
0.6
0.4
0.2
0.0
−0.5
0.0
0.5
correlation
alpha: 0.1, n: 0.1K, datasetNum:10K, cor=−0.738, mu: 0.83, pi: 0.002

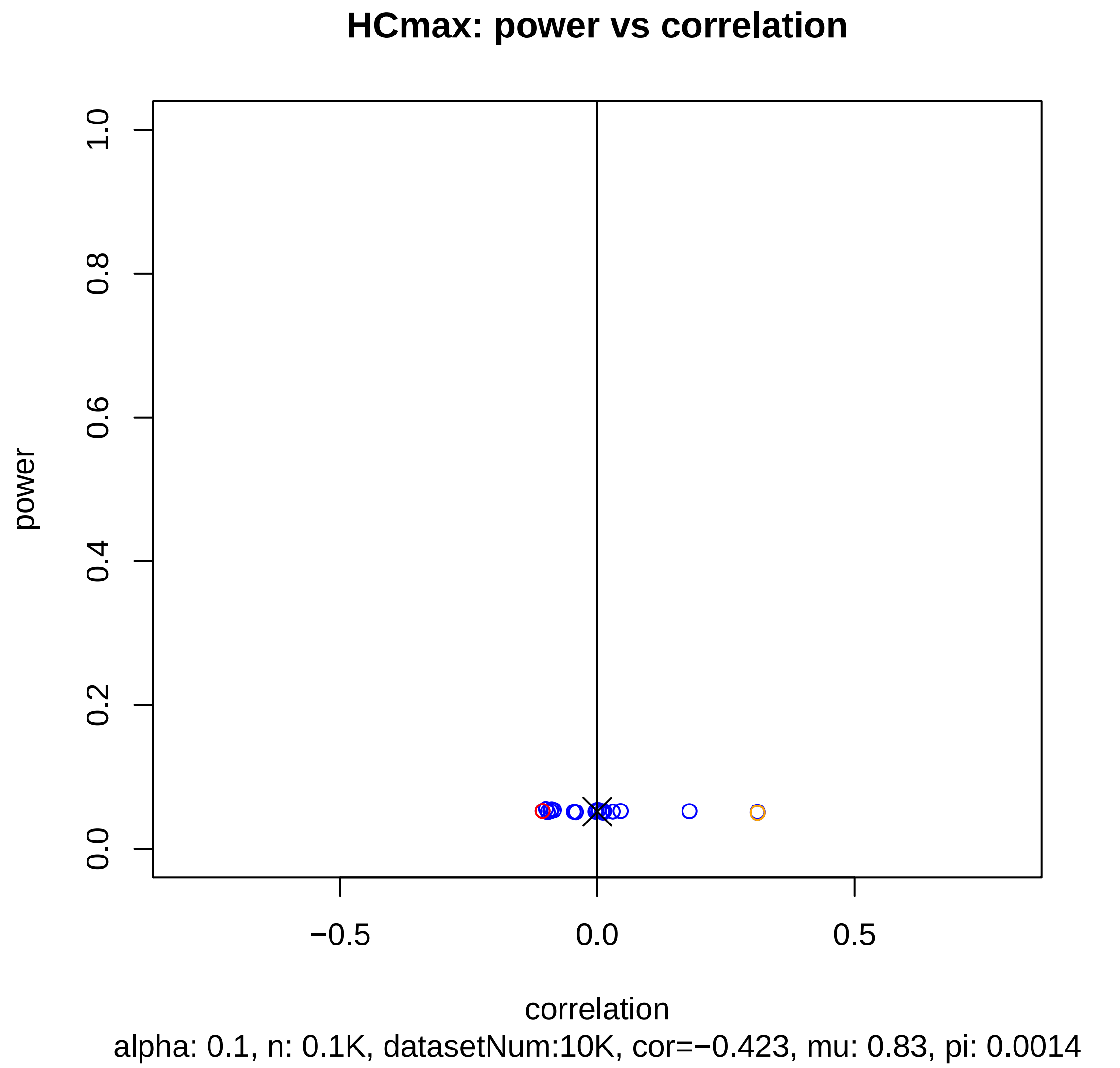

HCmax: power vs correlation
power
1.0
0.8
0.6
0.4
0.2
0.0
−0.5
0.0
0.5
correlation
alpha: 0.1, n: 0.1K, datasetNum:10K, cor=−0.423, mu: 0.83, pi: 0.0014

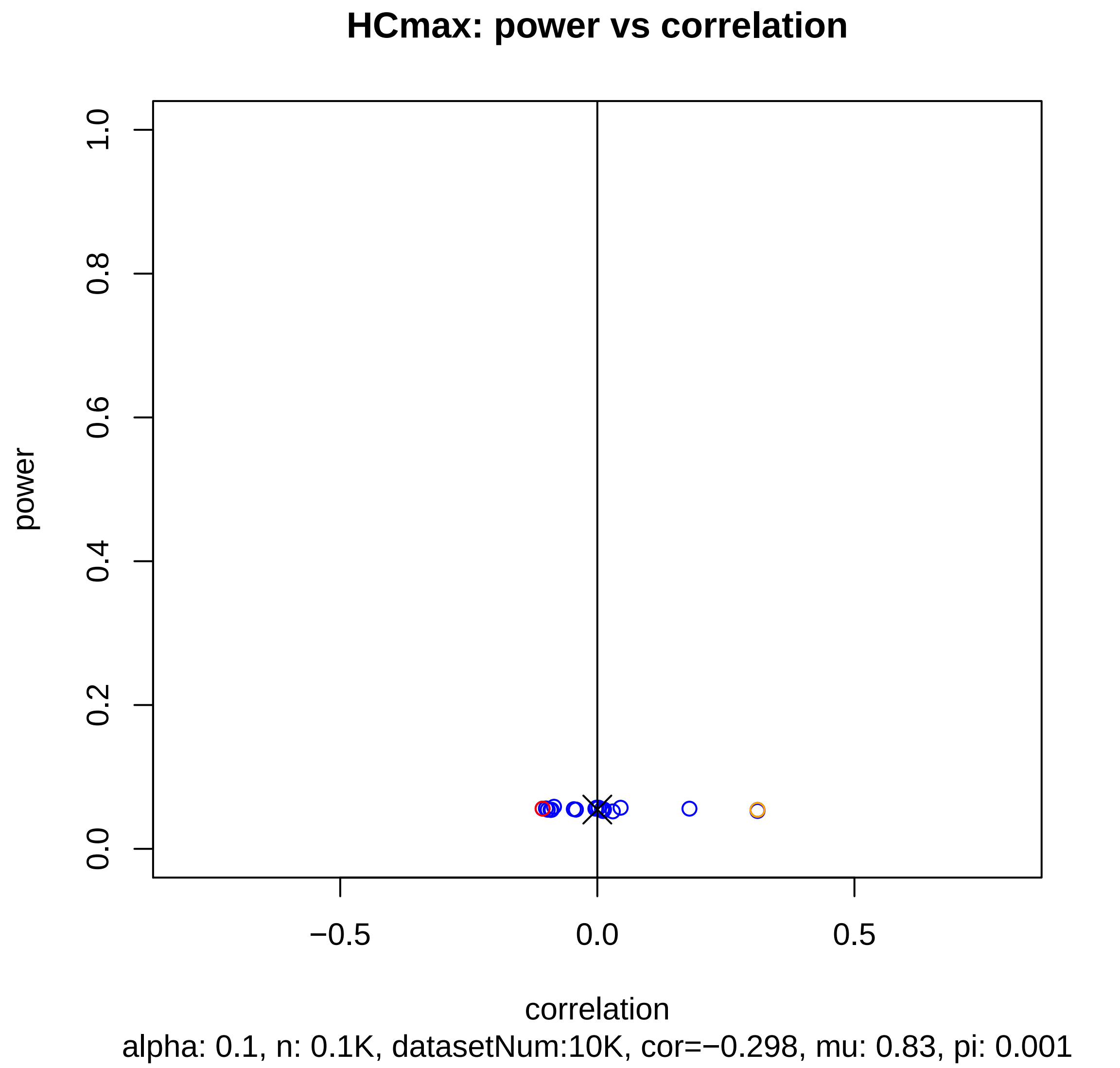
HCmax: power vs correlation
power
0.0
0.2
0.4
0.6
0.8
1.0
−0.5
0.0
0.5
correlation
alpha: 0.1, n: 0.1K, datasetNum:10K, cor=−0.298, mu: 0.83, pi: 0.001

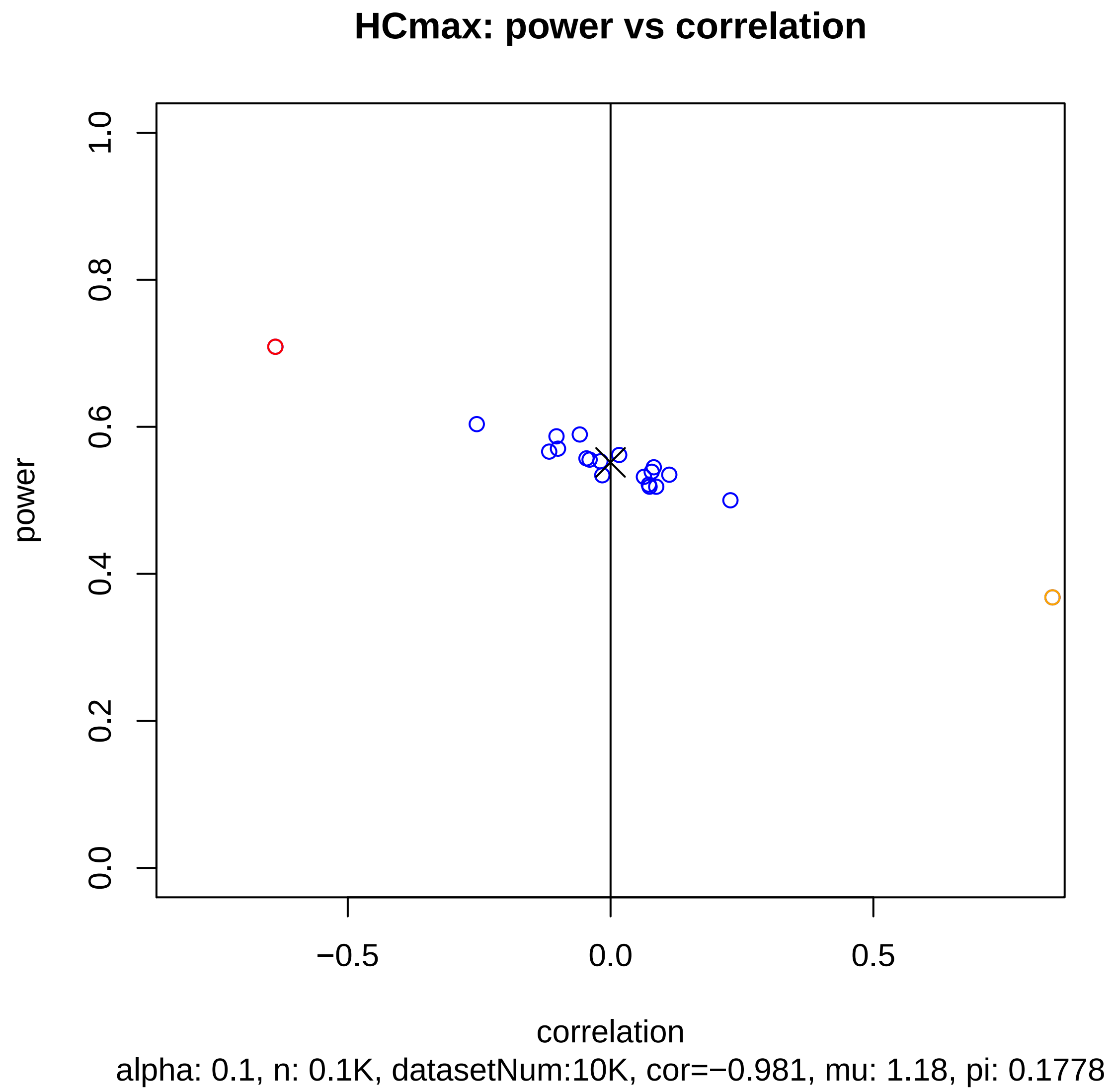
HCmax: power vs correlation
power
0.0
0.2
0.4
0.6
0.8
1.0
−0.5
0.0
0.5
correlation
alpha: 0.1, n: 0.1K, datasetNum:10K, cor=−0.981, mu: 1.18, pi: 0.1778

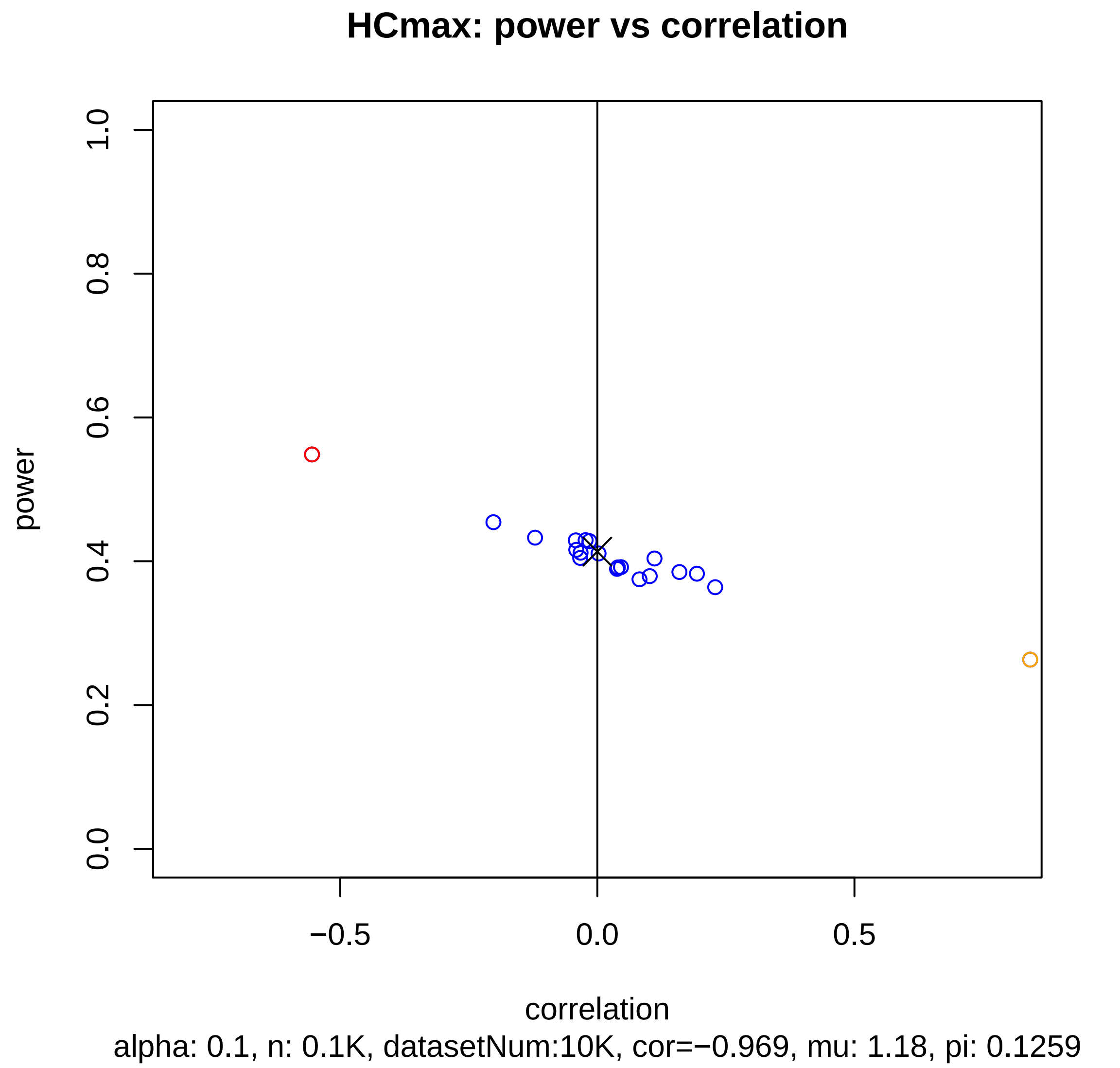
HCmax: power vs correlation
power
0.0
0.2
0.4
0.6
0.8
1.0
−0.5
0.0
0.5
correlation
alpha: 0.1, n: 0.1K, datasetNum:10K, cor=−0.969, mu: 1.18, pi: 0.1259

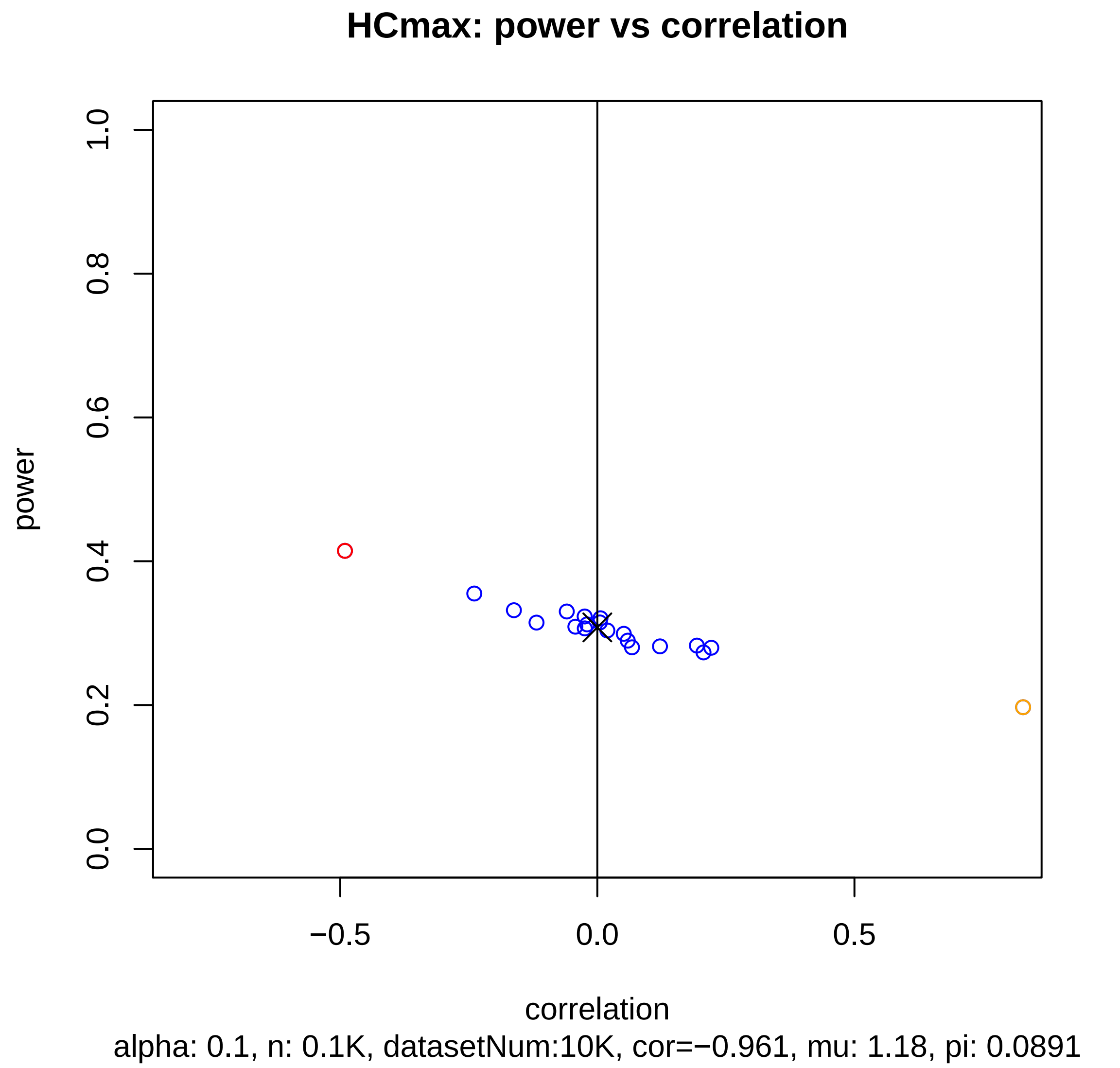

HCmax: power vs correlation
power
1.0
0.8
0.6
0.4
0.2
0.0
−0.5
0.0
0.5
correlation
alpha: 0.1, n: 0.1K, datasetNum:10K, cor=−0.961, mu: 1.18, pi: 0.0891

### HCmax: power vs correlation

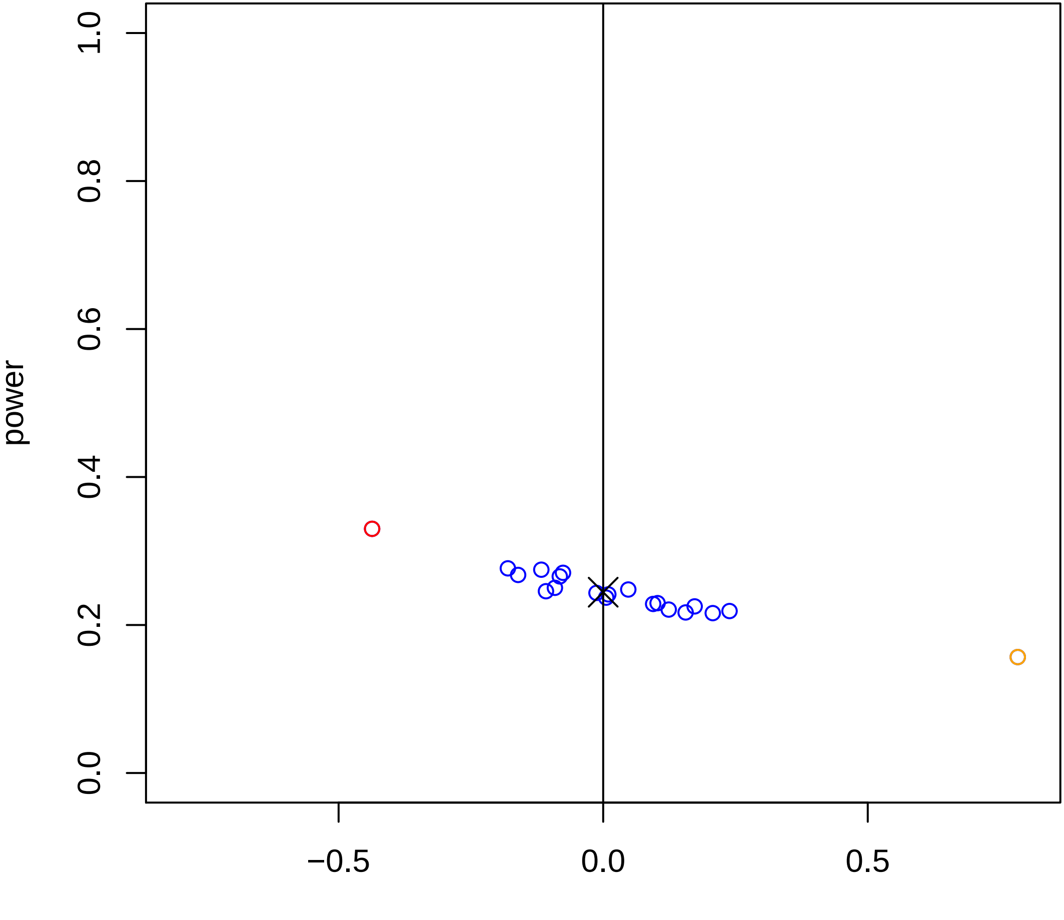


alpha: 0.1, n: 0.1K, datasetNum:10K, cor=−0.955, mu: 1.18, pi: 0.0631

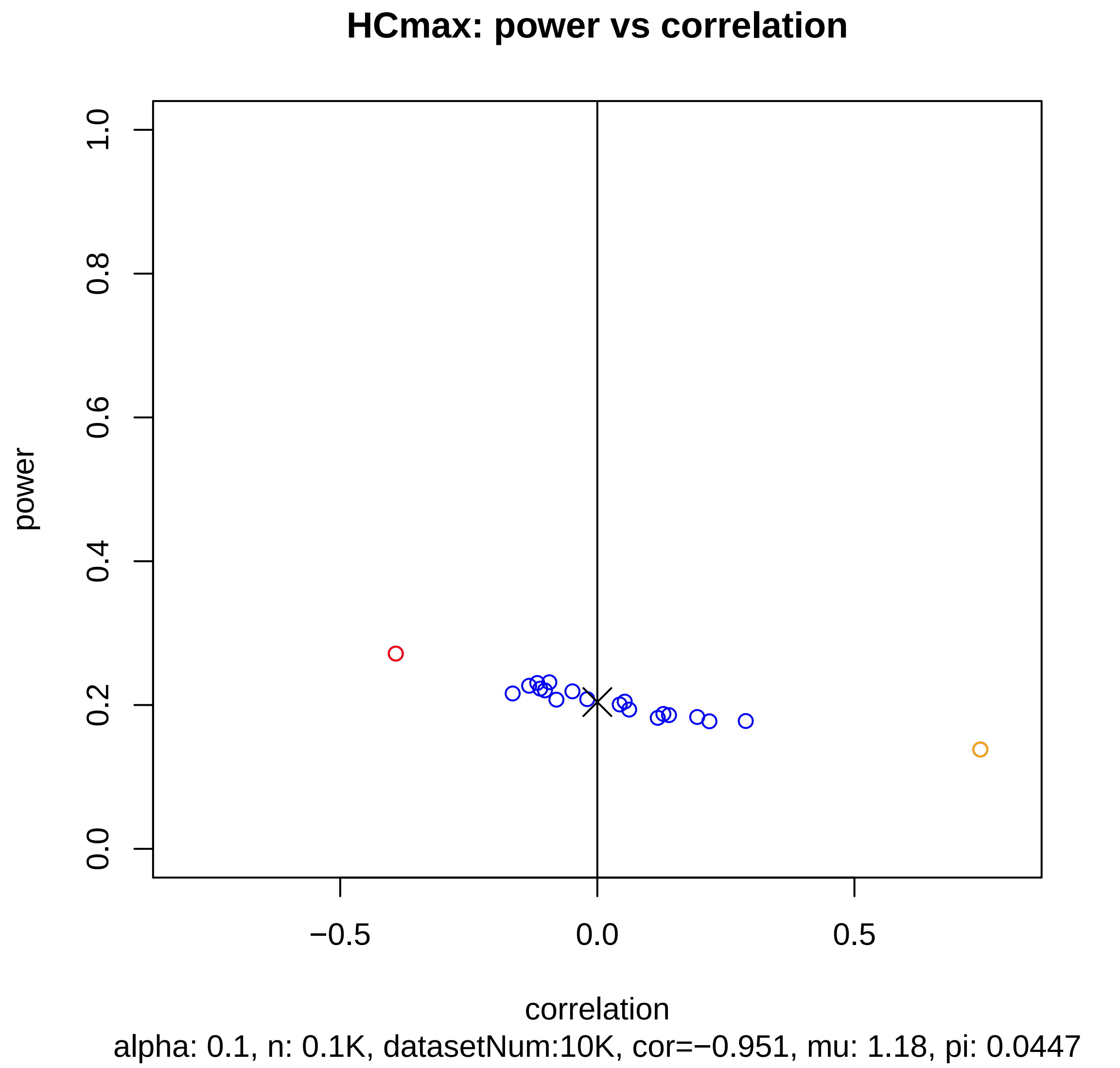
HCmax: power vs correlation
power
1.0
0.8
0.6
0.4
0.2
0.0
−0.5
0.0
0.5
correlation
alpha: 0.1, n: 0.1K, datasetNum:10K, cor=−0.951, mu: 1.18, pi: 0.0447

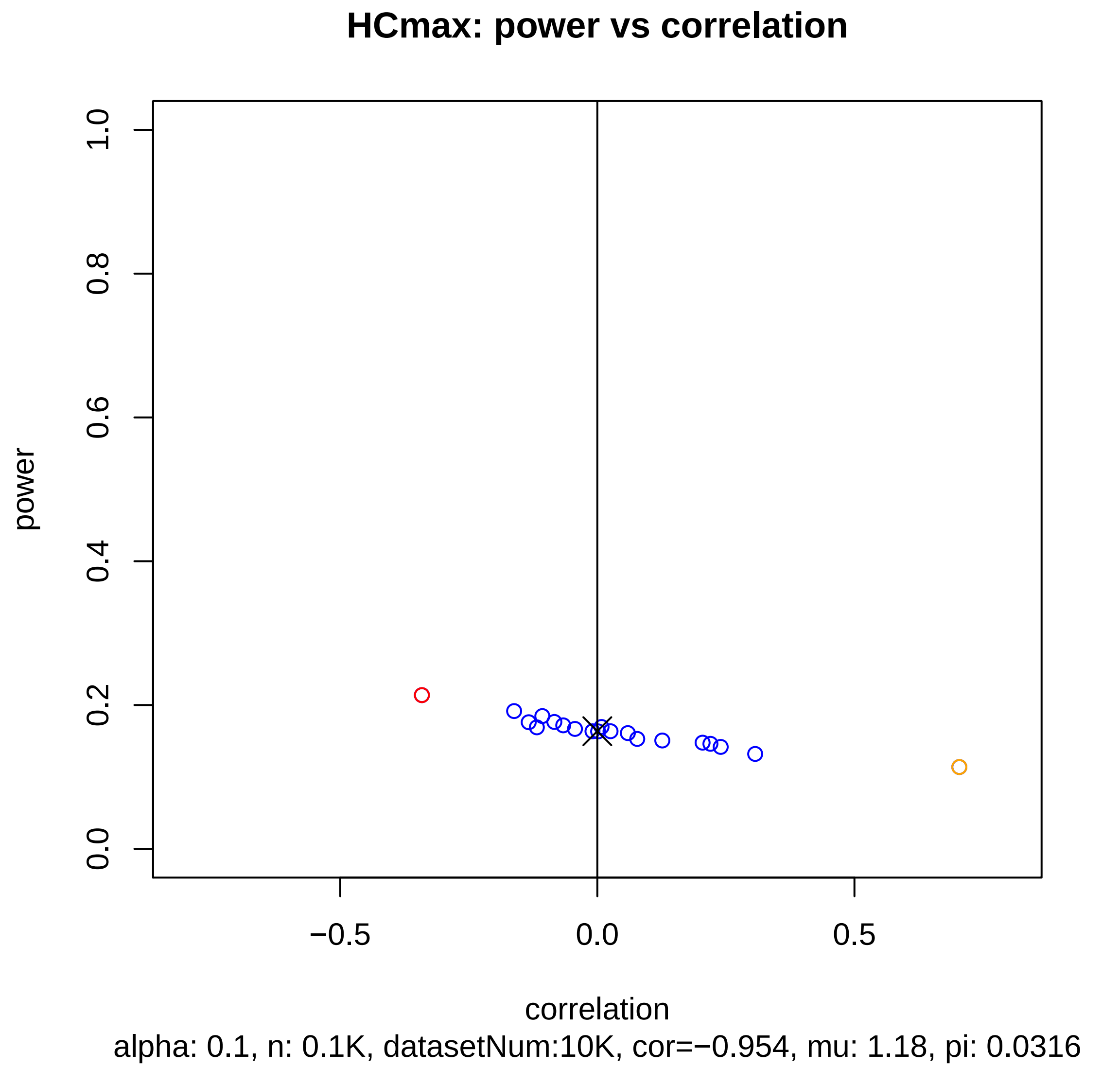

HCmax: power vs correlation
power
1.0
0.8
0.6
0.4
0.2
0.0
−0.5
0.0
0.5
correlation
alpha: 0.1, n: 0.1K, datasetNum:10K, cor=−0.954, mu: 1.18, pi: 0.0316

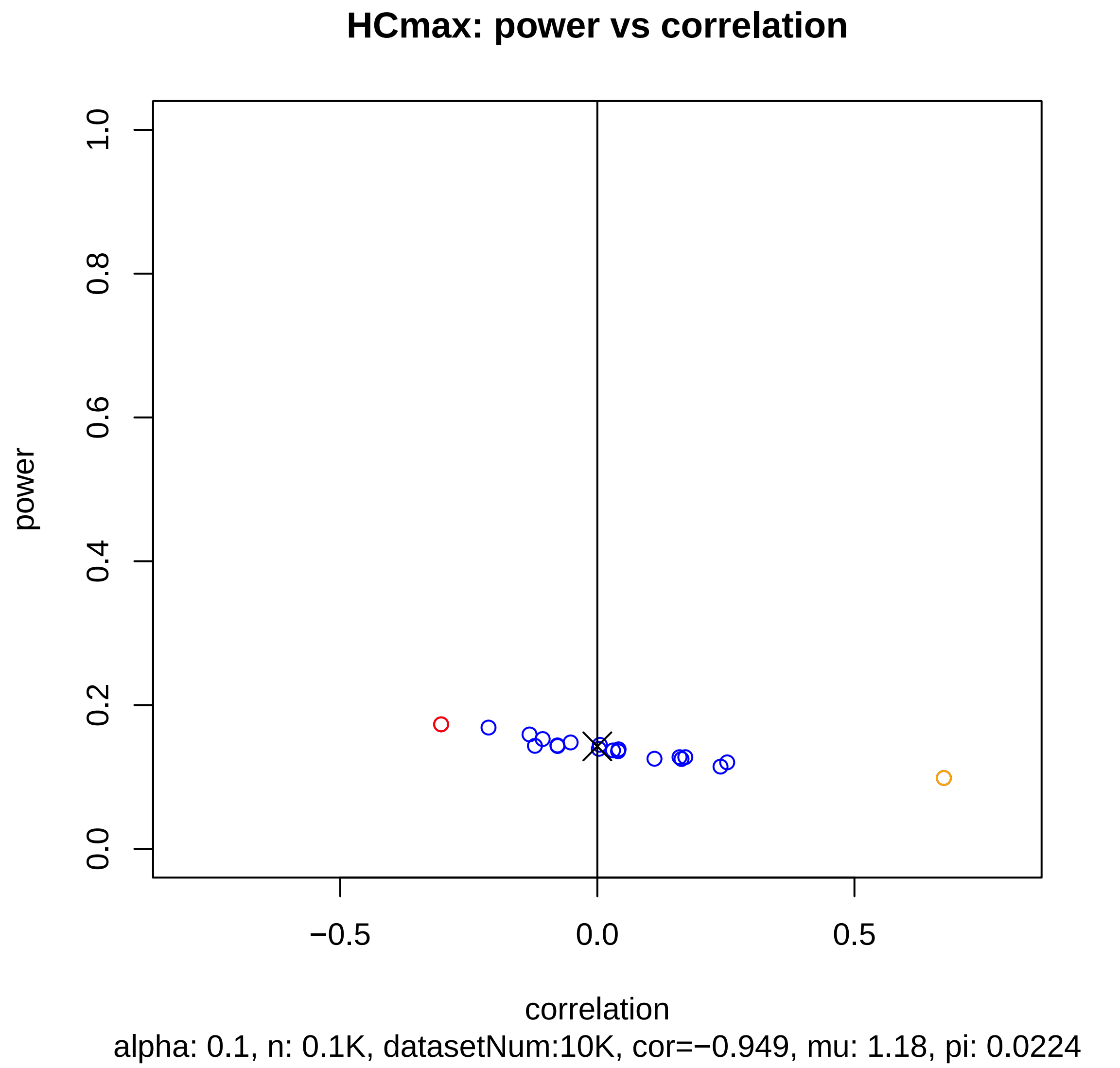
HCmax: power vs correlation
power
0.0
0.2
0.4
0.6
0.8
1.0
−0.5
0.0
0.5
correlation
alpha: 0.1, n: 0.1K, datasetNum:10K, cor=−0.949, mu: 1.18, pi: 0.0224

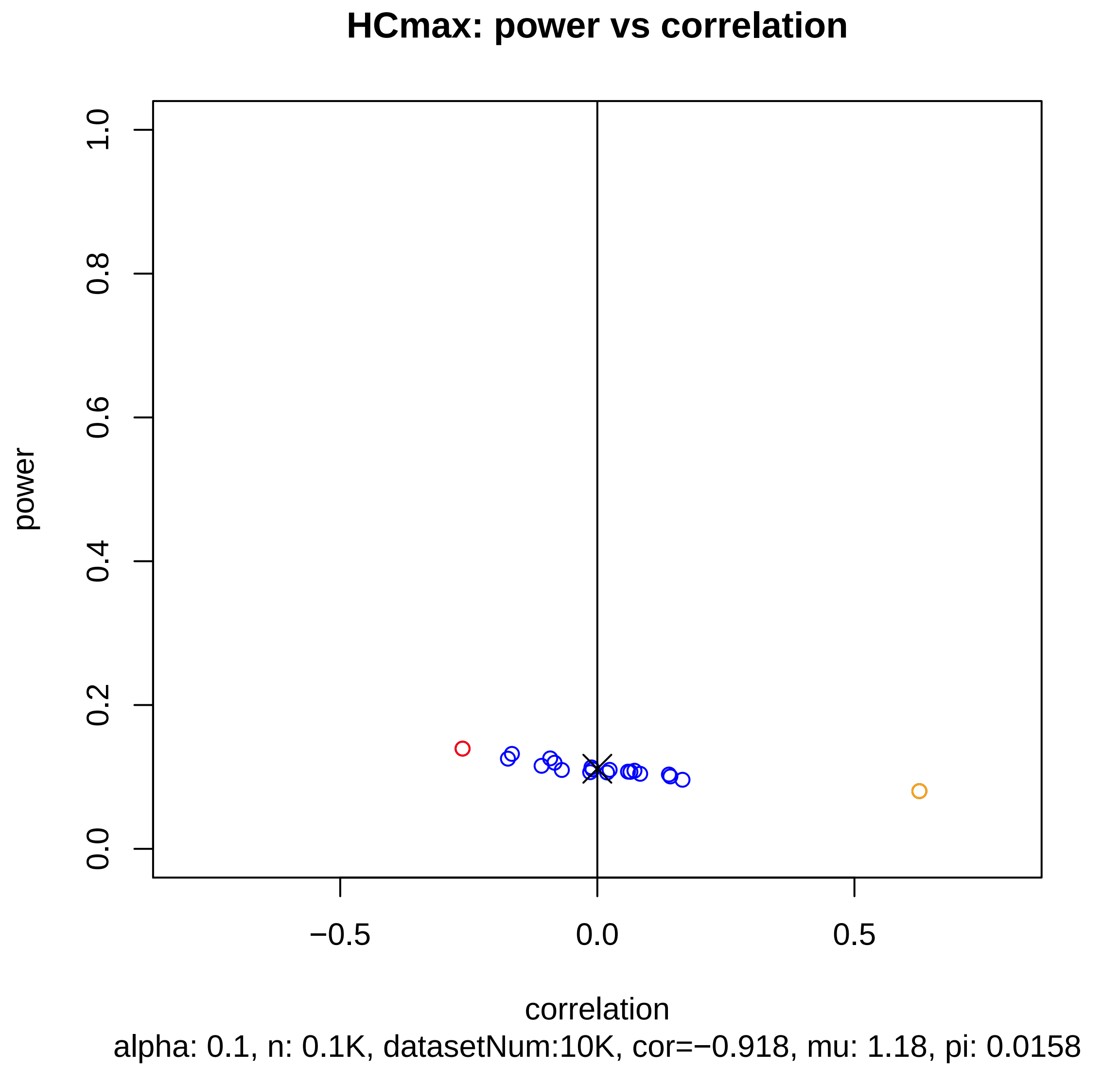
HCmax: power vs correlation
power
1.0
0.8
0.6
0.4
0.2
0.0
−0.5
0.0
0.5
correlation
alpha: 0.1, n: 0.1K, datasetNum:10K, cor=−0.918, mu: 1.18, pi: 0.0158

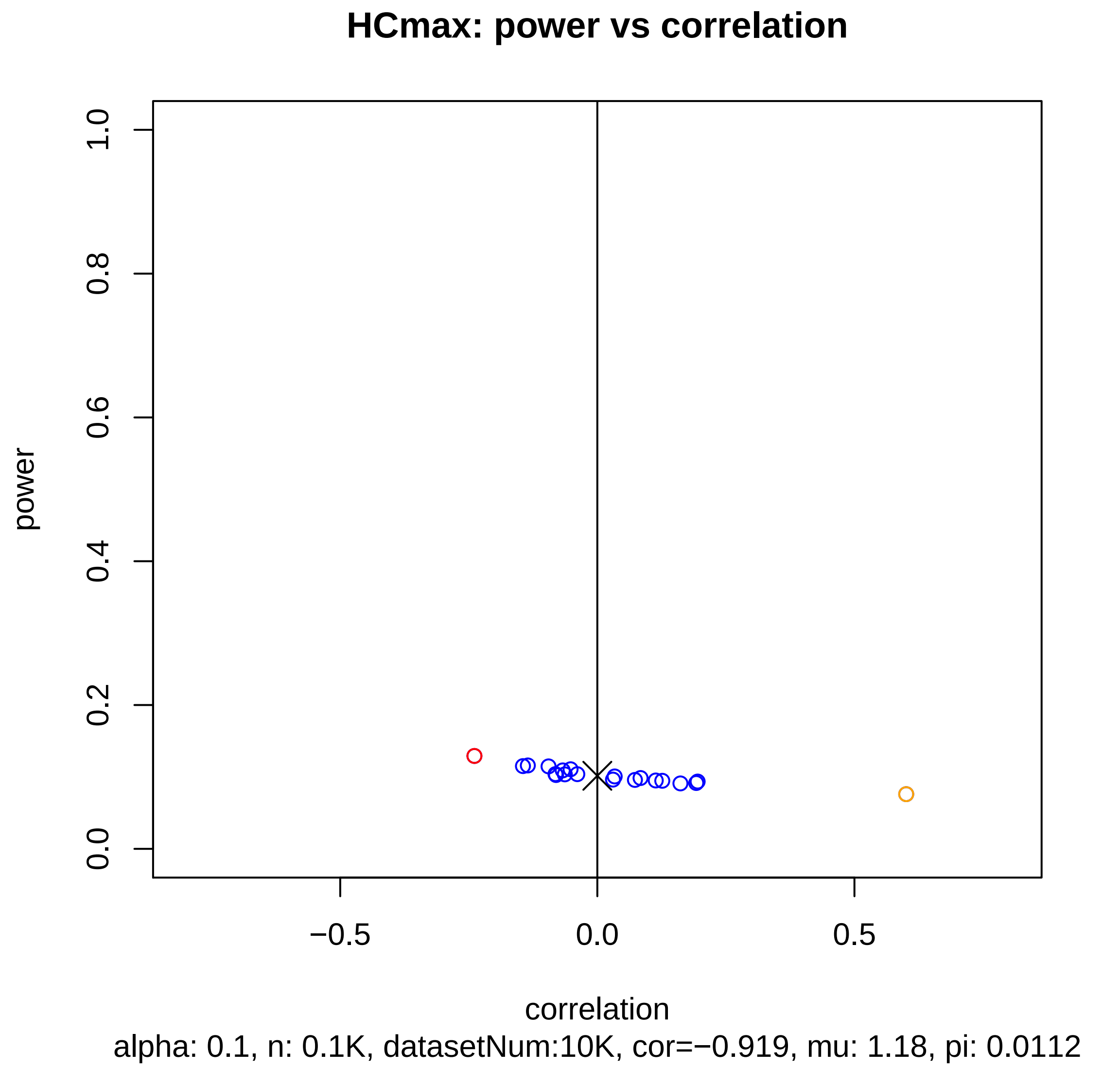
HCmax: power vs correlation
power
1.0
0.8
0.6
0.4
0.2
0.0
−0.5
0.0
0.5
correlation
alpha: 0.1, n: 0.1K, datasetNum:10K, cor=−0.919, mu: 1.18, pi: 0.0112

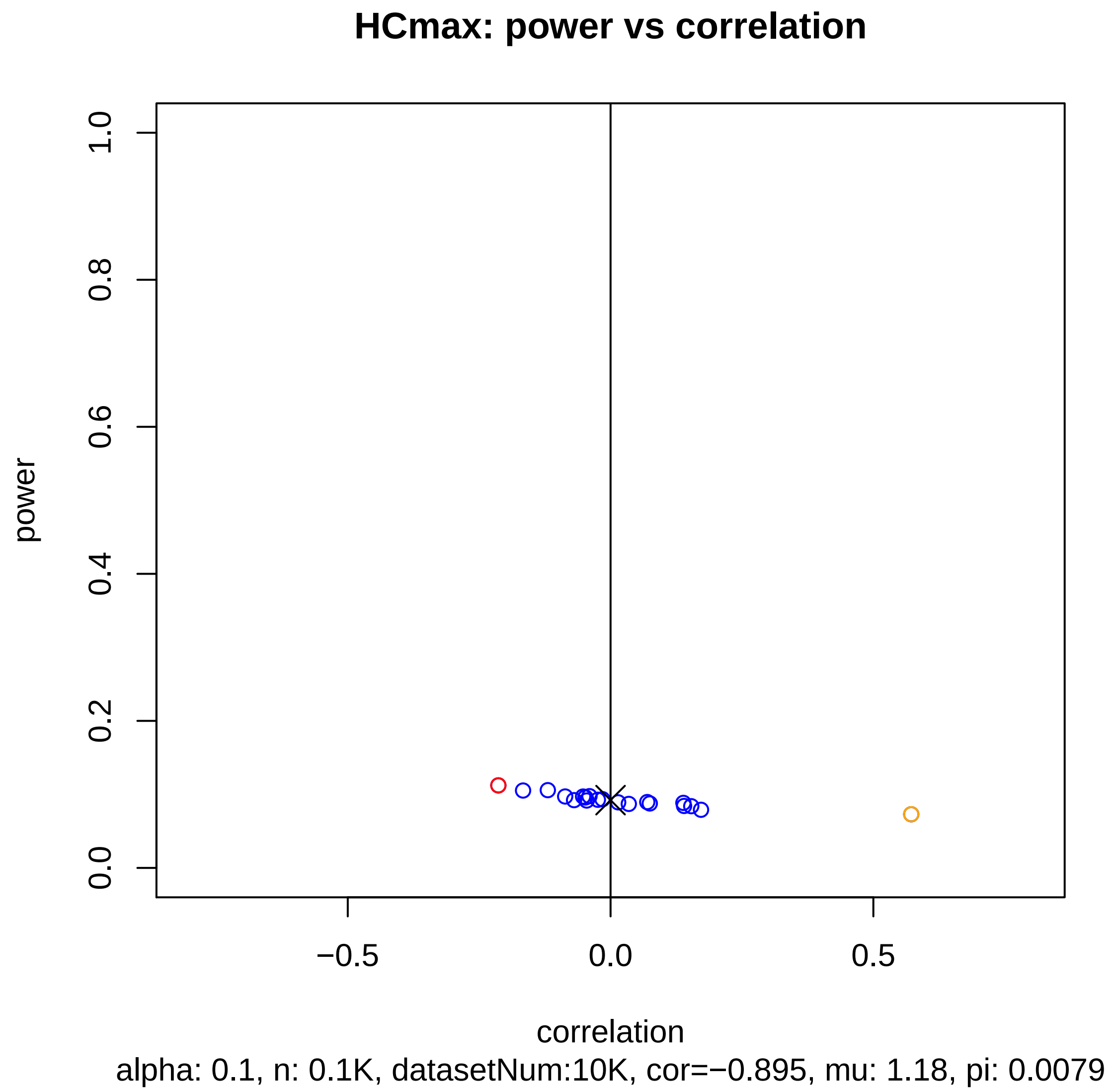

HCmax: power vs correlation
power
1.0
0.8
0.6
0.4
0.2
0.0
−0.5
0.0
0.5
correlation
alpha: 0.1, n: 0.1K, datasetNum:10K, cor=−0.895, mu: 1.18, pi: 0.0079

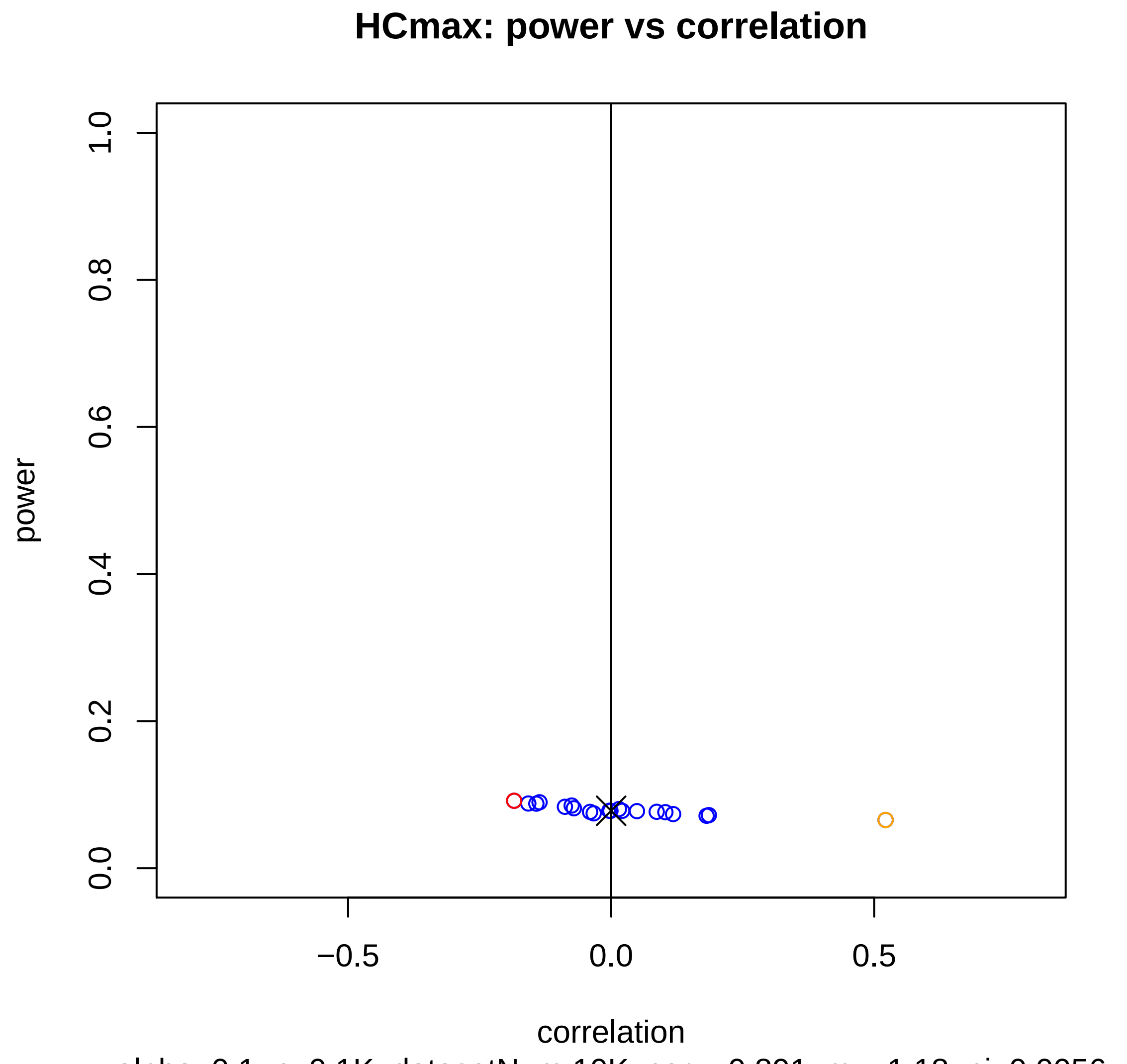

alpha: 0.1, n: 0.1K, datasetNum:10K, cor=−0.891, mu: 1.18, pi: 0.0056

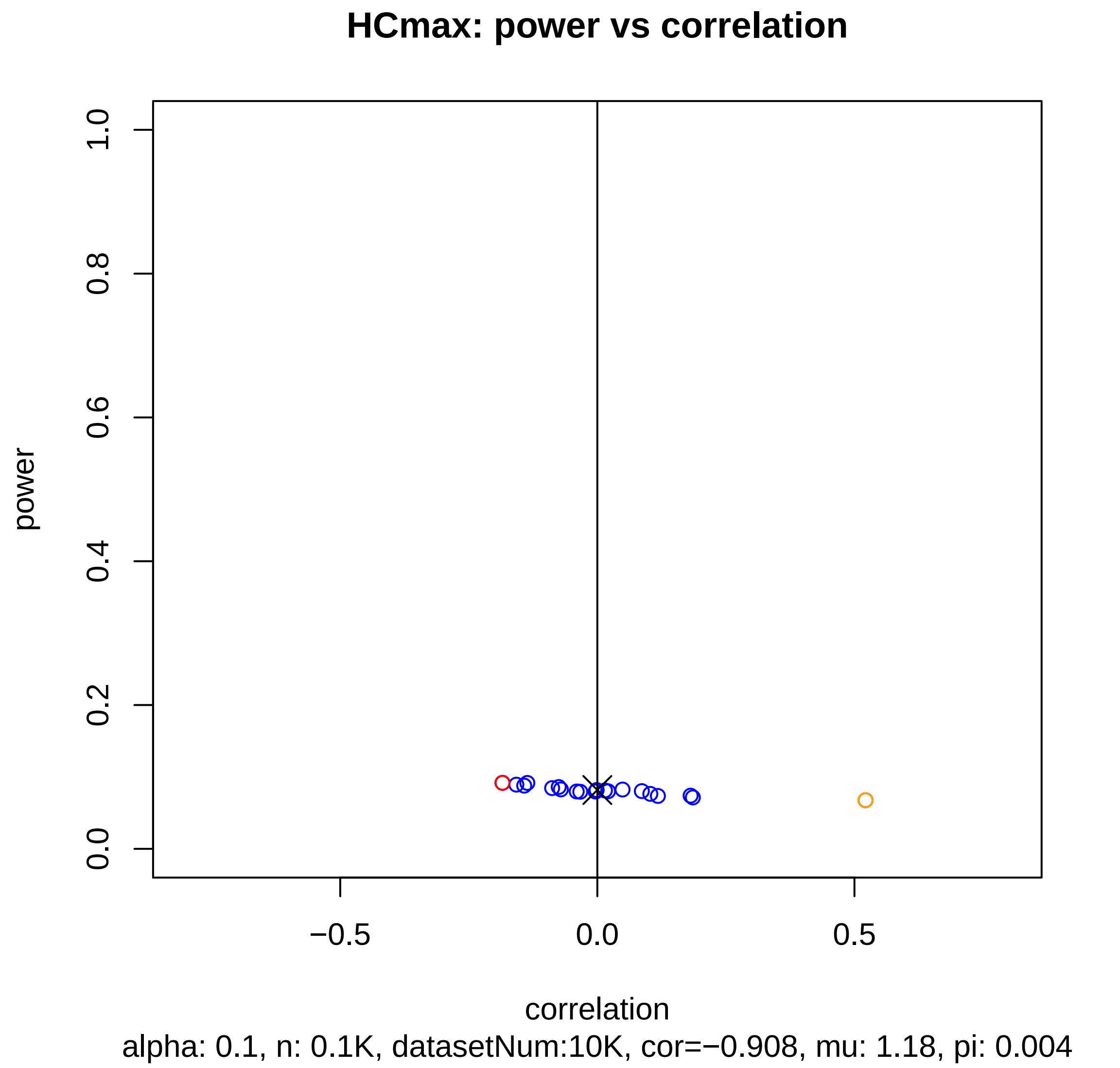
HCmax: power vs correlation
power
0.0
0.2
0.4
0.6
0.8
1.0
−0.5
0.0
0.5
correlation
alpha: 0.1, n: 0.1K, datasetNum:10K, cor=−0.908, mu: 1.18, pi: 0.004

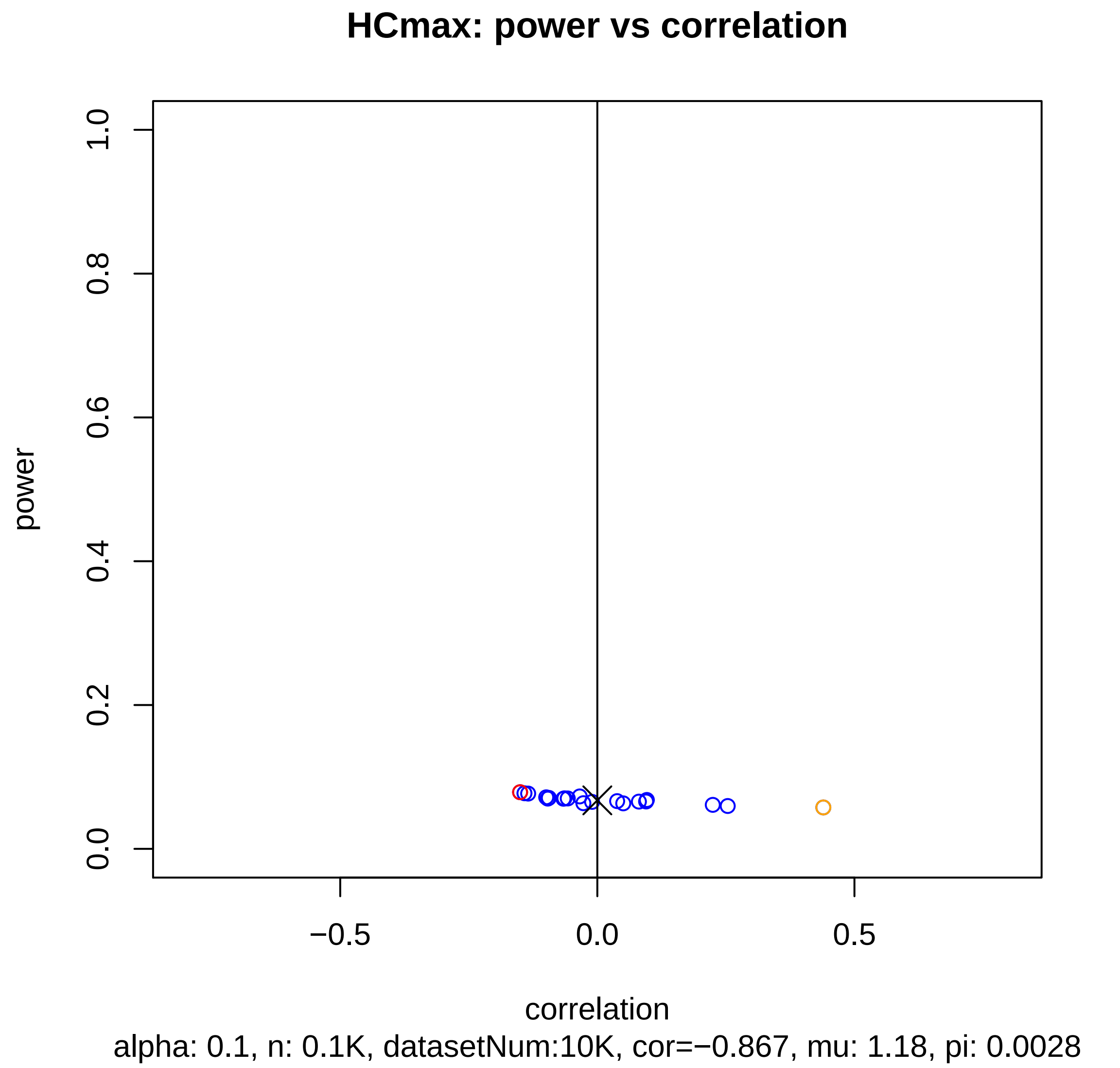
HCmax: power vs correlation
power
1.0
0.8
0.6
0.4
0.2
0.0
−0.5
0.0
0.5
correlation
alpha: 0.1, n: 0.1K, datasetNum:10K, cor=−0.867, mu: 1.18, pi: 0.0028

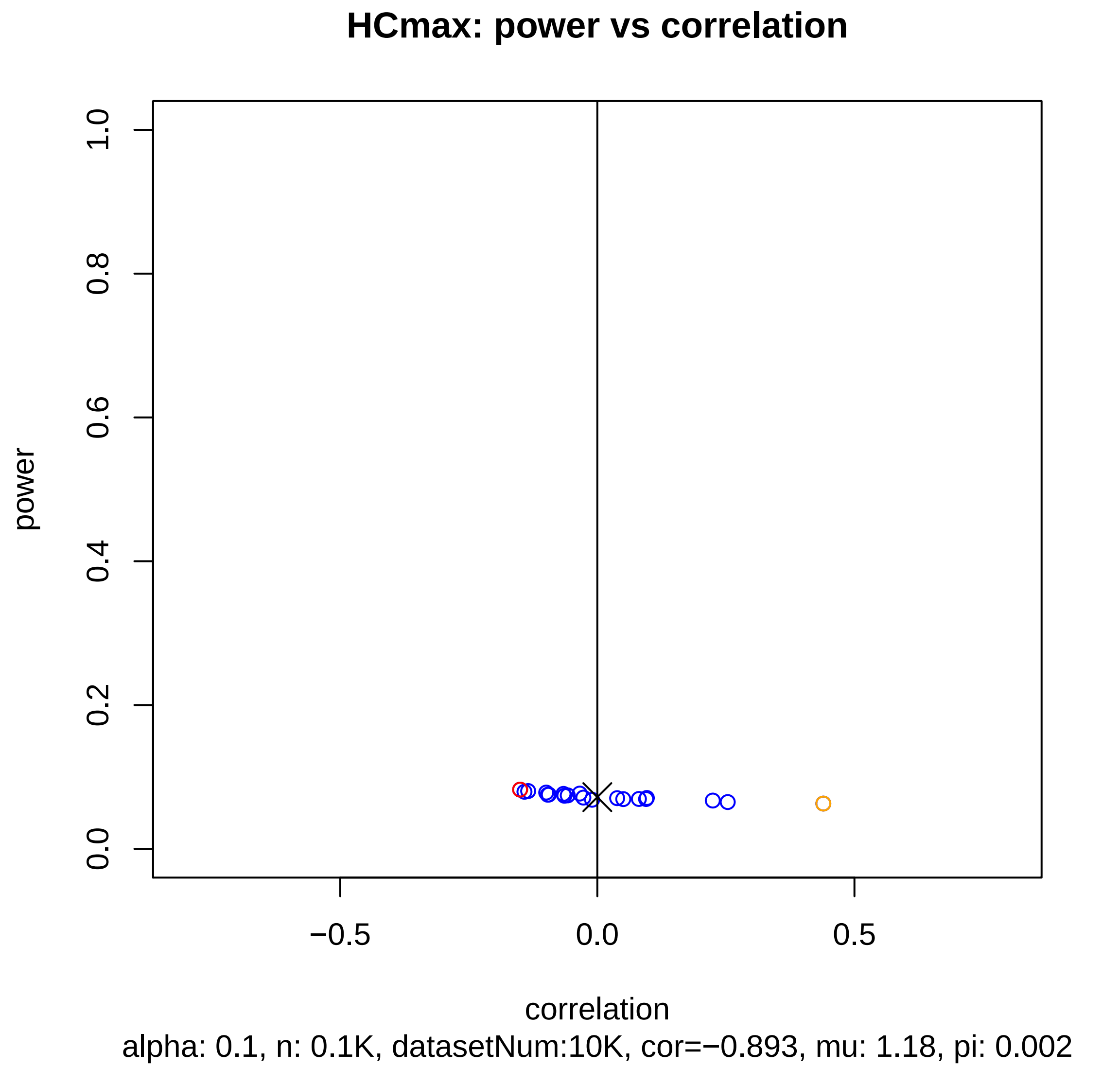
HCmax: power vs correlation
power
0.0
0.2
0.4
0.6
0.8
1.0
−0.5
0.0
0.5
correlation
alpha: 0.1, n: 0.1K, datasetNum:10K, cor=−0.893, mu: 1.18, pi: 0.002

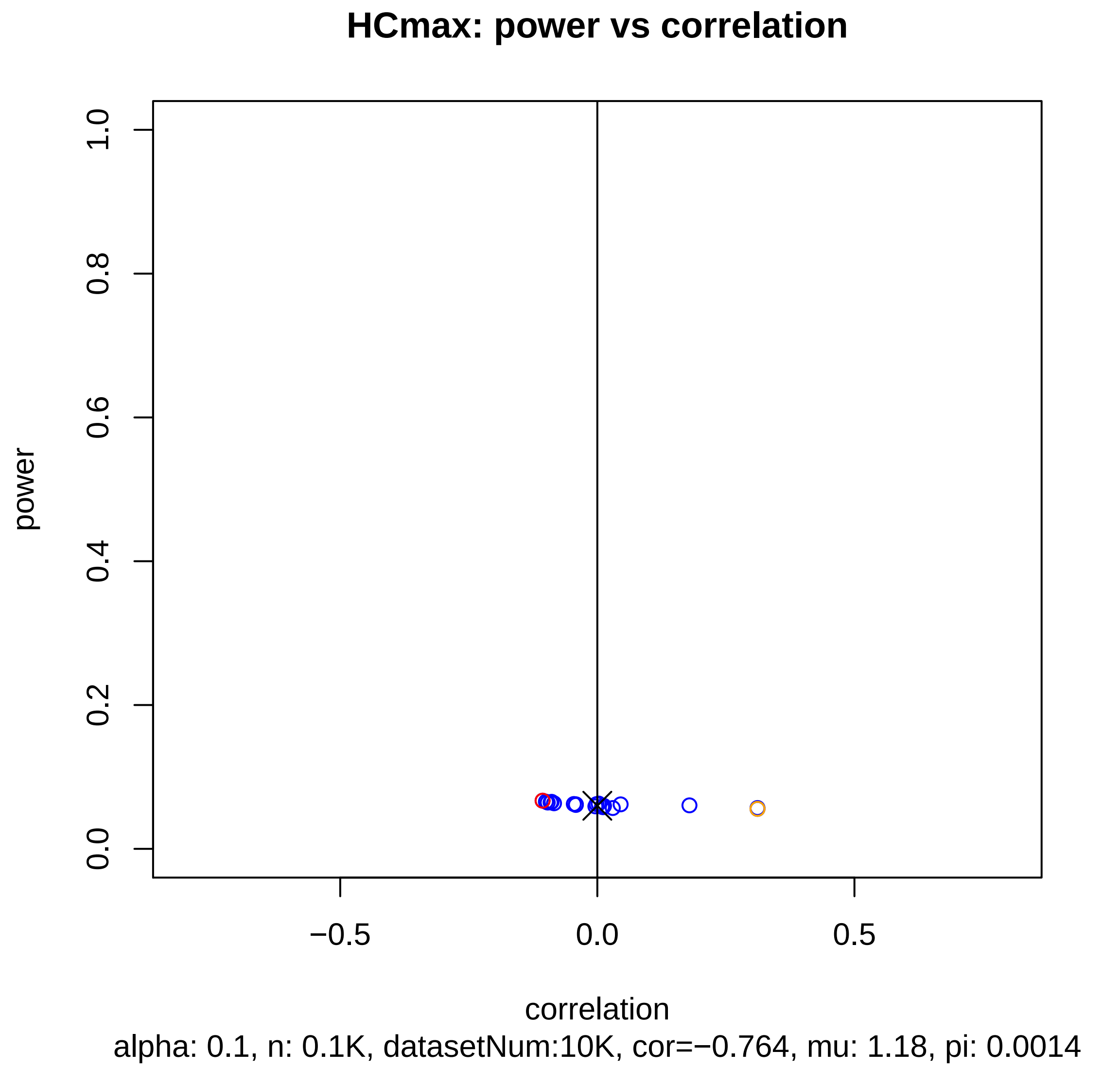
HCmax: power vs correlation
power
1.0
0.8
0.6
0.4
0.2
0.0
−0.5
0.0
0.5
correlation
alpha: 0.1, n: 0.1K, datasetNum:10K, cor=−0.764, mu: 1.18, pi: 0.0014

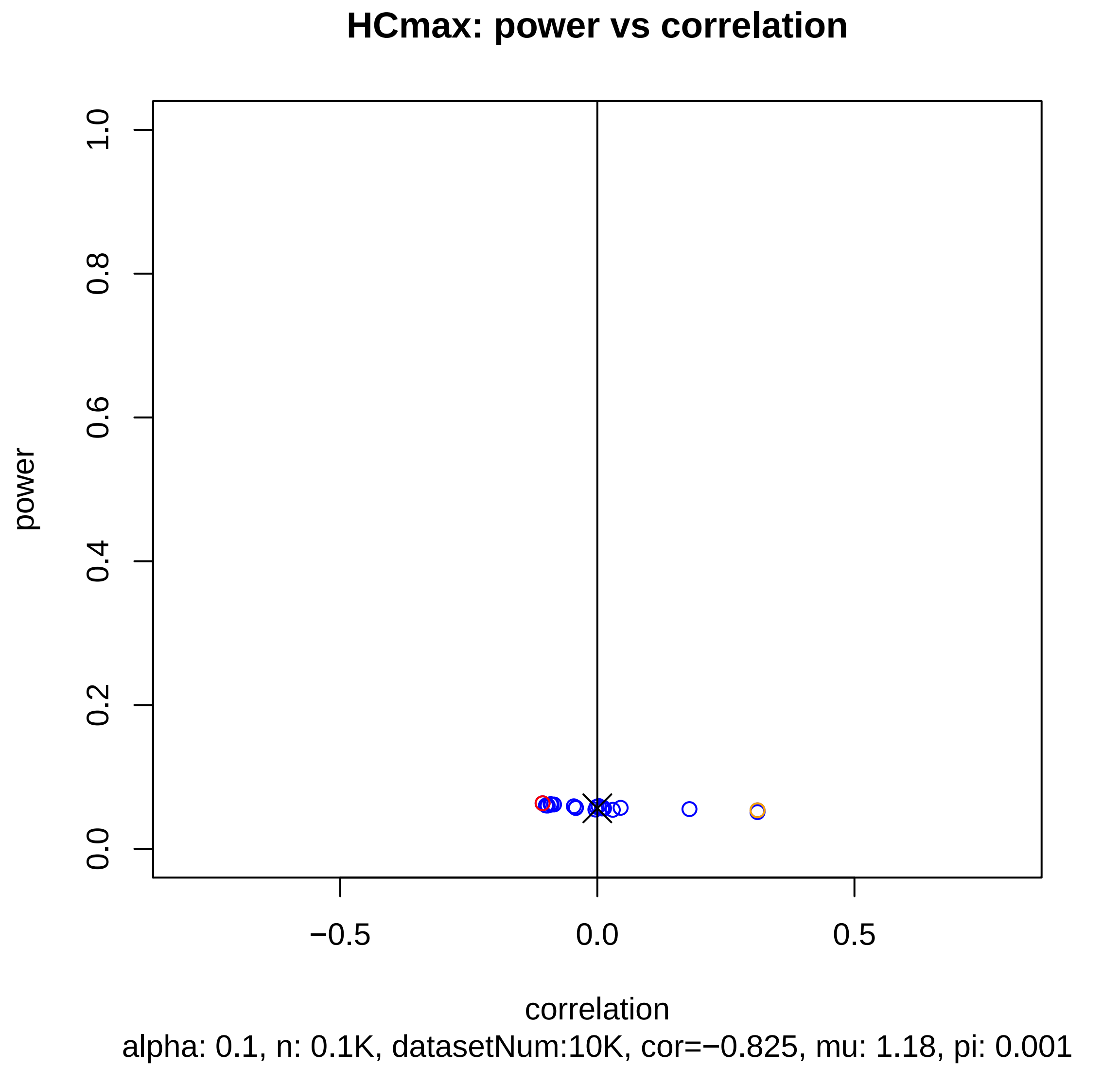
HCmax: power vs correlation
power
0.0
0.2
0.4
0.6
0.8
1.0
−0.5
0.0
0.5
correlation
alpha: 0.1, n: 0.1K, datasetNum:10K, cor=−0.825, mu: 1.18, pi: 0.001

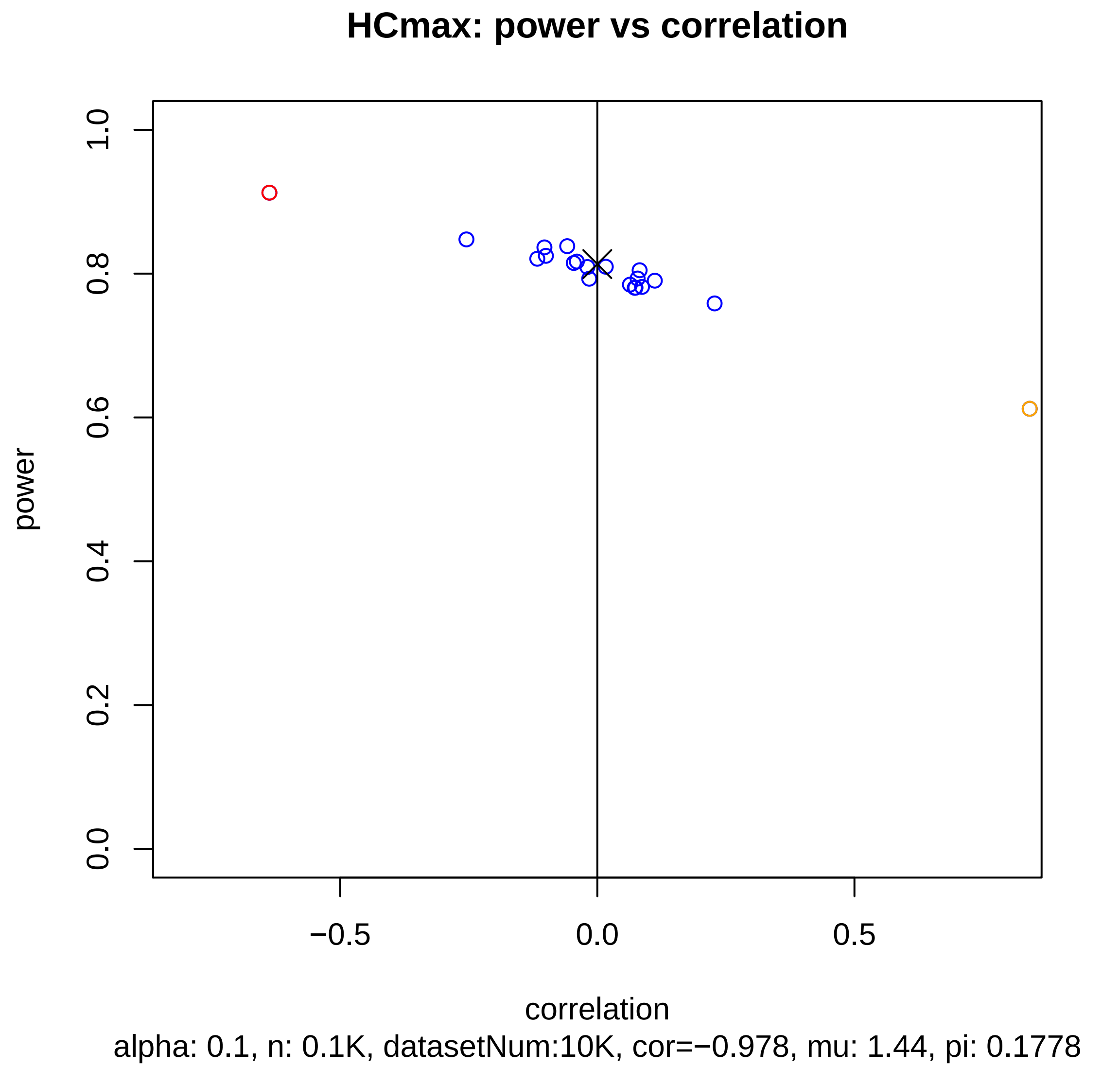

HCmax: power vs correlation
power
correlation
0.0
0.2
0.4
0.6
0.8
1.0
−0.5
0.0
0.5
alpha: 0.1, n: 0.1K, datasetNum:10K, cor=−0.978, mu: 1.44, pi: 0.1778

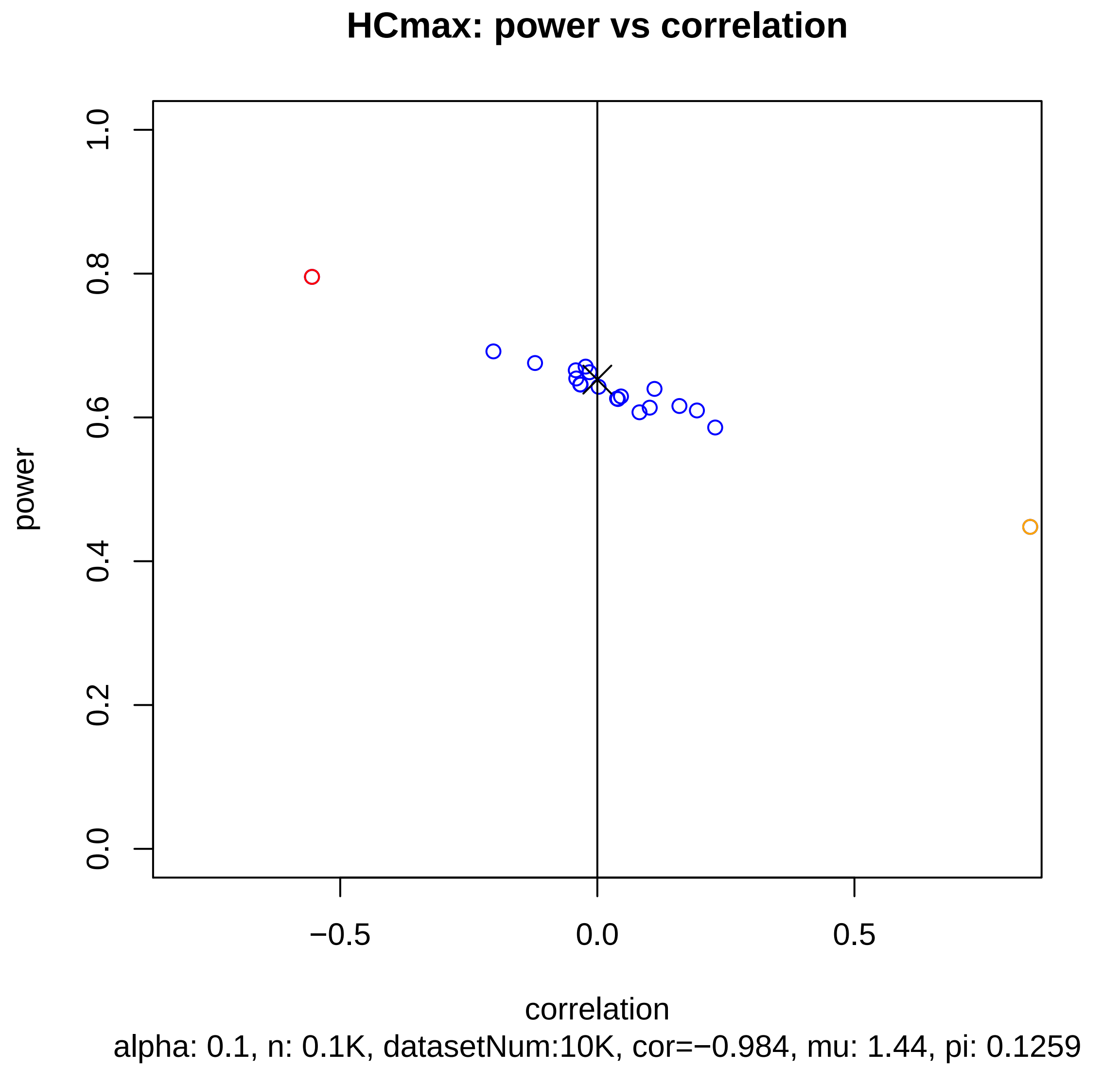
HCmax: power vs correlation
power
1.0
0.8
0.6
0.4
0.2
0.0
−0.5
0.0
0.5
correlation
alpha: 0.1, n: 0.1K, datasetNum:10K, cor=−0.984, mu: 1.44, pi: 0.1259

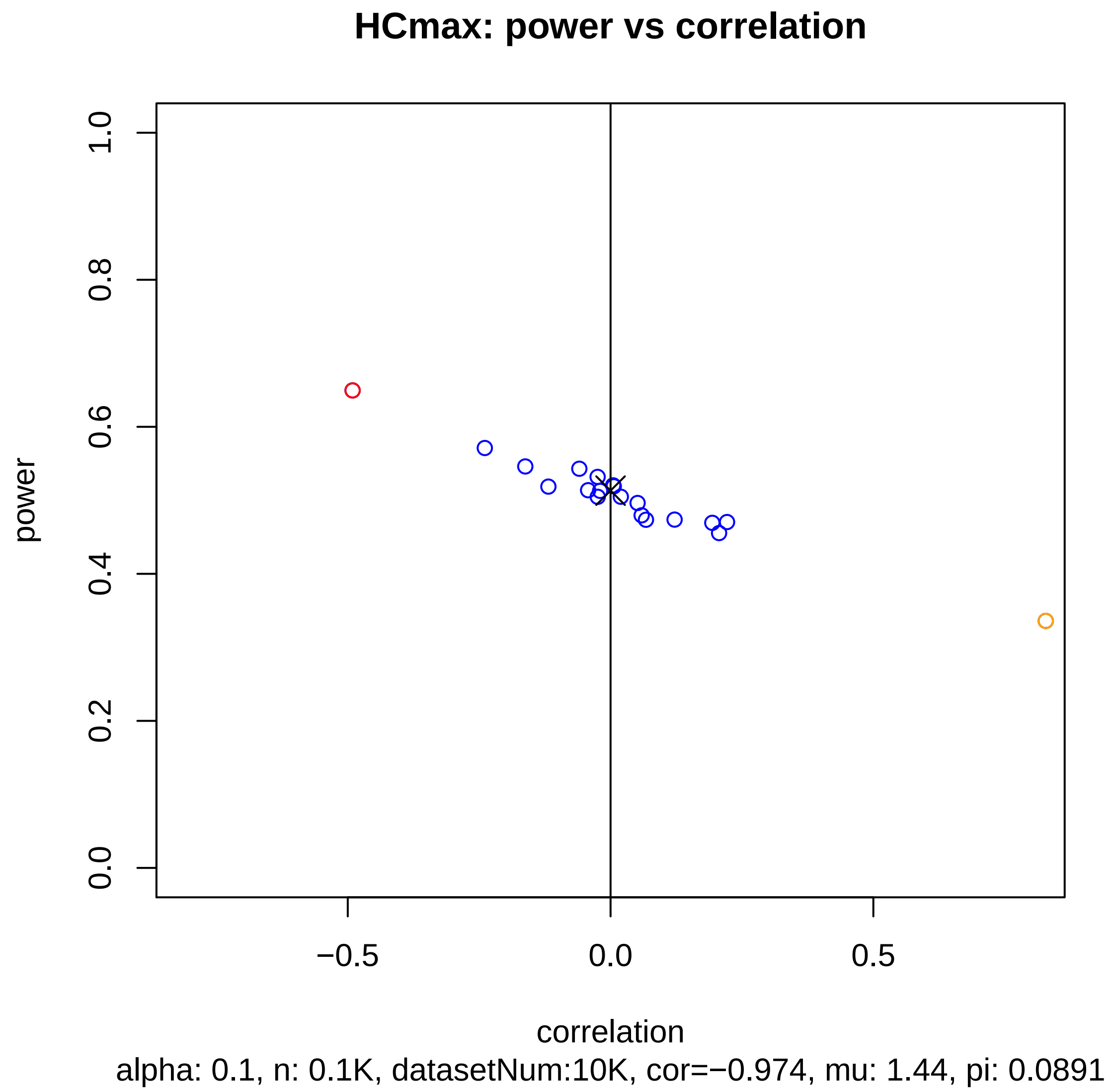
HCmax: power vs correlation
power
0.0
0.2
0.4
0.6
0.8
1.0
−0.5
0.0
0.5
correlation
alpha: 0.1, n: 0.1K, datasetNum:10K, cor=−0.974, mu: 1.44, pi: 0.0891

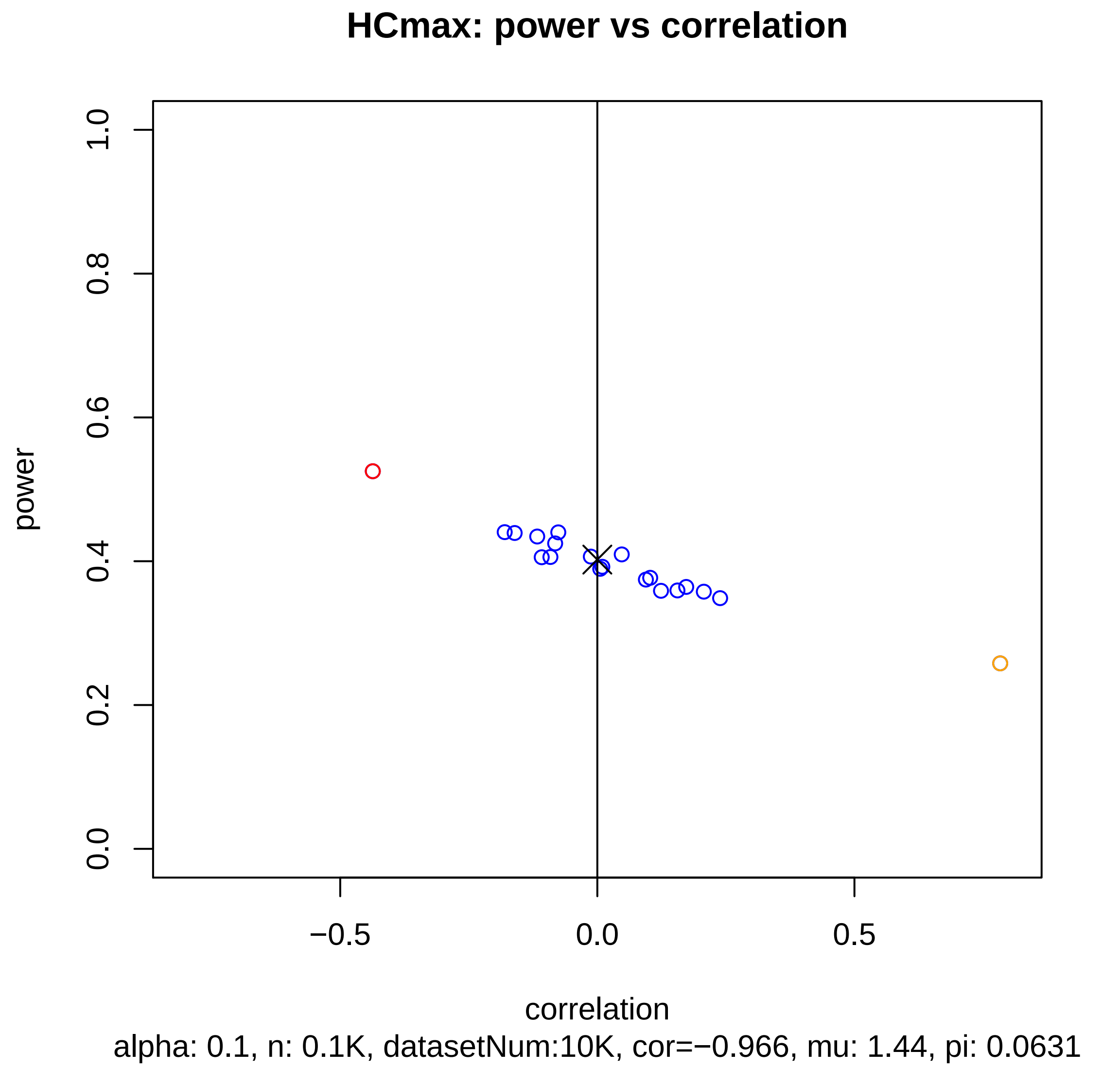
HCmax: power vs correlation
power
0.0
0.2
0.4
0.6
0.8
1.0
−0.5
0.0
0.5
correlation
alpha: 0.1, n: 0.1K, datasetNum:10K, cor=−0.966, mu: 1.44, pi: 0.0631

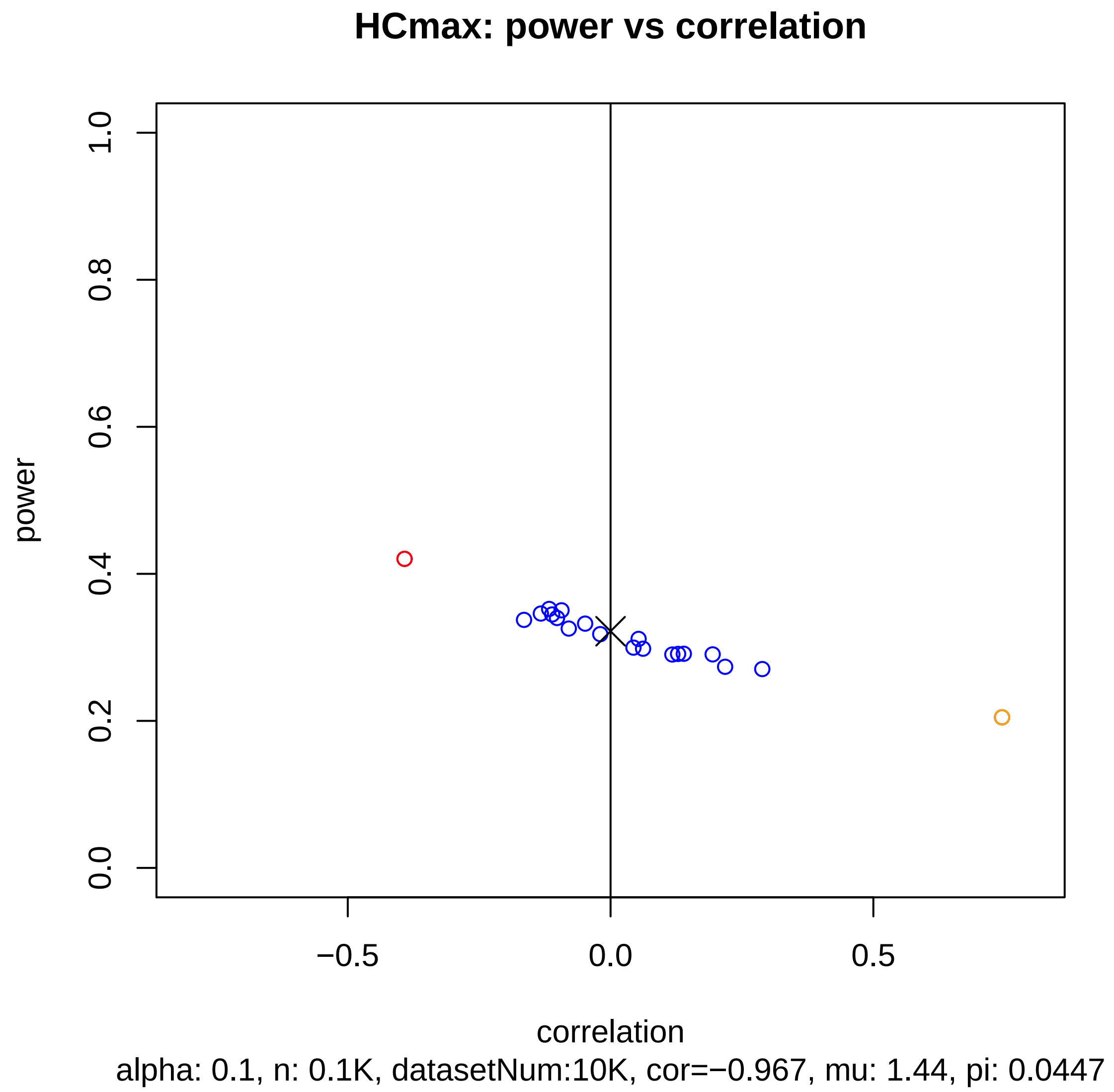
HCmax: power vs correlation
power
correlation
1.0
0.8
0.6
0.4
0.2
0.0
−0.5
0.0
0.5
alpha: 0.1, n: 0.1K, datasetNum:10K, cor=−0.967, mu: 1.44, pi: 0.0447

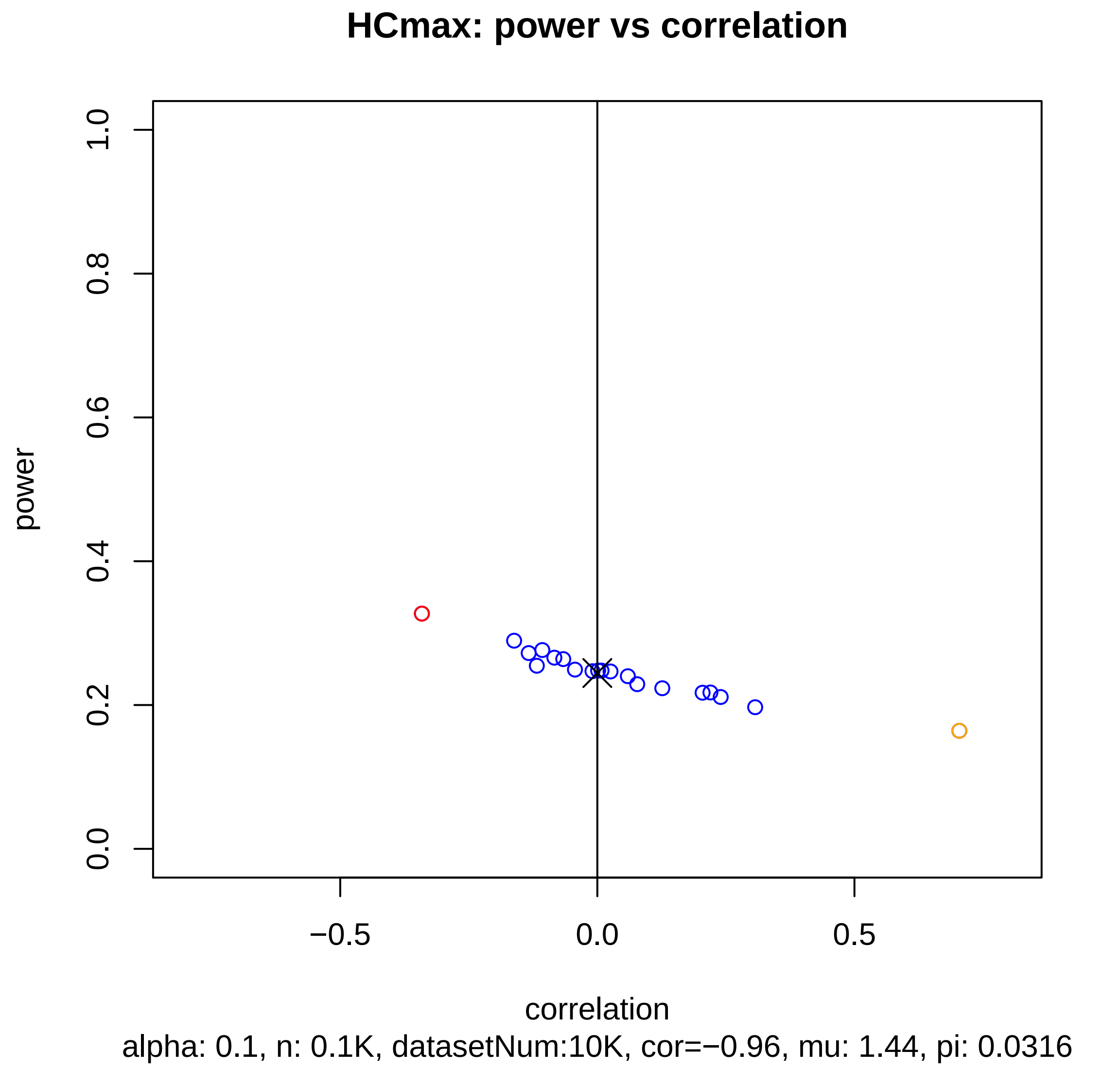
HCmax: power vs correlation
power
0.0
0.2
0.4
0.6
0.8
1.0
−0.5
0.0
0.5
correlation
alpha: 0.1, n: 0.1K, datasetNum:10K, cor=−0.96, mu: 1.44, pi: 0.0316

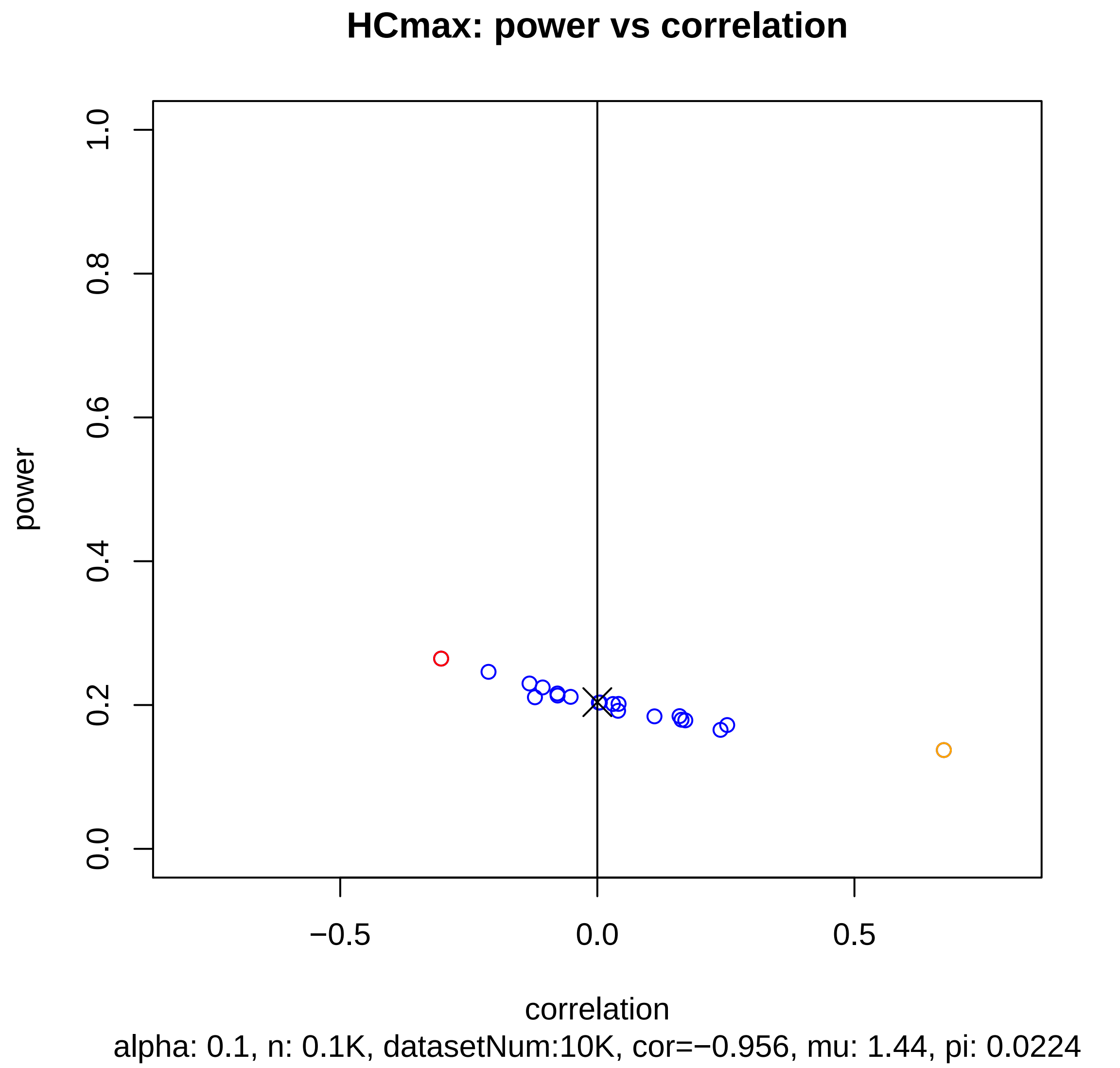
HCmax: power vs correlation
power
1.0
0.8
0.6
0.4
0.2
0.0
−0.5
0.0
0.5
correlation
alpha: 0.1, n: 0.1K, datasetNum:10K, cor=−0.956, mu: 1.44, pi: 0.0224

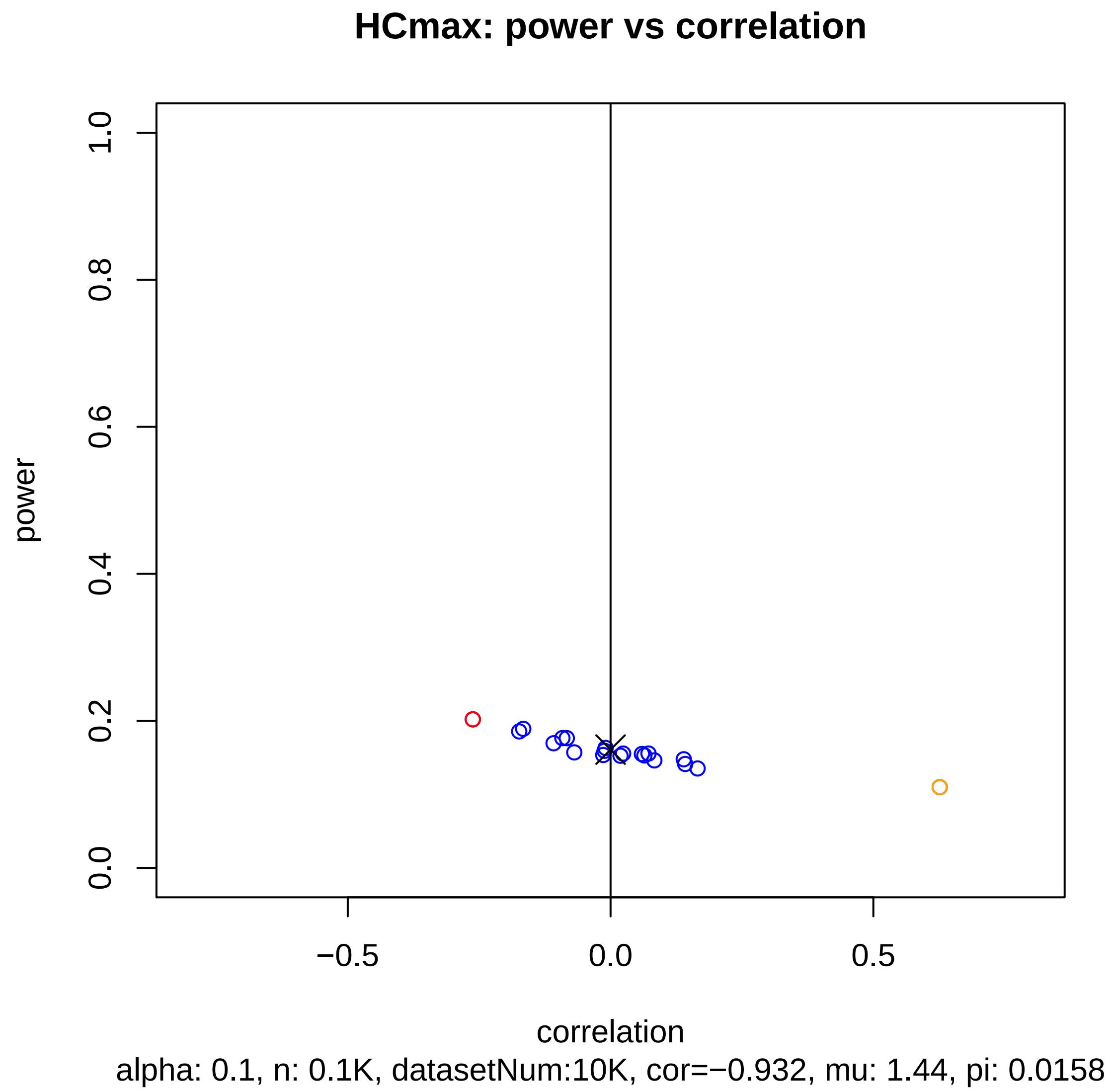
HCmax: power vs correlation
power
1.0
0.8
0.6
0.4
0.2
0.0
−0.5
0.0
0.5
correlation
alpha: 0.1, n: 0.1K, datasetNum:10K, cor=−0.932, mu: 1.44, pi: 0.0158

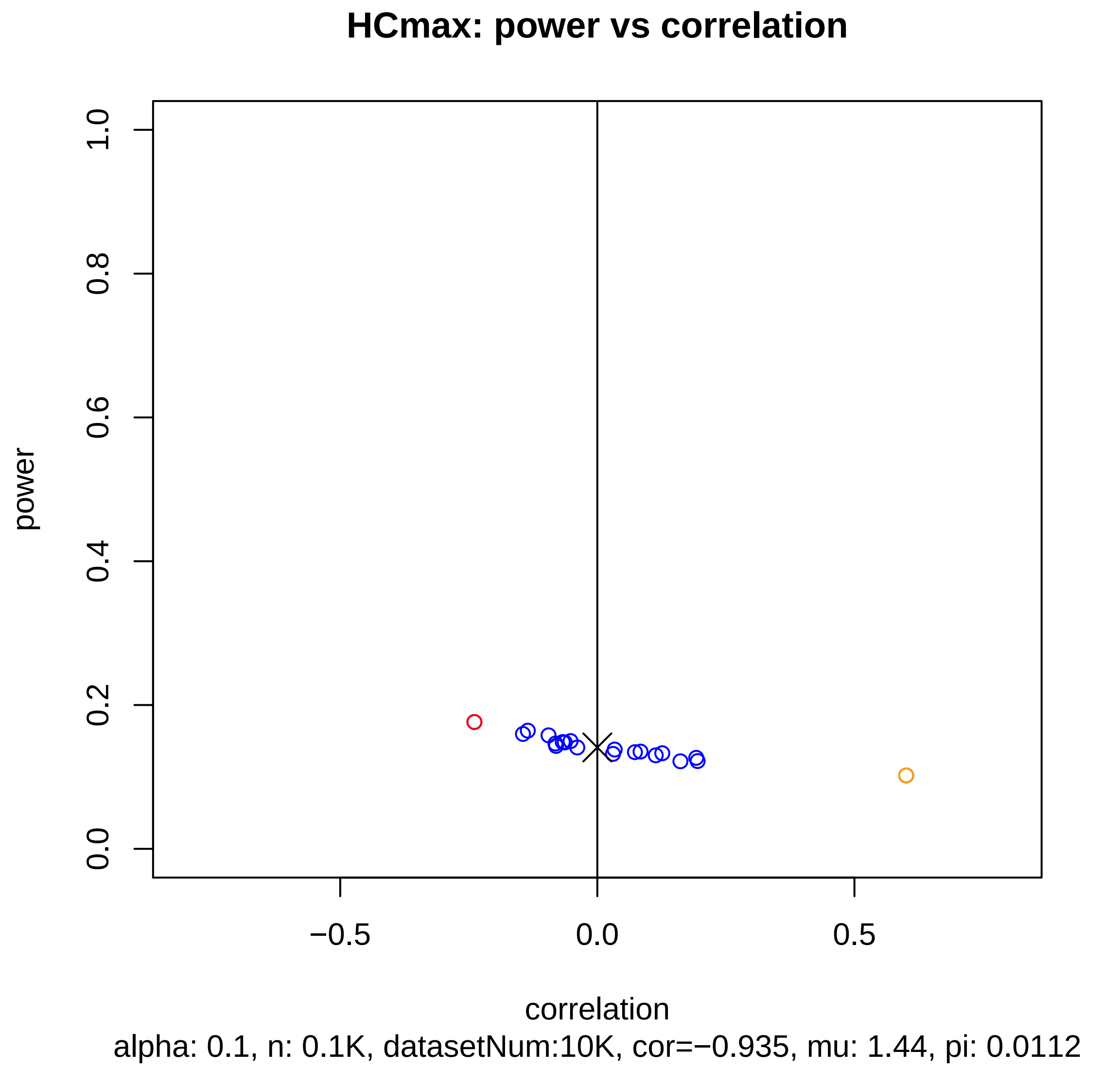
HCmax: power vs correlation
power
1.0
0.8
0.6
0.4
0.2
0.0
−0.5
0.0
0.5
correlation
alpha: 0.1, n: 0.1K, datasetNum:10K, cor=−0.935, mu: 1.44, pi: 0.0112

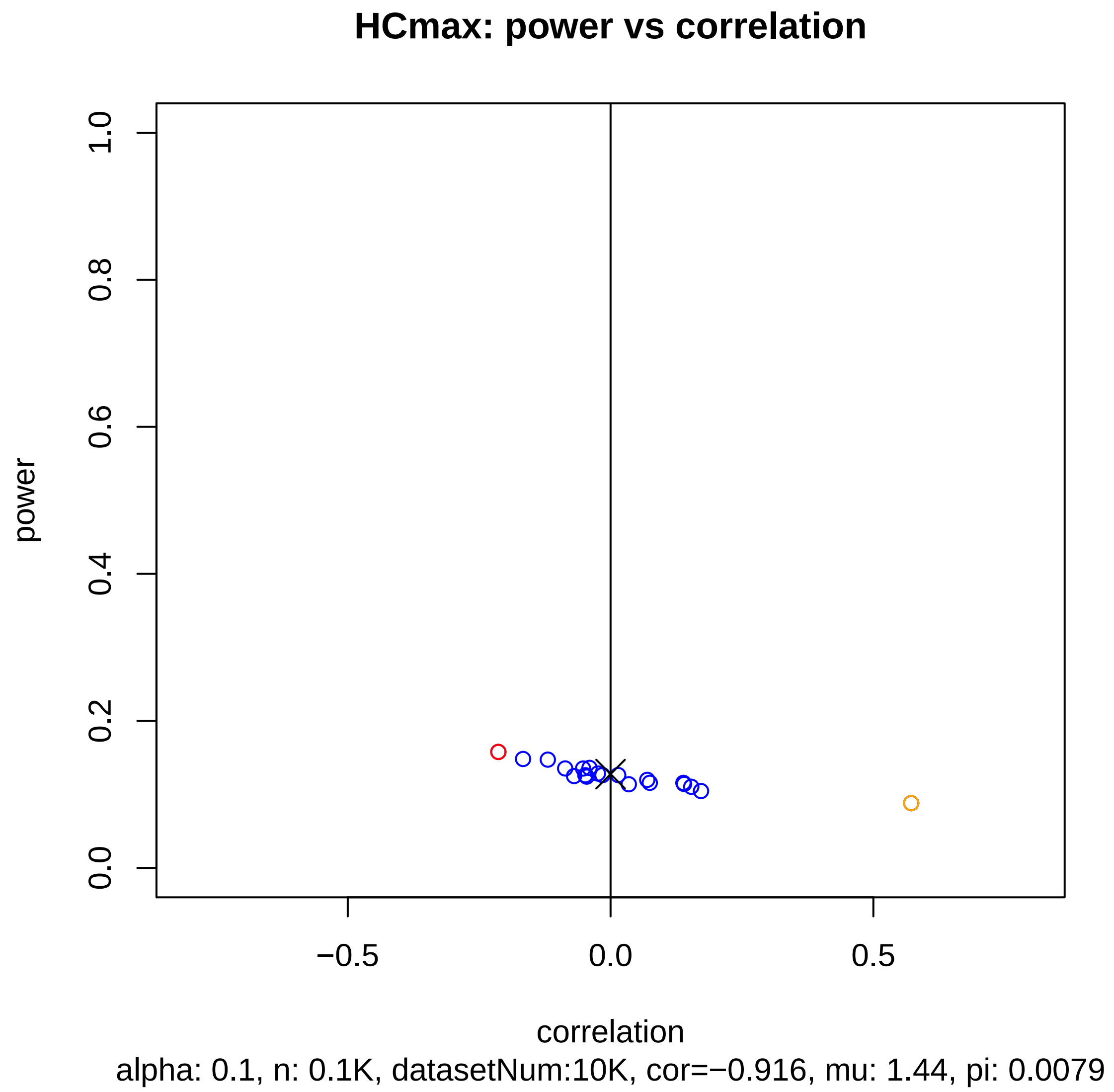
HCmax: power vs correlation
power
0.0
0.2
0.4
0.6
0.8
1.0
−0.5
0.0
0.5
correlation
alpha: 0.1, n: 0.1K, datasetNum:10K, cor=−0.916, mu: 1.44, pi: 0.0079

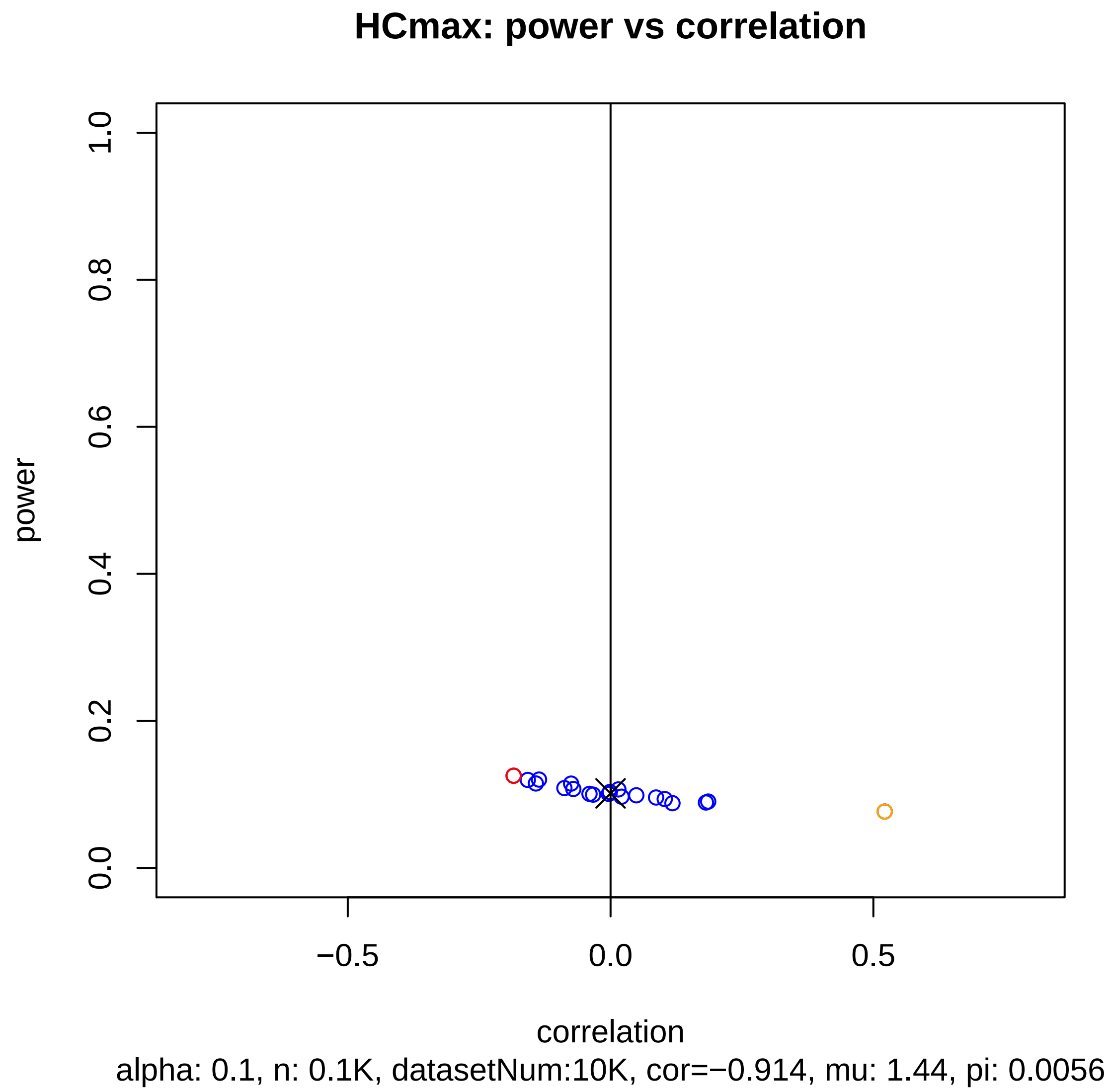
HCmax: power vs correlation
power
0.0
0.2
0.4
0.6
0.8
1.0
−0.5
0.0
0.5
correlation
alpha: 0.1, n: 0.1K, datasetNum:10K, cor=−0.914, mu: 1.44, pi: 0.0056

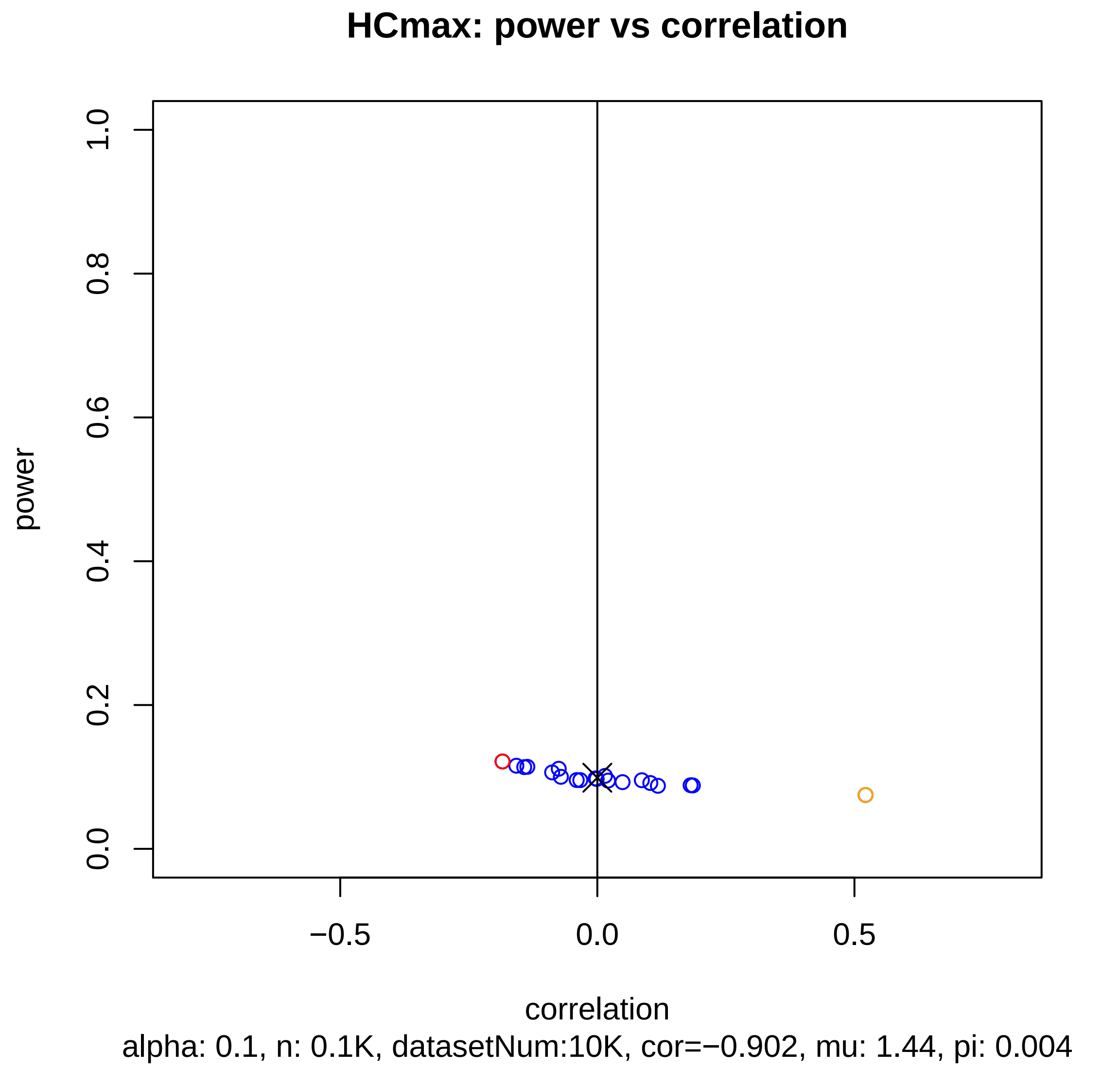
HCmax: power vs correlation
power
correlation
0.0
0.2
0.4
0.6
0.8
1.0
−0.5
0.0
0.5
alpha: 0.1, n: 0.1K, datasetNum:10K, cor=−0.902, mu: 1.44, pi: 0.004

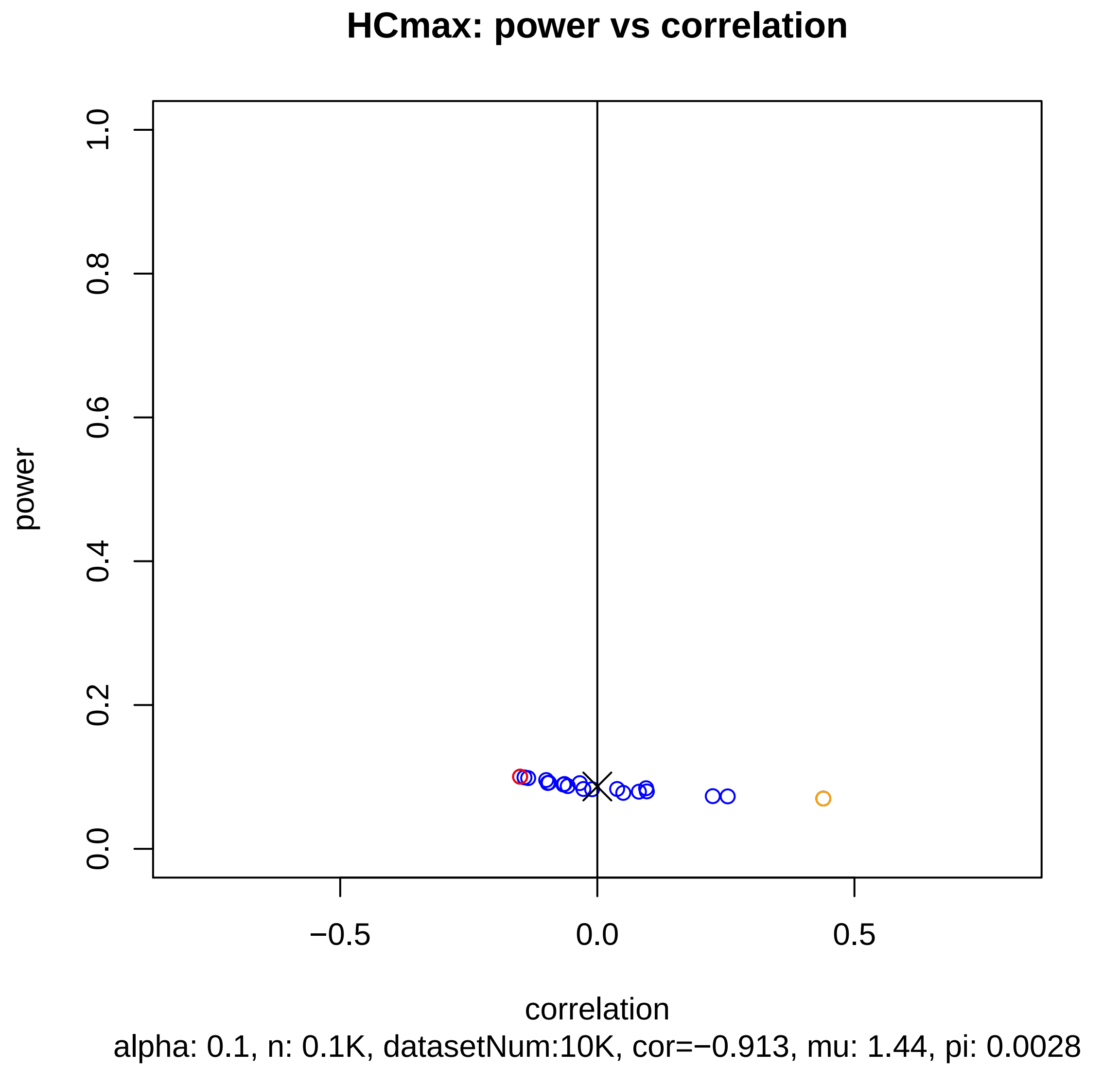

HCmax: power vs correlation
power
0.0
0.2
0.4
0.6
0.8
1.0
−0.5
0.0
0.5
correlation
alpha: 0.1, n: 0.1K, datasetNum:10K, cor=−0.913, mu: 1.44, pi: 0.0028

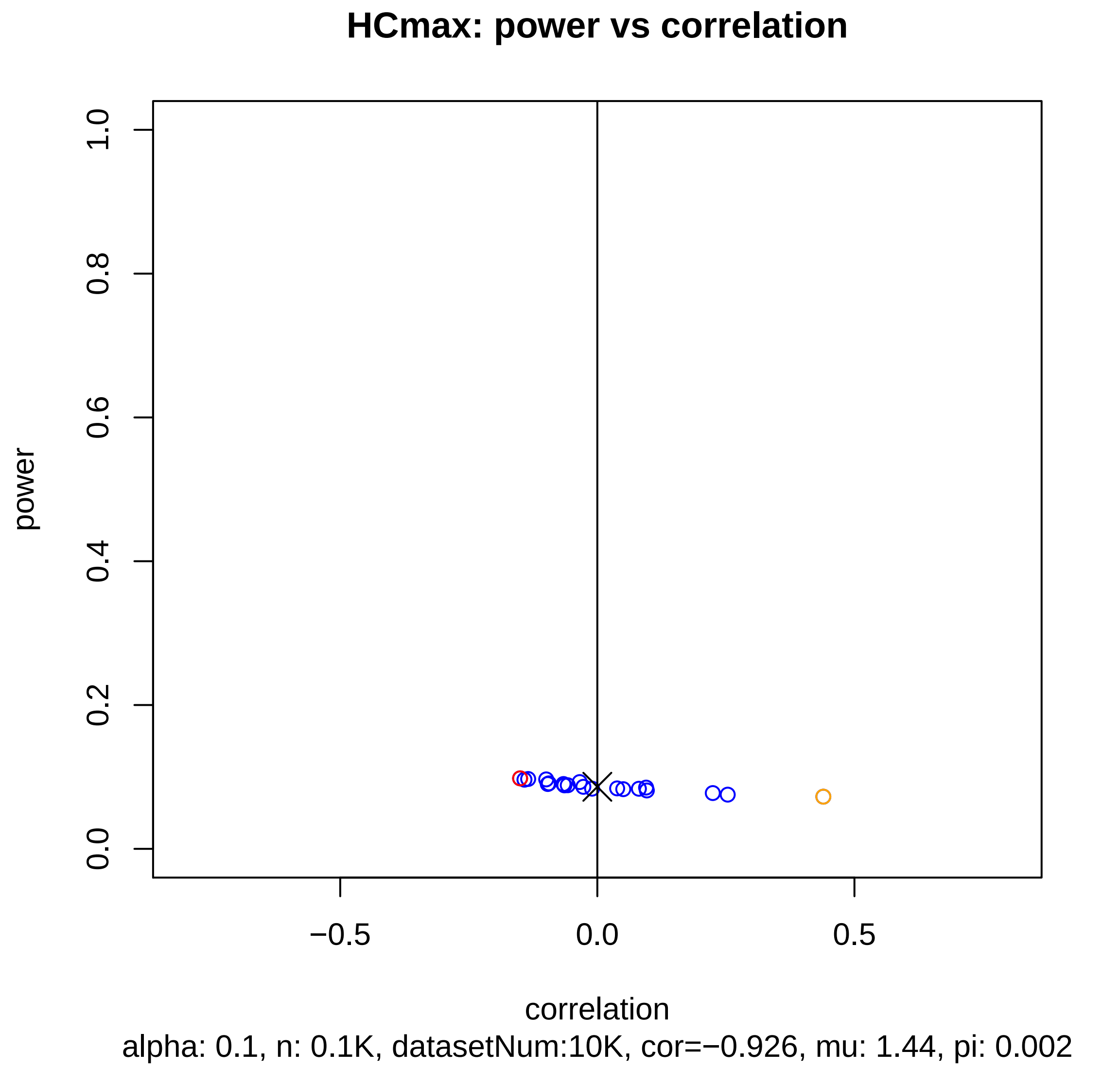

HCmax: power vs correlation
power
1.0
0.8
0.6
0.4
0.2
0.0
−0.5
0.0
0.5
correlation
alpha: 0.1, n: 0.1K, datasetNum:10K, cor=−0.926, mu: 1.44, pi: 0.002

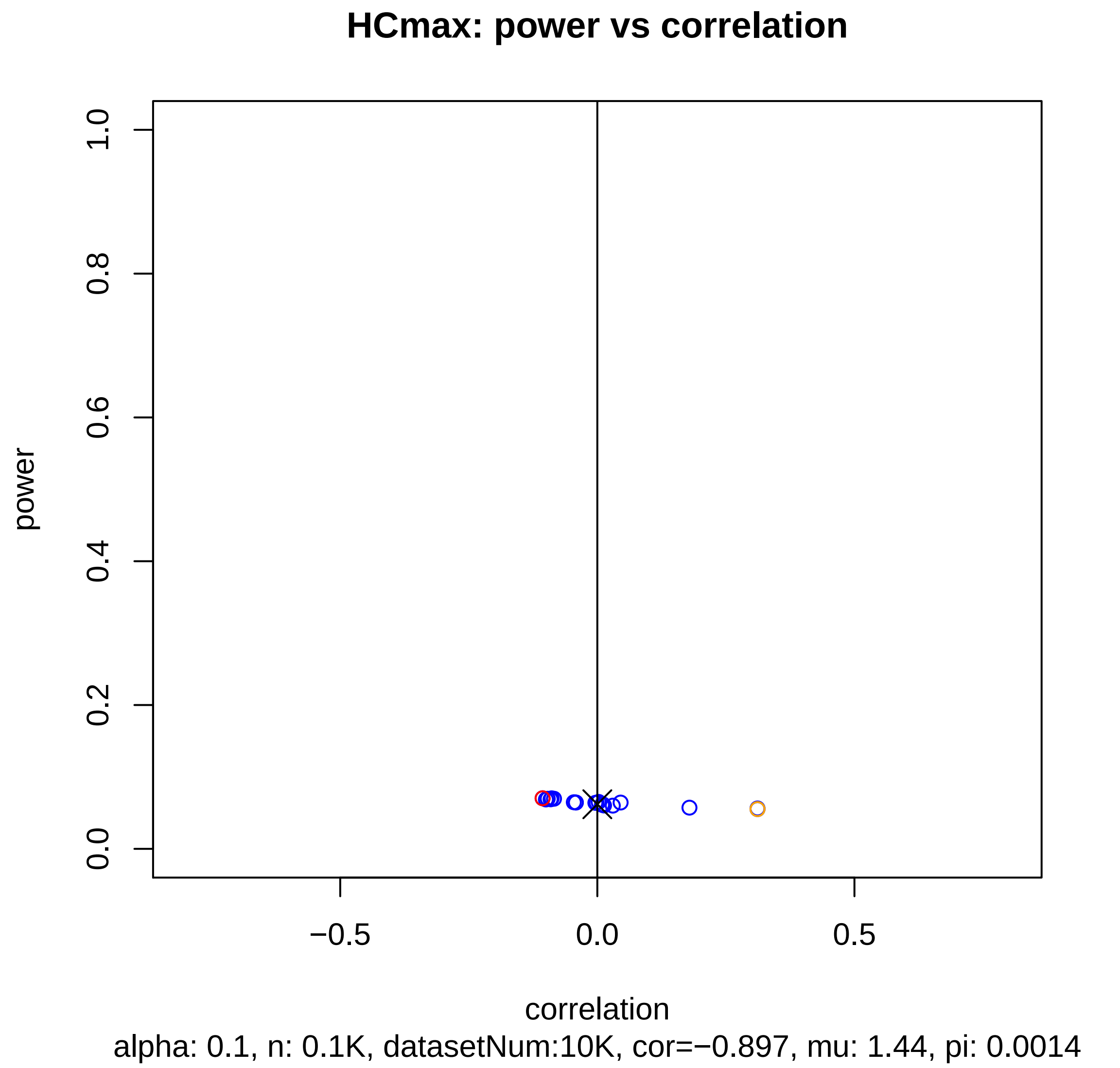

HCmax: power vs correlation
power
1.0
0.8
0.6
0.4
0.2
0.0
−0.5
0.0
0.5
correlation
alpha: 0.1, n: 0.1K, datasetNum:10K, cor=−0.897, mu: 1.44, pi: 0.0014

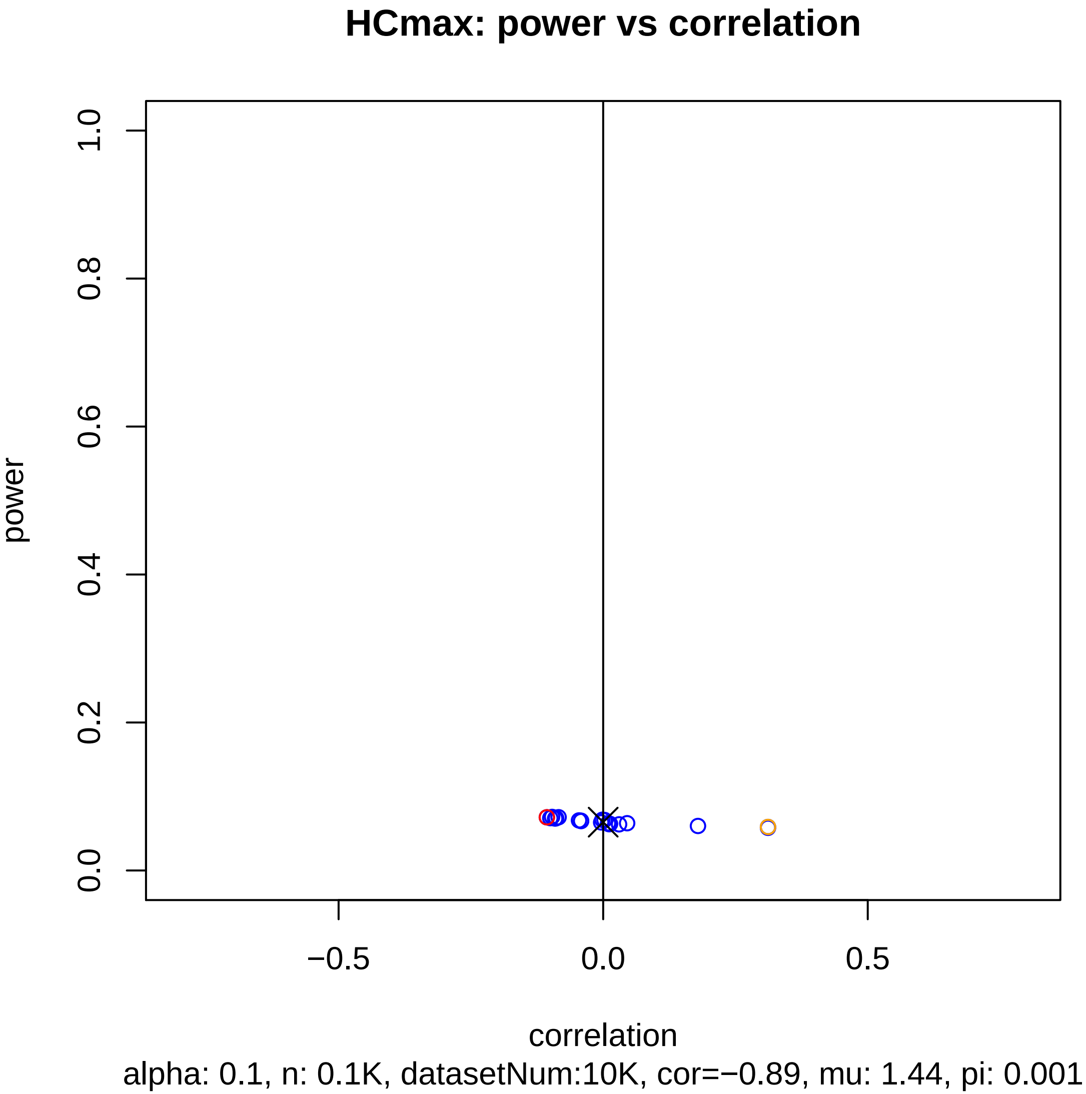
HCmax: power vs correlation
power
1.0
0.8
0.6
0.4
0.2
0.0
−0.5
0.0
0.5
correlation
alpha: 0.1, n: 0.1K, datasetNum:10K, cor=−0.89, mu: 1.44, pi: 0.001

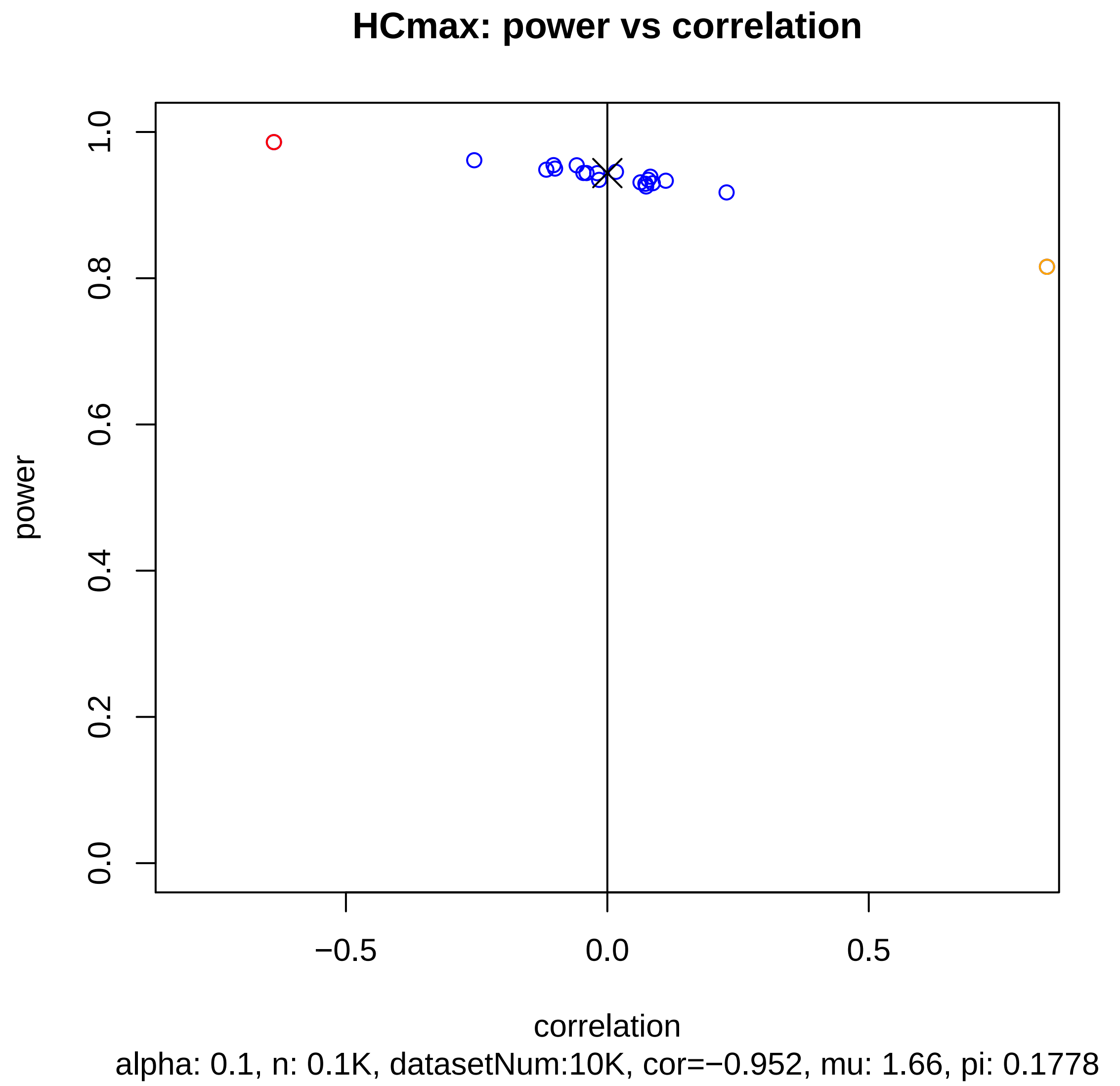
HCmax: power vs correlation
power
0.0
0.2
0.4
0.6
0.8
1.0
−0.5
0.0
0.5
correlation
alpha: 0.1, n: 0.1K, datasetNum:10K, cor=−0.952, mu: 1.66, pi: 0.1778

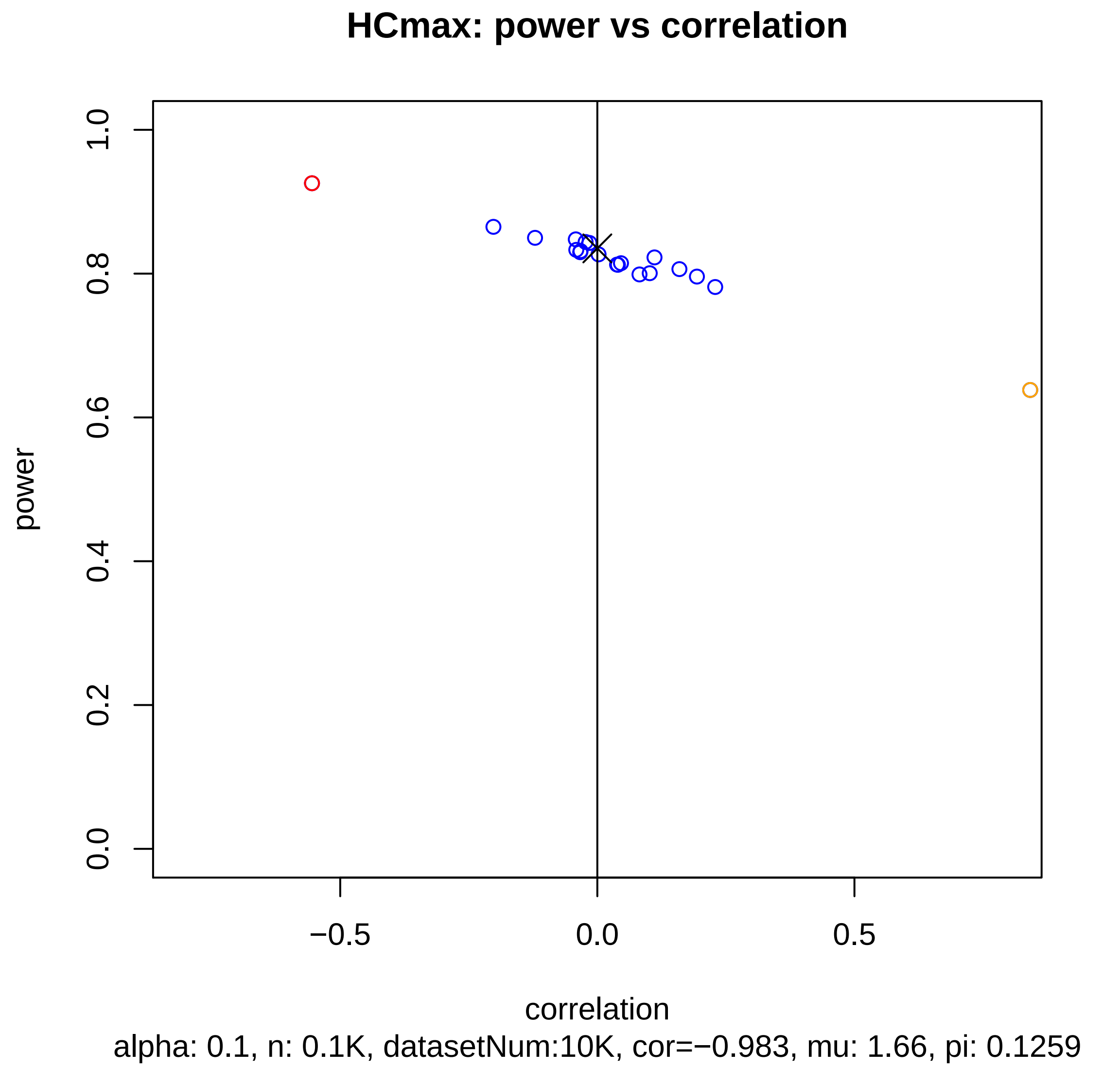
HCmax: power vs correlation
power
correlation
1.0
0.8
0.6
0.4
0.2
0.0
−0.5
0.0
0.5
alpha: 0.1, n: 0.1K, datasetNum:10K, cor=−0.983, mu: 1.66, pi: 0.1259

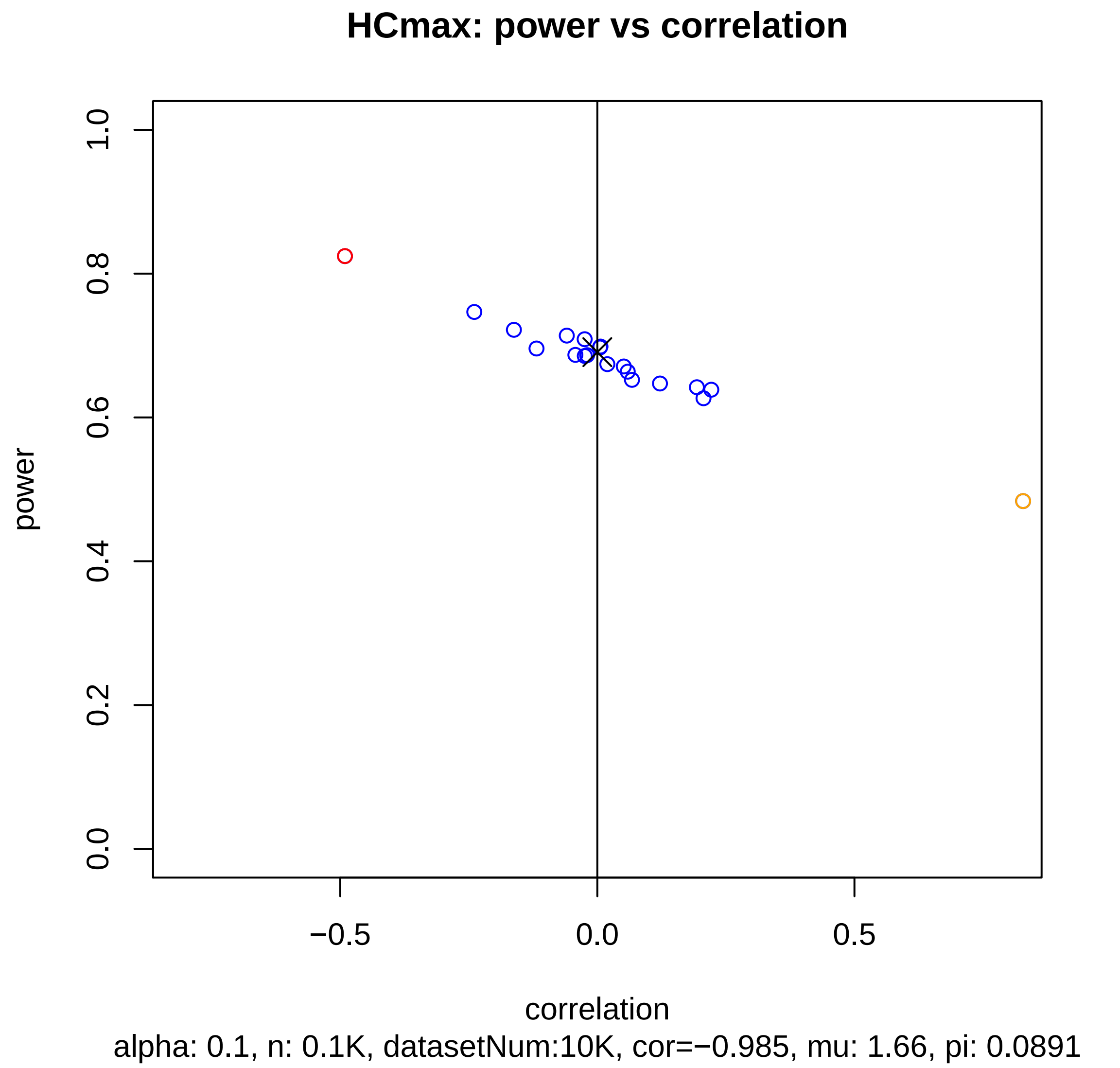

HCmax: power vs correlation
power
1.0
0.8
0.6
0.4
0.2
0.0
−0.5
0.0
0.5
correlation
alpha: 0.1, n: 0.1K, datasetNum:10K, cor=−0.985, mu: 1.66, pi: 0.0891

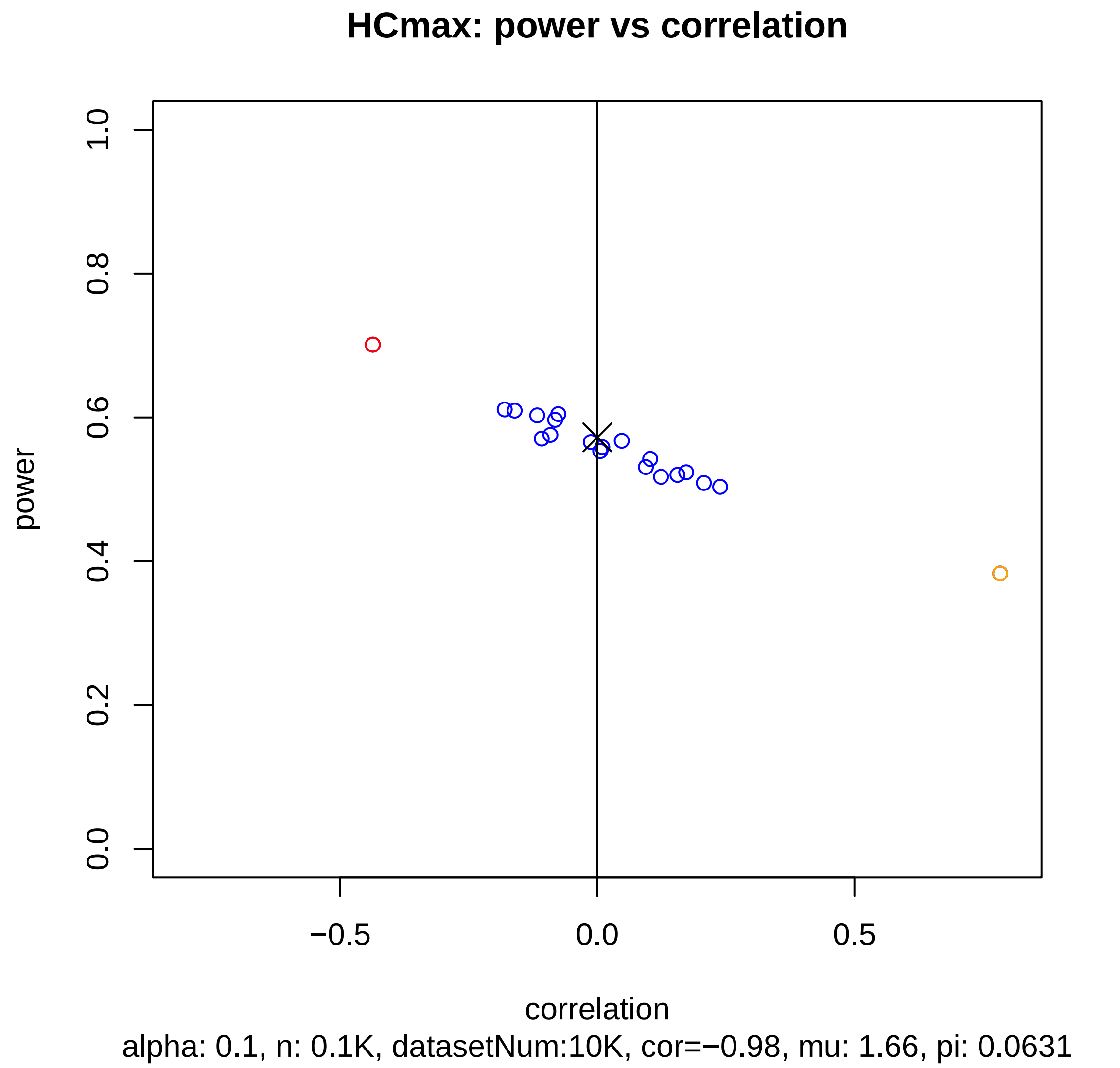
HCmax: power vs correlation
power
0.0
0.2
0.4
0.6
0.8
1.0
−0.5
0.0
0.5
correlation
alpha: 0.1, n: 0.1K, datasetNum:10K, cor=−0.98, mu: 1.66, pi: 0.0631

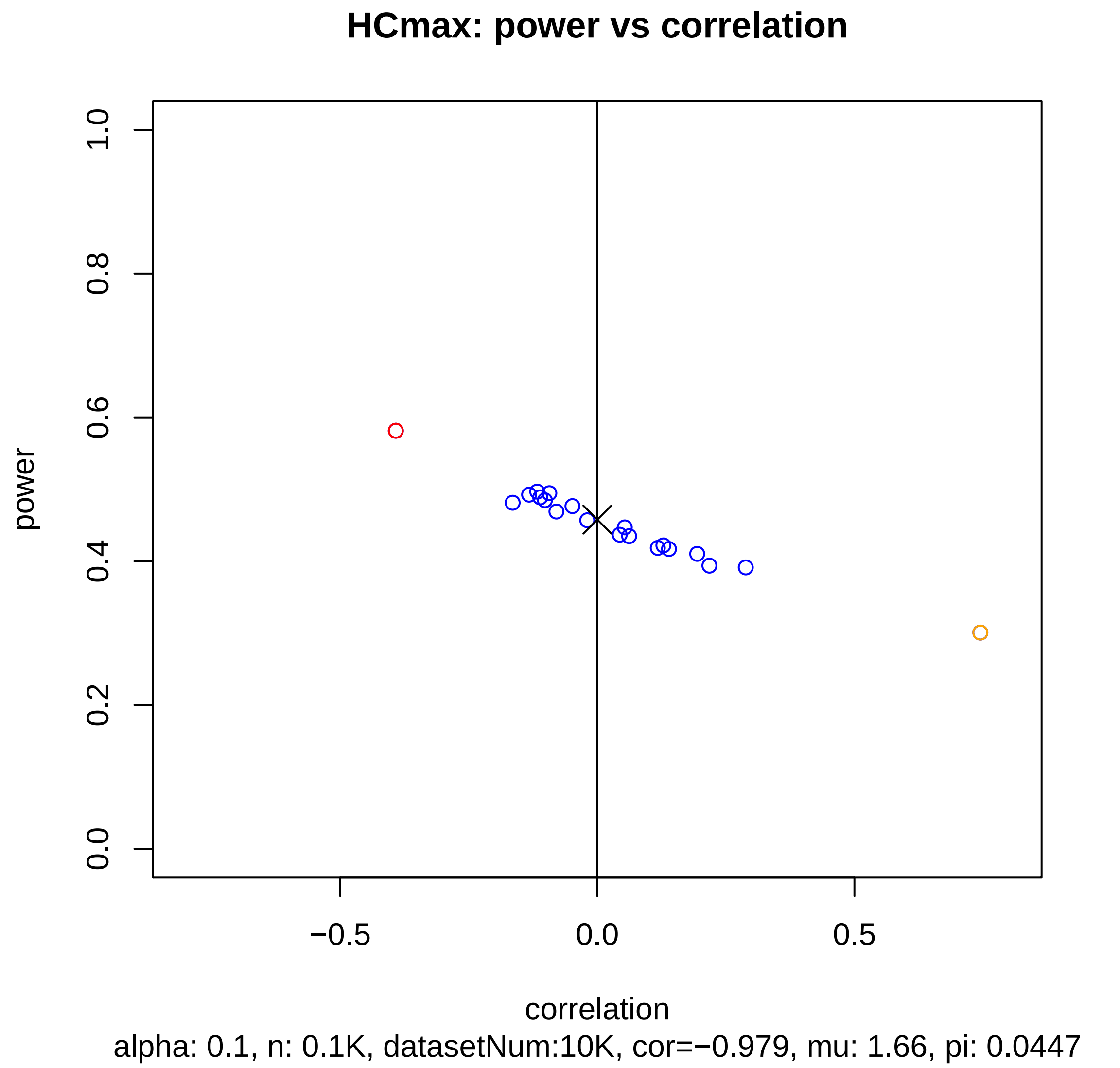
HCmax: power vs correlation
power
0.0
0.2
0.4
0.6
0.8
1.0
−0.5
0.0
0.5
correlation
alpha: 0.1, n: 0.1K, datasetNum:10K, cor=−0.979, mu: 1.66, pi: 0.0447

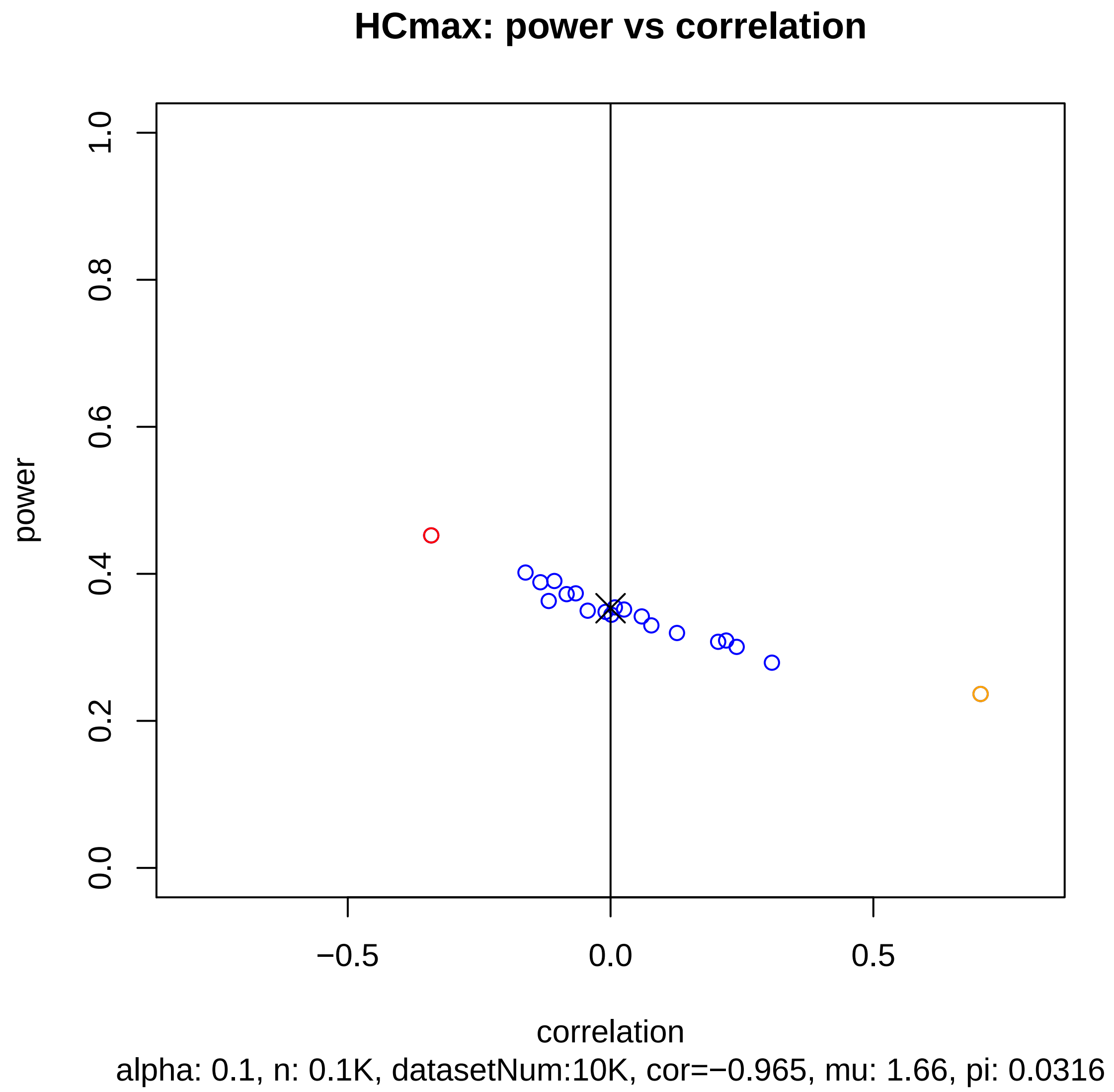
HCmax: power vs correlation
power
0.0
0.2
0.4
0.6
0.8
1.0
−0.5
0.0
0.5
correlation
alpha: 0.1, n: 0.1K, datasetNum:10K, cor=−0.965, mu: 1.66, pi: 0.0316

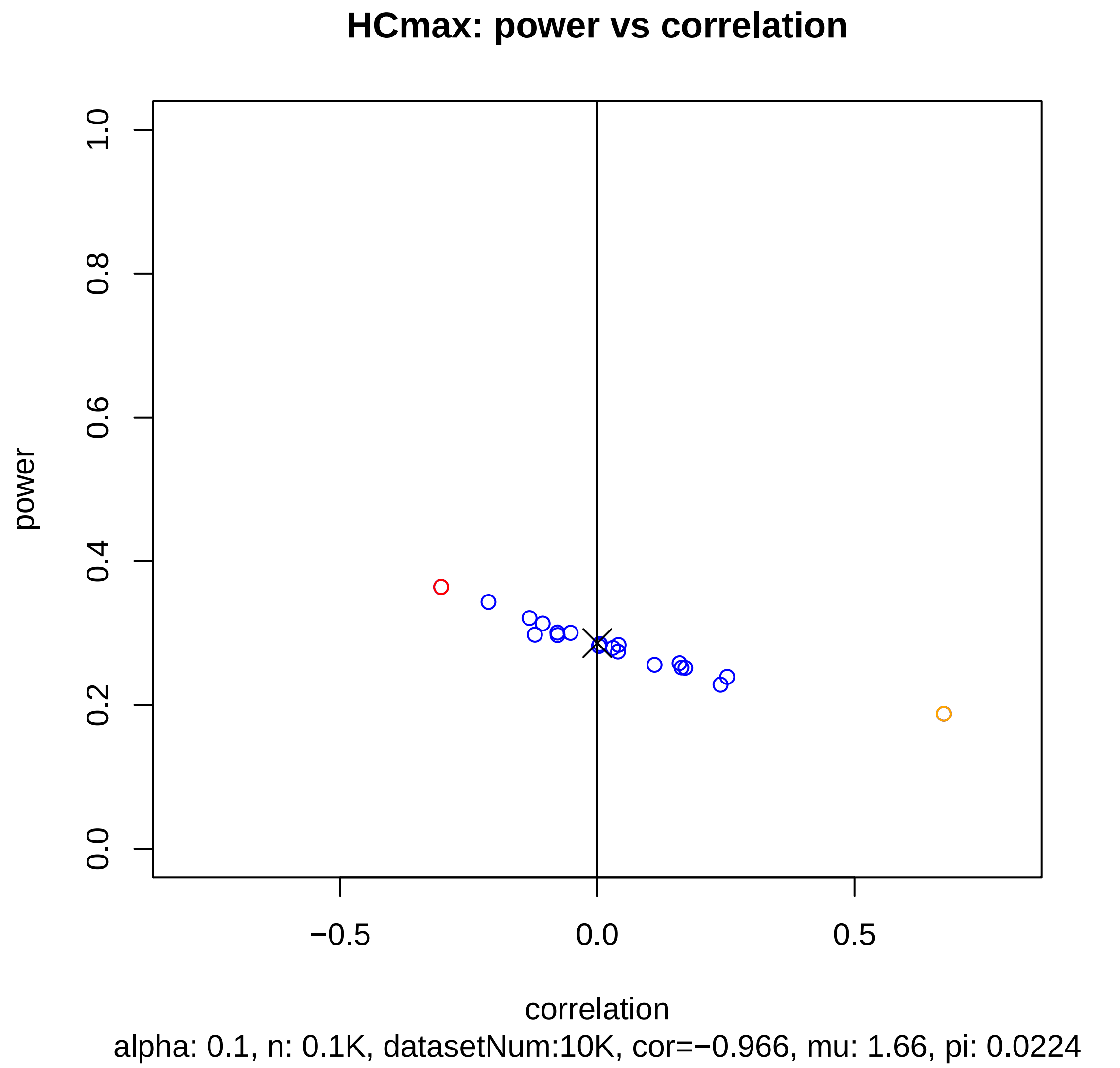

HCmax: power vs correlation
power
1.0
0.8
0.6
0.4
0.2
0.0
−0.5
0.0
0.5
correlation
alpha: 0.1, n: 0.1K, datasetNum:10K, cor=−0.966, mu: 1.66, pi: 0.0224

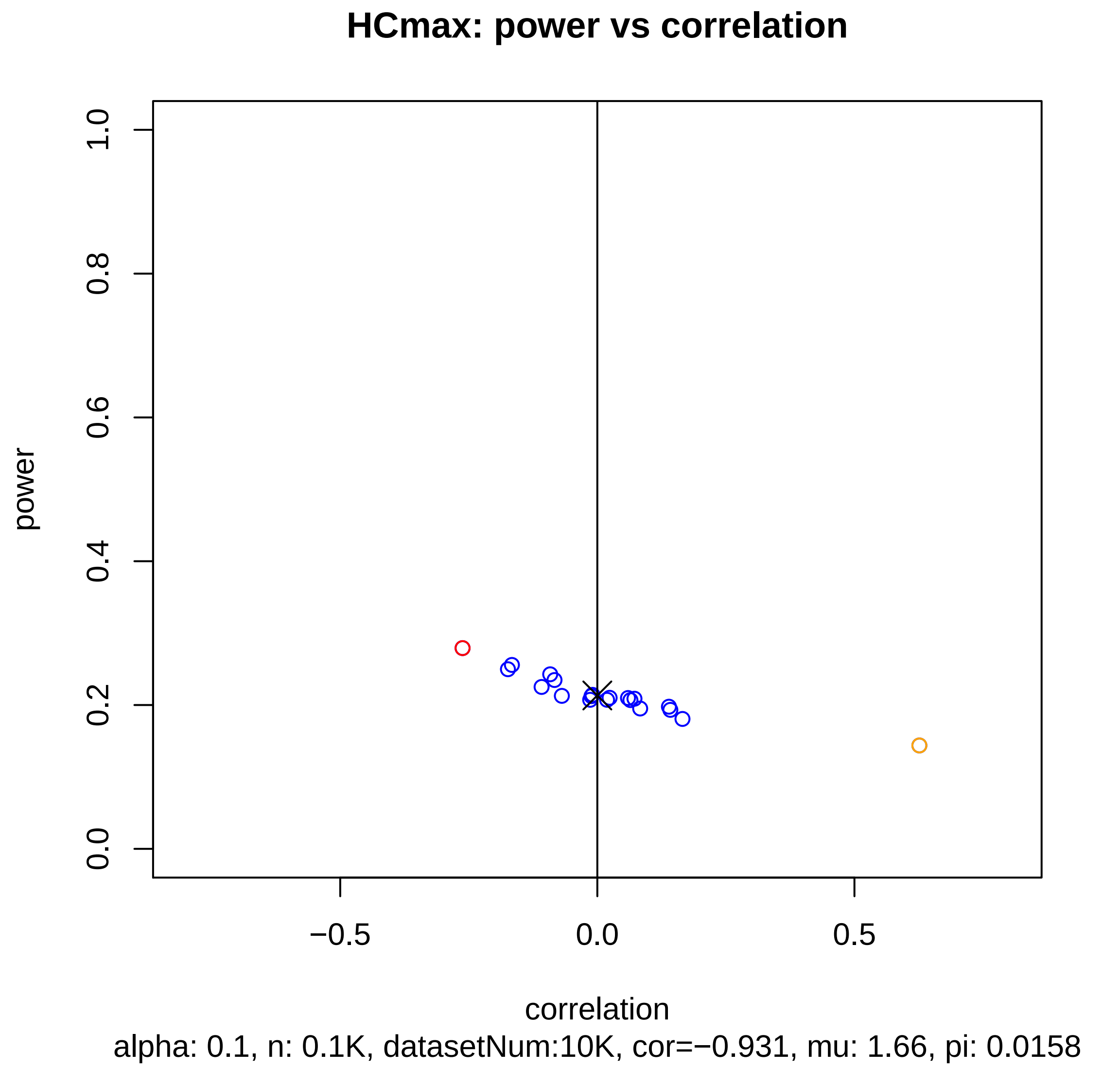
HCmax: power vs correlation
power
1.0
0.8
0.6
0.4
0.2
0.0
−0.5
0.0
0.5
correlation
alpha: 0.1, n: 0.1K, datasetNum:10K, cor=−0.931, mu: 1.66, pi: 0.0158

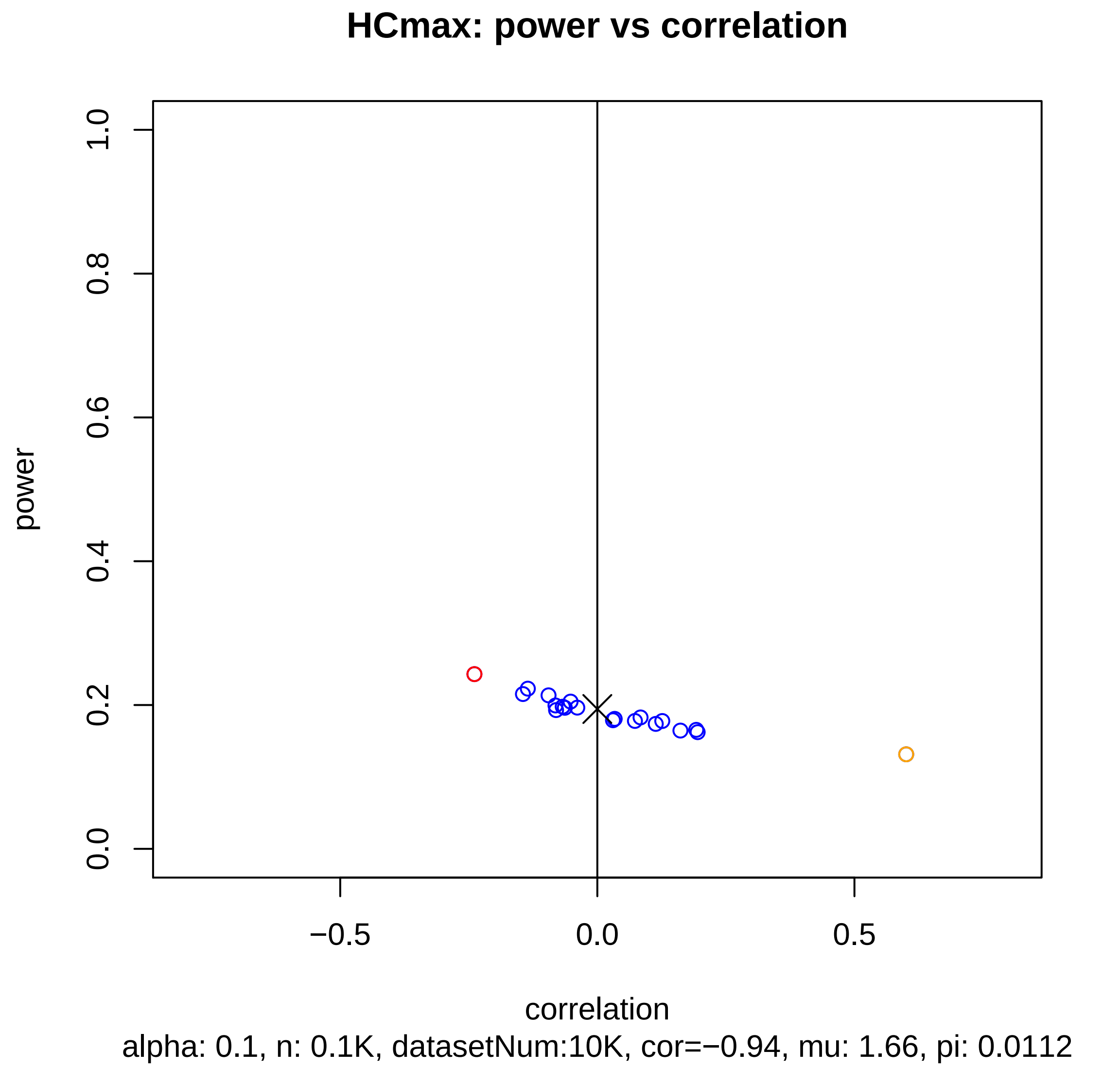
HCmax: power vs correlation
power
1.0
0.8
0.6
0.4
0.2
0.0
−0.5
0.0
0.5
correlation
alpha: 0.1, n: 0.1K, datasetNum:10K, cor=−0.94, mu: 1.66, pi: 0.0112

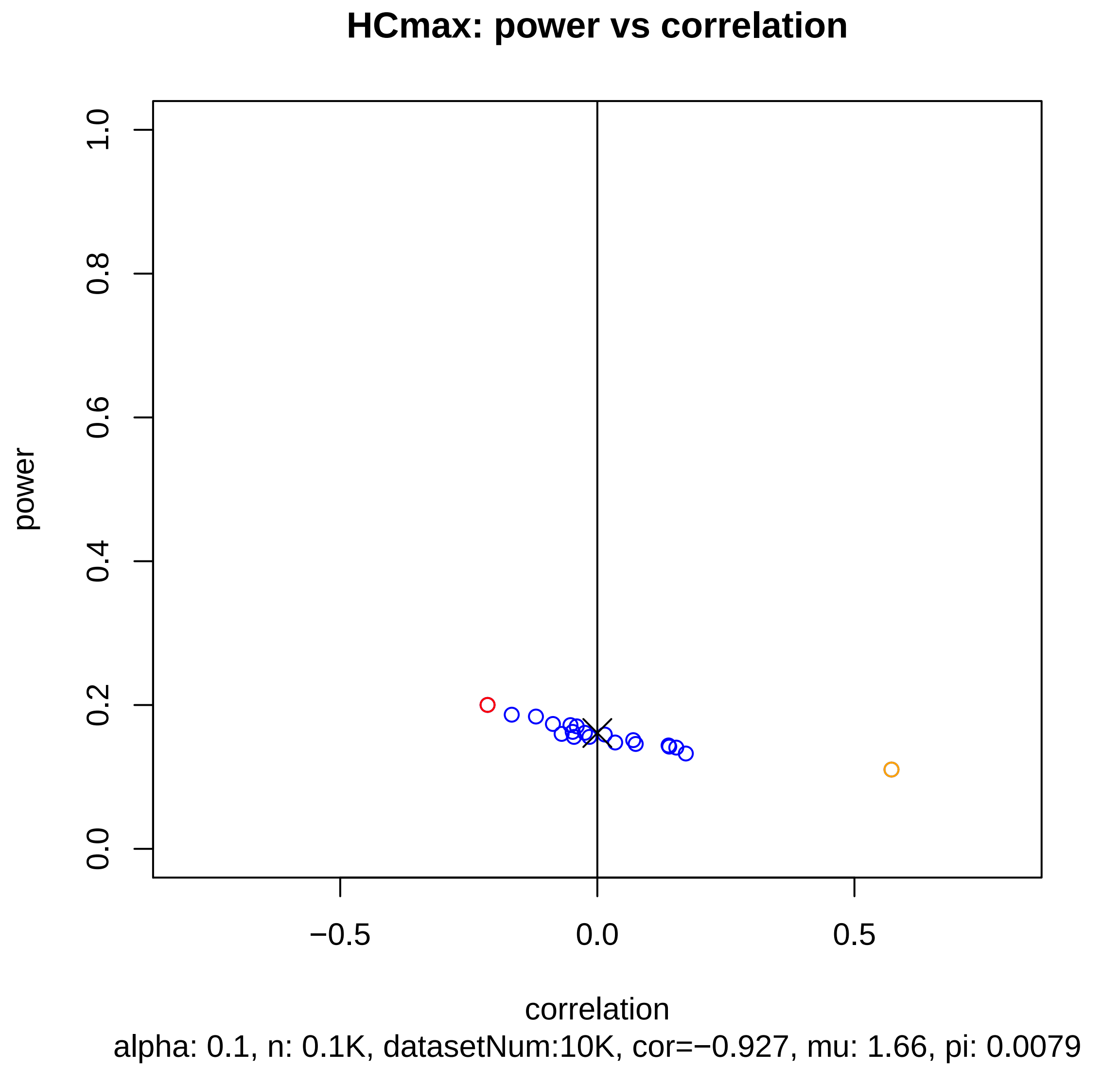
HCmax: power vs correlation
power
1.0
0.8
0.6
0.4
0.2
0.0
−0.5
0.0
0.5
correlation
alpha: 0.1, n: 0.1K, datasetNum:10K, cor=−0.927, mu: 1.66, pi: 0.0079

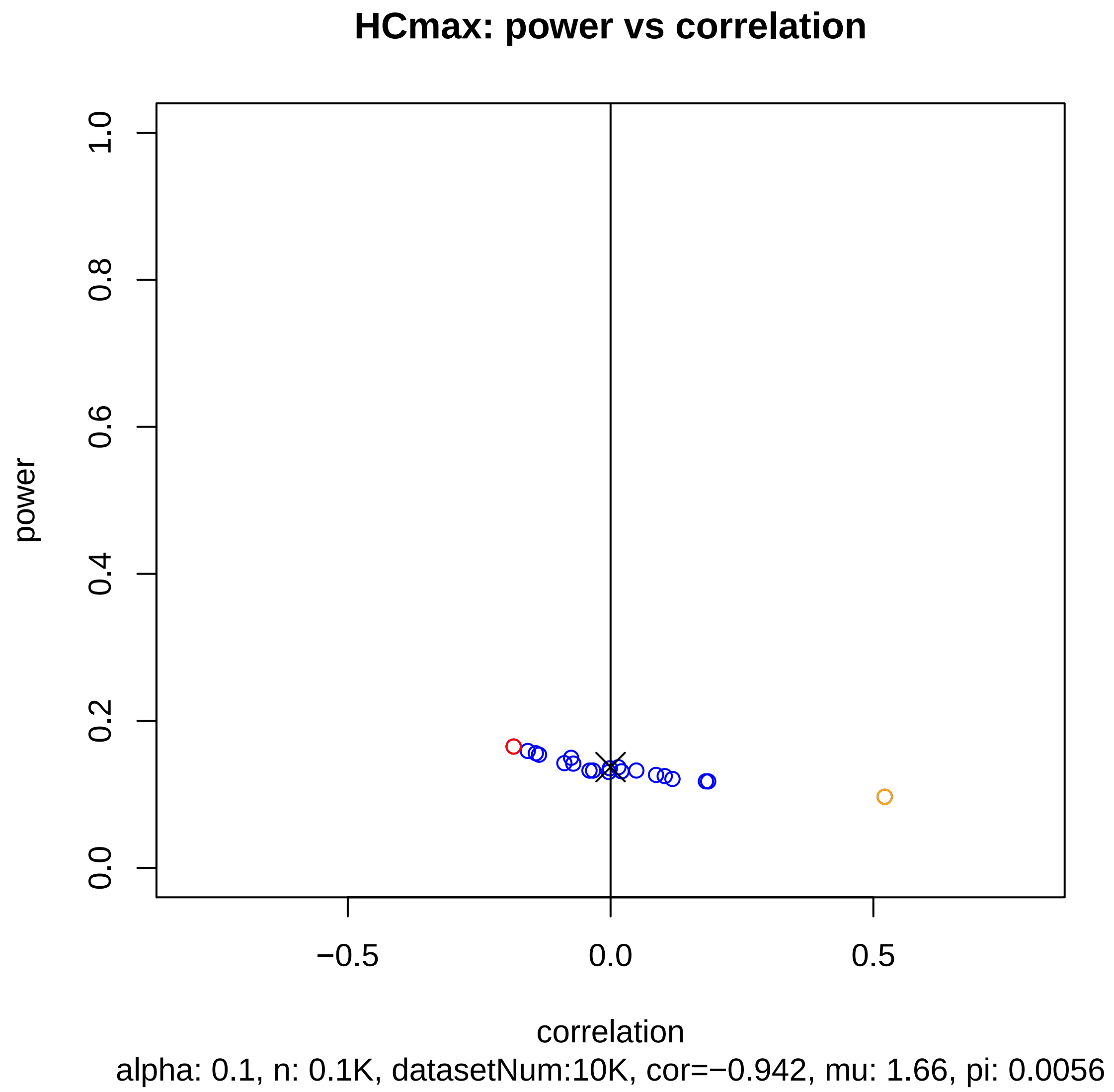
HCmax: power vs correlation
power
1.0
0.8
0.6
0.4
0.2
0.0
−0.5
0.0
0.5
correlation
alpha: 0.1, n: 0.1K, datasetNum:10K, cor=−0.942, mu: 1.66, pi: 0.0056

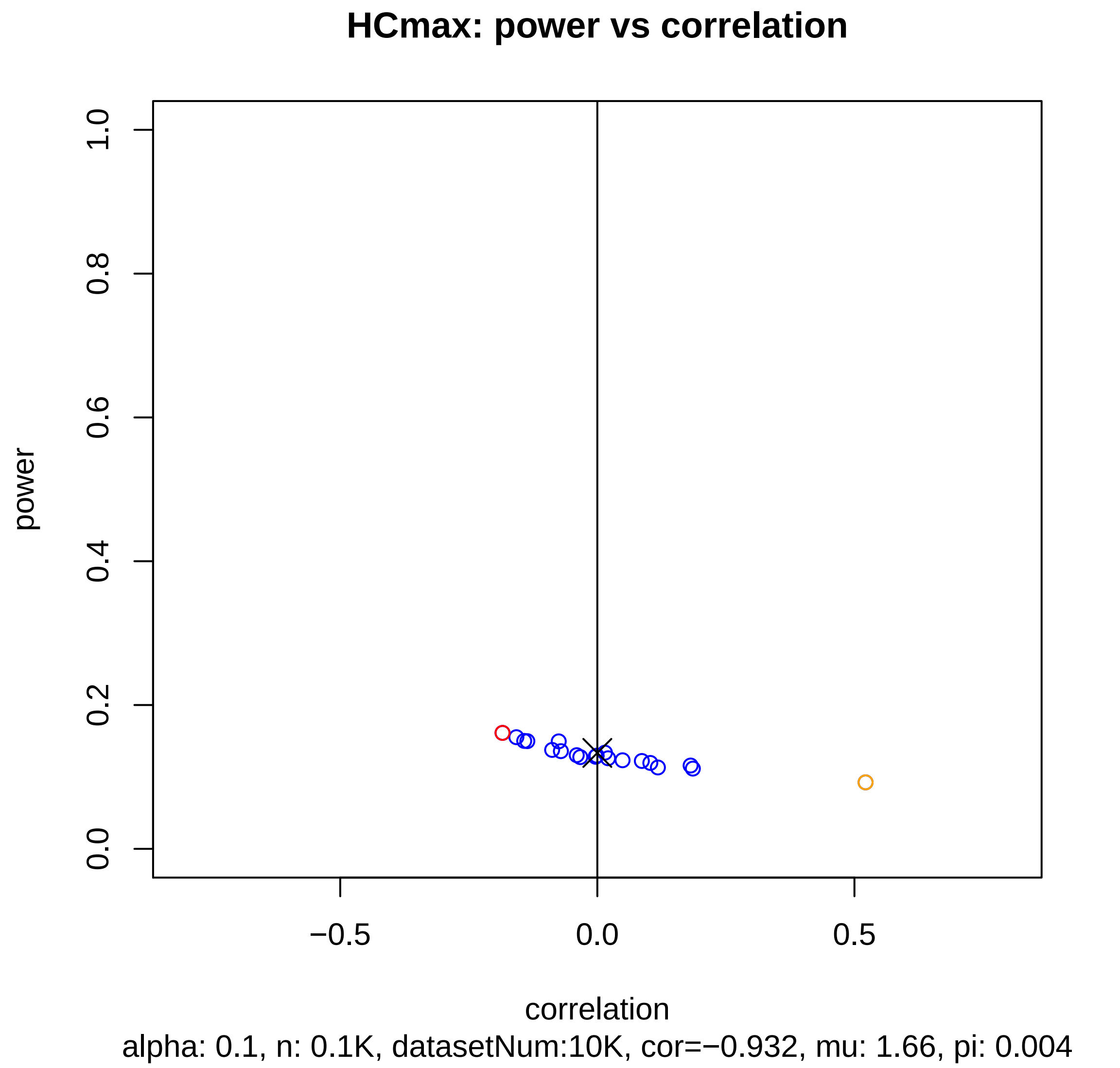
HCmax: power vs correlation
power
correlation
0.0
0.2
0.4
0.6
0.8
1.0
−0.5
0.0
0.5
alpha: 0.1, n: 0.1K, datasetNum:10K, cor=−0.932, mu: 1.66, pi: 0.004

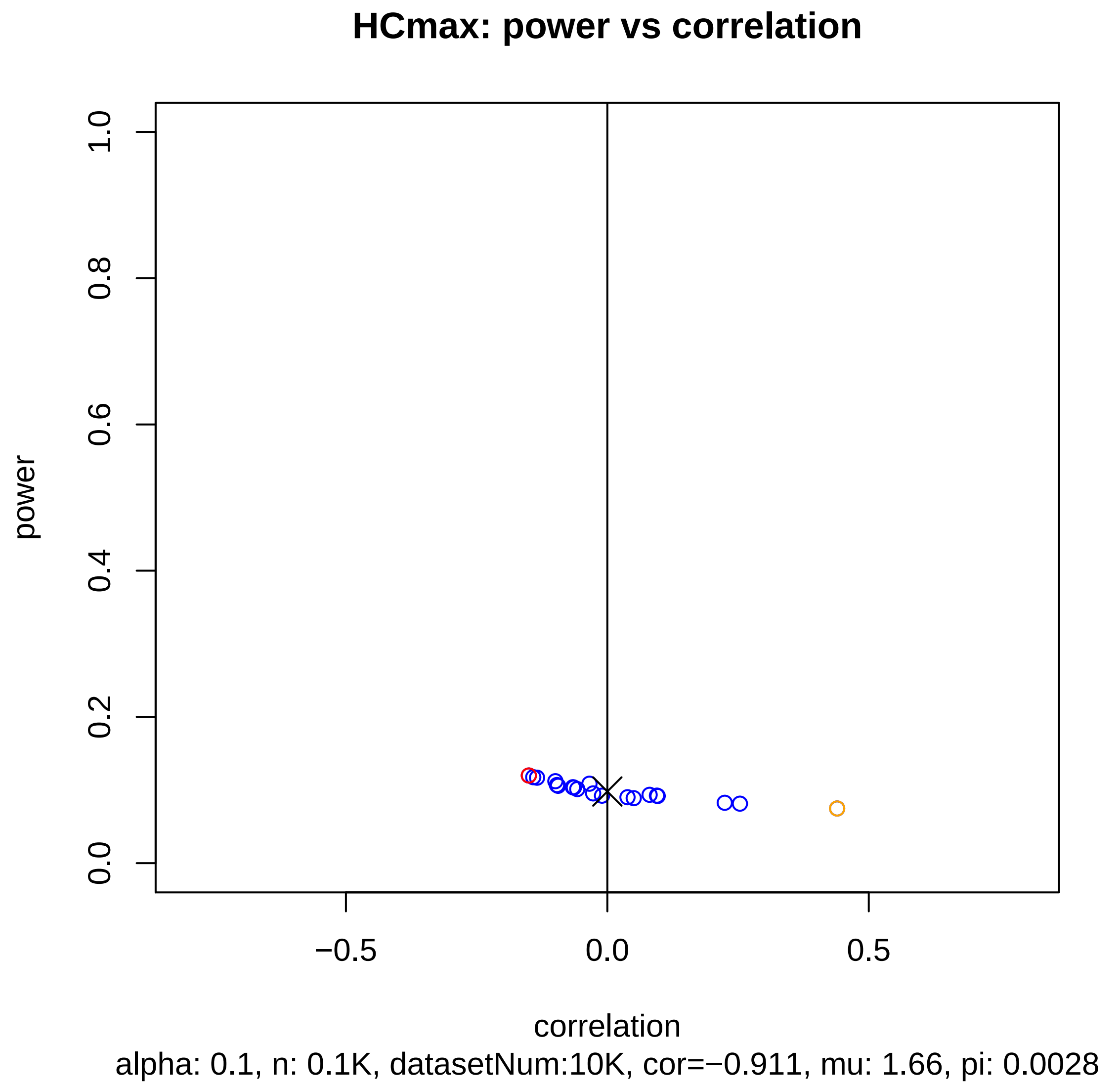
HCmax: power vs correlation
power
1.0
0.8
0.6
0.4
0.2
0.0
−0.5
0.0
0.5
correlation
alpha: 0.1, n: 0.1K, datasetNum:10K, cor=−0.911, mu: 1.66, pi: 0.0028

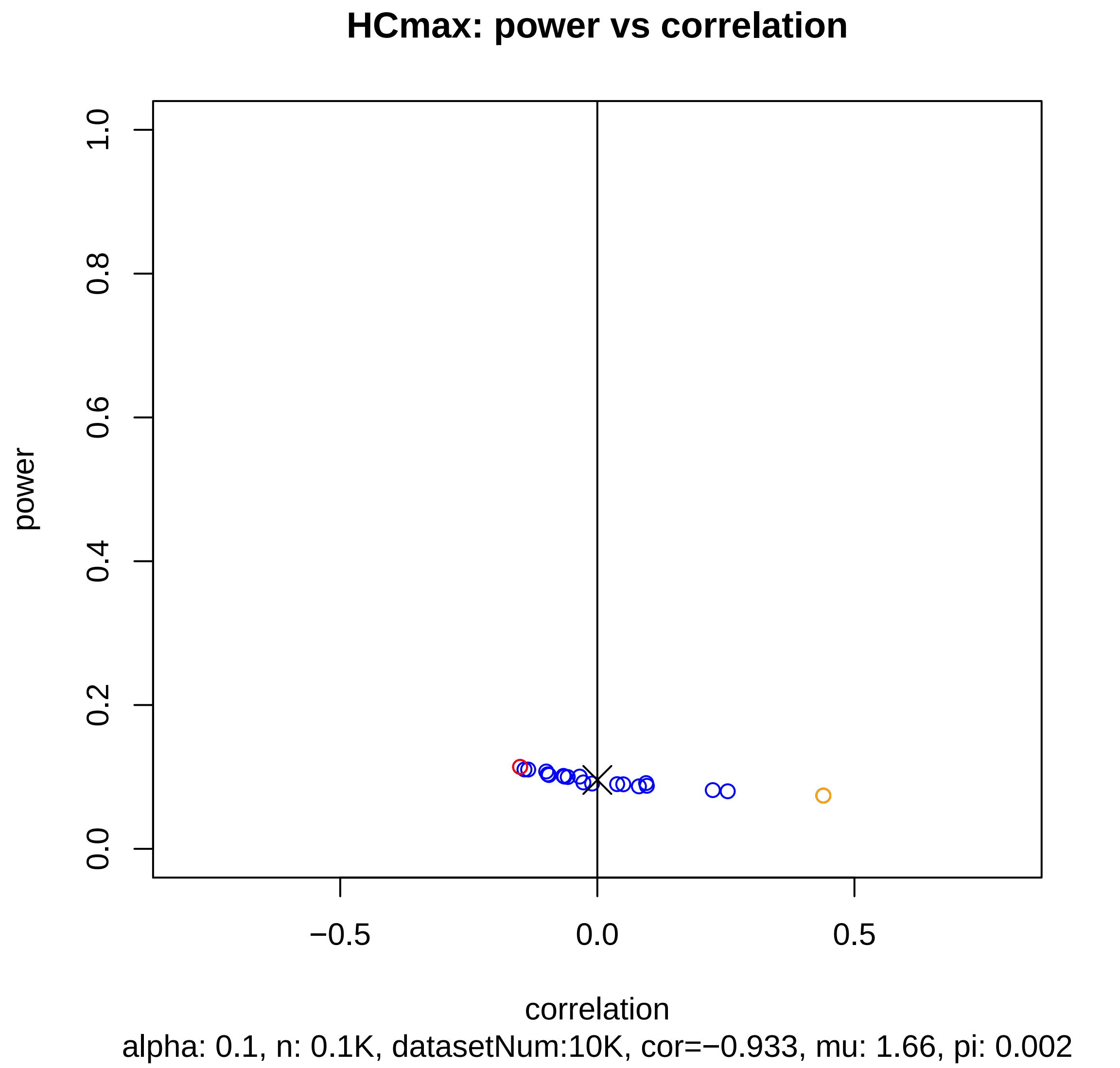
HCmax: power vs correlation
power
correlation
0.0
0.2
0.4
0.6
0.8
1.0
−0.5
0.0
0.5
alpha: 0.1, n: 0.1K, datasetNum:10K, cor=−0.933, mu: 1.66, pi: 0.002

### HCmax: power vs correlation

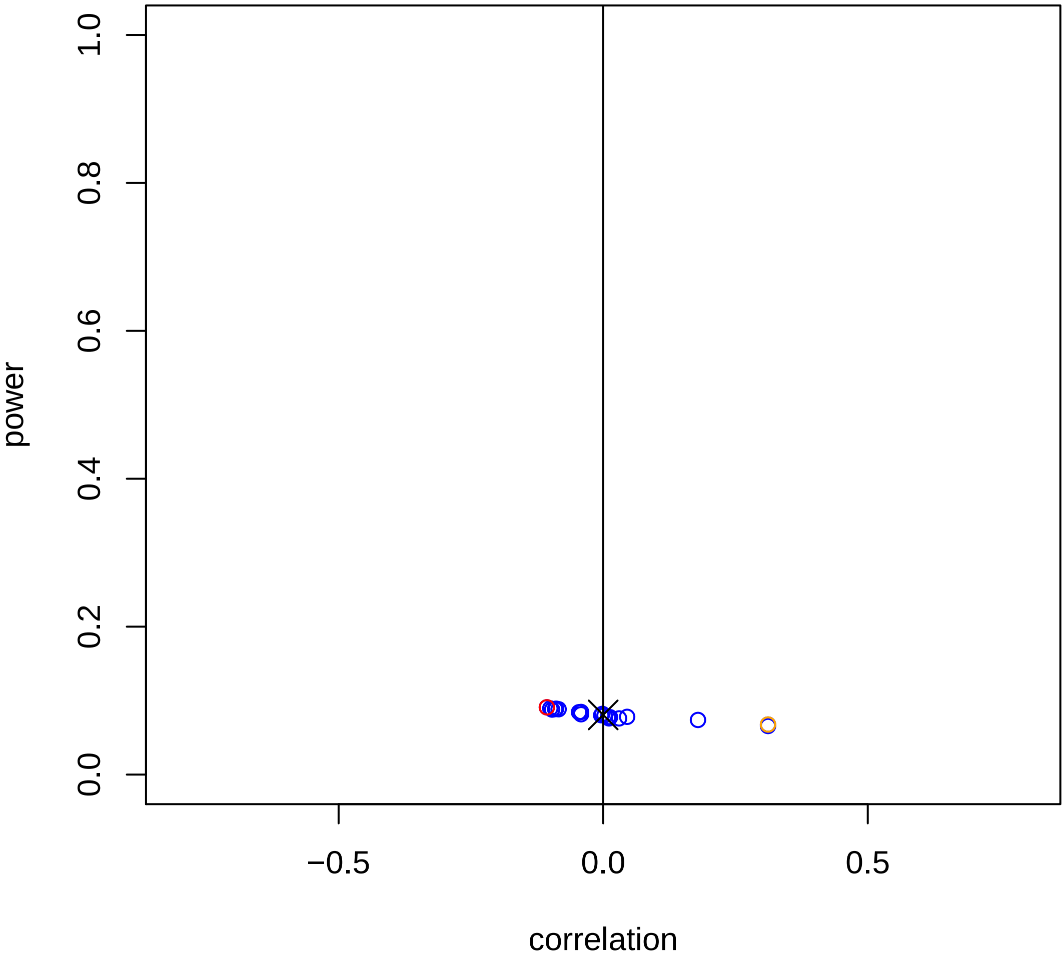


alpha: 0.1, n: 0.1K, datasetNum:10K, cor=−0.933, mu: 1.66, pi: 0.0014

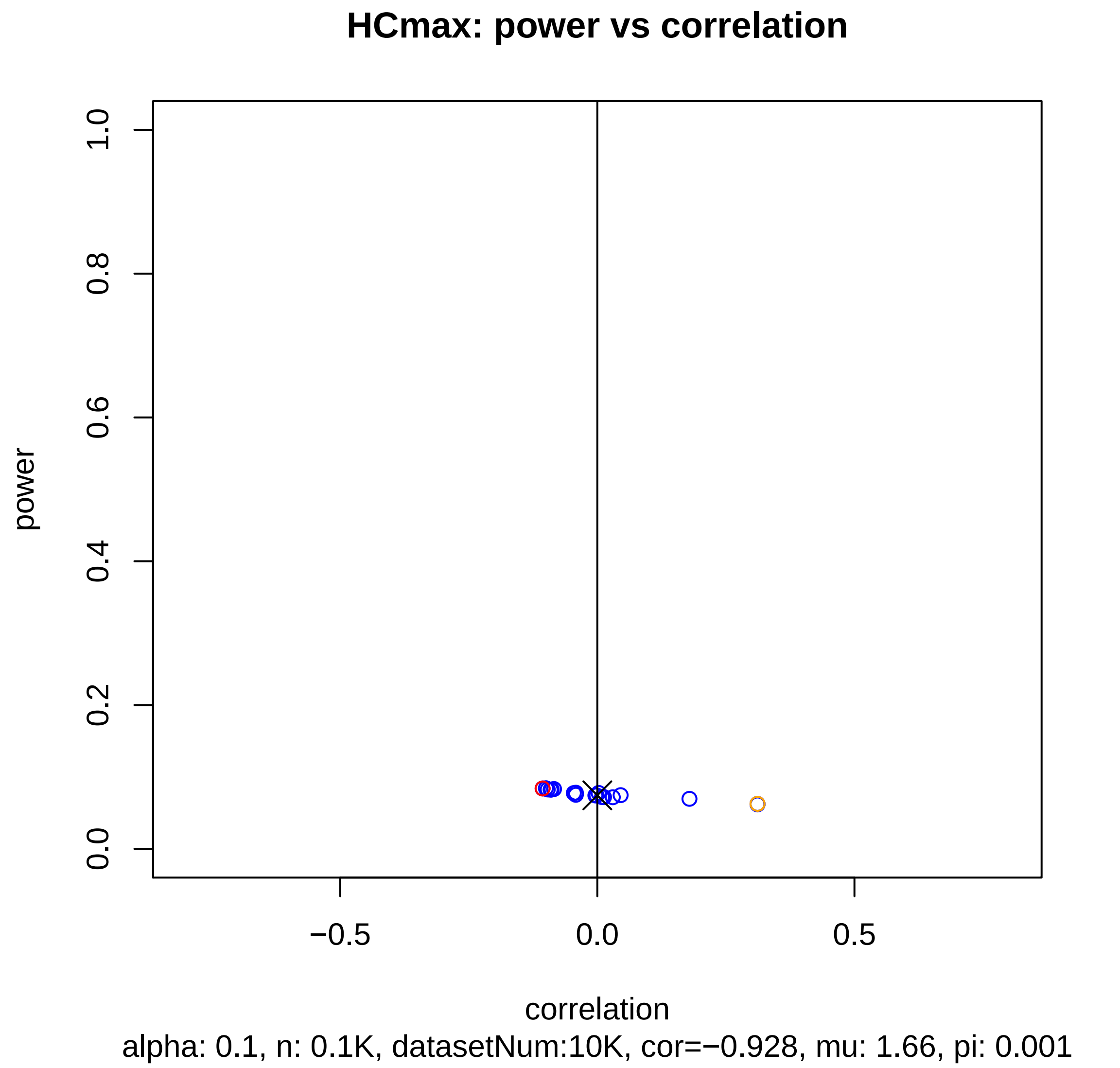
HCmax: power vs correlation
power
1.0
0.8
0.6
0.4
0.2
0.0
−0.5
0.0
0.5
correlation
alpha: 0.1, n: 0.1K, datasetNum:10K, cor=−0.928, mu: 1.66, pi: 0.001

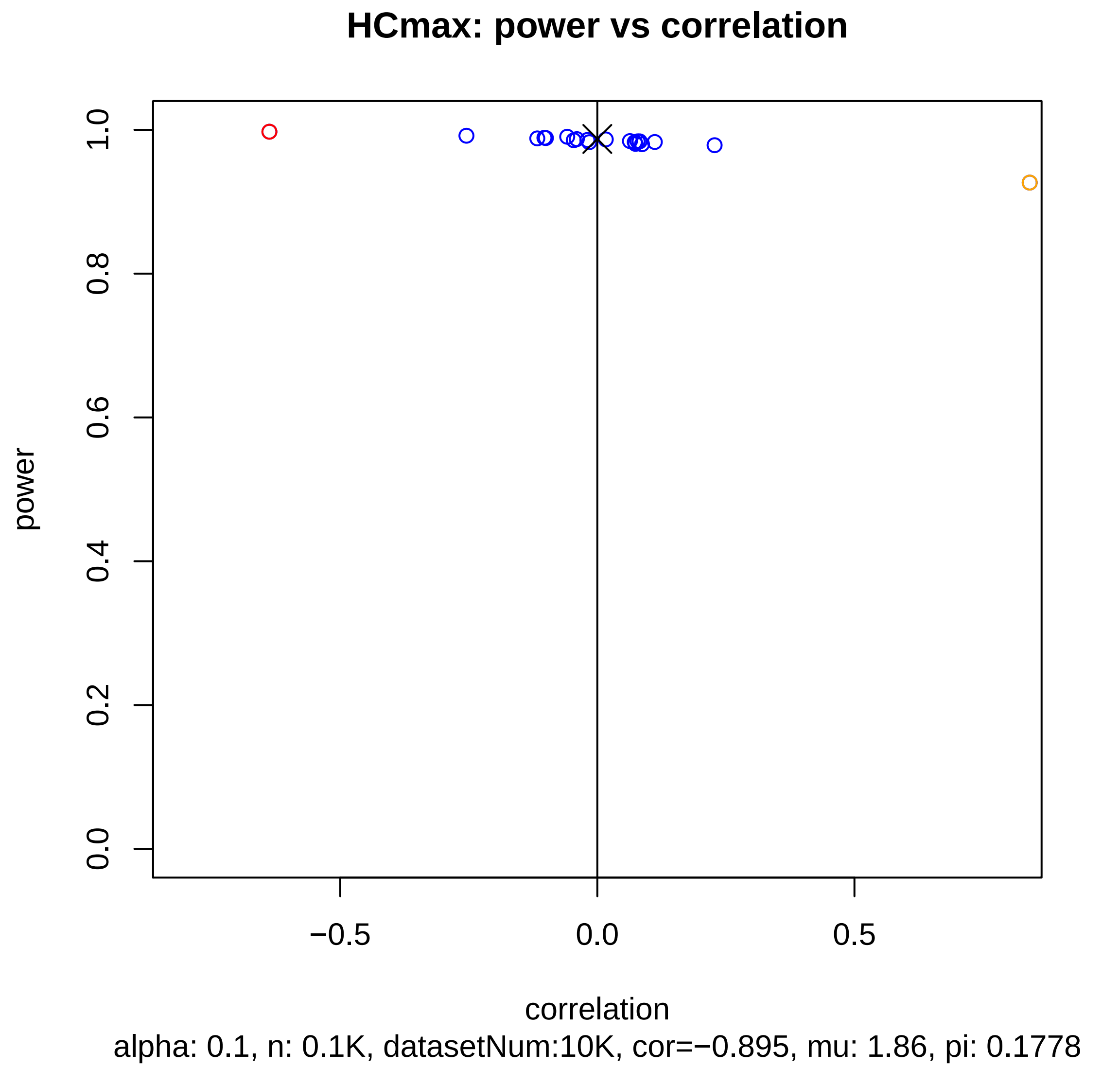

HCmax: power vs correlation
power
1.0
0.8
0.6
0.4
0.2
0.0
−0.5
0.0
0.5
correlation
alpha: 0.1, n: 0.1K, datasetNum:10K, cor=−0.895, mu: 1.86, pi: 0.1778

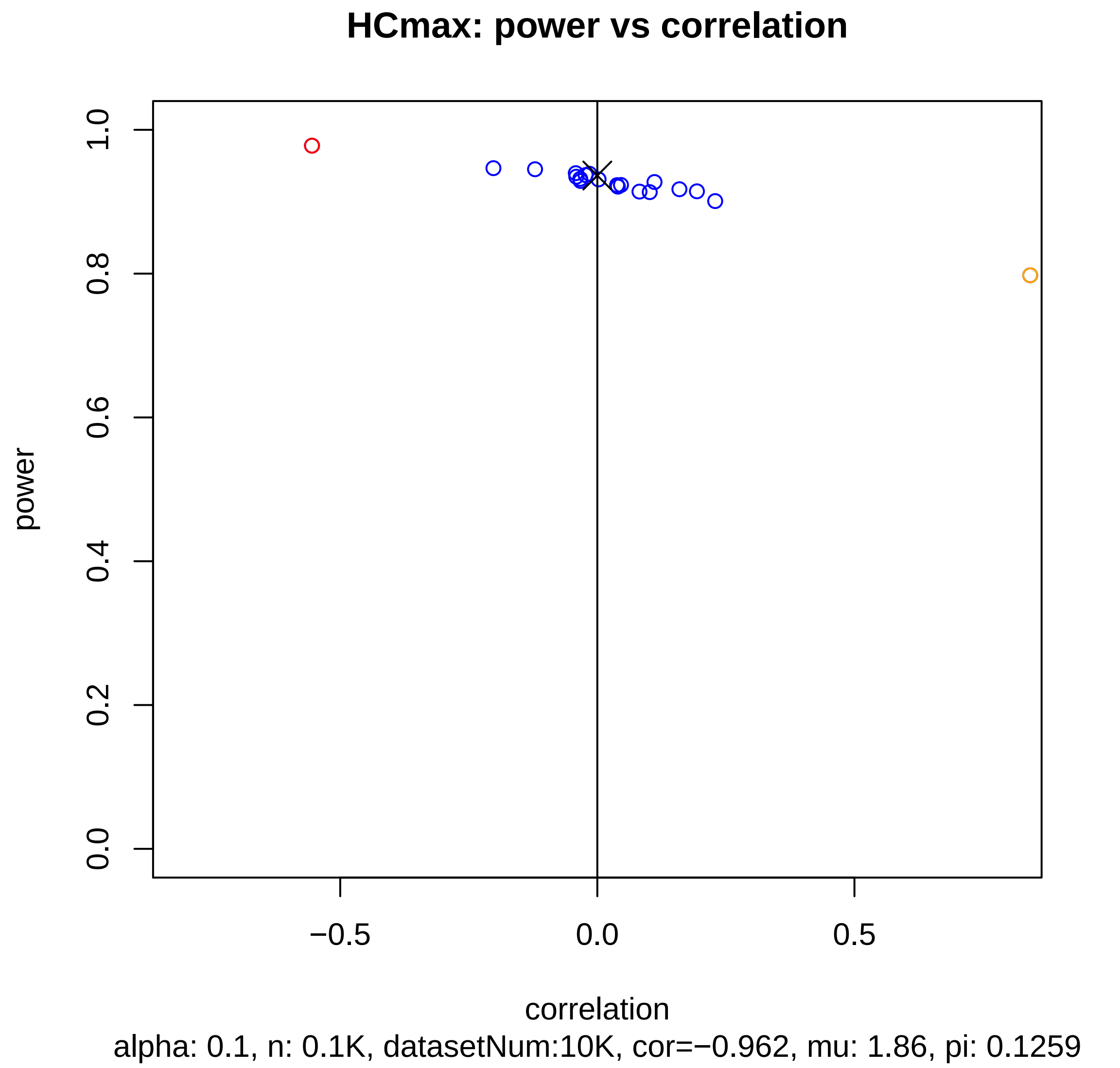

HCmax: power vs correlation
power
1.0
0.8
0.6
0.4
0.2
0.0
−0.5
0.0
0.5
correlation
alpha: 0.1, n: 0.1K, datasetNum:10K, cor=−0.962, mu: 1.86, pi: 0.1259

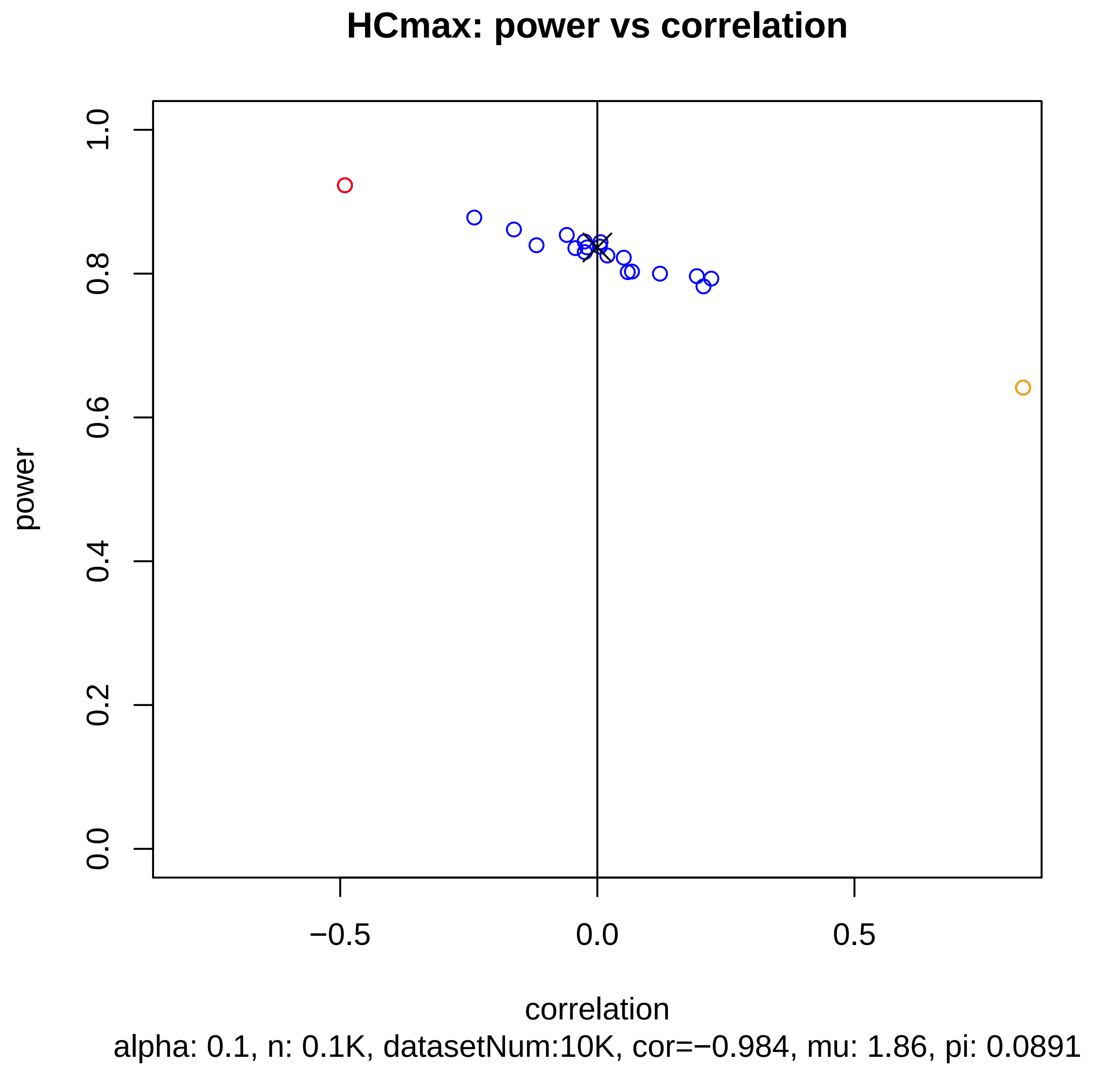
HCmax: power vs correlation
power
1.0
0.8
0.6
0.4
0.2
0.0
−0.5
0.0
0.5
correlation
alpha: 0.1, n: 0.1K, datasetNum:10K, cor=−0.984, mu: 1.86, pi: 0.0891

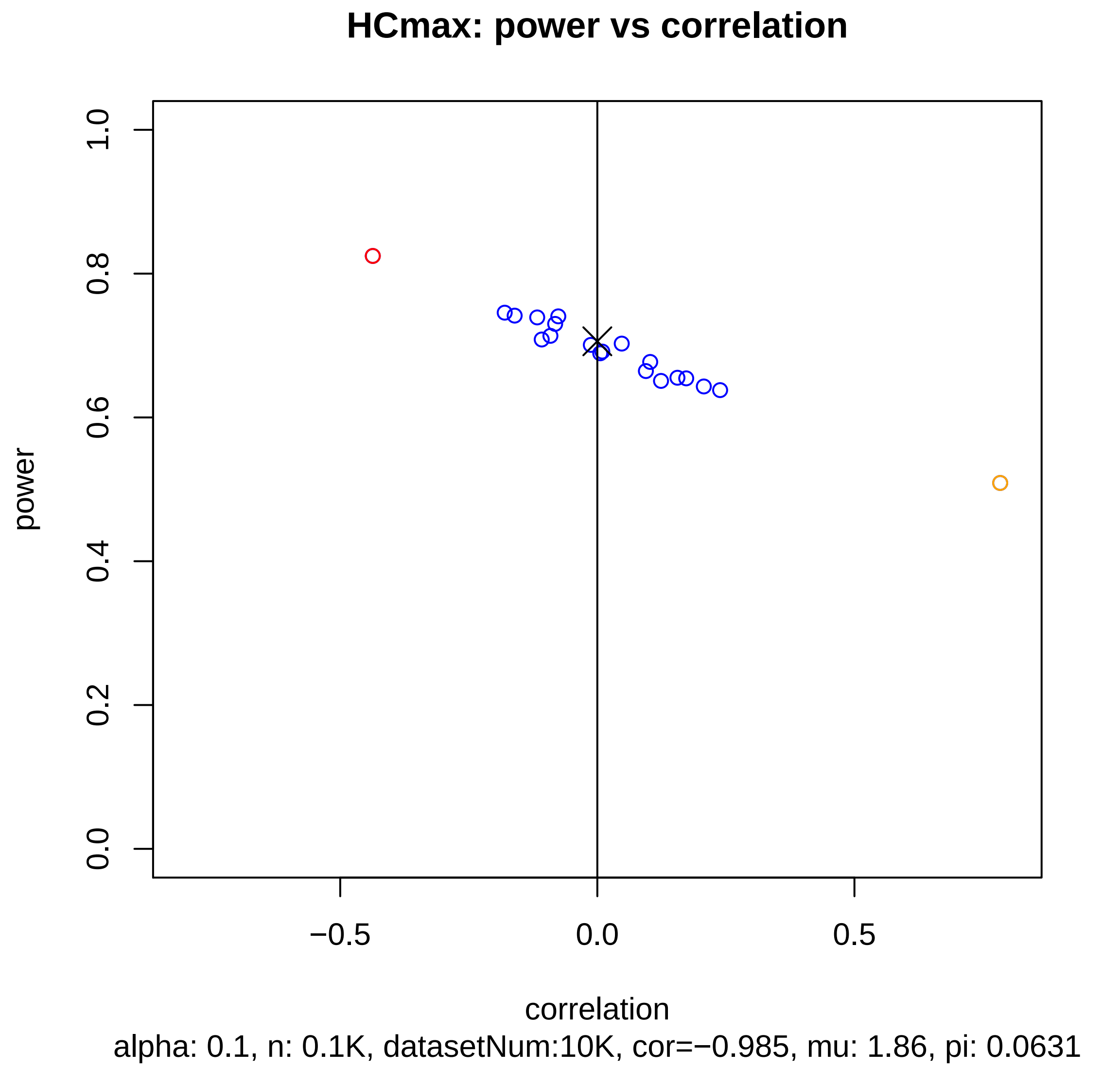
HCmax: power vs correlation
power
1.0
0.8
0.6
0.4
0.2
0.0
−0.5
0.0
0.5
correlation
alpha: 0.1, n: 0.1K, datasetNum:10K, cor=−0.985, mu: 1.86, pi: 0.0631

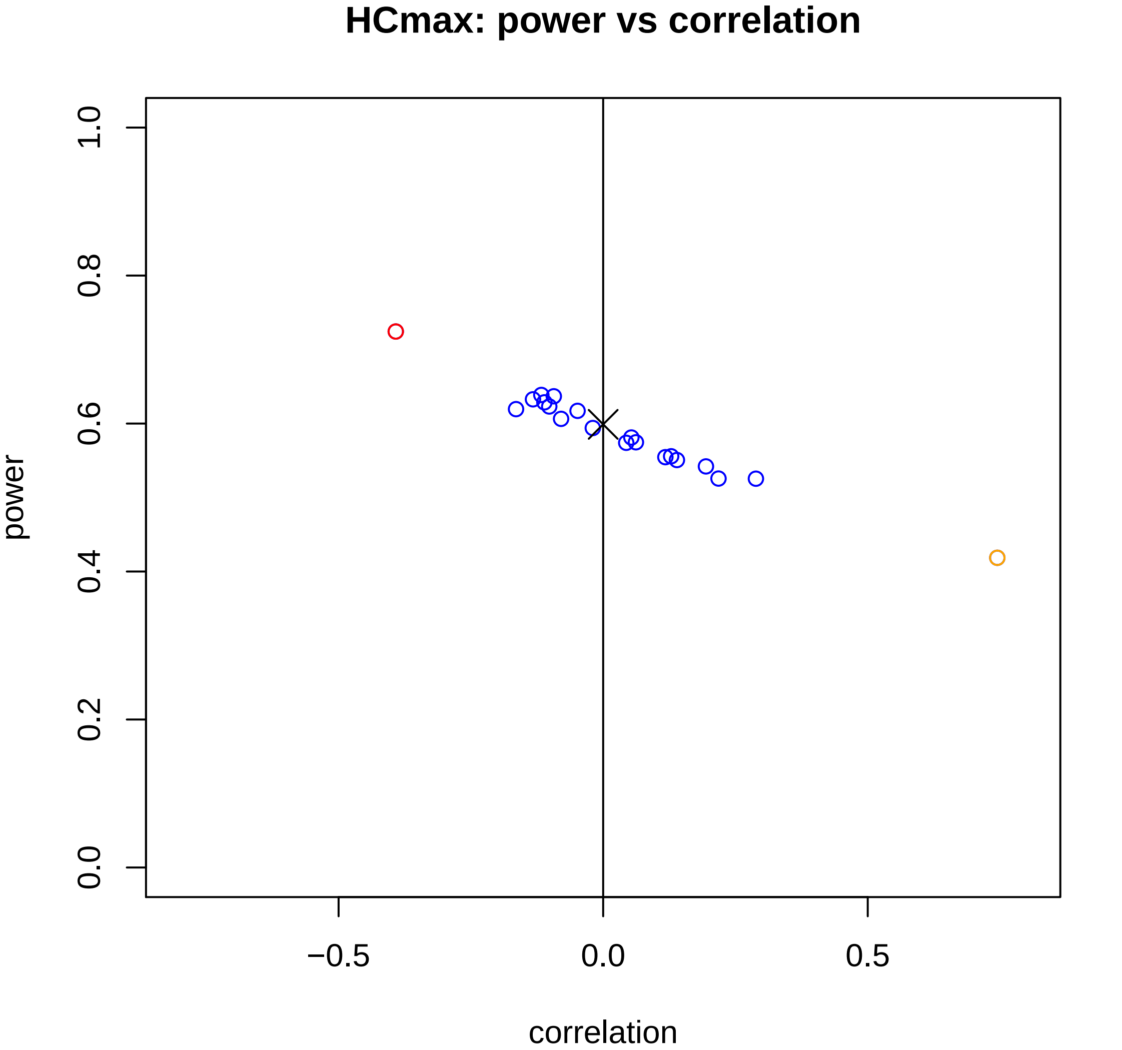

alpha: 0.1, n: 0.1K, datasetNum:10K, cor=−0.982, mu: 1.86, pi: 0.0447

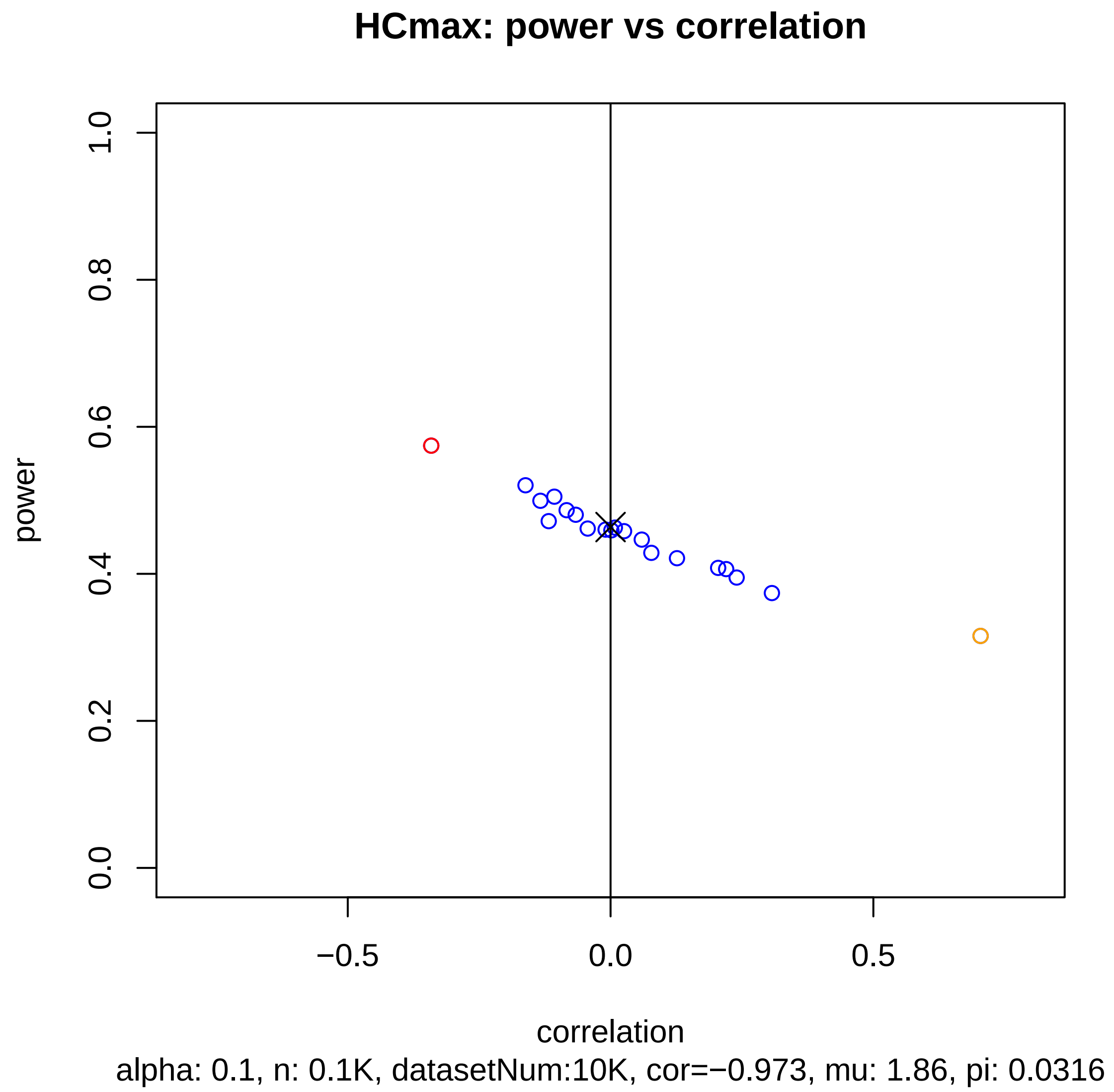
HCmax: power vs correlation
power
0.0
0.2
0.4
0.6
0.8
1.0
−0.5
0.0
0.5
correlation
alpha: 0.1, n: 0.1K, datasetNum:10K, cor=−0.973, mu: 1.86, pi: 0.0316

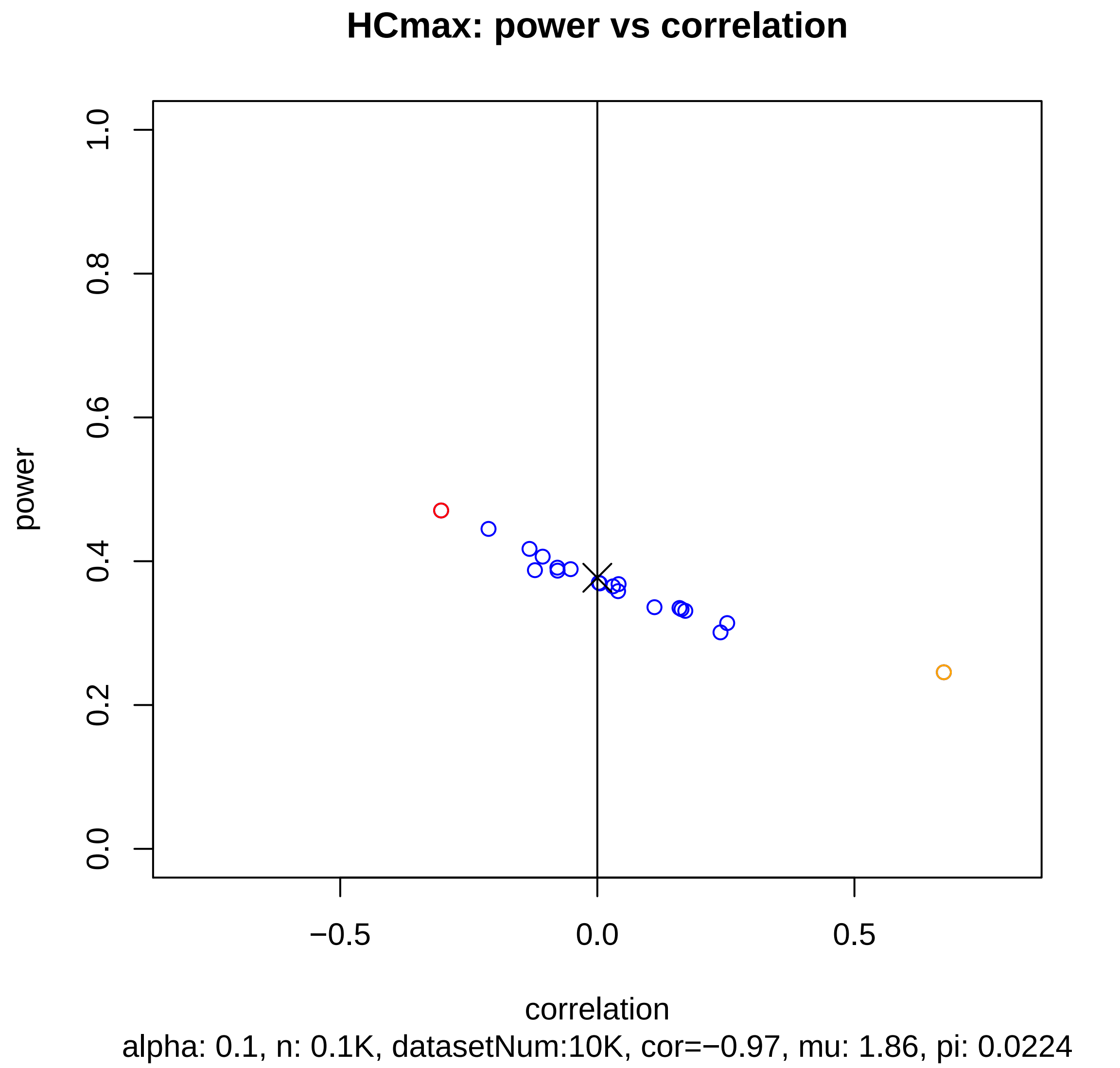

HCmax: power vs correlation
power
1.0
0.8
0.6
0.4
0.2
0.0
−0.5
0.0
0.5
correlation
alpha: 0.1, n: 0.1K, datasetNum:10K, cor=−0.97, mu: 1.86, pi: 0.0224

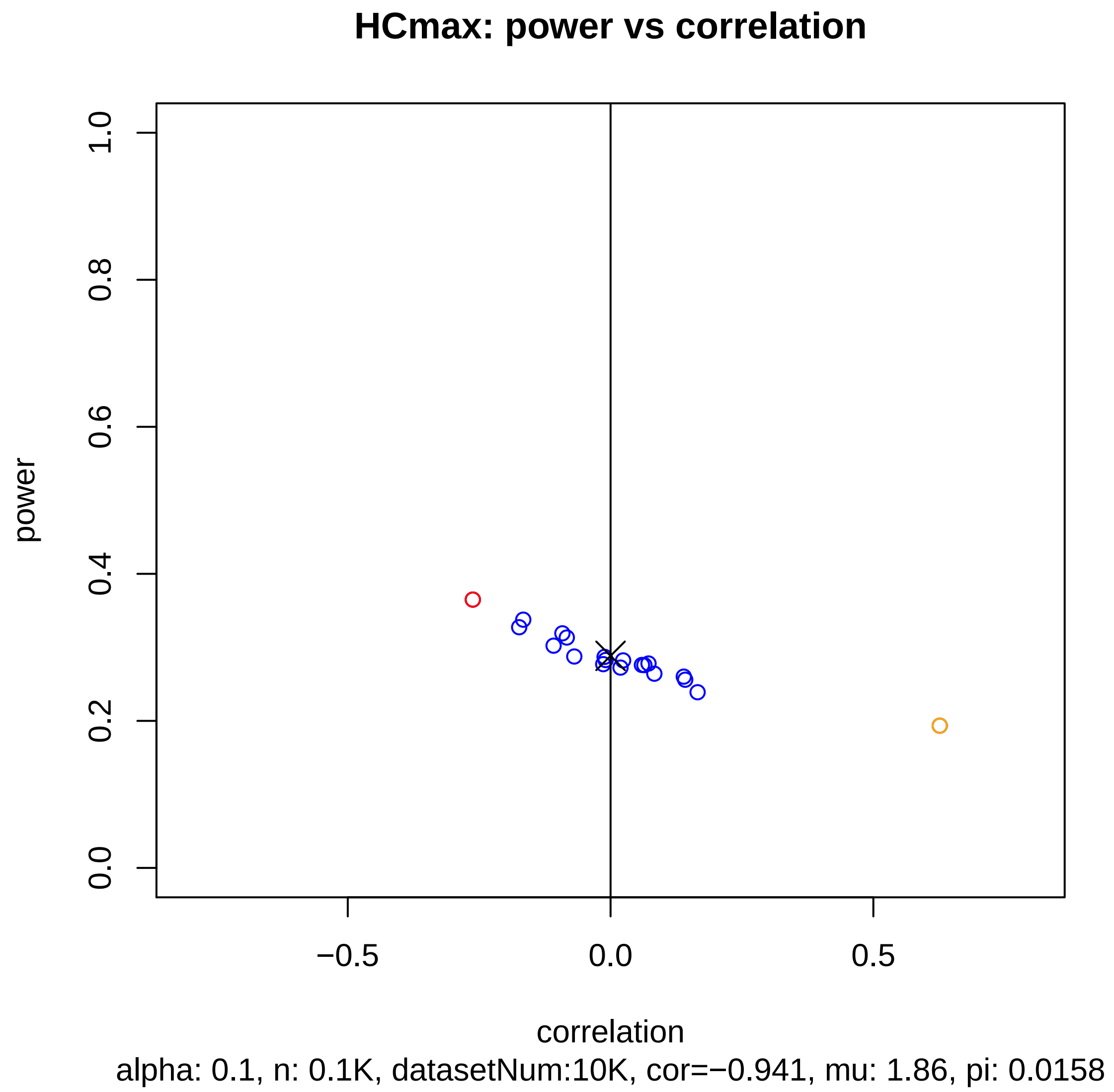

HCmax: power vs correlation
power
0.0
0.2
0.4
0.6
0.8
1.0
−0.5
0.0
0.5
correlation
alpha: 0.1, n: 0.1K, datasetNum:10K, cor=−0.941, mu: 1.86, pi: 0.0158

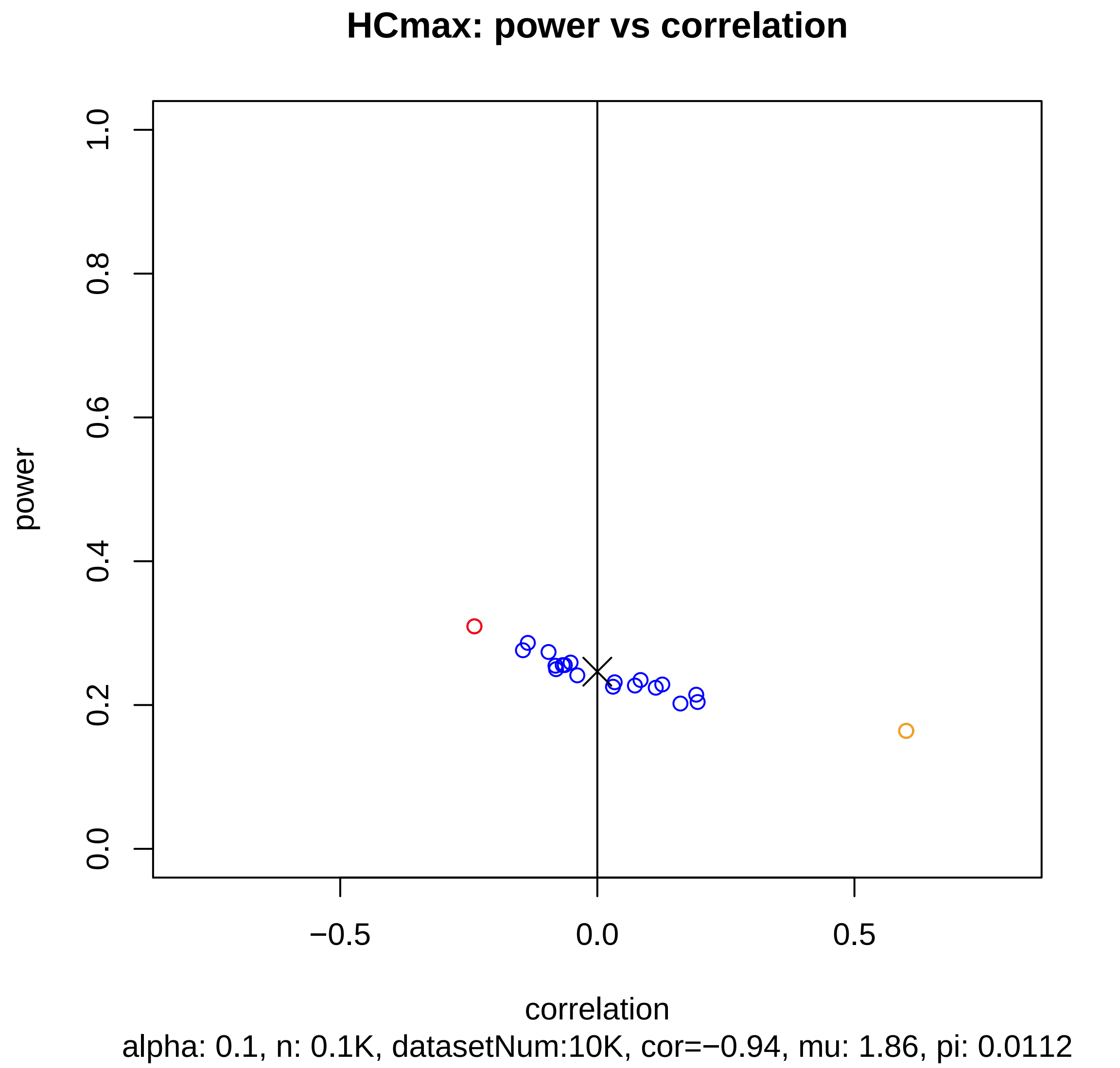
HCmax: power vs correlation
power
0.0
0.2
0.4
0.6
0.8
1.0
−0.5
0.0
0.5
correlation
alpha: 0.1, n: 0.1K, datasetNum:10K, cor=−0.94, mu: 1.86, pi: 0.0112

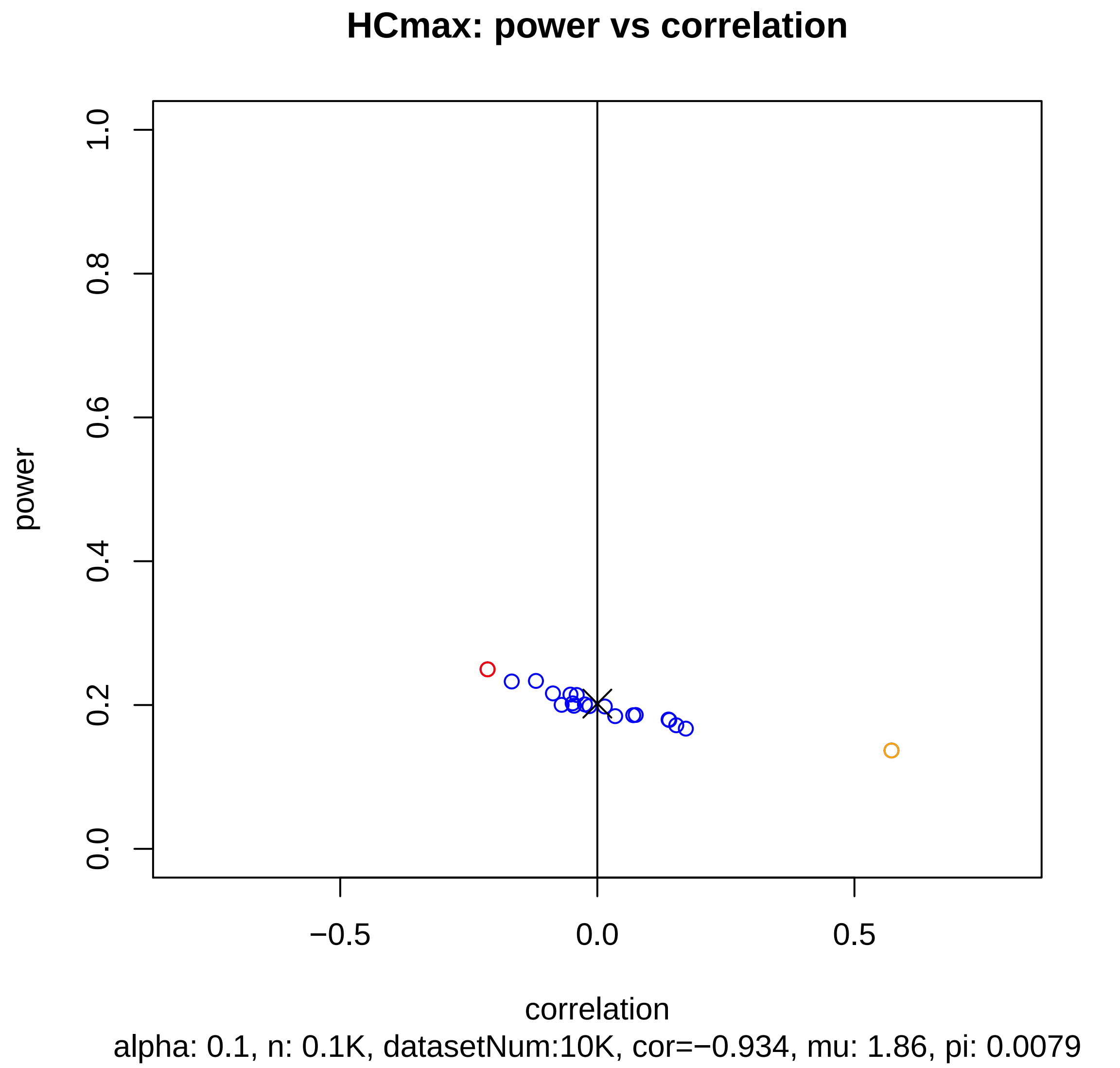
HCmax: power vs correlation
power
1.0
0.8
0.6
0.4
0.2
0.0
−0.5
0.0
0.5
correlation
alpha: 0.1, n: 0.1K, datasetNum:10K, cor=−0.934, mu: 1.86, pi: 0.0079

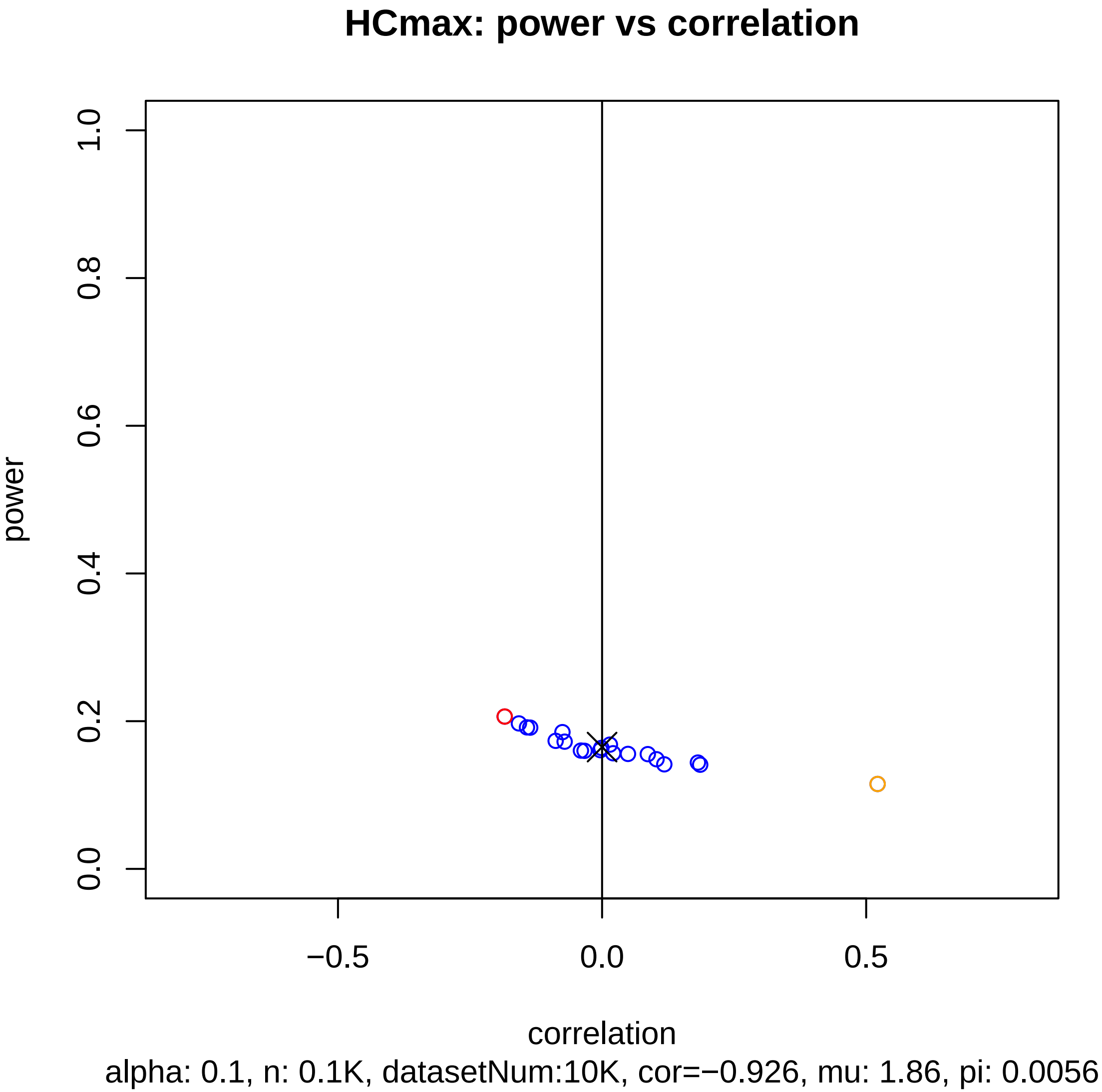

HCmax: power vs correlation
power
1.0
0.8
0.6
0.4
0.2
0.0
−0.5
0.0
0.5
correlation
alpha: 0.1, n: 0.1K, datasetNum:10K, cor=−0.926, mu: 1.86, pi: 0.0056

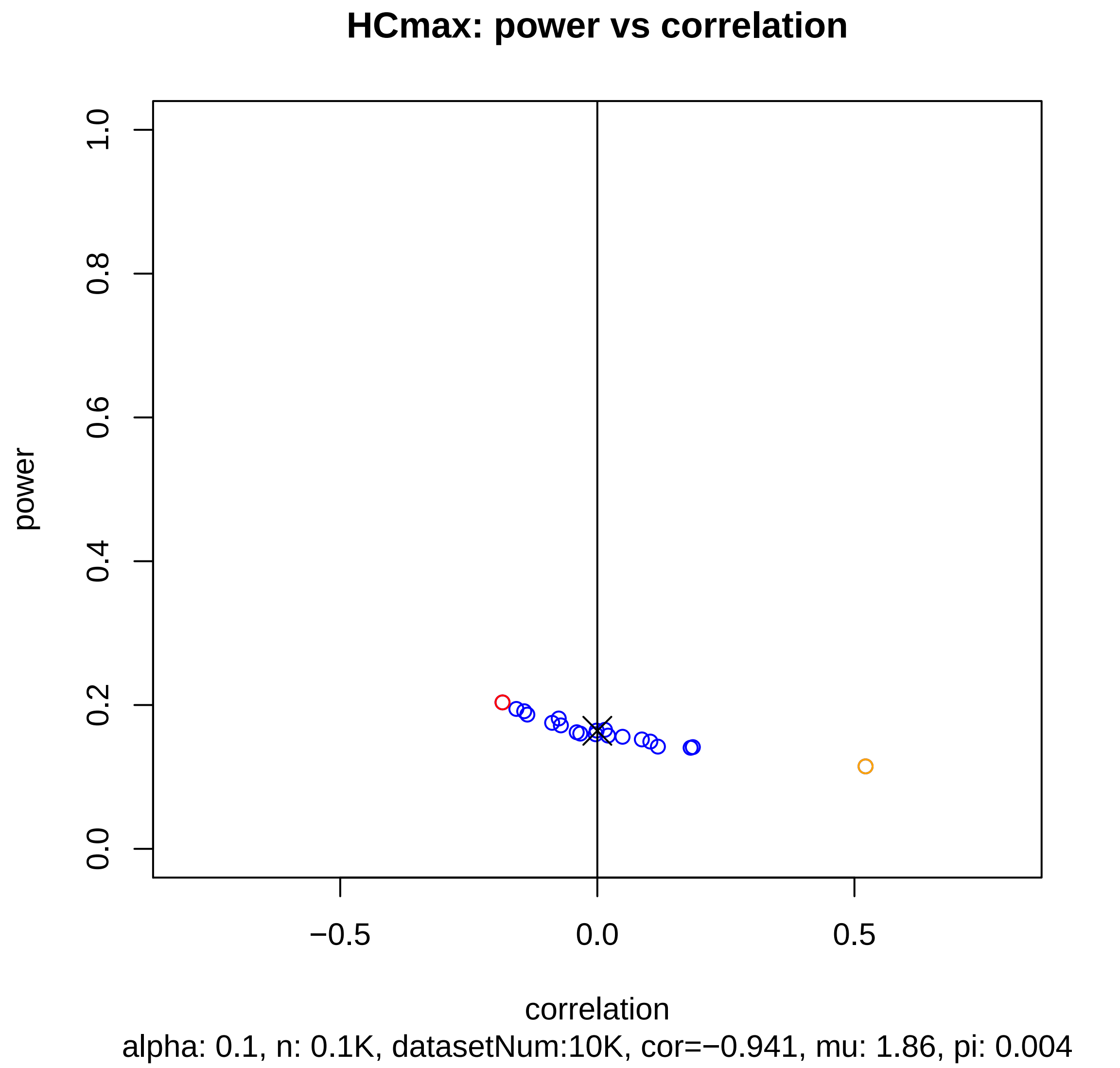
HCmax: power vs correlation
power
1.0
0.8
0.6
0.4
0.2
0.0
−0.5
0.0
0.5
correlation
alpha: 0.1, n: 0.1K, datasetNum:10K, cor=−0.941, mu: 1.86, pi: 0.004

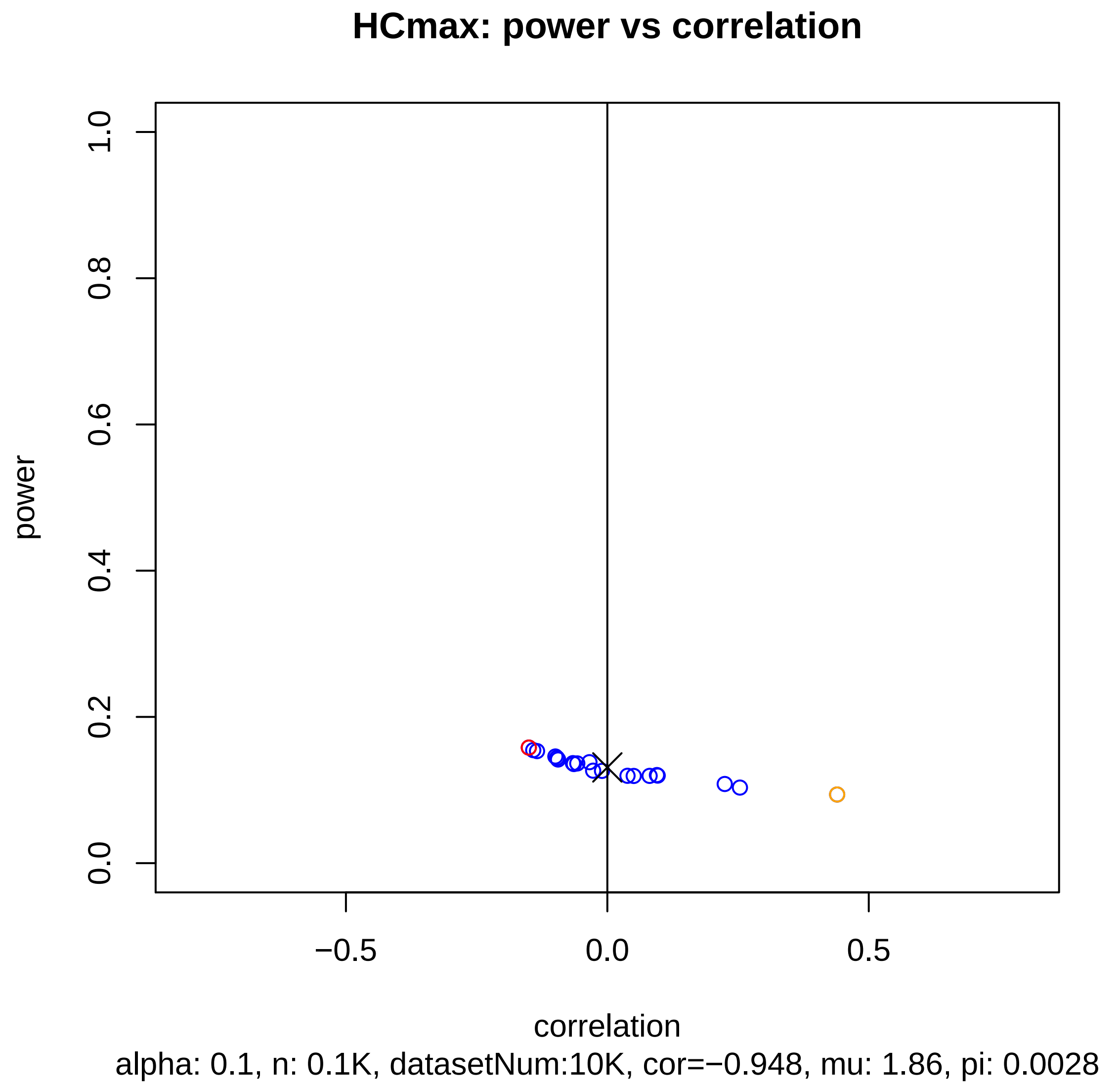
HCmax: power vs correlation
power
1.0
0.8
0.6
0.4
0.2
0.0
−0.5
0.0
0.5
correlation
alpha: 0.1, n: 0.1K, datasetNum:10K, cor=−0.948, mu: 1.86, pi: 0.0028

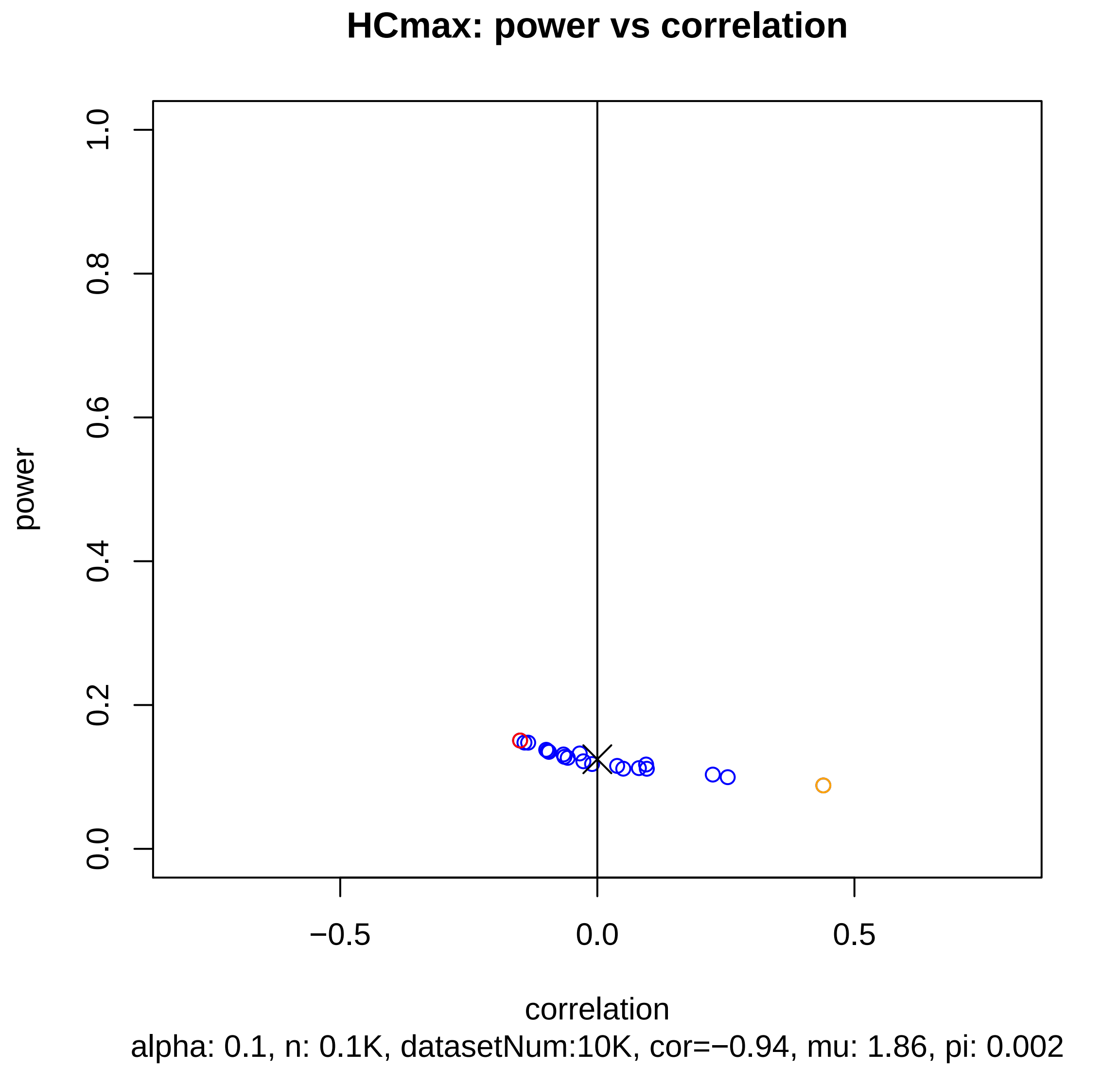

HCmax: power vs correlation
power
0.0
0.2
0.4
0.6
0.8
1.0
−0.5
0.0
0.5
correlation
alpha: 0.1, n: 0.1K, datasetNum:10K, cor=−0.94, mu: 1.86, pi: 0.002

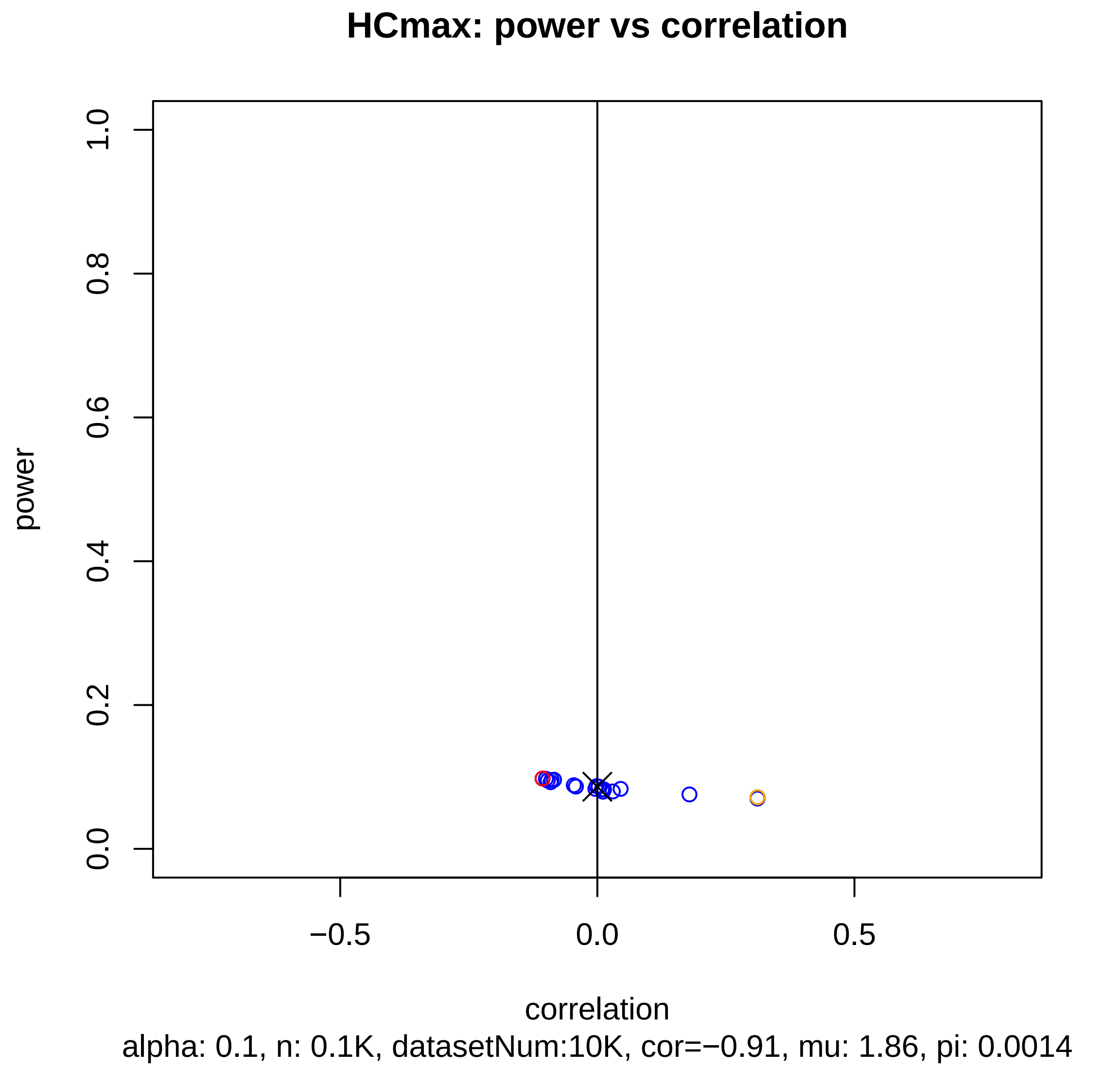
HCmax: power vs correlation
power
1.0
0.8
0.6
0.4
0.2
0.0
−0.5
0.0
0.5
correlation
alpha: 0.1, n: 0.1K, datasetNum:10K, cor=−0.91, mu: 1.86, pi: 0.0014

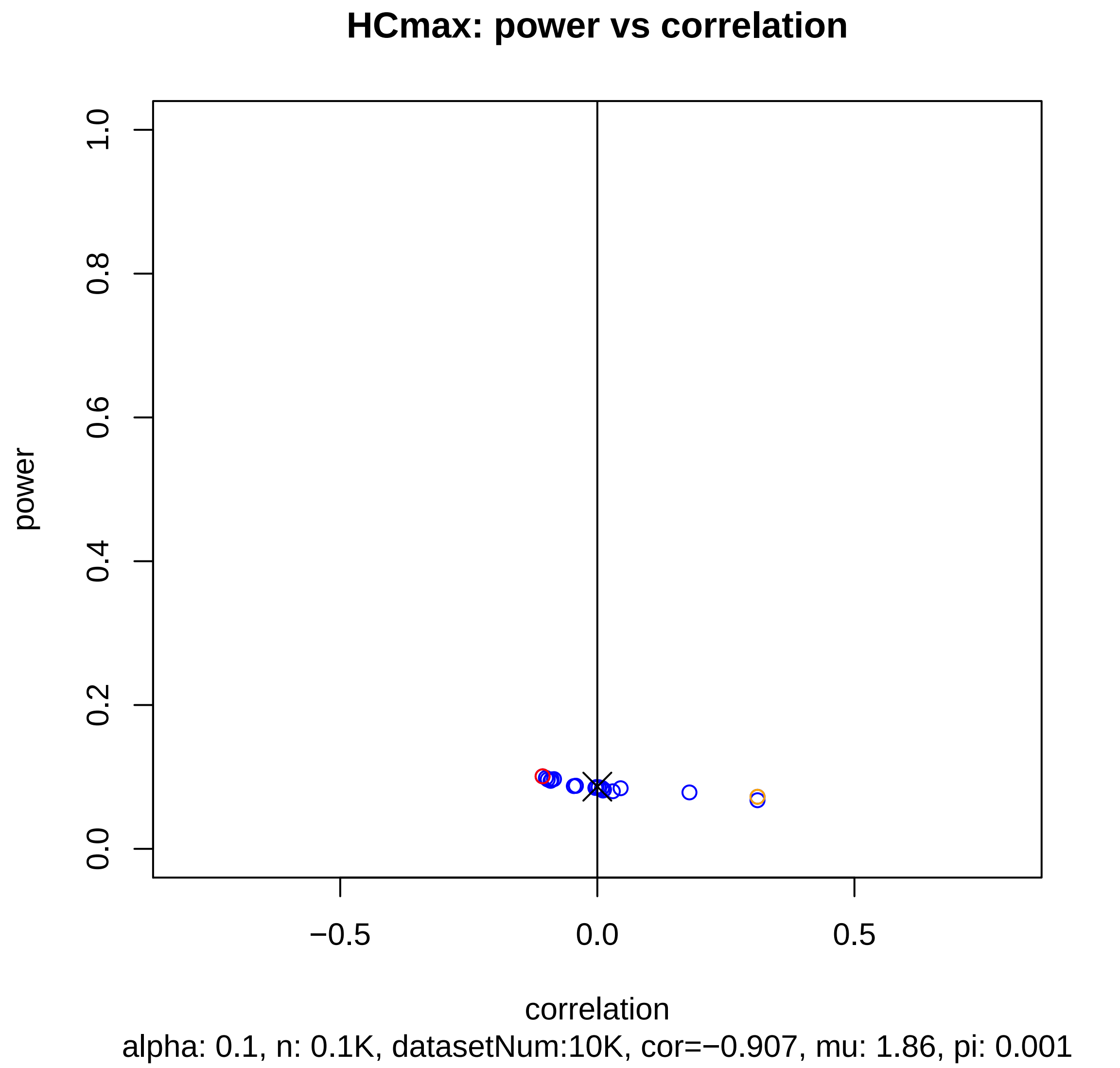

HCmax: power vs correlation
power
1.0
0.8
0.6
0.4
0.2
0.0
−0.5
0.0
0.5
correlation
alpha: 0.1, n: 0.1K, datasetNum:10K, cor=−0.907, mu: 1.86, pi: 0.001

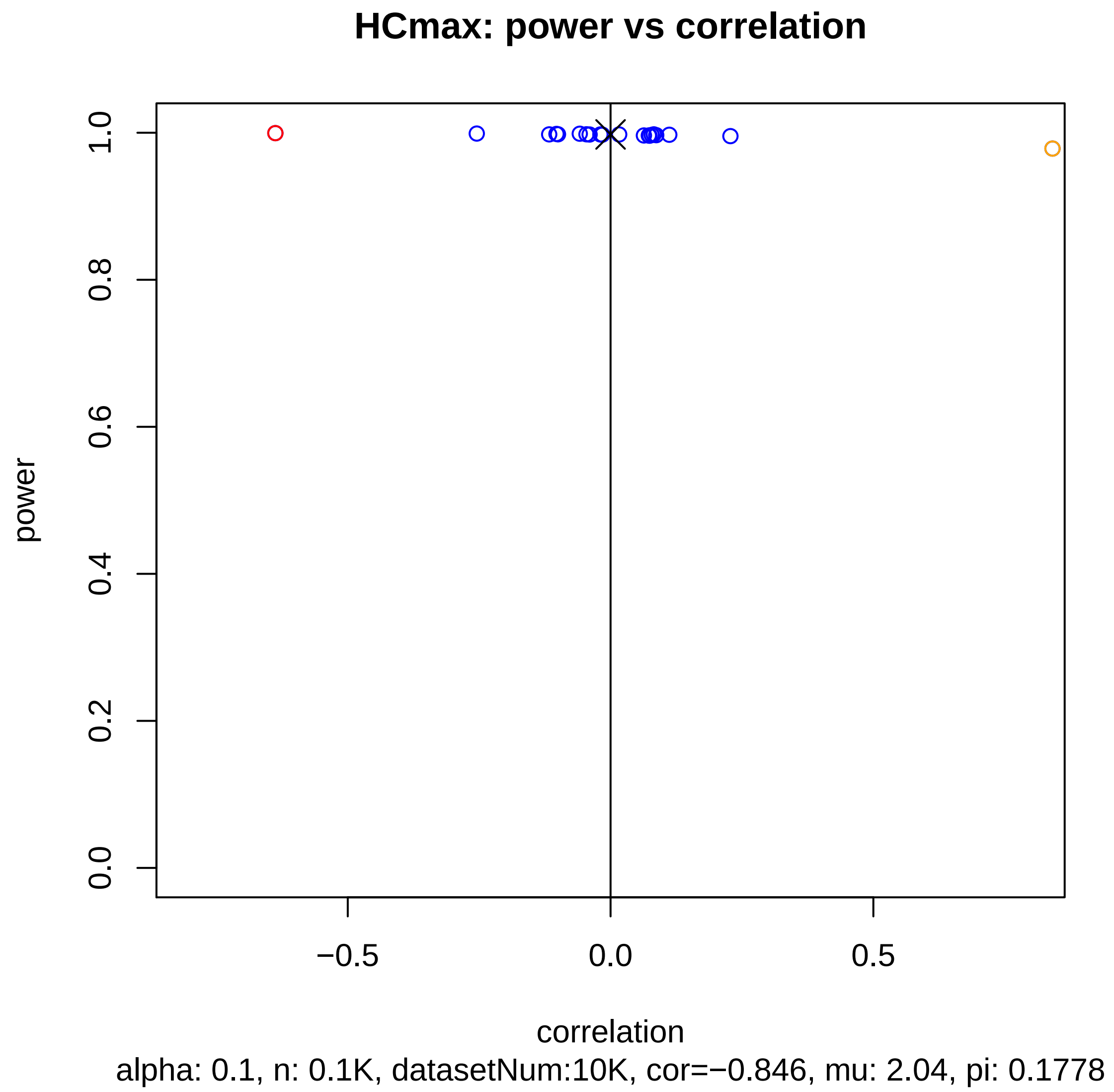
HCmax: power vs correlation
power
1.0
0.8
0.6
0.4
0.2
0.0
−0.5
0.0
0.5
correlation
alpha: 0.1, n: 0.1K, datasetNum:10K, cor=−0.846, mu: 2.04, pi: 0.1778

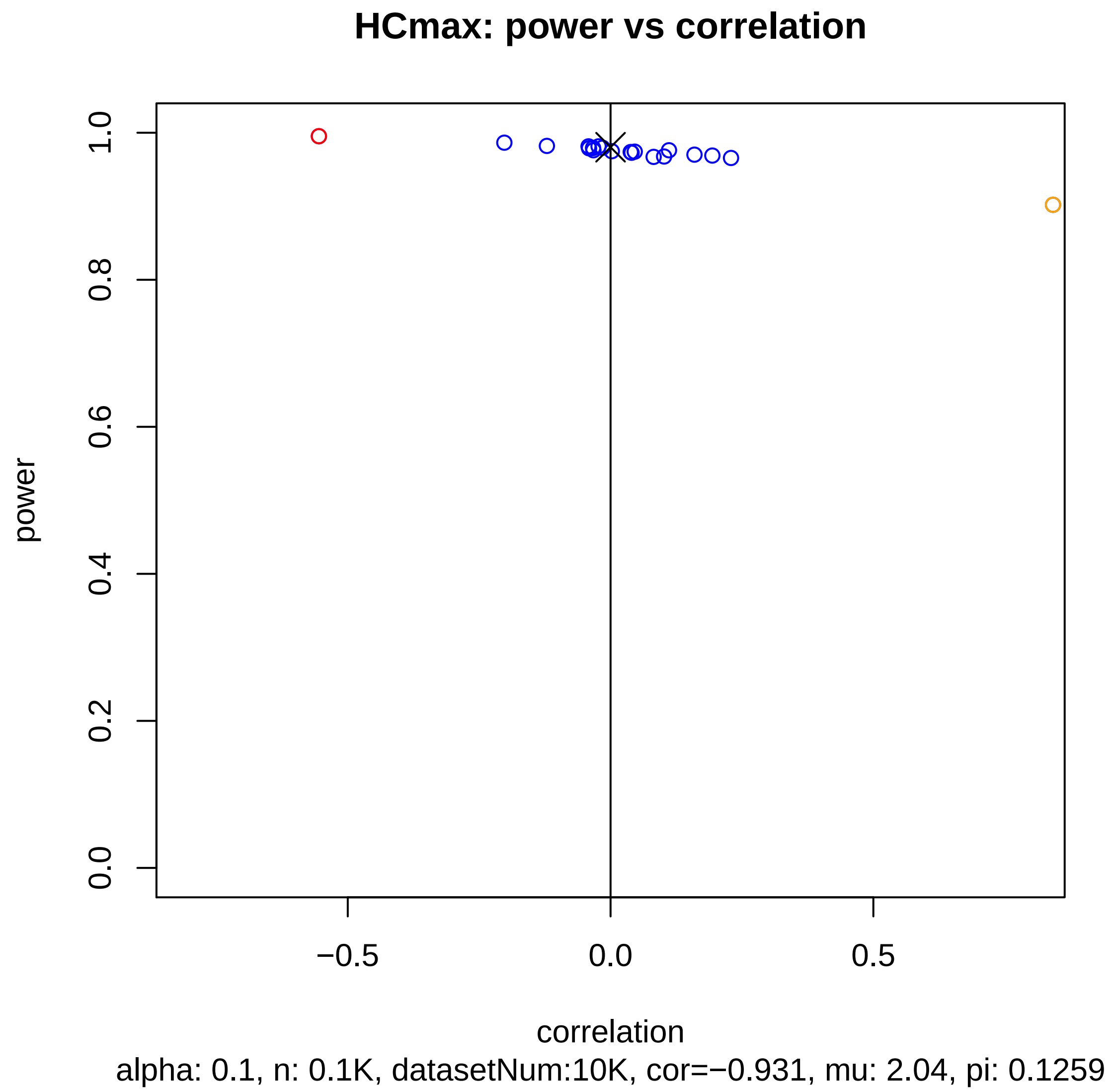
HCmax: power vs correlation
power
1.0
0.8
0.6
0.4
0.2
0.0
−0.5
0.0
0.5
correlation
alpha: 0.1, n: 0.1K, datasetNum:10K, cor=−0.931, mu: 2.04, pi: 0.1259

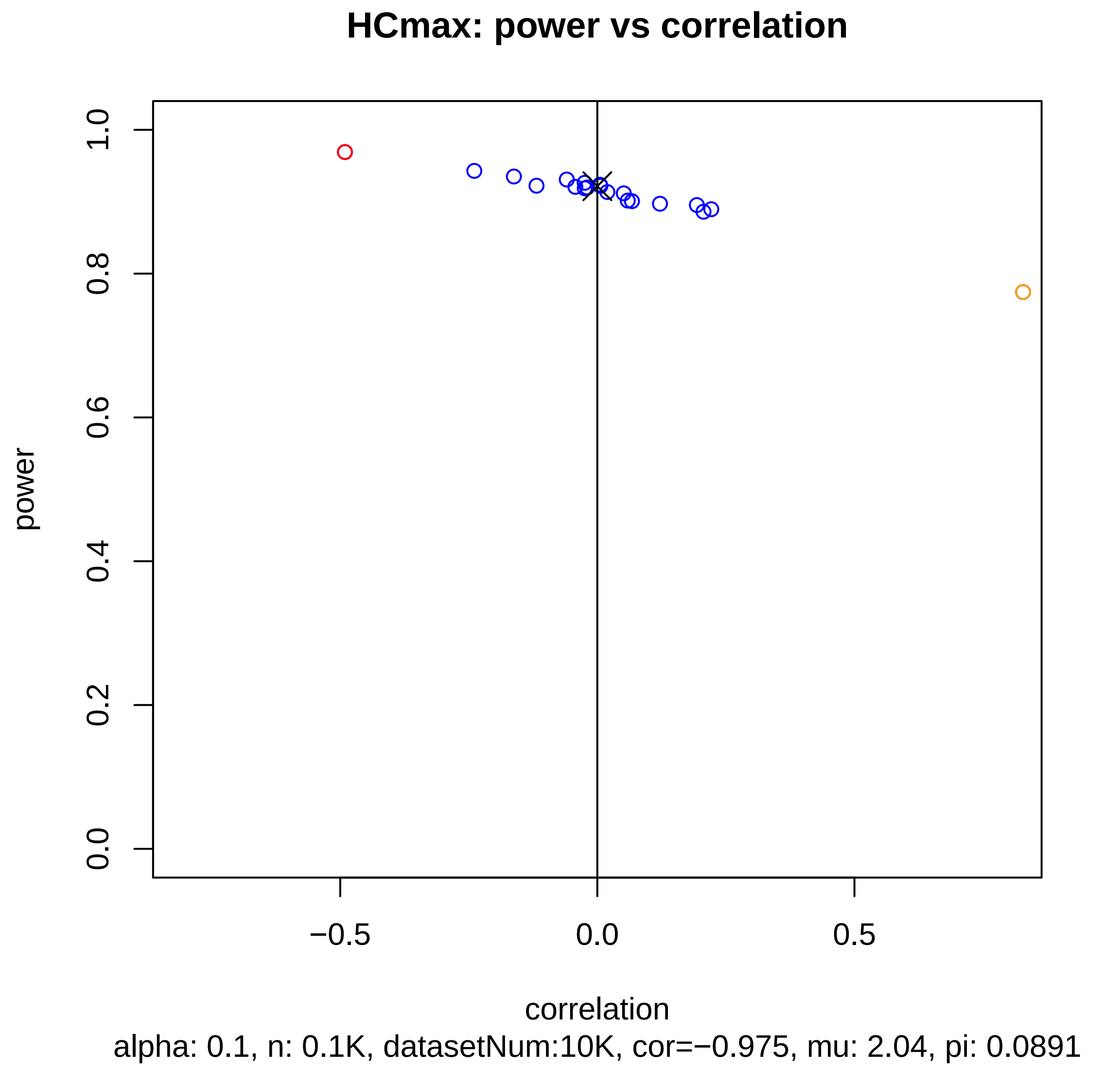

HCmax: power vs correlation
power
1.0
0.8
0.6
0.4
0.2
0.0
−0.5
0.0
0.5
correlation
alpha: 0.1, n: 0.1K, datasetNum:10K, cor=−0.975, mu: 2.04, pi: 0.0891

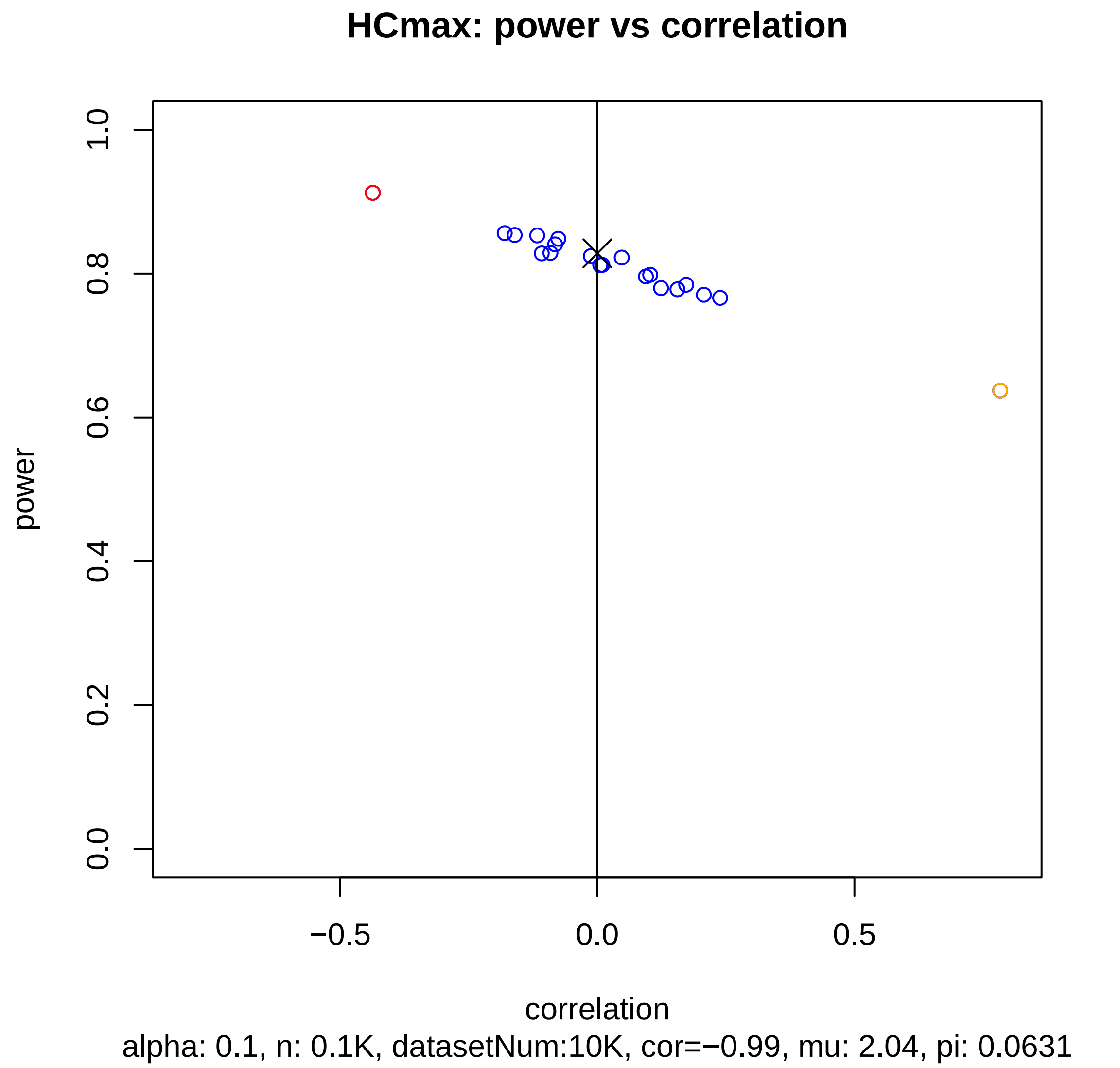

HCmax: power vs correlation
power
1.0
0.8
0.6
0.4
0.2
0.0
−0.5
0.0
0.5
correlation
alpha: 0.1, n: 0.1K, datasetNum:10K, cor=−0.99, mu: 2.04, pi: 0.0631

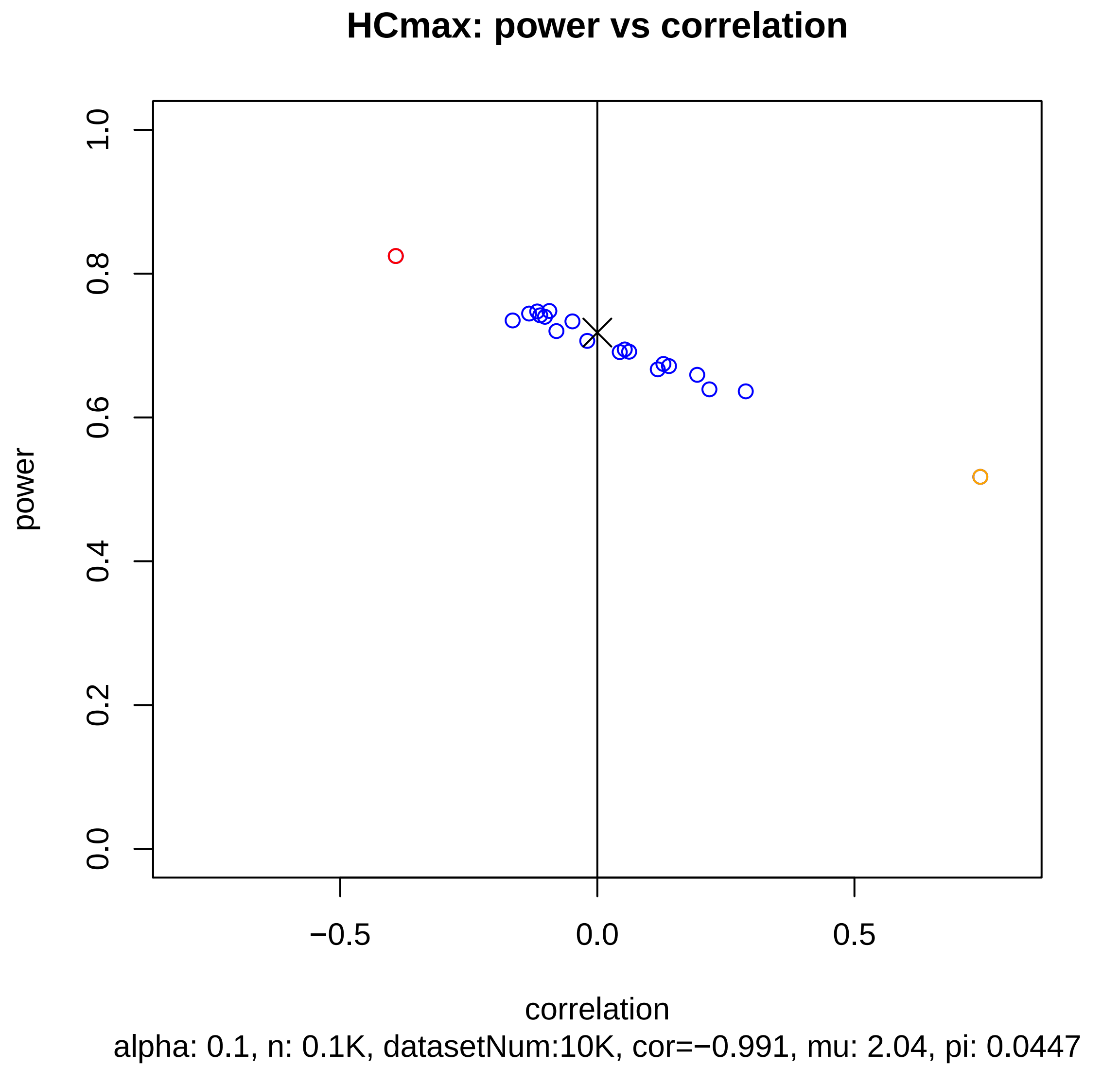
HCmax: power vs correlation
power
1.0
0.8
0.6
0.4
0.2
0.0
−0.5
0.0
0.5
correlation
alpha: 0.1, n: 0.1K, datasetNum:10K, cor=−0.991, mu: 2.04, pi: 0.0447

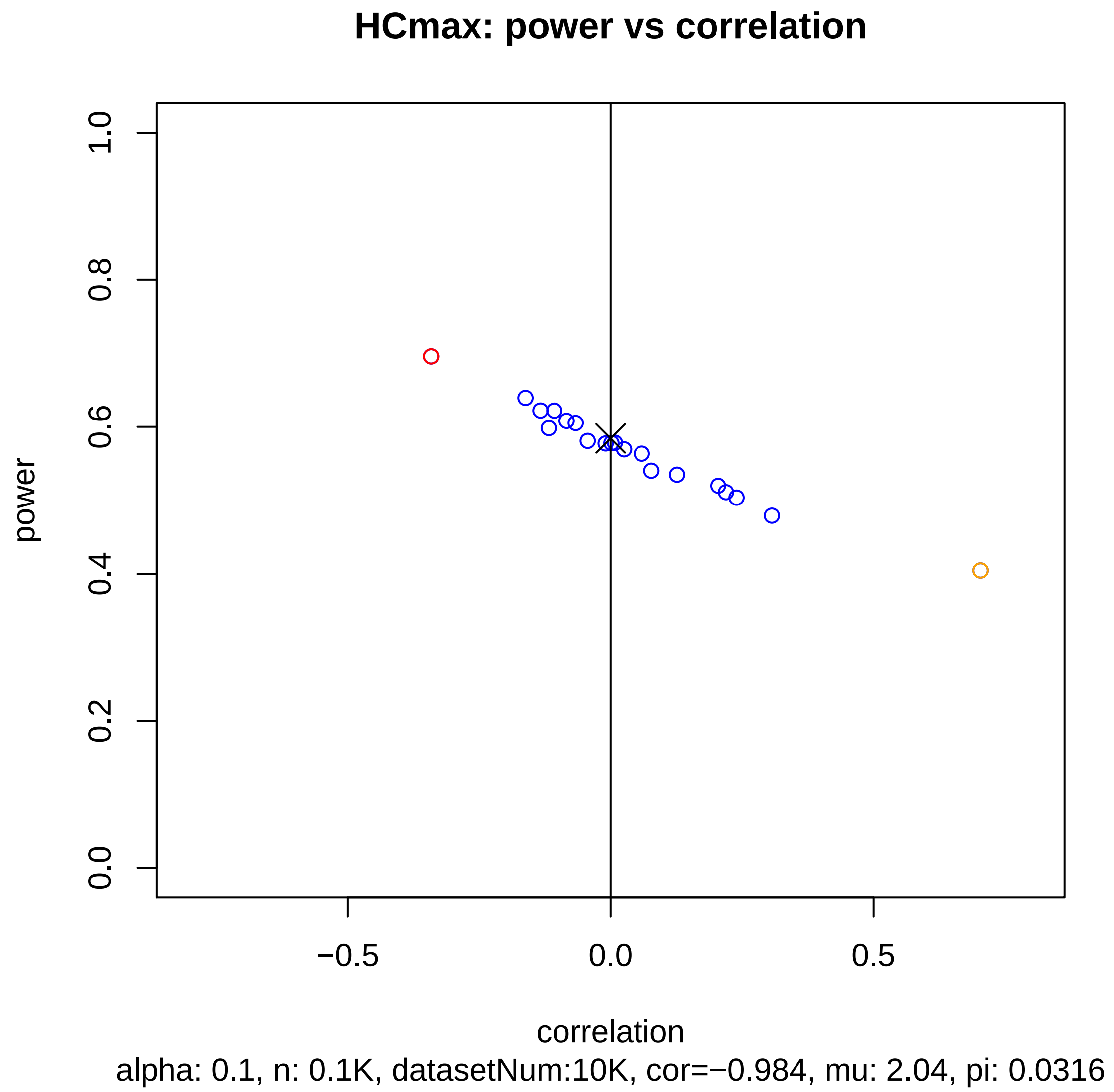
HCmax: power vs correlation
power
0.0
0.2
0.4
0.6
0.8
1.0
−0.5
0.0
0.5
correlation
alpha: 0.1, n: 0.1K, datasetNum:10K, cor=−0.984, mu: 2.04, pi: 0.0316

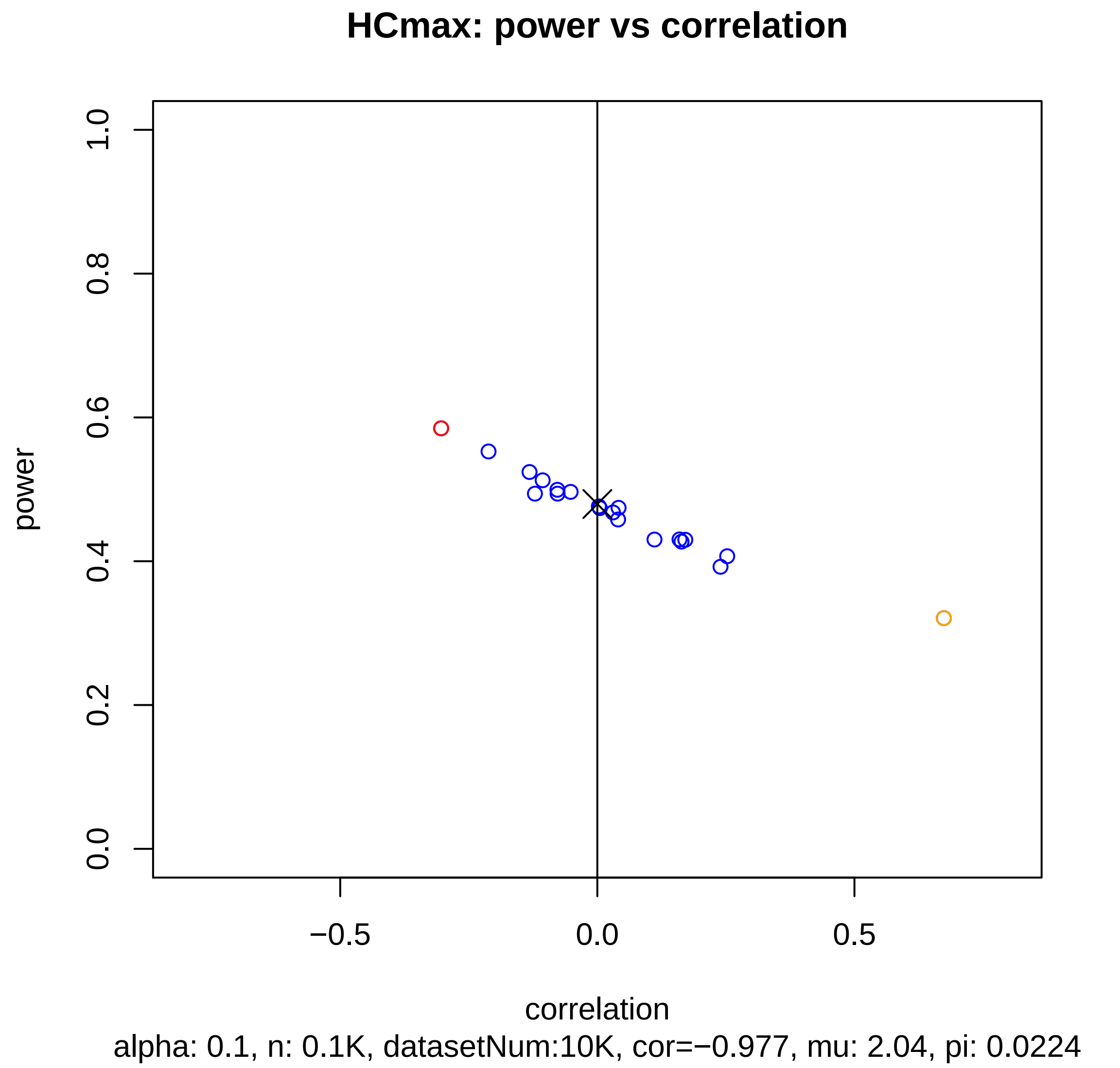
HCmax: power vs correlation
power
1.0
0.8
0.6
0.4
0.2
0.0
−0.5
0.0
0.5
correlation
alpha: 0.1, n: 0.1K, datasetNum:10K, cor=−0.977, mu: 2.04, pi: 0.0224

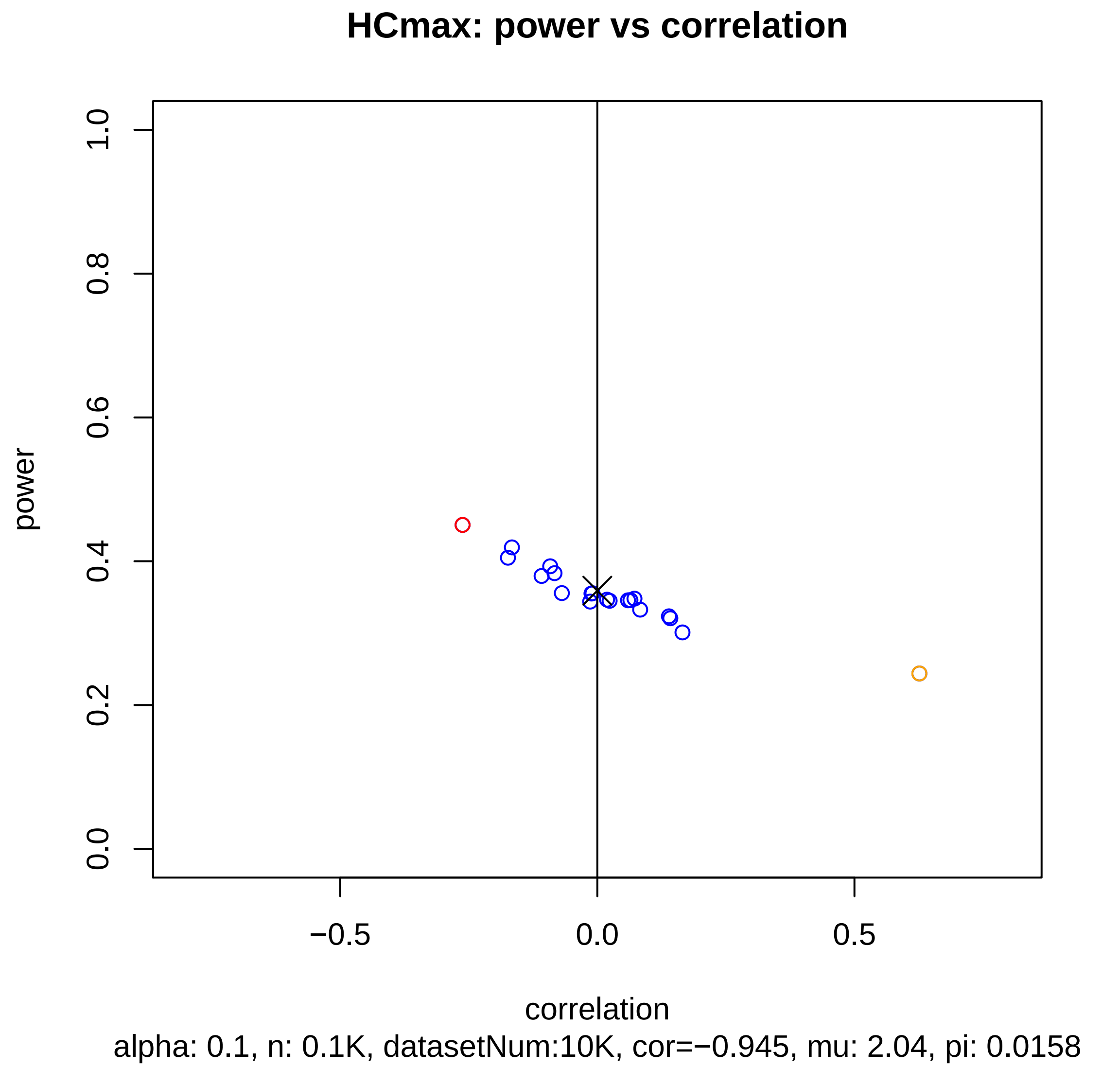
HCmax: power vs correlation
power
1.0
0.8
0.6
0.4
0.2
0.0
−0.5
0.0
0.5
correlation
alpha: 0.1, n: 0.1K, datasetNum:10K, cor=−0.945, mu: 2.04, pi: 0.0158

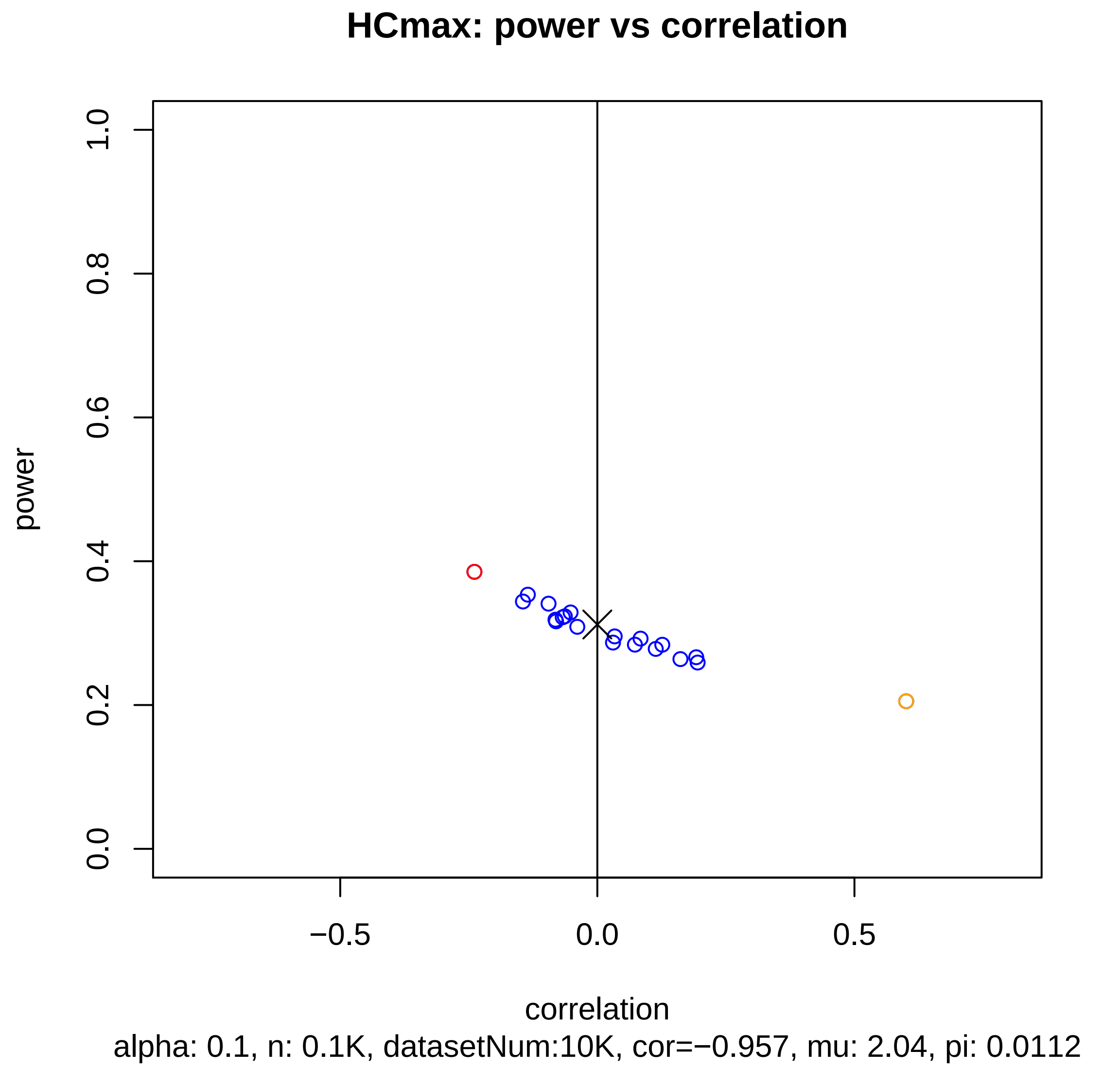
HCmax: power vs correlation
power
1.0
0.8
0.6
0.4
0.2
0.0
−0.5
0.0
0.5
correlation
alpha: 0.1, n: 0.1K, datasetNum:10K, cor=−0.957, mu: 2.04, pi: 0.0112

#### HCmax: power vs correlation

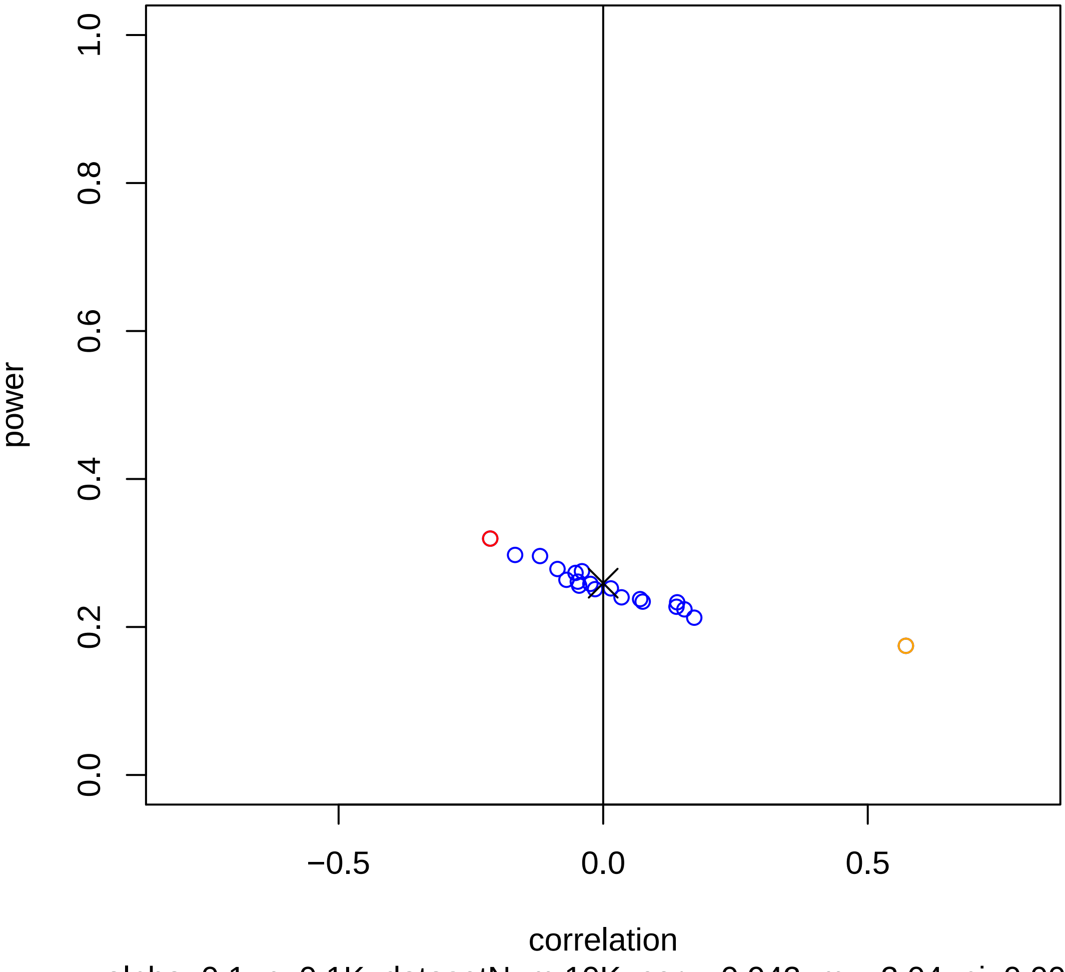


alpha: 0.1, n: 0.1K, datasetNum:10K, cor=−0.942, mu: 2.04, pi: 0.0079

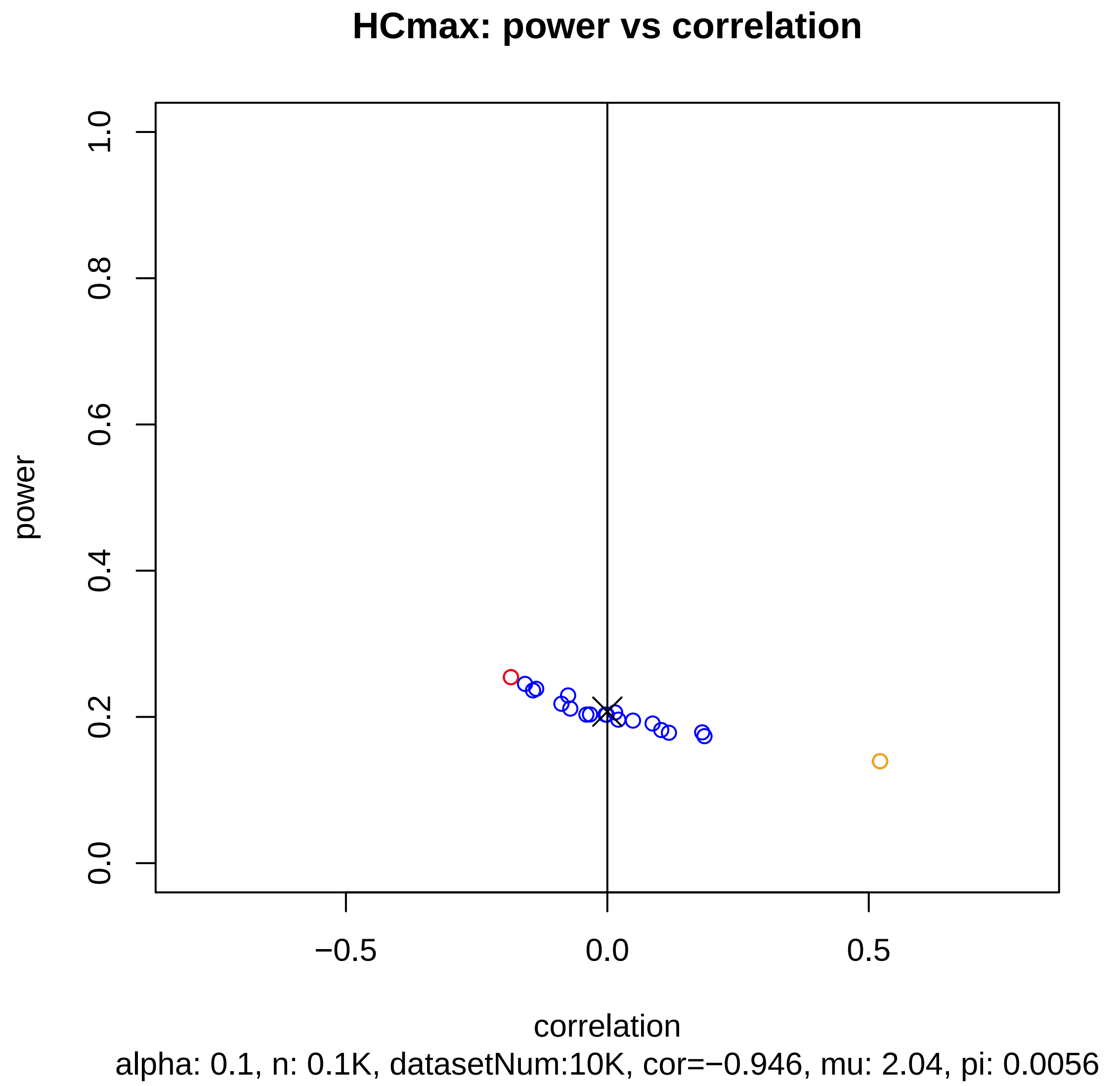

HCmax: power vs correlation
power
1.0
0.8
0.6
0.4
0.2
0.0
−0.5
0.0
0.5
correlation
alpha: 0.1, n: 0.1K, datasetNum:10K, cor=−0.946, mu: 2.04, pi: 0.0056

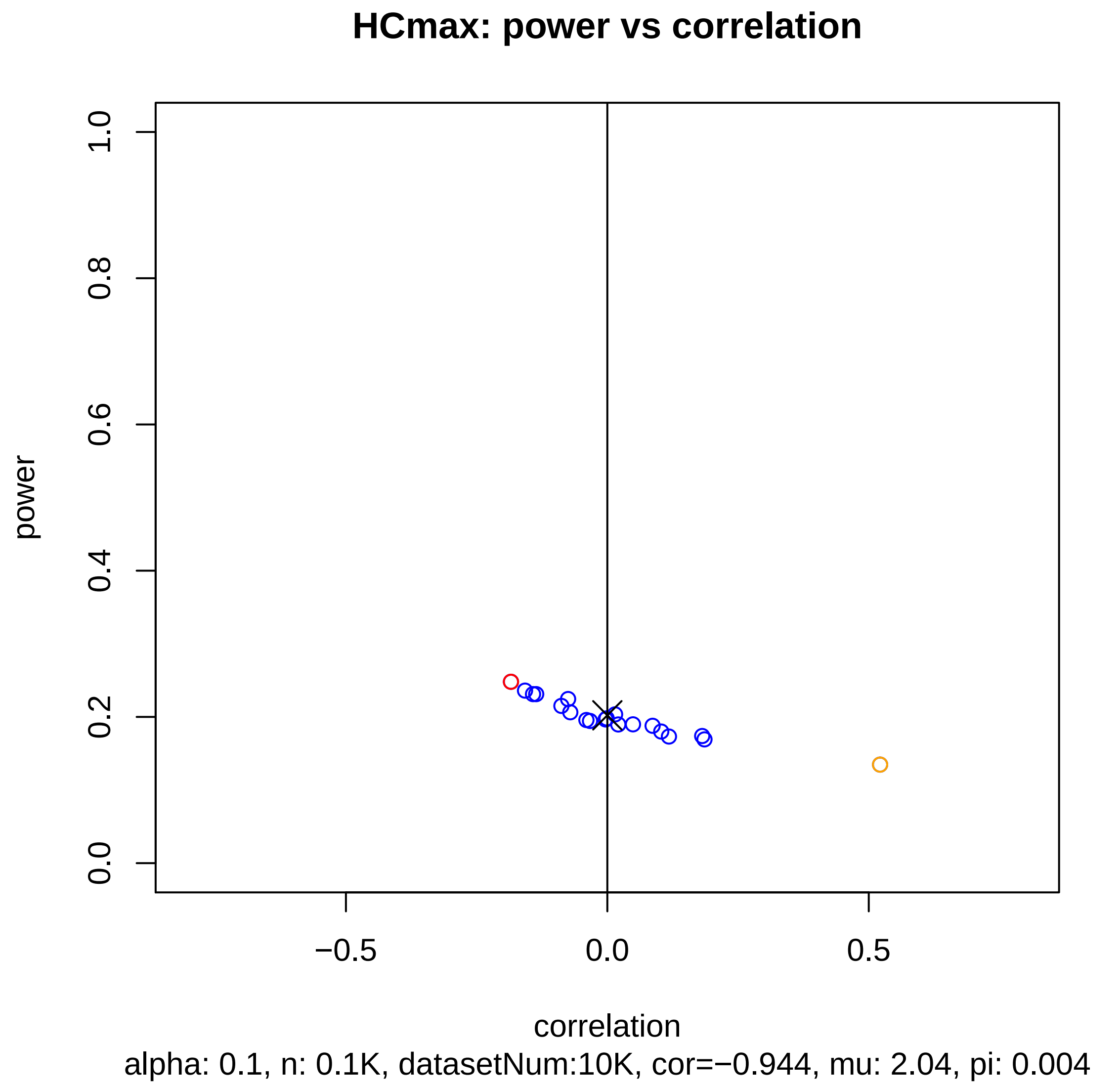
HCmax: power vs correlation
power
1.0
0.8
0.6
0.4
0.2
0.0
−0.5
0.0
0.5
correlation
alpha: 0.1, n: 0.1K, datasetNum:10K, cor=−0.944, mu: 2.04, pi: 0.004

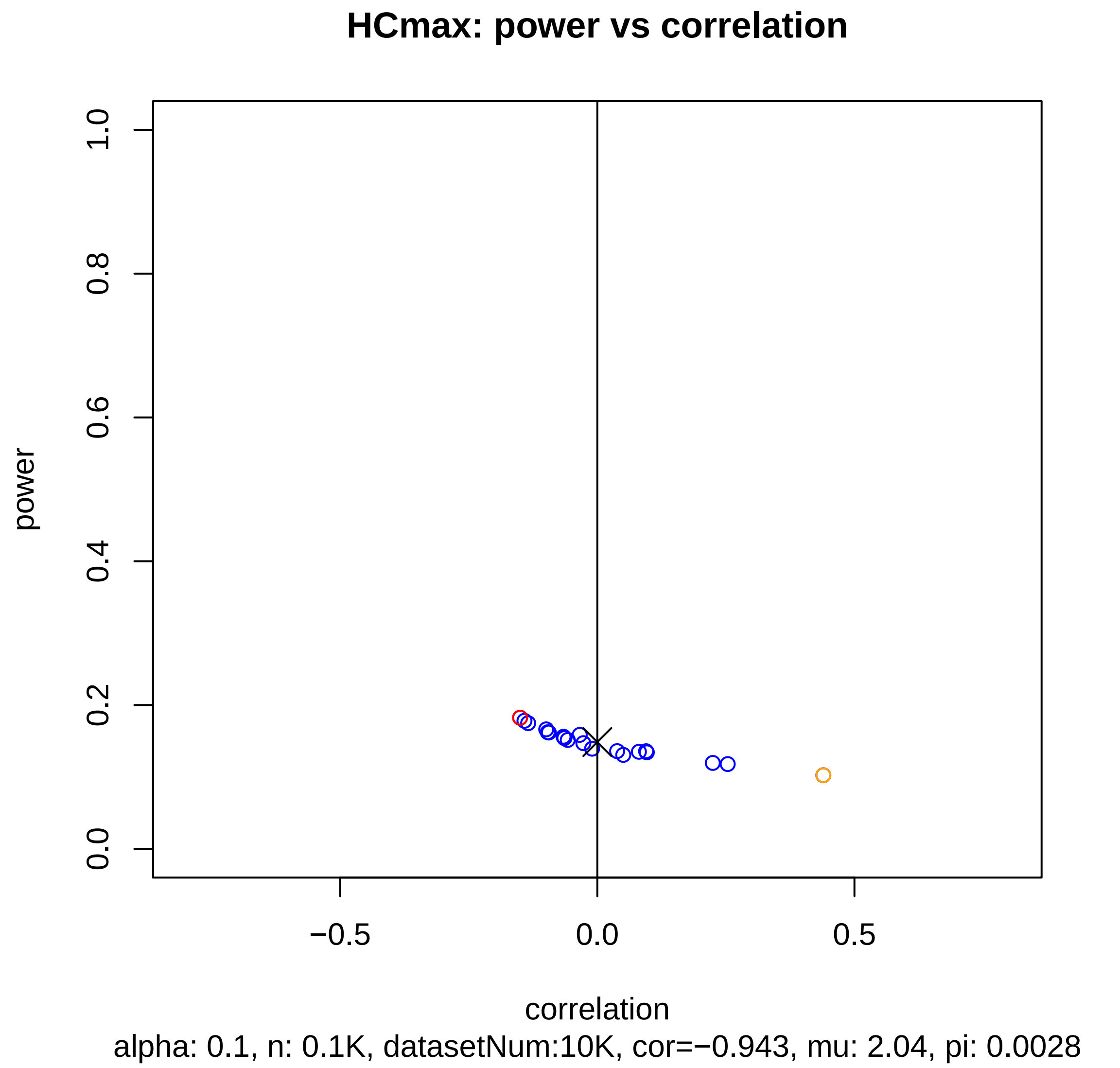
HCmax: power vs correlation
power
1.0
0.8
0.6
0.4
0.2
0.0
−0.5
0.0
0.5
correlation
alpha: 0.1, n: 0.1K, datasetNum:10K, cor=−0.943, mu: 2.04, pi: 0.0028

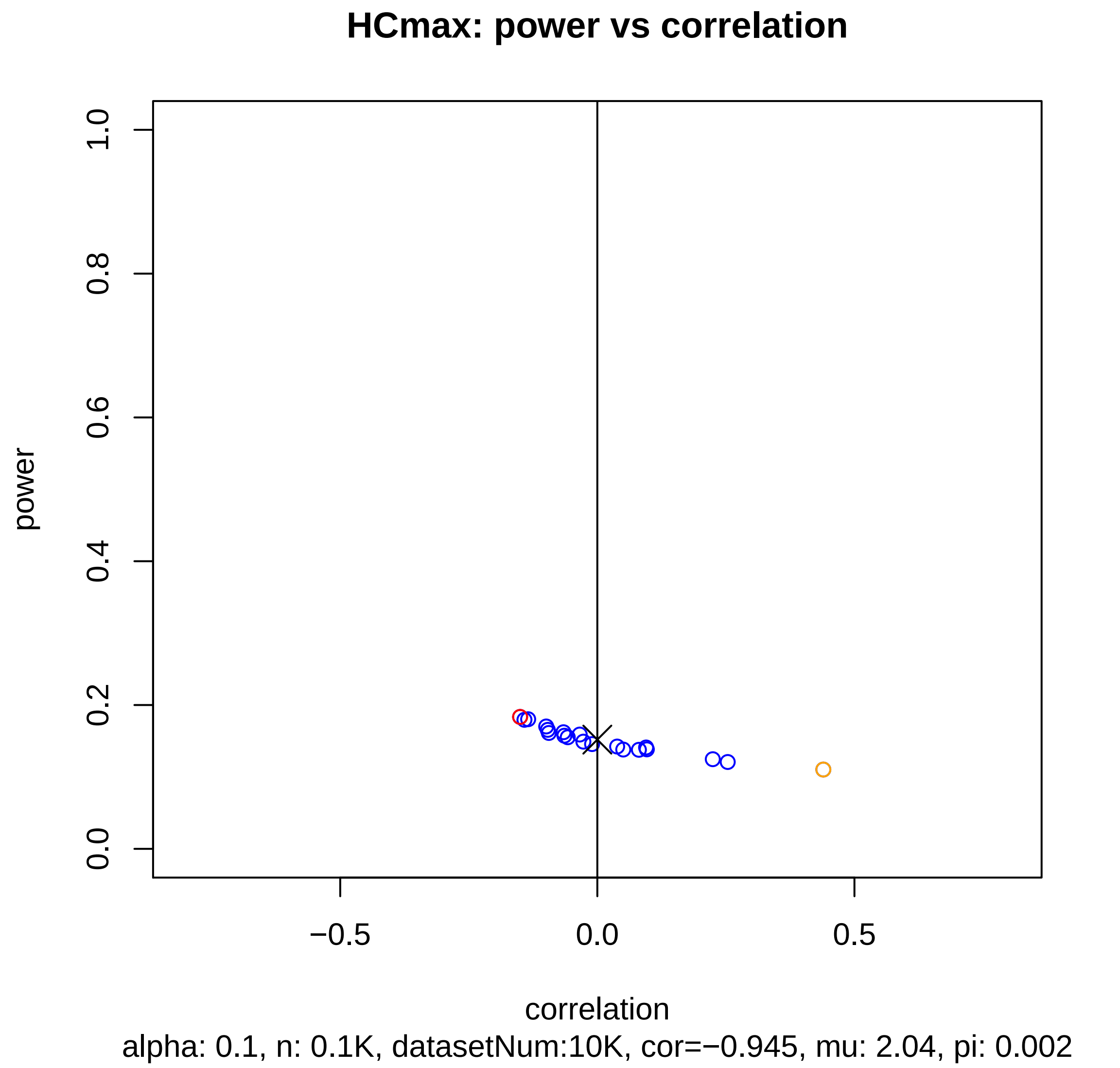
HCmax: power vs correlation
power
1.0
0.8
0.6
0.4
0.2
0.0
−0.5
0.0
0.5
correlation
alpha: 0.1, n: 0.1K, datasetNum:10K, cor=−0.945, mu: 2.04, pi: 0.002

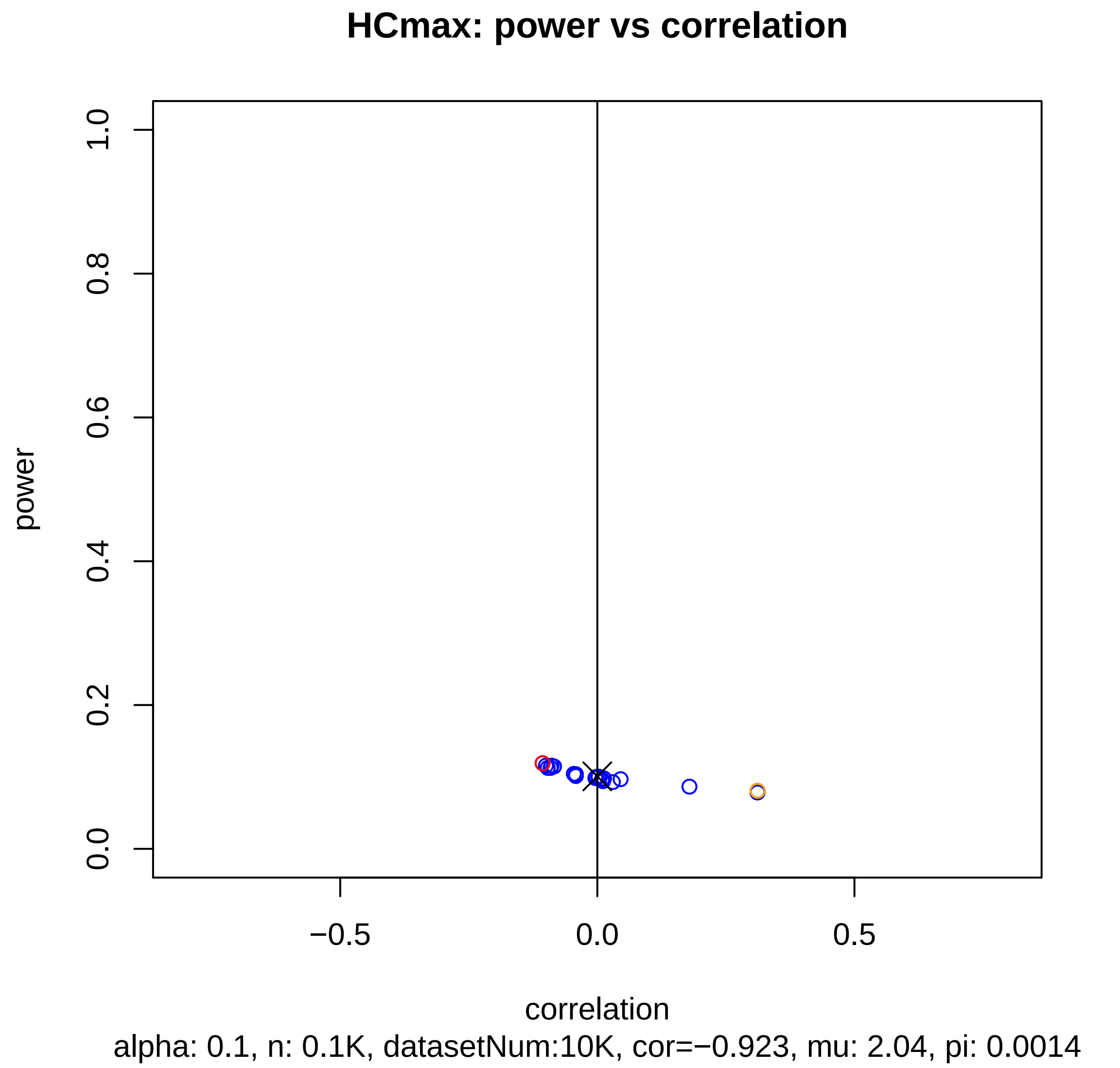
HCmax: power vs correlation
power
0.0
0.2
0.4
0.6
0.8
1.0
−0.5
0.0
0.5
correlation
alpha: 0.1, n: 0.1K, datasetNum:10K, cor=−0.923, mu: 2.04, pi: 0.0014

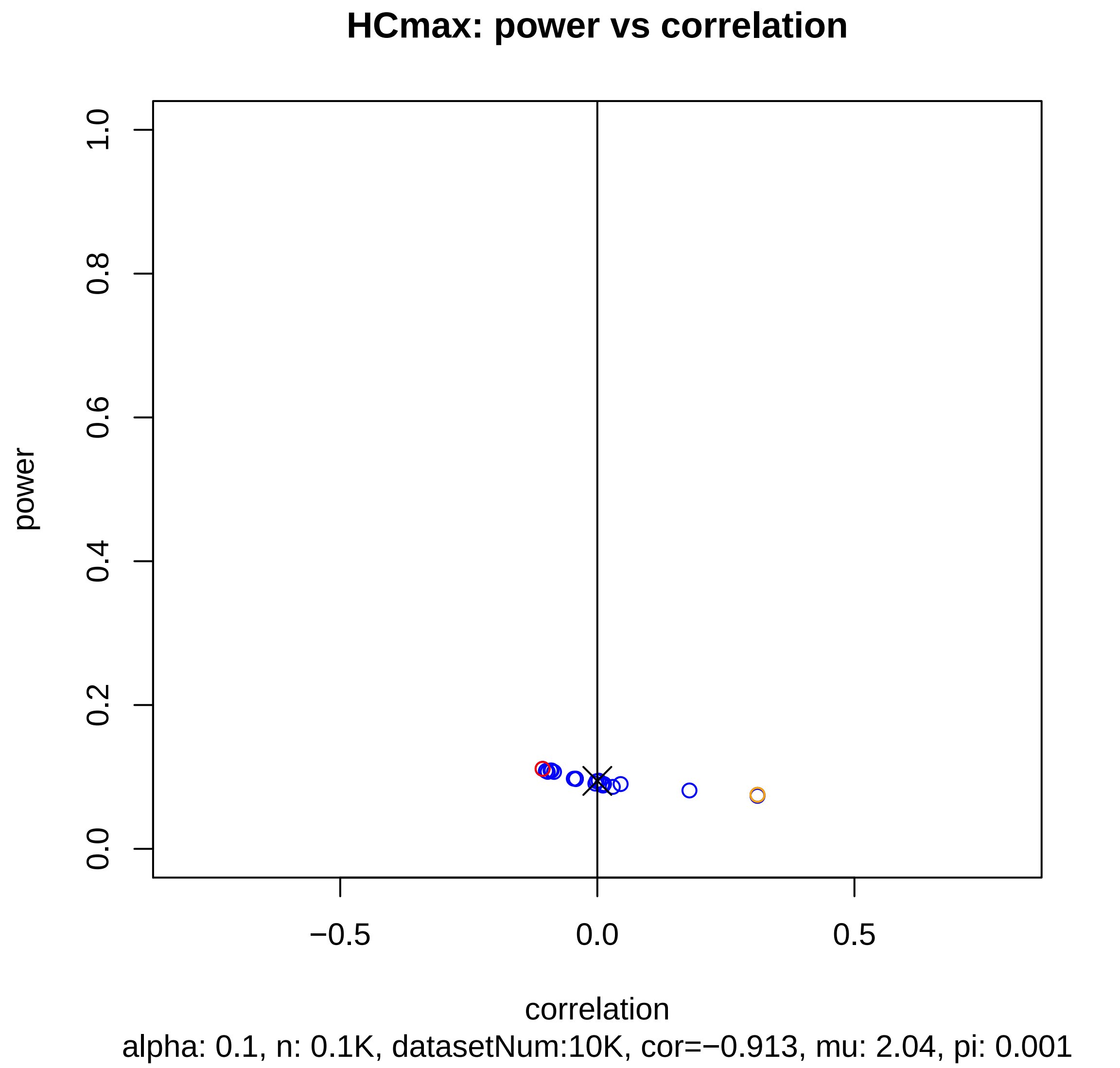
HCmax: power vs correlation
power
1.0
0.8
0.6
0.4
0.2
0.0
−0.5
0.0
0.5
correlation
alpha: 0.1, n: 0.1K, datasetNum:10K, cor=−0.913, mu: 2.04, pi: 0.001

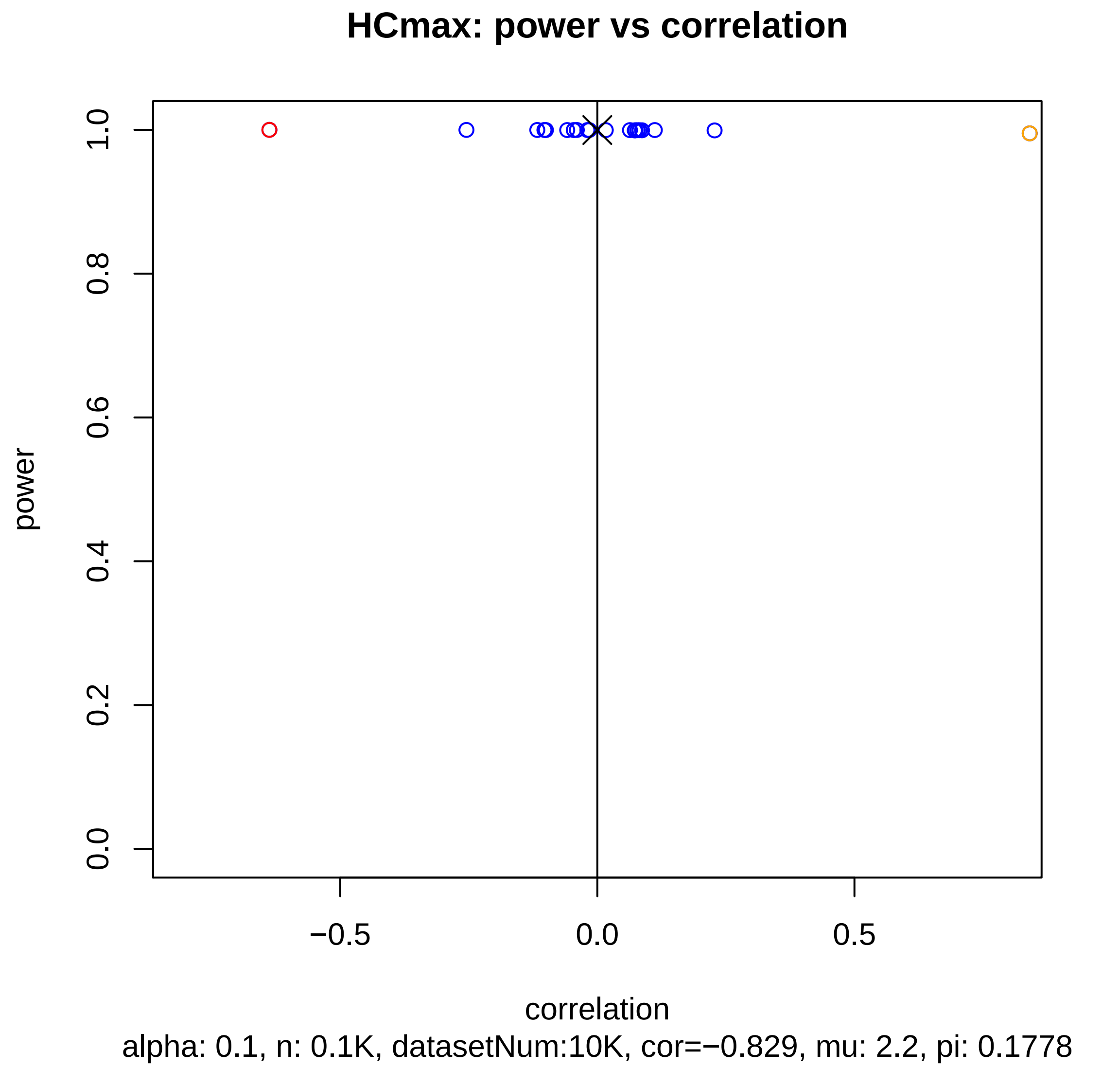
HCmax: power vs correlation
power
1.0
0.8
0.6
0.4
0.2
0.0
−0.5
0.0
0.5
correlation
alpha: 0.1, n: 0.1K, datasetNum:10K, cor=−0.829, mu: 2.2, pi: 0.1778

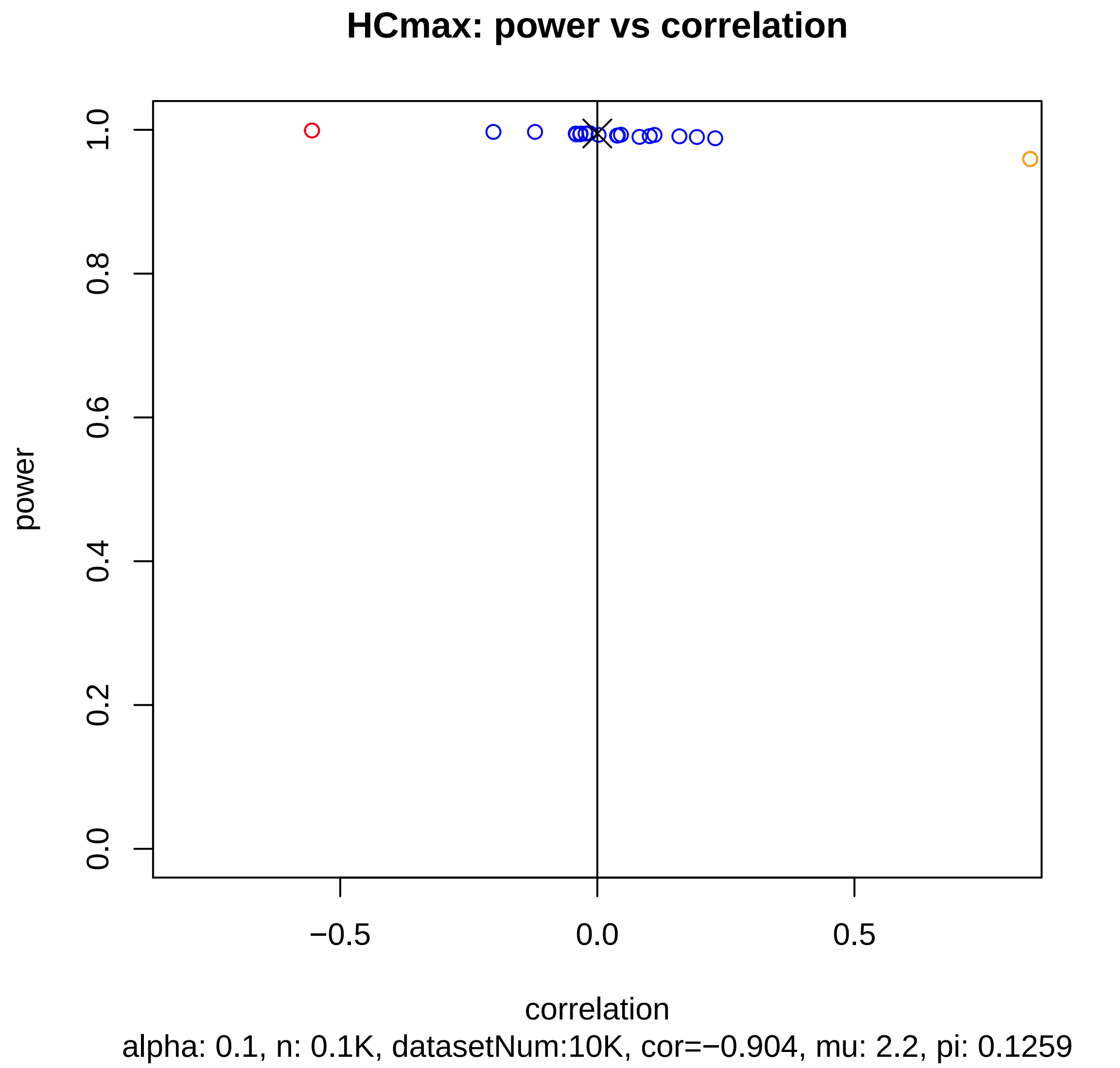
HCmax: power vs correlation
power
1.0
0.8
0.6
0.4
0.2
0.0
−0.5
0.0
0.5
correlation
alpha: 0.1, n: 0.1K, datasetNum:10K, cor=−0.904, mu: 2.2, pi: 0.1259

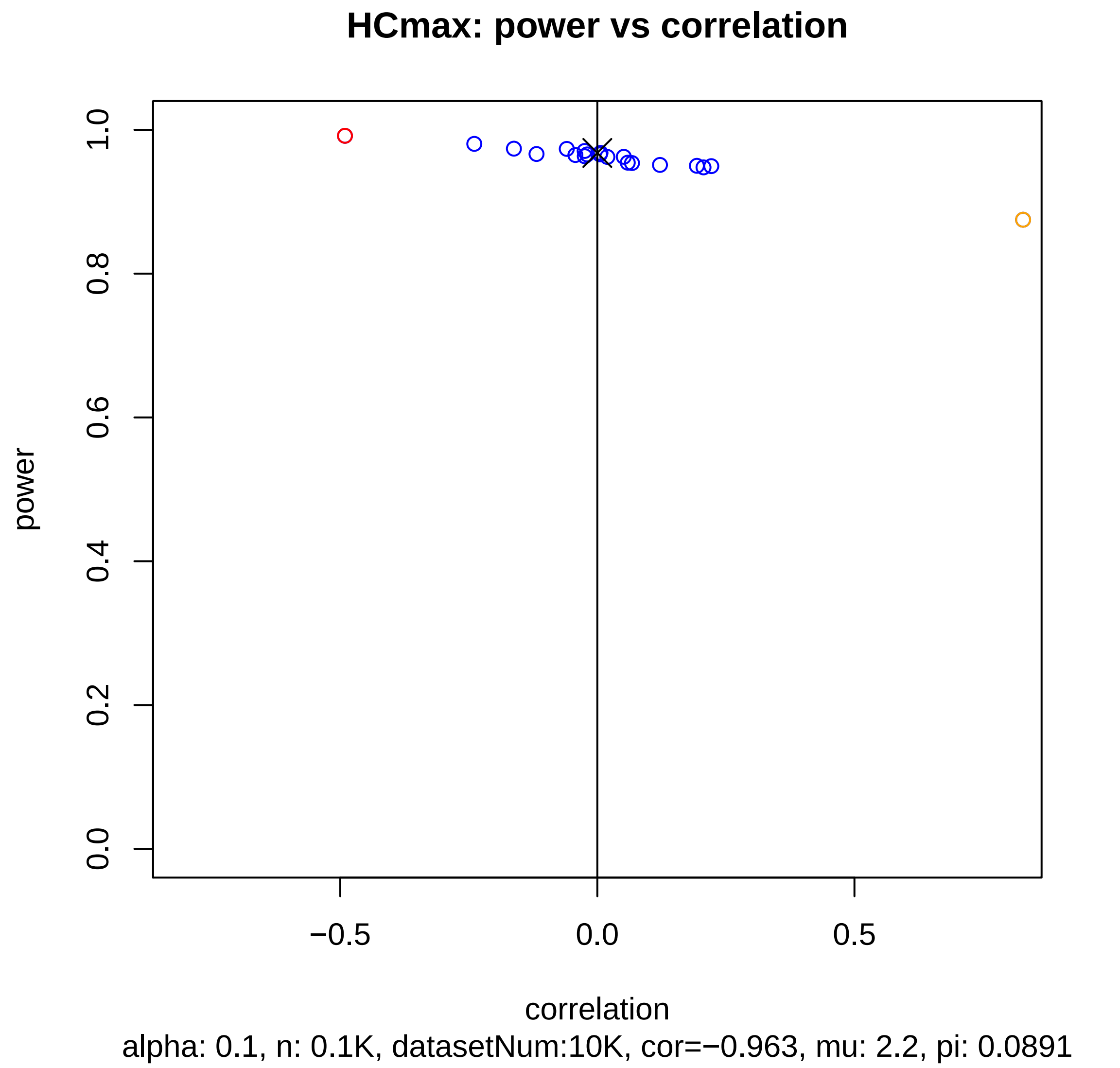

HCmax: power vs correlation
power
0.0
0.2
0.4
0.6
0.8
1.0
−0.5
0.0
0.5
correlation
alpha: 0.1, n: 0.1K, datasetNum:10K, cor=−0.963, mu: 2.2, pi: 0.0891

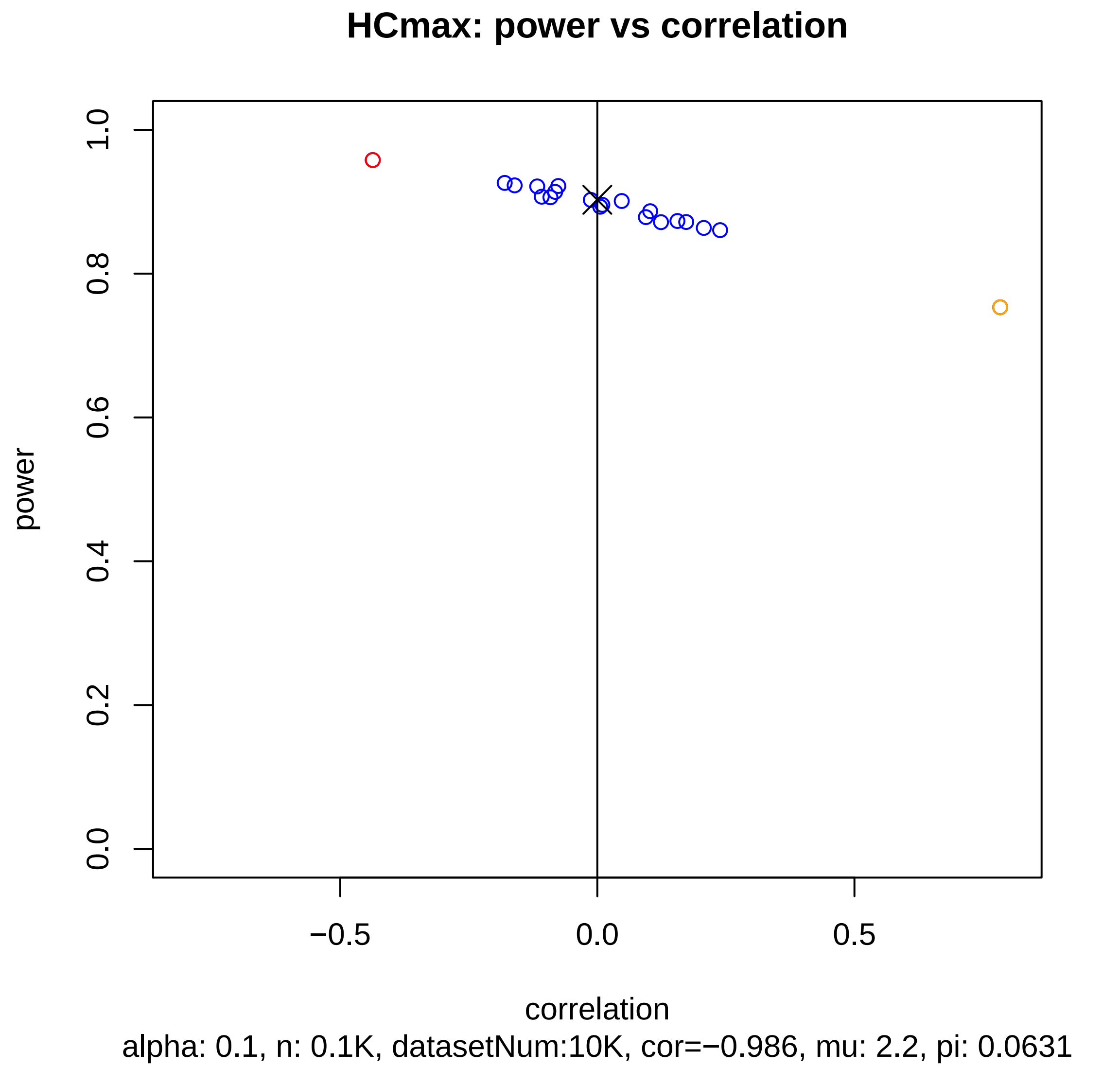
HCmax: power vs correlation
power
1.0
0.8
0.6
0.4
0.2
0.0
−0.5
0.0
0.5
correlation
alpha: 0.1, n: 0.1K, datasetNum:10K, cor=−0.986, mu: 2.2, pi: 0.0631

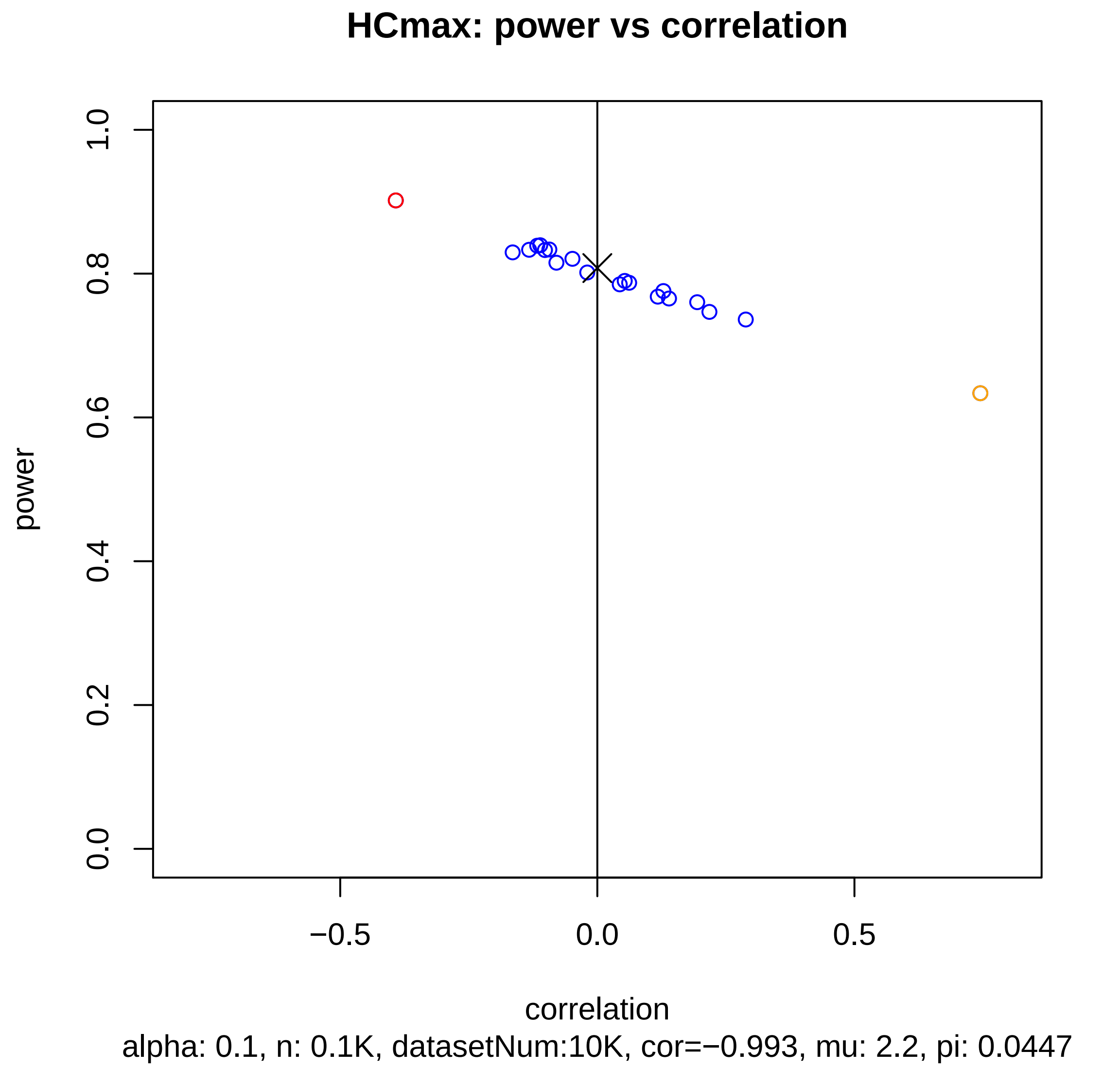

HCmax: power vs correlation
power
1.0
0.8
0.6
0.4
0.2
0.0
−0.5
0.0
0.5
correlation
alpha: 0.1, n: 0.1K, datasetNum:10K, cor=−0.993, mu: 2.2, pi: 0.0447

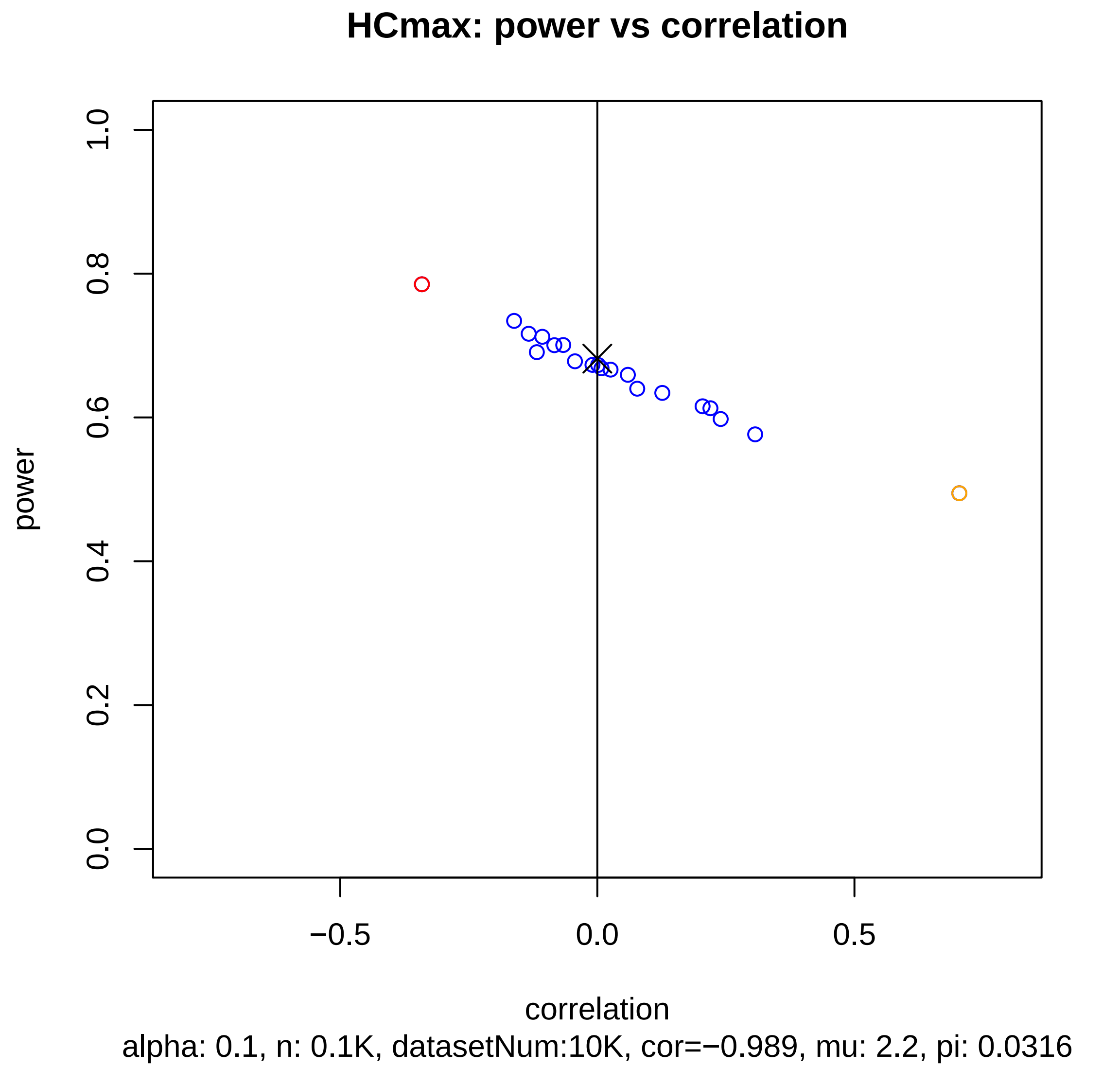
HCmax: power vs correlation
power
0.0
0.2
0.4
0.6
0.8
1.0
−0.5
0.0
0.5
correlation
alpha: 0.1, n: 0.1K, datasetNum:10K, cor=−0.989, mu: 2.2, pi: 0.0316

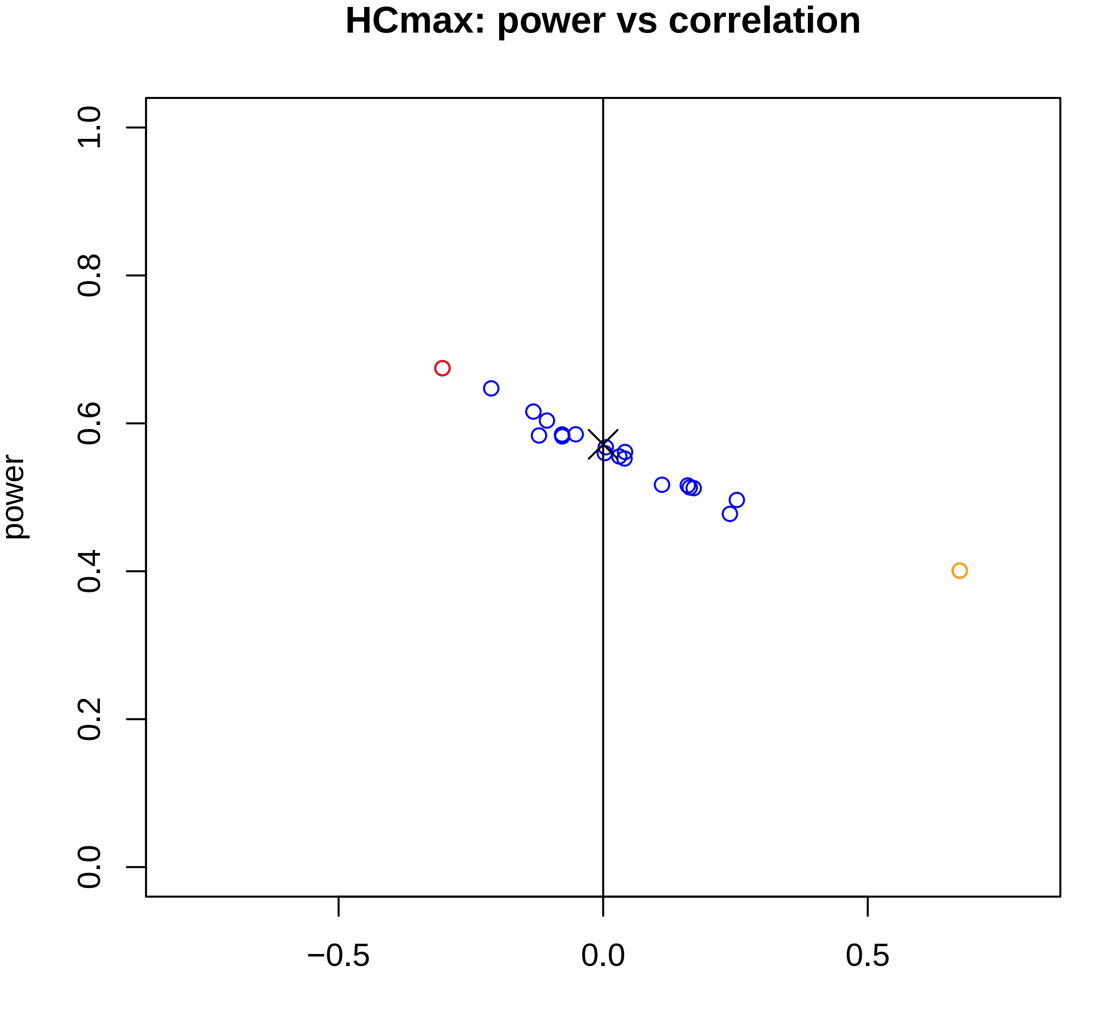

alpha: 0.1, n: 0.1K, datasetNum:10K, cor=−0.979, mu: 2.2, pi: 0.0224

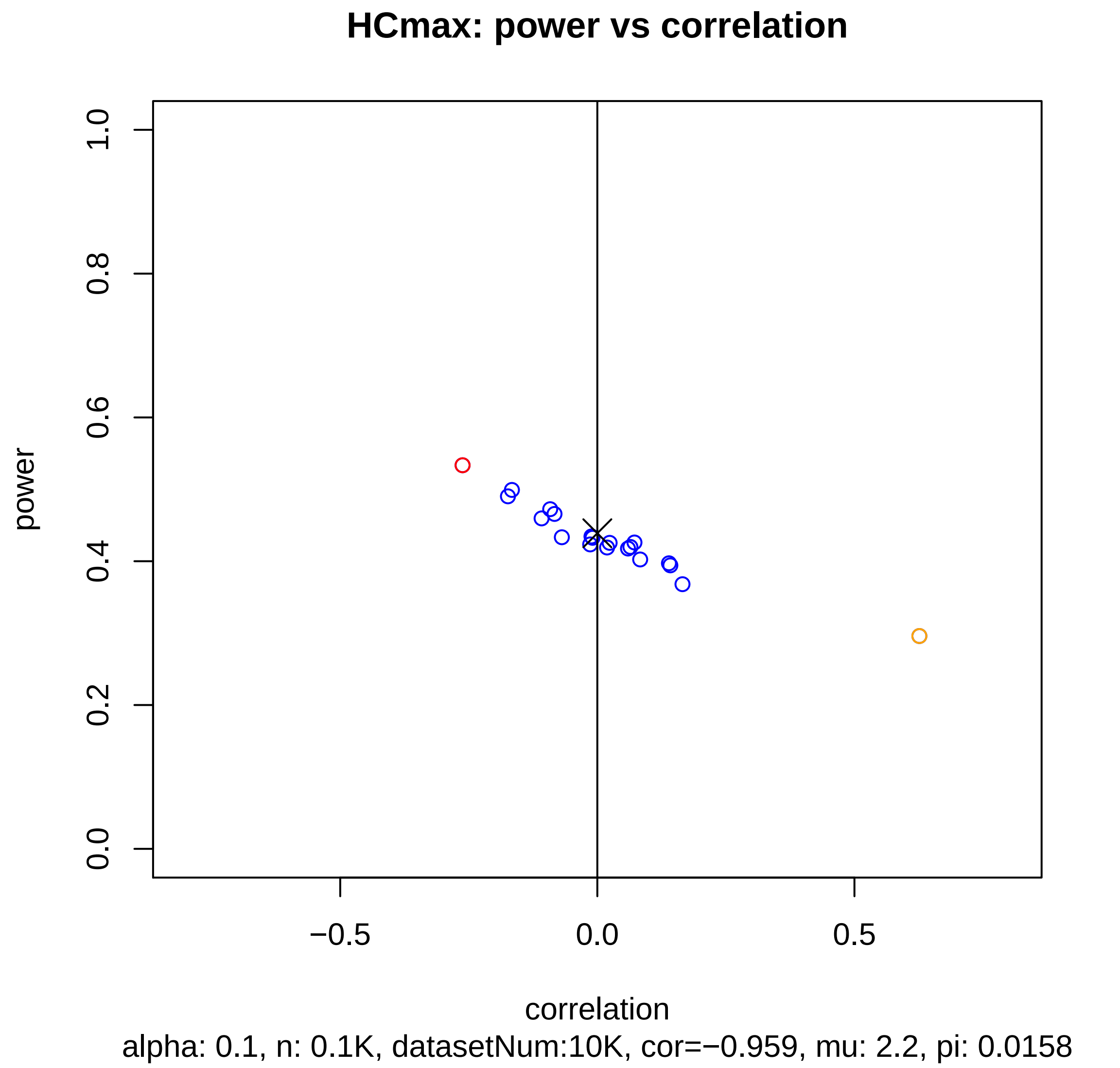
HCmax: power vs correlation
power
0.0
0.2
0.4
0.6
0.8
1.0
−0.5
0.0
0.5
correlation
alpha: 0.1, n: 0.1K, datasetNum:10K, cor=−0.959, mu: 2.2, pi: 0.0158

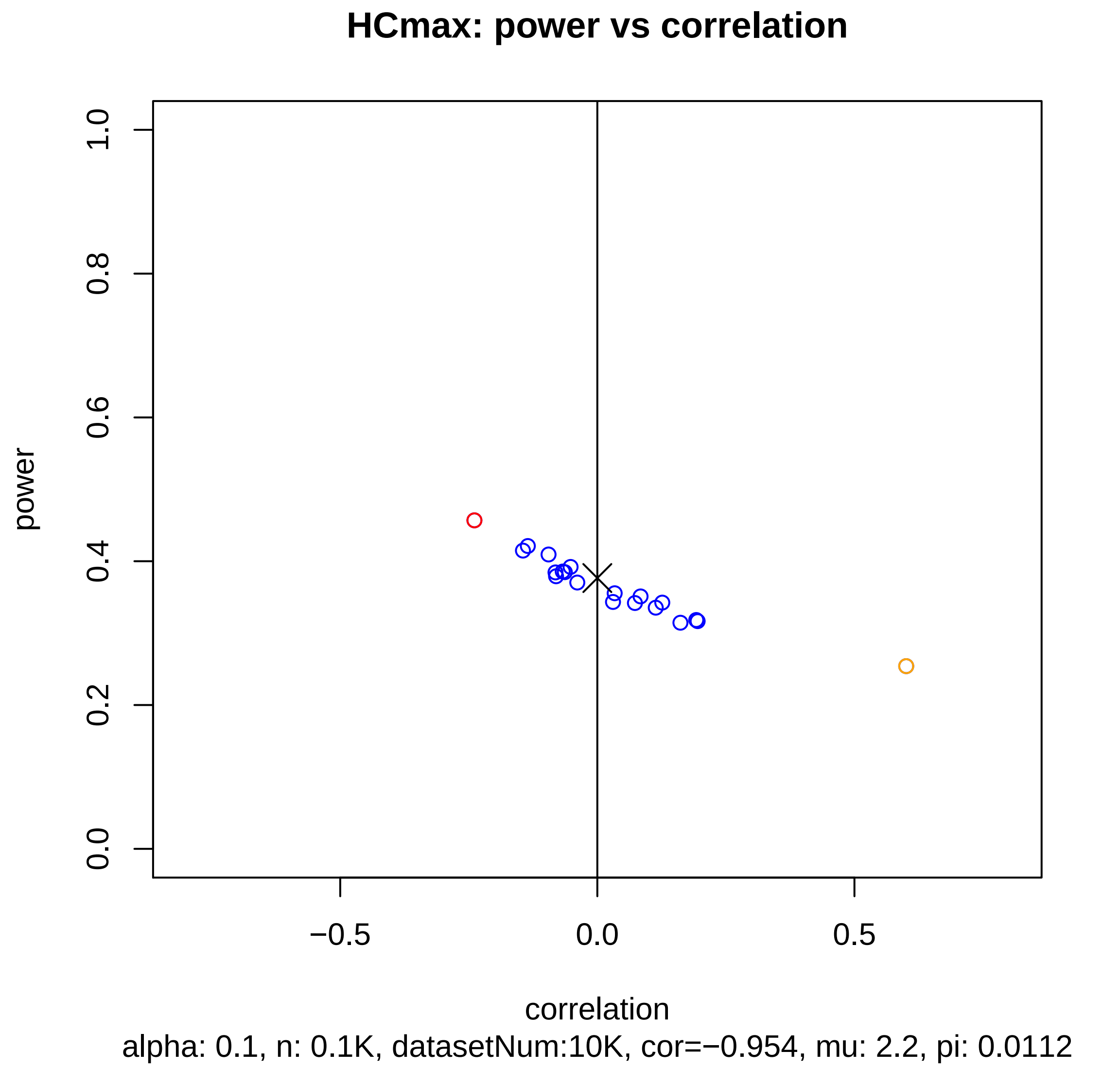

HCmax: power vs correlation
power
1.0
0.8
0.6
0.4
0.2
0.0
−0.5
0.0
0.5
correlation
alpha: 0.1, n: 0.1K, datasetNum:10K, cor=−0.954, mu: 2.2, pi: 0.0112

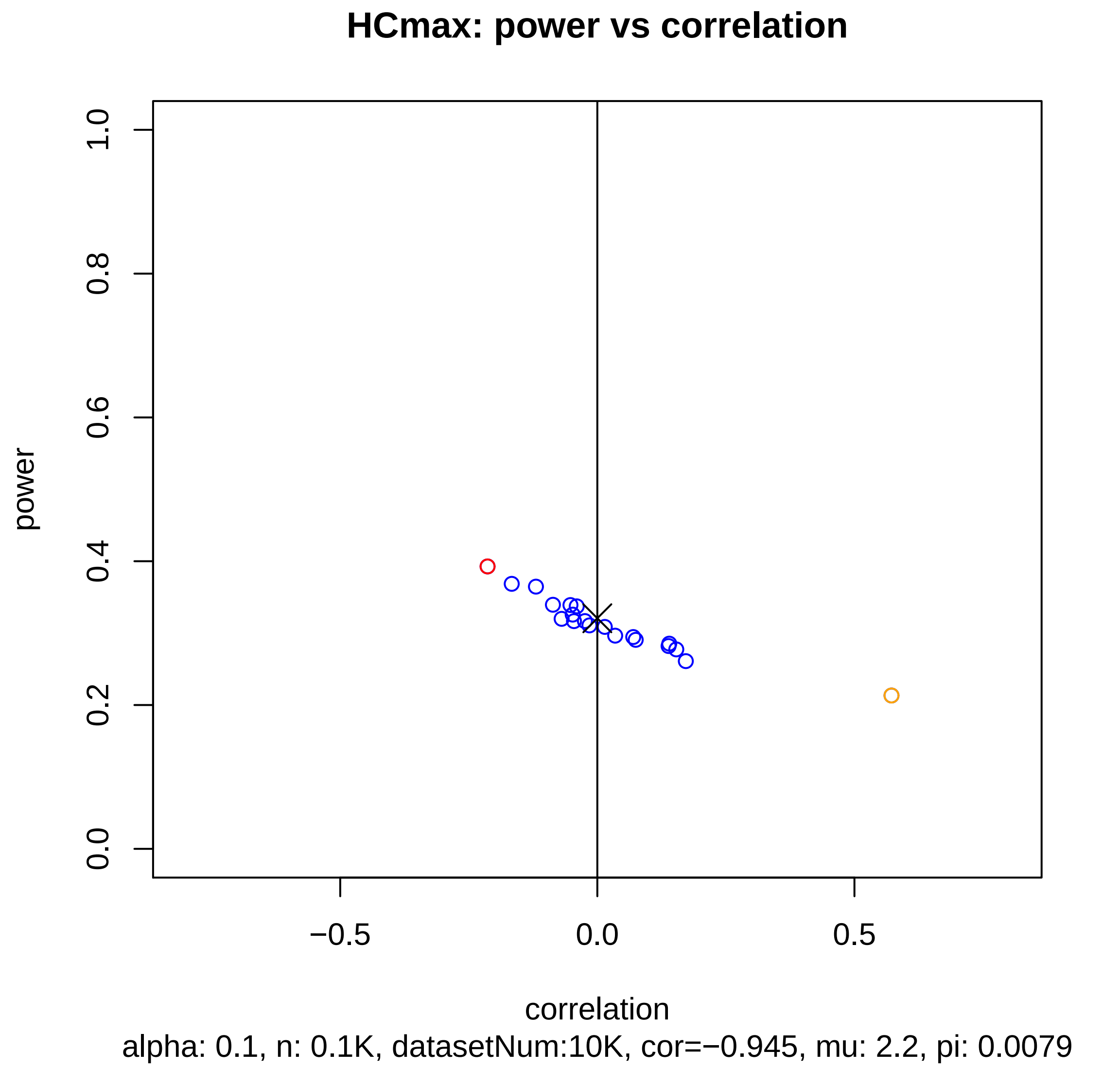
HCmax: power vs correlation
power
0.0
0.2
0.4
0.6
0.8
1.0
−0.5
0.0
0.5
correlation
alpha: 0.1, n: 0.1K, datasetNum:10K, cor=−0.945, mu: 2.2, pi: 0.0079

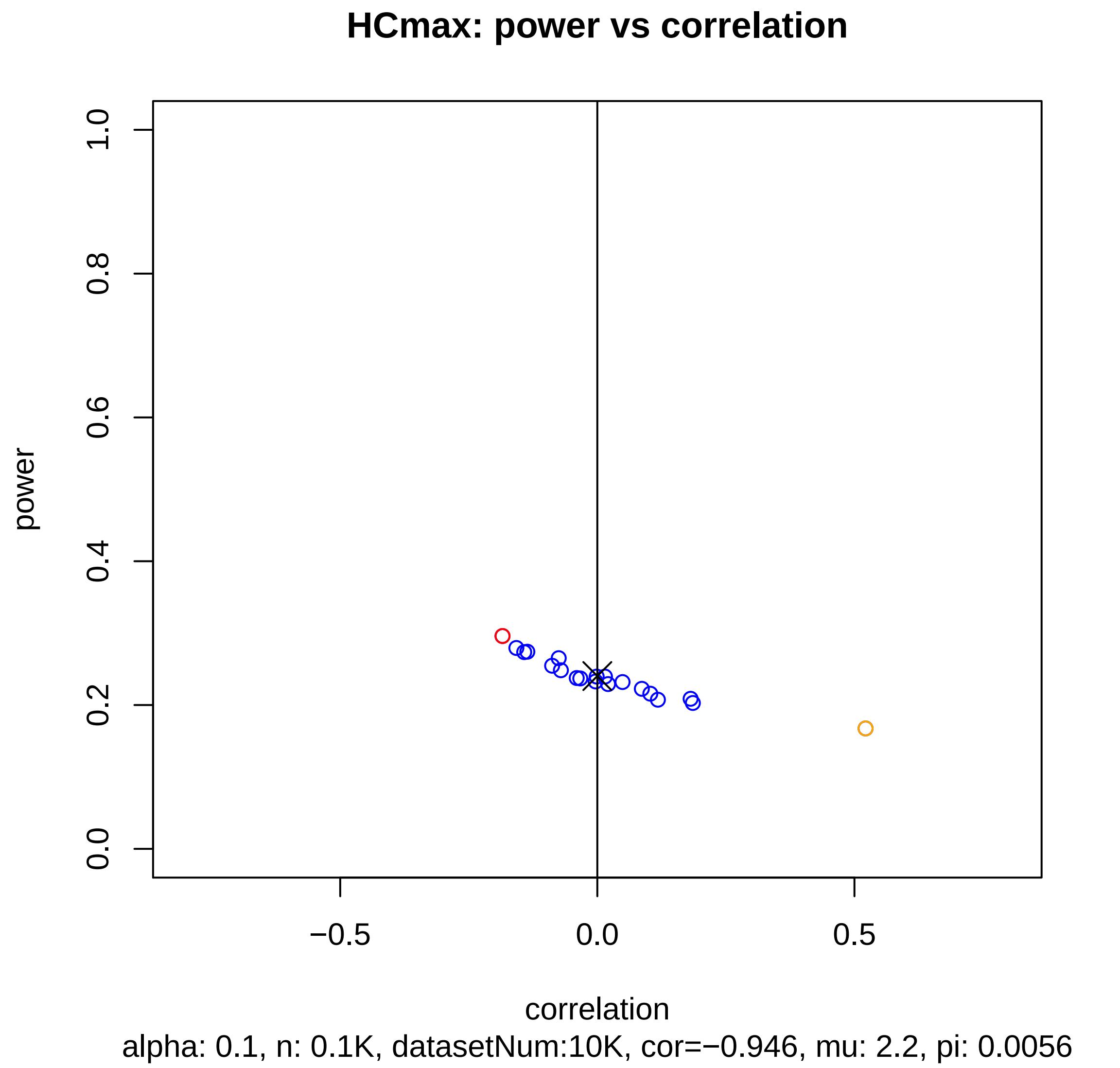
HCmax: power vs correlation
power
1.0
0.8
0.6
0.4
0.2
0.0
−0.5
0.0
0.5
correlation
alpha: 0.1, n: 0.1K, datasetNum:10K, cor=−0.946, mu: 2.2, pi: 0.0056

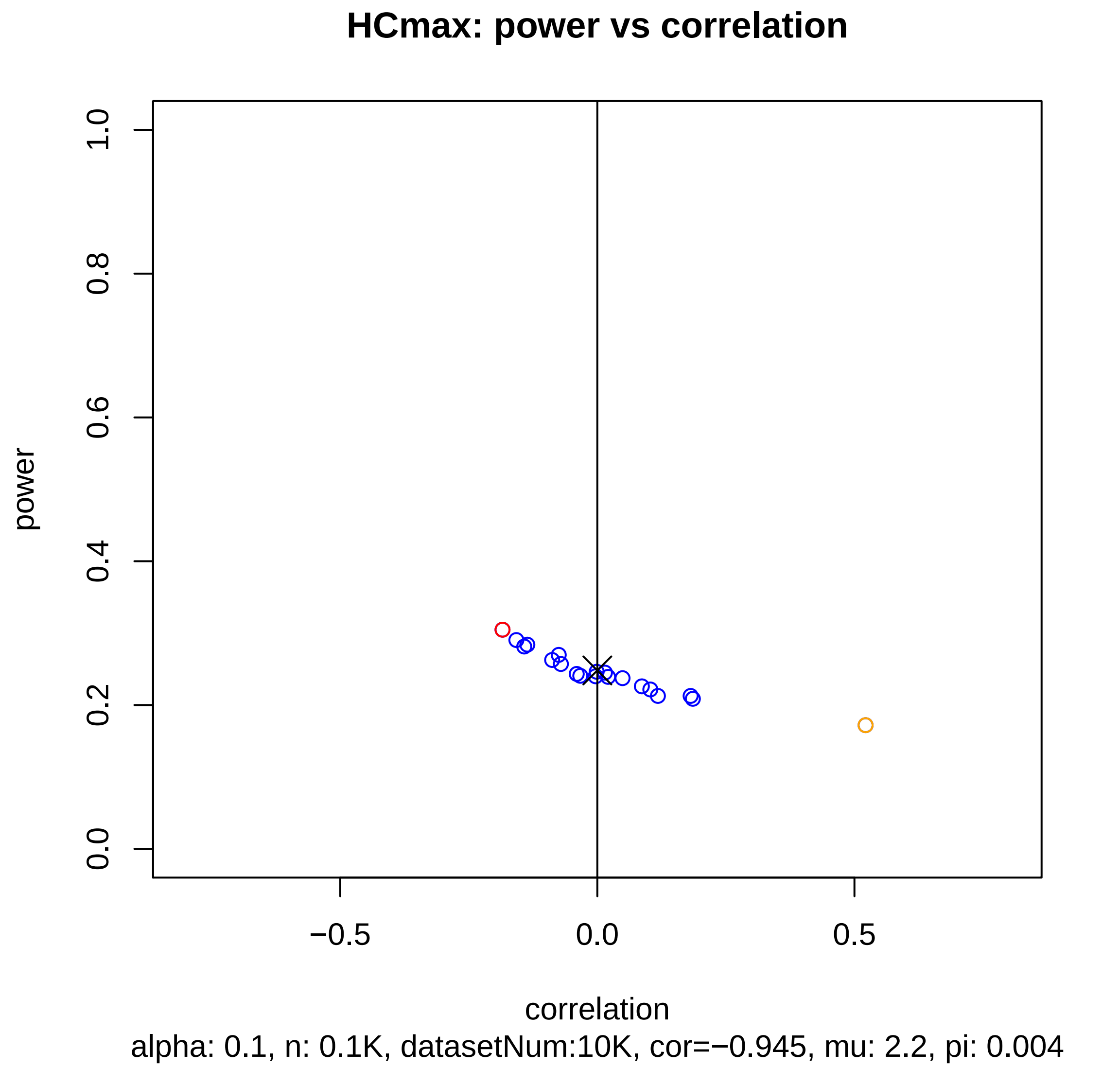

HCmax: power vs correlation
power
0.0
0.2
0.4
0.6
0.8
1.0
−0.5
0.0
0.5
correlation
alpha: 0.1, n: 0.1K, datasetNum:10K, cor=−0.945, mu: 2.2, pi: 0.004

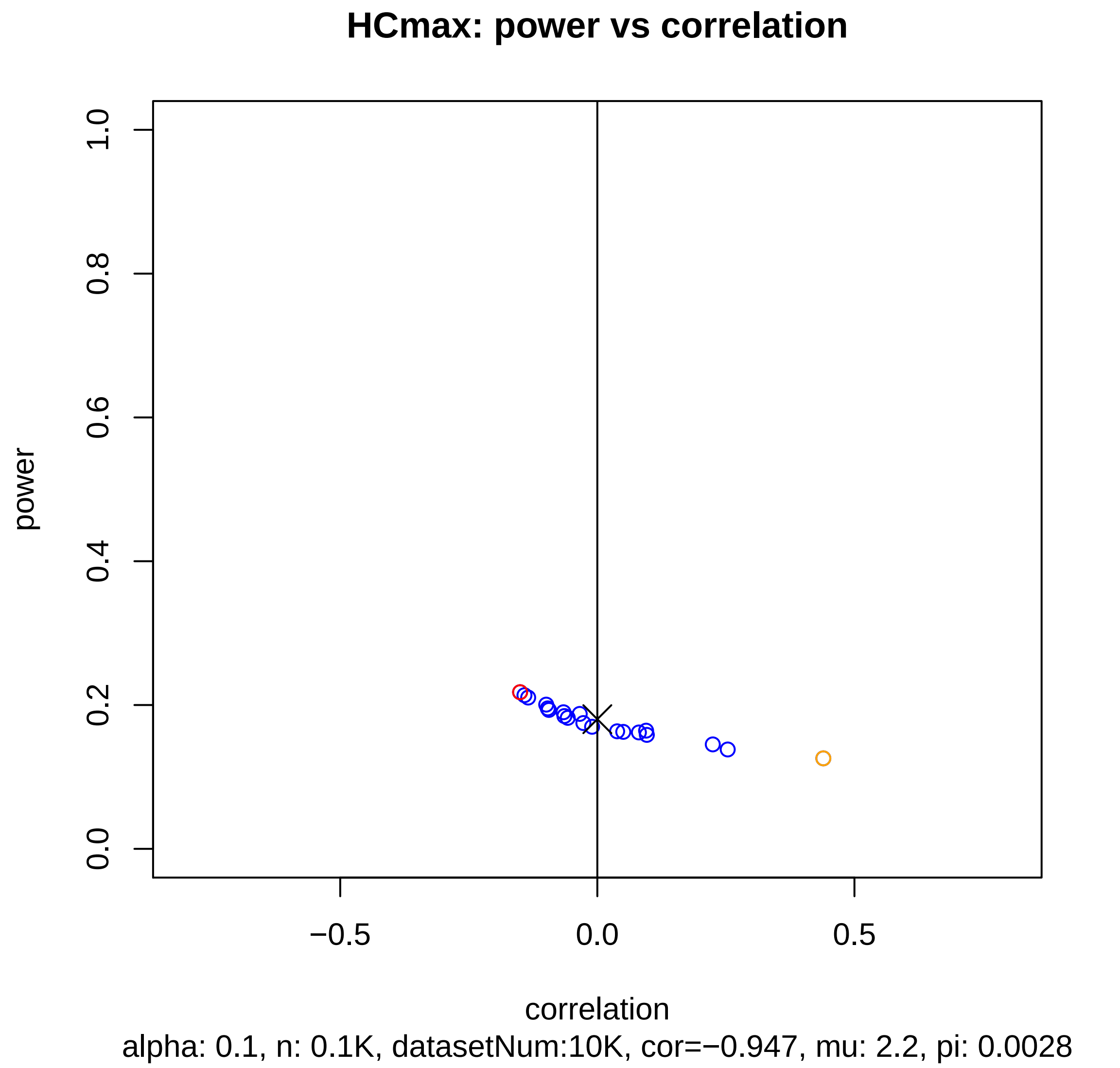
HCmax: power vs correlation
power
1.0
0.8
0.6
0.4
0.2
0.0
−0.5
0.0
0.5
correlation
alpha: 0.1, n: 0.1K, datasetNum:10K, cor=−0.947, mu: 2.2, pi: 0.0028

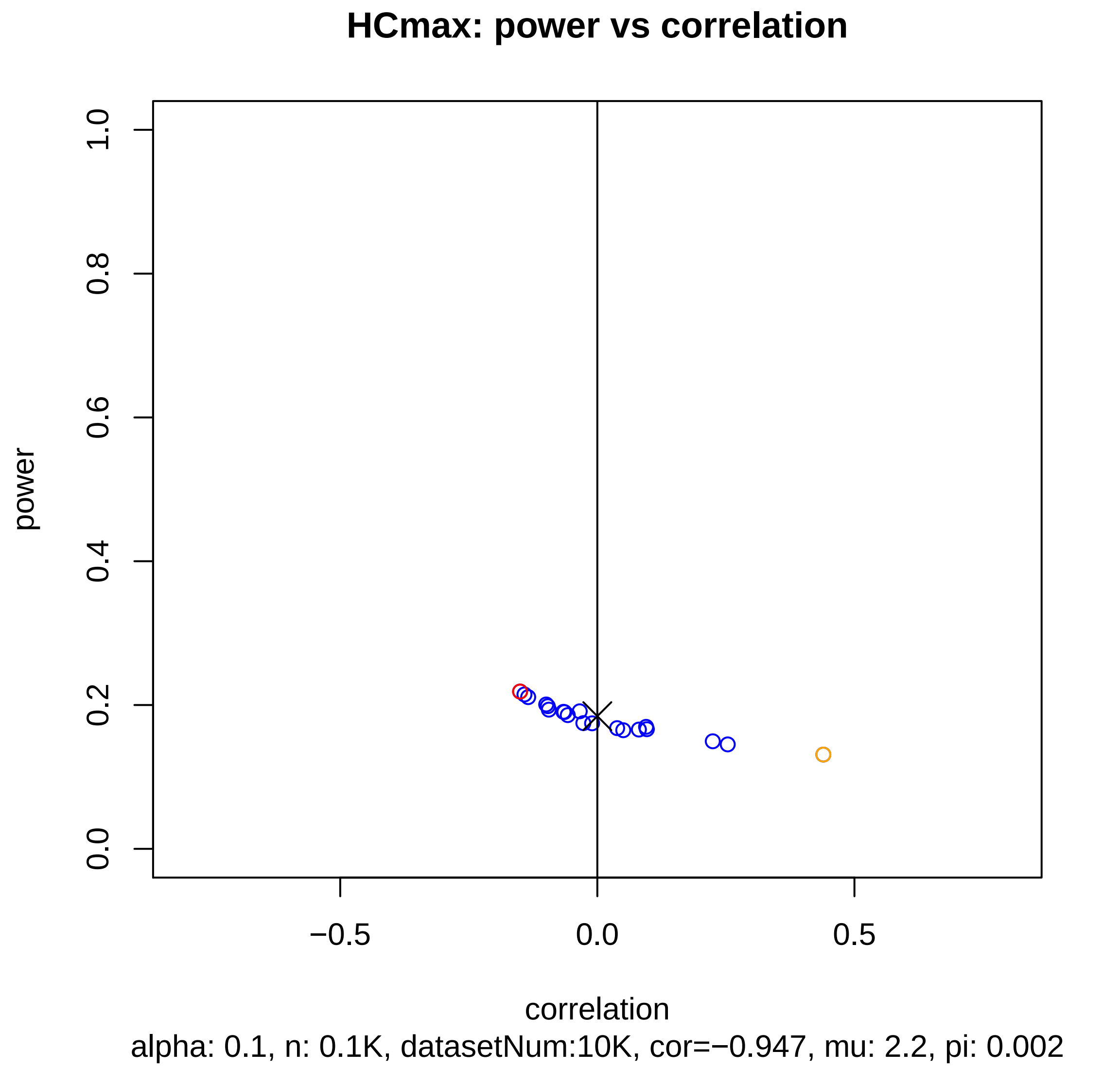
HCmax: power vs correlation
power
0.0
0.2
0.4
0.6
0.8
1.0
−0.5
0.0
0.5
correlation
alpha: 0.1, n: 0.1K, datasetNum:10K, cor=−0.947, mu: 2.2, pi: 0.002

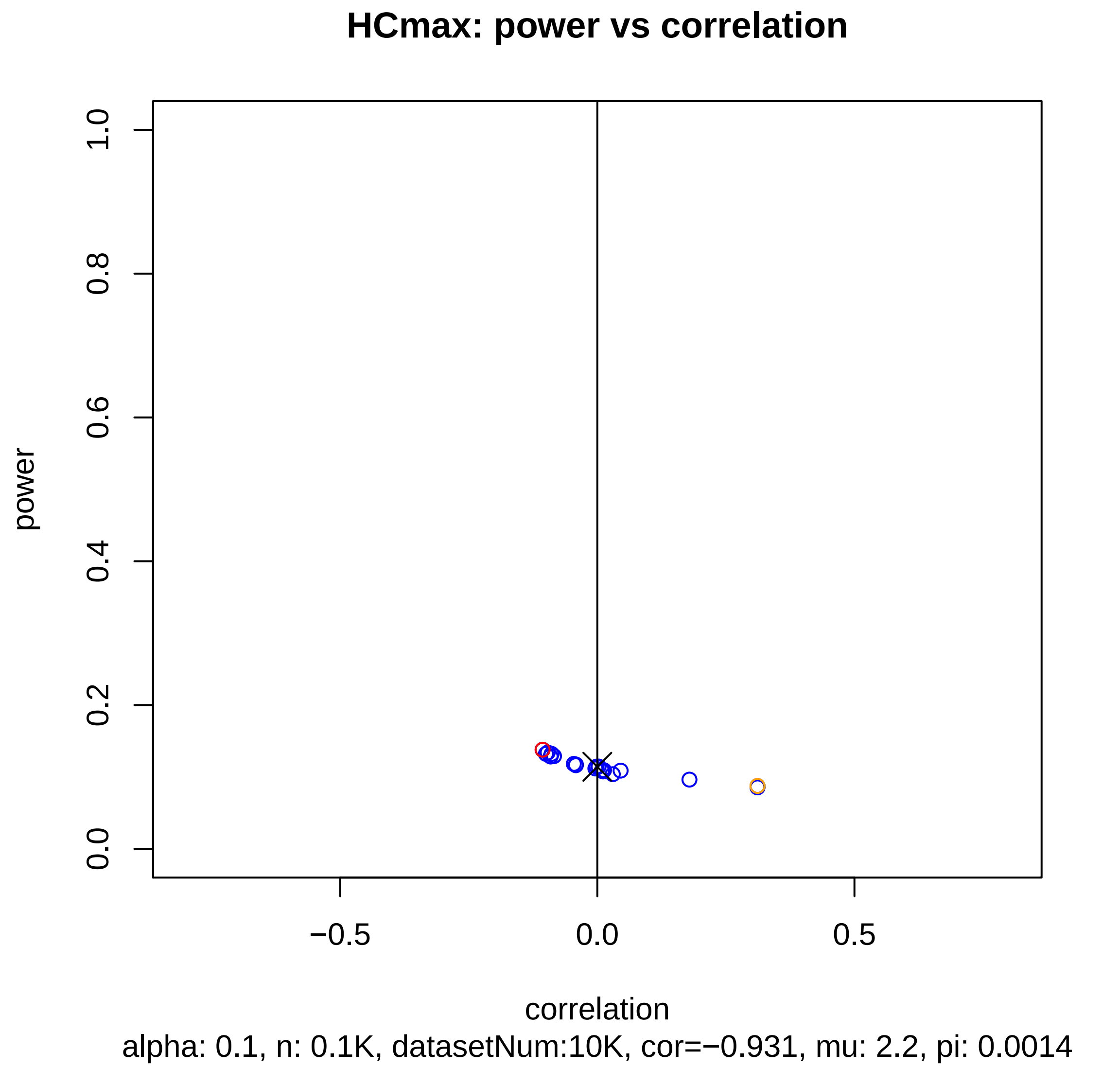
HCmax: power vs correlation
power
1.0
0.8
0.6
0.4
0.2
0.0
−0.5
0.0
0.5
correlation
alpha: 0.1, n: 0.1K, datasetNum:10K, cor=−0.931, mu: 2.2, pi: 0.0014

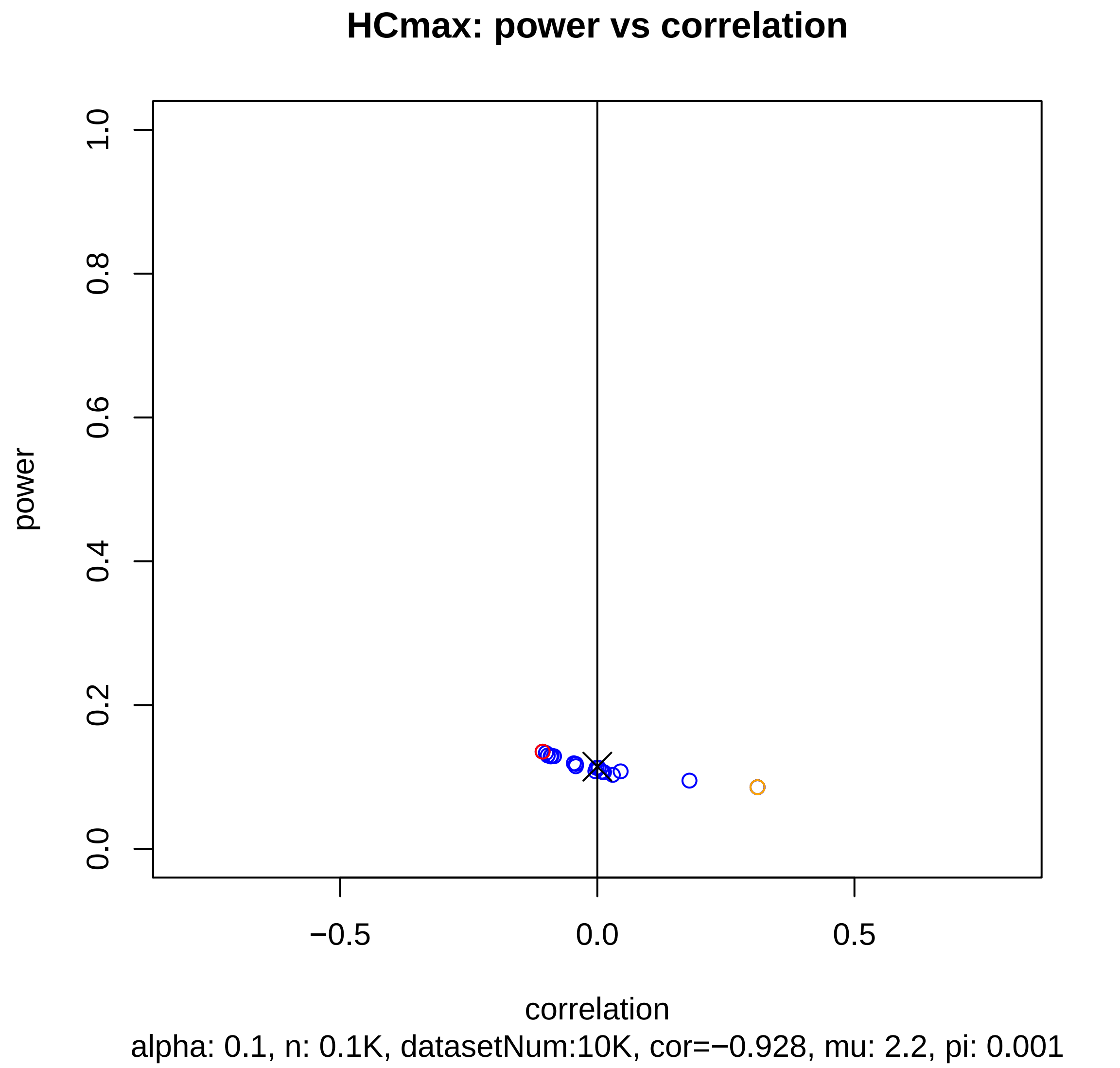
HCmax: power vs correlation
power
1.0
0.8
0.6
0.4
0.2
0.0
−0.5
0.0
0.5
correlation
alpha: 0.1, n: 0.1K, datasetNum:10K, cor=−0.928, mu: 2.2, pi: 0.001

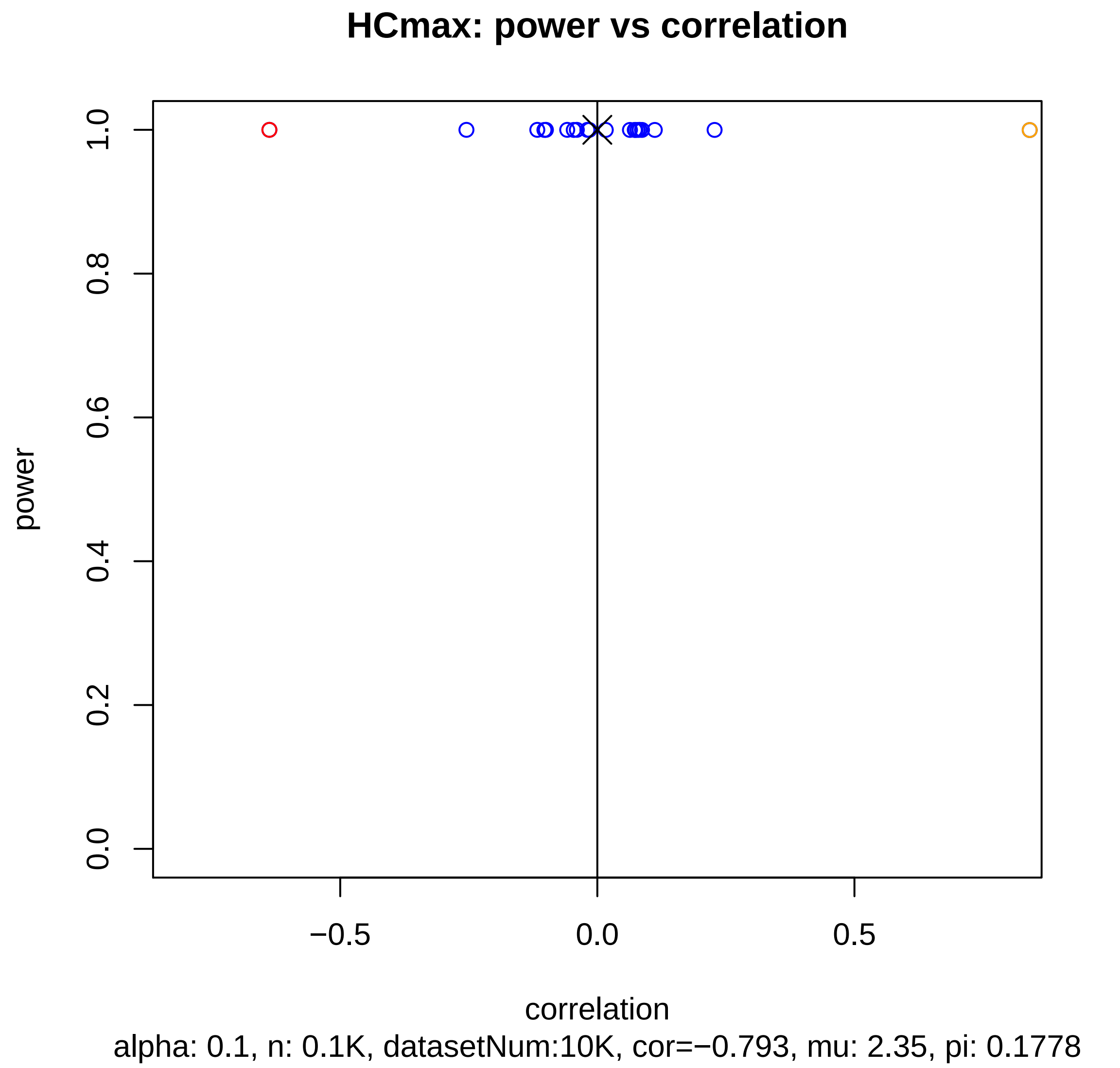
HCmax: power vs correlation
power
0.0
0.2
0.4
0.6
0.8
1.0
−0.5
0.0
0.5
correlation
alpha: 0.1, n: 0.1K, datasetNum:10K, cor=−0.793, mu: 2.35, pi: 0.1778

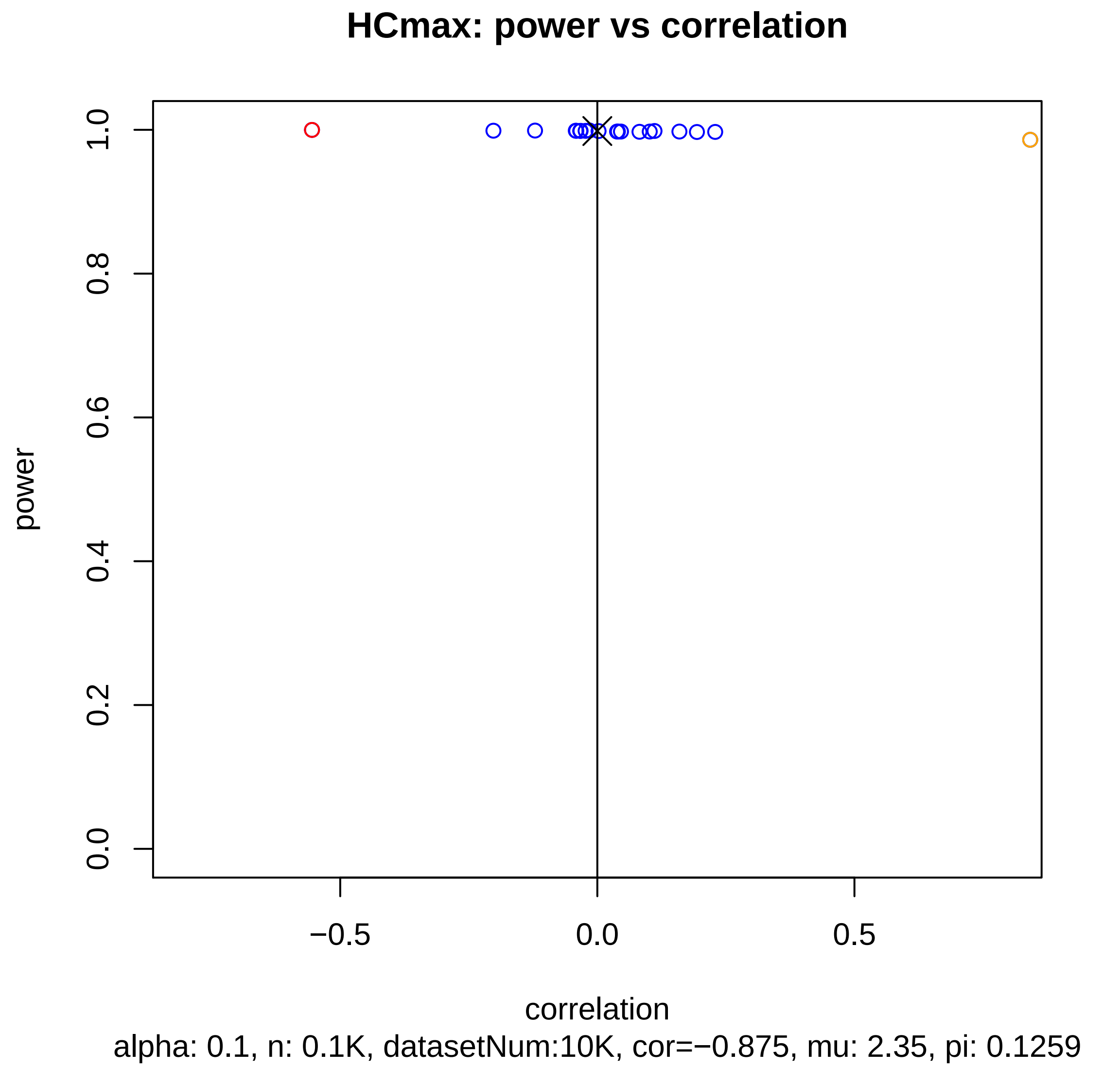

HCmax: power vs correlation
1.0
0.8
0.6
0.4
0.2
0.0
power
−0.5
0.0
0.5
correlation
alpha: 0.1, n: 0.1K, datasetNum:10K, cor=−0.875, mu: 2.35, pi: 0.1259

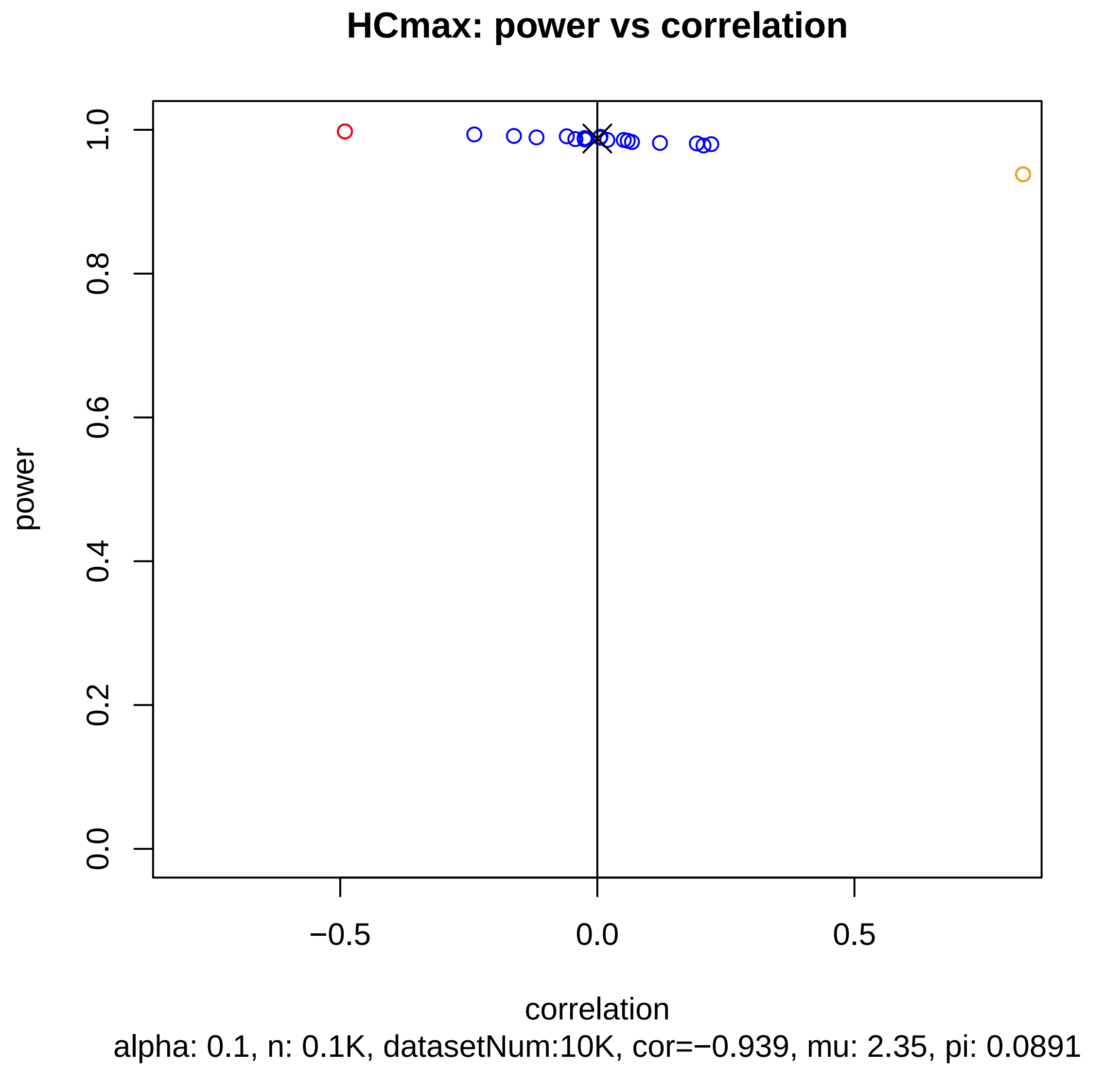
HCmax: power vs correlation
power
0.0
0.2
0.4
0.6
0.8
1.0
−0.5
0.0
0.5
correlation
alpha: 0.1, n: 0.1K, datasetNum:10K, cor=−0.939, mu: 2.35, pi: 0.0891

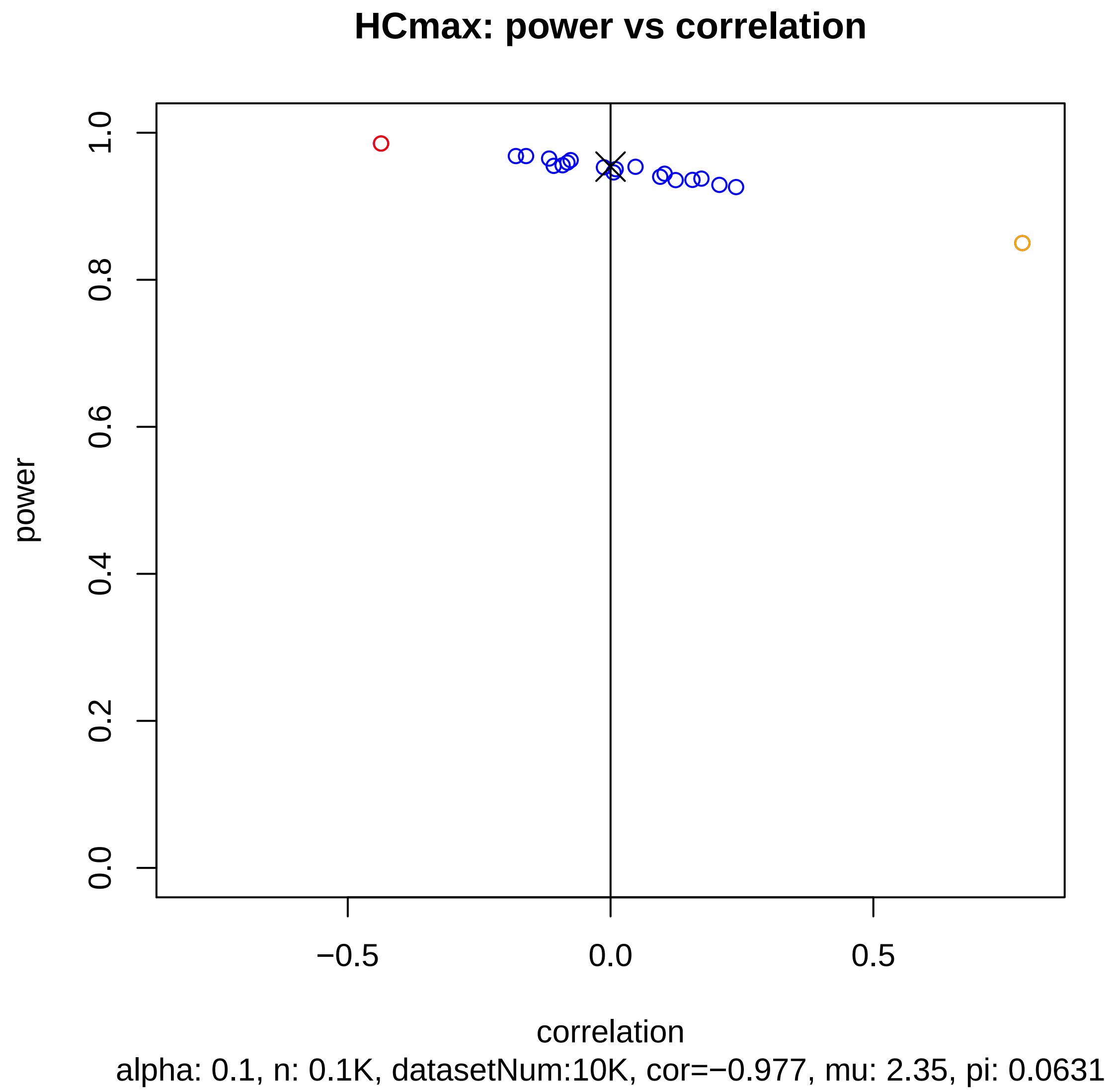
HCmax: power vs correlation
1.0
0.8
0.6
0.4
0.2
0.0
power
−0.5
0.0
0.5
correlation
alpha: 0.1, n: 0.1K, datasetNum:10K, cor=−0.977, mu: 2.35, pi: 0.0631

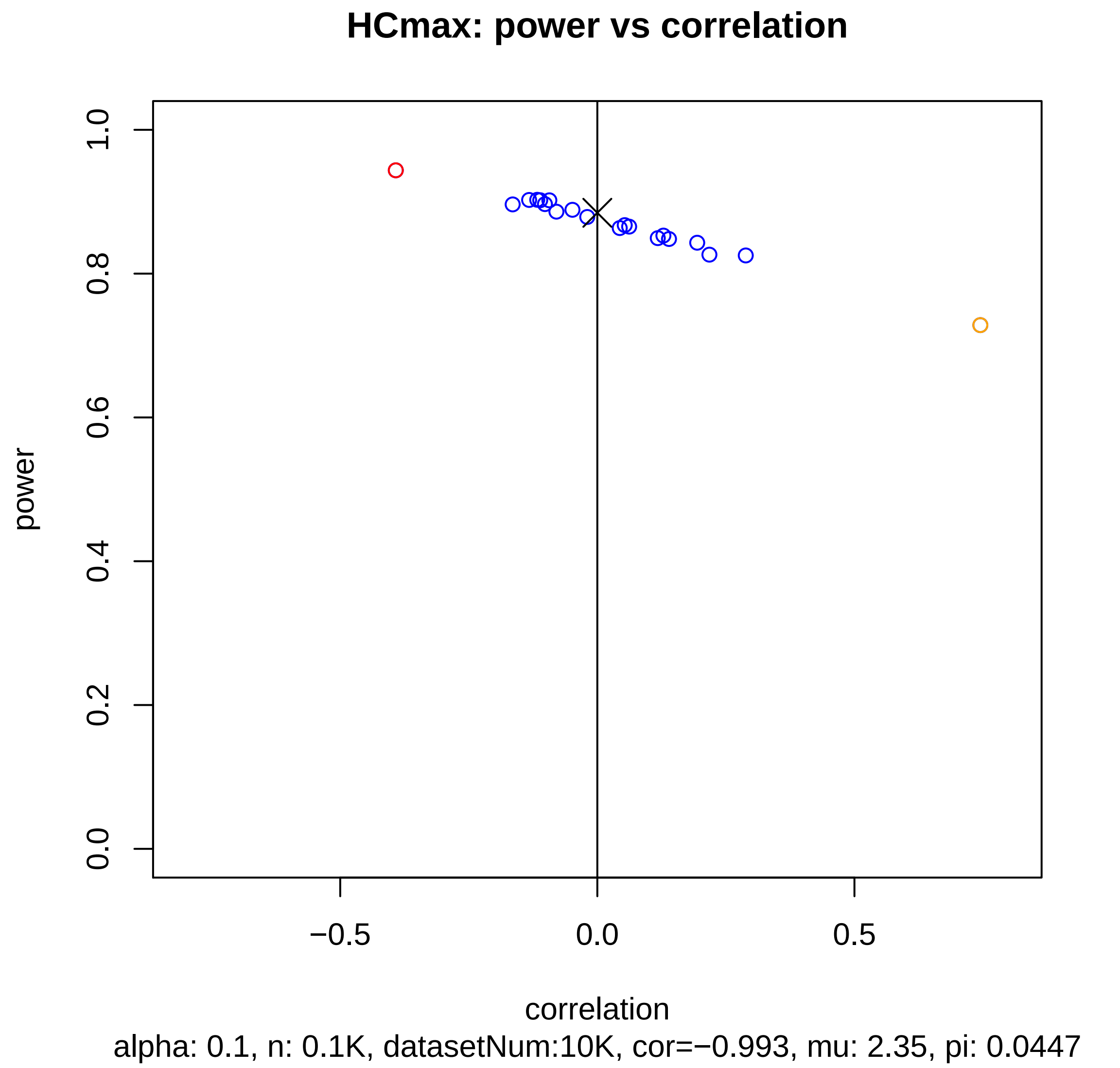

HCmax: power vs correlation
power
0.0
0.2
0.4
0.6
0.8
1.0
−0.5
0.0
0.5
correlation
alpha: 0.1, n: 0.1K, datasetNum:10K, cor=−0.993, mu: 2.35, pi: 0.0447

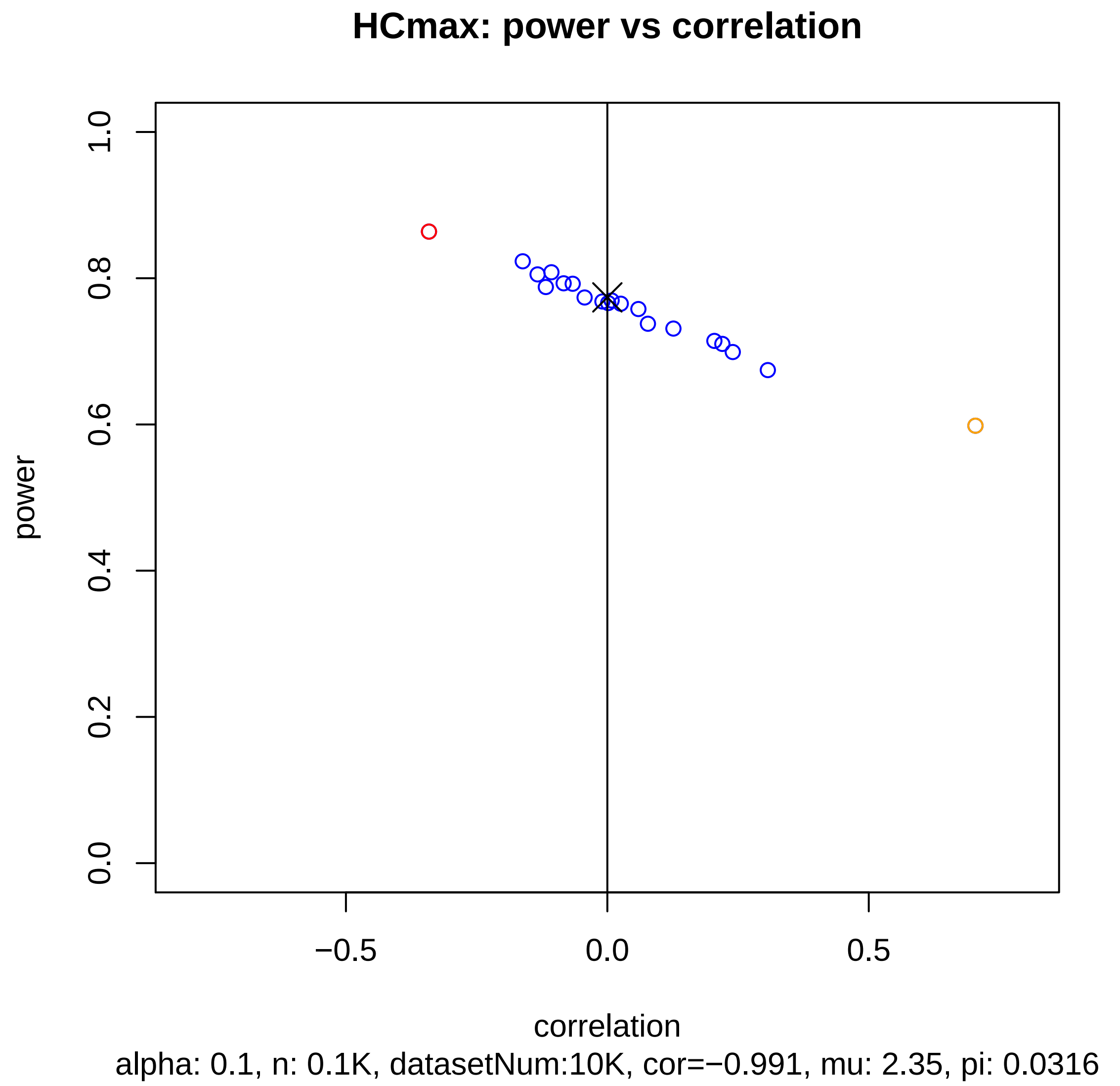
HCmax: power vs correlation
power
1.0
0.8
0.6
0.4
0.2
0.0
−0.5
0.0
0.5
correlation
alpha: 0.1, n: 0.1K, datasetNum:10K, cor=−0.991, mu: 2.35, pi: 0.0316

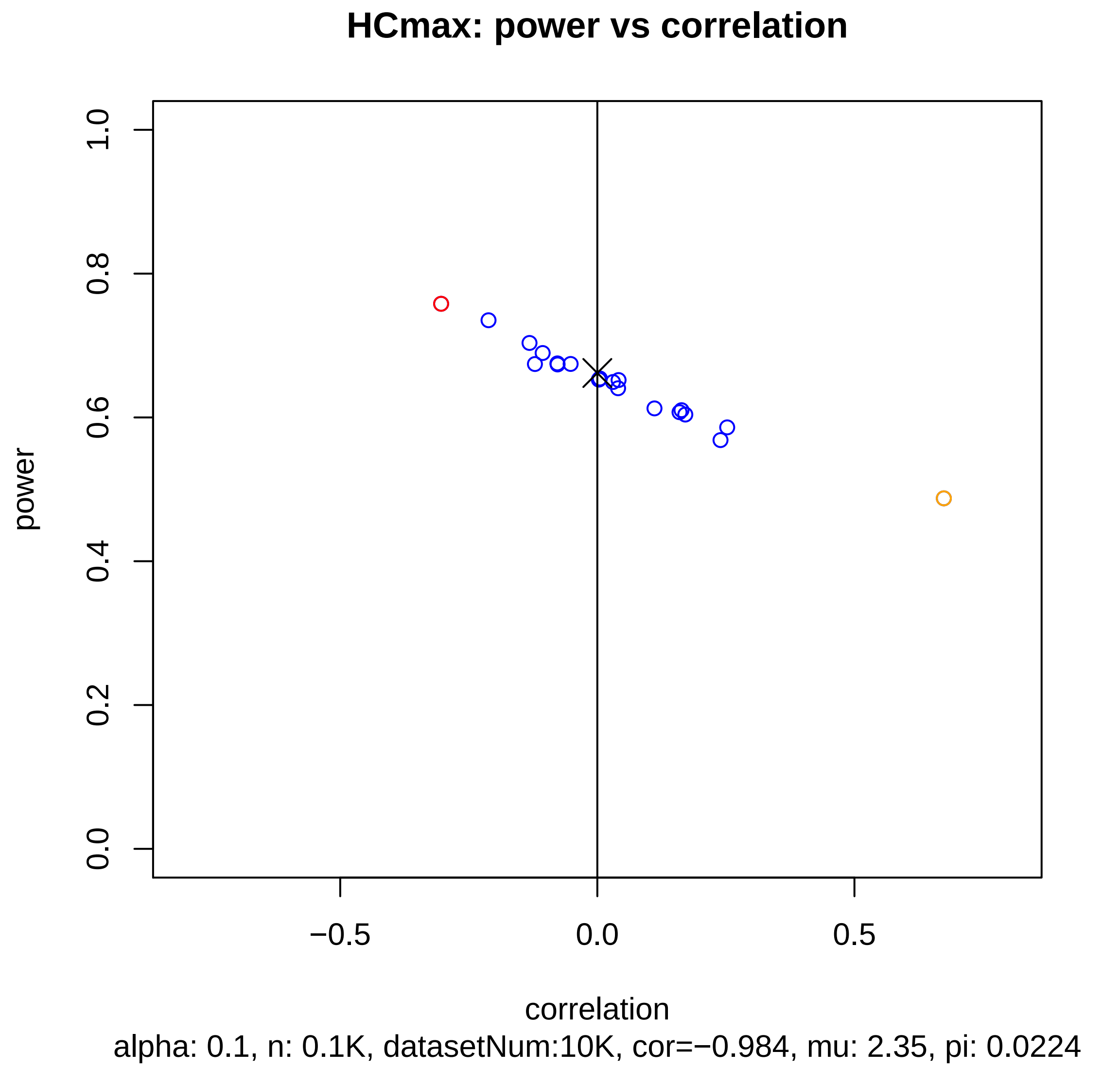
HCmax: power vs correlation
power
1.0
0.8
0.6
0.4
0.2
0.0
−0.5
0.0
0.5
correlation
alpha: 0.1, n: 0.1K, datasetNum:10K, cor=−0.984, mu: 2.35, pi: 0.0224

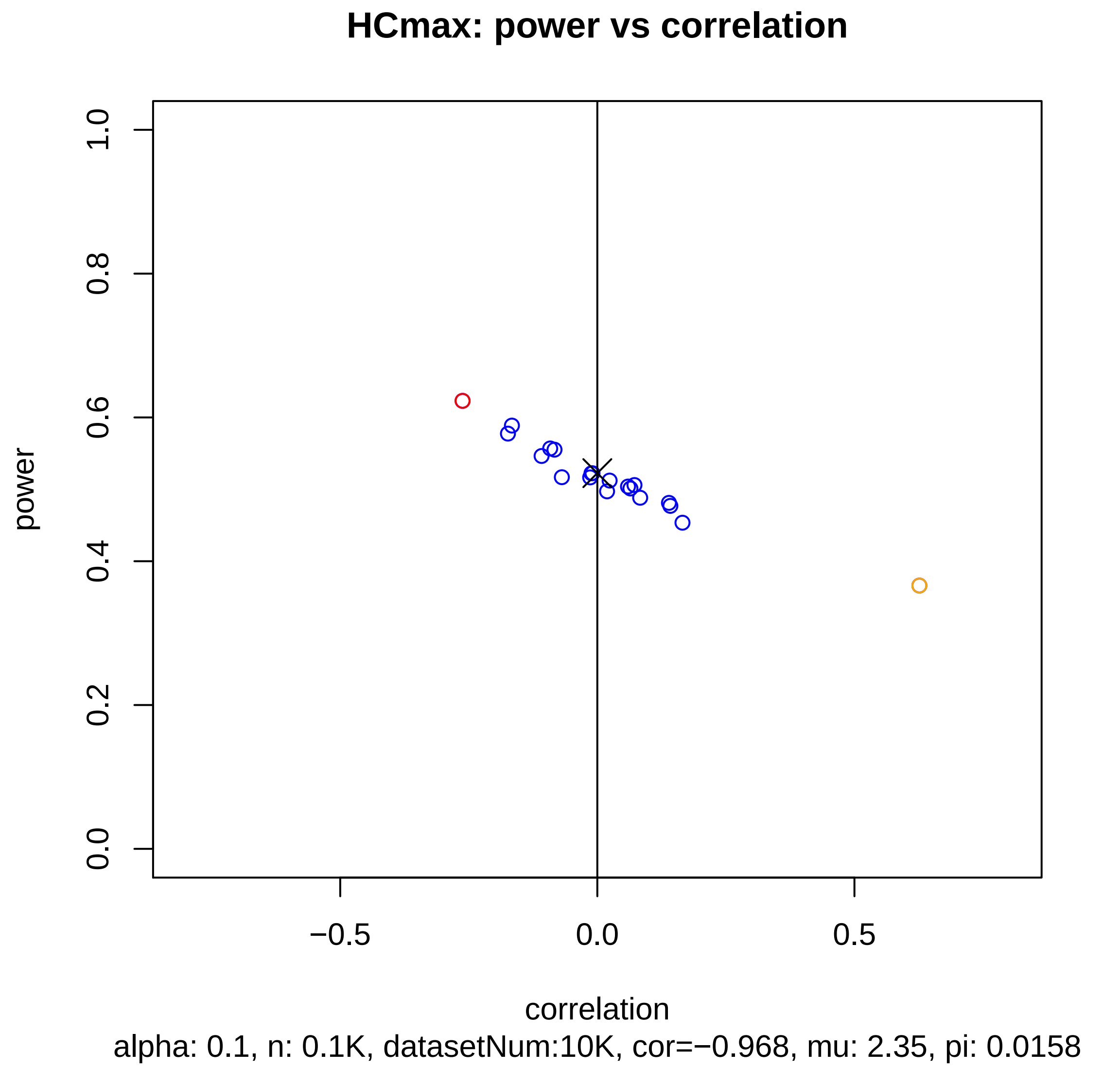

HCmax: power vs correlation
power
0.0
0.2
0.4
0.6
0.8
1.0
−0.5
0.0
0.5
correlation
alpha: 0.1, n: 0.1K, datasetNum:10K, cor=−0.968, mu: 2.35, pi: 0.0158

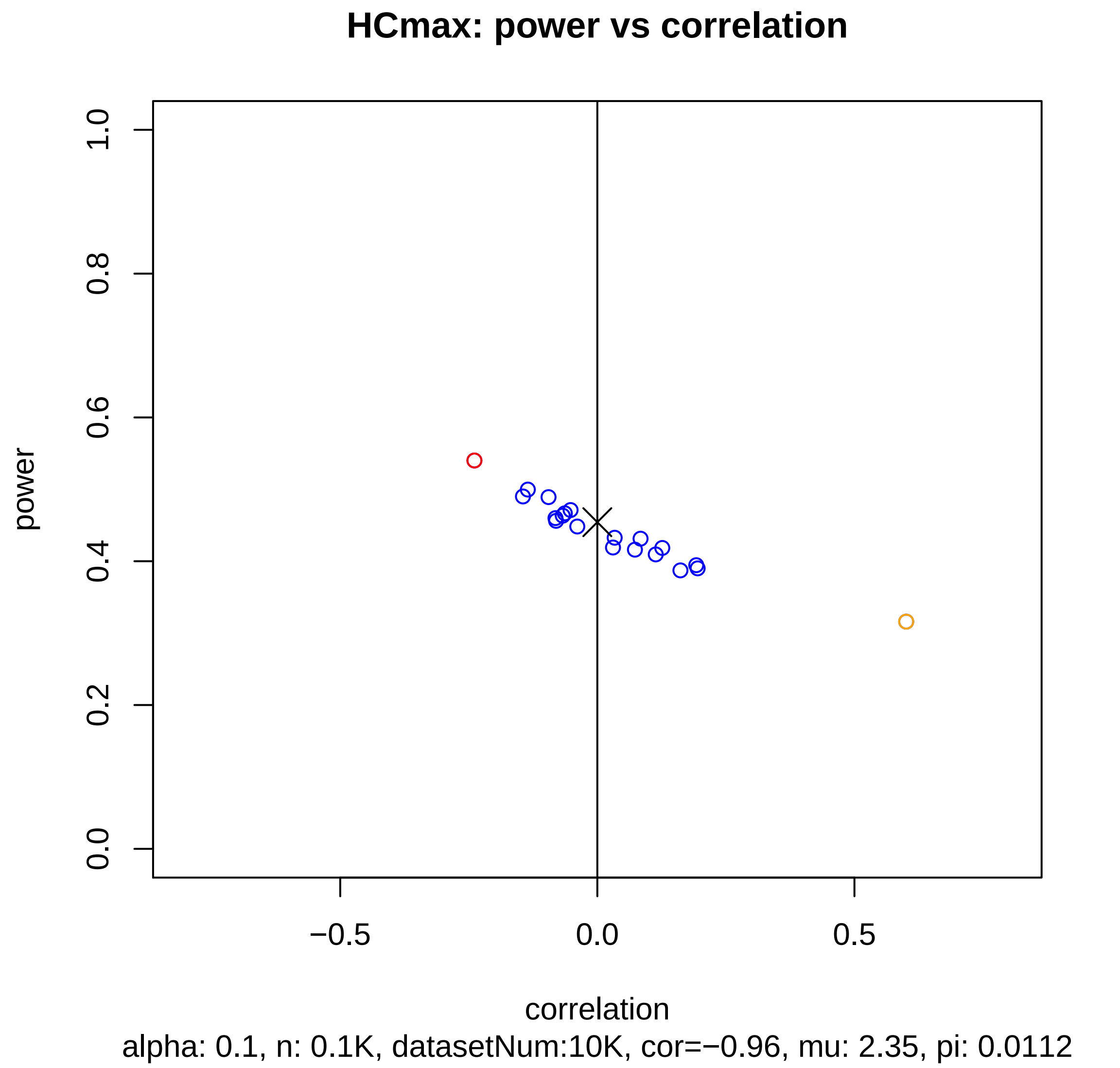

HCmax: power vs correlation
power
0.0
0.2
0.4
0.6
0.8
1.0
−0.5
0.0
0.5
correlation
alpha: 0.1, n: 0.1K, datasetNum:10K, cor=−0.96, mu: 2.35, pi: 0.0112

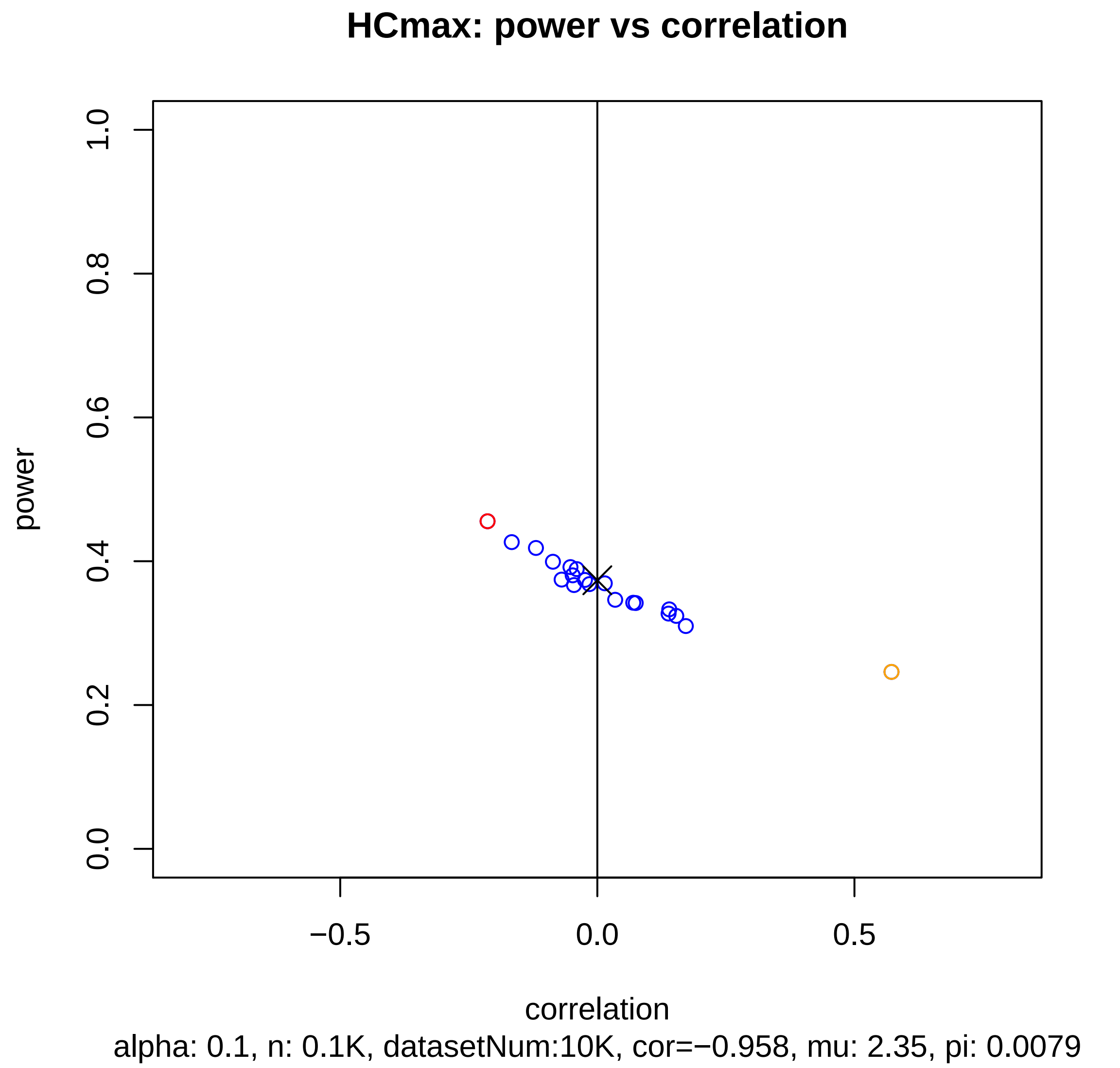
HCmax: power vs correlation
power
1.0
0.8
0.6
0.4
0.2
0.0
−0.5
0.0
0.5
correlation
alpha: 0.1, n: 0.1K, datasetNum:10K, cor=−0.958, mu: 2.35, pi: 0.0079

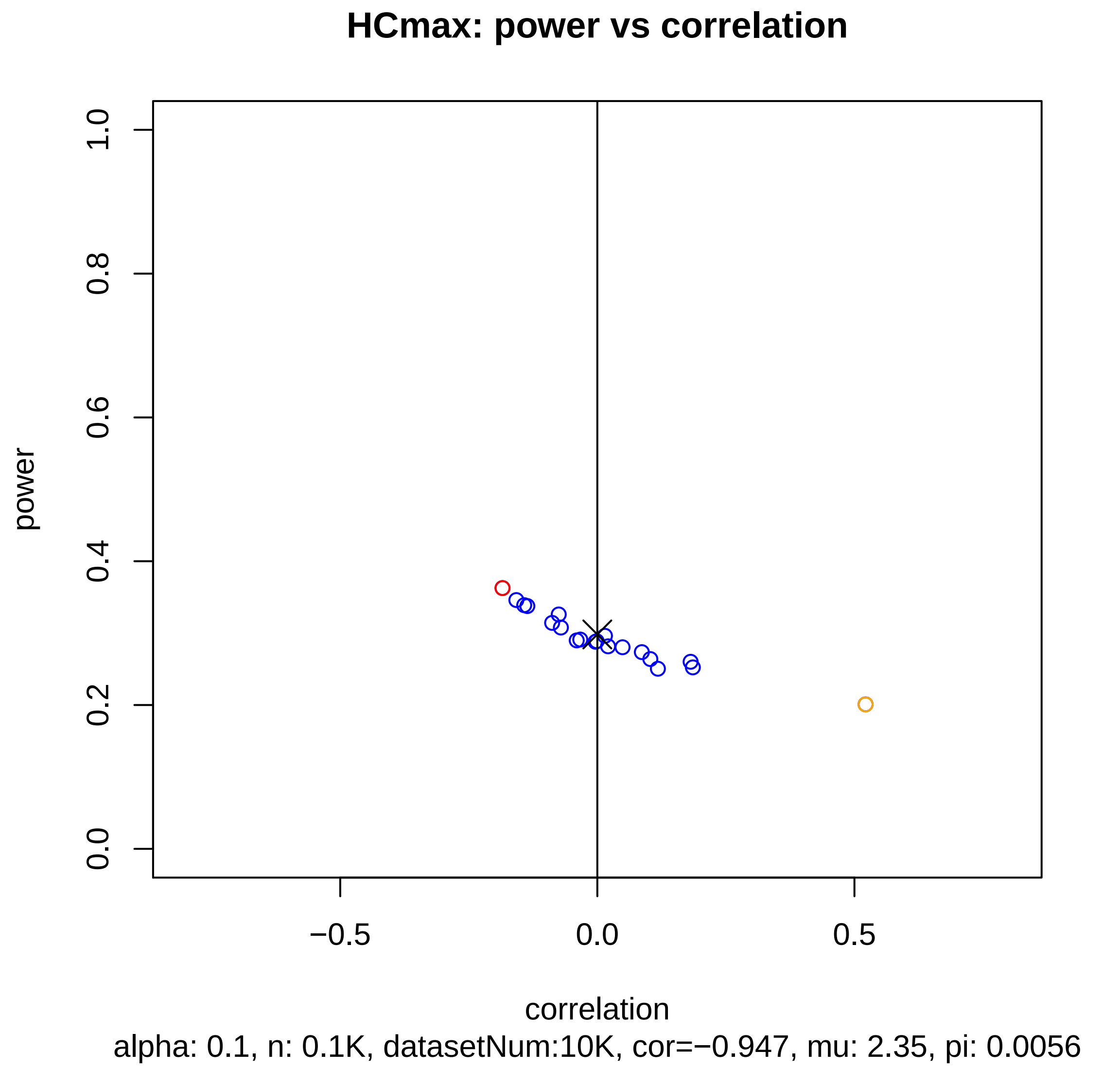
HCmax: power vs correlation
power
1.0
0.8
0.6
0.4
0.2
0.0
−0.5
0.0
0.5
correlation
alpha: 0.1, n: 0.1K, datasetNum:10K, cor=−0.947, mu: 2.35, pi: 0.0056

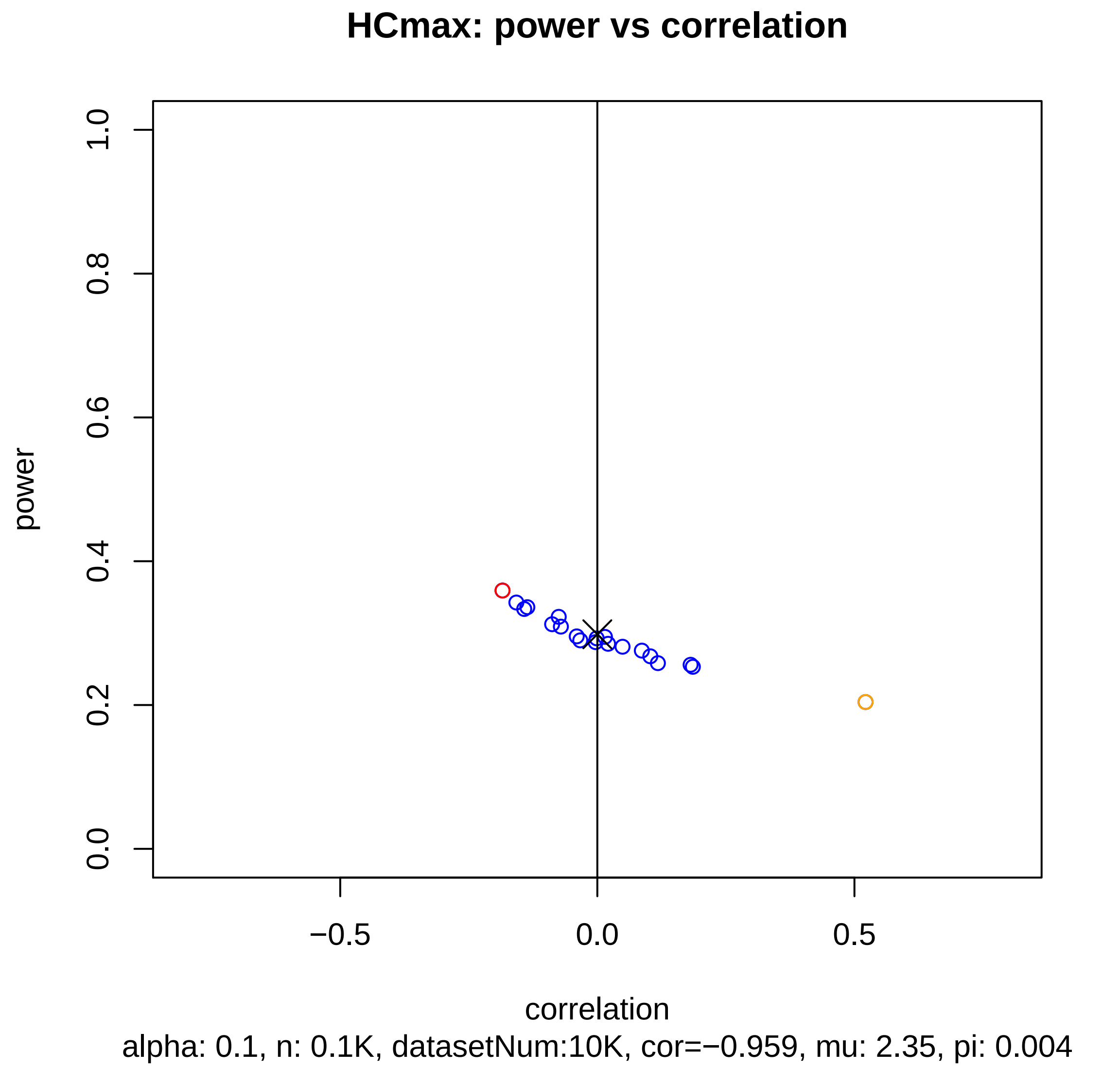
HCmax: power vs correlation
power
1.0
0.8
0.6
0.4
0.2
0.0
−0.5
0.0
0.5
correlation
alpha: 0.1, n: 0.1K, datasetNum:10K, cor=−0.959, mu: 2.35, pi: 0.004

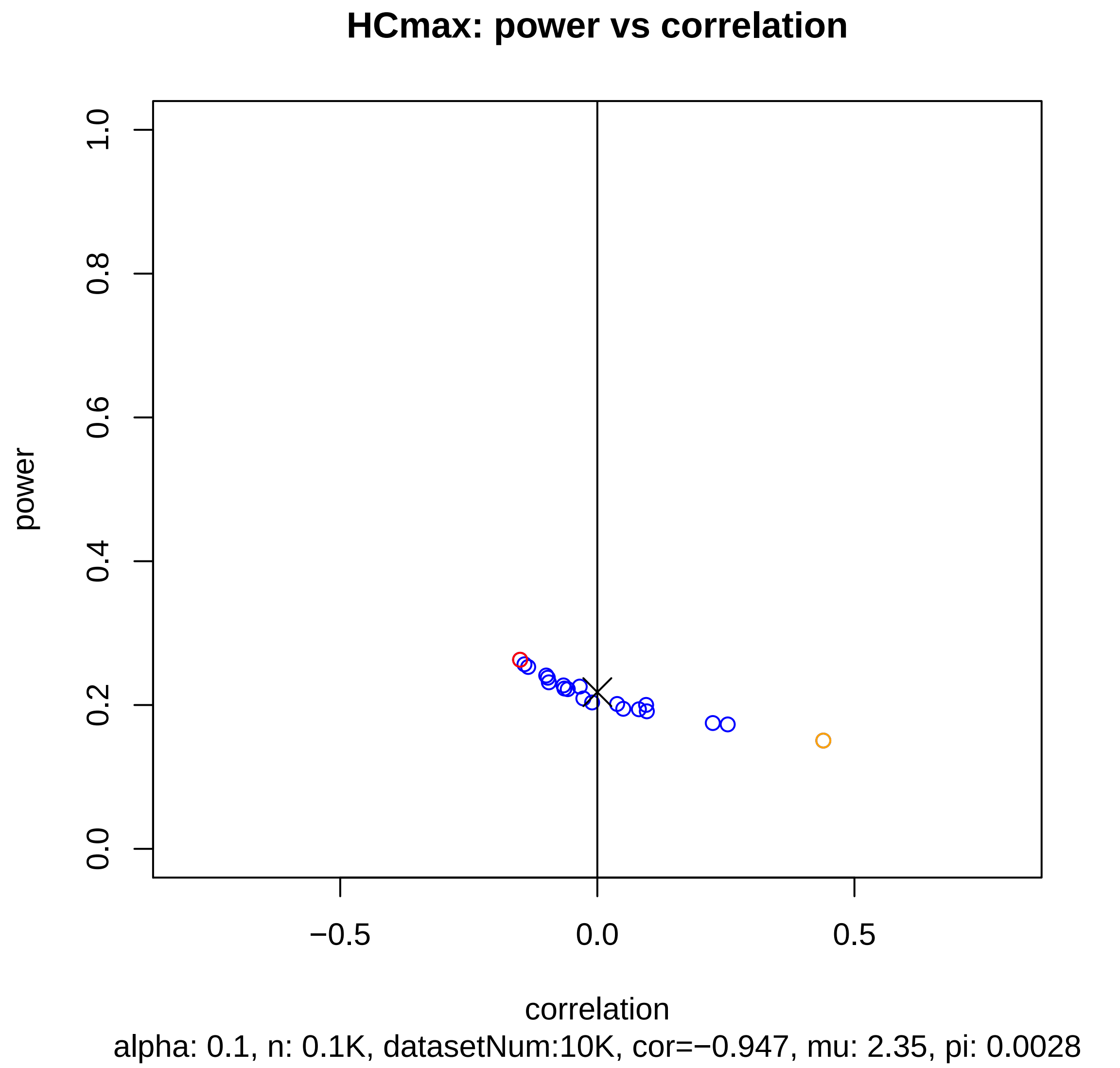
HCmax: power vs correlation
power
1.0
0.8
0.6
0.4
0.2
0.0
−0.5
0.0
0.5
correlation
alpha: 0.1, n: 0.1K, datasetNum:10K, cor=−0.947, mu: 2.35, pi: 0.0028

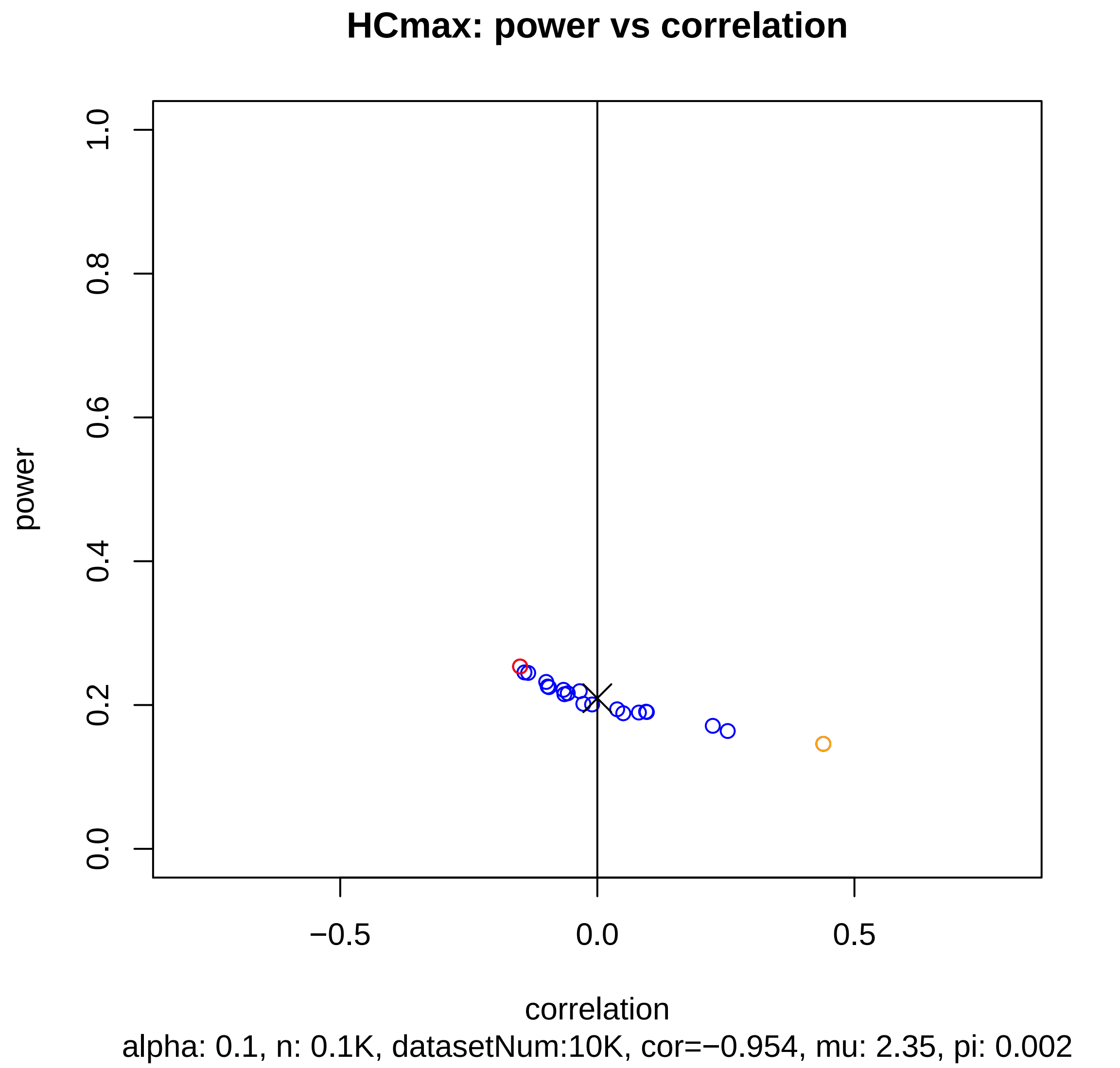
HCmax: power vs correlation
power
1.0
0.8
0.6
0.4
0.2
0.0
−0.5
0.0
0.5
correlation
alpha: 0.1, n: 0.1K, datasetNum:10K, cor=−0.954, mu: 2.35, pi: 0.002

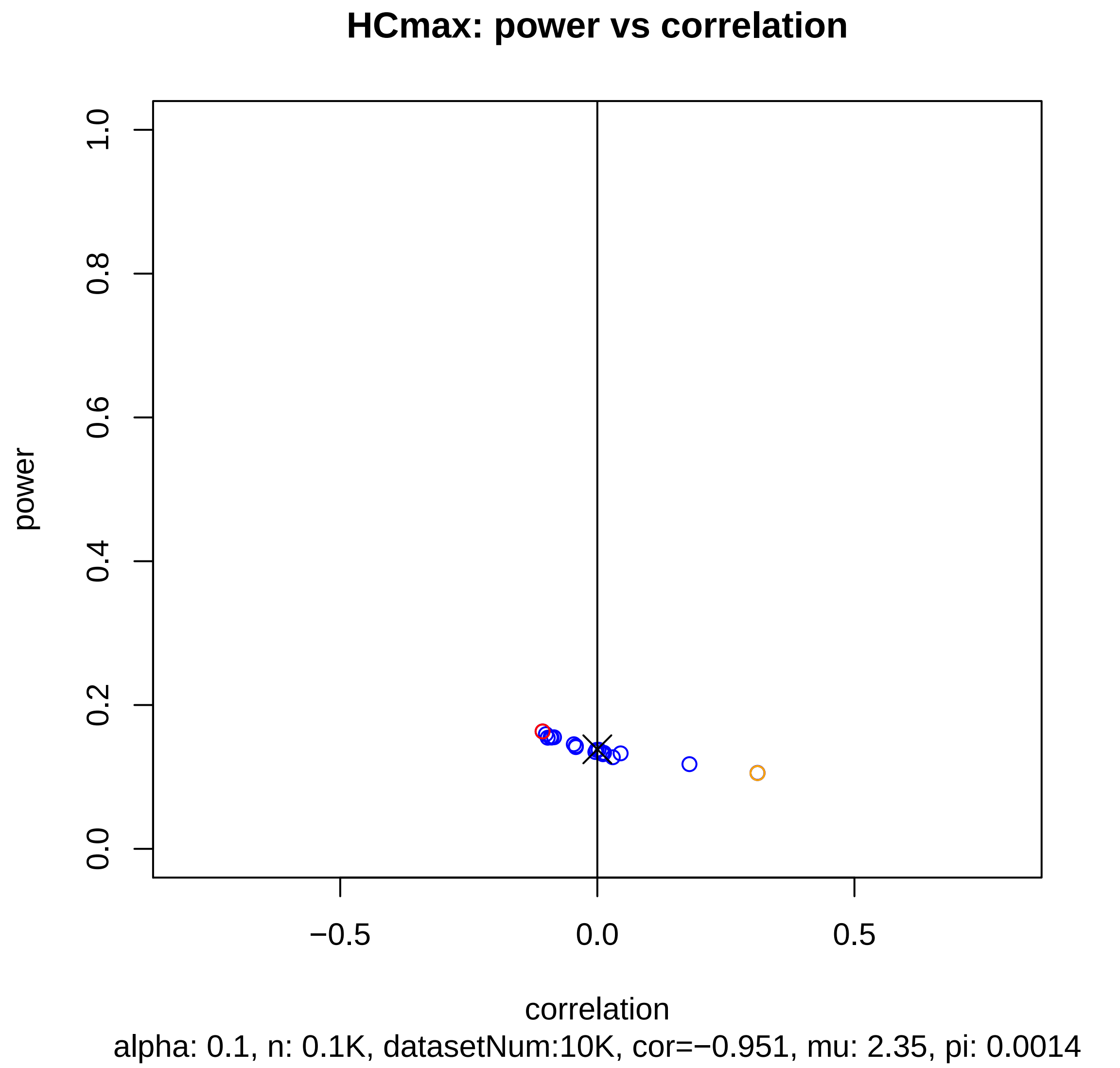

HCmax: power vs correlation
power
0.0
0.2
0.4
0.6
0.8
1.0
−0.5
0.0
0.5
correlation
alpha: 0.1, n: 0.1K, datasetNum:10K, cor=−0.951, mu: 2.35, pi: 0.0014

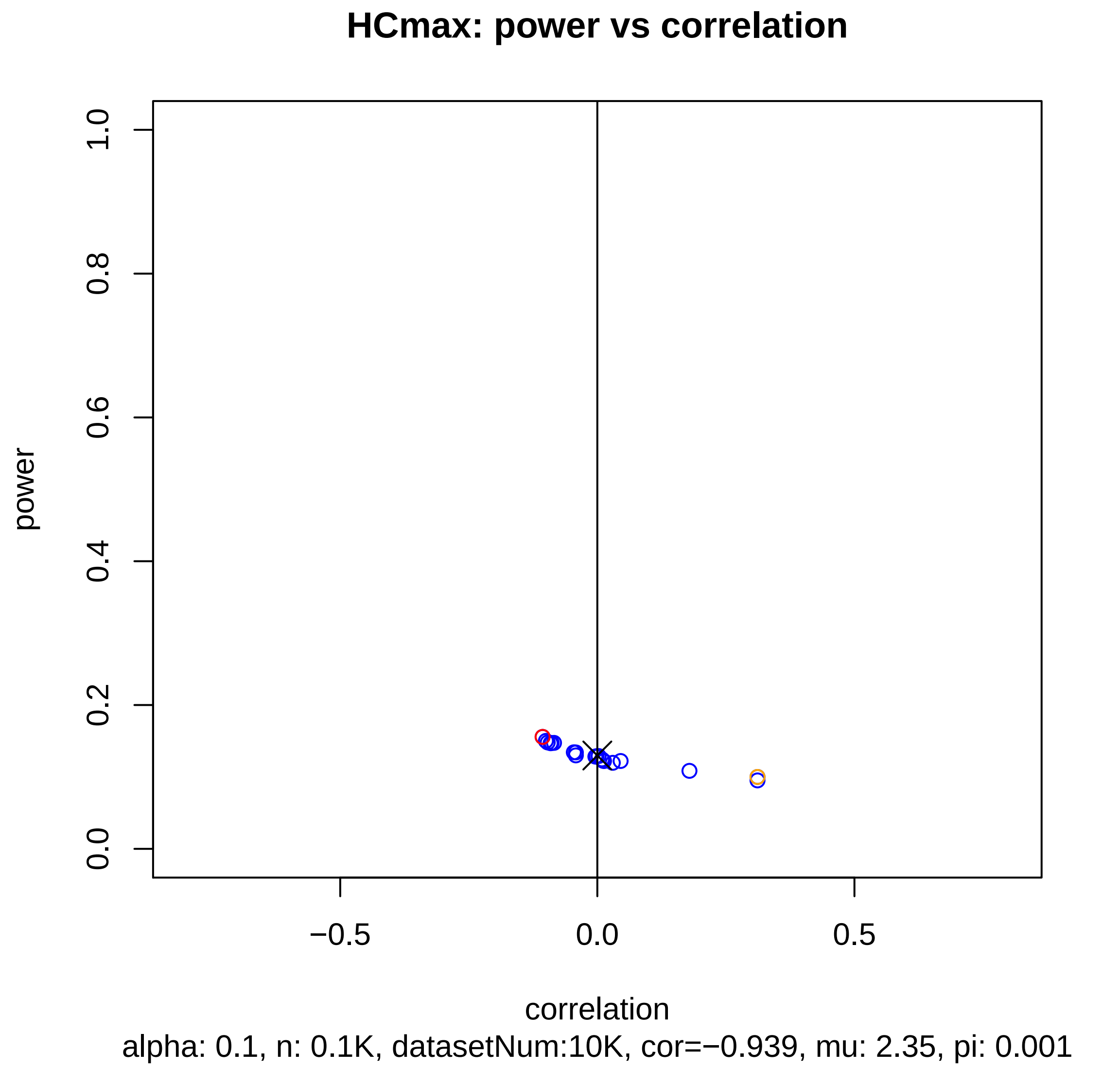
HCmax: power vs correlation
power
1.0
0.8
0.6
0.4
0.2
0.0
−0.5
0.0
0.5
correlation
alpha: 0.1, n: 0.1K, datasetNum:10K, cor=−0.939, mu: 2.35, pi: 0.001

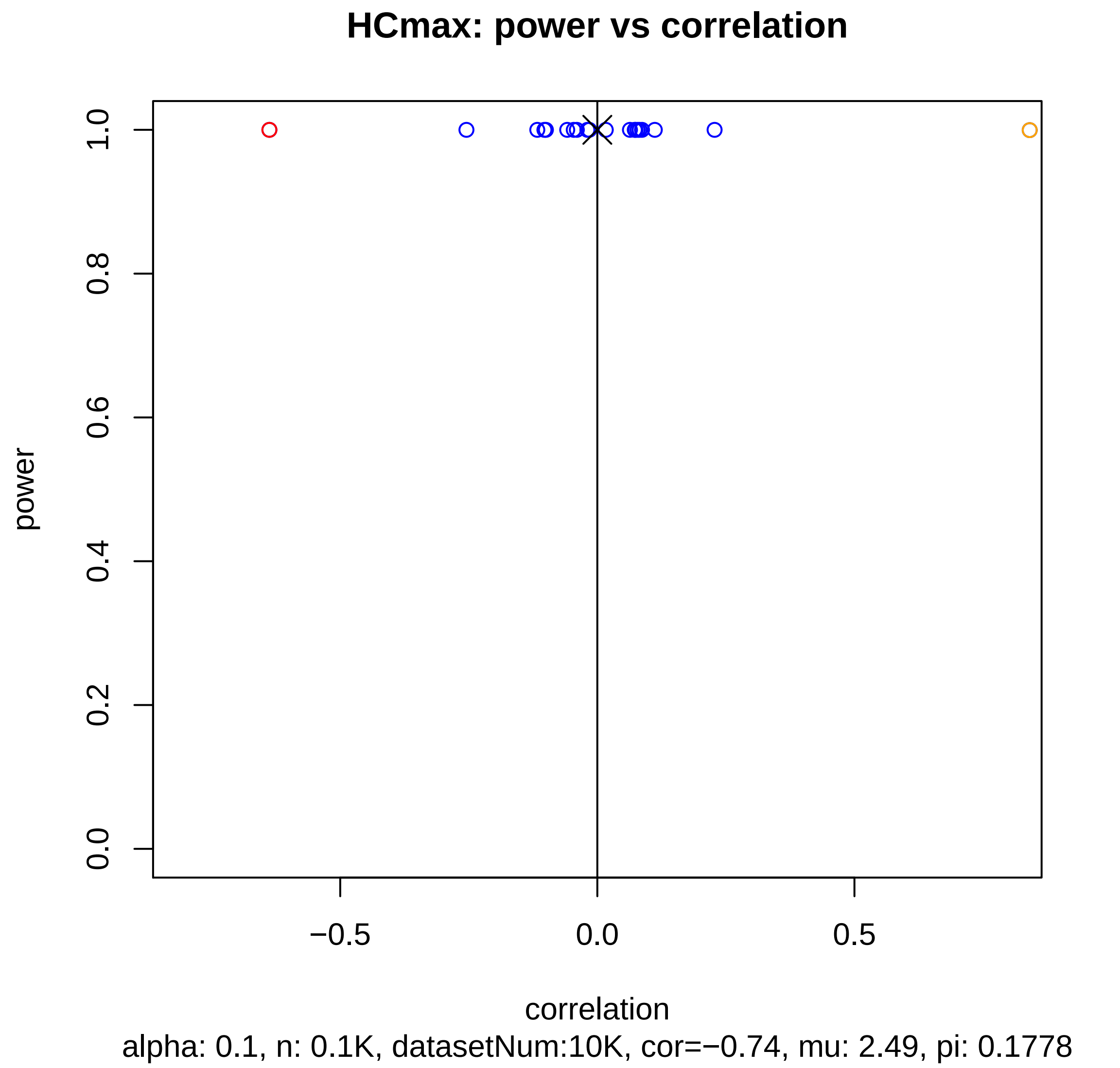
HCmax: power vs correlation
1.0
0.8
0.6
0.4
0.2
0.0
power
−0.5
0.0
0.5
correlation
alpha: 0.1, n: 0.1K, datasetNum:10K, cor=−0.74, mu: 2.49, pi: 0.1778

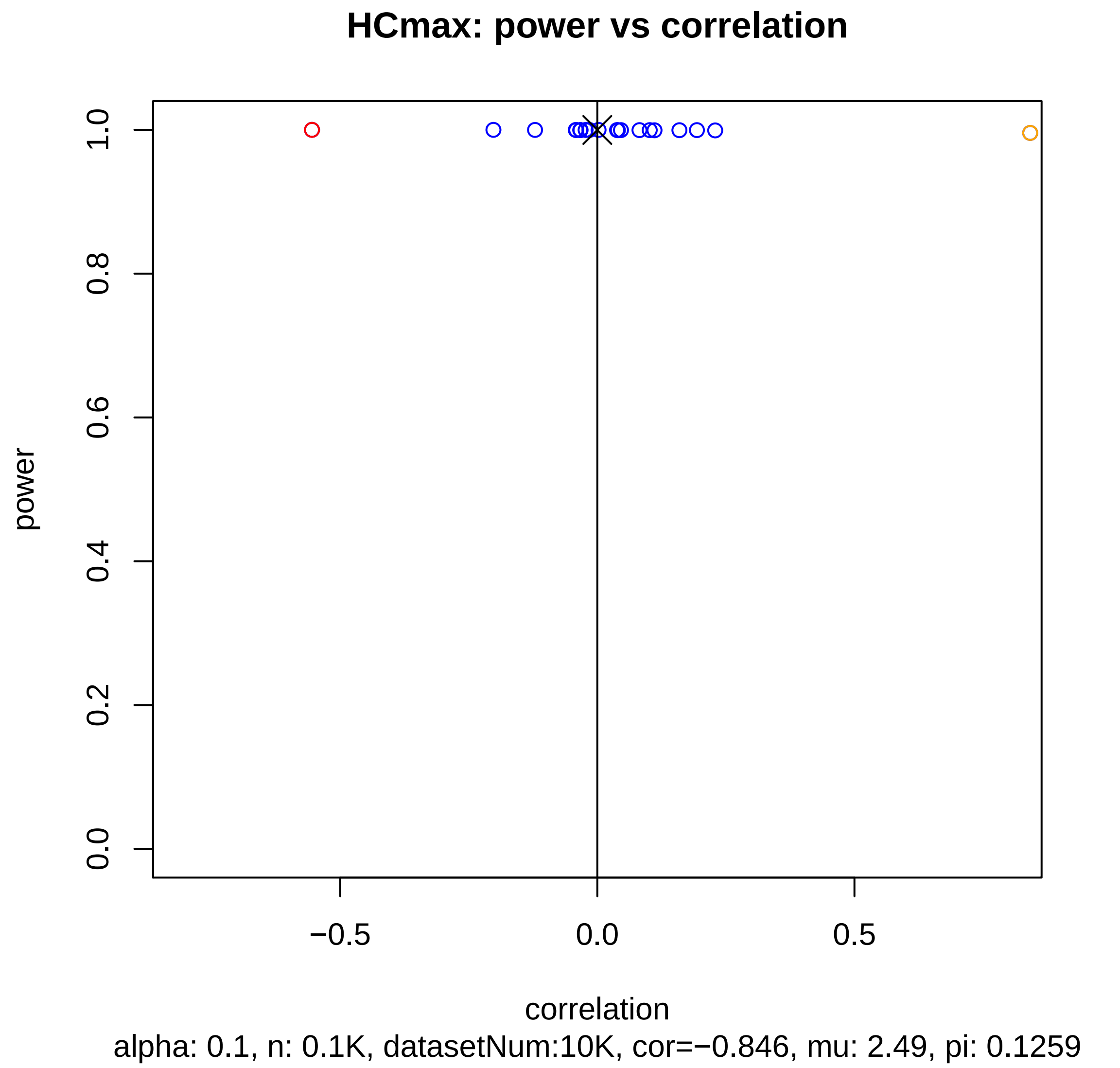
HCmax: power vs correlation
power
0.0
0.2
0.4
0.6
0.8
1.0
−0.5
0.0
0.5
correlation
alpha: 0.1, n: 0.1K, datasetNum:10K, cor=−0.846, mu: 2.49, pi: 0.1259

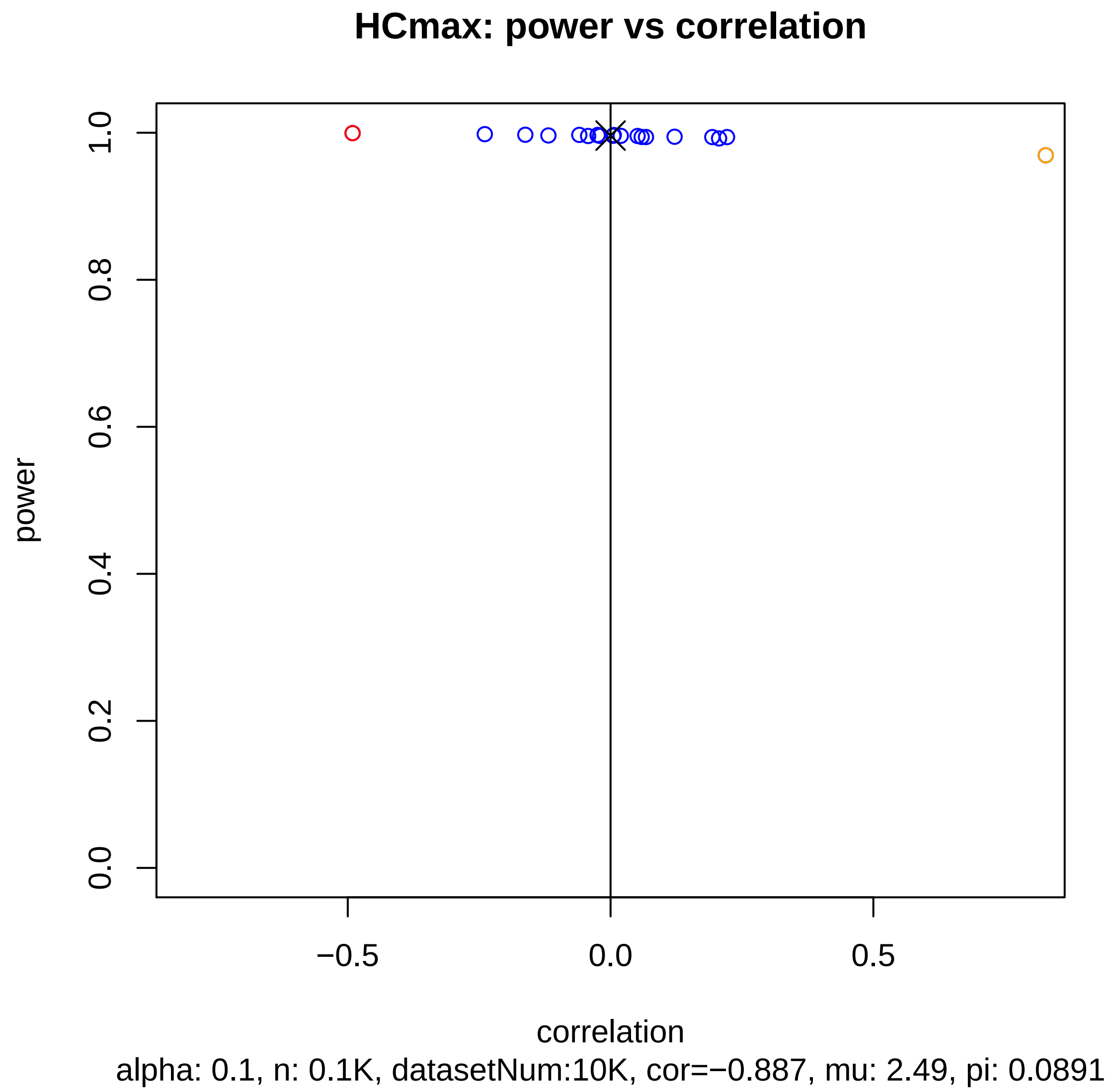

HCmax: power vs correlation
power
correlation
0.0
0.2
0.4
0.6
0.8
1.0
−0.5
0.0
0.5
alpha: 0.1, n: 0.1K, datasetNum:10K, cor=−0.887, mu: 2.49, pi: 0.0891

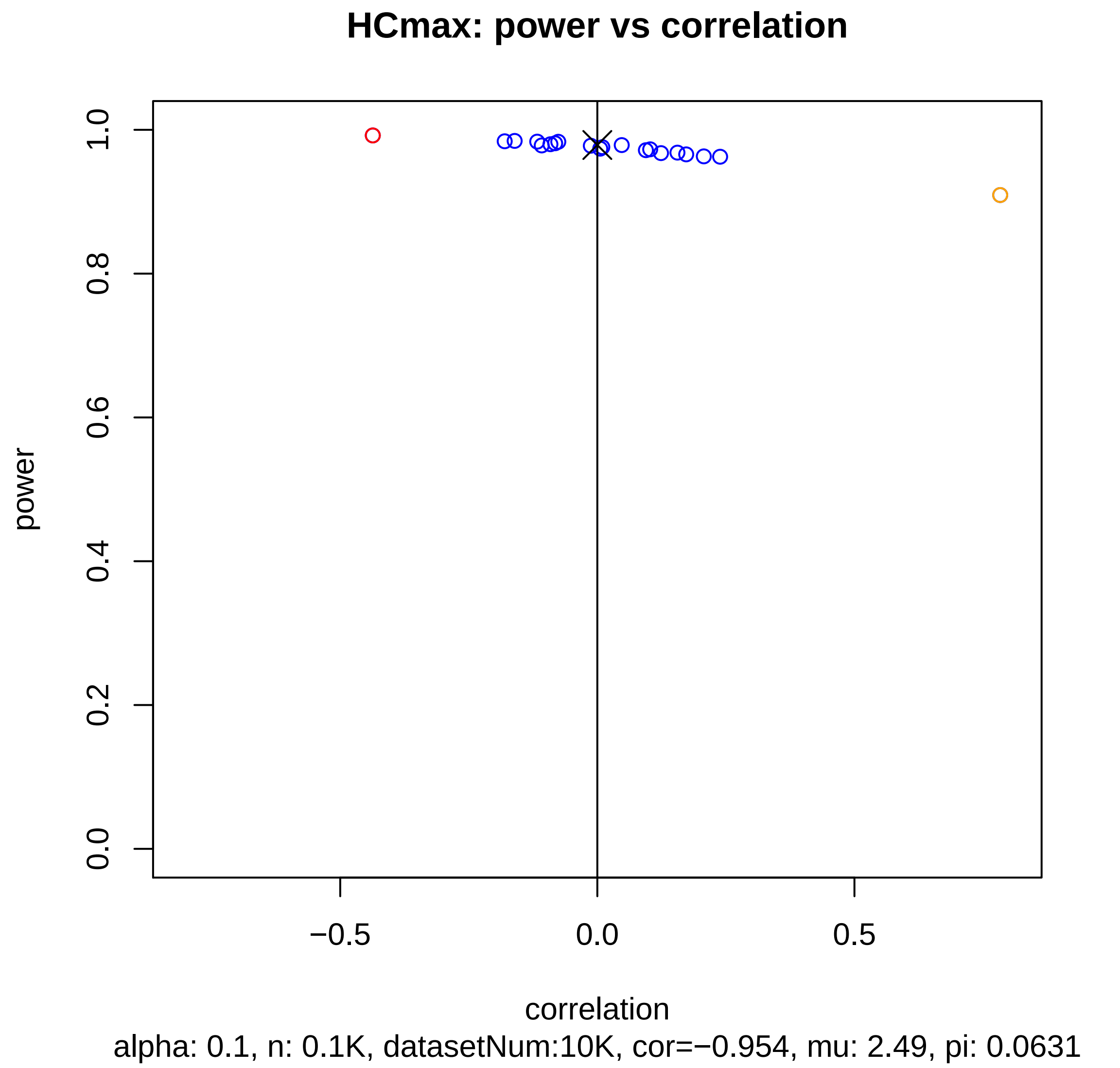
HCmax: power vs correlation
power
1.0
0.8
0.6
0.4
0.2
0.0
−0.5
0.0
0.5
correlation
alpha: 0.1, n: 0.1K, datasetNum:10K, cor=−0.954, mu: 2.49, pi: 0.0631

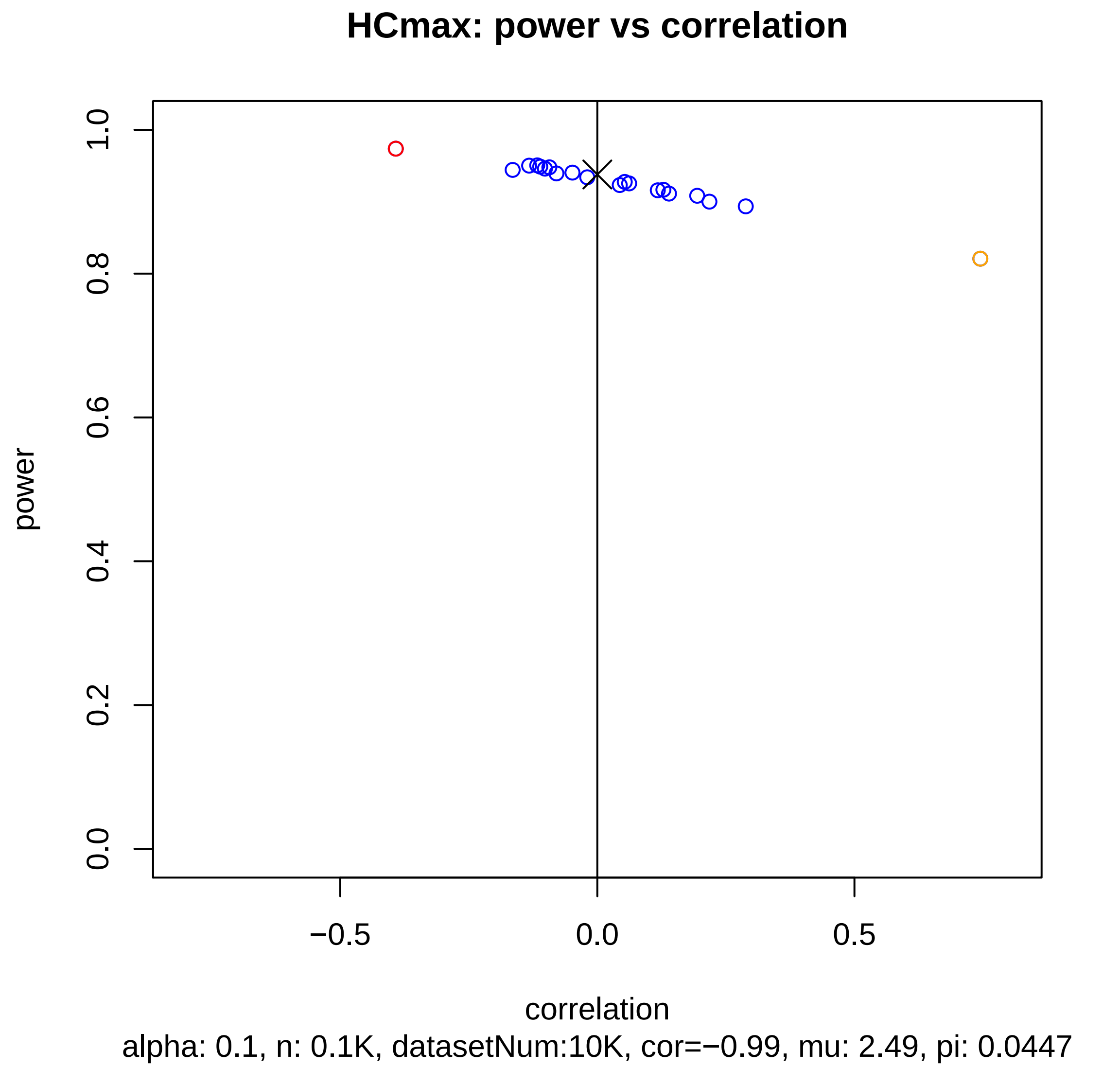
HCmax: power vs correlation
power
1.0
0.8
0.6
0.4
0.2
0.0
−0.5
0.0
0.5
correlation
alpha: 0.1, n: 0.1K, datasetNum:10K, cor=−0.99, mu: 2.49, pi: 0.0447

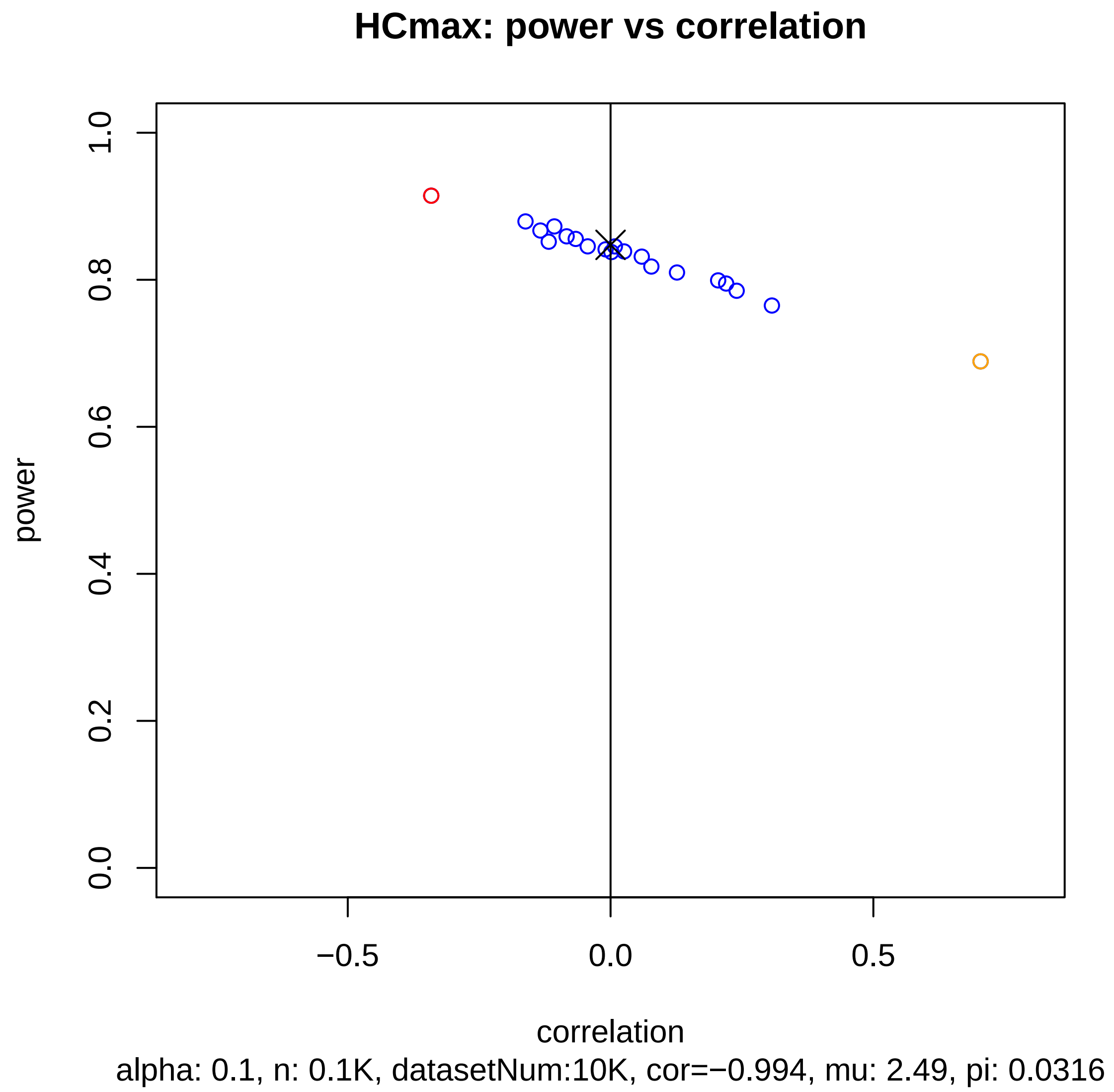

HCmax: power vs correlation
power
1.0
0.8
0.6
0.4
0.2
0.0
−0.5
0.0
0.5
correlation
alpha: 0.1, n: 0.1K, datasetNum:10K, cor=−0.994, mu: 2.49, pi: 0.0316

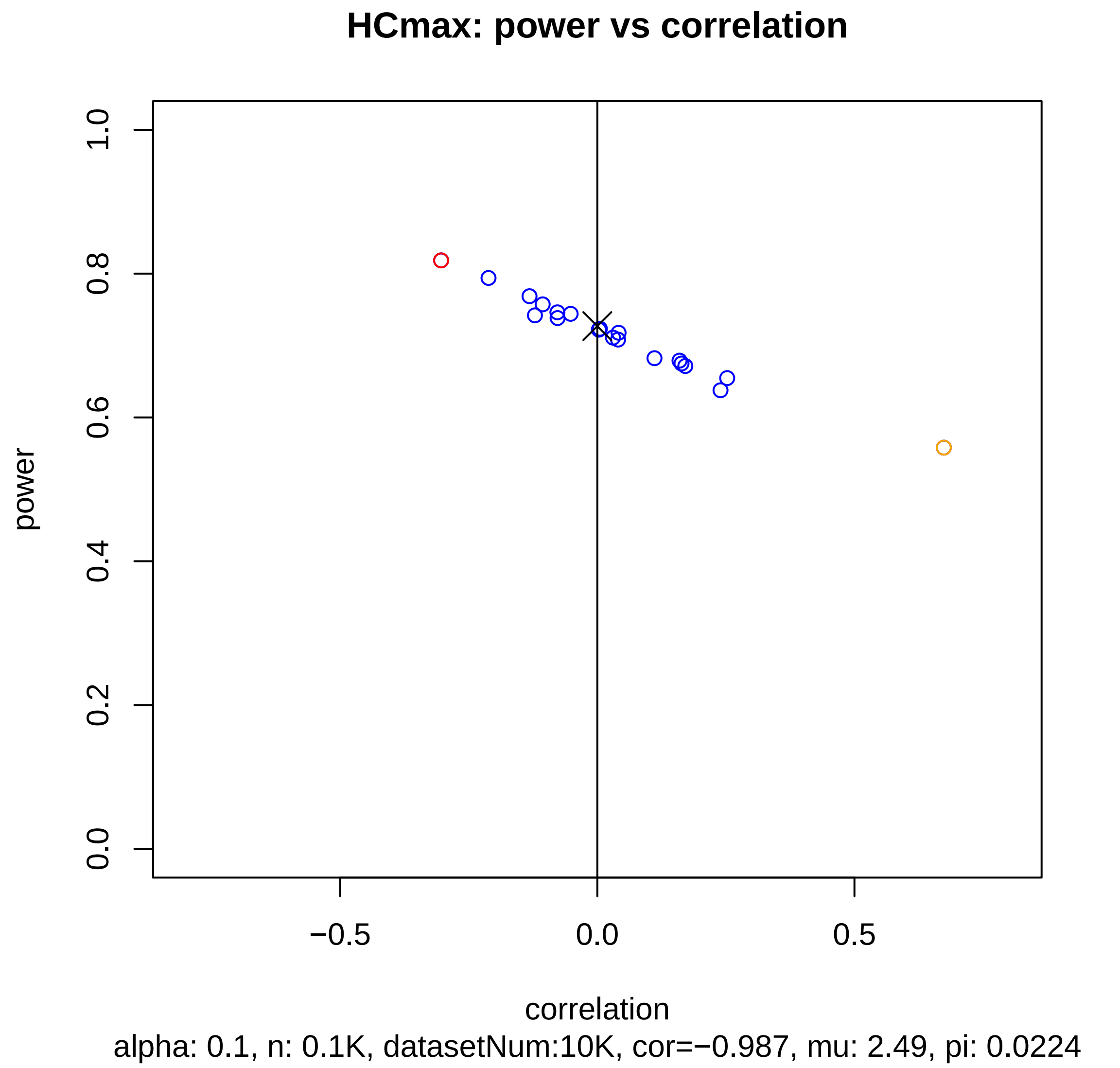
HCmax: power vs correlation
power
0.0
0.2
0.4
0.6
0.8
1.0
−0.5
0.0
0.5
correlation
alpha: 0.1, n: 0.1K, datasetNum:10K, cor=−0.987, mu: 2.49, pi: 0.0224

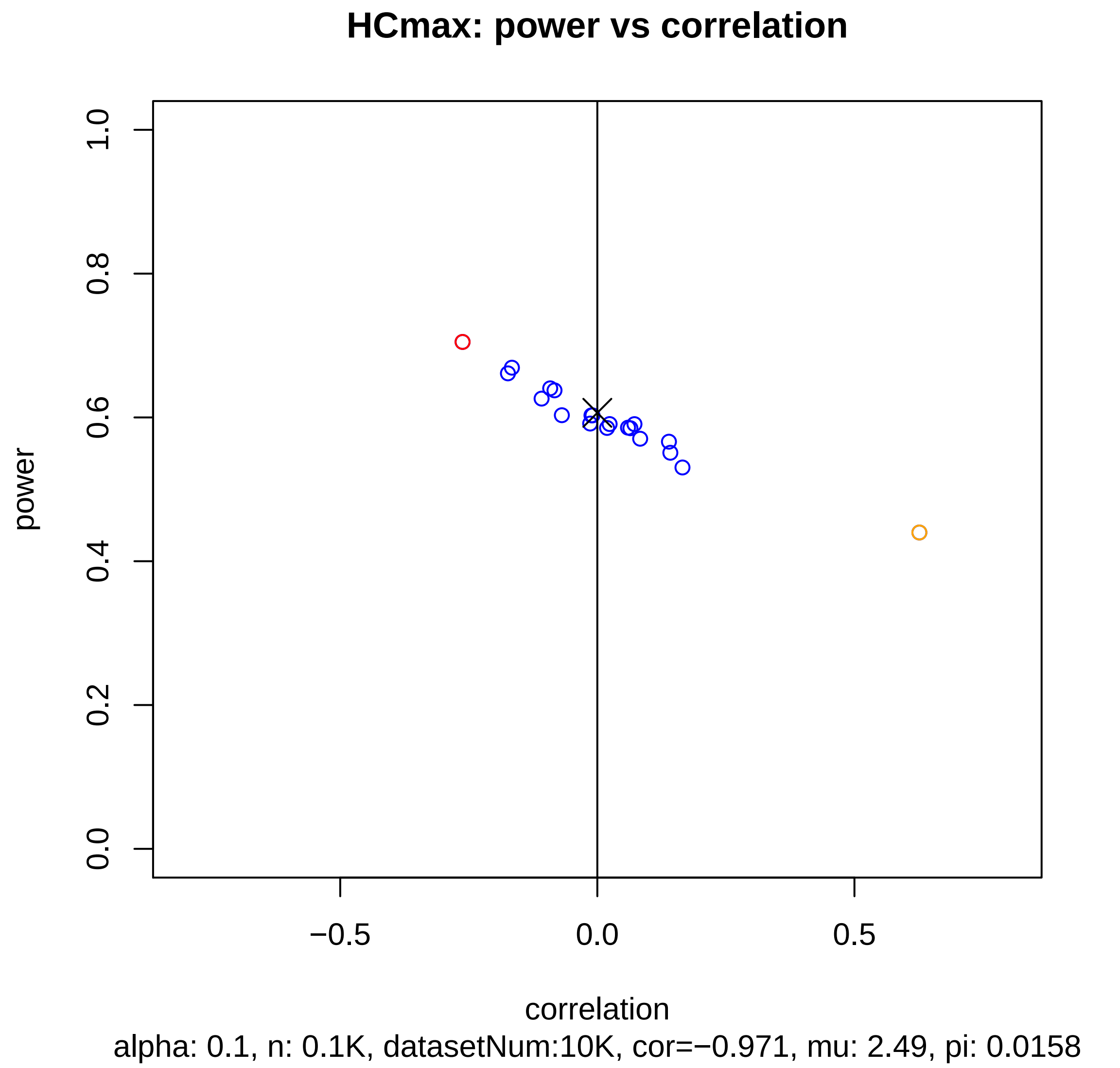

HCmax: power vs correlation
power
0.0
0.2
0.4
0.6
0.8
1.0
−0.5
0.0
0.5
correlation
alpha: 0.1, n: 0.1K, datasetNum:10K, cor=−0.971, mu: 2.49, pi: 0.0158

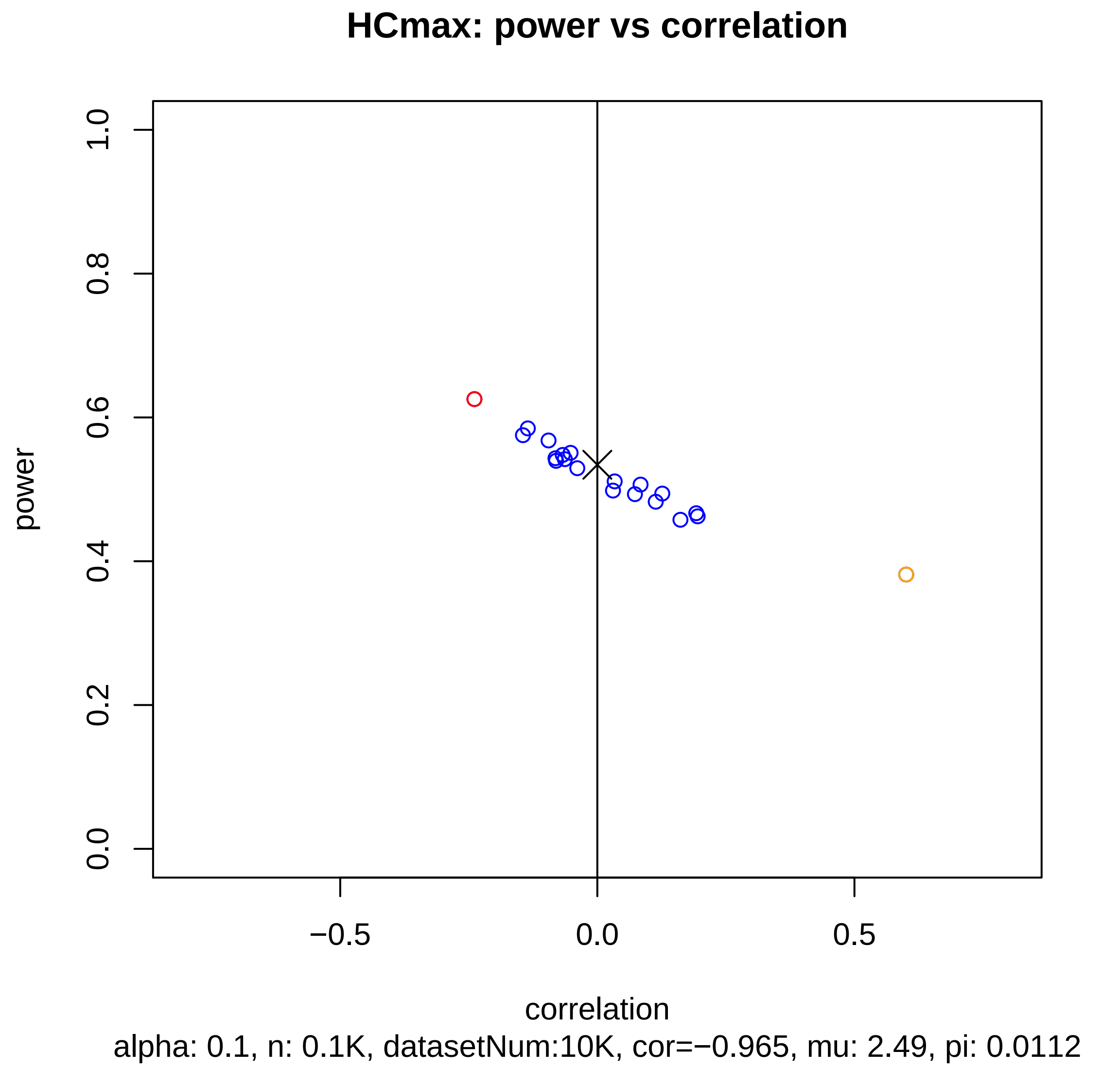

HCmax: power vs correlation
power
1.0
0.8
0.6
0.4
0.2
0.0
−0.5
0.0
0.5
correlation
alpha: 0.1, n: 0.1K, datasetNum:10K, cor=−0.965, mu: 2.49, pi: 0.0112

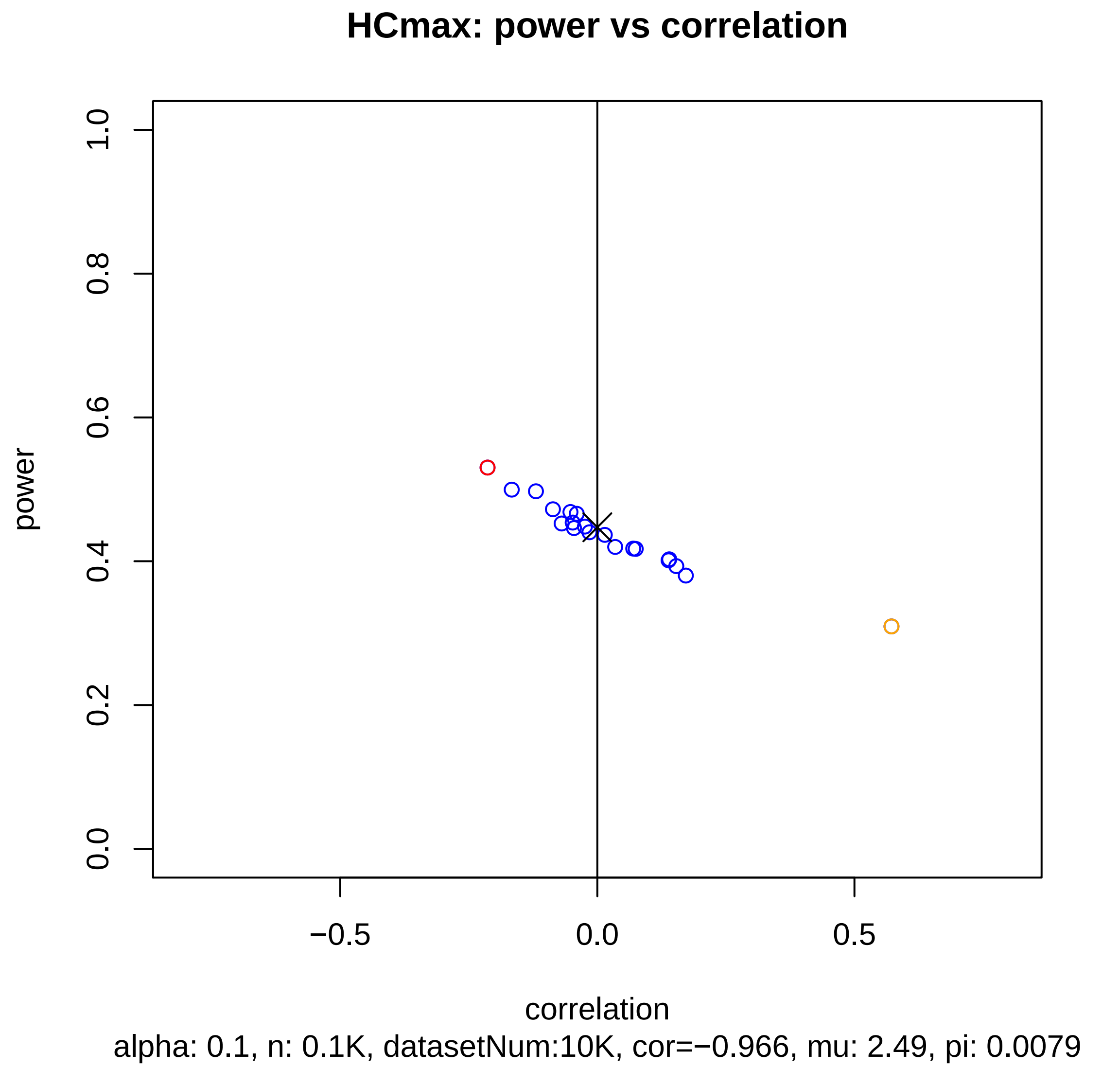
HCmax: power vs correlation
power
1.0
0.8
0.6
0.4
0.2
0.0
−0.5
0.0
0.5
correlation
alpha: 0.1, n: 0.1K, datasetNum:10K, cor=−0.966, mu: 2.49, pi: 0.0079

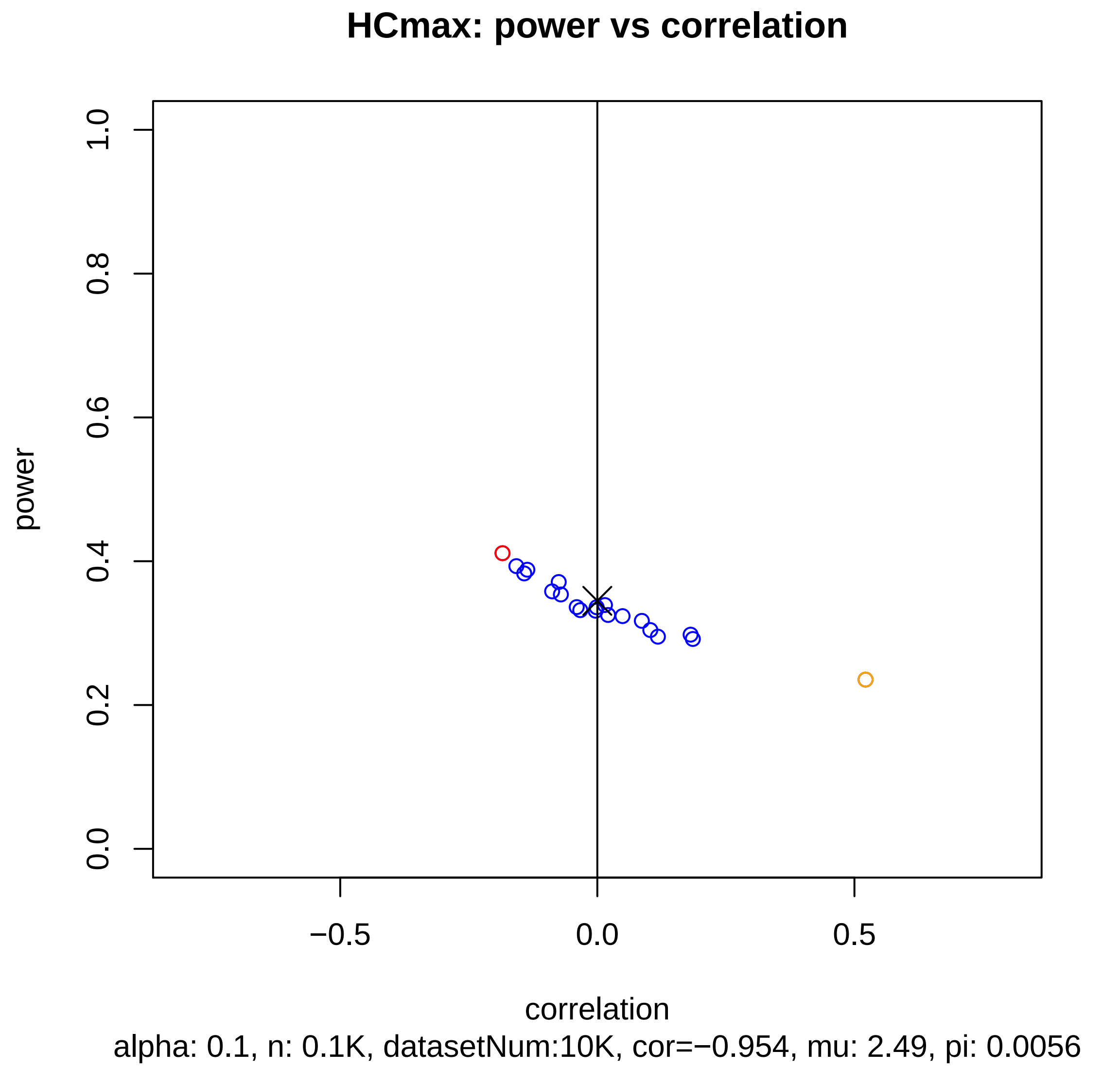

HCmax: power vs correlation
power
1.0
0.8
0.6
0.4
0.2
0.0
−0.5
0.0
0.5
correlation
alpha: 0.1, n: 0.1K, datasetNum:10K, cor=−0.954, mu: 2.49, pi: 0.0056

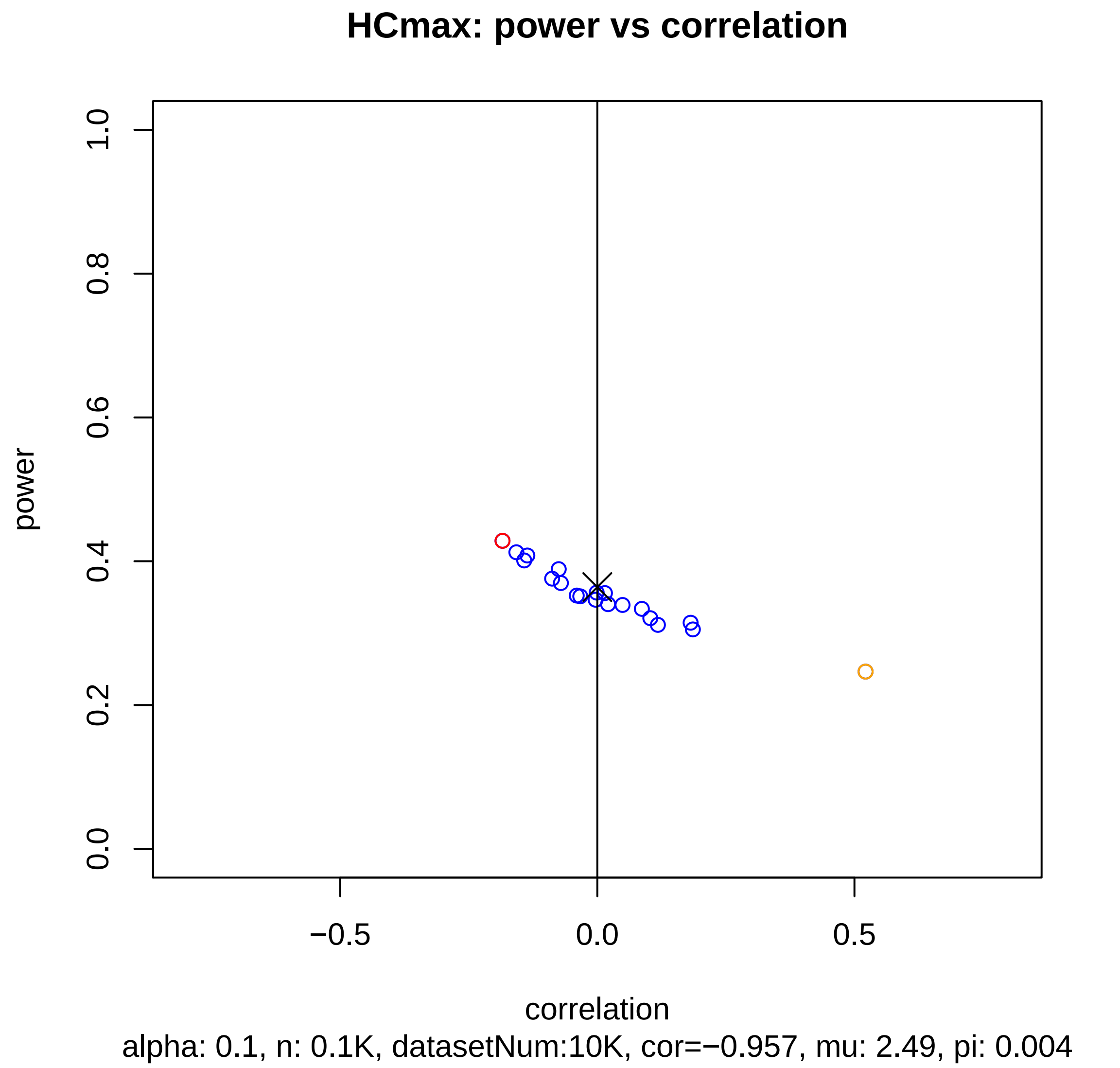
HCmax: power vs correlation
power
0.0
0.2
0.4
0.6
0.8
1.0
−0.5
0.0
0.5
correlation
alpha: 0.1, n: 0.1K, datasetNum:10K, cor=−0.957, mu: 2.49, pi: 0.004

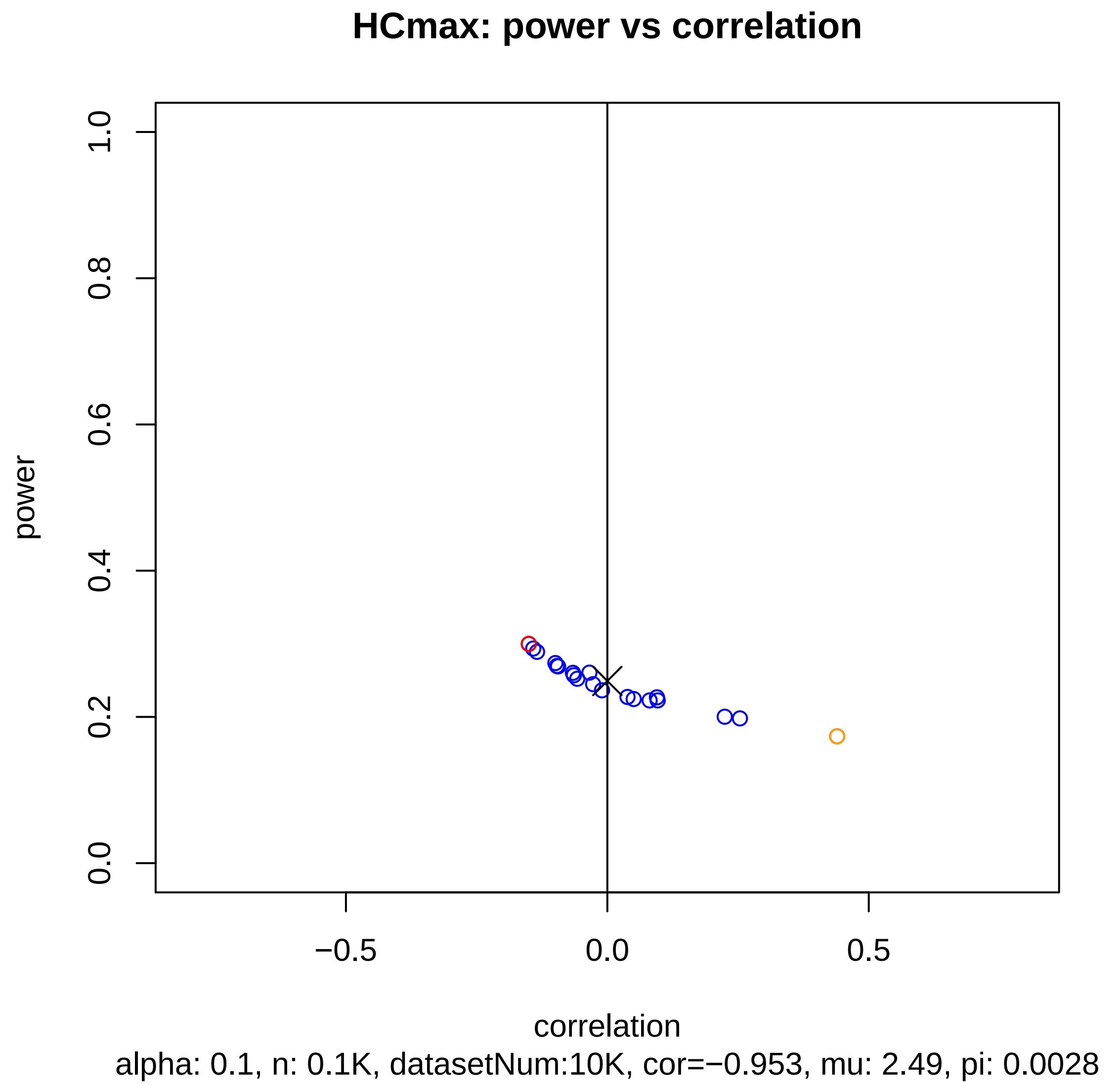
HCmax: power vs correlation
power
0.0
0.2
0.4
0.6
0.8
1.0
−0.5
0.0
0.5
correlation
alpha: 0.1, n: 0.1K, datasetNum:10K, cor=−0.953, mu: 2.49, pi: 0.0028

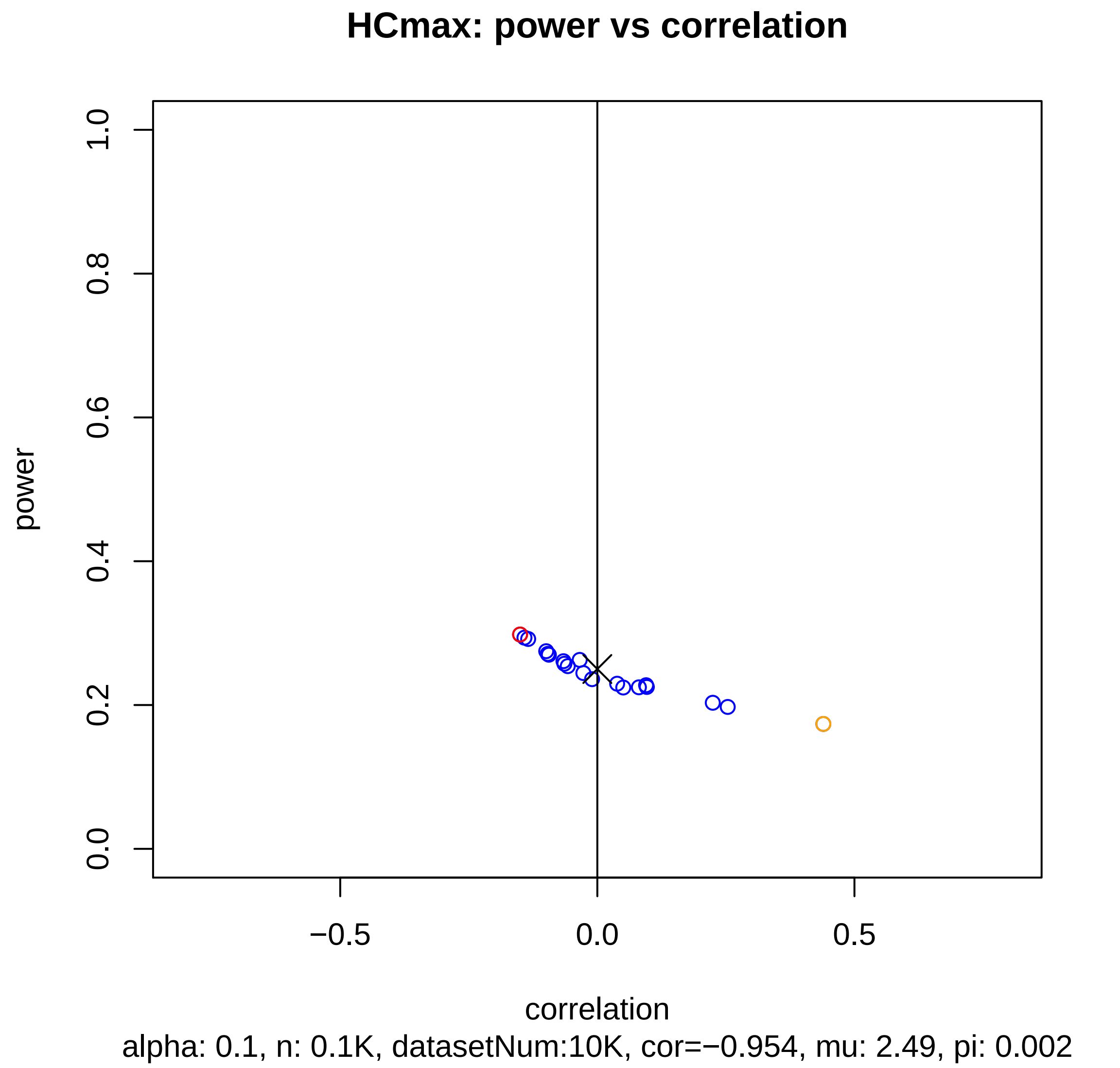

HCmax: power vs correlation
power
1.0
0.8
0.6
0.4
0.2
0.0
−0.5
0.0
0.5
correlation
alpha: 0.1, n: 0.1K, datasetNum:10K, cor=−0.954, mu: 2.49, pi: 0.002

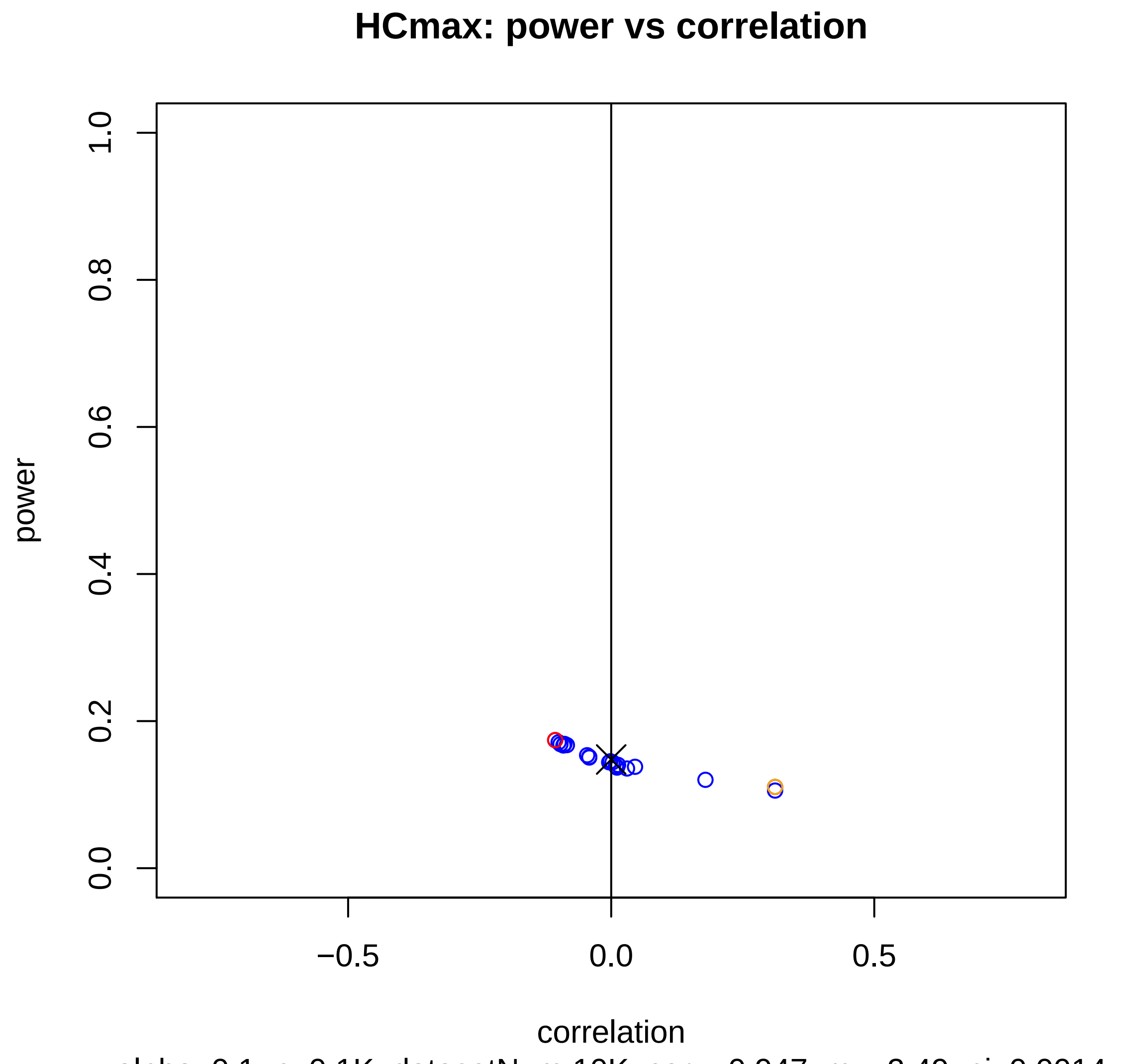

alpha: 0.1, n: 0.1K, datasetNum:10K, cor=−0.947, mu: 2.49, pi: 0.0014

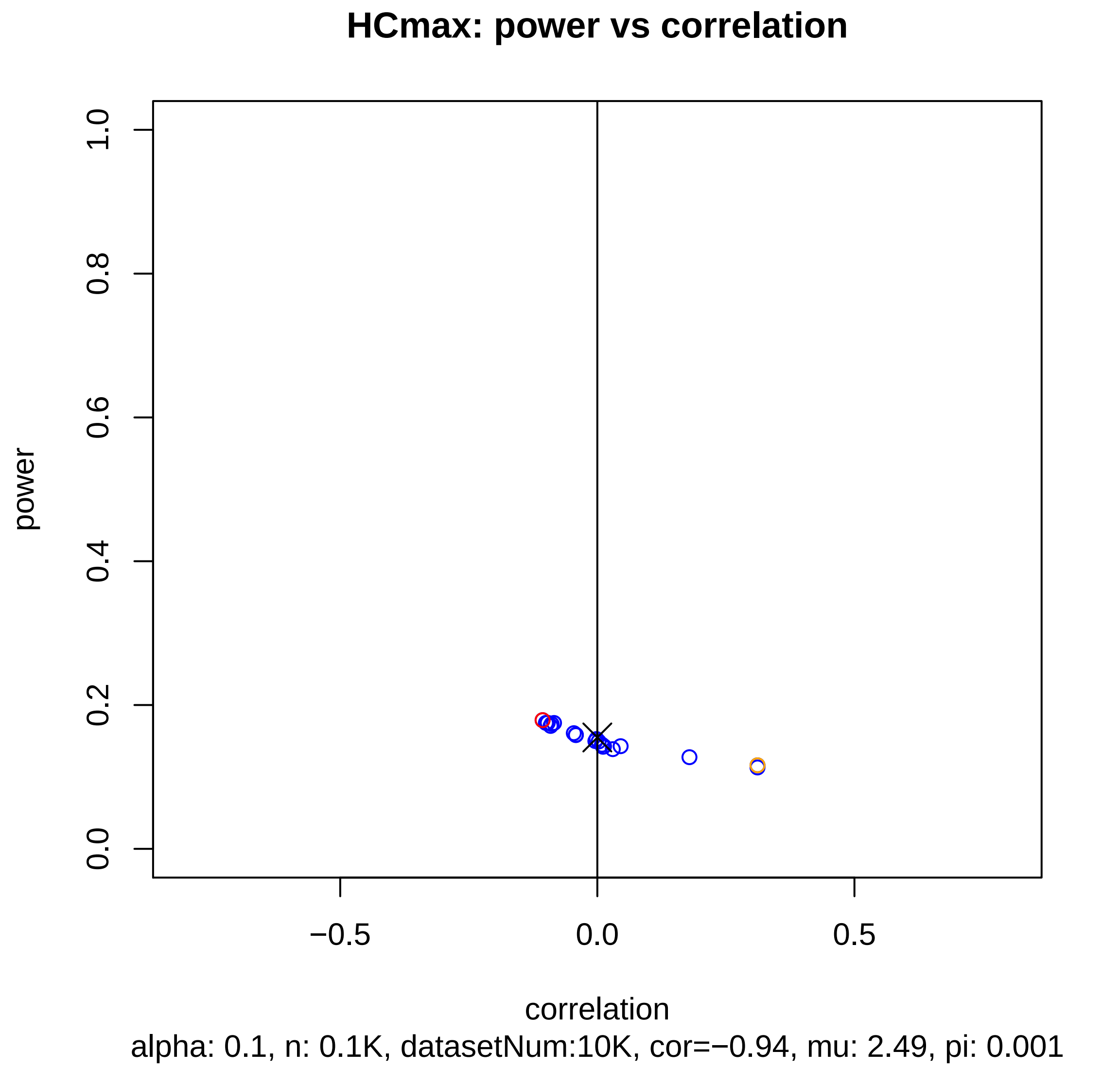
HCmax: power vs correlation
power
1.0
0.8
0.6
0.4
0.2
0.0
−0.5
0.0
0.5
correlation
alpha: 0.1, n: 0.1K, datasetNum:10K, cor=−0.94, mu: 2.49, pi: 0.001

**HCmax: power vs correlation**

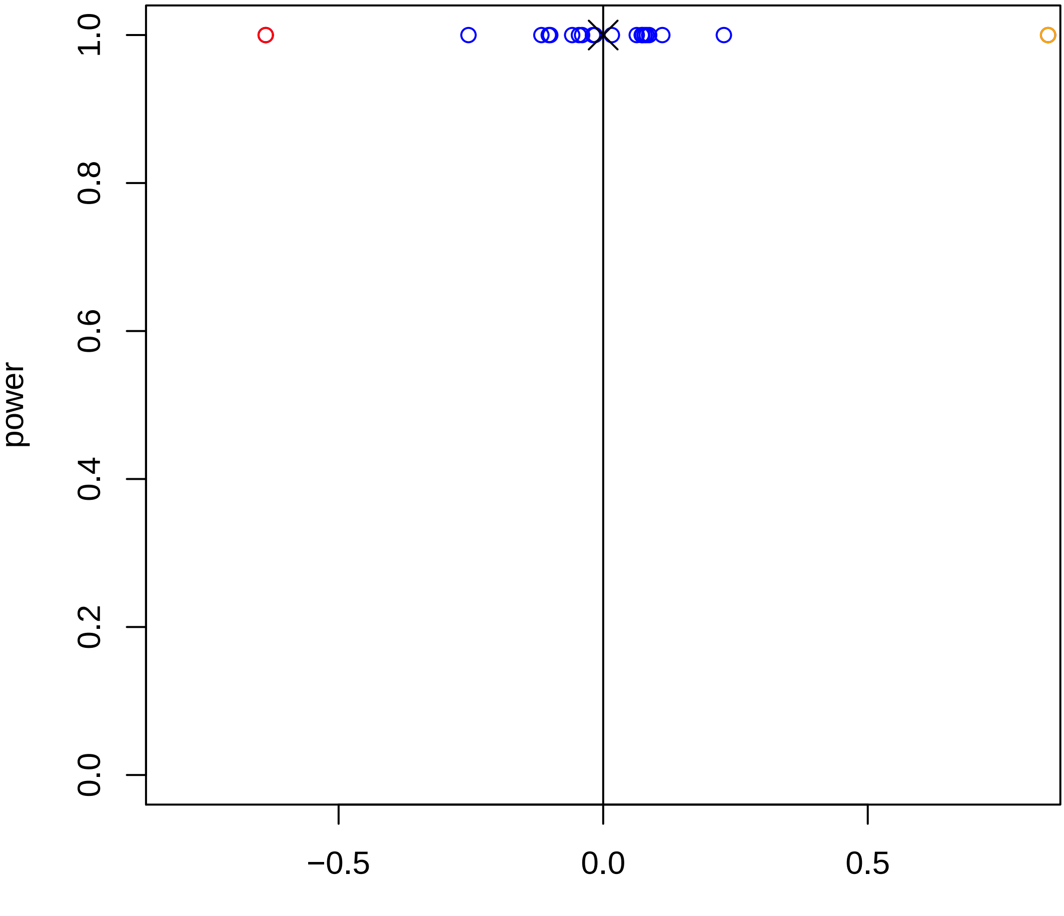


alpha: 0.1, n: 0.1K, datasetNum:10K, cor=NA, mu: 2.63, pi: 0.1778

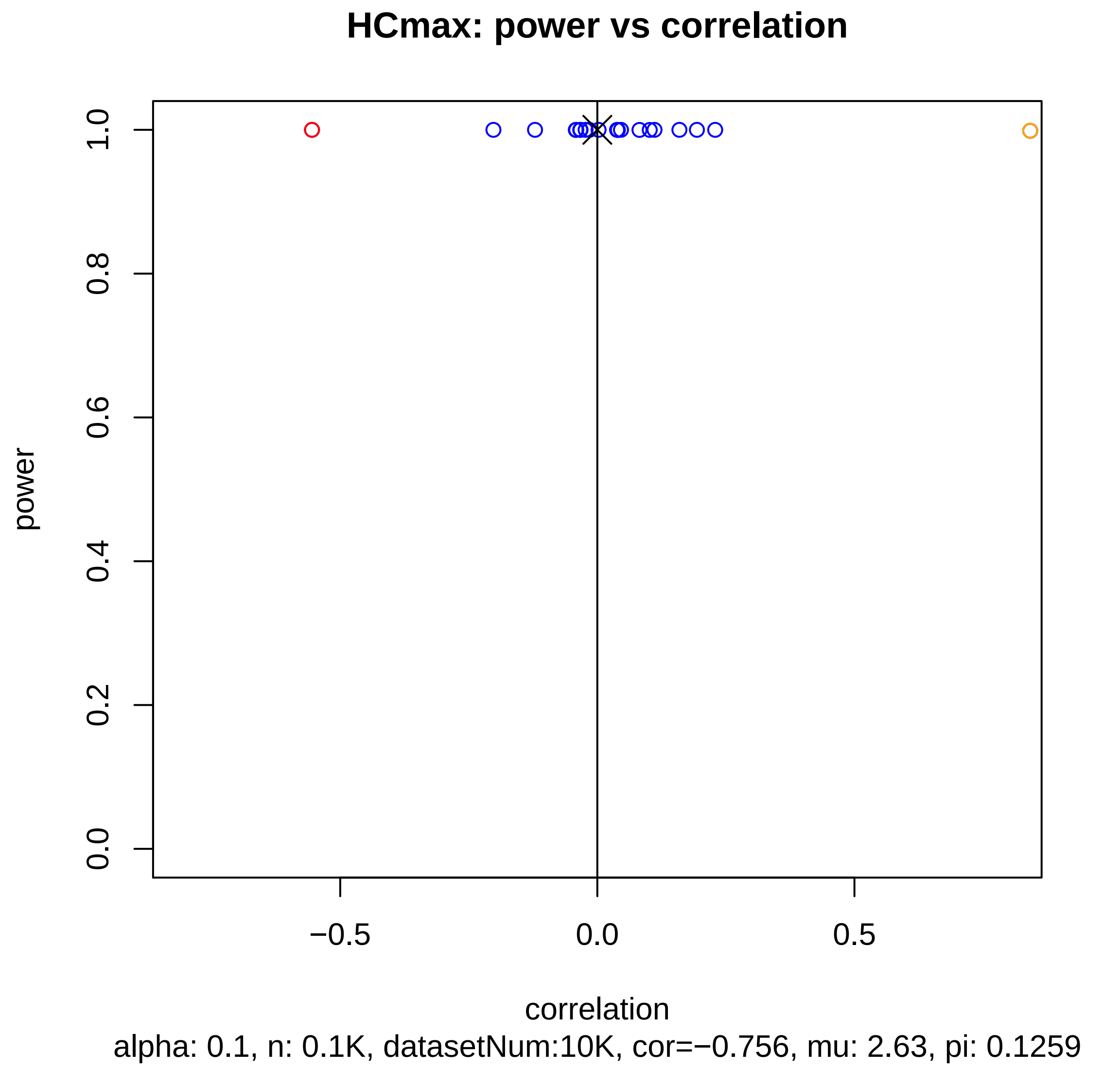
HCmax: power vs correlation
power
1.0
0.8
0.6
0.4
0.2
0.0
−0.5
0.0
0.5
correlation
alpha: 0.1, n: 0.1K, datasetNum:10K, cor=−0.756, mu: 2.63, pi: 0.1259

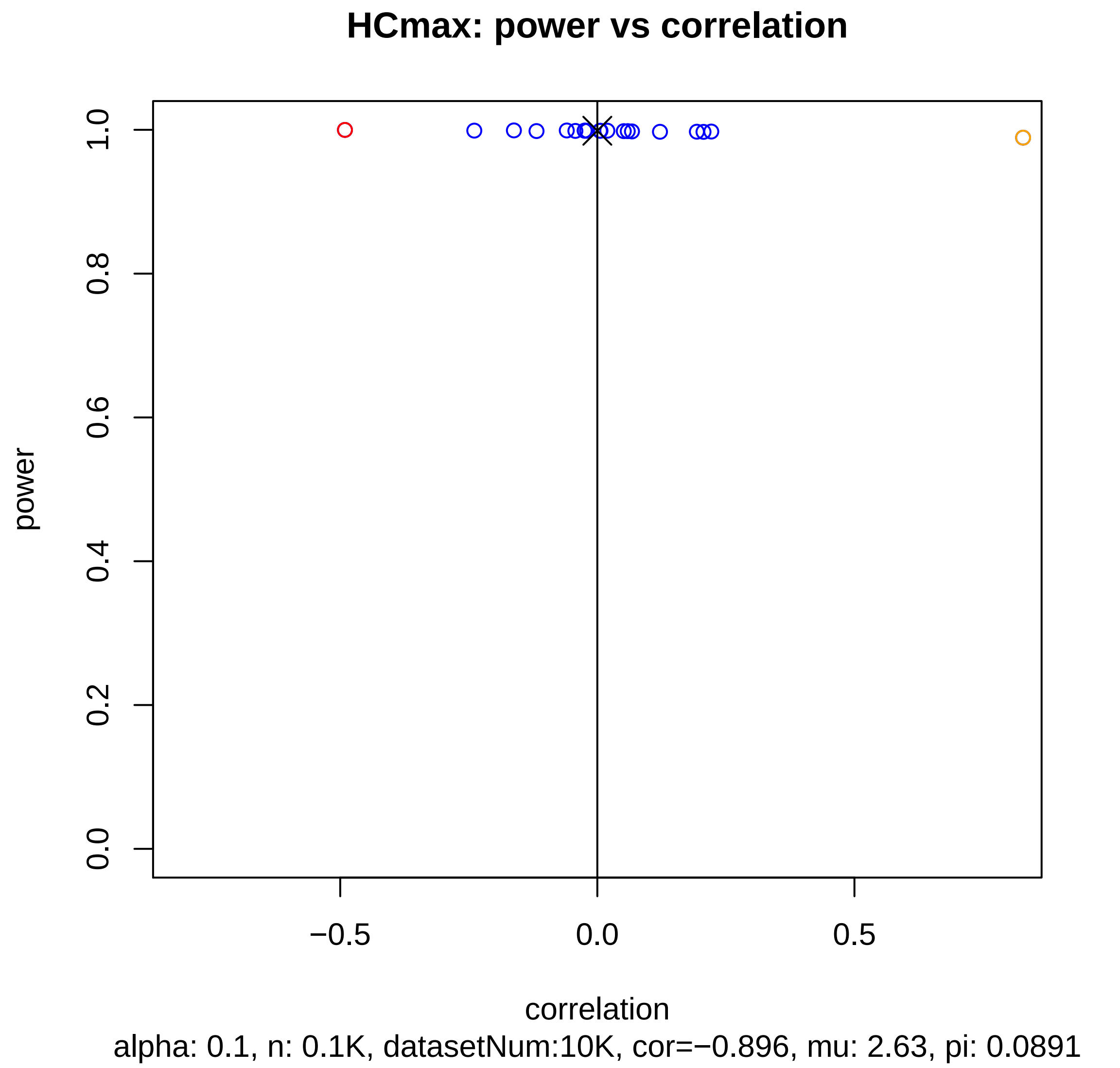
HCmax: power vs correlation
power
1.0
0.8
0.6
0.4
0.2
0.0
−0.5
0.0
0.5
correlation
alpha: 0.1, n: 0.1K, datasetNum:10K, cor=−0.896, mu: 2.63, pi: 0.0891

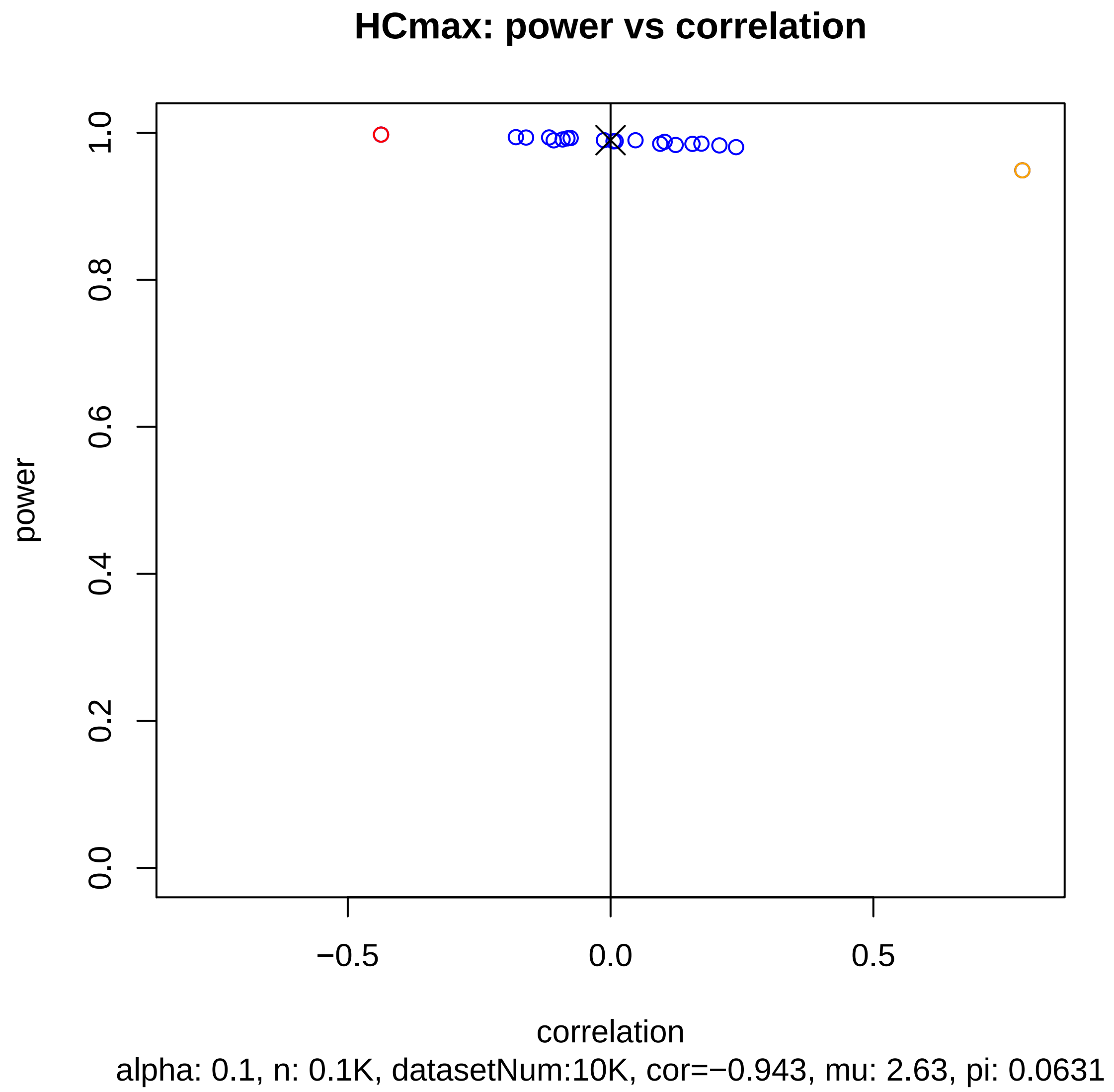
HCmax: power vs correlation
power
1.0
0.8
0.6
0.4
0.2
0.0
−0.5
0.0
0.5
correlation
alpha: 0.1, n: 0.1K, datasetNum:10K, cor=−0.943, mu: 2.63, pi: 0.0631

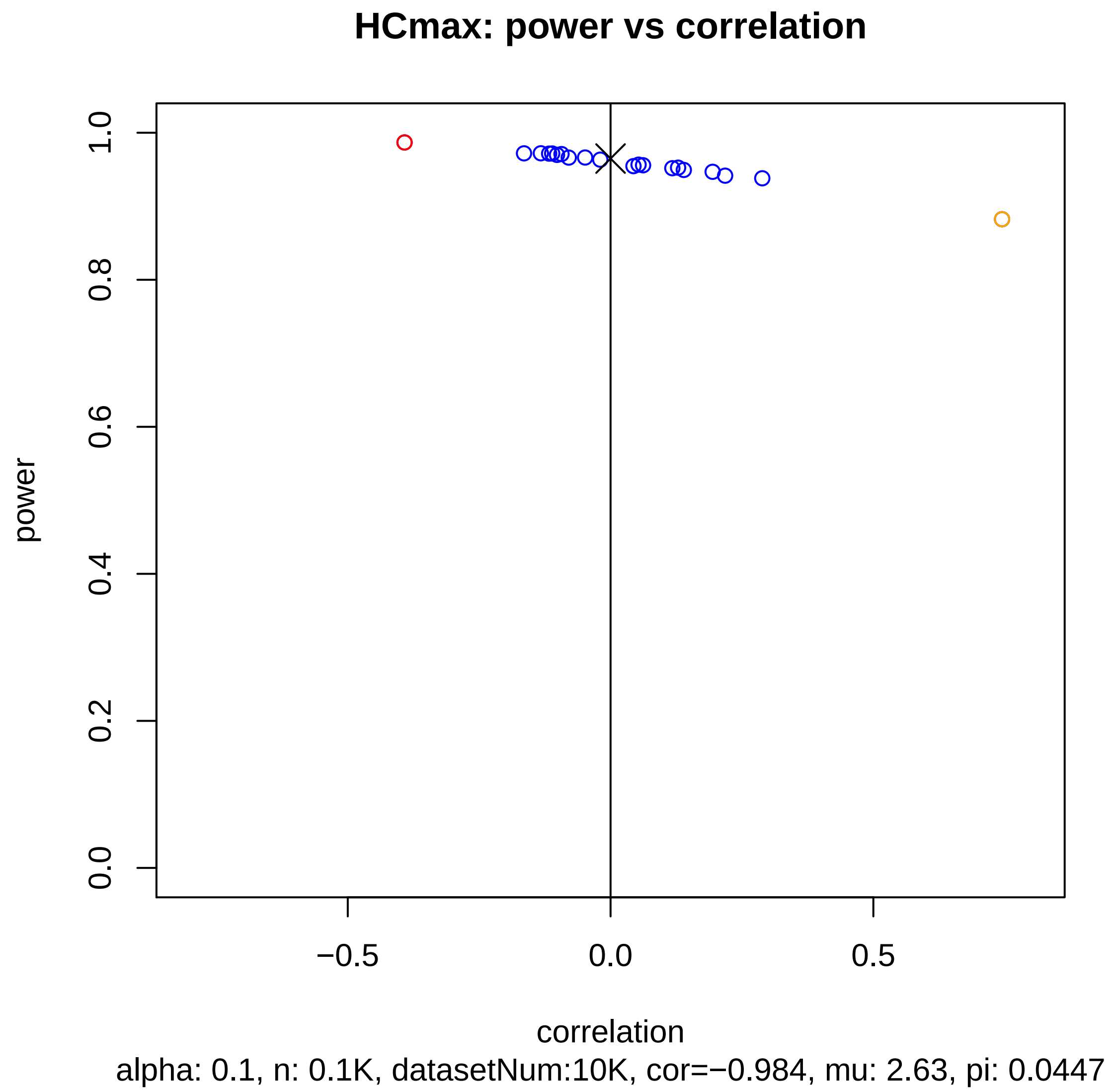

HCmax: power vs correlation
power
1.0
0.8
0.6
0.4
0.2
0.0
−0.5
0.0
0.5
correlation
alpha: 0.1, n: 0.1K, datasetNum:10K, cor=−0.984, mu: 2.63, pi: 0.0447

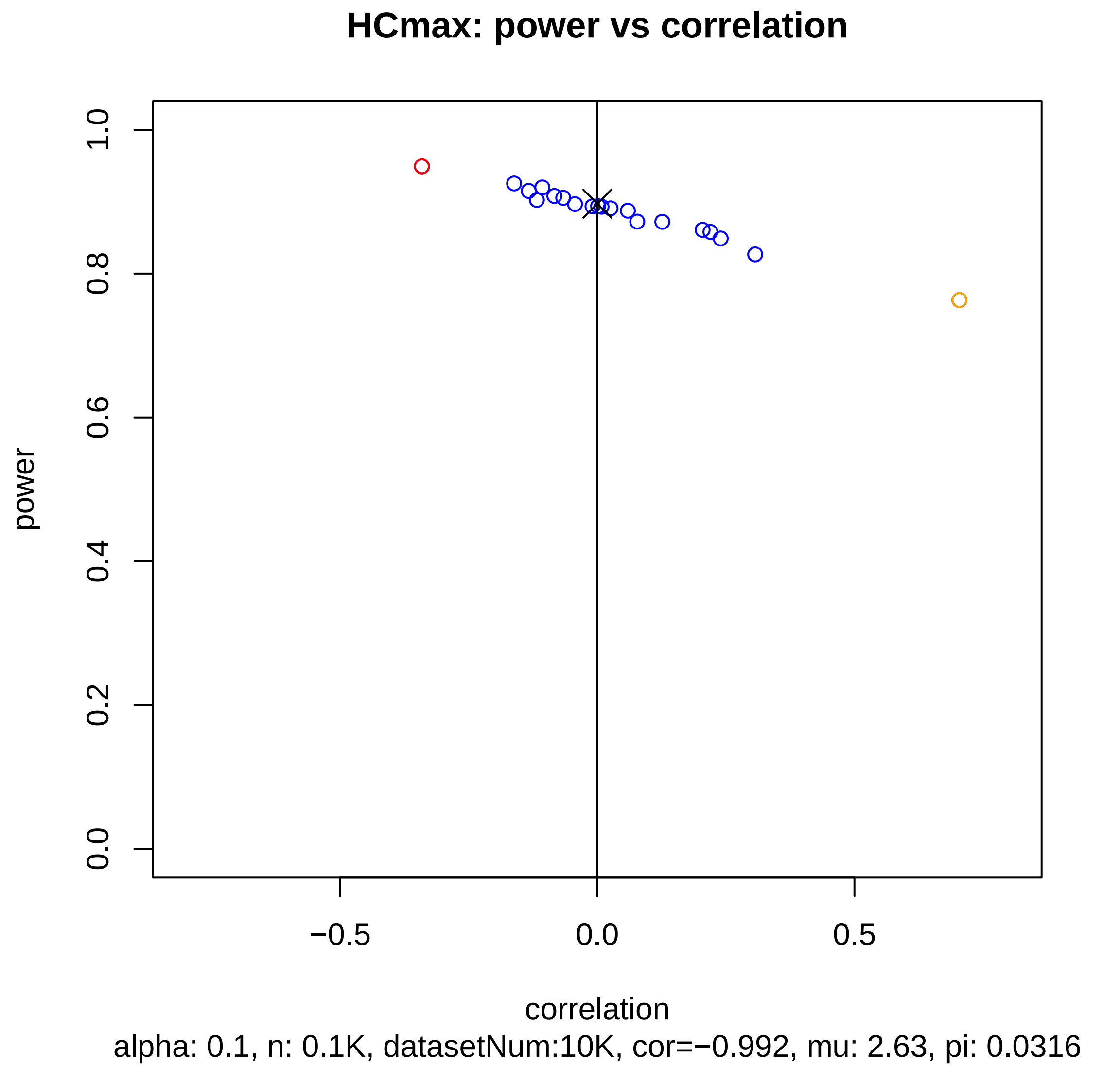
HCmax: power vs correlation
power
1.0
0.8
0.6
0.4
0.2
0.0
−0.5
0.0
0.5
correlation
alpha: 0.1, n: 0.1K, datasetNum:10K, cor=−0.992, mu: 2.63, pi: 0.0316

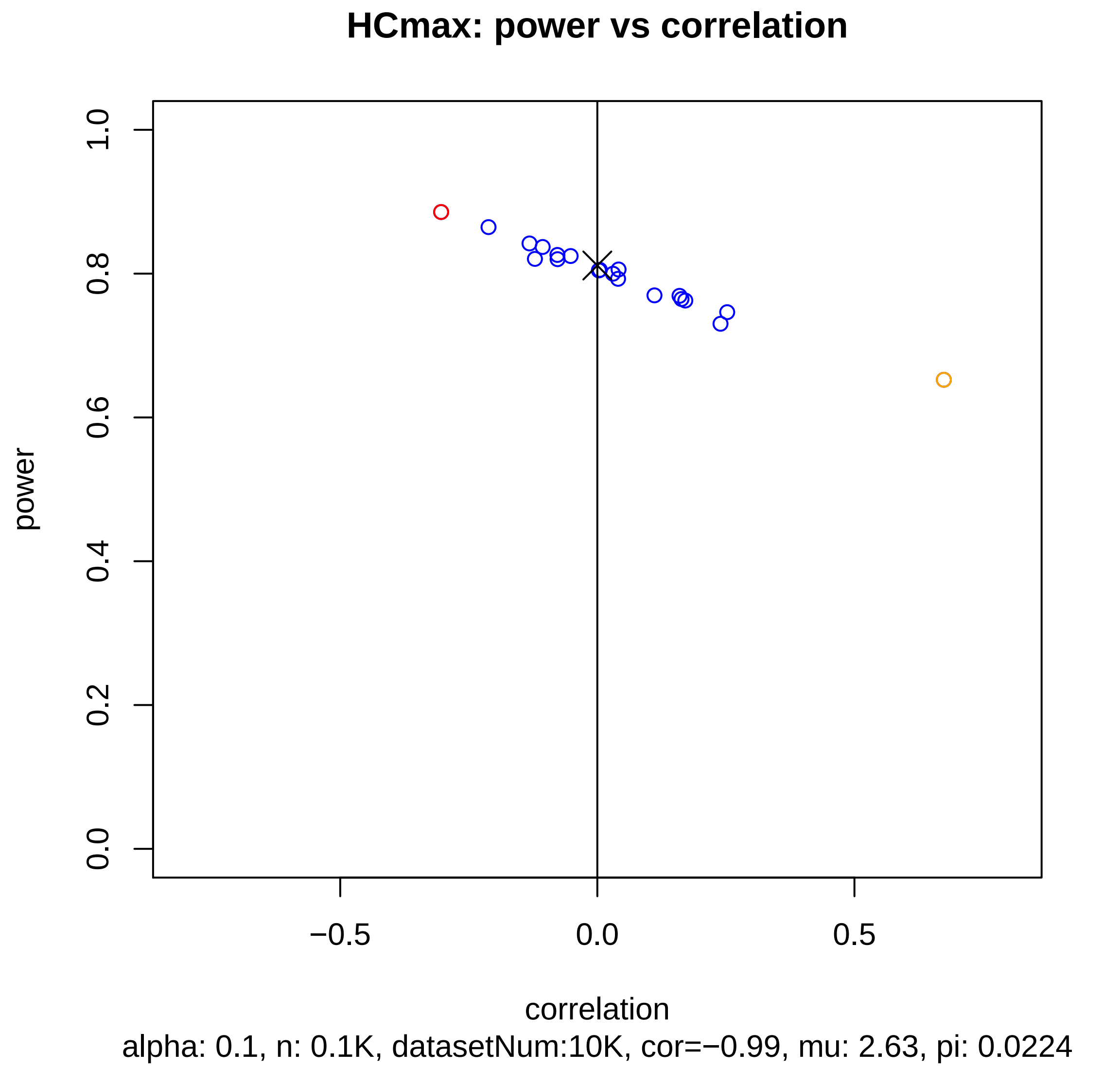

HCmax: power vs correlation
power
0.0
0.2
0.4
0.6
0.8
1.0
−0.5
0.0
0.5
correlation
alpha: 0.1, n: 0.1K, datasetNum:10K, cor=−0.99, mu: 2.63, pi: 0.0224

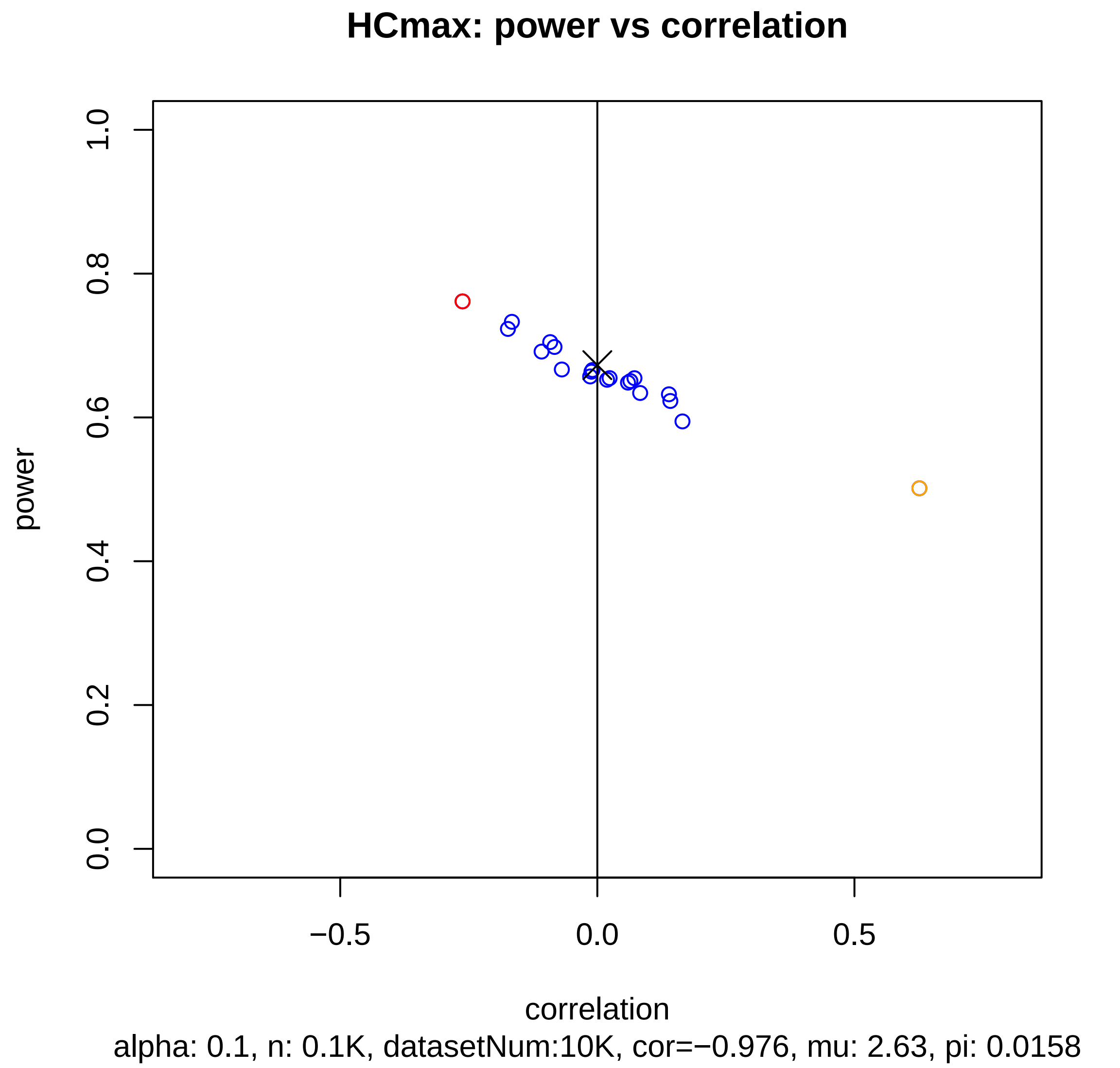
HCmax: power vs correlation
power
0.0
0.2
0.4
0.6
0.8
1.0
−0.5
0.0
0.5
correlation
alpha: 0.1, n: 0.1K, datasetNum:10K, cor=−0.976, mu: 2.63, pi: 0.0158

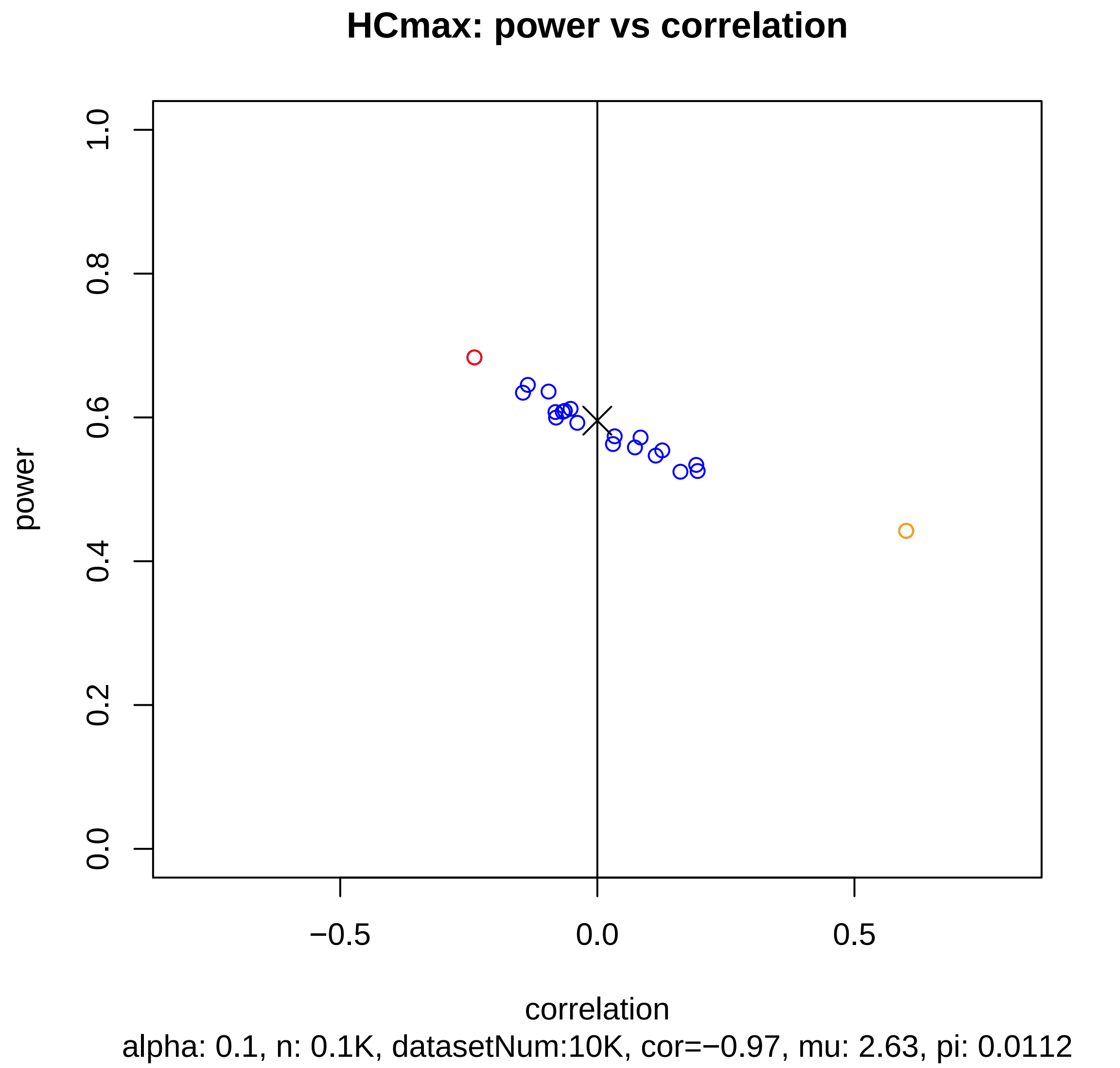

HCmax: power vs correlation
power
0.0
0.2
0.4
0.6
0.8
1.0
−0.5
0.0
0.5
correlation
alpha: 0.1, n: 0.1K, datasetNum:10K, cor=−0.97, mu: 2.63, pi: 0.0112

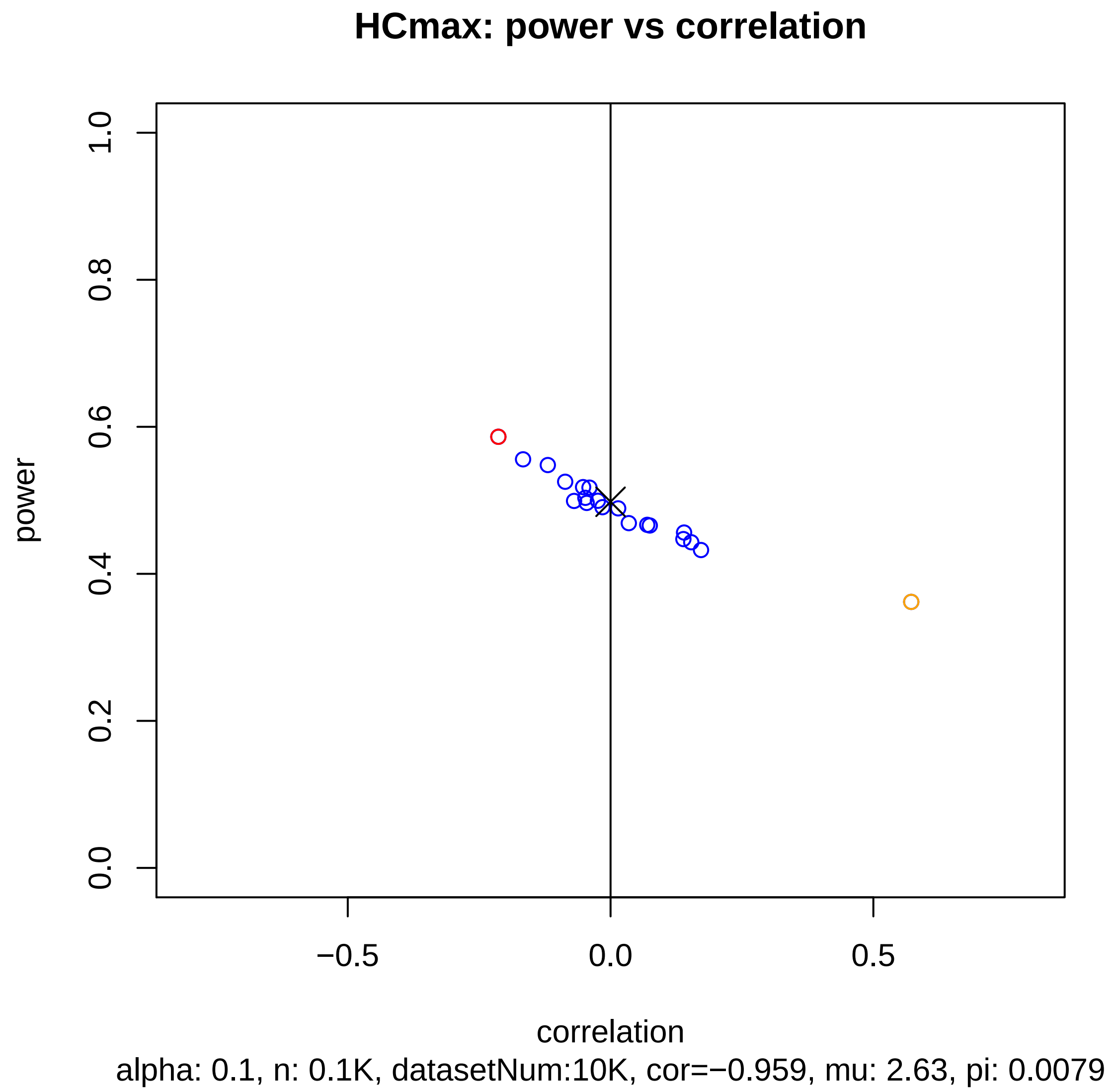
HCmax: power vs correlation
power
correlation
1.0
0.8
0.6
0.4
0.2
0.0
−0.5
0.0
0.5
alpha: 0.1, n: 0.1K, datasetNum:10K, cor=−0.959, mu: 2.63, pi: 0.0079

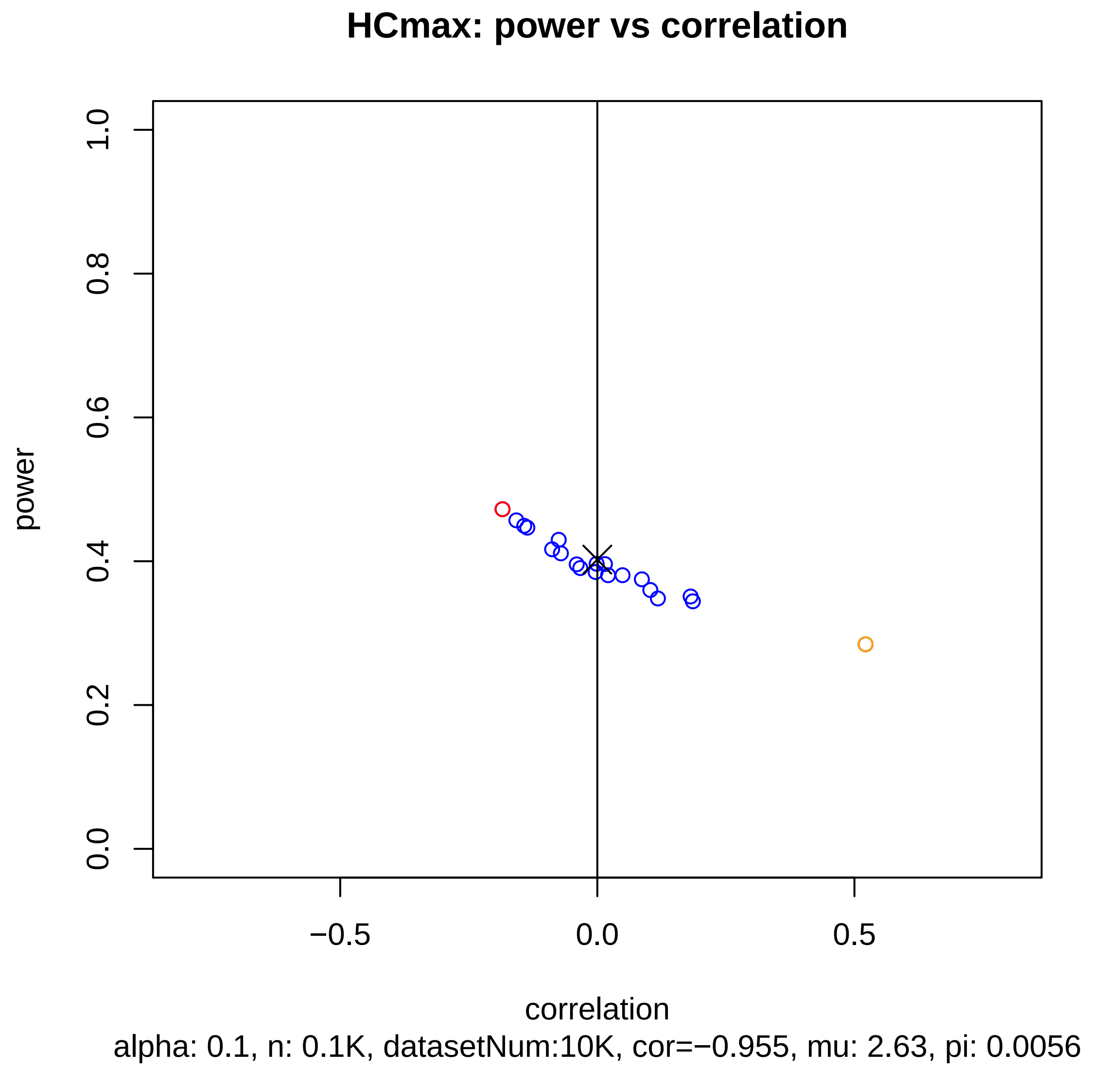
HCmax: power vs correlation
power
1.0
0.8
0.6
0.4
0.2
0.0
−0.5
0.0
0.5
correlation
alpha: 0.1, n: 0.1K, datasetNum:10K, cor=−0.955, mu: 2.63, pi: 0.0056

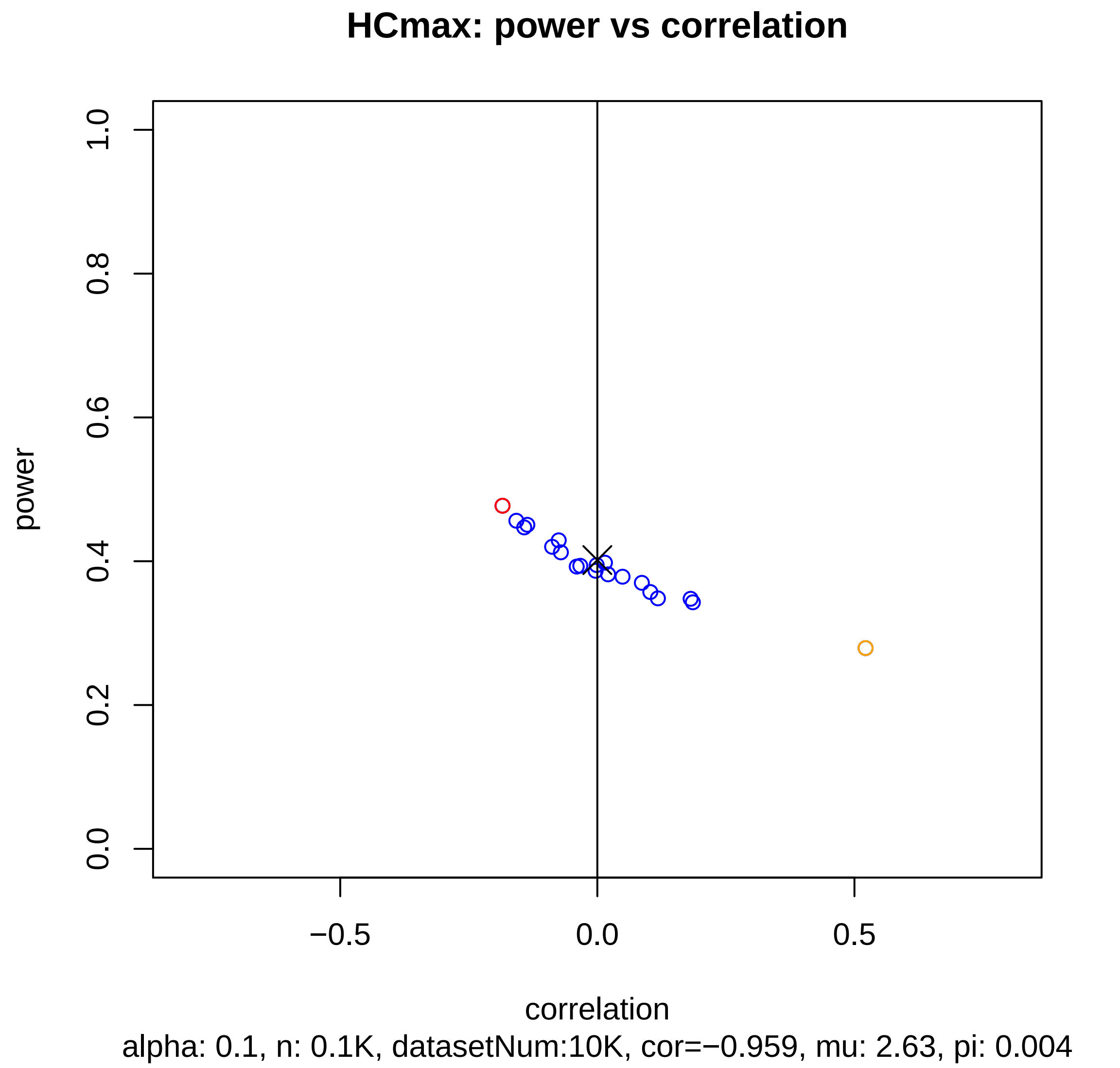
HCmax: power vs correlation
power
1.0
0.8
0.6
0.4
0.2
0.0
−0.5
0.0
0.5
correlation
alpha: 0.1, n: 0.1K, datasetNum:10K, cor=−0.959, mu: 2.63, pi: 0.004

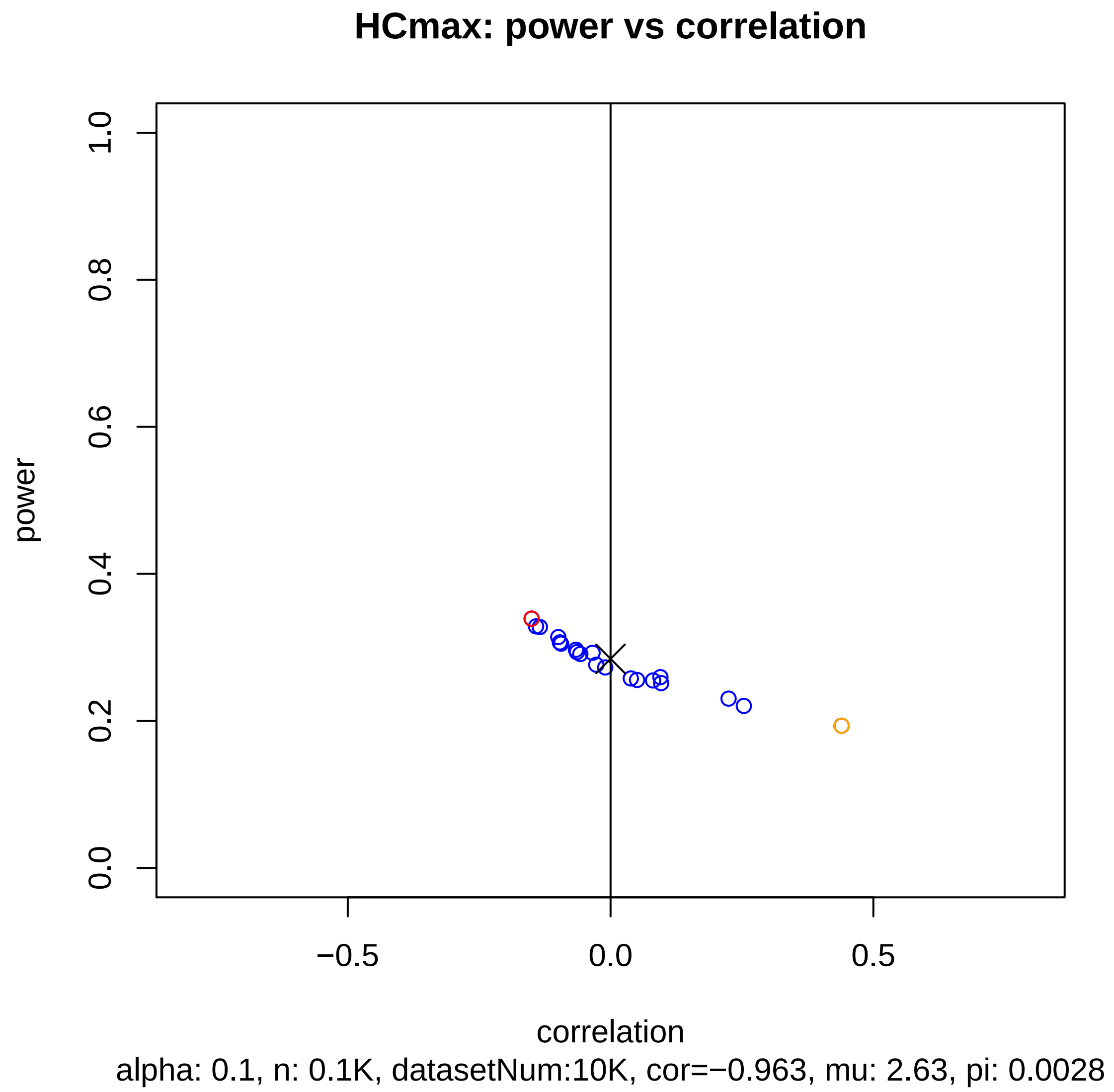

HCmax: power vs correlation
power
0.0
0.2
0.4
0.6
0.8
1.0
−0.5
0.0
0.5
correlation
alpha: 0.1, n: 0.1K, datasetNum:10K, cor=−0.963, mu: 2.63, pi: 0.0028

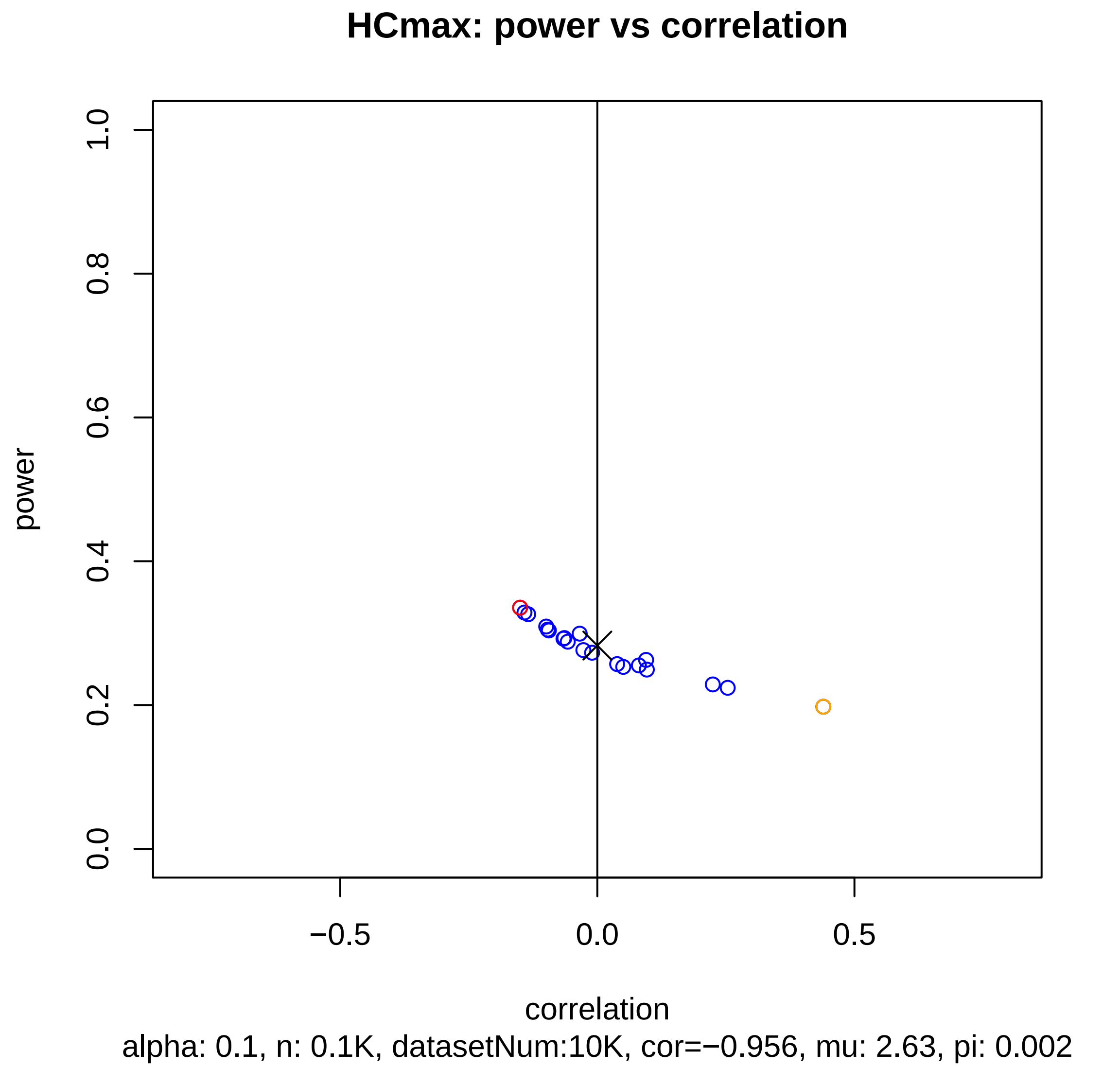

HCmax: power vs correlation
power
0.0
0.2
0.4
0.6
0.8
1.0
−0.5
0.0
0.5
correlation
alpha: 0.1, n: 0.1K, datasetNum:10K, cor=−0.956, mu: 2.63, pi: 0.002

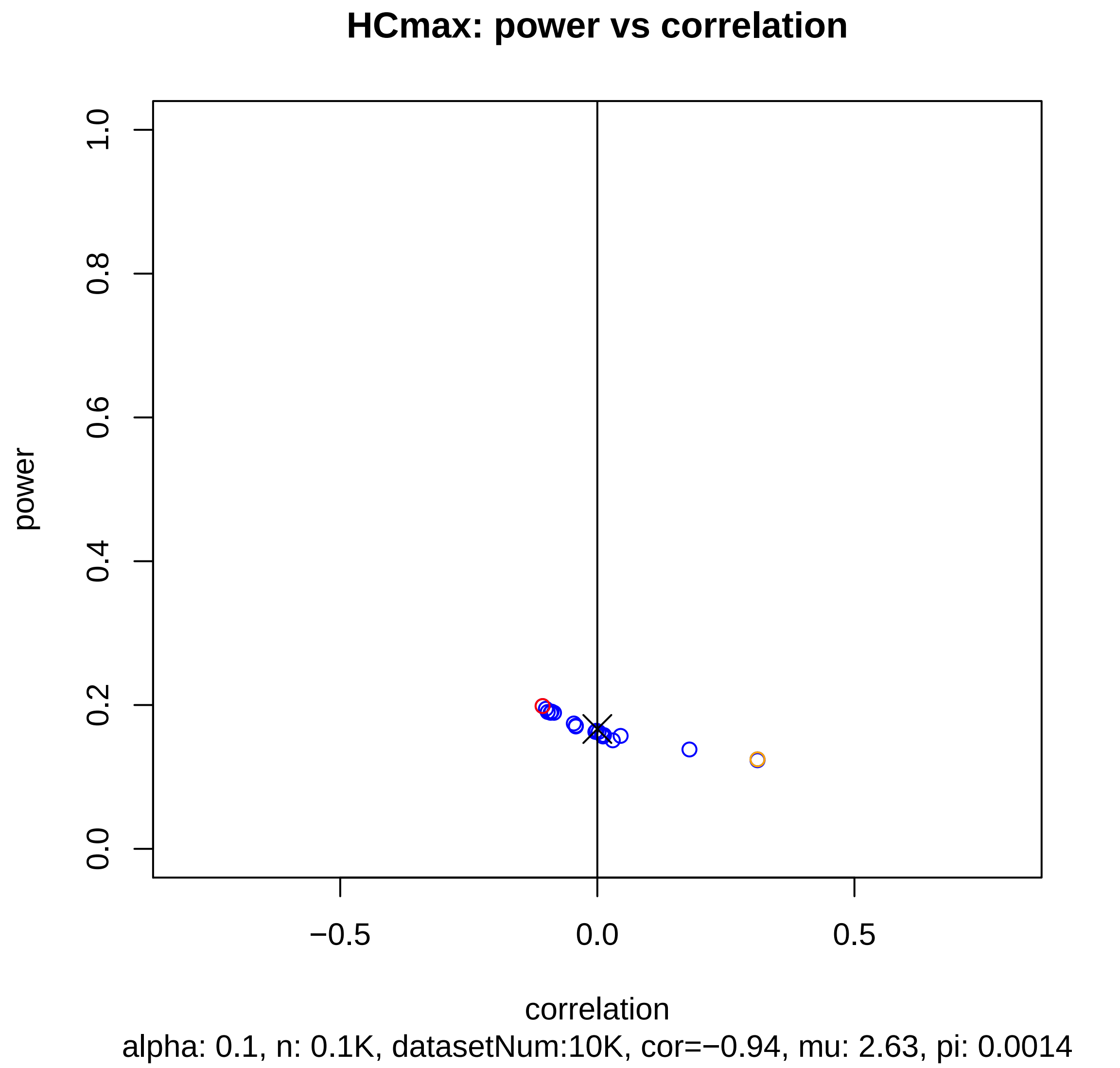
HCmax: power vs correlation
power
0.0
0.2
0.4
0.6
0.8
1.0
−0.5
0.0
0.5
correlation
alpha: 0.1, n: 0.1K, datasetNum:10K, cor=−0.94, mu: 2.63, pi: 0.0014

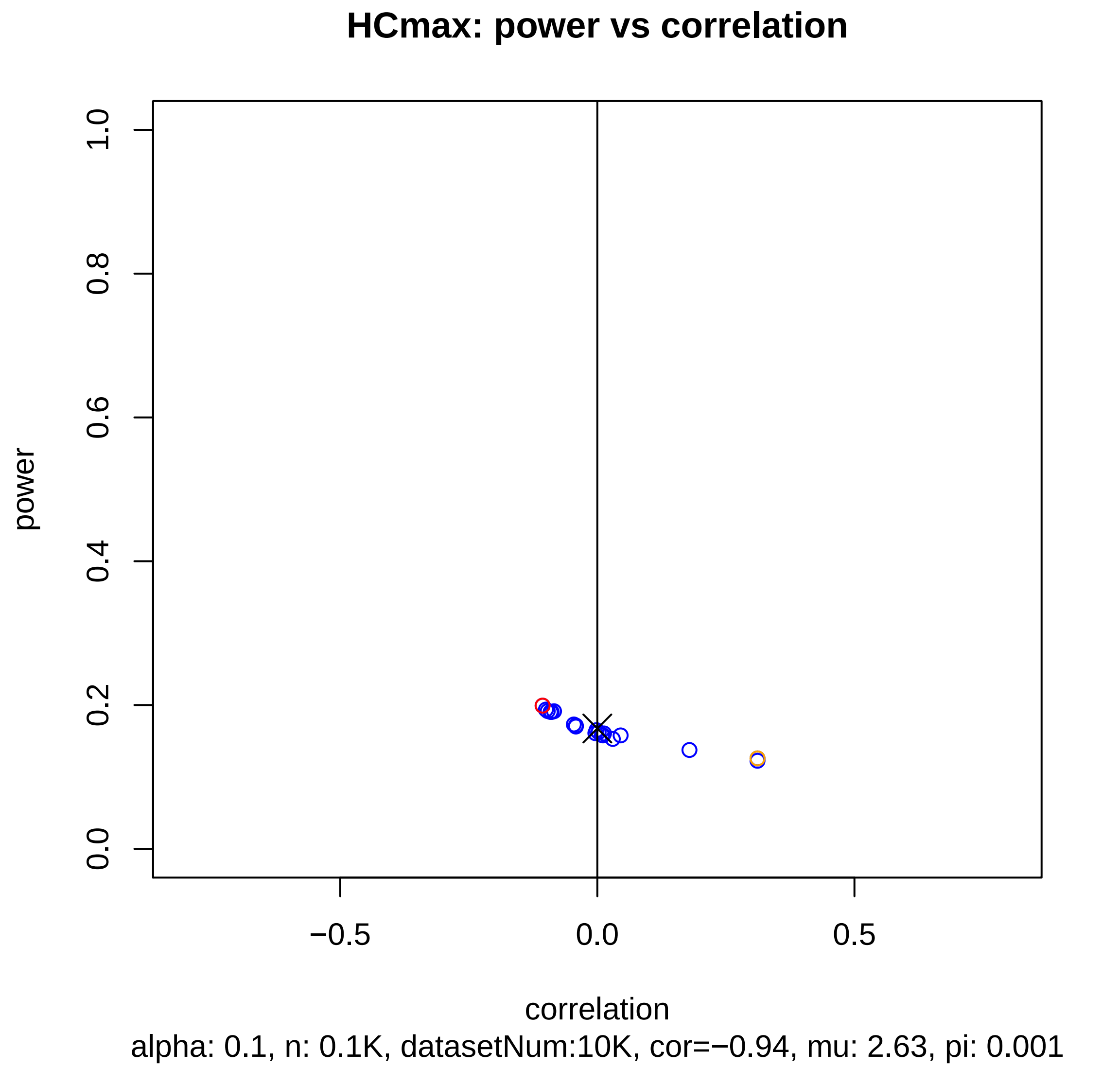

HCmax: power vs correlation
power
0.0
0.2
0.4
0.6
0.8
1.0
−0.5
0.0
0.5
correlation
alpha: 0.1, n: 0.1K, datasetNum:10K, cor=−0.94, mu: 2.63, pi: 0.001

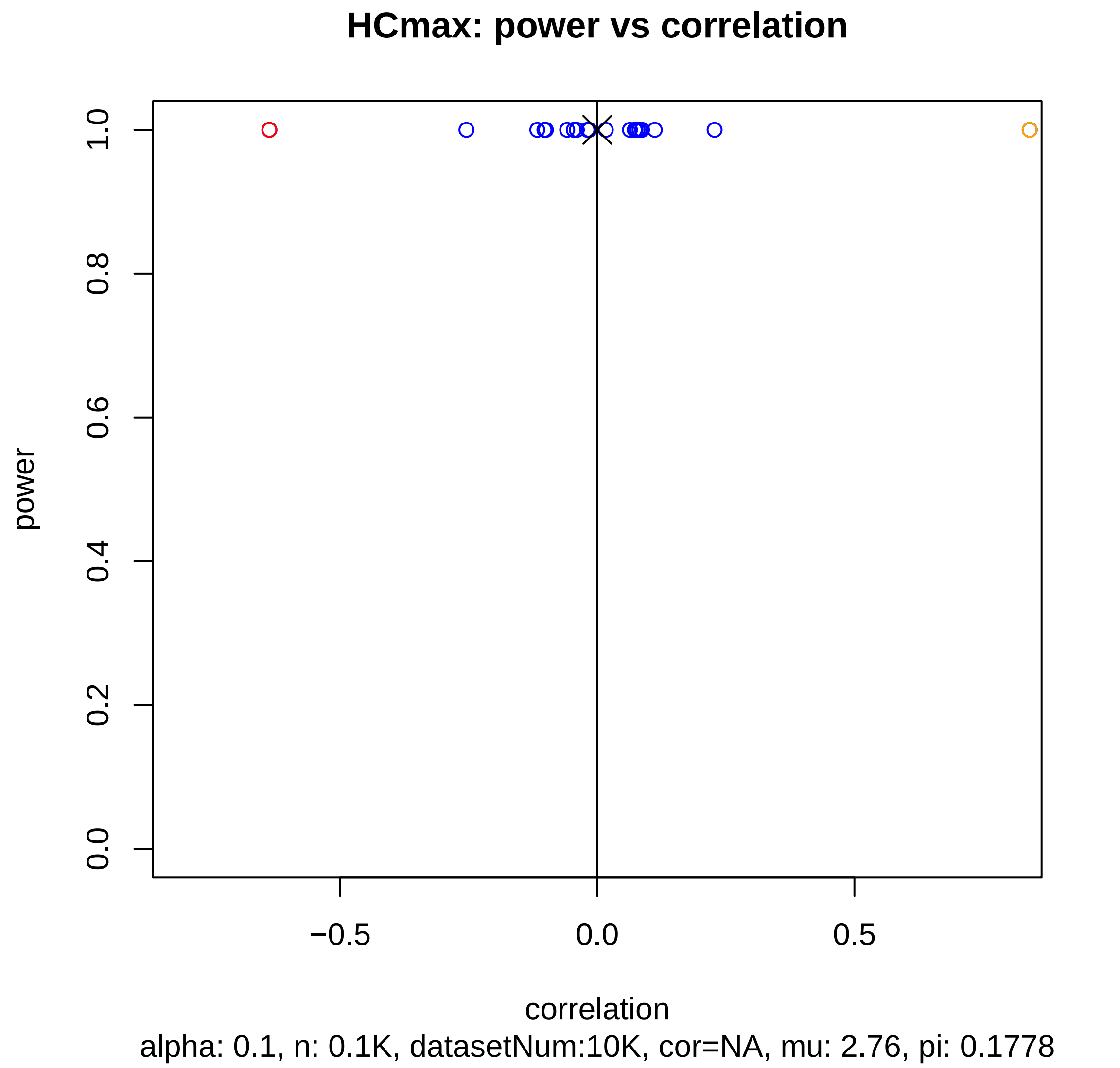
HCmax: power vs correlation
power
1.0
0.8
0.6
0.4
0.2
0.0
−0.5
0.0
0.5
correlation
alpha: 0.1, n: 0.1K, datasetNum:10K, cor=NA, mu: 2.76, pi: 0.1778

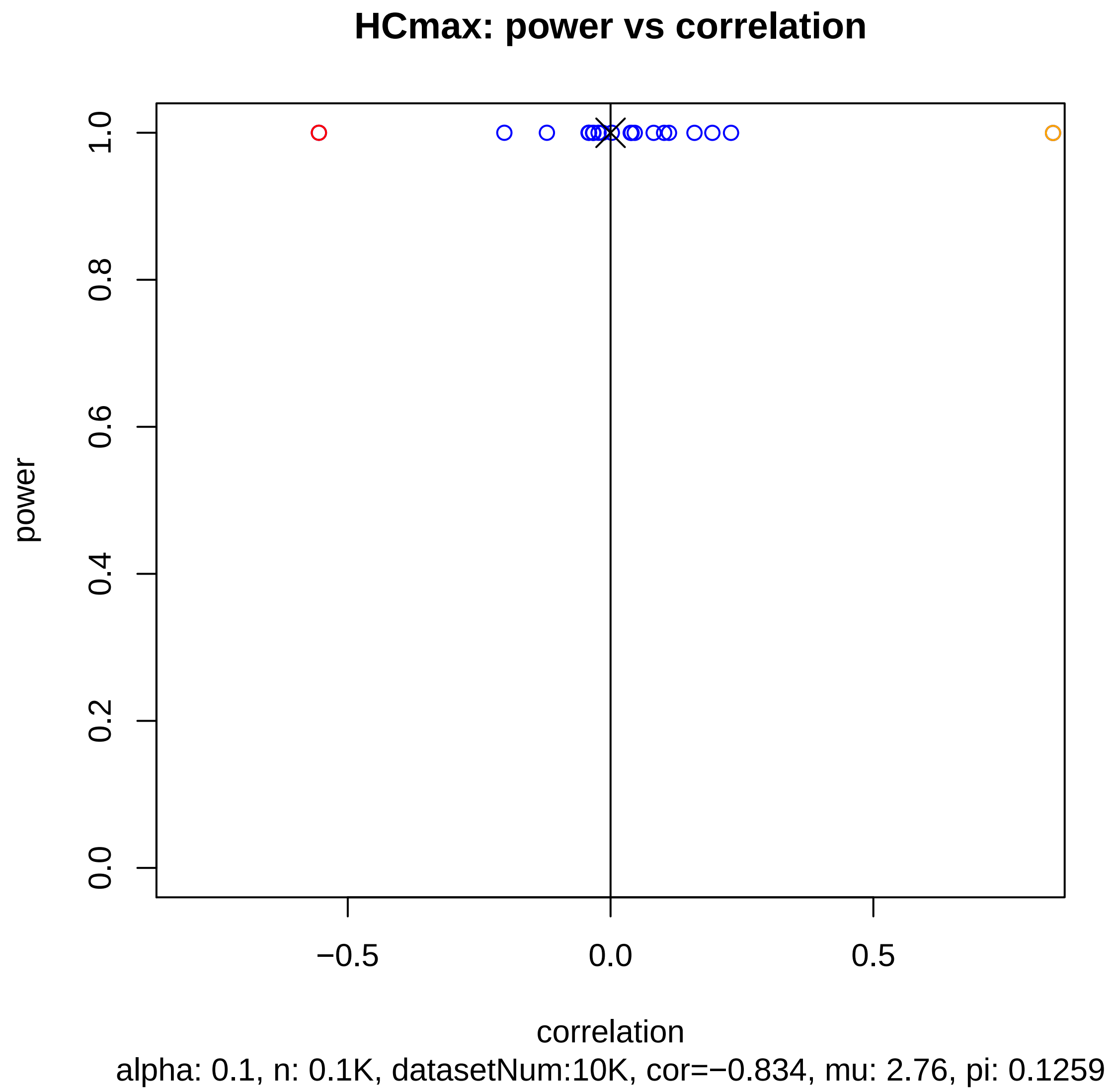
HCmax: power vs correlation
power
1.0
0.8
0.6
0.4
0.2
0.0
−0.5
0.0
0.5
correlation
alpha: 0.1, n: 0.1K, datasetNum:10K, cor=−0.834, mu: 2.76, pi: 0.1259

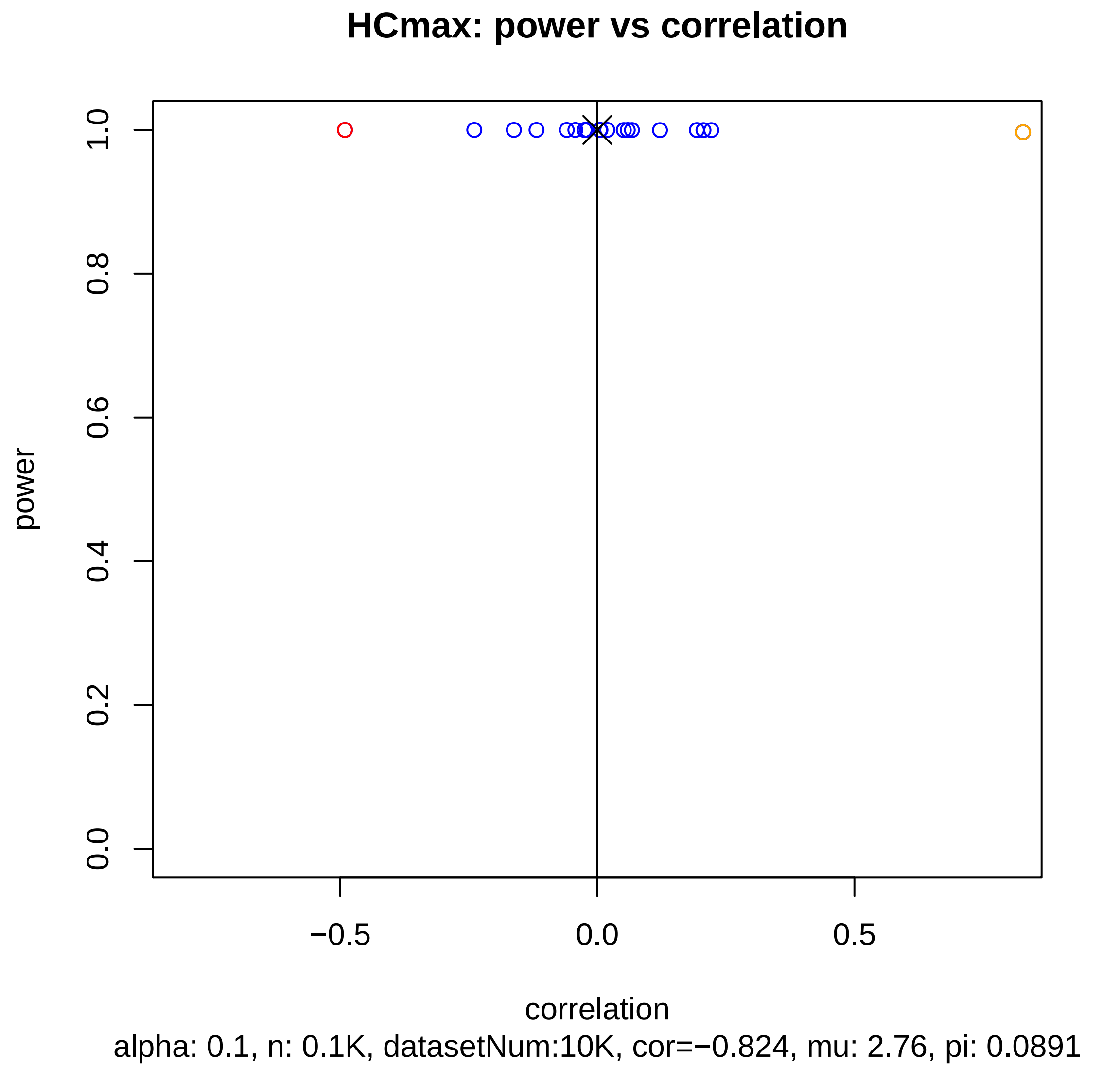

HCmax: power vs correlation
power
1.0
0.8
0.6
0.4
0.2
0.0
−0.5
0.0
0.5
correlation
alpha: 0.1, n: 0.1K, datasetNum:10K, cor=−0.824, mu: 2.76, pi: 0.0891

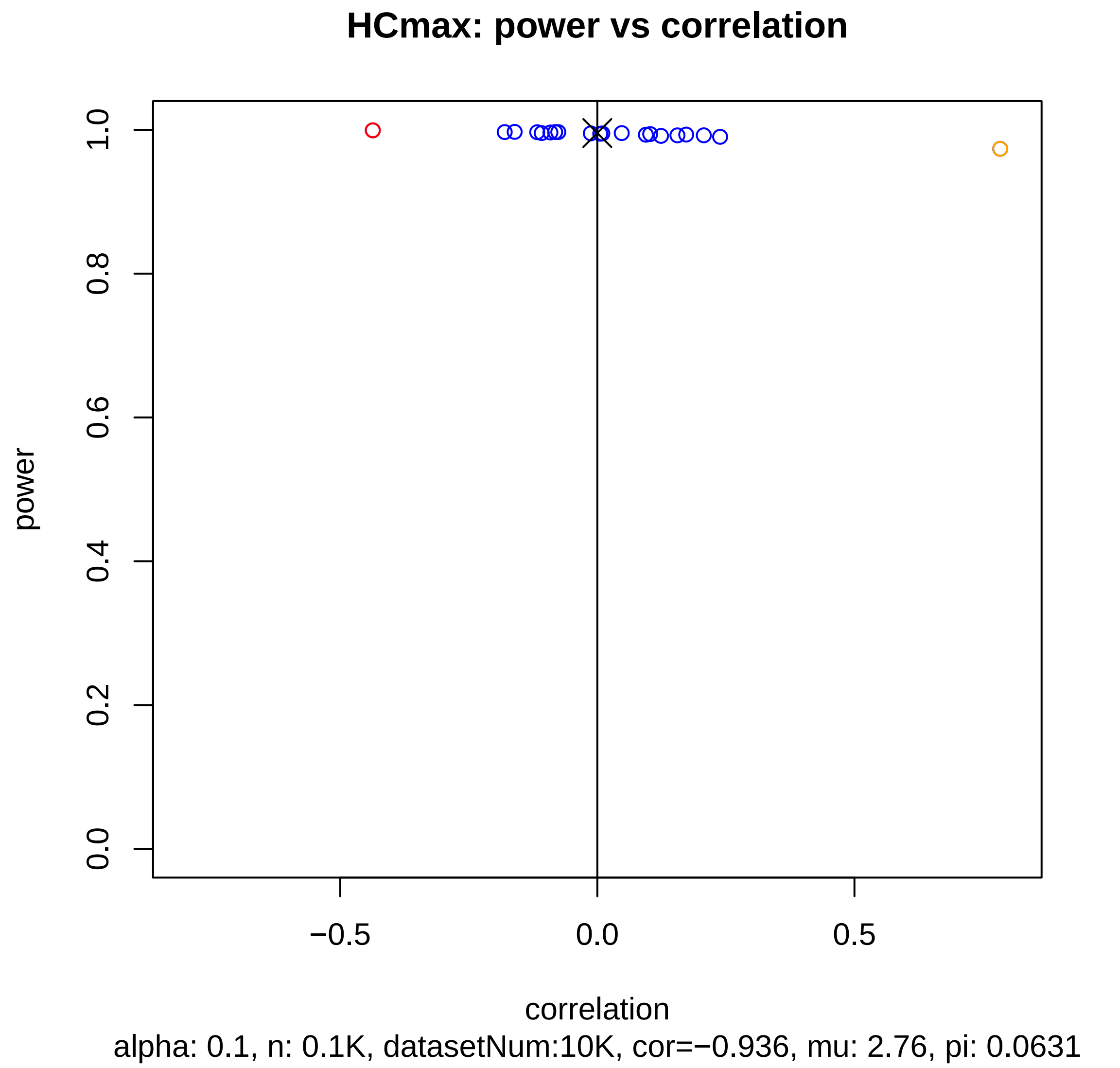

HCmax: power vs correlation
power
1.0
0.8
0.6
0.4
0.2
0.0
−0.5
0.0
0.5
correlation
alpha: 0.1, n: 0.1K, datasetNum:10K, cor=−0.936, mu: 2.76, pi: 0.0631

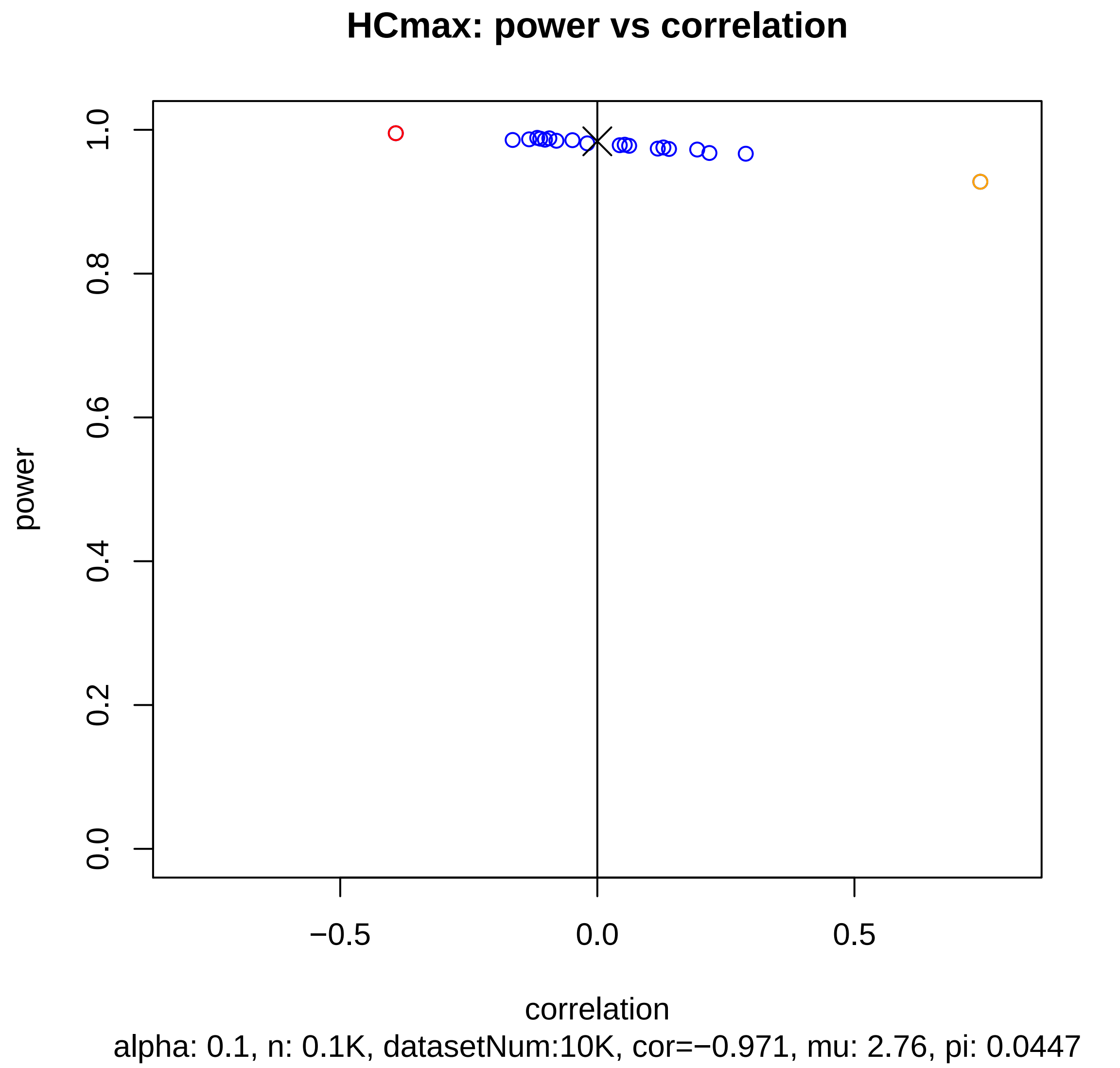
HCmax: power vs correlation
power
0.0
0.2
0.4
0.6
0.8
1.0
−0.5
0.0
0.5
correlation
alpha: 0.1, n: 0.1K, datasetNum:10K, cor=−0.971, mu: 2.76, pi: 0.0447

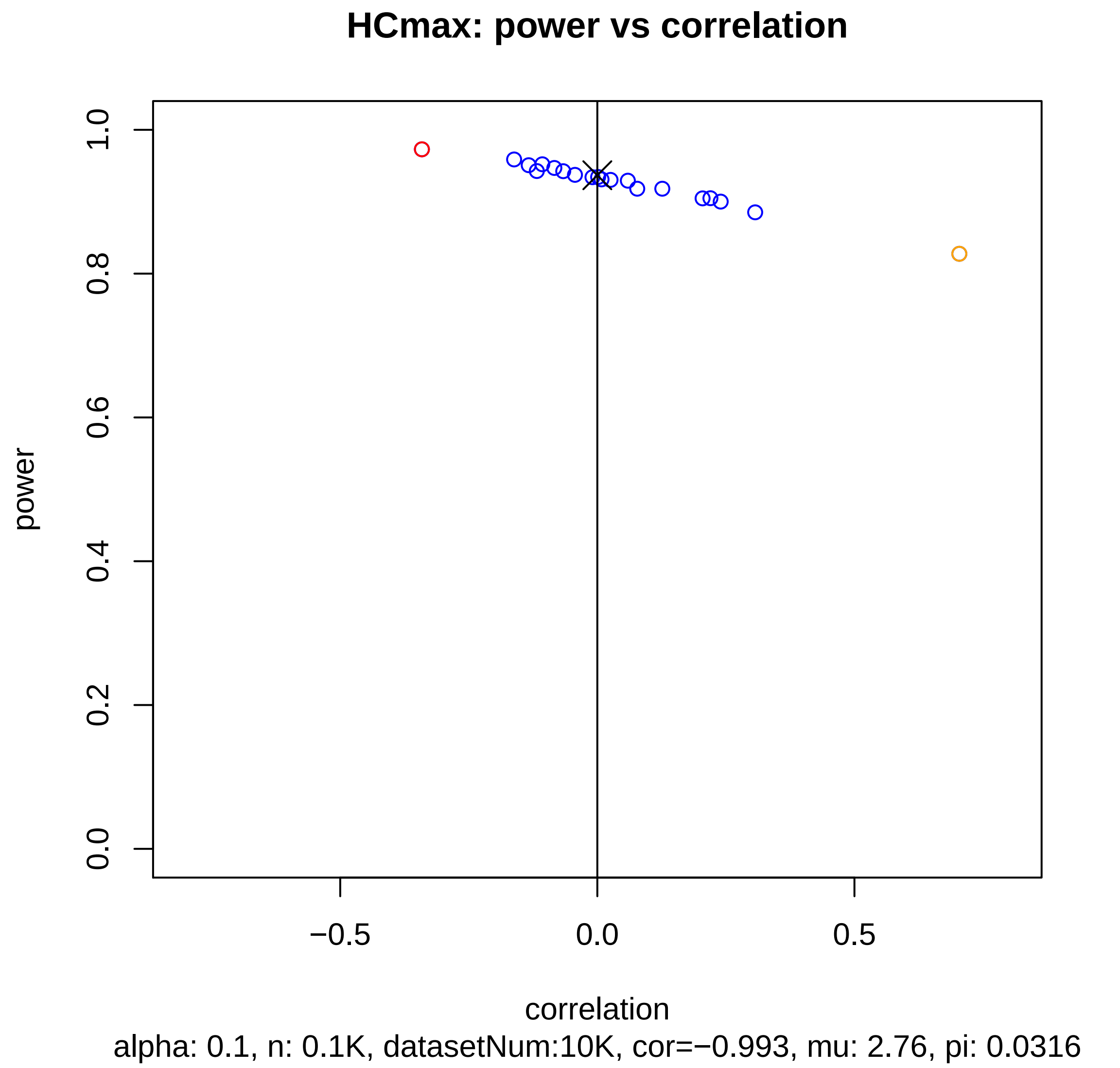

HCmax: power vs correlation
power
1.0
0.8
0.6
0.4
0.2
0.0
−0.5
0.0
0.5
correlation
alpha: 0.1, n: 0.1K, datasetNum:10K, cor=−0.993, mu: 2.76, pi: 0.0316

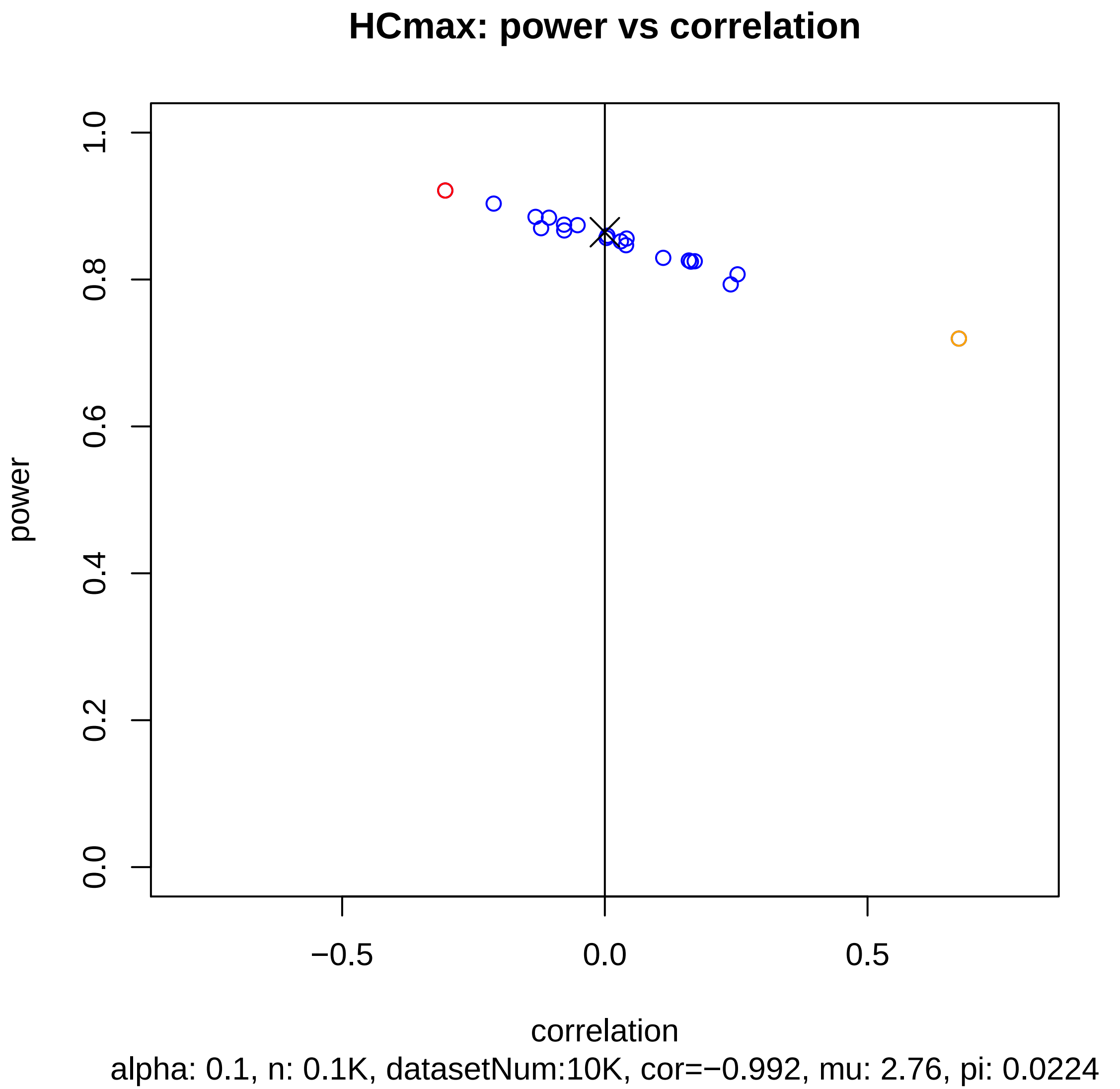
HCmax: power vs correlation
power
1.0
0.8
0.6
0.4
0.2
0.0
−0.5
0.0
0.5
correlation
alpha: 0.1, n: 0.1K, datasetNum:10K, cor=−0.992, mu: 2.76, pi: 0.0224

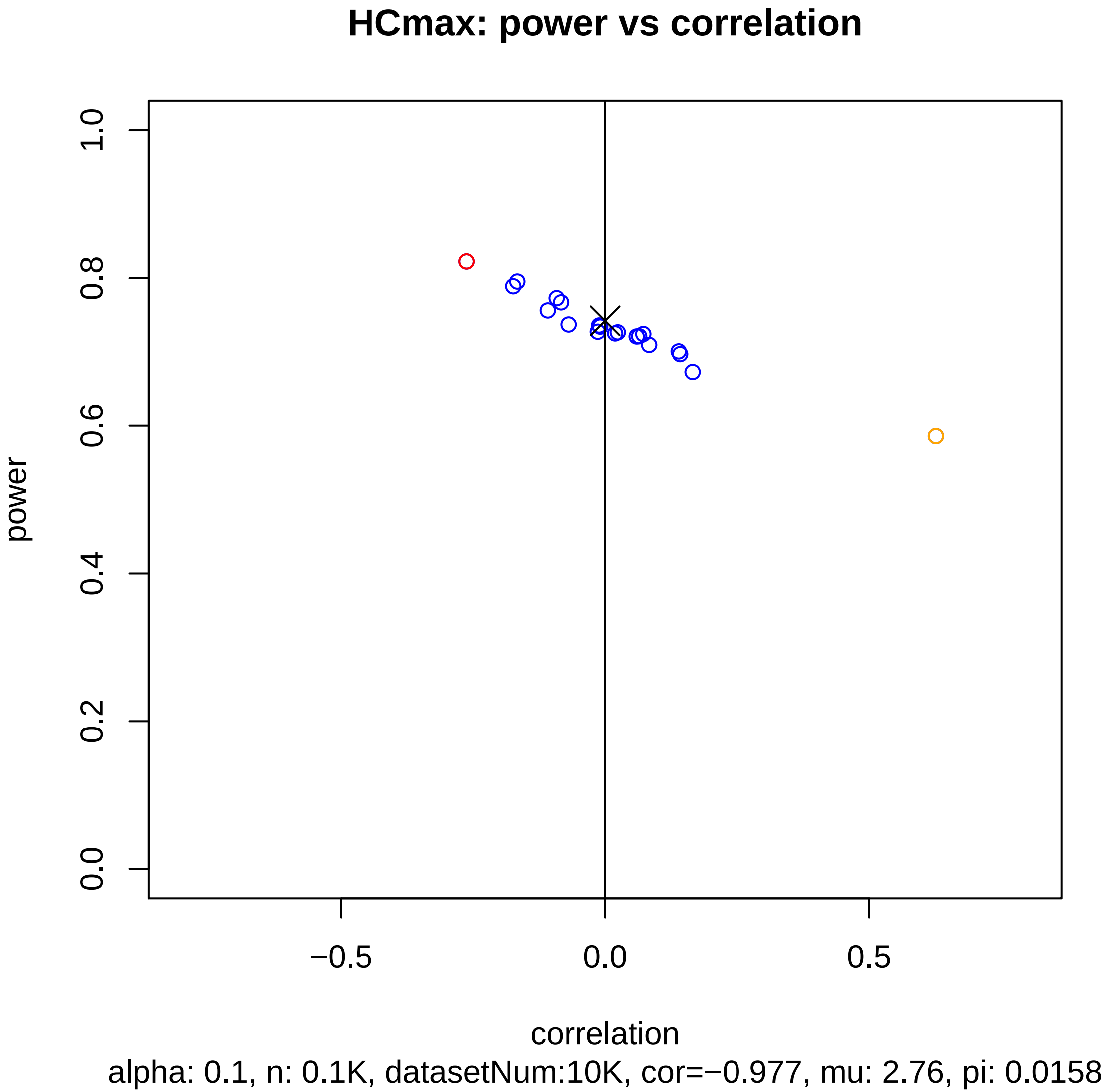
HCmax: power vs correlation
power
0.0
0.2
0.4
0.6
0.8
1.0
−0.5
0.0
0.5
correlation
alpha: 0.1, n: 0.1K, datasetNum:10K, cor=−0.977, mu: 2.76, pi: 0.0158

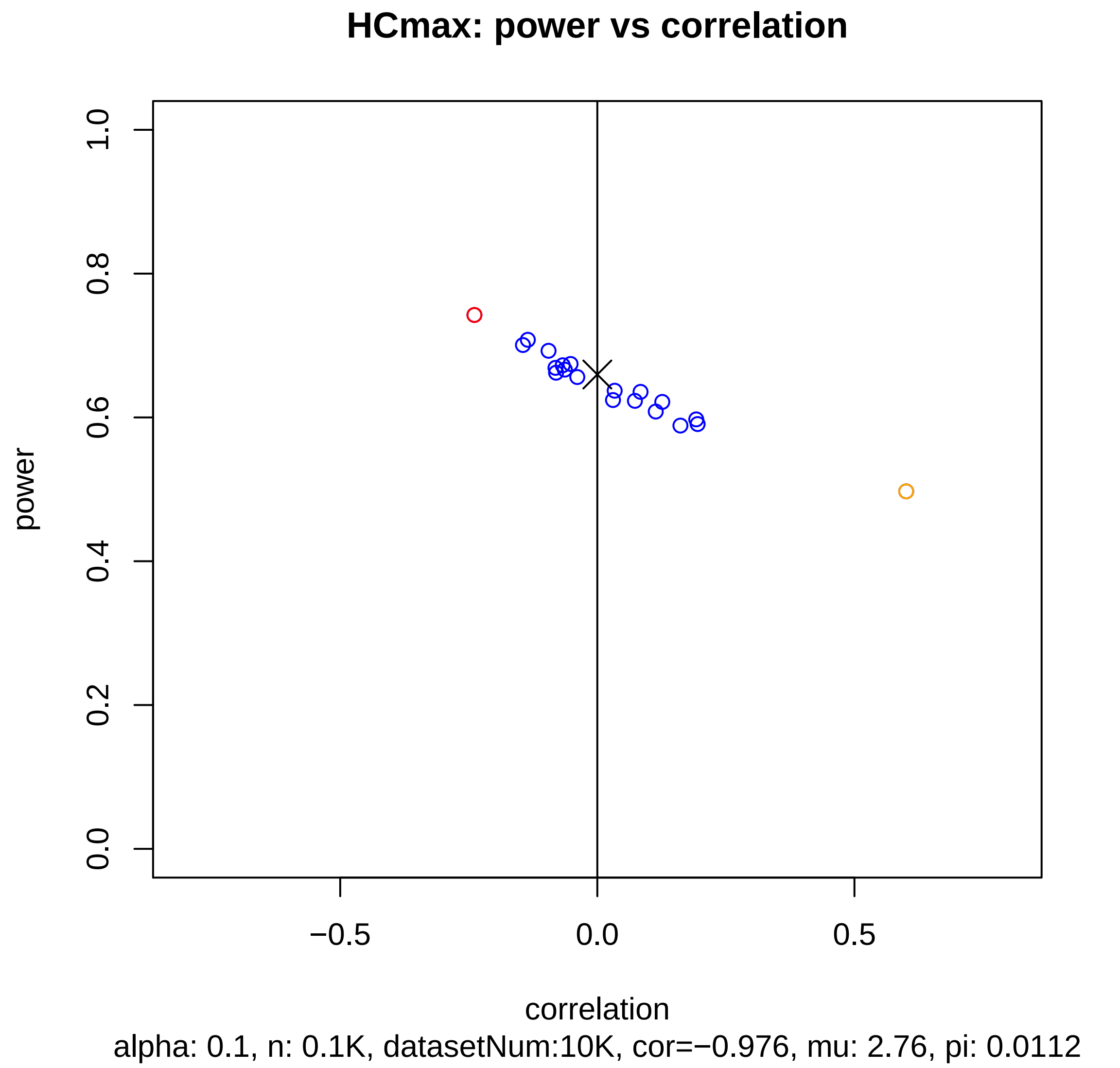
HCmax: power vs correlation
power
1.0
0.8
0.6
0.4
0.2
0.0
−0.5
0.0
0.5
correlation
alpha: 0.1, n: 0.1K, datasetNum:10K, cor=−0.976, mu: 2.76, pi: 0.0112

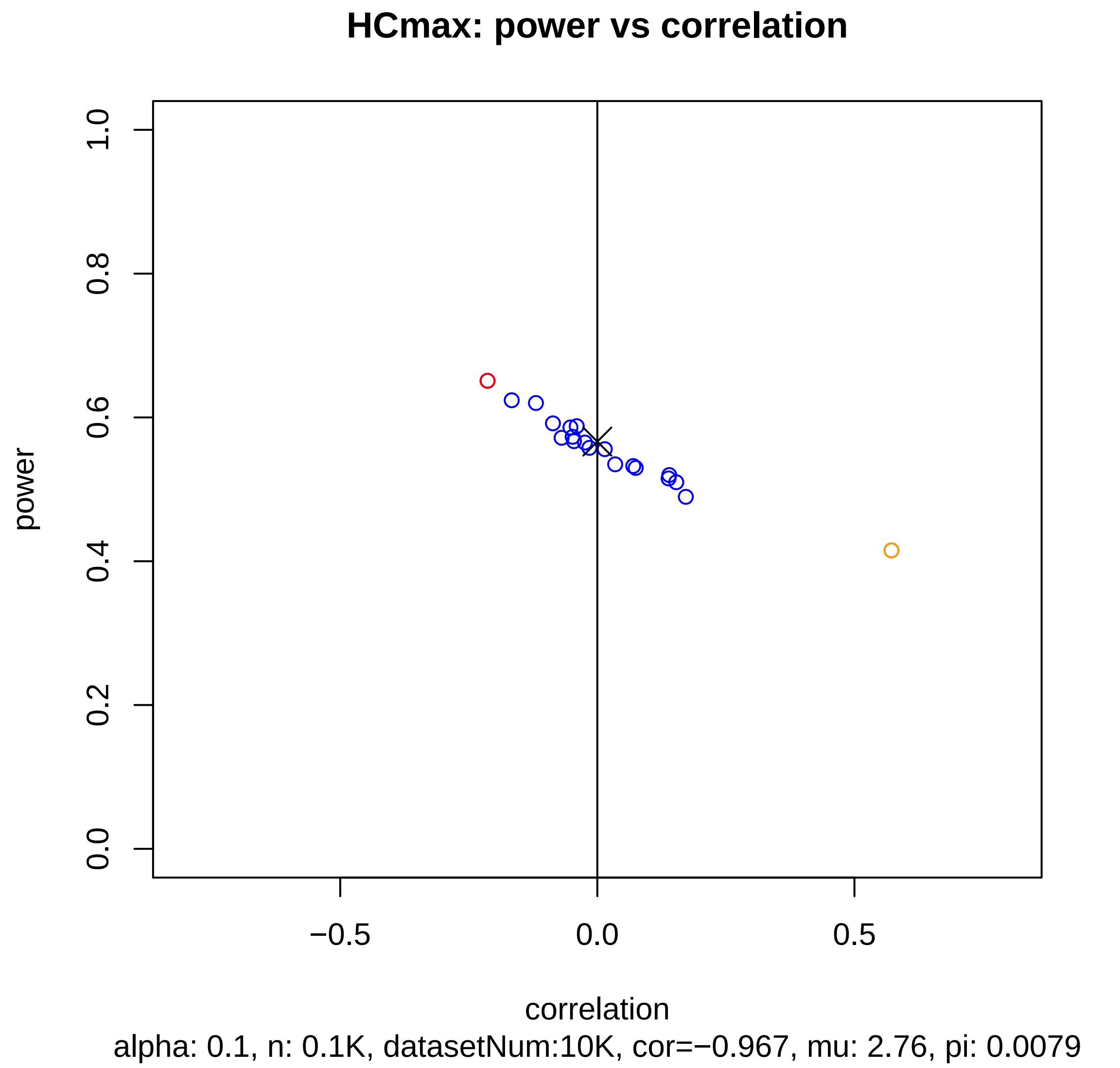
HCmax: power vs correlation
power
0.0
0.2
0.4
0.6
0.8
1.0
−0.5
0.0
0.5
correlation
alpha: 0.1, n: 0.1K, datasetNum:10K, cor=−0.967, mu: 2.76, pi: 0.0079

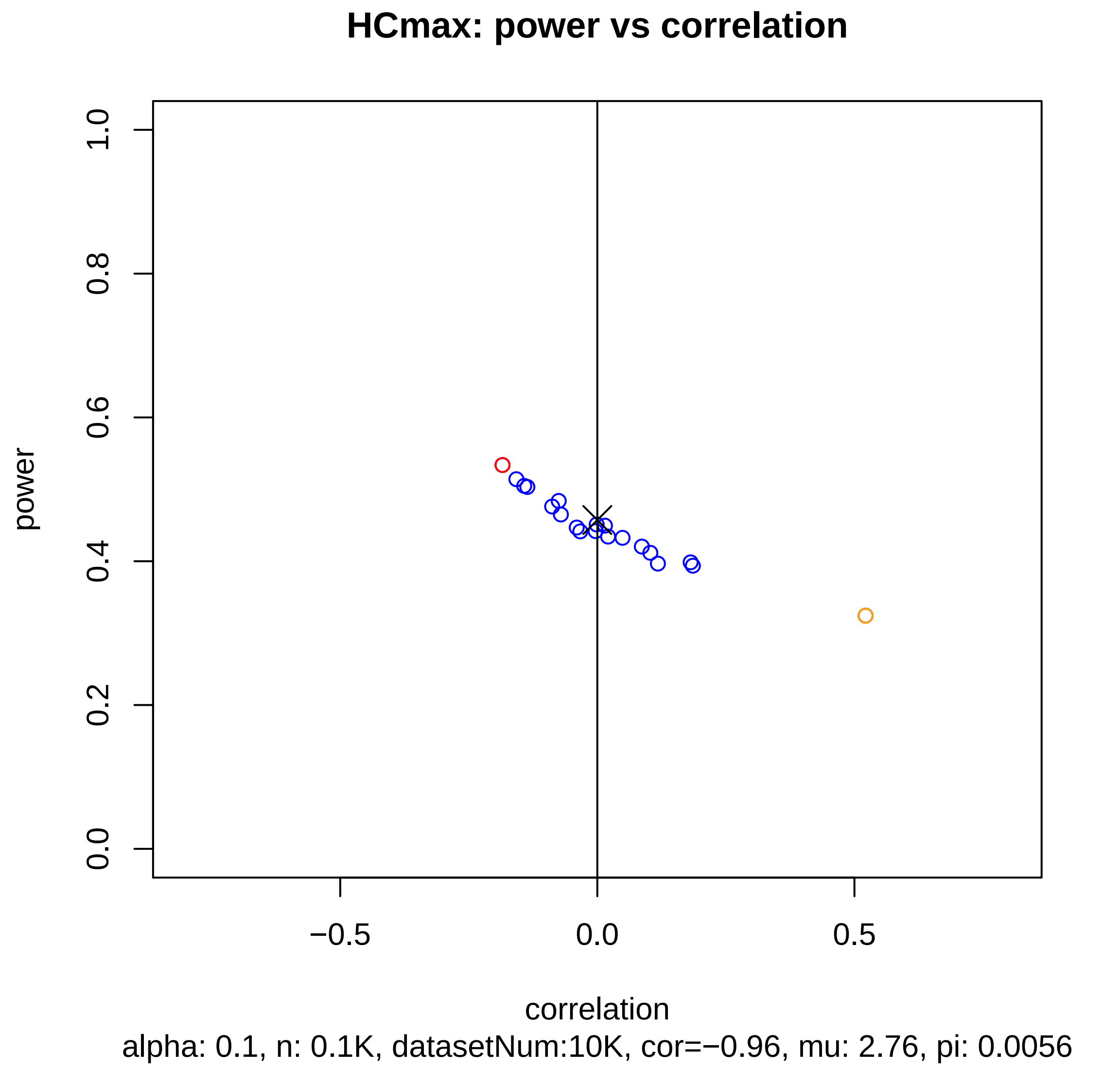
HCmax: power vs correlation
power
1.0
0.8
0.6
0.4
0.2
0.0
−0.5
0.0
0.5
correlation
alpha: 0.1, n: 0.1K, datasetNum:10K, cor=−0.96, mu: 2.76, pi: 0.0056

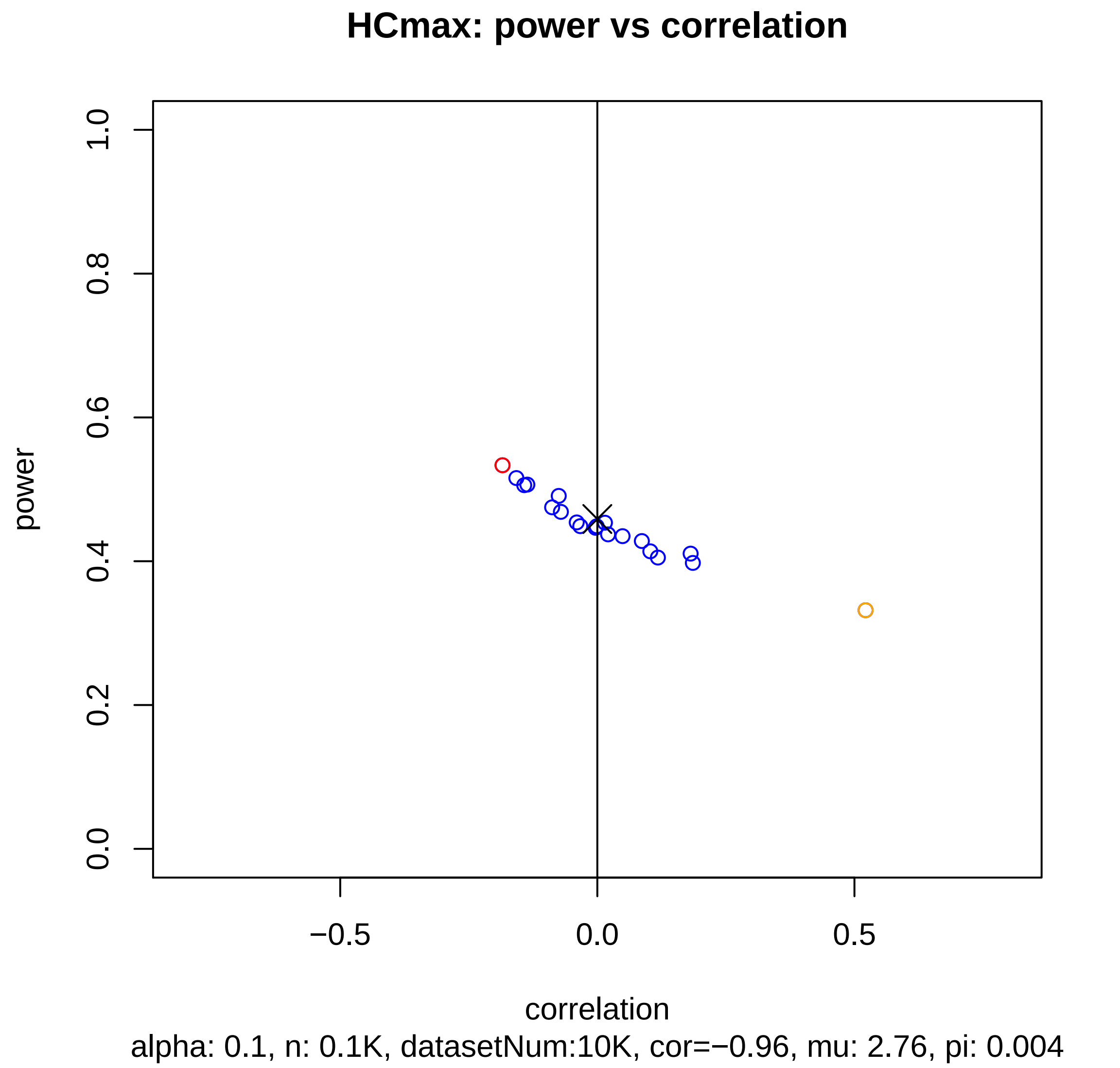
HCmax: power vs correlation
power
1.0
0.8
0.6
0.4
0.2
0.0
−0.5
0.0
0.5
correlation
alpha: 0.1, n: 0.1K, datasetNum:10K, cor=−0.96, mu: 2.76, pi: 0.004

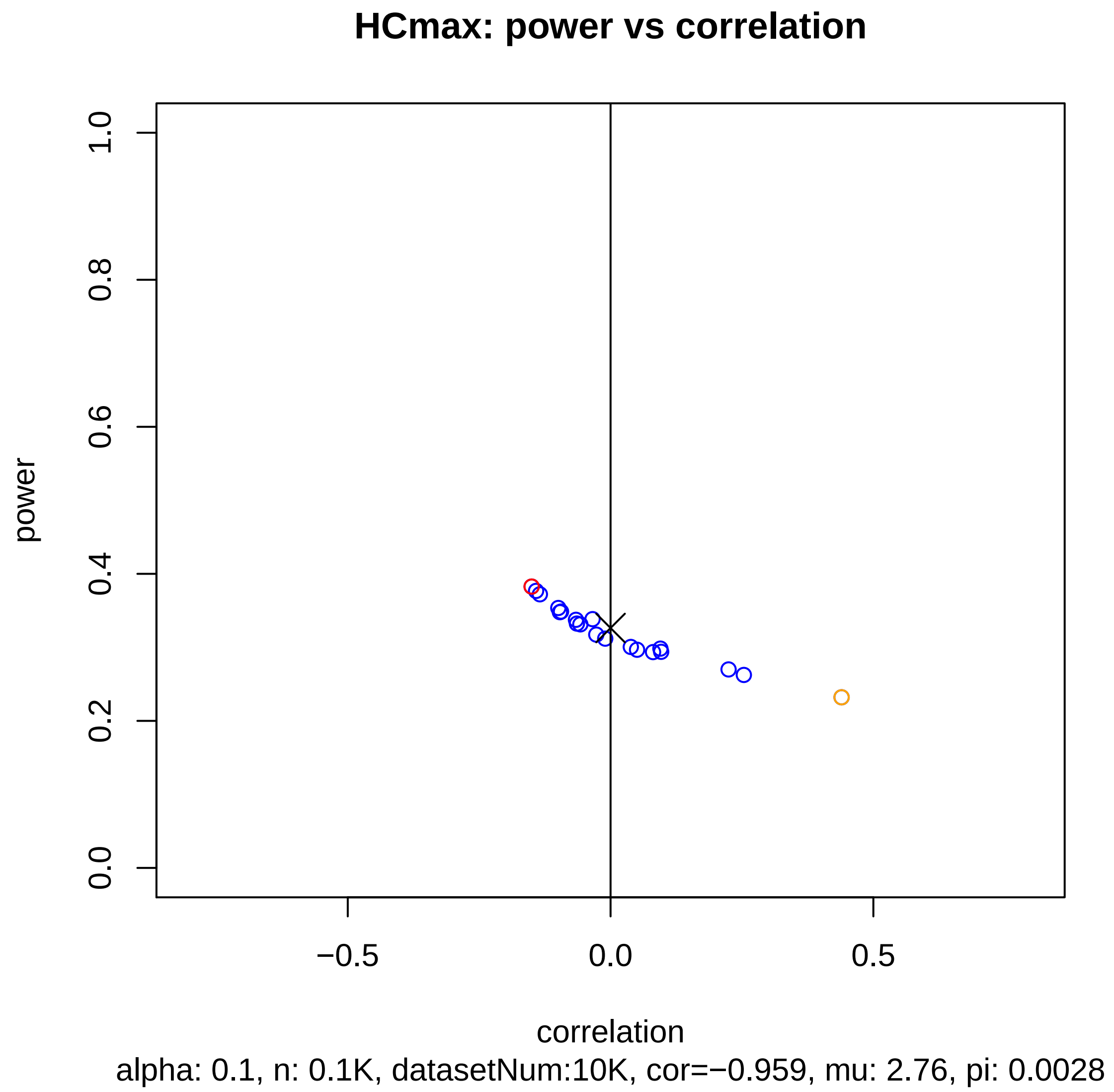
HCmax: power vs correlation
power
correlation
0.0
0.2
0.4
0.6
0.8
1.0
−0.5
0.0
0.5
alpha: 0.1, n: 0.1K, datasetNum:10K, cor=−0.959, mu: 2.76, pi: 0.0028

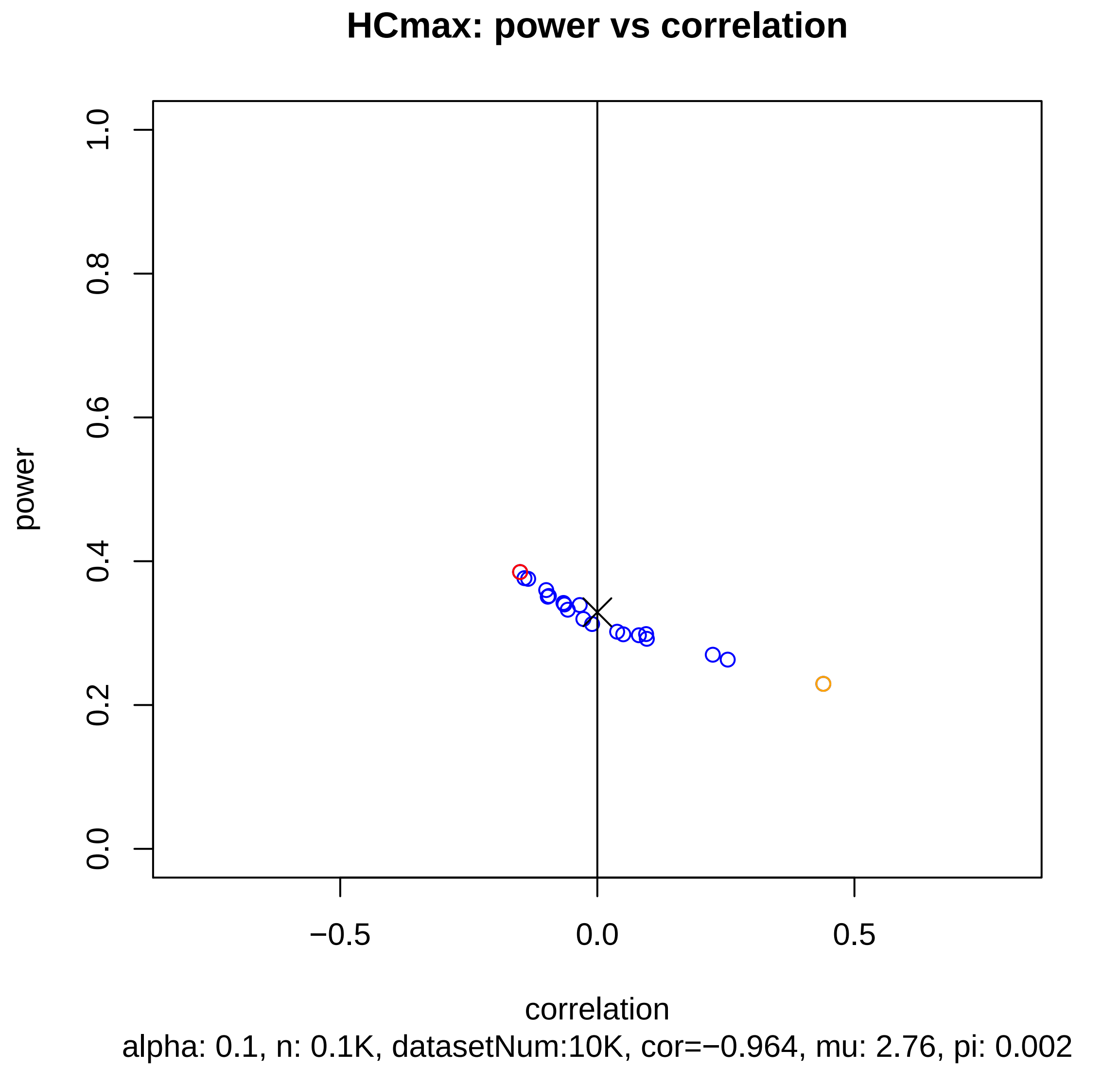
HCmax: power vs correlation
power
1.0
0.8
0.6
0.4
0.2
0.0
−0.5
0.0
0.5
correlation
alpha: 0.1, n: 0.1K, datasetNum:10K, cor=−0.964, mu: 2.76, pi: 0.002

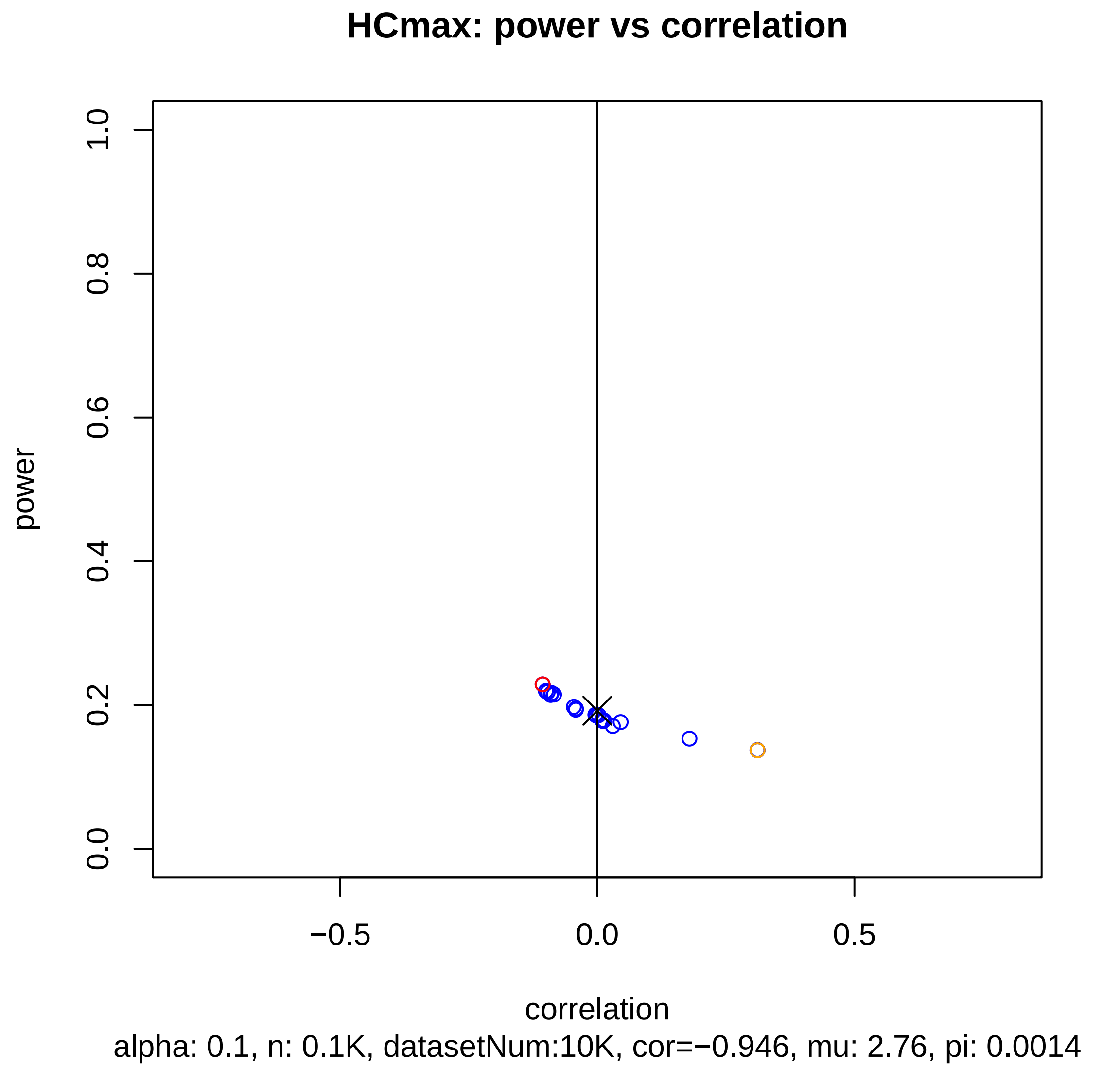

HCmax: power vs correlation
power
1.0
0.8
0.6
0.4
0.2
0.0
−0.5
0.0
0.5
correlation
alpha: 0.1, n: 0.1K, datasetNum:10K, cor=−0.946, mu: 2.76, pi: 0.0014

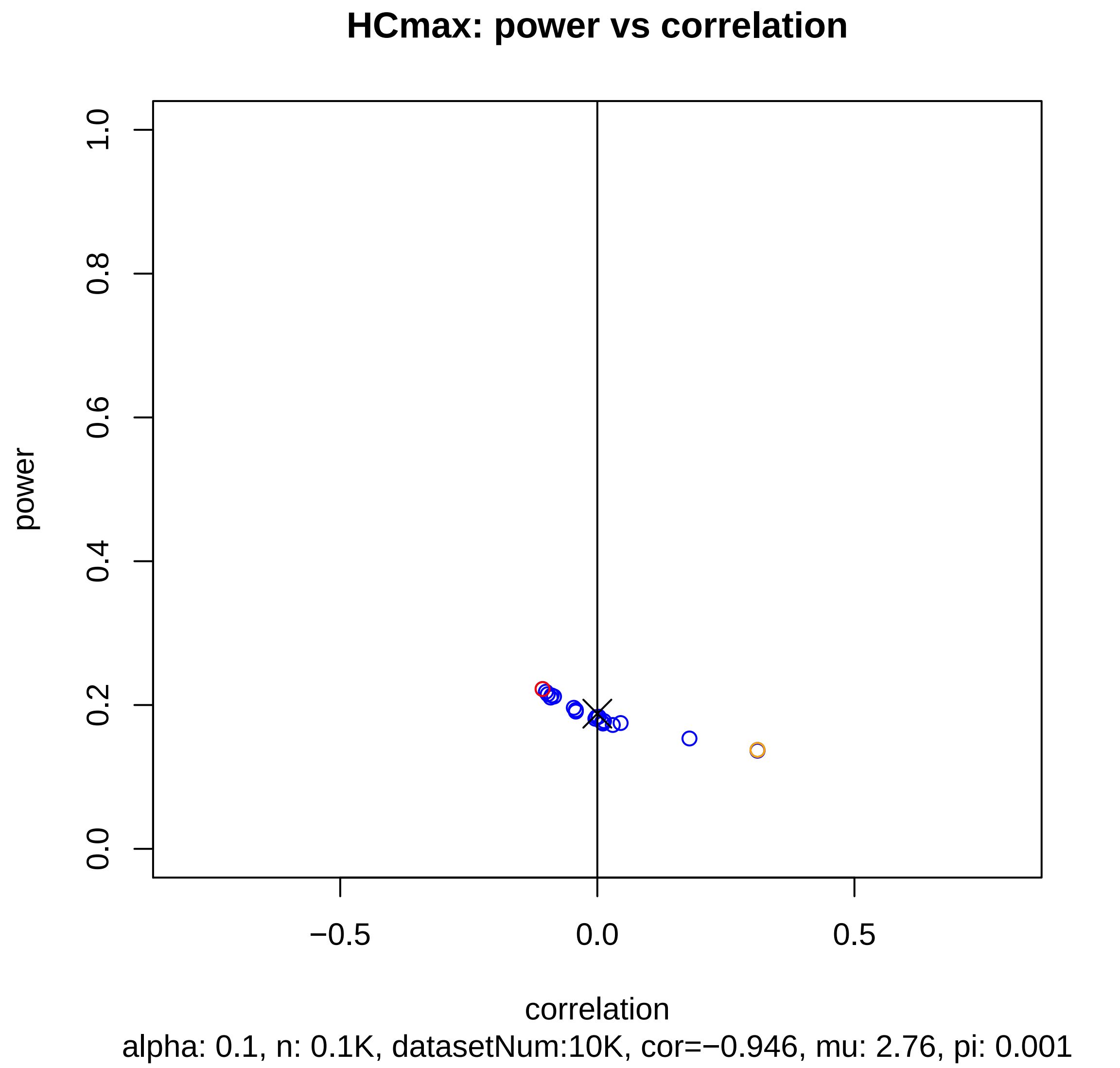

HCmax: power vs correlation
power
0.0
0.2
0.4
0.6
0.8
1.0
−0.5
0.0
0.5
correlation
alpha: 0.1, n: 0.1K, datasetNum:10K, cor=−0.946, mu: 2.76, pi: 0.001

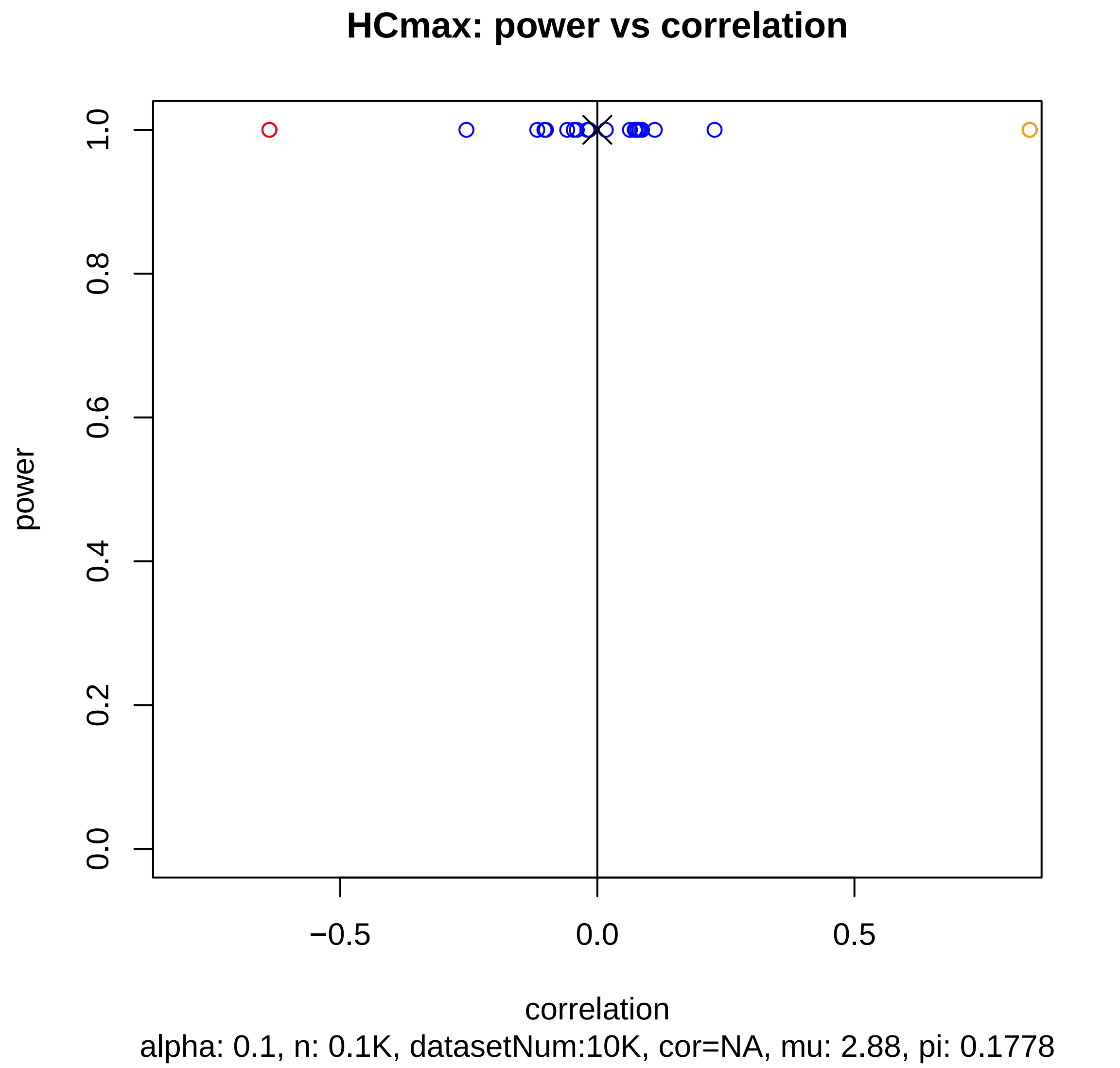
HCmax: power vs correlation
power
1.0
0.8
0.6
0.4
0.2
0.0
−0.5
0.0
0.5
correlation
alpha: 0.1, n: 0.1K, datasetNum:10K, cor=NA, mu: 2.88, pi: 0.1778

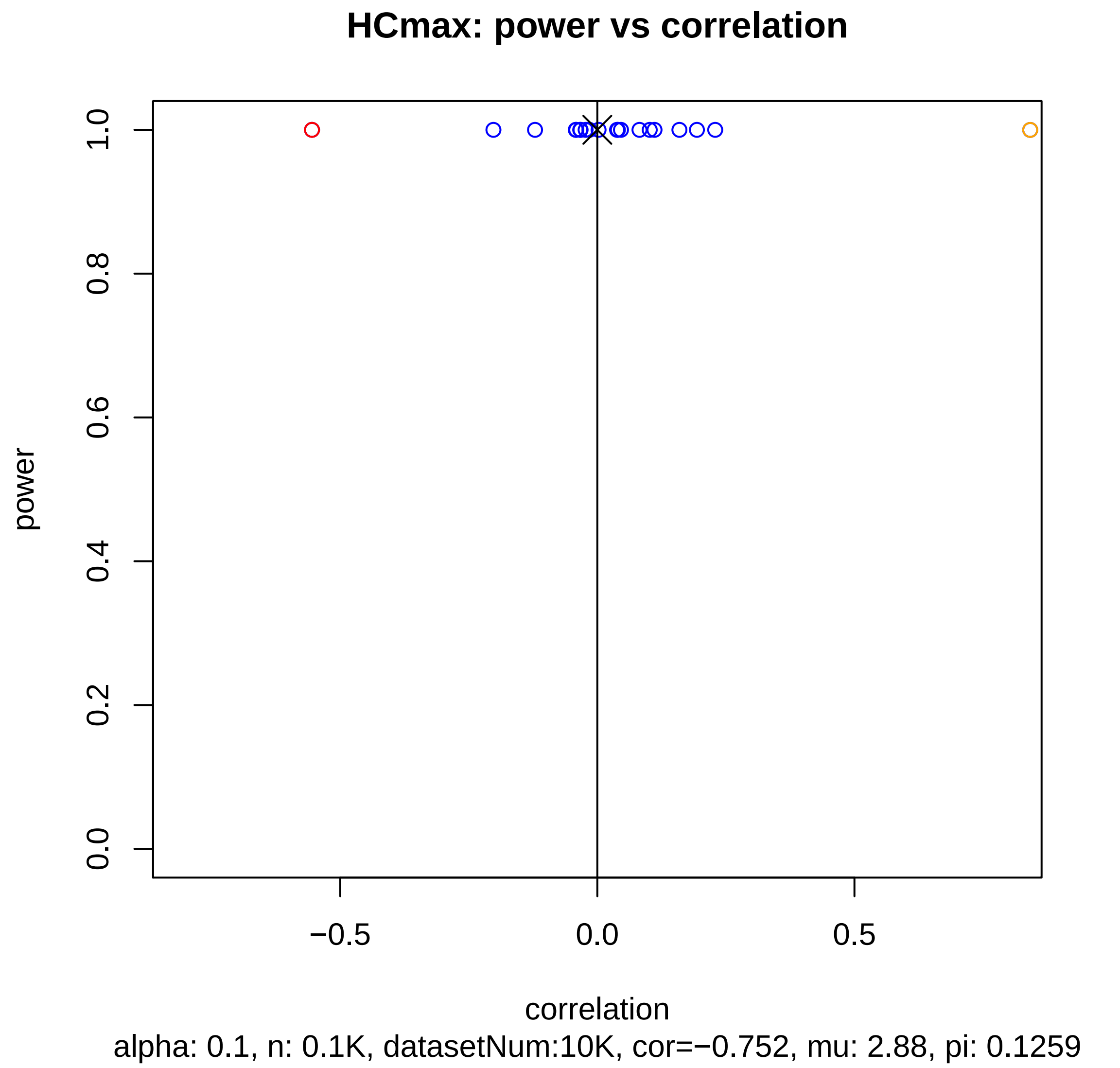
HCmax: power vs correlation
power
0.0
0.2
0.4
0.6
0.8
1.0
−0.5
0.0
0.5
correlation
alpha: 0.1, n: 0.1K, datasetNum:10K, cor=−0.752, mu: 2.88, pi: 0.1259

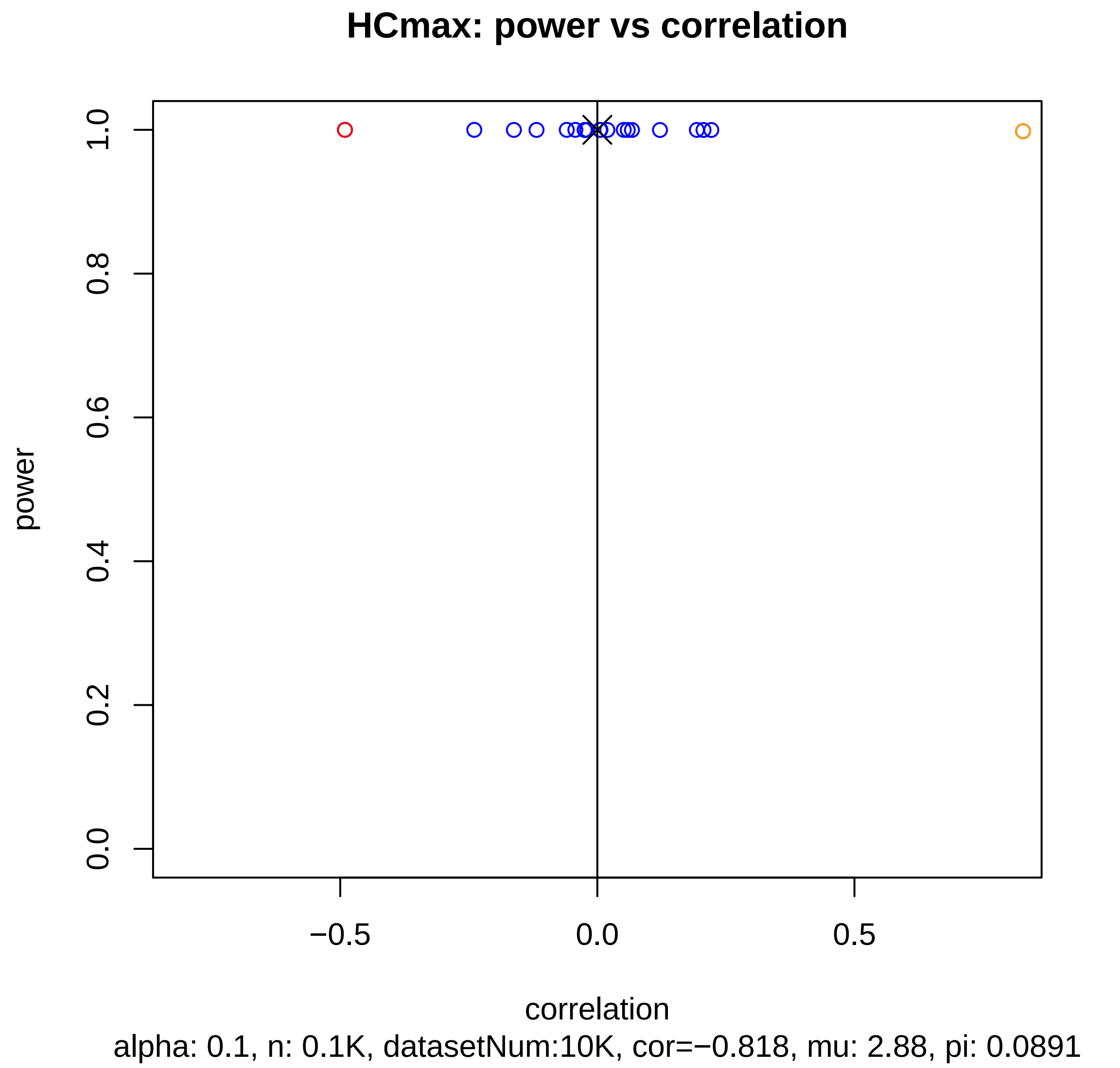

HCmax: power vs correlation
power
1.0
0.8
0.6
0.4
0.2
0.0
−0.5
0.0
0.5
correlation
alpha: 0.1, n: 0.1K, datasetNum:10K, cor=−0.818, mu: 2.88, pi: 0.0891

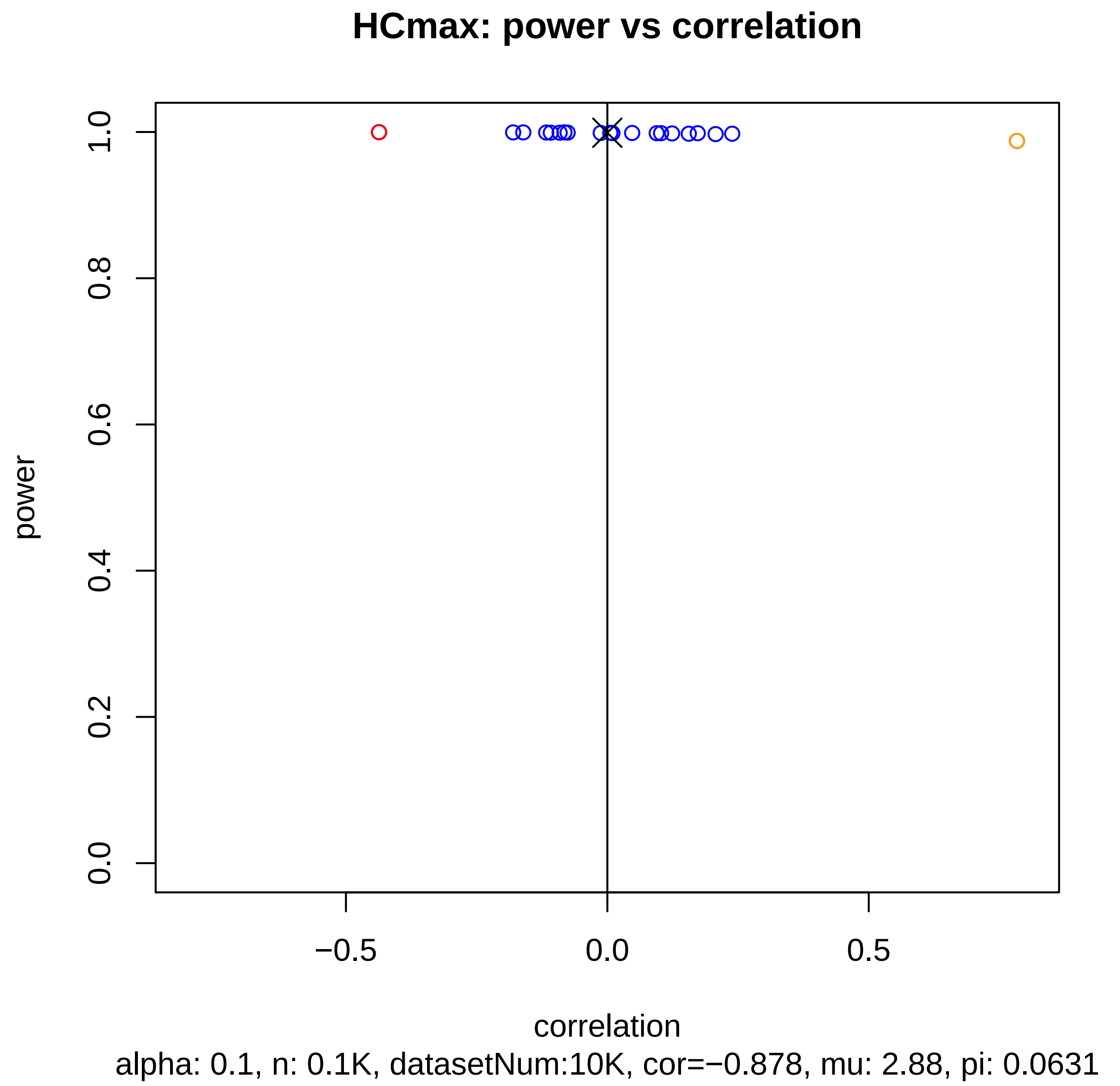
HCmax: power vs correlation
power
0.0
0.2
0.4
0.6
0.8
1.0
−0.5
0.0
0.5
correlation
alpha: 0.1, n: 0.1K, datasetNum:10K, cor=−0.878, mu: 2.88, pi: 0.0631

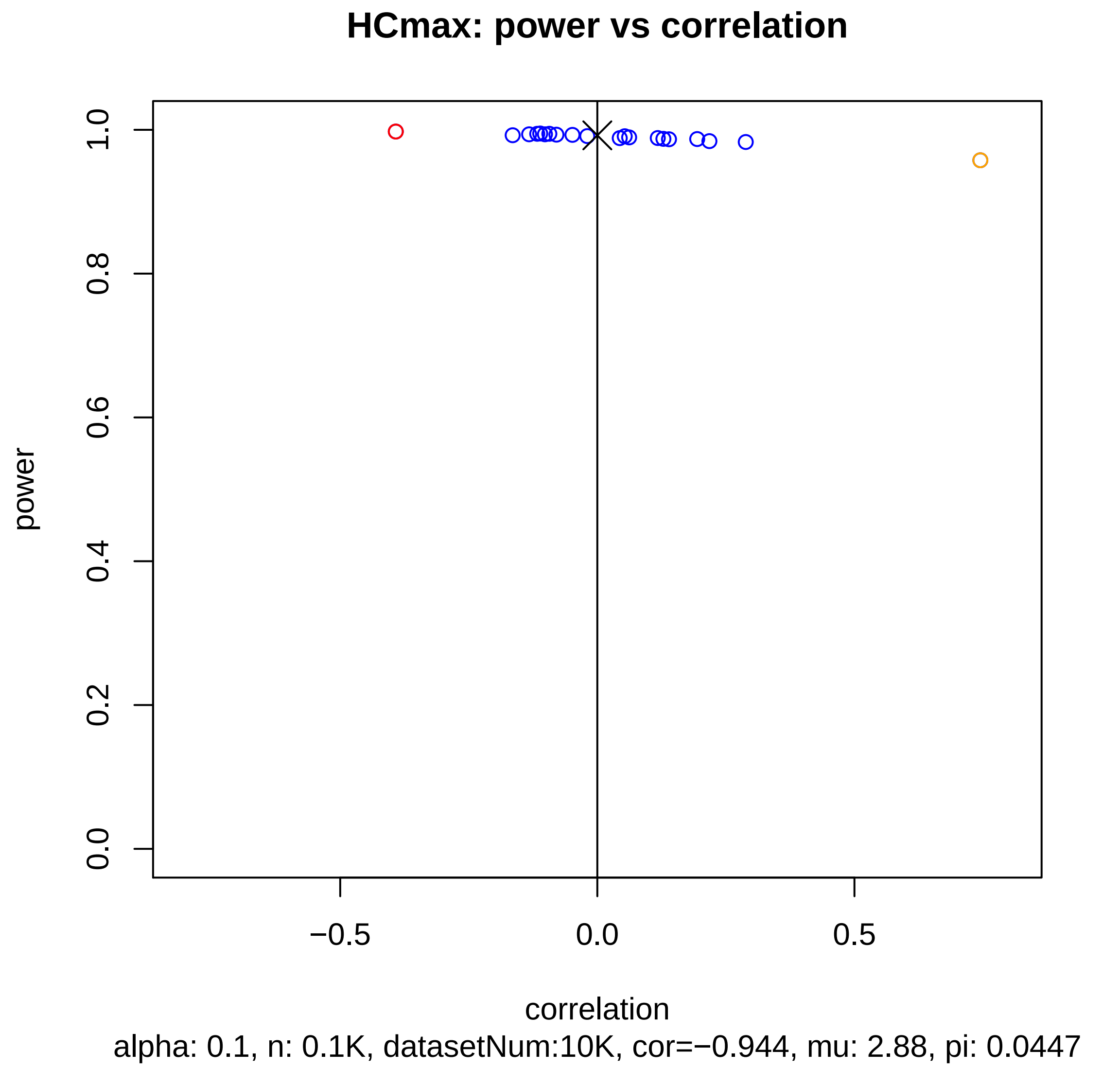
HCmax: power vs correlation
power
correlation
0.0
0.2
0.4
0.6
0.8
1.0
−0.5
0.0
0.5
alpha: 0.1, n: 0.1K, datasetNum:10K, cor=−0.944, mu: 2.88, pi: 0.0447

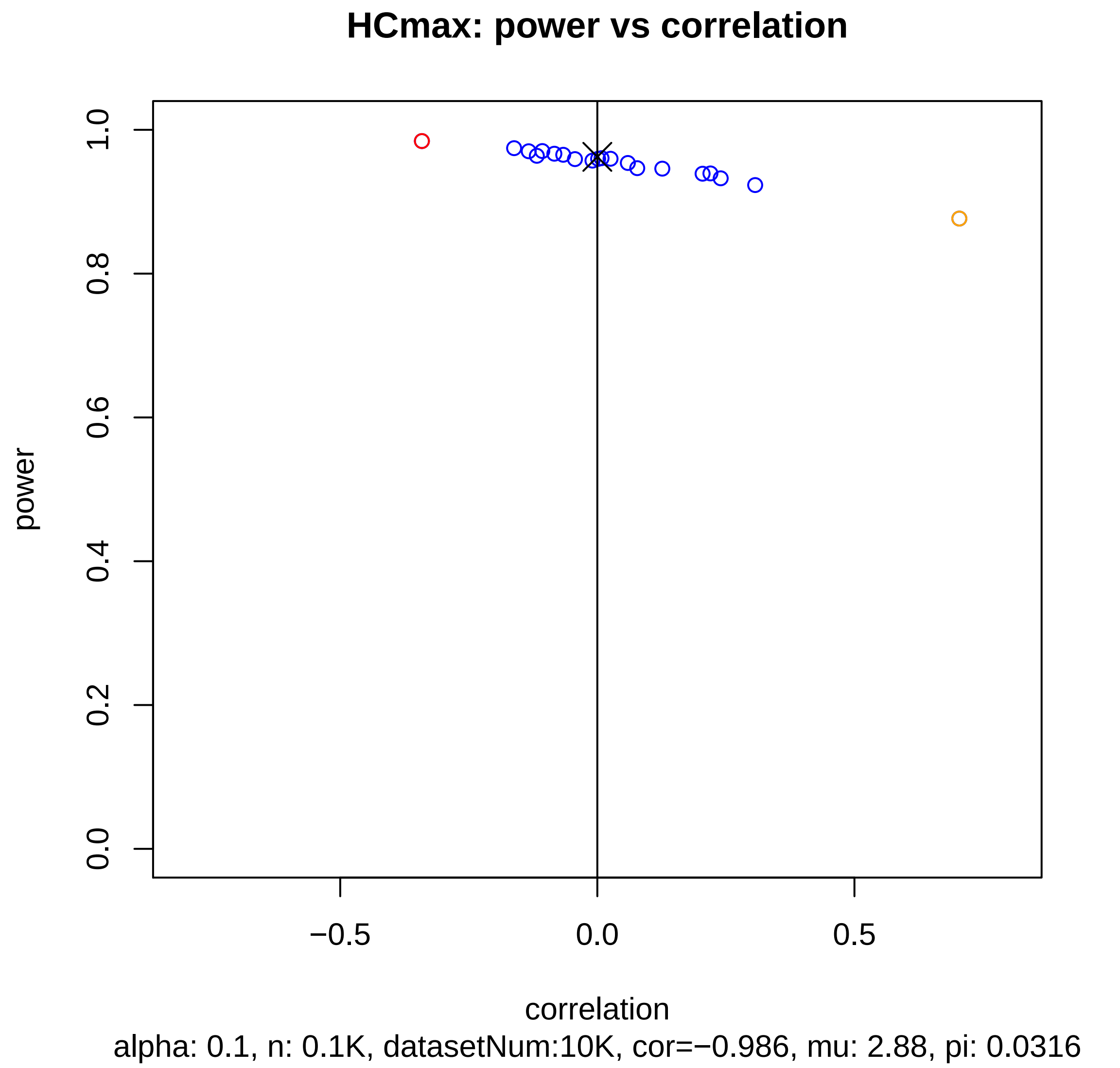
HCmax: power vs correlation
power
0.0
0.2
0.4
0.6
0.8
1.0
−0.5
0.0
0.5
correlation
alpha: 0.1, n: 0.1K, datasetNum:10K, cor=−0.986, mu: 2.88, pi: 0.0316

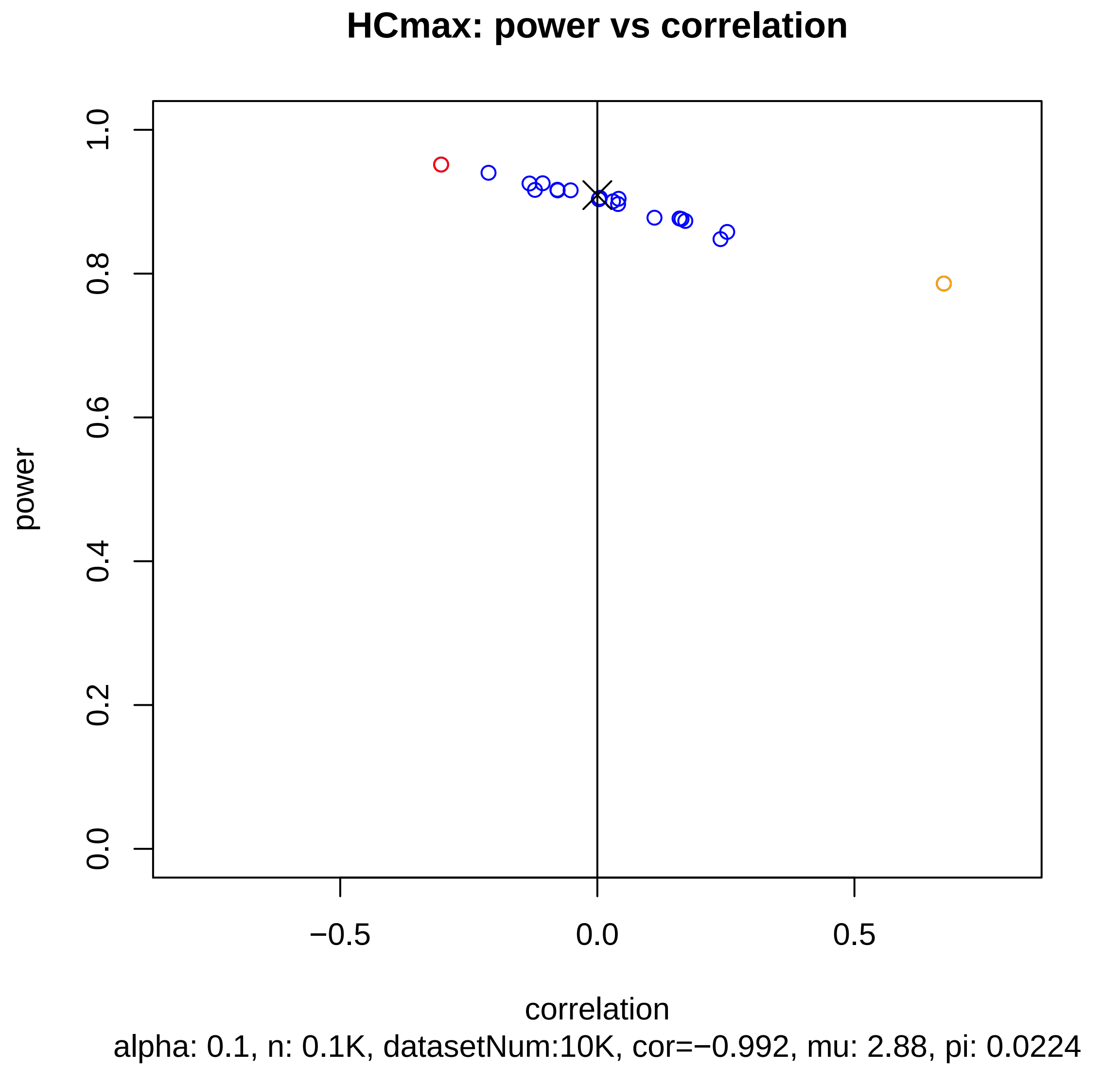
HCmax: power vs correlation
power
1.0
0.8
0.6
0.4
0.2
0.0
−0.5
0.0
0.5
correlation
alpha: 0.1, n: 0.1K, datasetNum:10K, cor=−0.992, mu: 2.88, pi: 0.0224

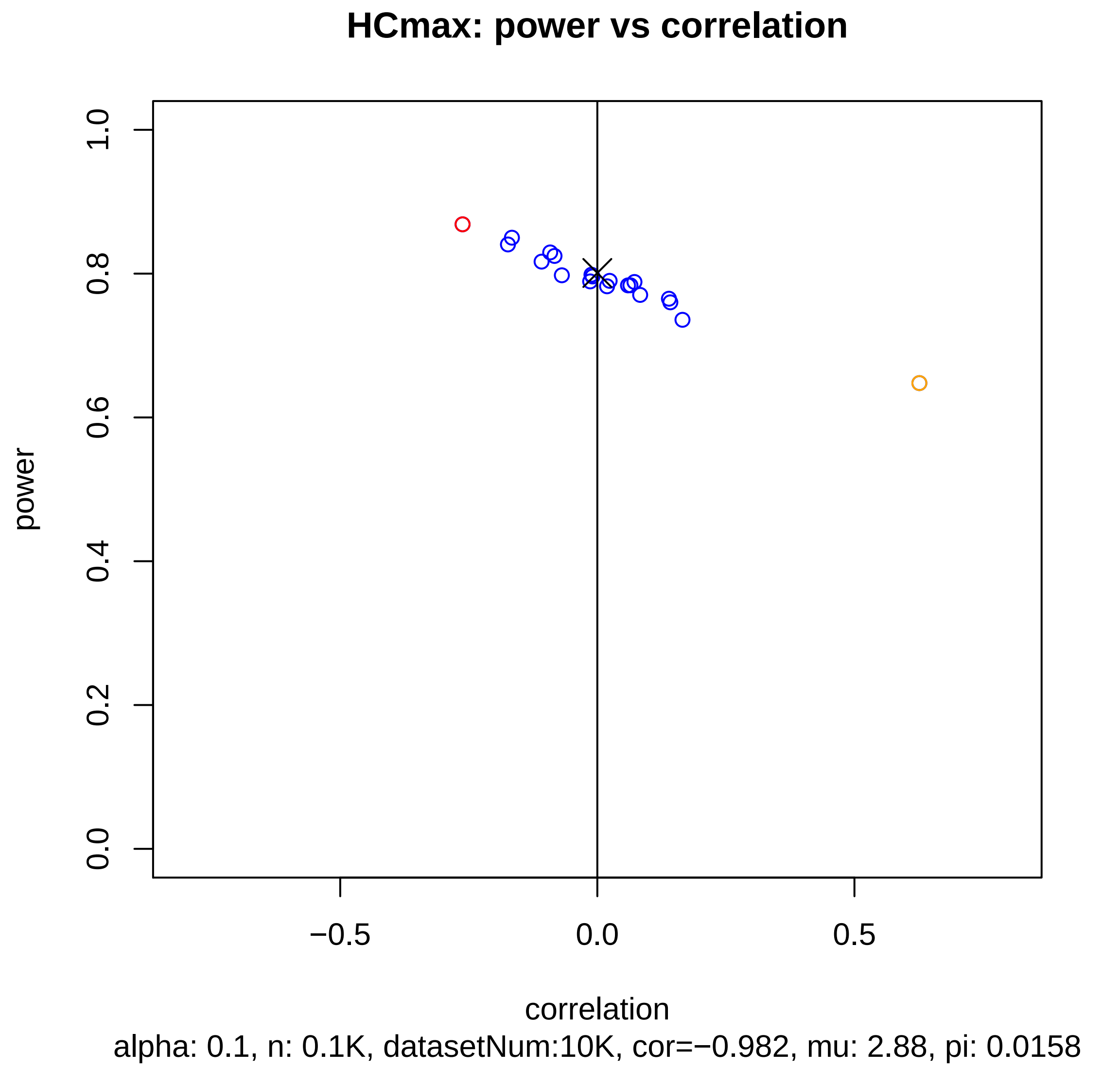
HCmax: power vs correlation
power
1.0
0.8
0.6
0.4
0.2
0.0
−0.5
0.0
0.5
correlation
alpha: 0.1, n: 0.1K, datasetNum:10K, cor=−0.982, mu: 2.88, pi: 0.0158

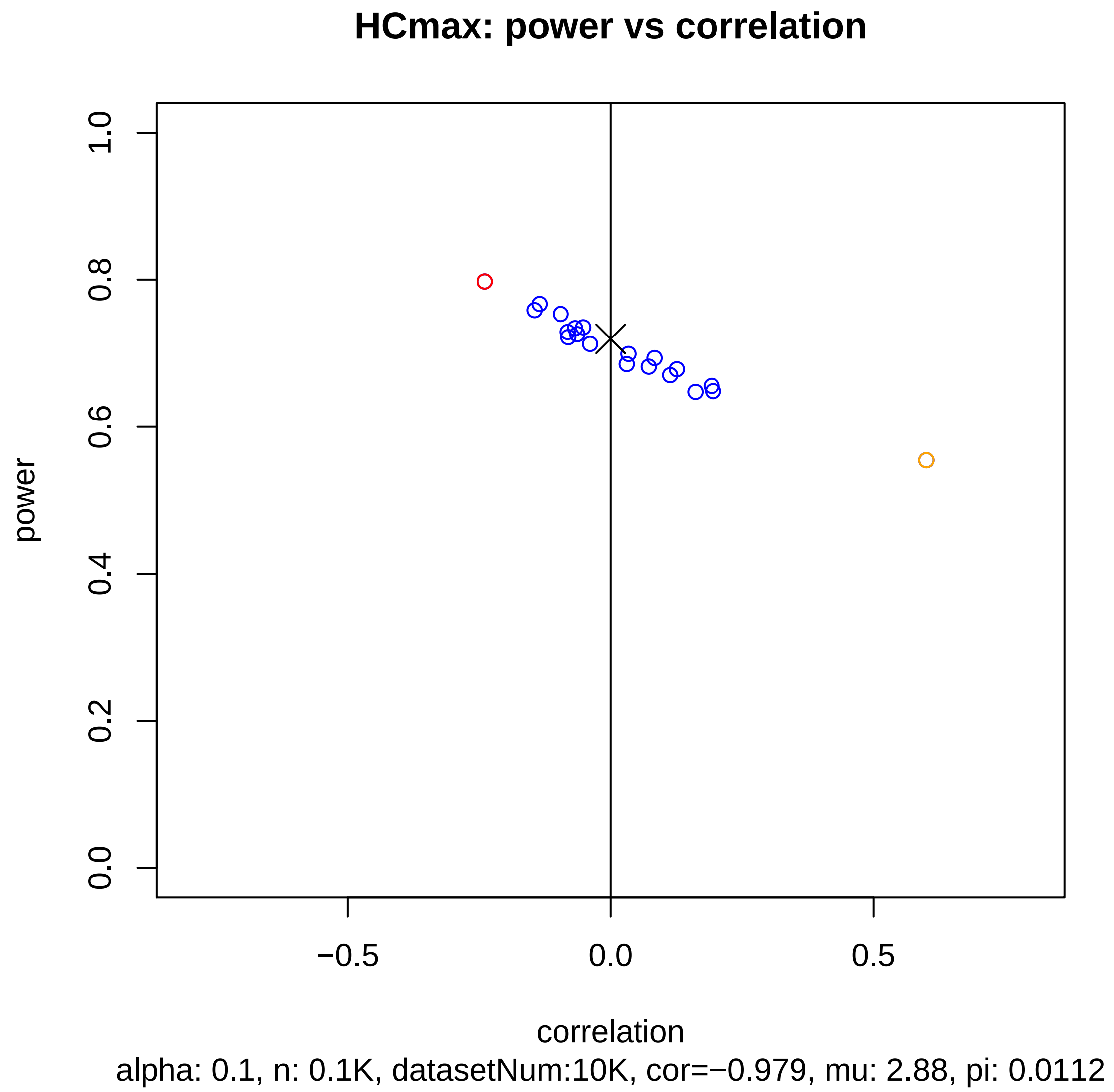
HCmax: power vs correlation
power
0.0
0.2
0.4
0.6
0.8
1.0
−0.5
0.0
0.5
correlation
alpha: 0.1, n: 0.1K, datasetNum:10K, cor=−0.979, mu: 2.88, pi: 0.0112

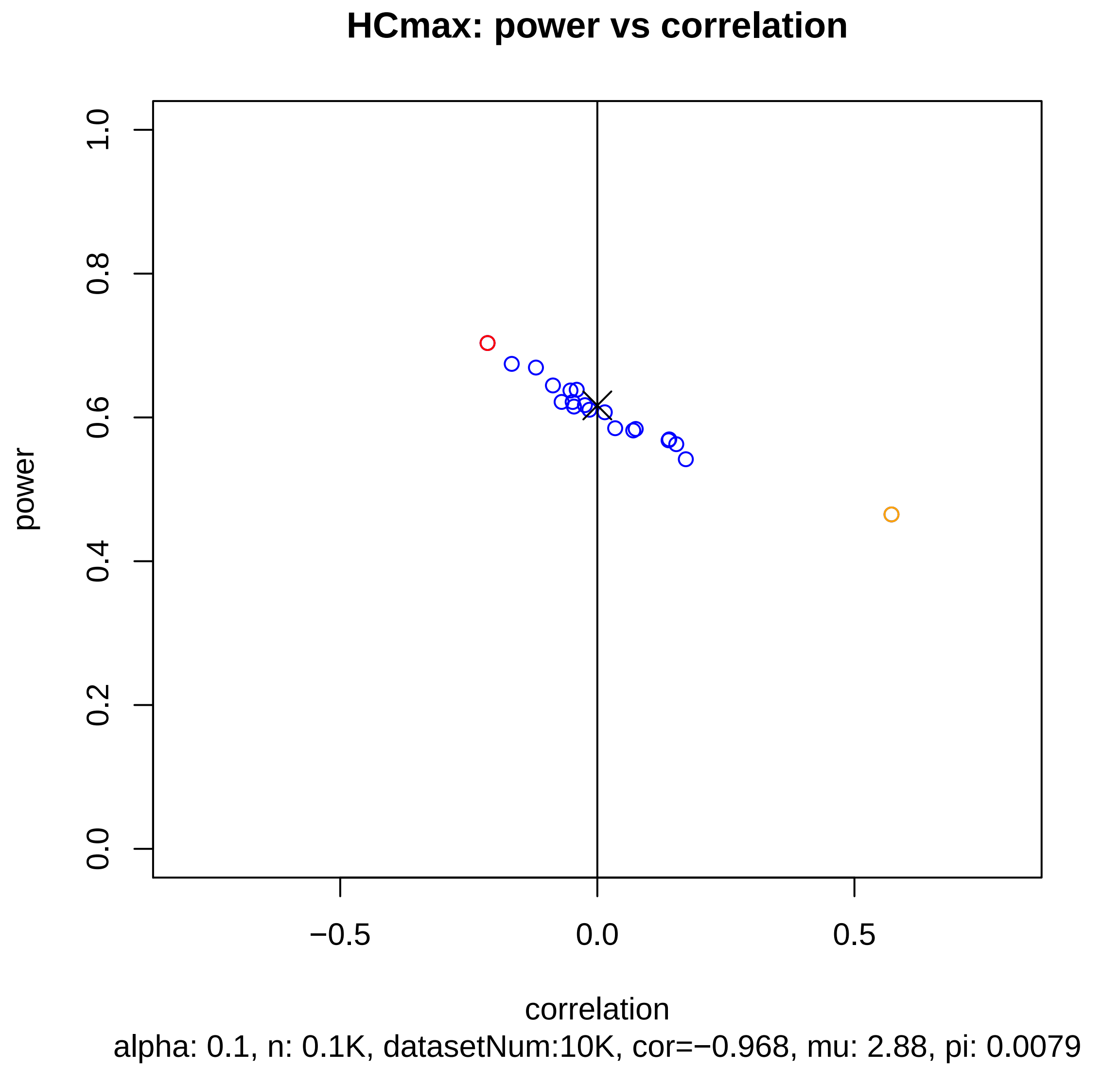

HCmax: power vs correlation
power
correlation
0.0
0.2
0.4
0.6
0.8
1.0
−0.5
0.0
0.5
alpha: 0.1, n: 0.1K, datasetNum:10K, cor=−0.968, mu: 2.88, pi: 0.0079

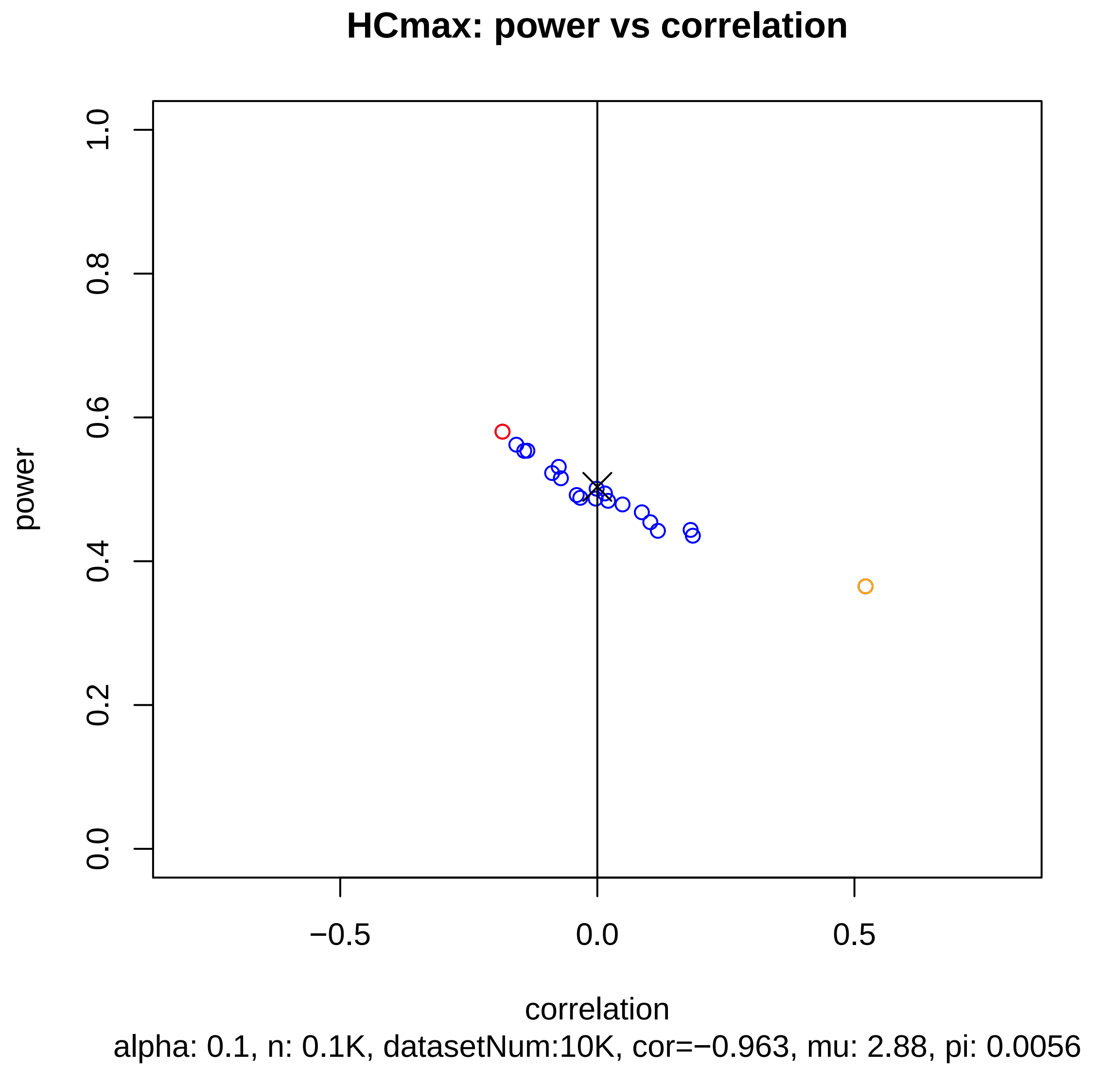
HCmax: power vs correlation
power
correlation
1.0
0.8
0.6
0.4
0.2
0.0
−0.5
0.0
0.5
alpha: 0.1, n: 0.1K, datasetNum:10K, cor=−0.963, mu: 2.88, pi: 0.0056

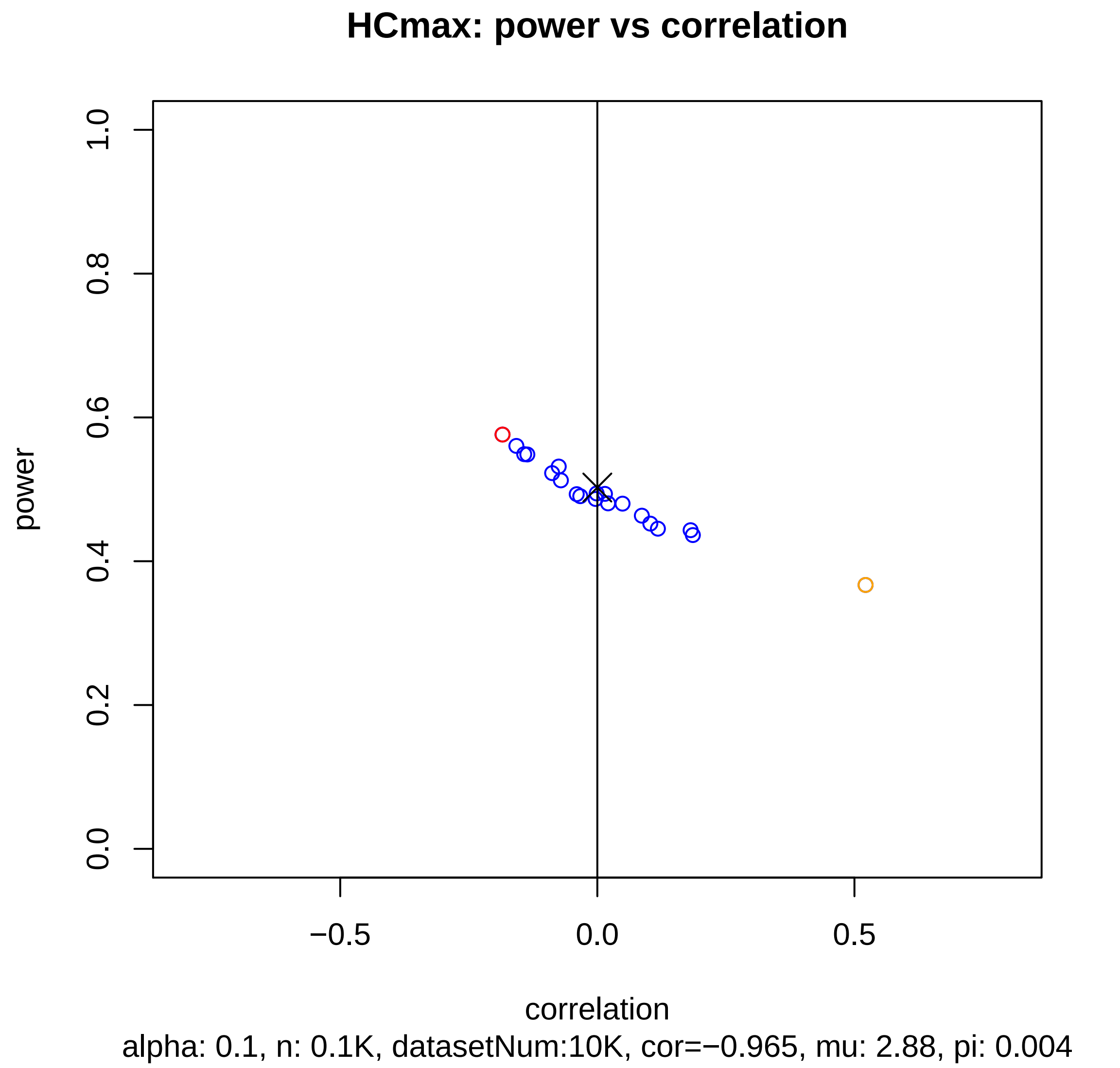
HCmax: power vs correlation
power
0.0
0.2
0.4
0.6
0.8
1.0
−0.5
0.0
0.5
correlation
alpha: 0.1, n: 0.1K, datasetNum:10K, cor=−0.965, mu: 2.88, pi: 0.004

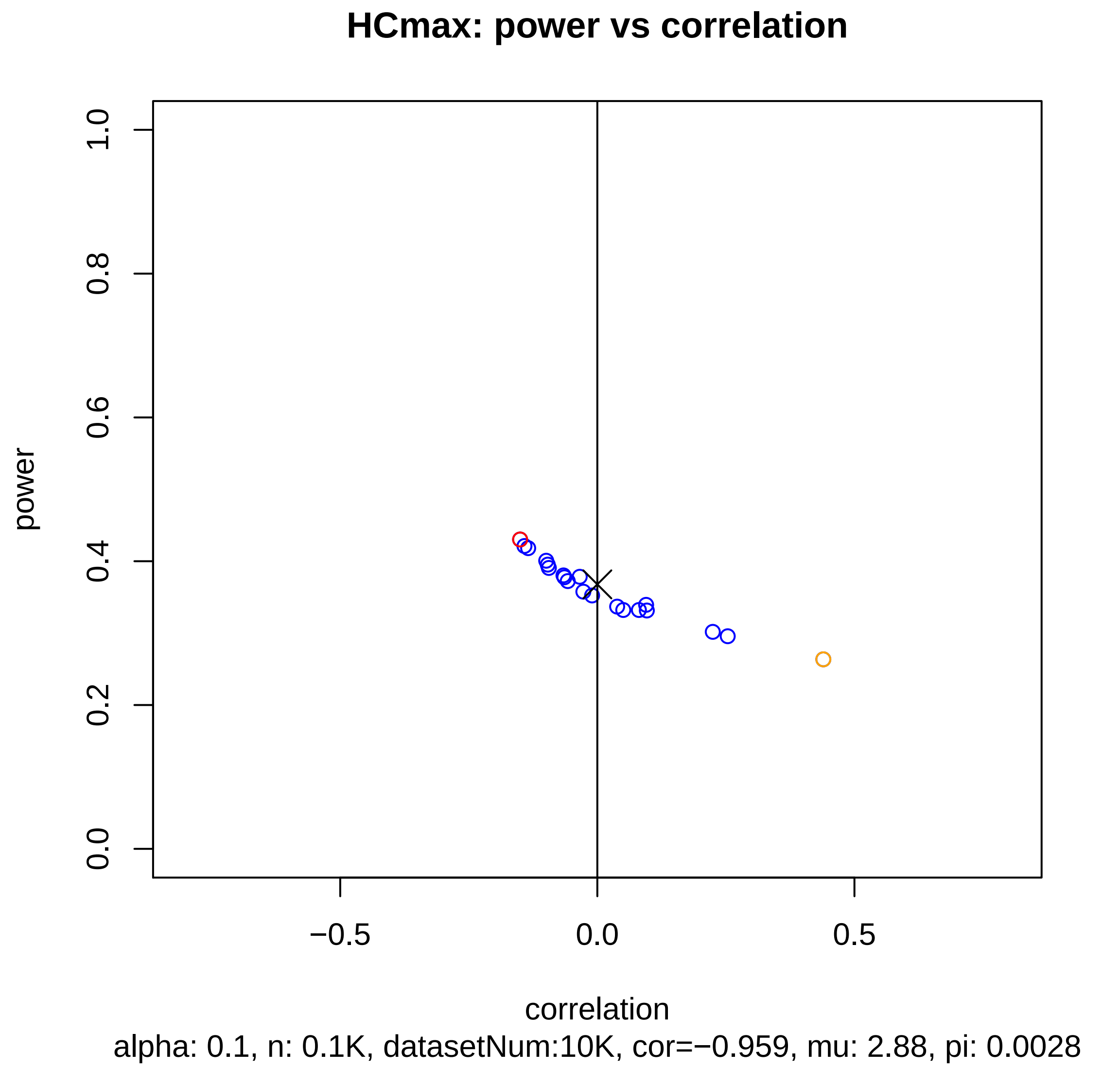
HCmax: power vs correlation
power
0.0
0.2
0.4
0.6
0.8
1.0
−0.5
0.0
0.5
correlation
alpha: 0.1, n: 0.1K, datasetNum:10K, cor=−0.959, mu: 2.88, pi: 0.0028

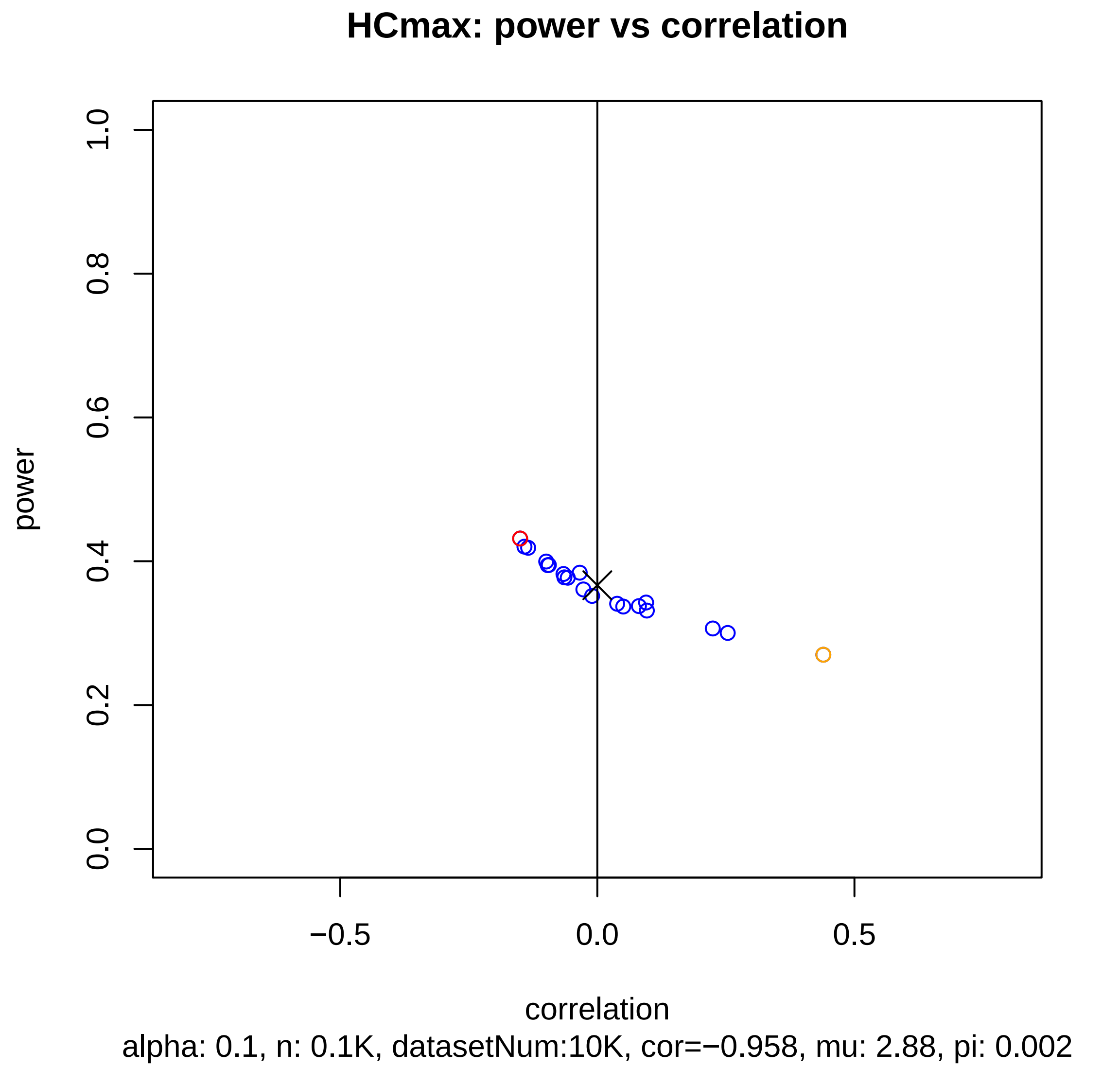
HCmax: power vs correlation
power
1.0
0.8
0.6
0.4
0.2
0.0
−0.5
0.0
0.5
correlation
alpha: 0.1, n: 0.1K, datasetNum:10K, cor=−0.958, mu: 2.88, pi: 0.002

### HCmax: power vs correlation

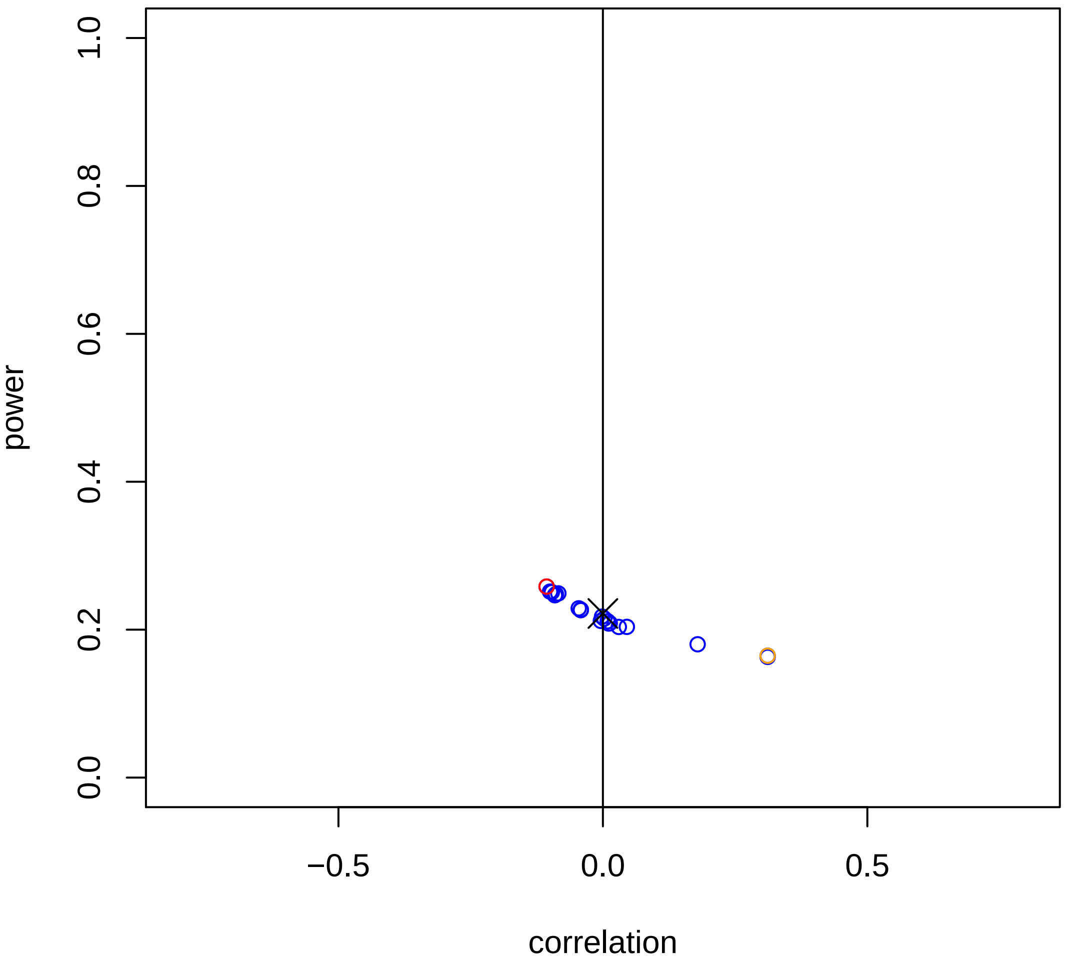


alpha: 0.1, n: 0.1K, datasetNum:10K, cor=−0.952, mu: 2.88, pi: 0.0014

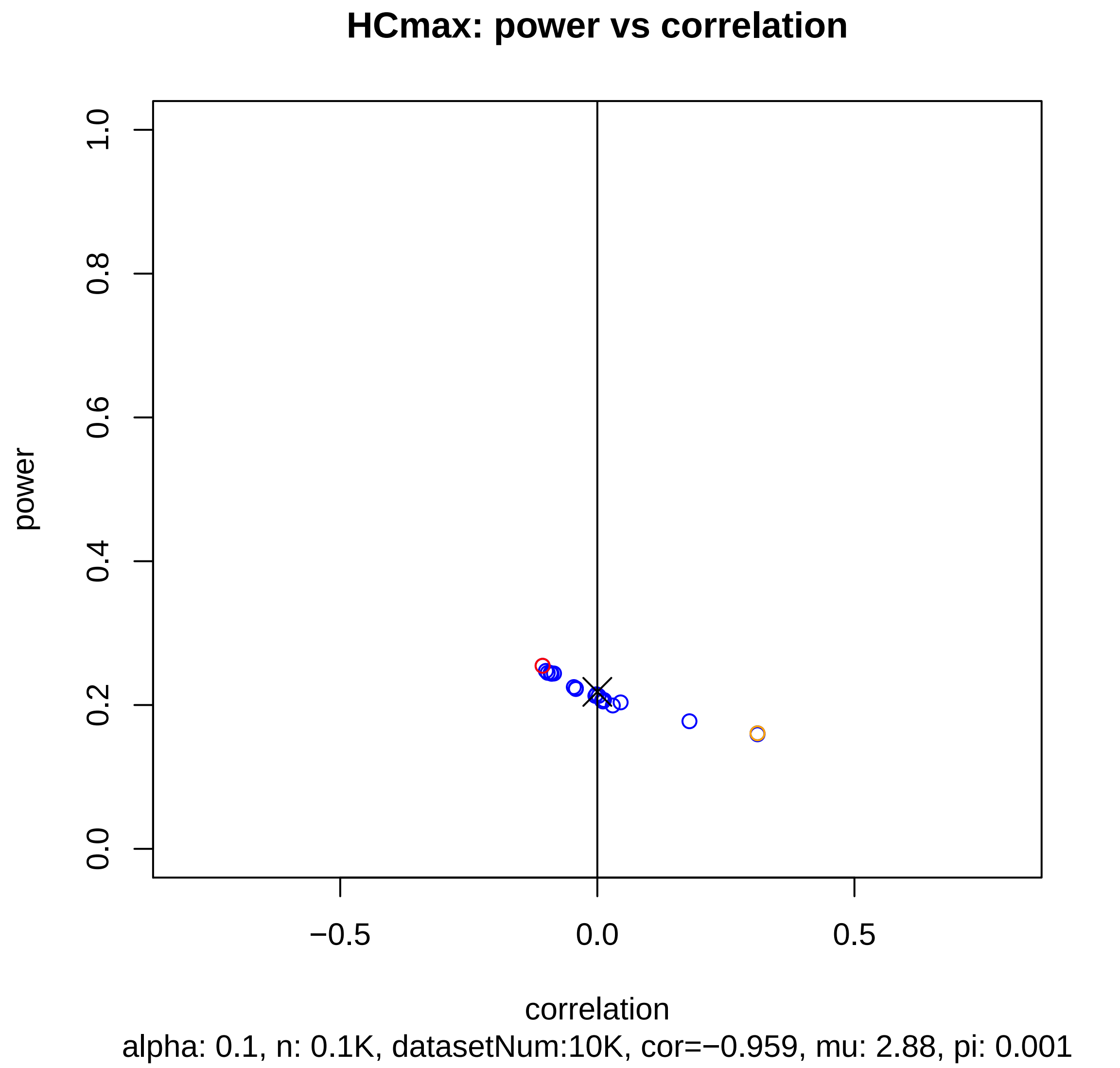
HCmax: power vs correlation
power
0.0
0.2
0.4
0.6
0.8
1.0
−0.5
0.0
0.5
correlation
alpha: 0.1, n: 0.1K, datasetNum:10K, cor=−0.959, mu: 2.88, pi: 0.001

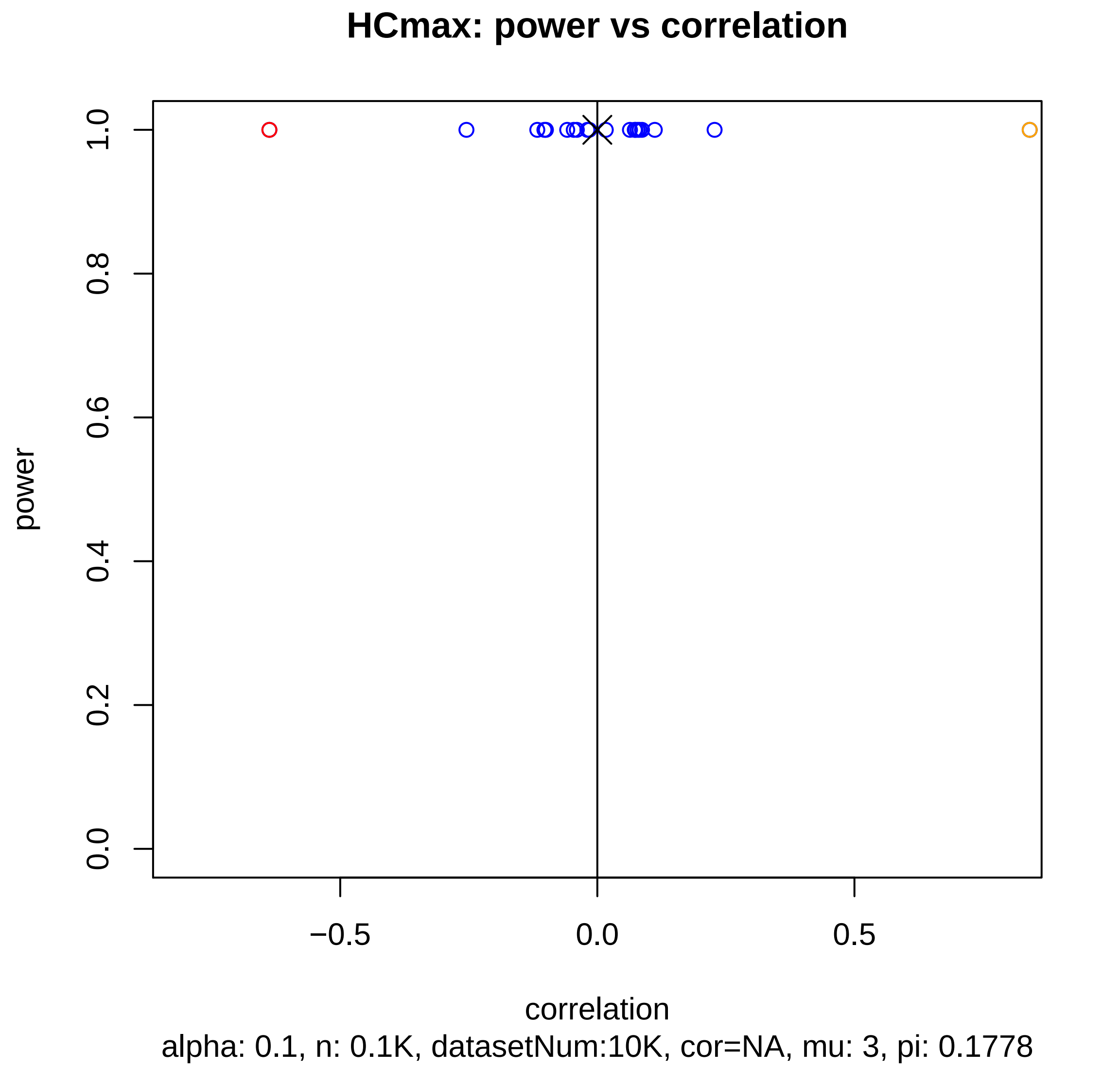
HCmax: power vs correlation
power
correlation
1.0
0.8
0.6
0.4
0.2
0.0
−0.5
0.0
0.5
alpha: 0.1, n: 0.1K, datasetNum:10K, cor=NA, mu: 3, pi: 0.1778

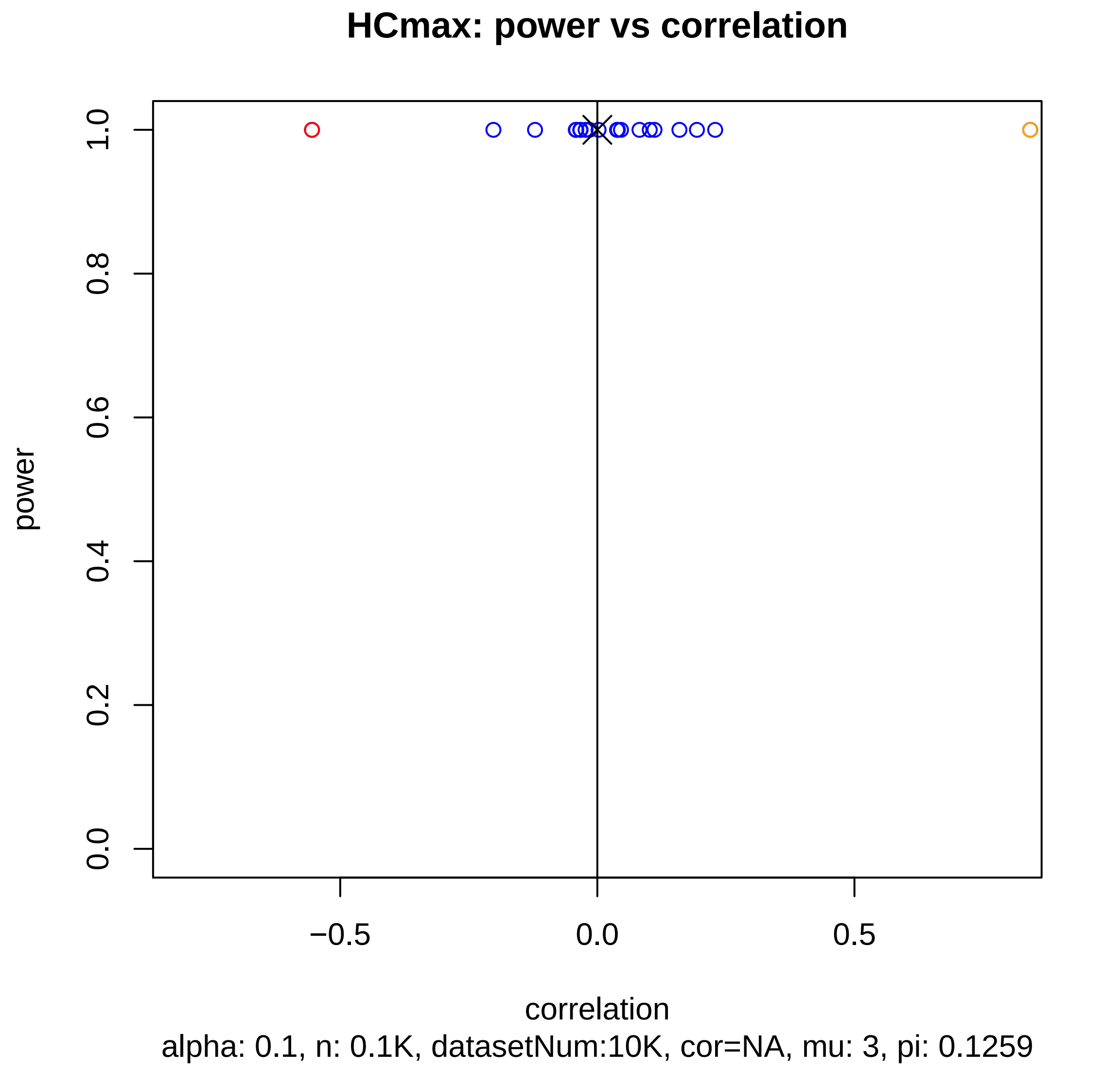
HCmax: power vs correlation
power
1.0
0.8
0.6
0.4
0.2
0.0
−0.5
0.0
0.5
correlation
alpha: 0.1, n: 0.1K, datasetNum:10K, cor=NA, mu: 3, pi: 0.1259

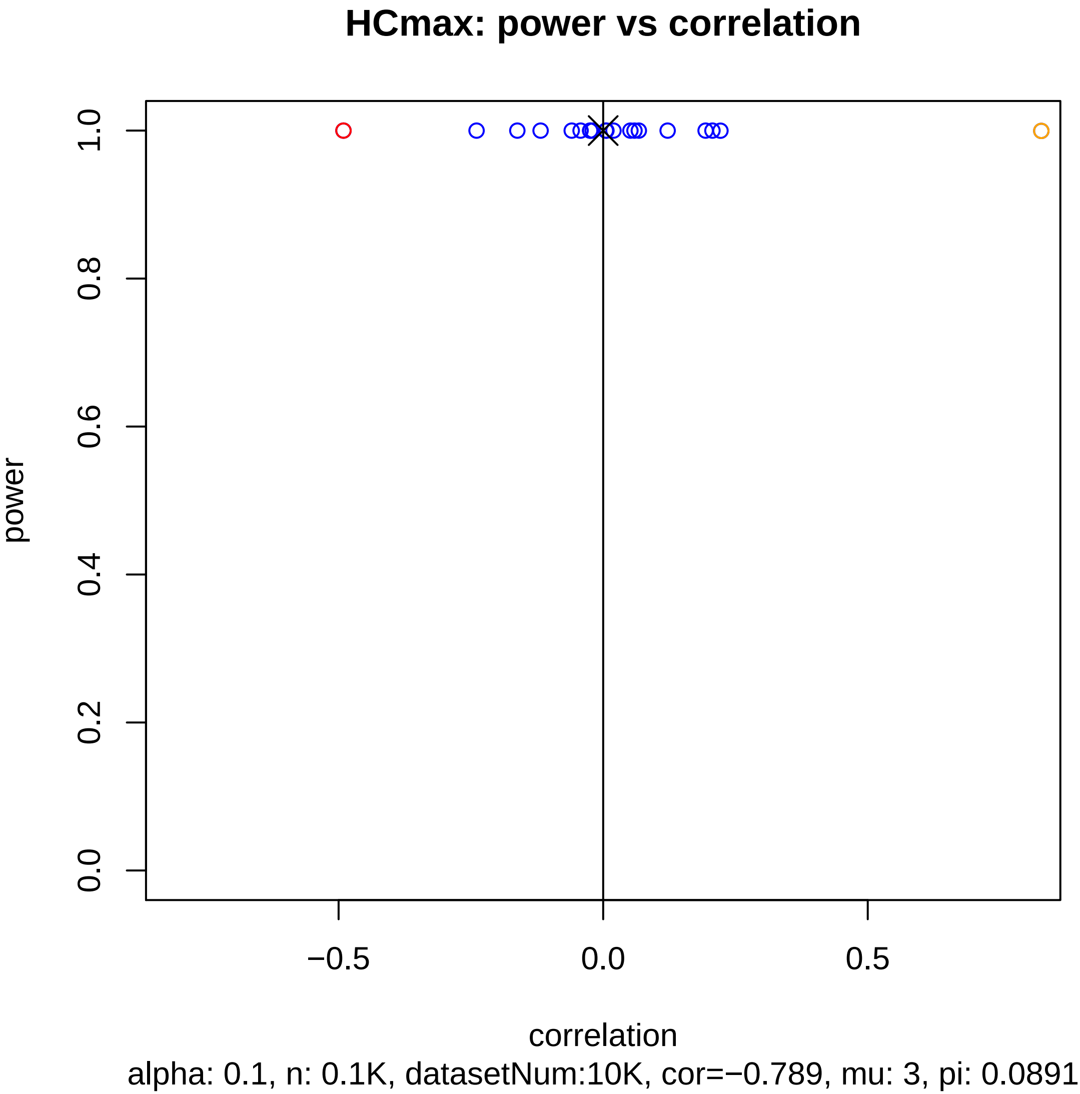
HCmax: power vs correlation
power
1.0
0.8
0.6
0.4
0.2
0.0
−0.5
0.0
0.5
correlation
alpha: 0.1, n: 0.1K, datasetNum:10K, cor=−0.789, mu: 3, pi: 0.0891

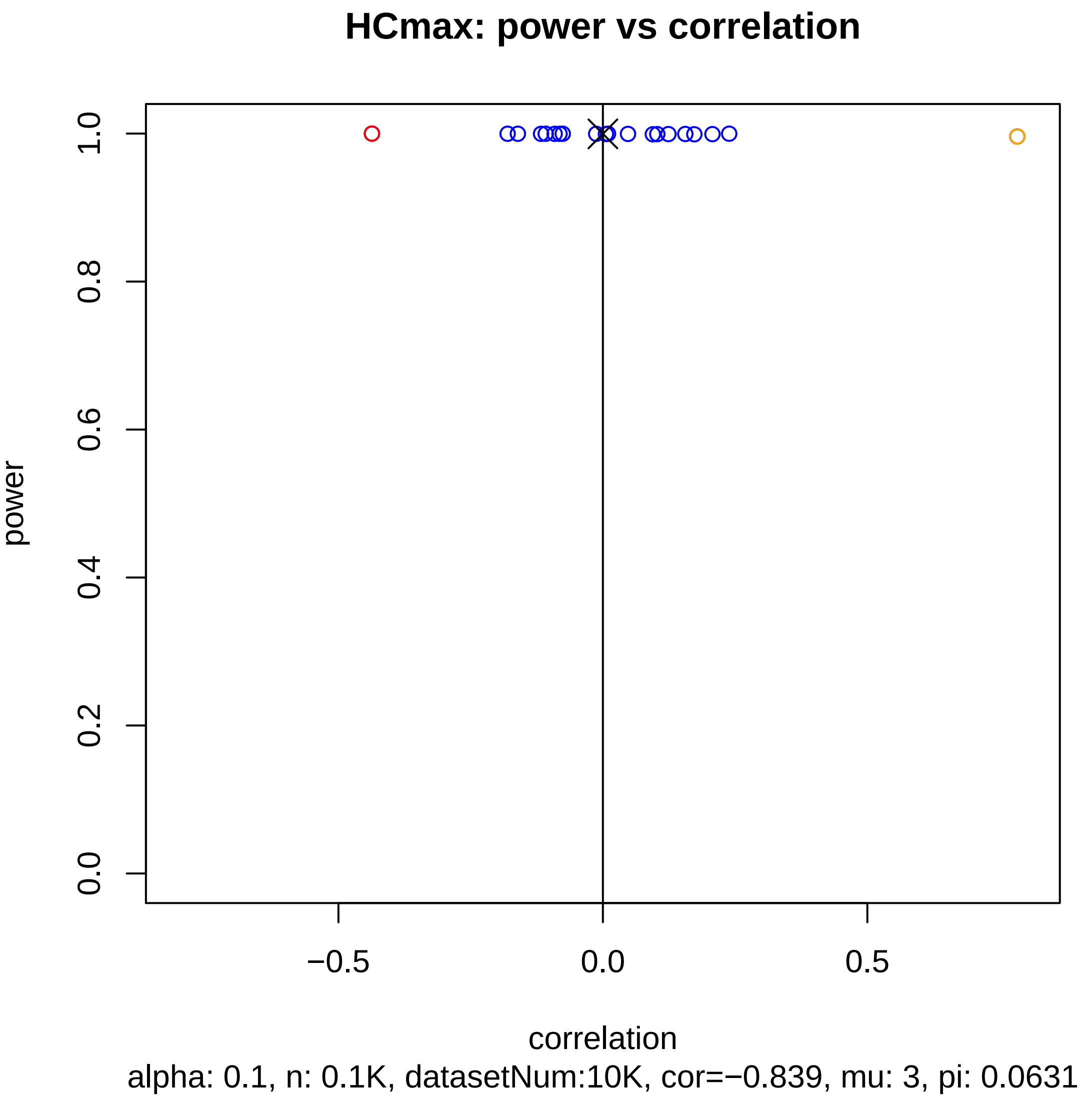

HCmax: power vs correlation
power
1.0
0.8
0.6
0.4
0.2
0.0
−0.5
0.0
0.5
correlation
alpha: 0.1, n: 0.1K, datasetNum:10K, cor=−0.839, mu: 3, pi: 0.0631

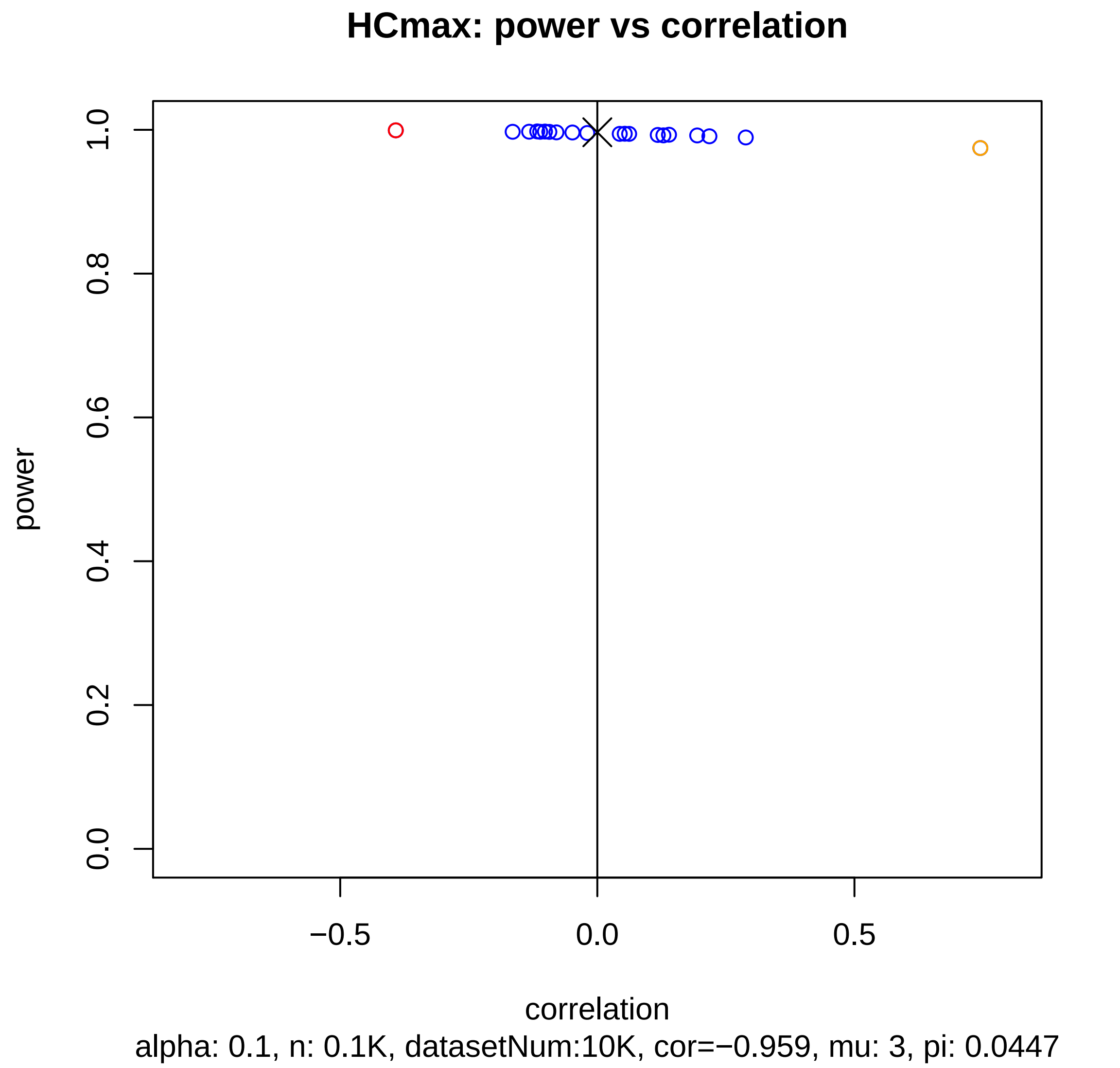
HCmax: power vs correlation
power
1.0
0.8
0.6
0.4
0.2
0.0
−0.5
0.0
0.5
correlation
alpha: 0.1, n: 0.1K, datasetNum:10K, cor=−0.959, mu: 3, pi: 0.0447

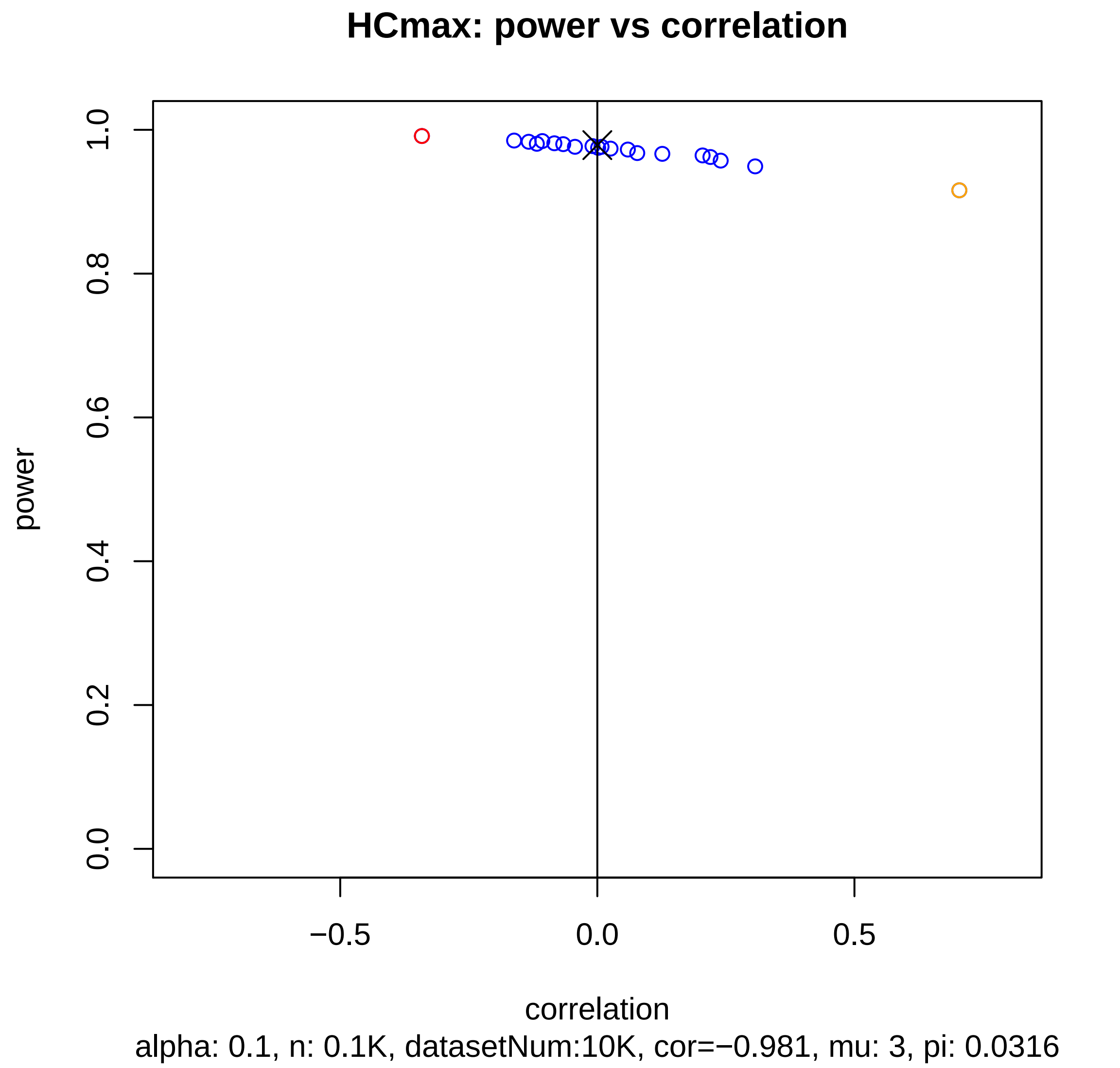
HCmax: power vs correlation
power
1.0
0.8
0.6
0.4
0.2
0.0
−0.5
0.0
0.5
correlation
alpha: 0.1, n: 0.1K, datasetNum:10K, cor=−0.981, mu: 3, pi: 0.0316

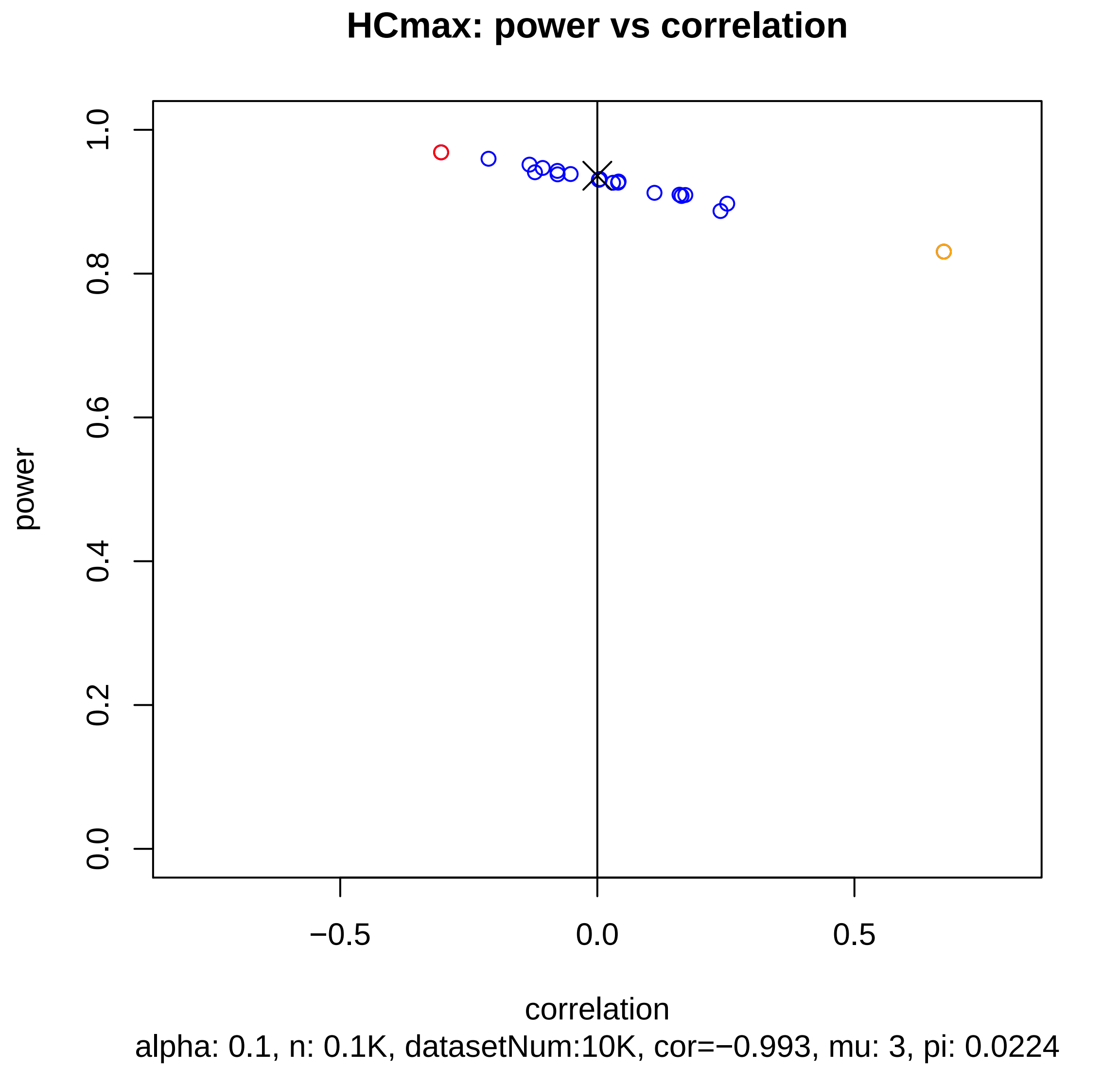

HCmax: power vs correlation
power
1.0
0.8
0.6
0.4
0.2
0.0
−0.5
0.0
0.5
correlation
alpha: 0.1, n: 0.1K, datasetNum:10K, cor=−0.993, mu: 3, pi: 0.0224

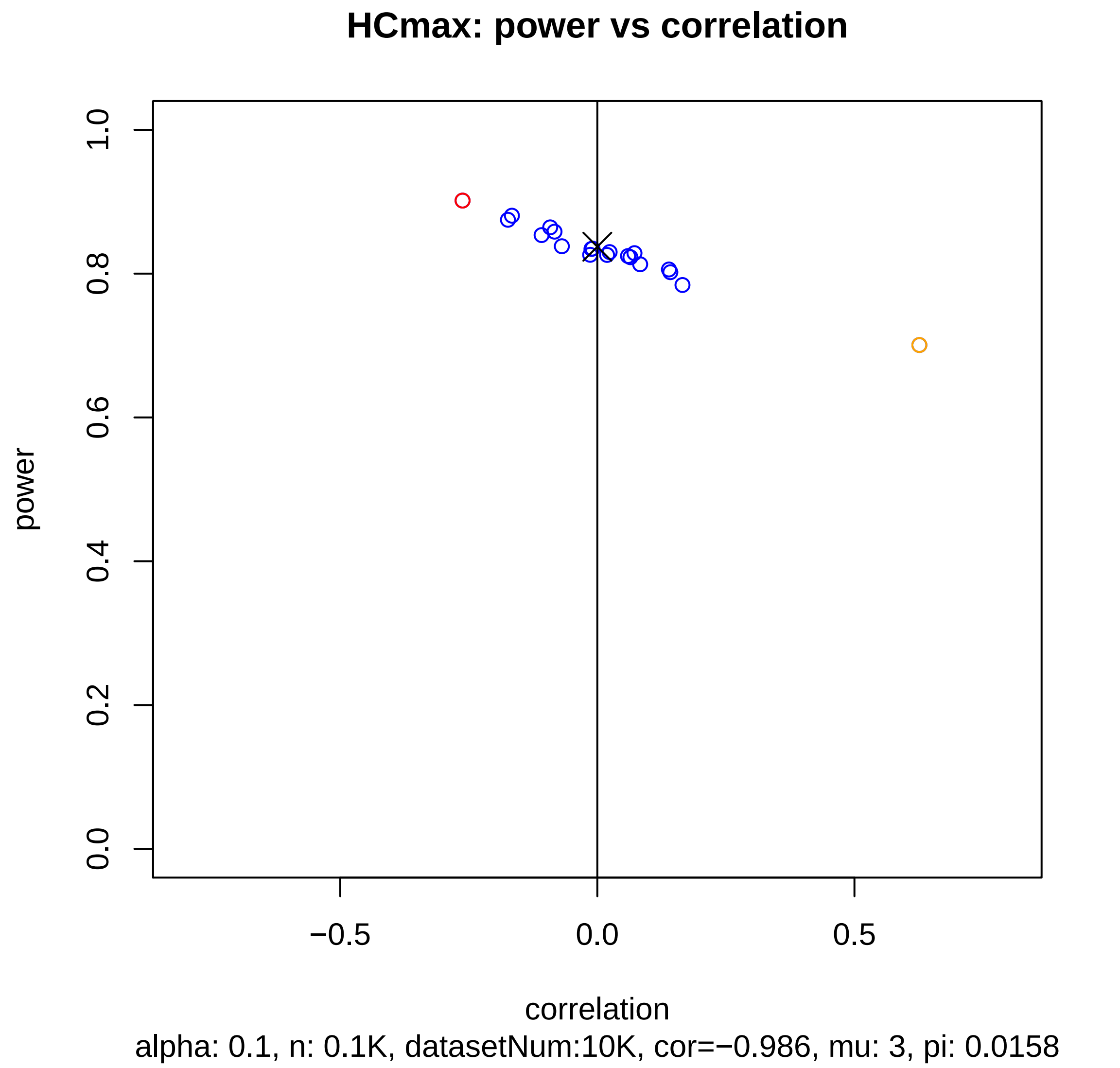
HCmax: power vs correlation
power
1.0
0.8
0.6
0.4
0.2
0.0
−0.5
0.0
0.5
correlation
alpha: 0.1, n: 0.1K, datasetNum:10K, cor=−0.986, mu: 3, pi: 0.0158

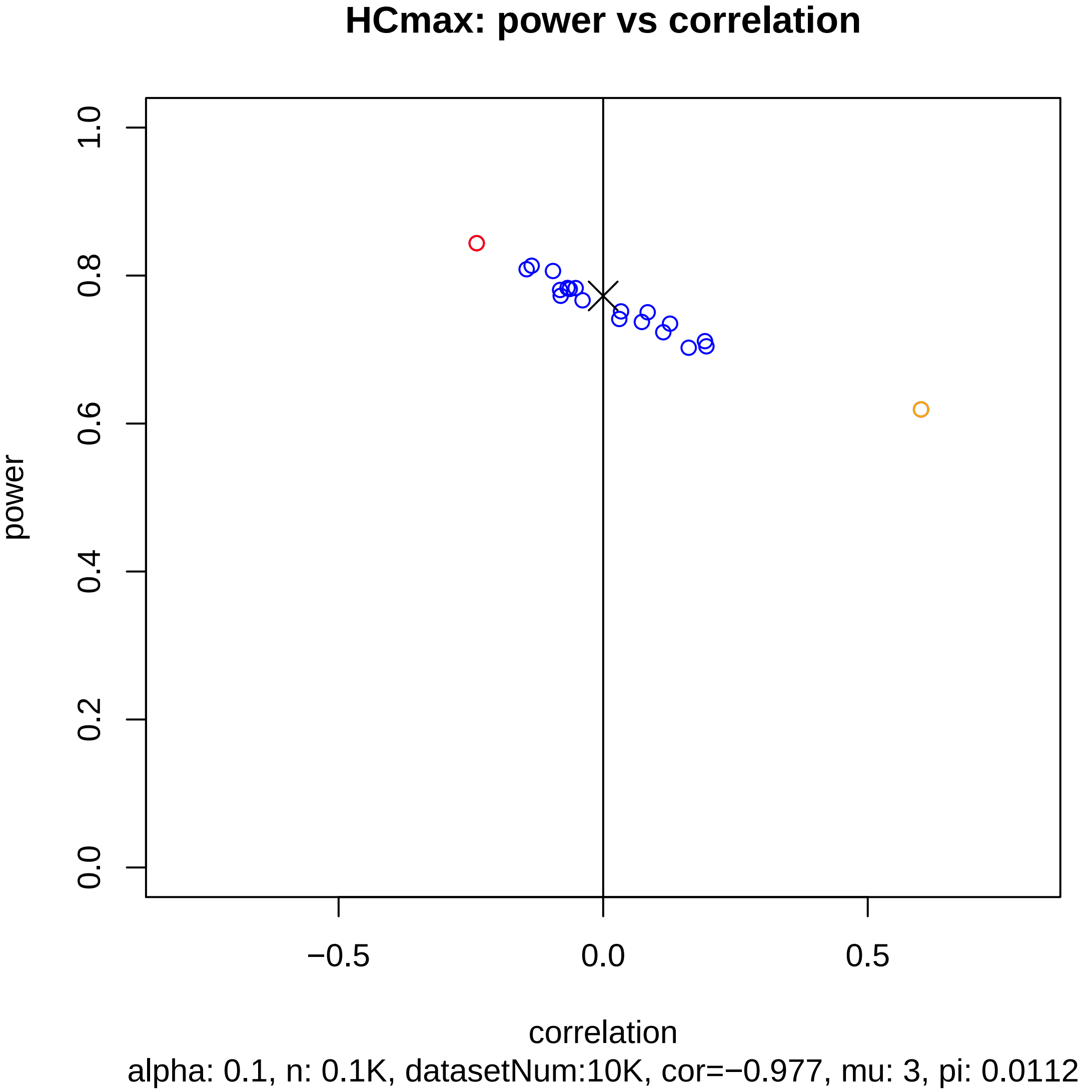
HCmax: power vs correlation
power
1.0
0.8
0.6
0.4
0.2
0.0
−0.5
0.0
0.5
correlation
alpha: 0.1, n: 0.1K, datasetNum:10K, cor=−0.977, mu: 3, pi: 0.0112

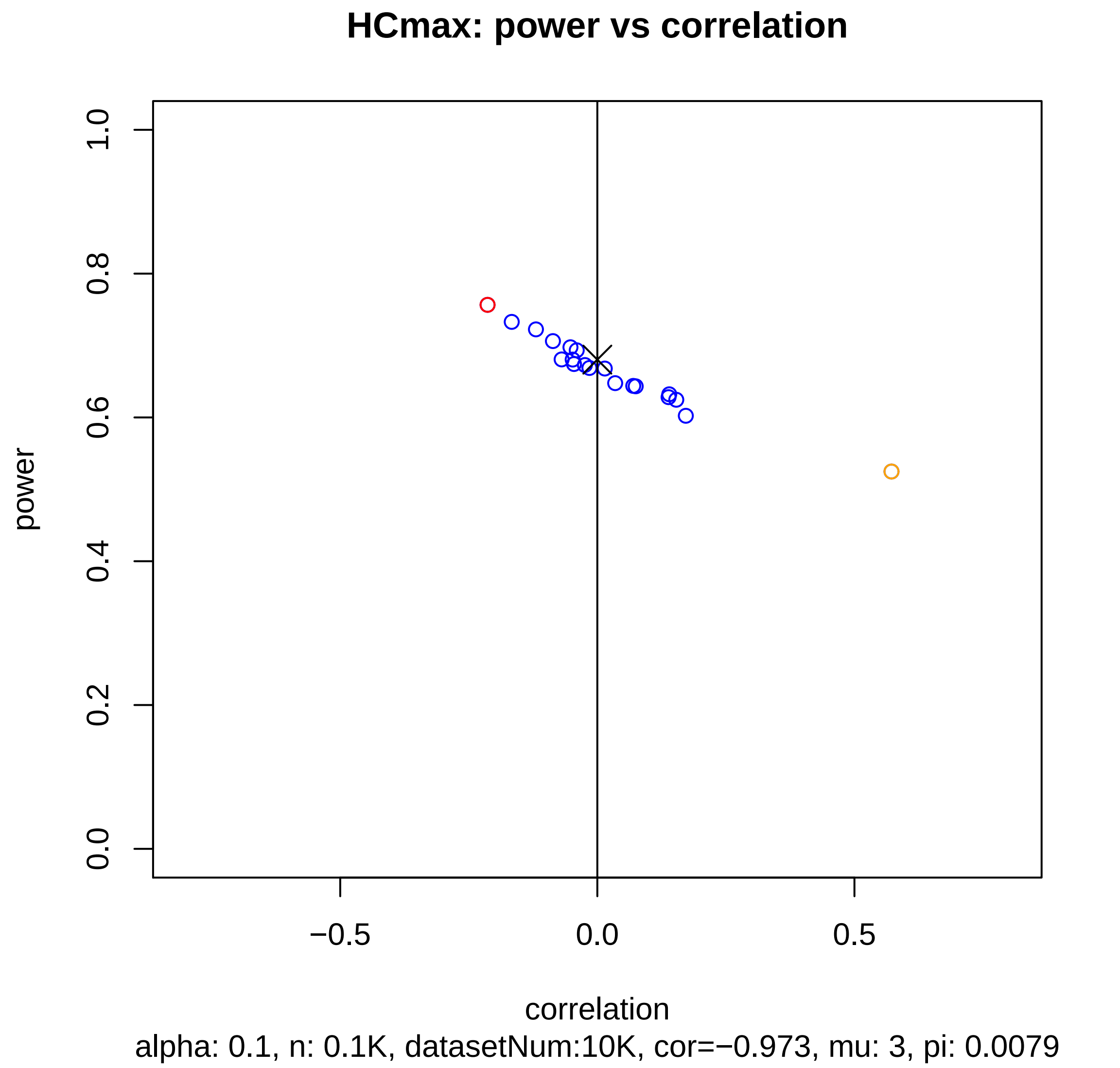

HCmax: power vs correlation
power
1.0
0.8
0.6
0.4
0.2
0.0
−0.5
0.0
0.5
correlation
alpha: 0.1, n: 0.1K, datasetNum:10K, cor=−0.973, mu: 3, pi: 0.0079

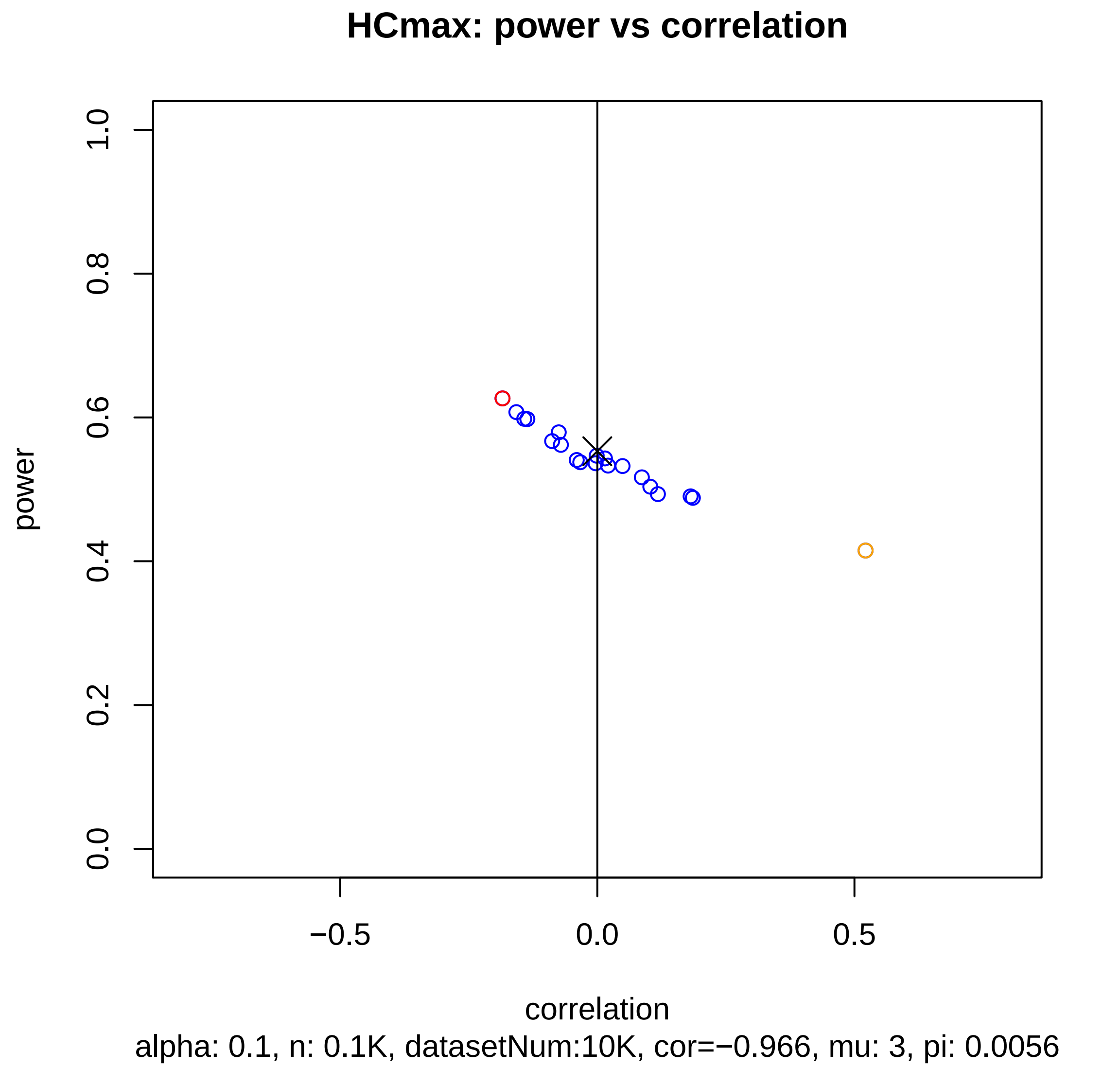
HCmax: power vs correlation
power
1.0
0.8
0.6
0.4
0.2
0.0
−0.5
0.0
0.5
correlation
alpha: 0.1, n: 0.1K, datasetNum:10K, cor=−0.966, mu: 3, pi: 0.0056

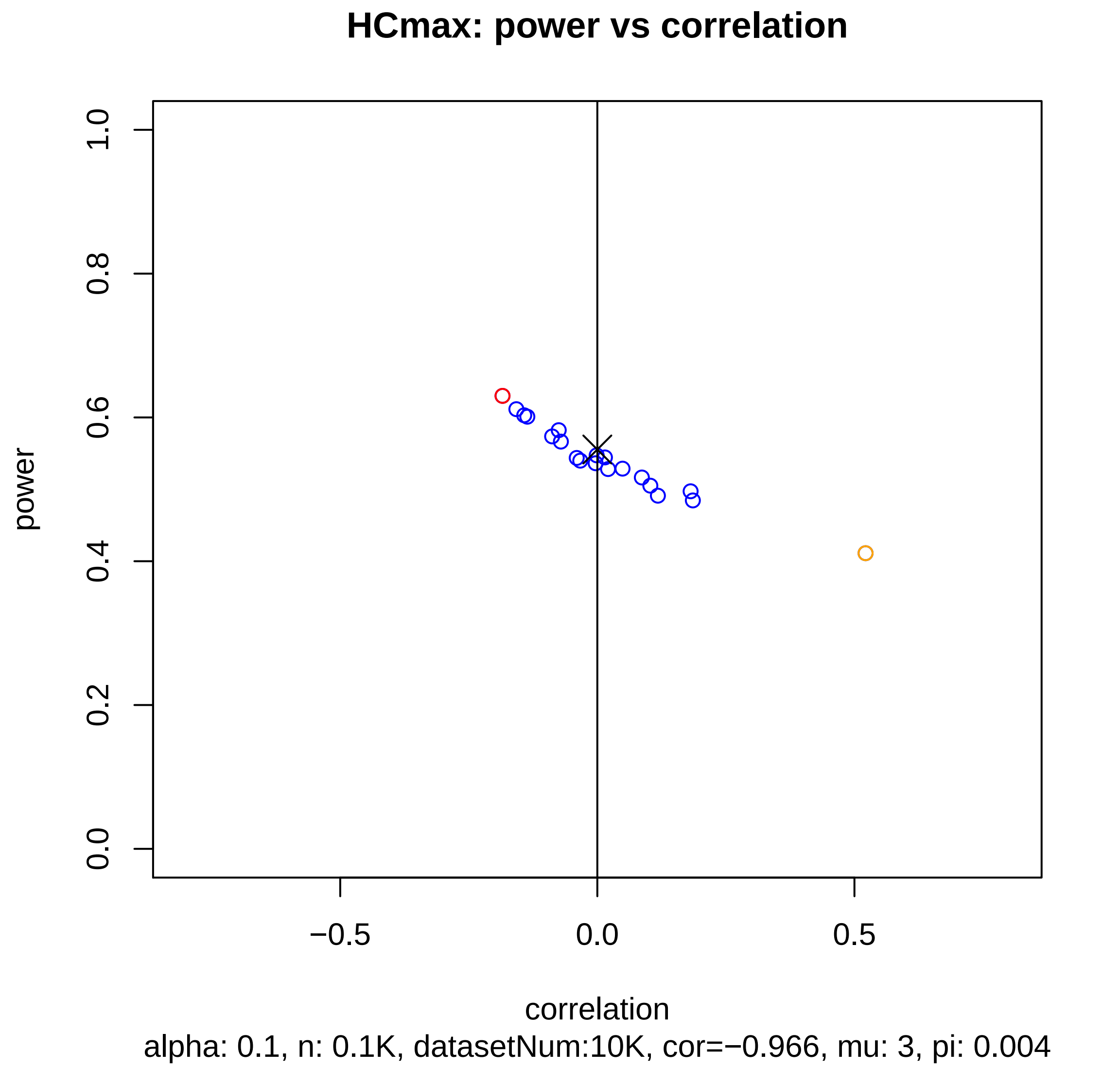
HCmax: power vs correlation
power
1.0
0.8
0.6
0.4
0.2
0.0
−0.5
0.0
0.5
correlation
alpha: 0.1, n: 0.1K, datasetNum:10K, cor=−0.966, mu: 3, pi: 0.004

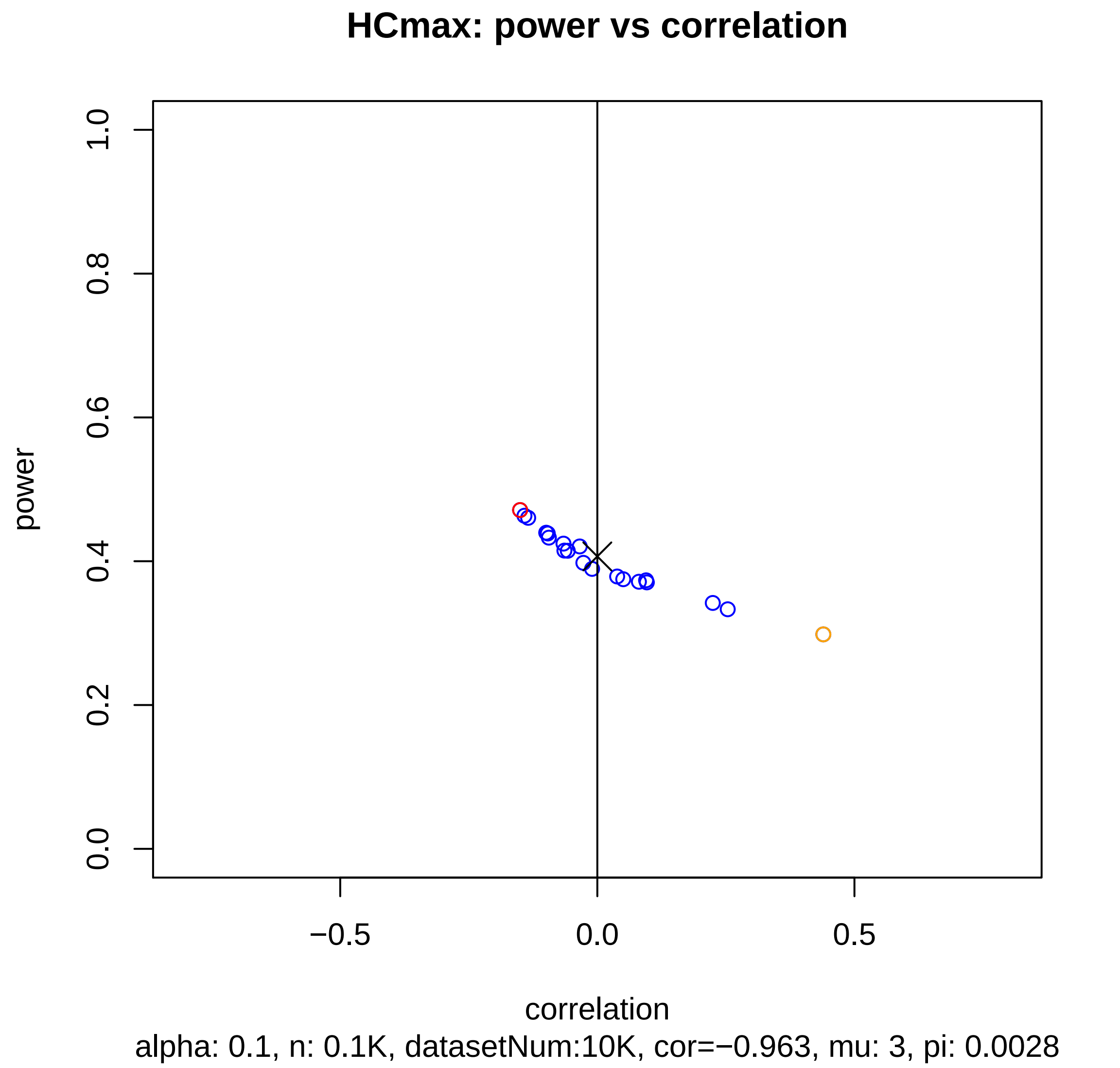
HCmax: power vs correlation
power
1.0
0.8
0.6
0.4
0.2
0.0
−0.5
0.0
0.5
correlation
alpha: 0.1, n: 0.1K, datasetNum:10K, cor=−0.963, mu: 3, pi: 0.0028

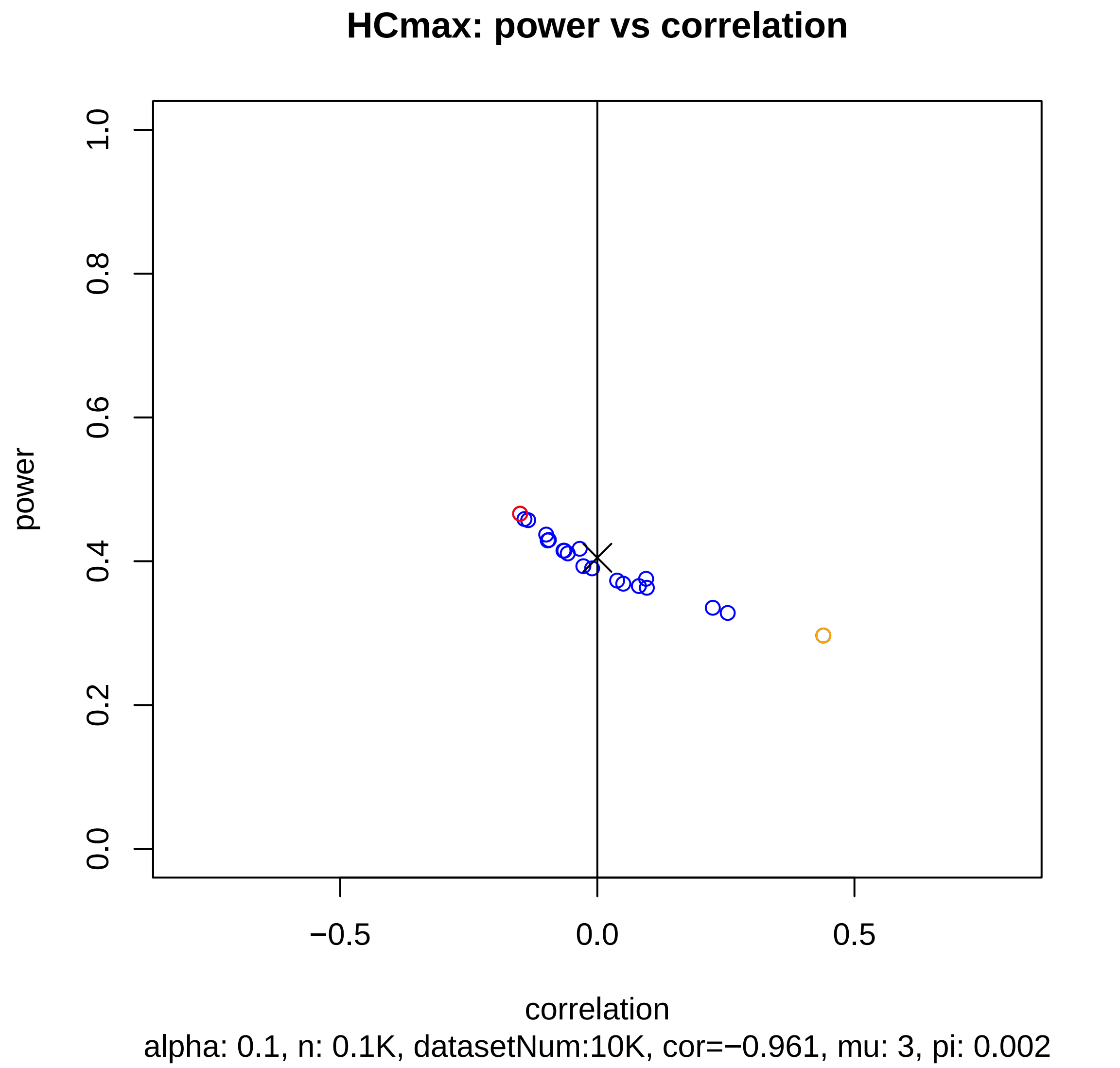
HCmax: power vs correlation
power
1.0
0.8
0.6
0.4
0.2
0.0
−0.5
0.0
0.5
correlation
alpha: 0.1, n: 0.1K, datasetNum:10K, cor=−0.961, mu: 3, pi: 0.002

**HCmax: power vs correlation**

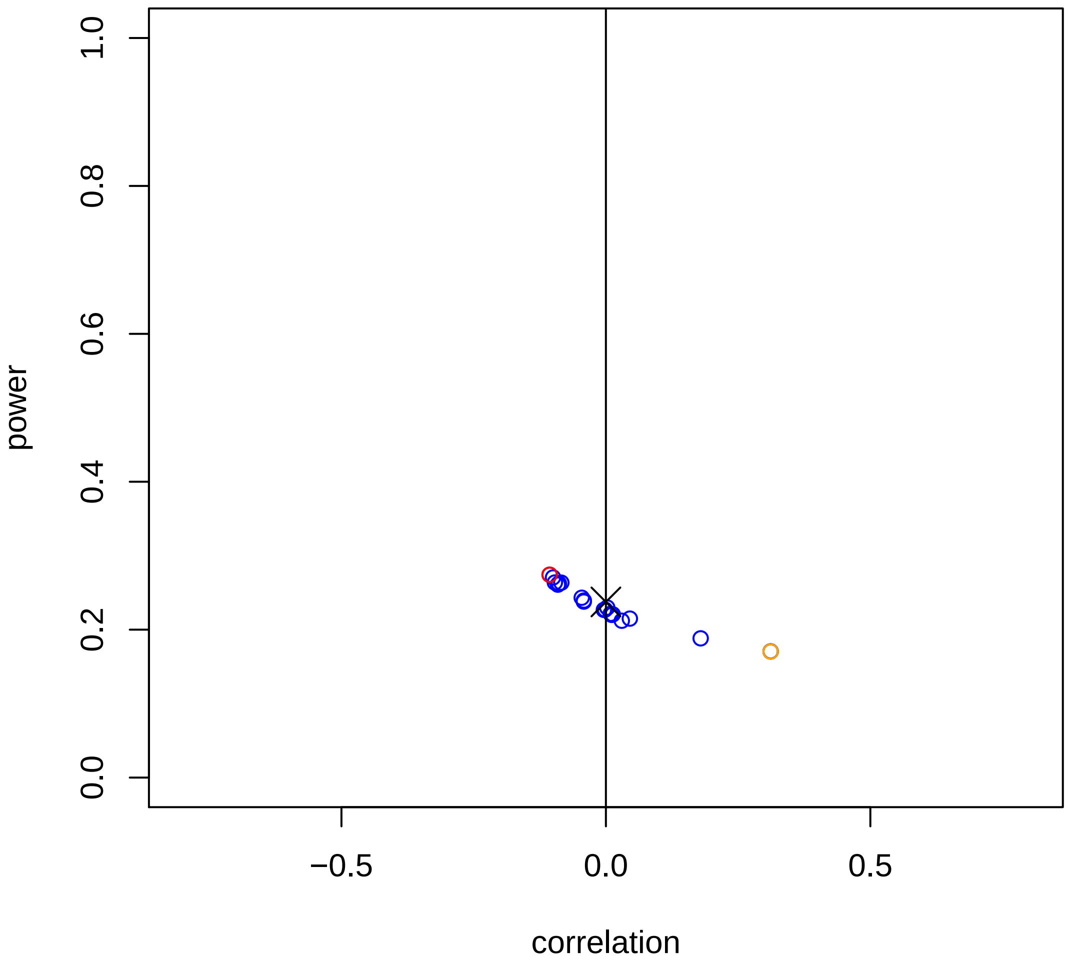


alpha: 0.1, n: 0.1K, datasetNum:10K, cor=−0.952, mu: 3, pi: 0.0014

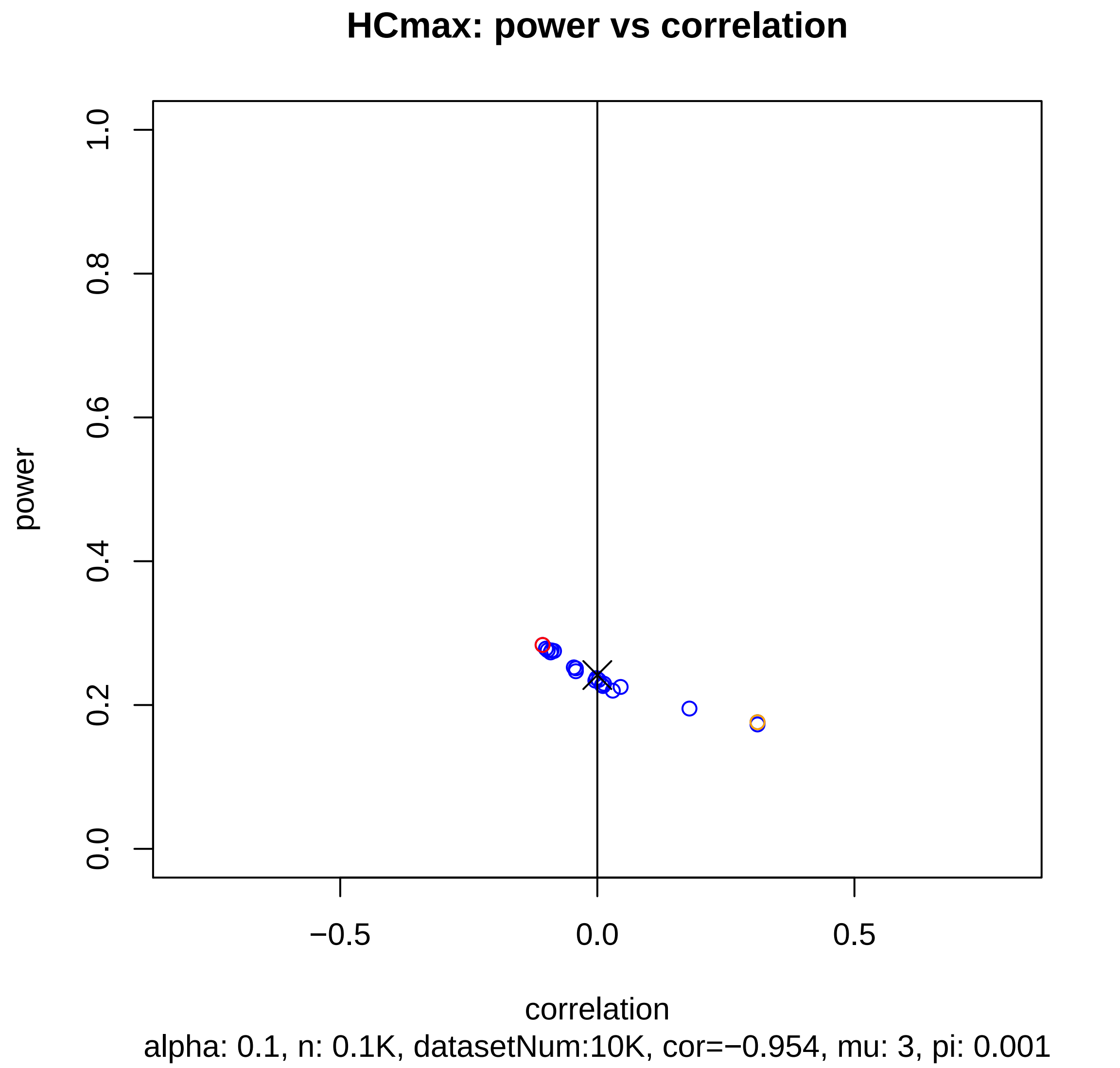
HCmax: power vs correlation
power
1.0
0.8
0.6
0.4
0.2
0.0
−0.5
0.0
0.5
correlation
alpha: 0.1, n: 0.1K, datasetNum:10K, cor=−0.954, mu: 3, pi: 0.001

**HCmax: power vs correlation**

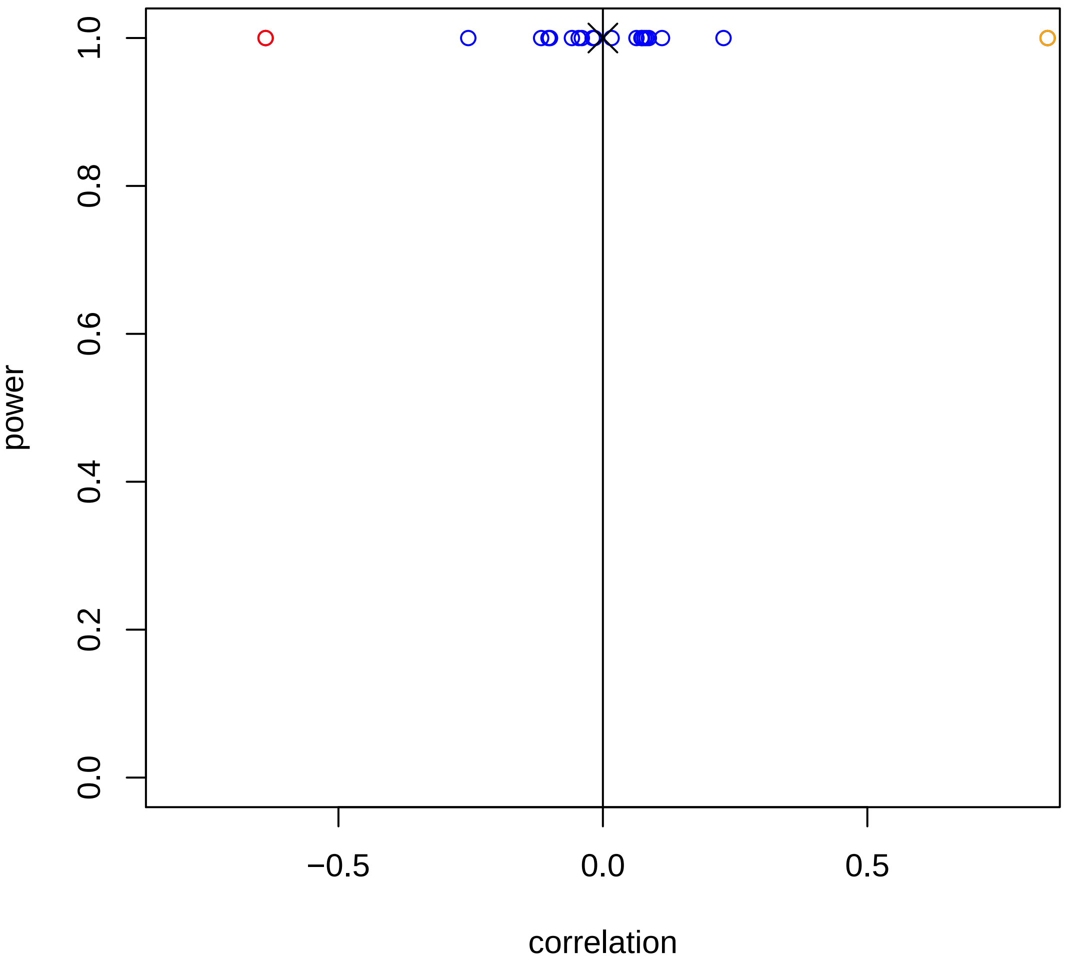


alpha: 0.1, n: 0.1K, datasetNum:10K, cor=NA, mu: 3.11, pi: 0.1778

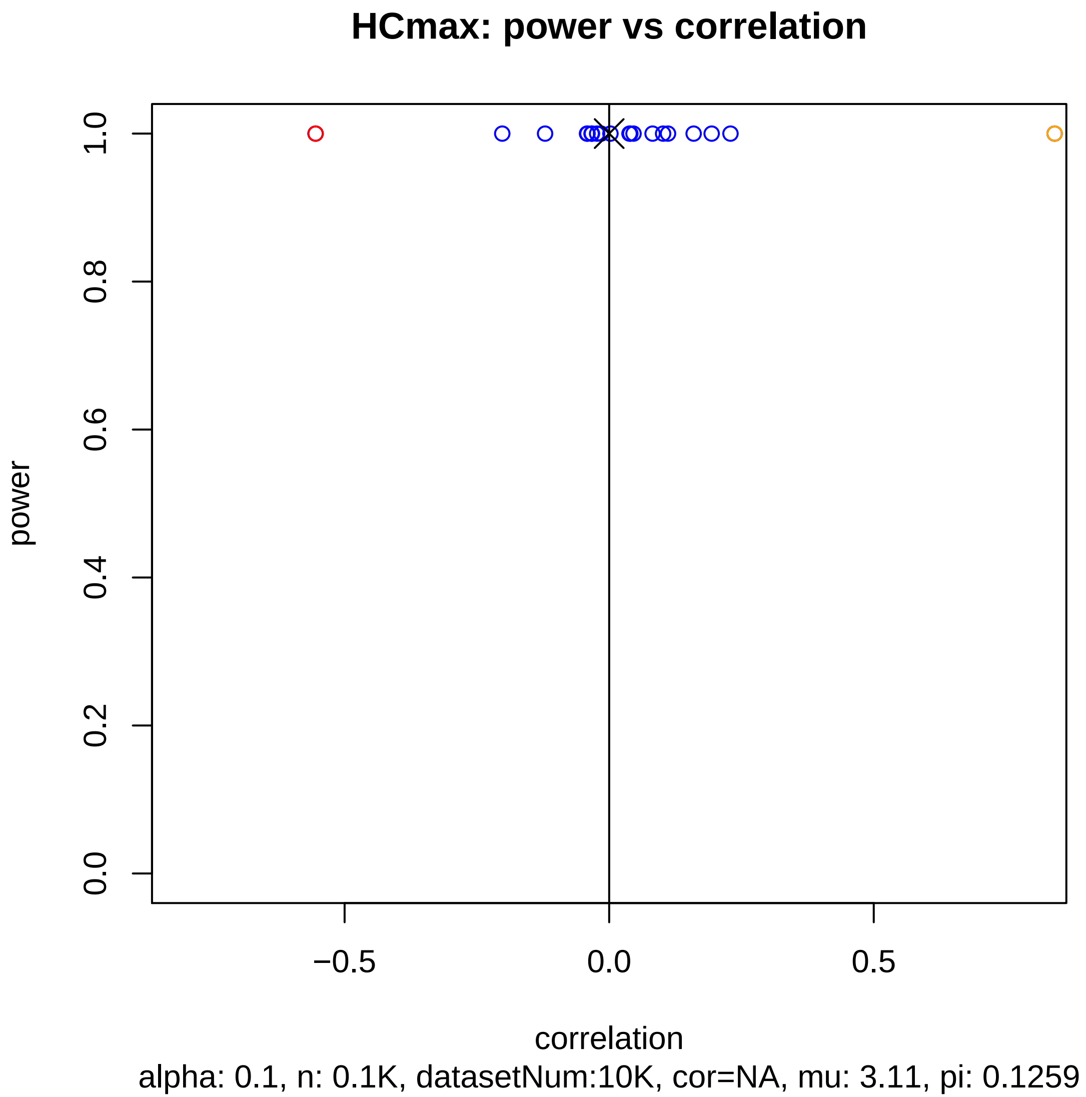
HCmax: power vs correlation
power
1.0
0.8
0.6
0.4
0.2
0.0
−0.5
0.0
0.5
correlation
alpha: 0.1, n: 0.1K, datasetNum:10K, cor=NA, mu: 3.11, pi: 0.1259

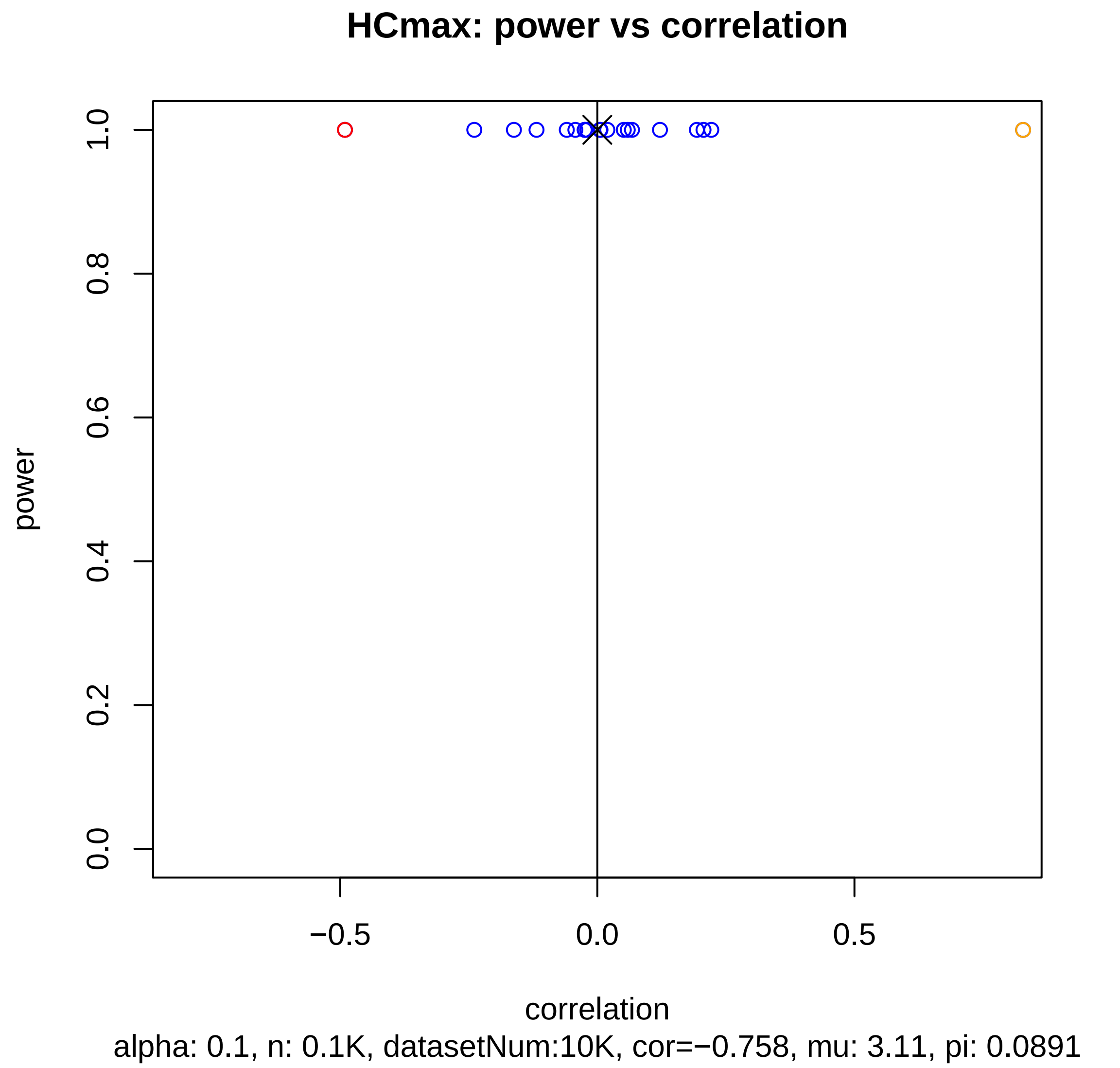
HCmax: power vs correlation
power
1.0
0.8
0.6
0.4
0.2
0.0
−0.5
0.0
0.5
correlation
alpha: 0.1, n: 0.1K, datasetNum:10K, cor=−0.758, mu: 3.11, pi: 0.0891

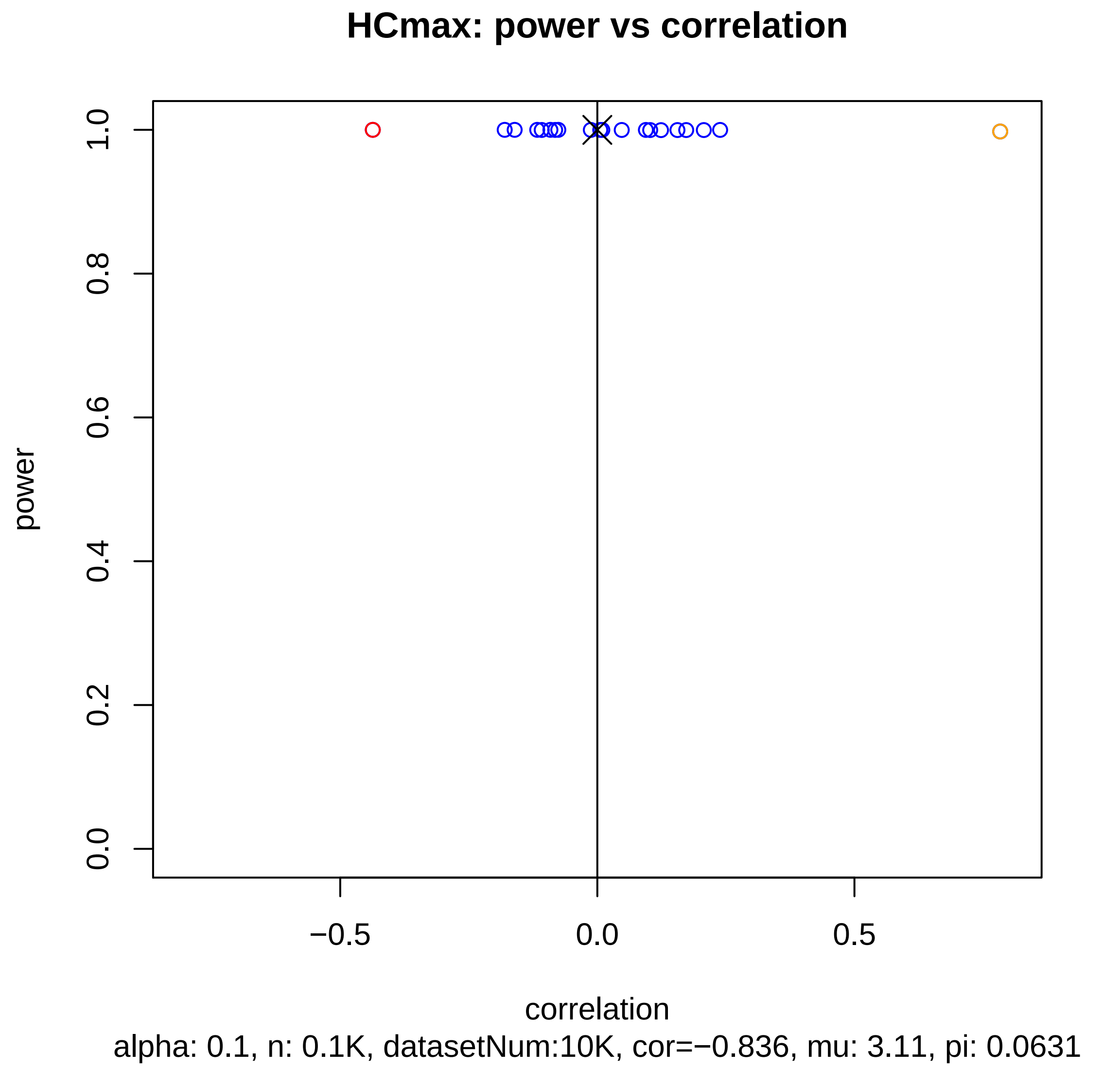
HCmax: power vs correlation
power
1.0
0.8
0.6
0.4
0.2
0.0
−0.5
0.0
0.5
correlation
alpha: 0.1, n: 0.1K, datasetNum:10K, cor=−0.836, mu: 3.11, pi: 0.0631

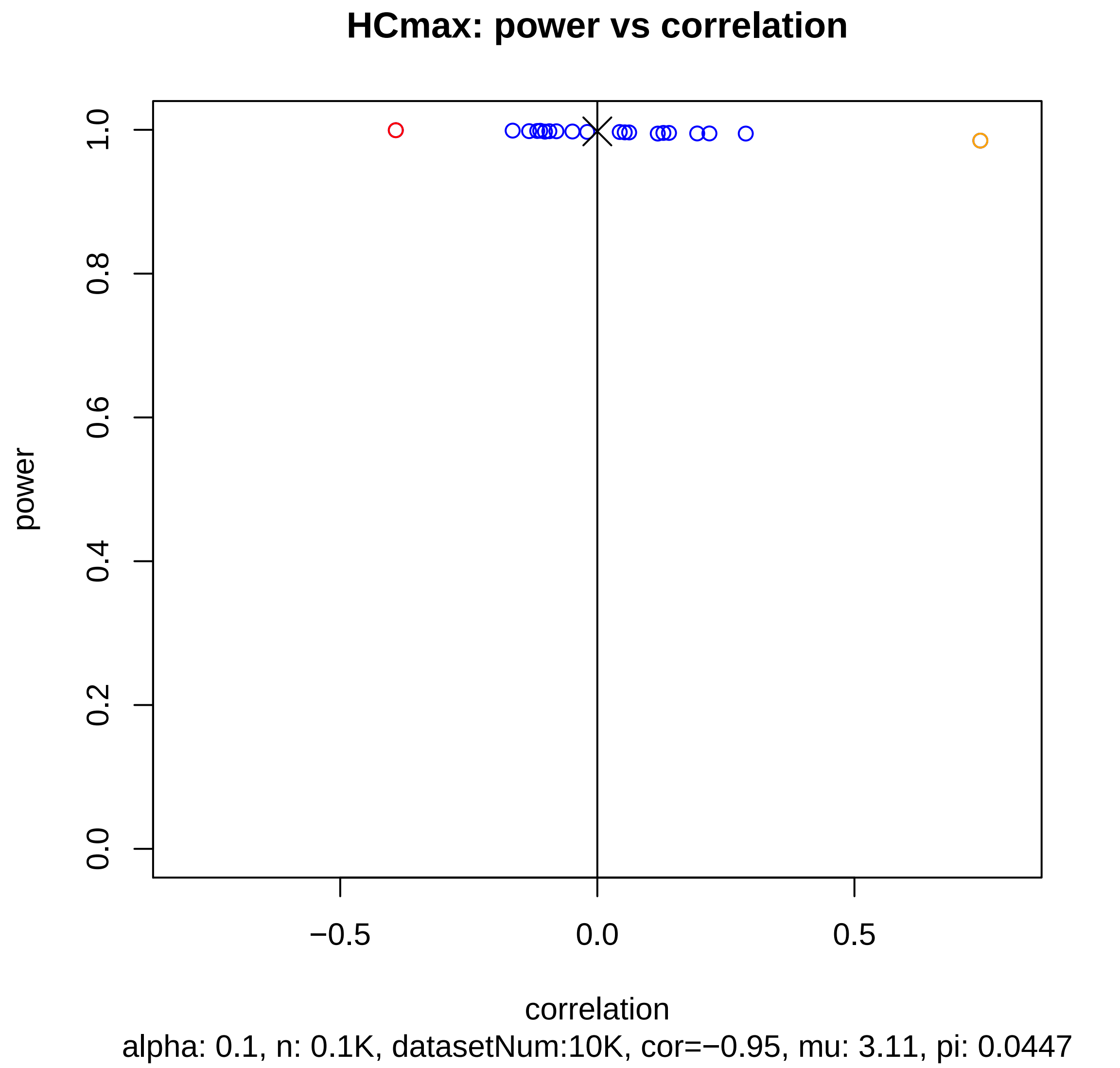
HCmax: power vs correlation
power
1.0
0.8
0.6
0.4
0.2
0.0
−0.5
0.0
0.5
correlation
alpha: 0.1, n: 0.1K, datasetNum:10K, cor=−0.95, mu: 3.11, pi: 0.0447

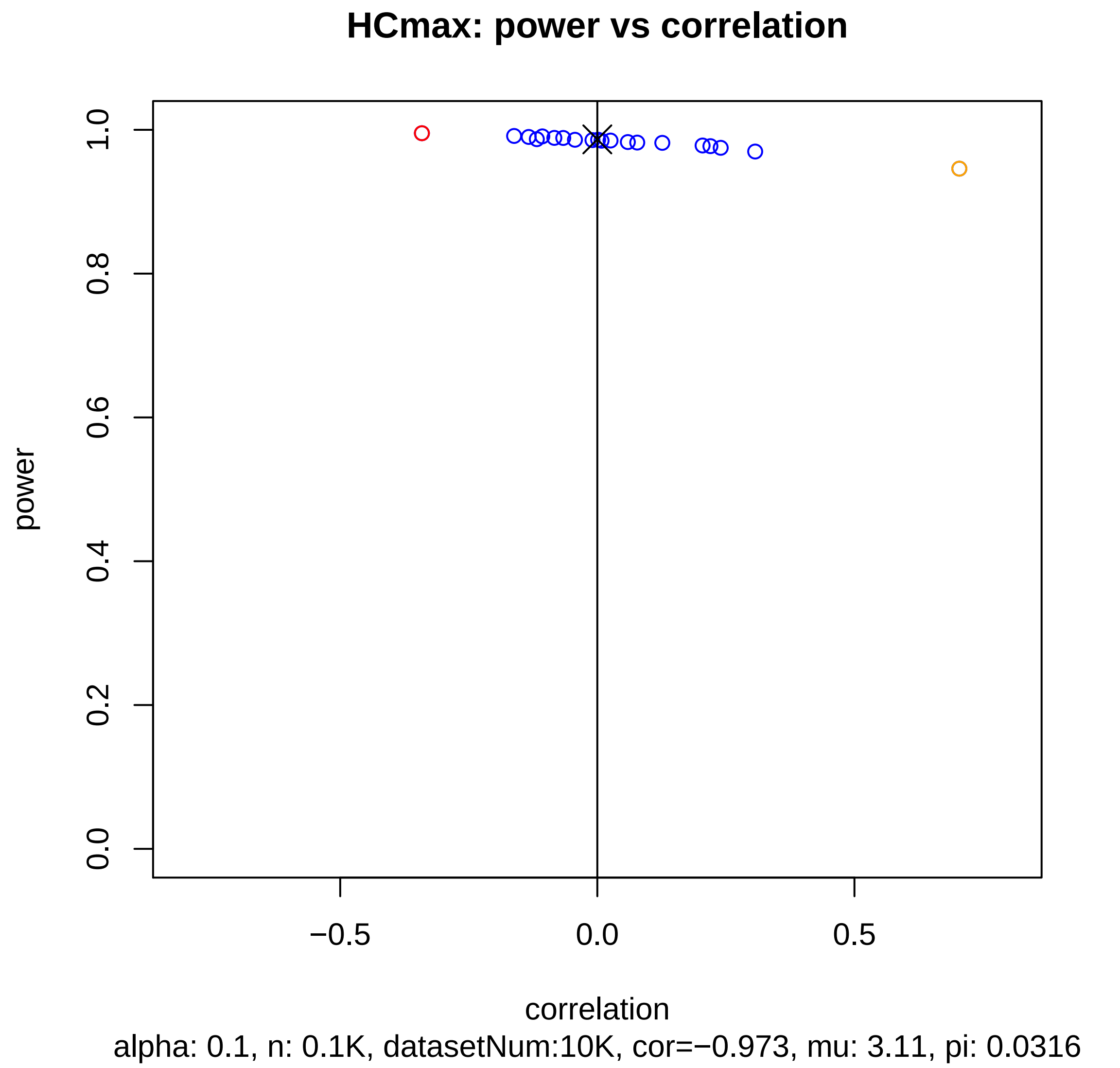
HCmax: power vs correlation
power
1.0
0.8
0.6
0.4
0.2
0.0
−0.5
0.0
0.5
correlation
alpha: 0.1, n: 0.1K, datasetNum:10K, cor=−0.973, mu: 3.11, pi: 0.0316

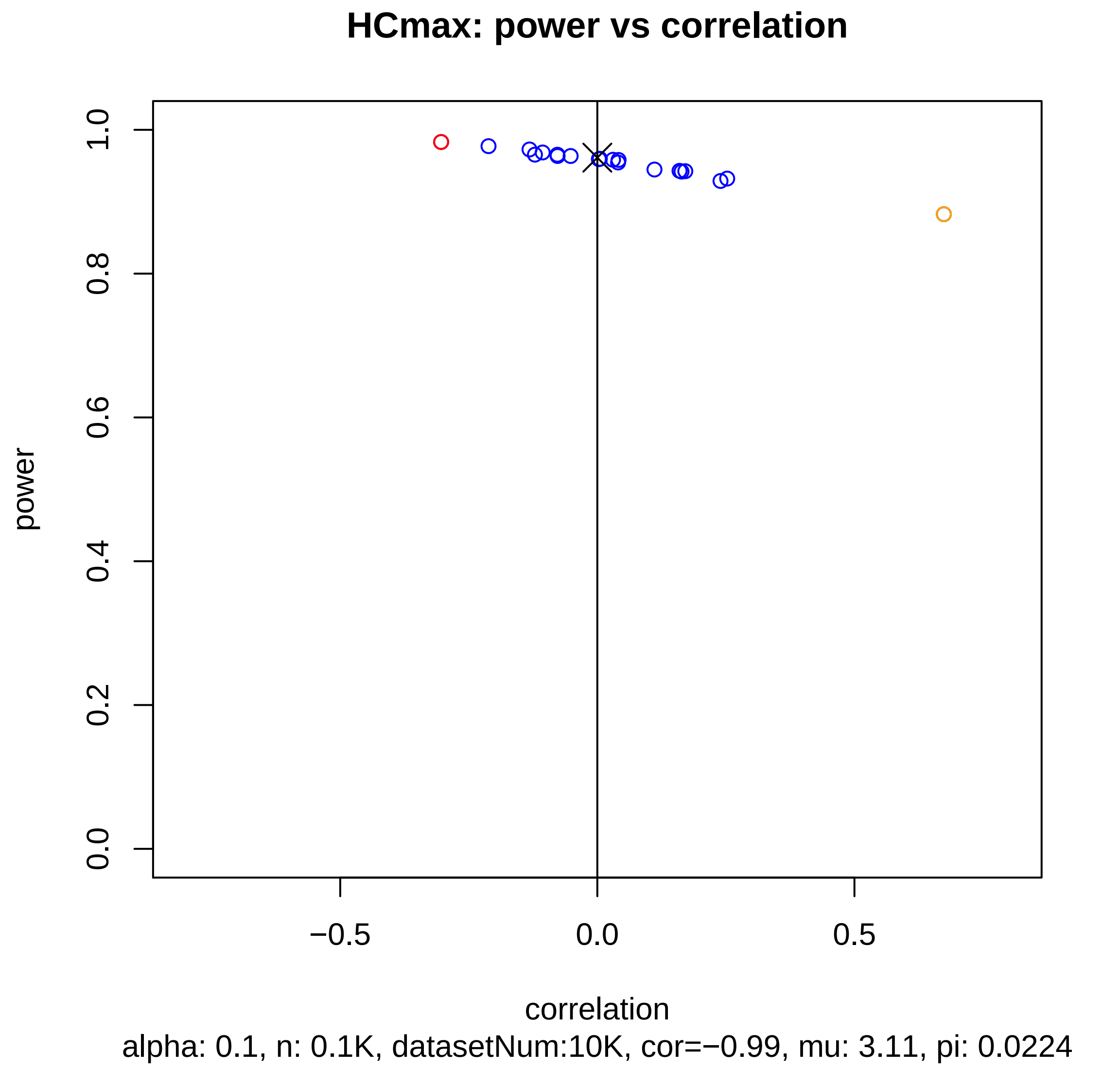
HCmax: power vs correlation
power
1.0
0.8
0.6
0.4
0.2
0.0
−0.5
0.0
0.5
correlation
alpha: 0.1, n: 0.1K, datasetNum:10K, cor=−0.99, mu: 3.11, pi: 0.0224

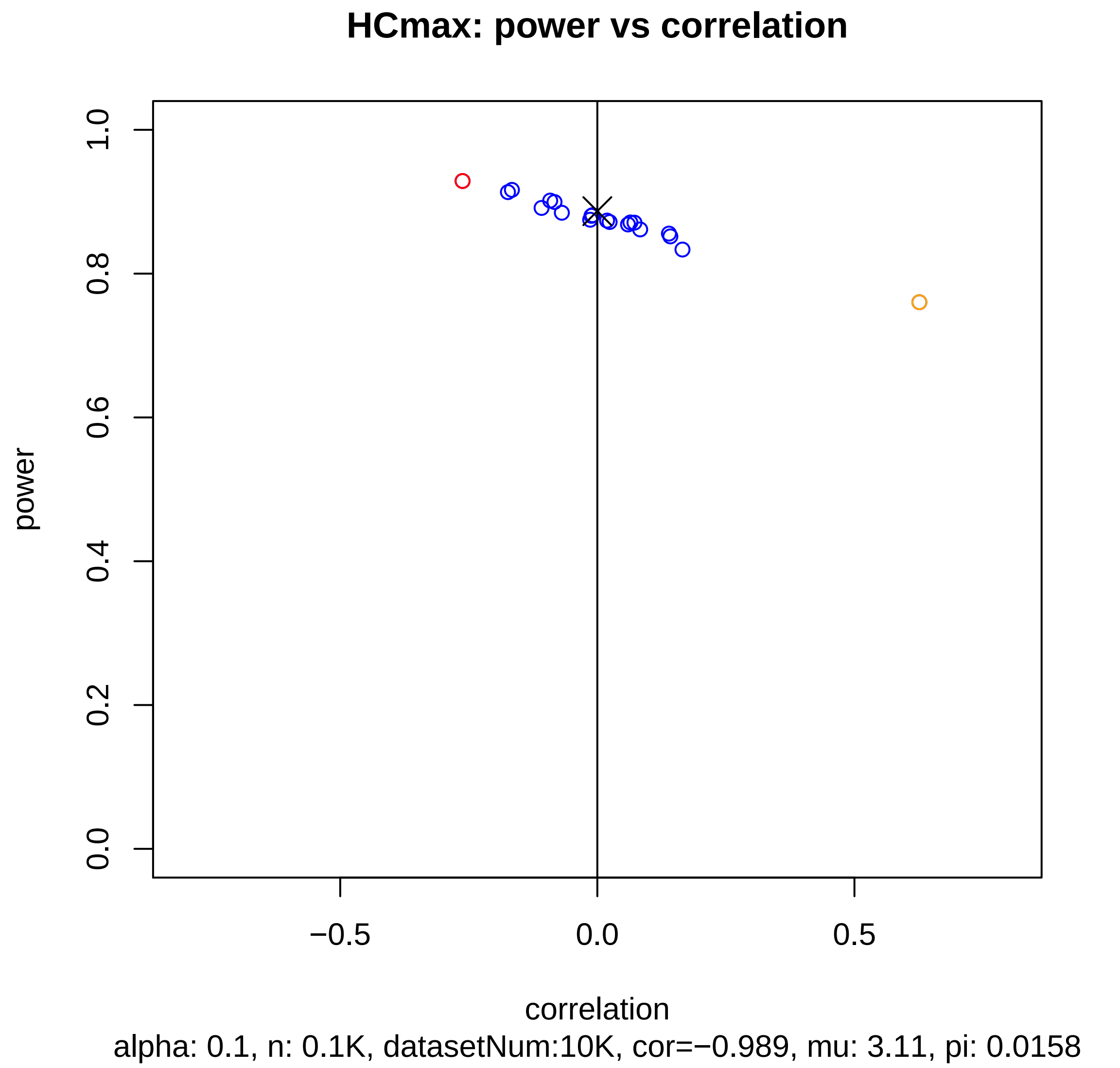

HCmax: power vs correlation
power
1.0
0.8
0.6
0.4
0.2
0.0
−0.5
0.0
0.5
correlation
alpha: 0.1, n: 0.1K, datasetNum:10K, cor=−0.989, mu: 3.11, pi: 0.0158

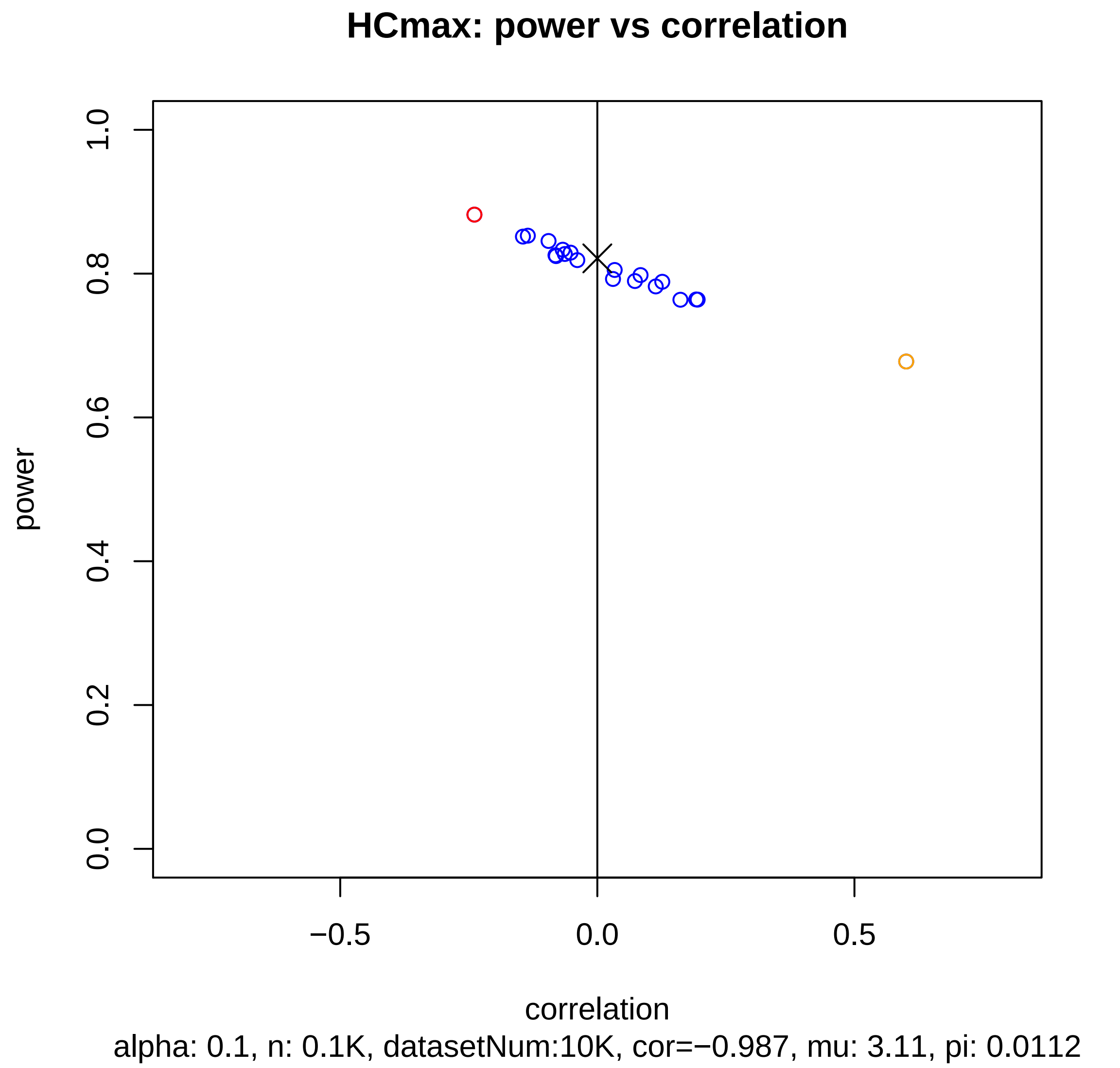
HCmax: power vs correlation
power
1.0
0.8
0.6
0.4
0.2
0.0
−0.5
0.0
0.5
correlation
alpha: 0.1, n: 0.1K, datasetNum:10K, cor=−0.987, mu: 3.11, pi: 0.0112

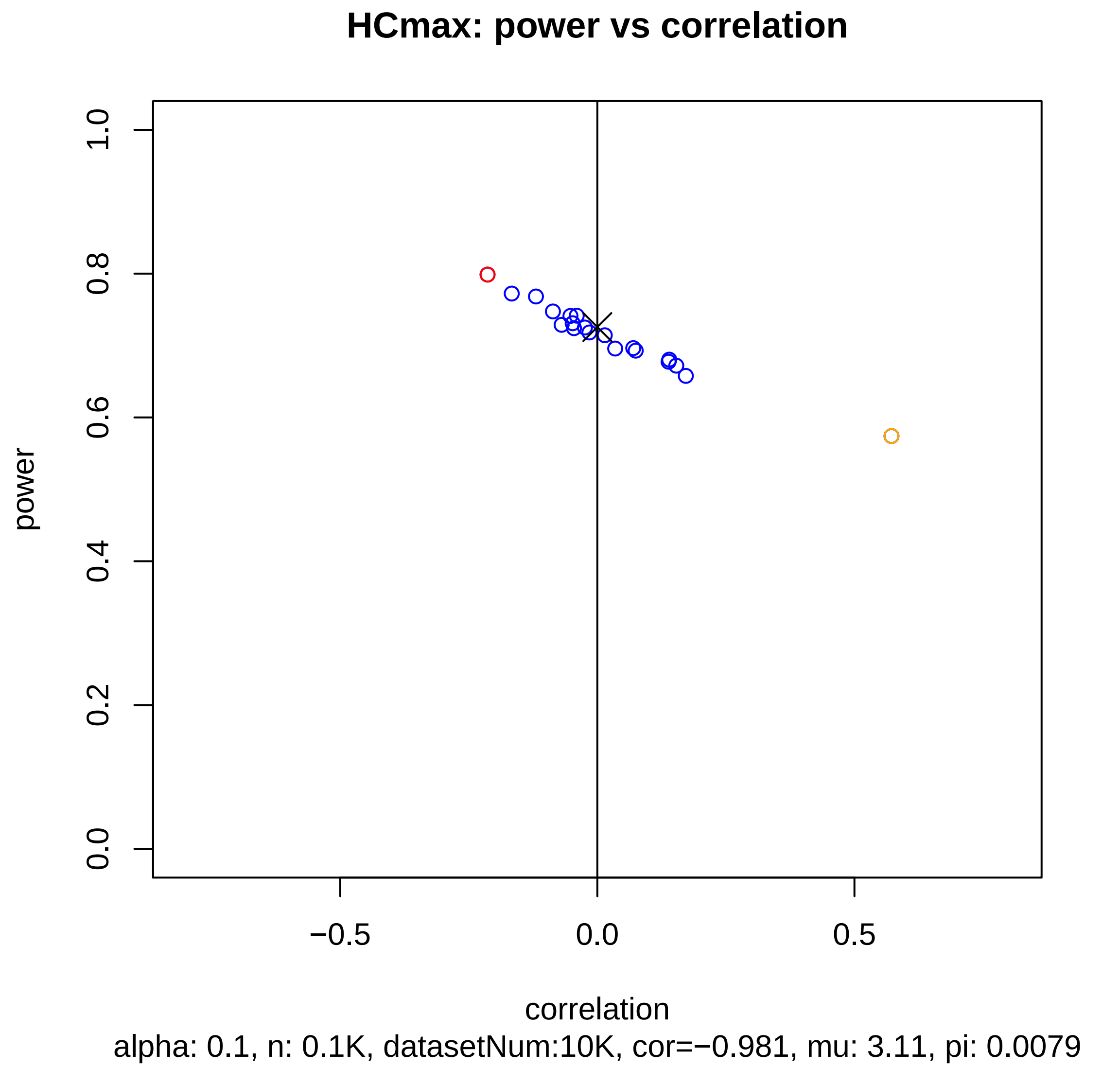
HCmax: power vs correlation
power
1.0
0.8
0.6
0.4
0.2
0.0
−0.5
0.0
0.5
correlation
alpha: 0.1, n: 0.1K, datasetNum:10K, cor=−0.981, mu: 3.11, pi: 0.0079

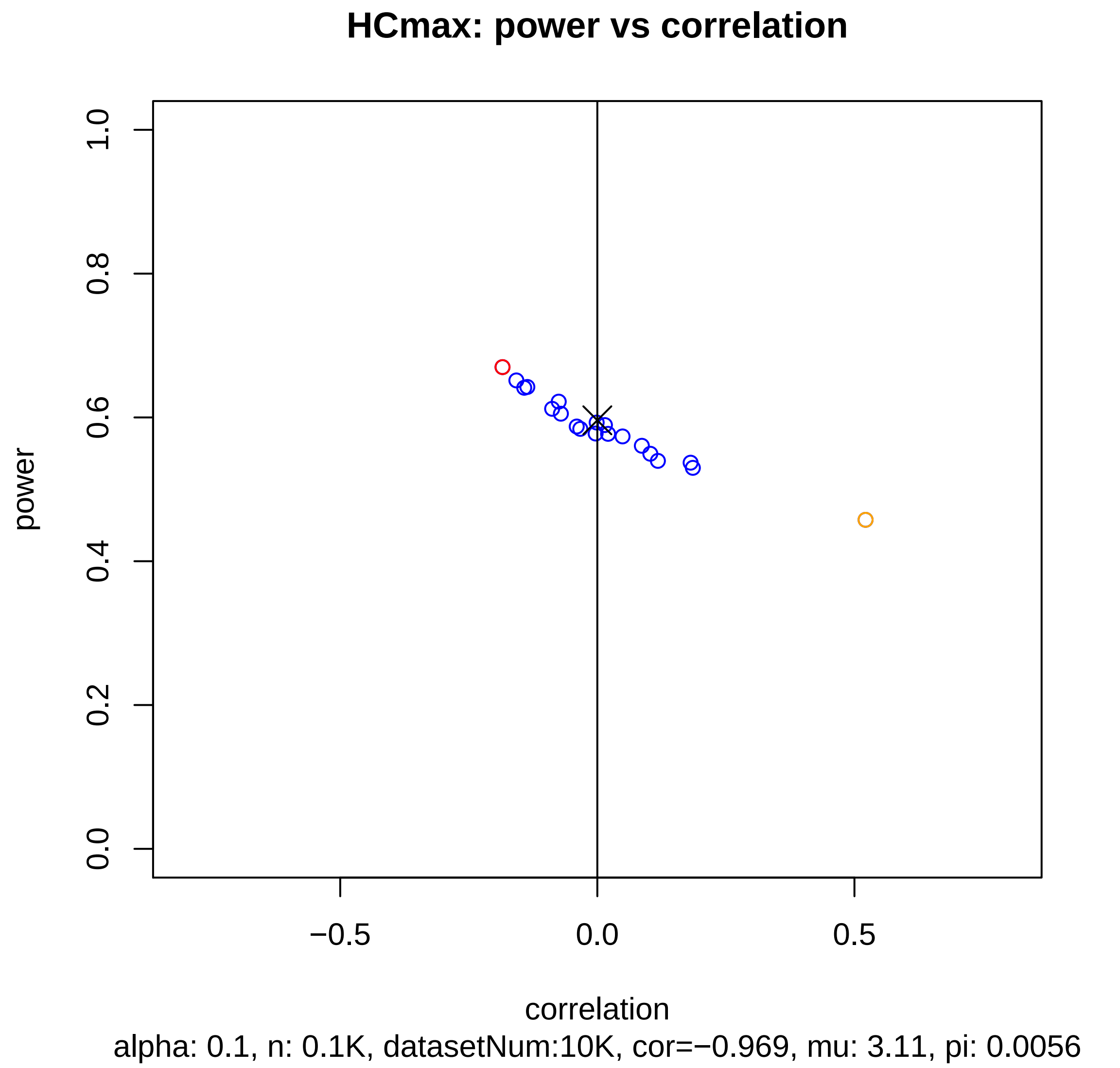
HCmax: power vs correlation
power
1.0
0.8
0.6
0.4
0.2
0.0
−0.5
0.0
0.5
correlation
alpha: 0.1, n: 0.1K, datasetNum:10K, cor=−0.969, mu: 3.11, pi: 0.0056

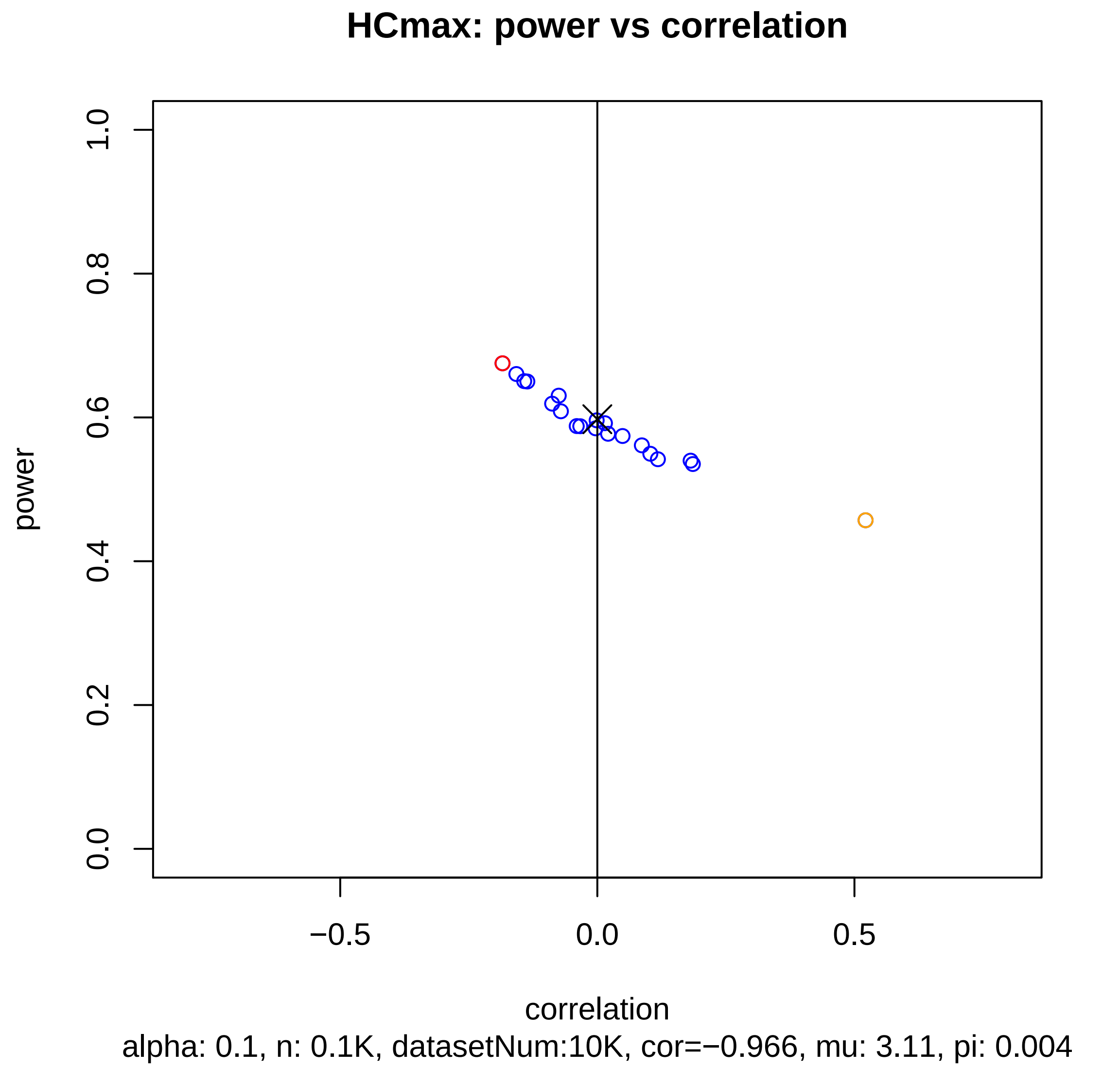
HCmax: power vs correlation
power
1.0
0.8
0.6
0.4
0.2
0.0
−0.5
0.0
0.5
correlation
alpha: 0.1, n: 0.1K, datasetNum:10K, cor=−0.966, mu: 3.11, pi: 0.004

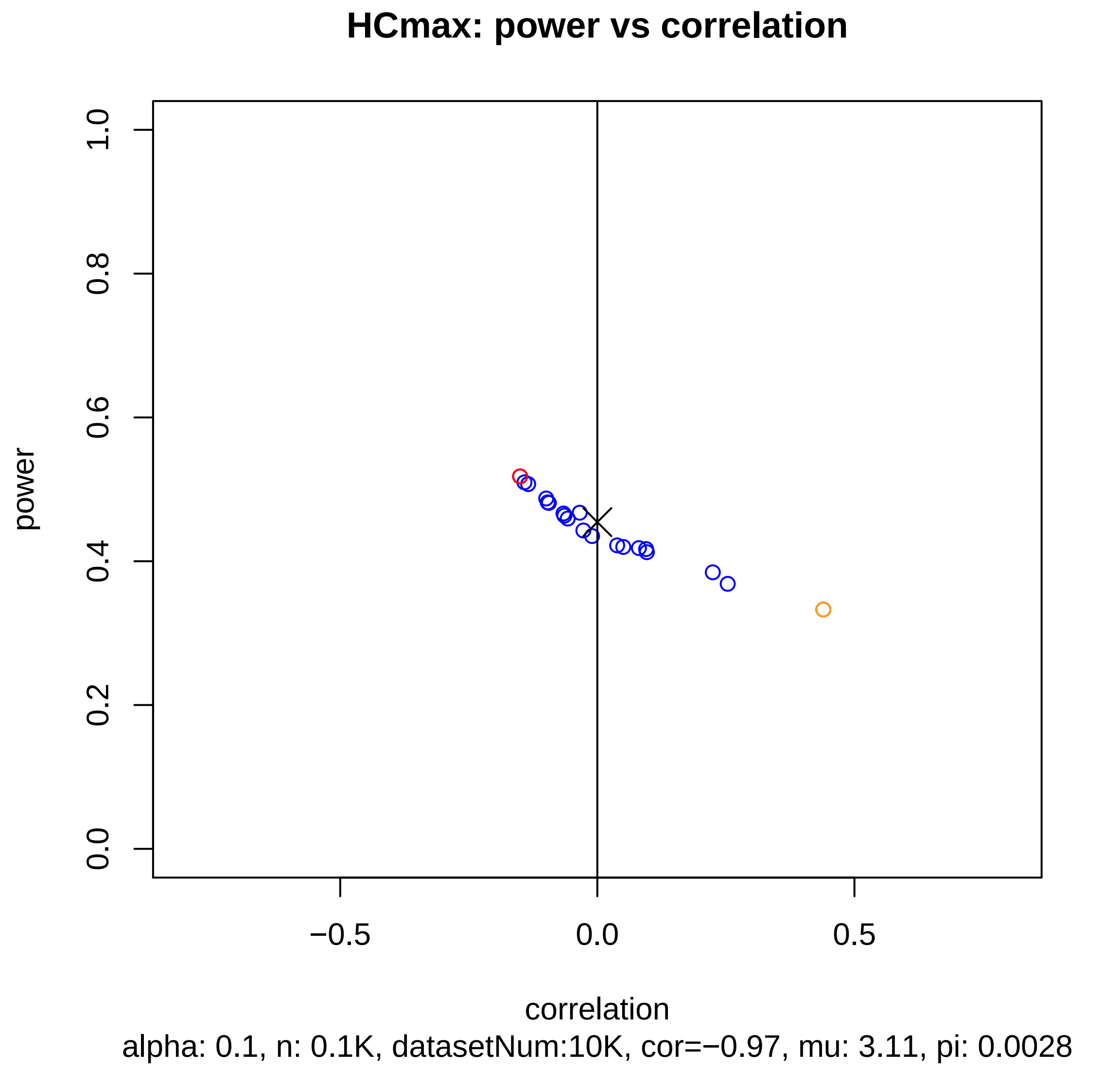

HCmax: power vs correlation
power
0.0
0.2
0.4
0.6
0.8
1.0
−0.5
0.0
0.5
correlation
alpha: 0.1, n: 0.1K, datasetNum:10K, cor=−0.97, mu: 3.11, pi: 0.0028

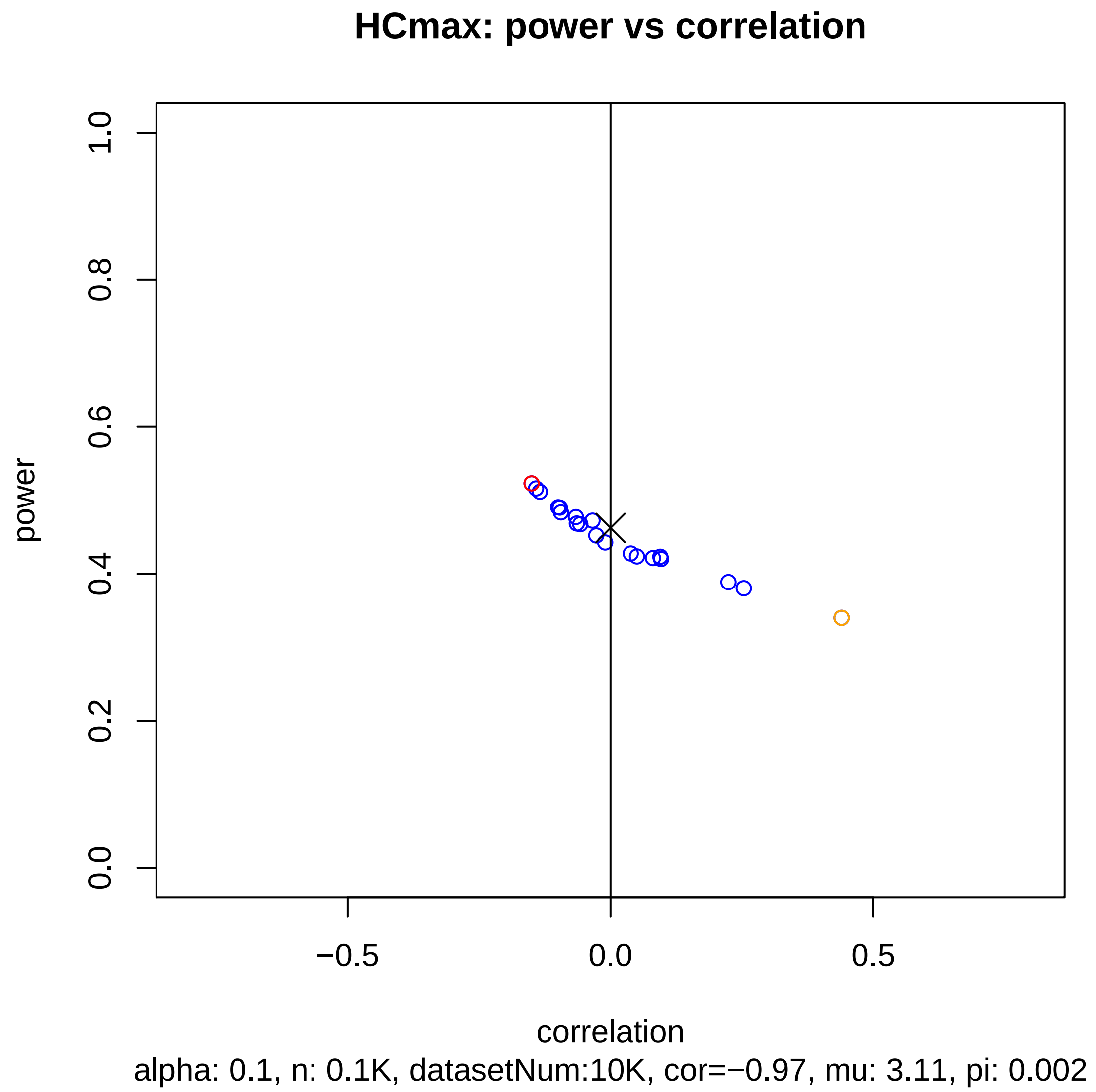
HCmax: power vs correlation
power
1.0
0.8
0.6
0.4
0.2
0.0
−0.5
0.0
0.5
correlation
alpha: 0.1, n: 0.1K, datasetNum:10K, cor=−0.97, mu: 3.11, pi: 0.002

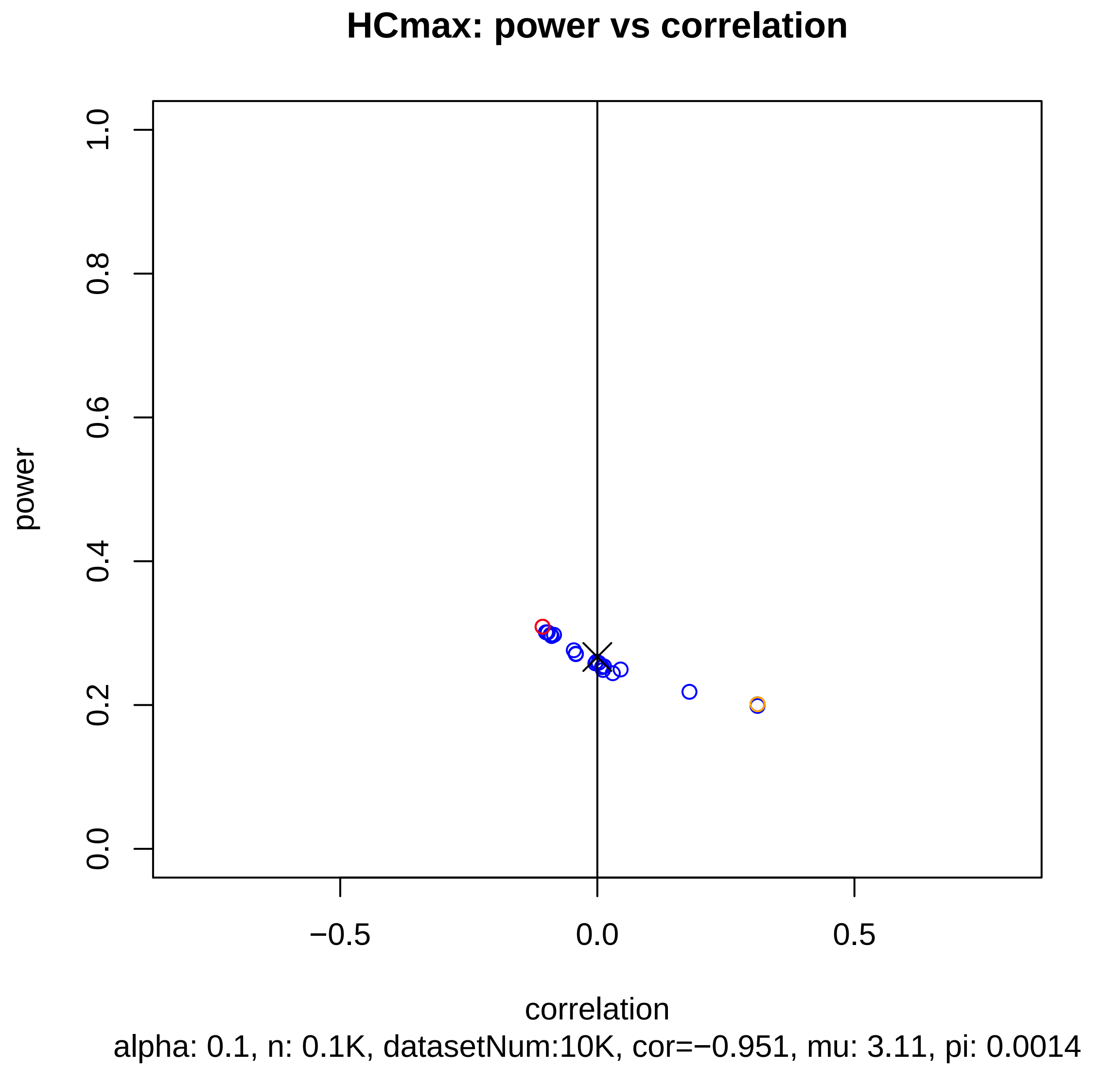
HCmax: power vs correlation
power
0.0
0.2
0.4
0.6
0.8
1.0
−0.5
0.0
0.5
correlation
alpha: 0.1, n: 0.1K, datasetNum:10K, cor=−0.951, mu: 3.11, pi: 0.0014

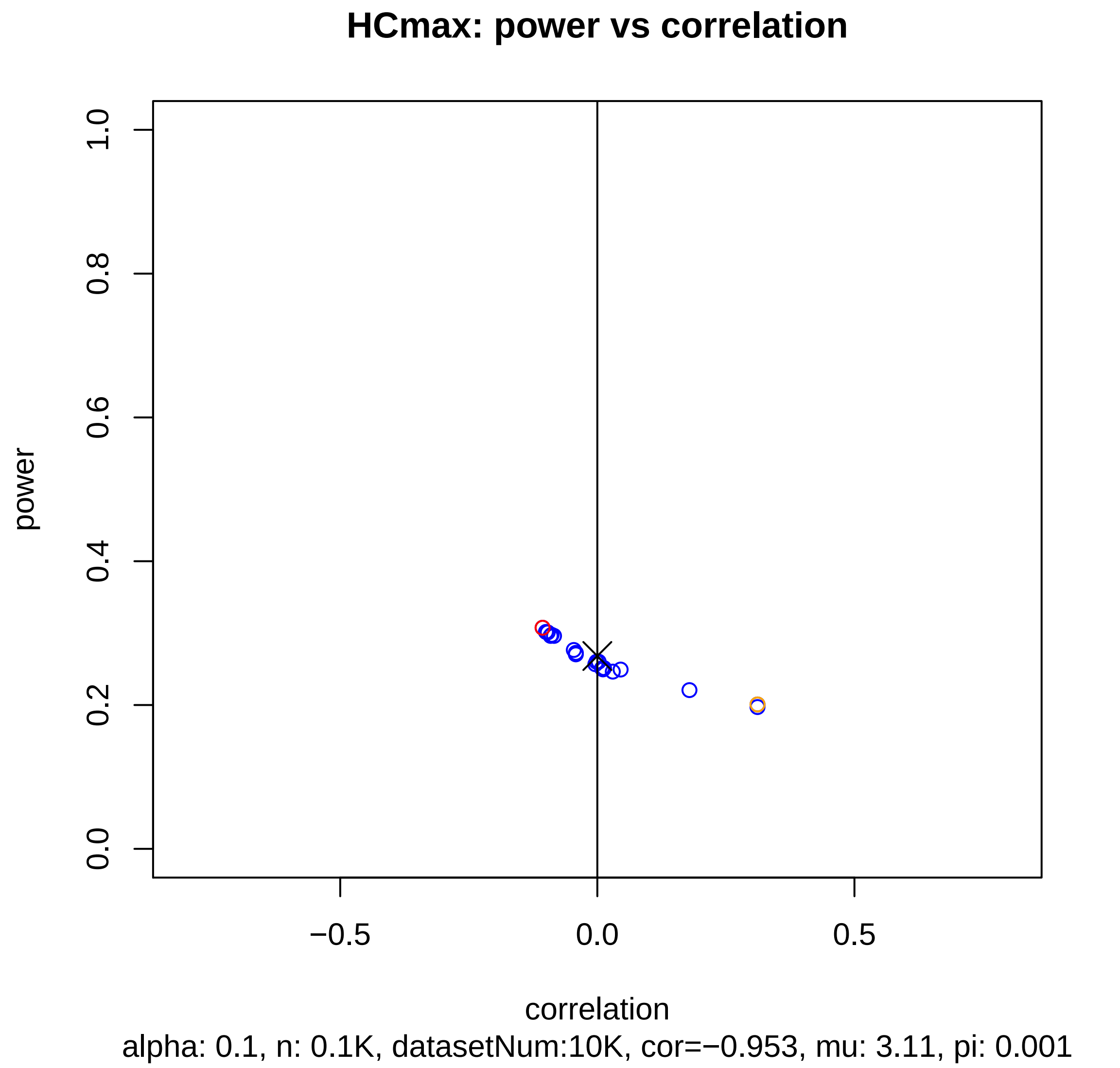

HCmax: power vs correlation
power
1.0
0.8
0.6
0.4
0.2
0.0
−0.5
0.0
0.5
correlation
alpha: 0.1, n: 0.1K, datasetNum:10K, cor=−0.953, mu: 3.11, pi: 0.001

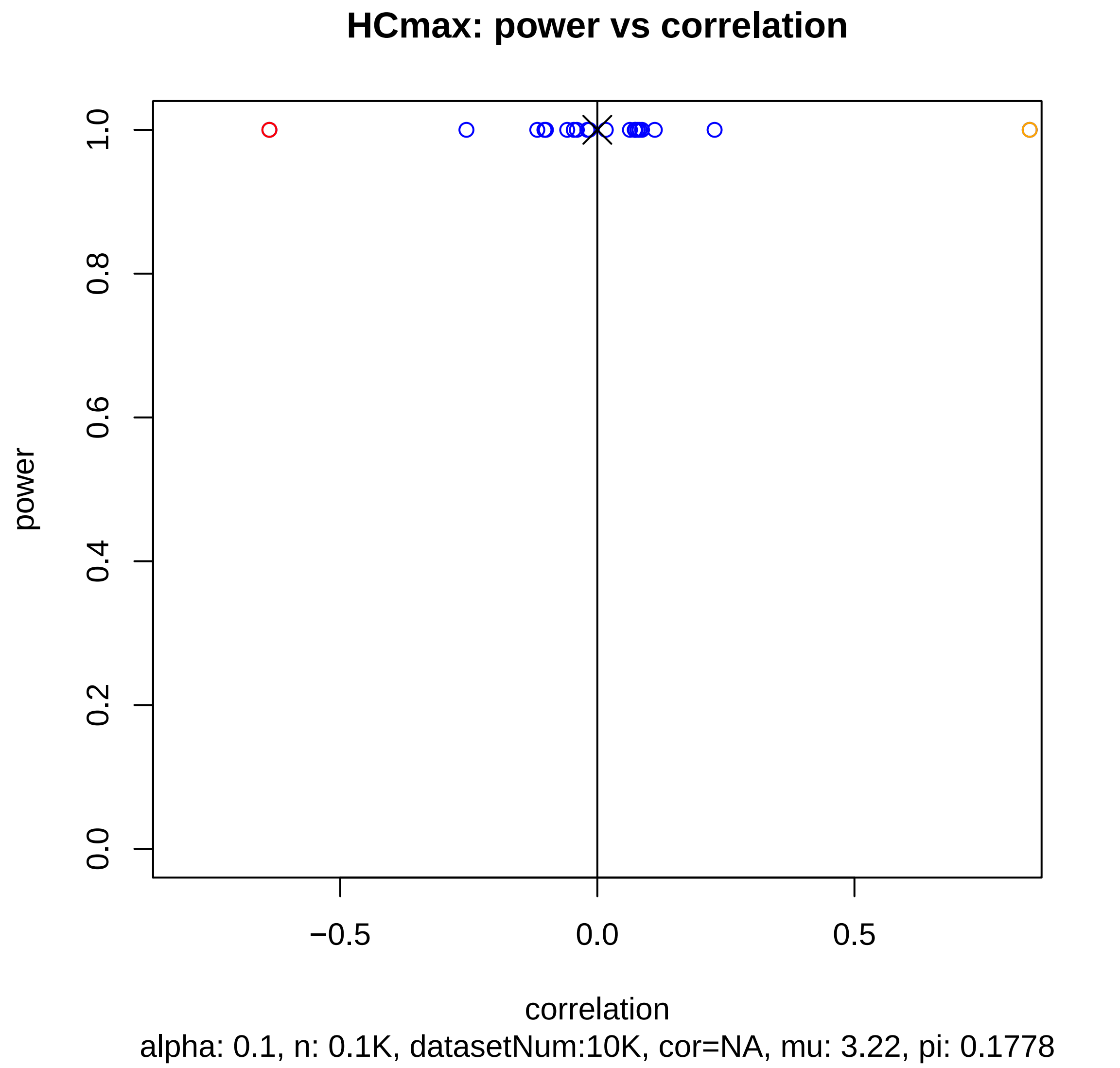

HCmax: power vs correlation
power
1.0
0.8
0.6
0.4
0.2
0.0
−0.5
0.0
0.5
correlation
alpha: 0.1, n: 0.1K, datasetNum:10K, cor=NA, mu: 3.22, pi: 0.1778

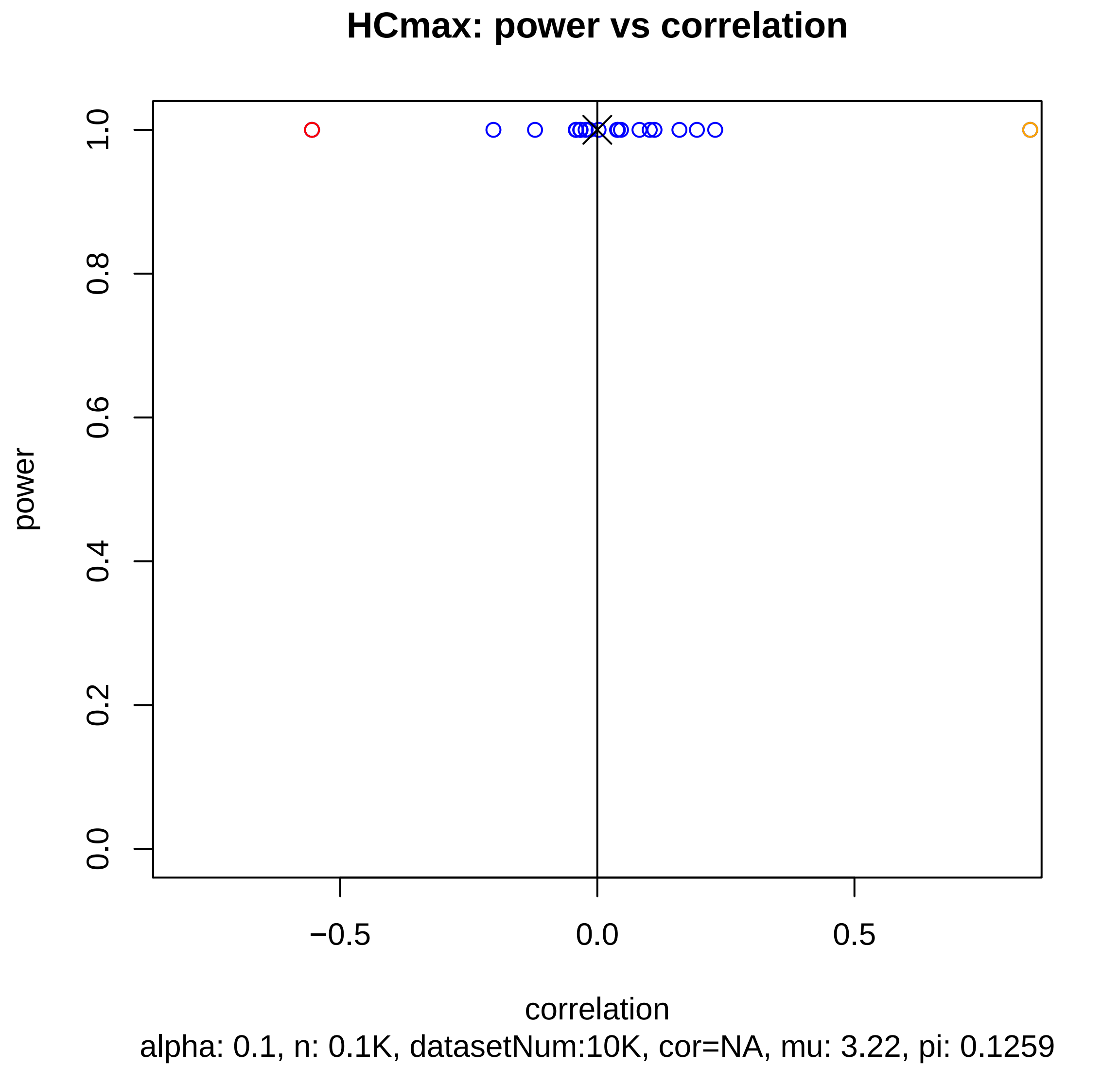
HCmax: power vs correlation
power
1.0
0.8
0.6
0.4
0.2
0.0
−0.5
0.0
0.5
correlation
alpha: 0.1, n: 0.1K, datasetNum:10K, cor=NA, mu: 3.22, pi: 0.1259

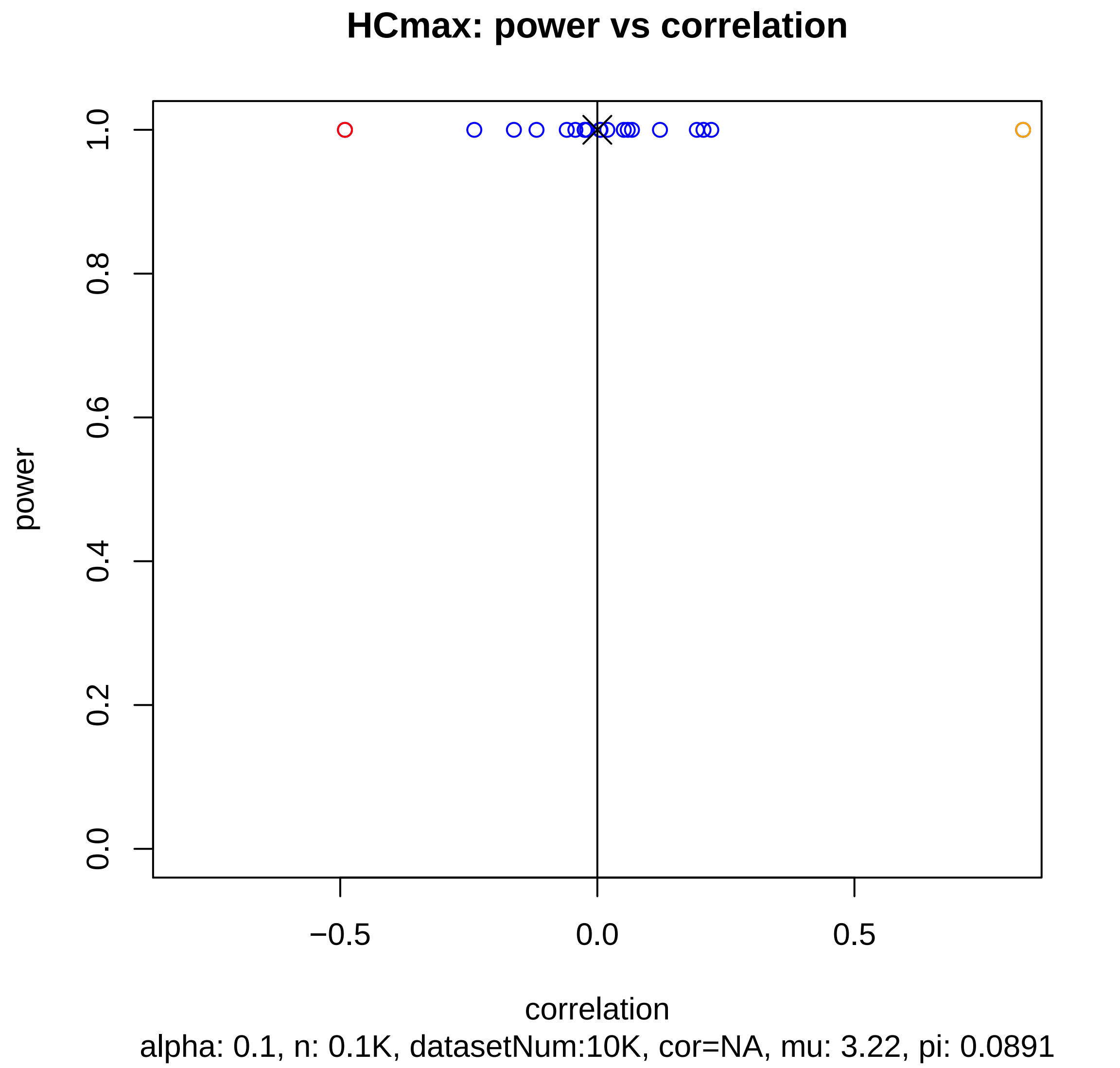
HCmax: power vs correlation
power
1.0
0.8
0.6
0.4
0.2
0.0
−0.5
0.0
0.5
correlation
alpha: 0.1, n: 0.1K, datasetNum:10K, cor=NA, mu: 3.22, pi: 0.0891

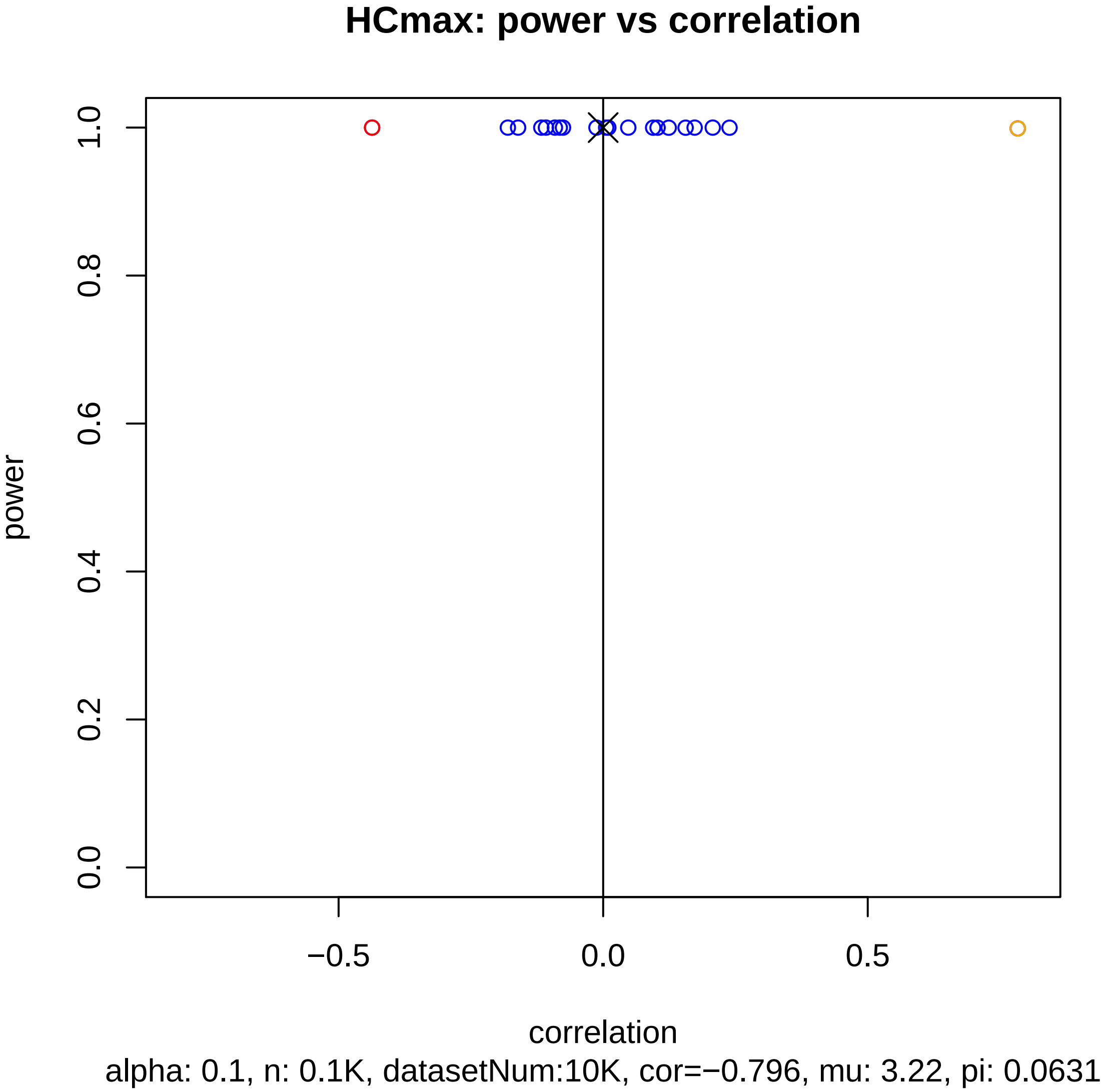
HCmax: power vs correlation
power
0.0
0.2
0.4
0.6
0.8
1.0
−0.5
0.0
0.5
correlation
alpha: 0.1, n: 0.1K, datasetNum:10K, cor=−0.796, mu: 3.22, pi: 0.0631

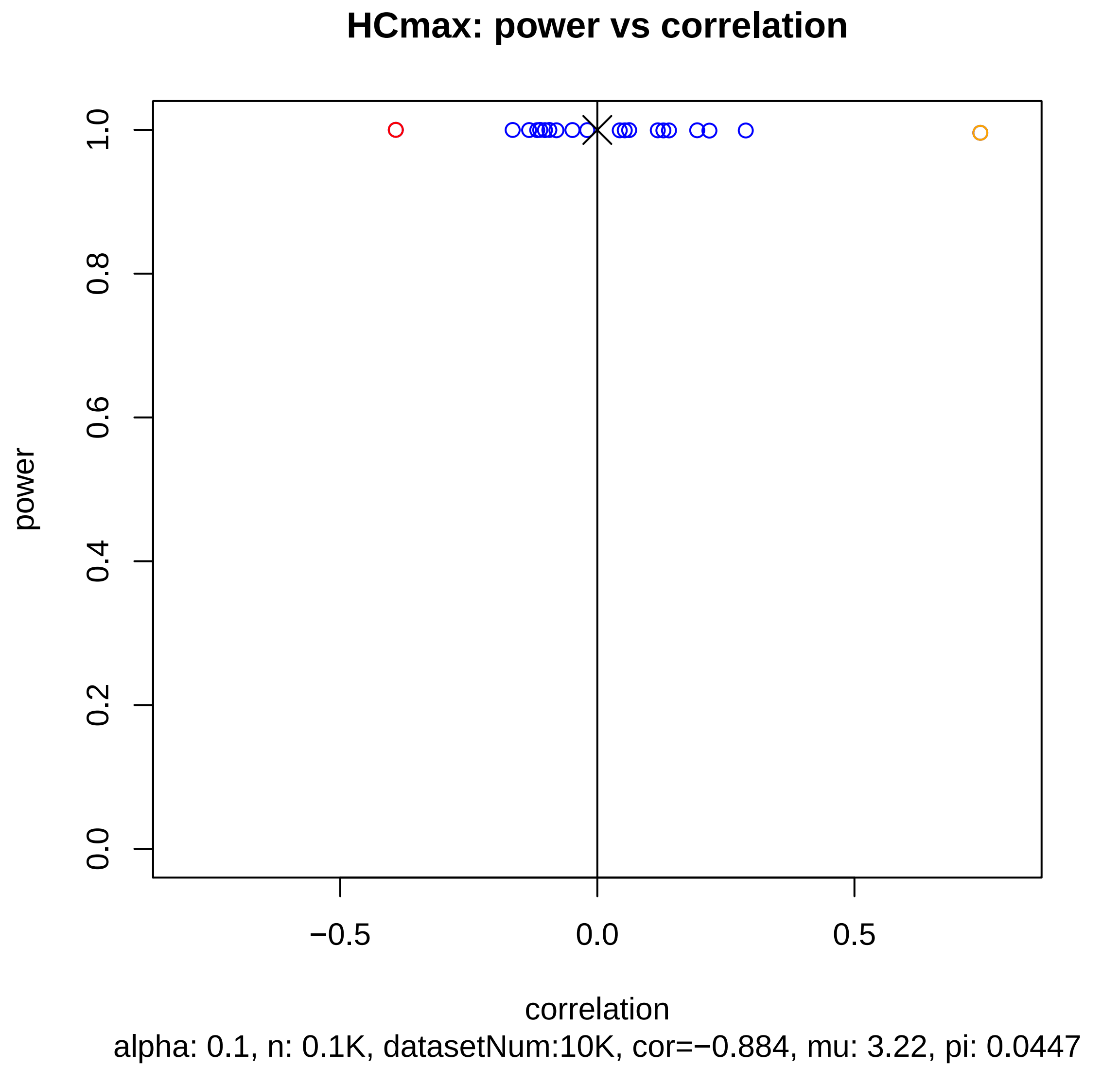
HCmax: power vs correlation
power
1.0
0.8
0.6
0.4
0.2
0.0
−0.5
0.0
0.5
correlation
alpha: 0.1, n: 0.1K, datasetNum:10K, cor=−0.884, mu: 3.22, pi: 0.0447

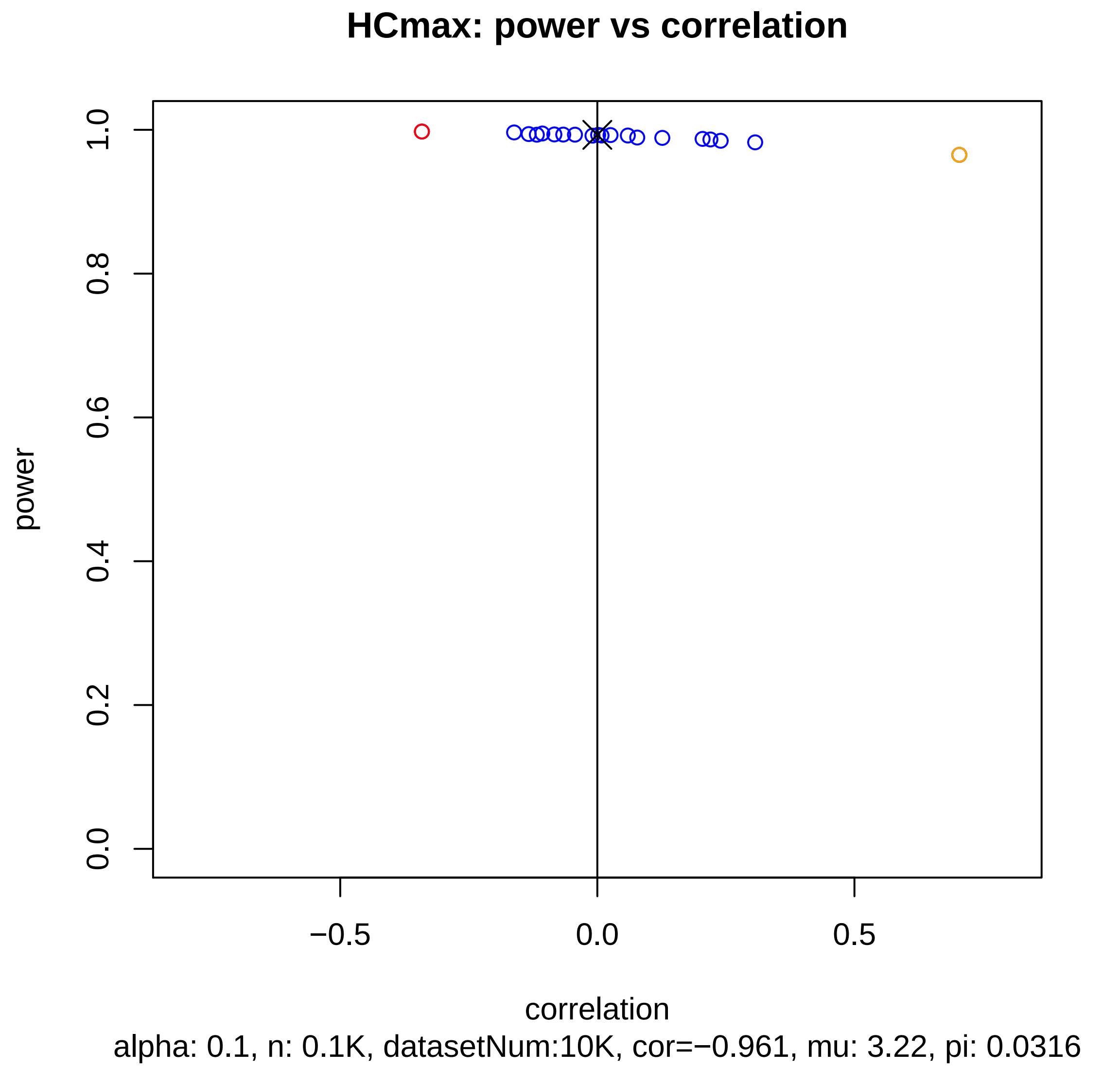

HCmax: power vs correlation
power
1.0
0.8
0.6
0.4
0.2
0.0
−0.5
0.0
0.5
correlation
alpha: 0.1, n: 0.1K, datasetNum:10K, cor=−0.961, mu: 3.22, pi: 0.0316

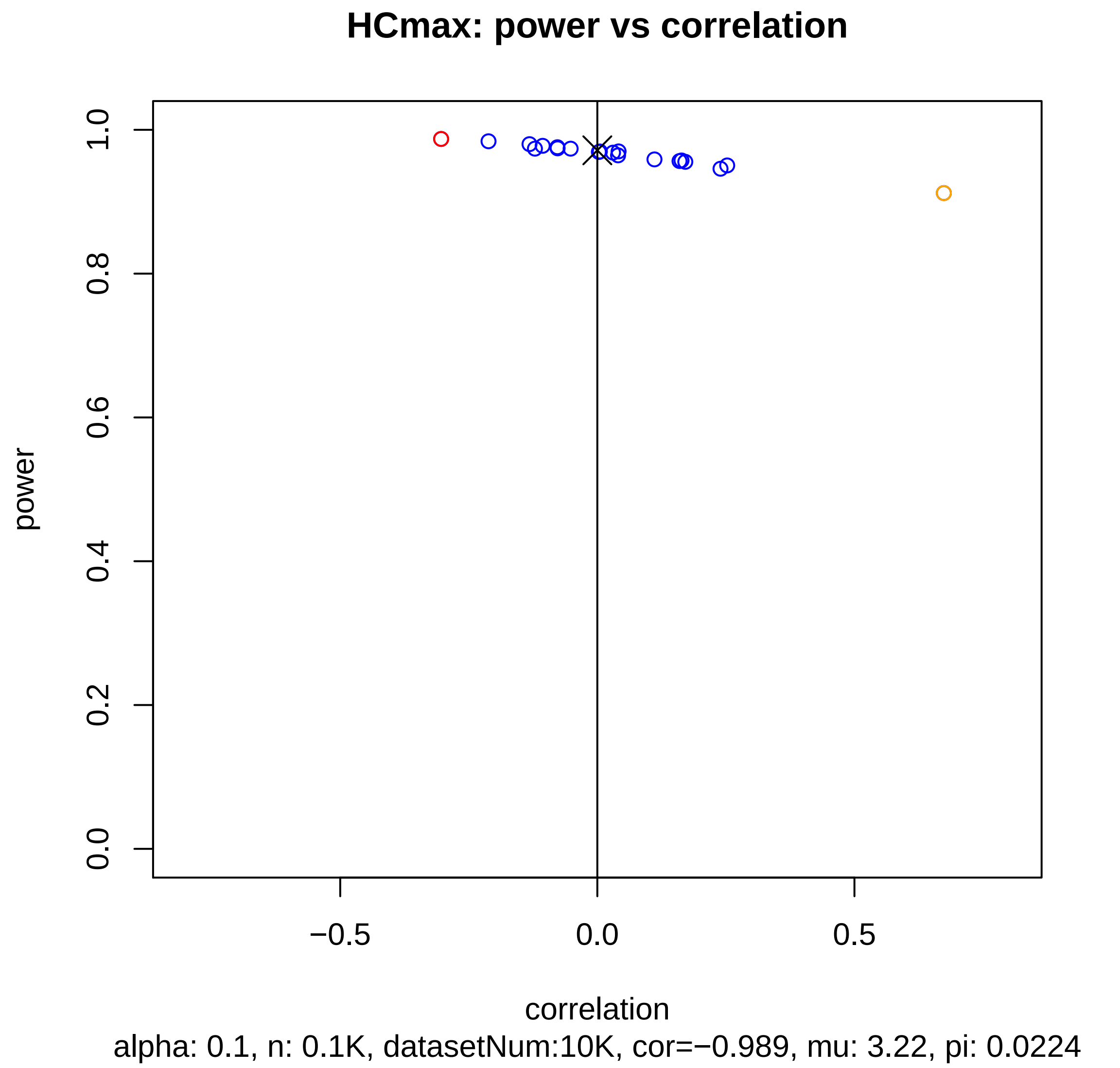

HCmax: power vs correlation
power
correlation
0.0
0.2
0.4
0.6
0.8
1.0
−0.5
0.0
0.5
alpha: 0.1, n: 0.1K, datasetNum:10K, cor=−0.989, mu: 3.22, pi: 0.0224

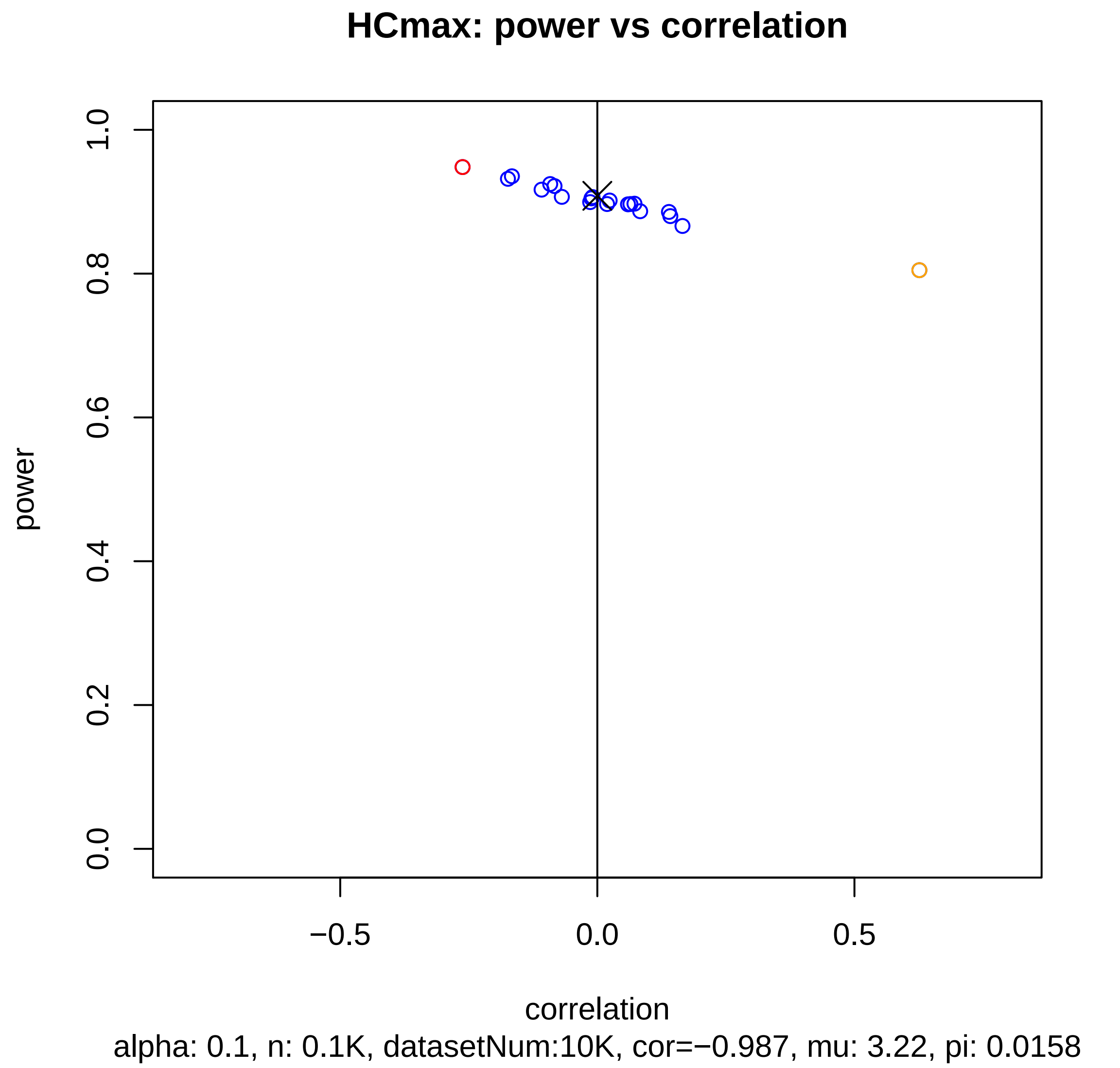
HCmax: power vs correlation
power
1.0
0.8
0.6
0.4
0.2
0.0
−0.5
0.0
0.5
correlation
alpha: 0.1, n: 0.1K, datasetNum:10K, cor=−0.987, mu: 3.22, pi: 0.0158

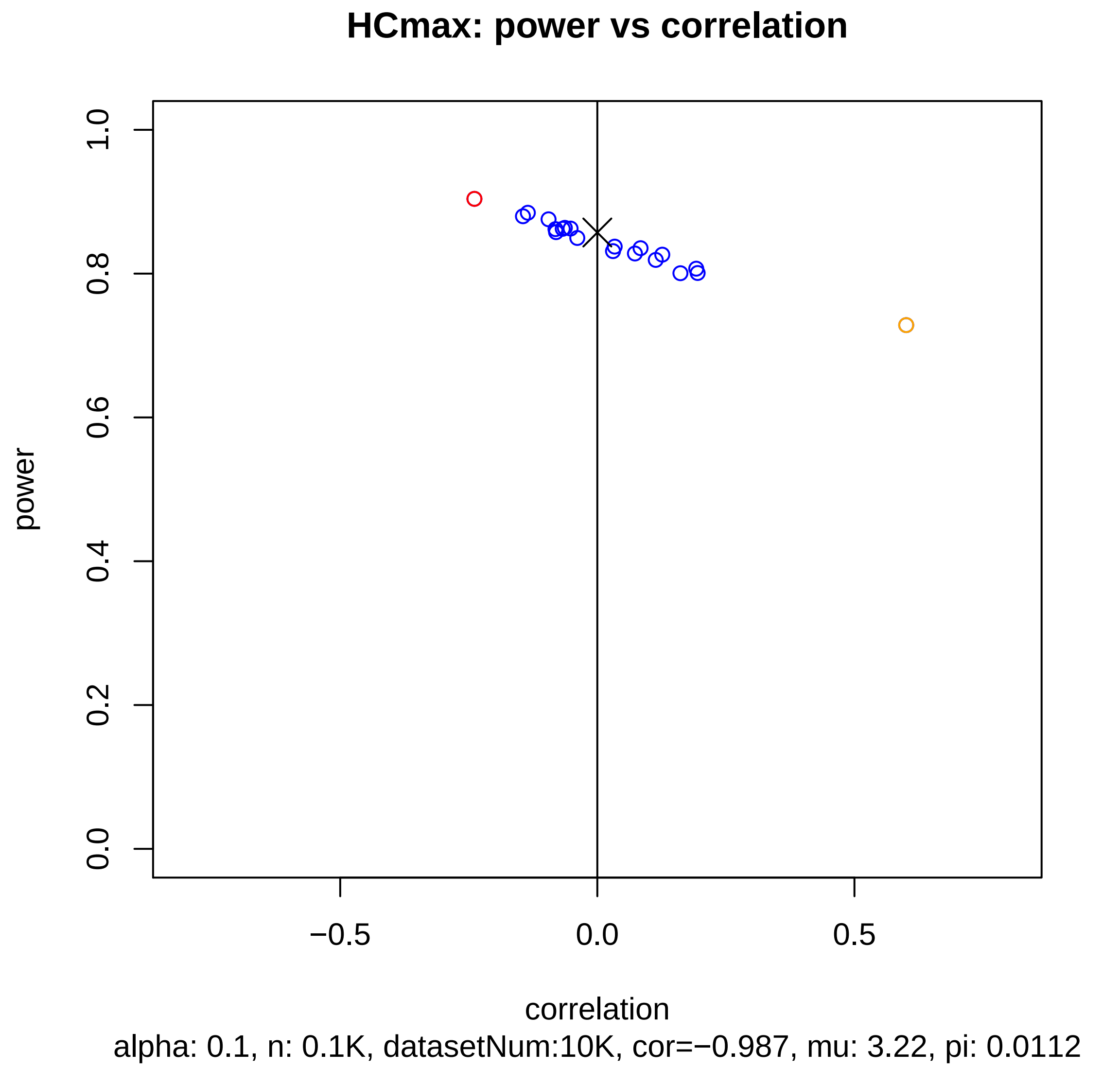

HCmax: power vs correlation
power
0.0
0.2
0.4
0.6
0.8
1.0
−0.5
0.0
0.5
correlation
alpha: 0.1, n: 0.1K, datasetNum:10K, cor=−0.987, mu: 3.22, pi: 0.0112

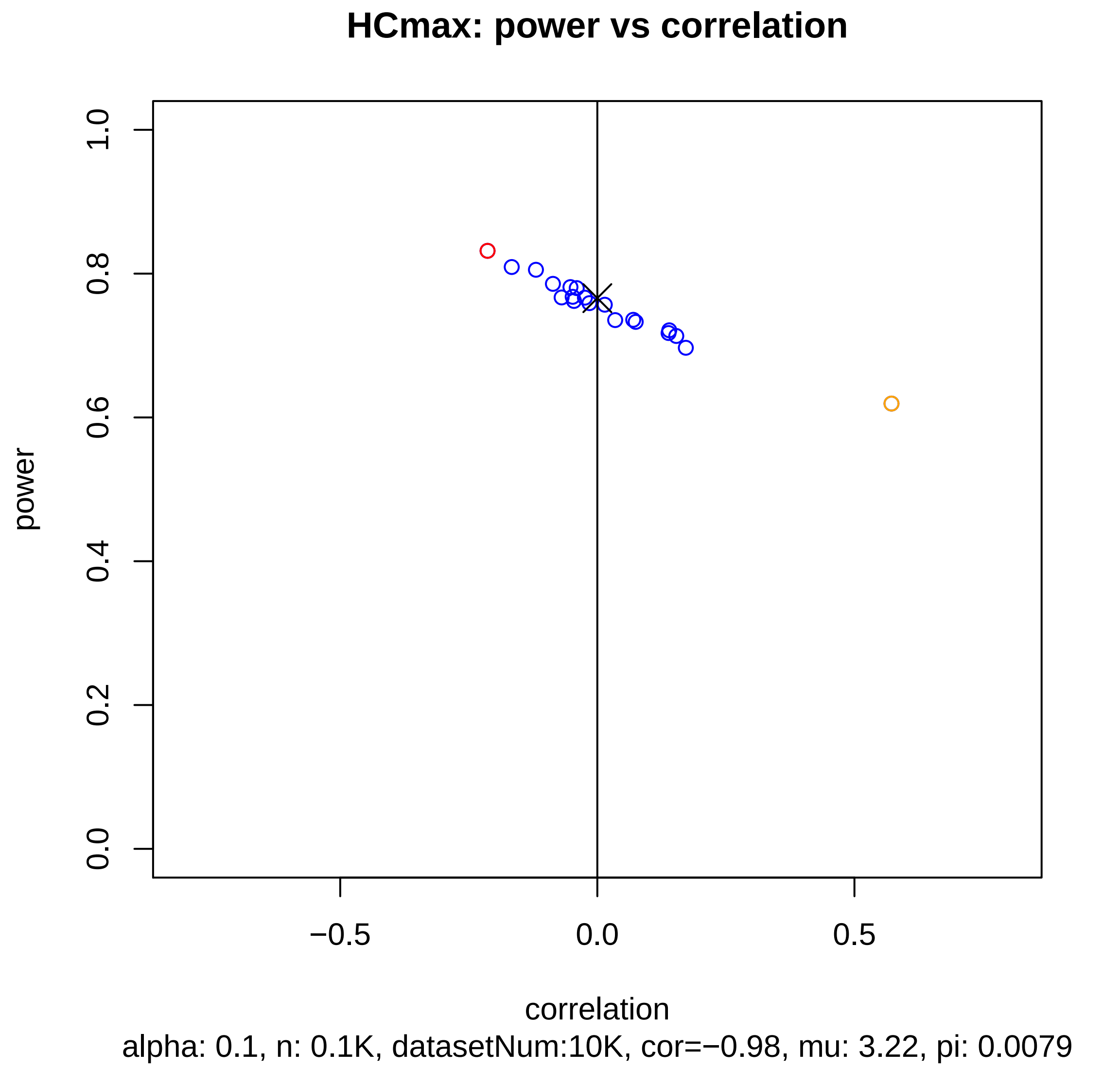

HCmax: power vs correlation
power
1.0
0.8
0.6
0.4
0.2
0.0
−0.5
0.0
0.5
correlation
alpha: 0.1, n: 0.1K, datasetNum:10K, cor=−0.98, mu: 3.22, pi: 0.0079

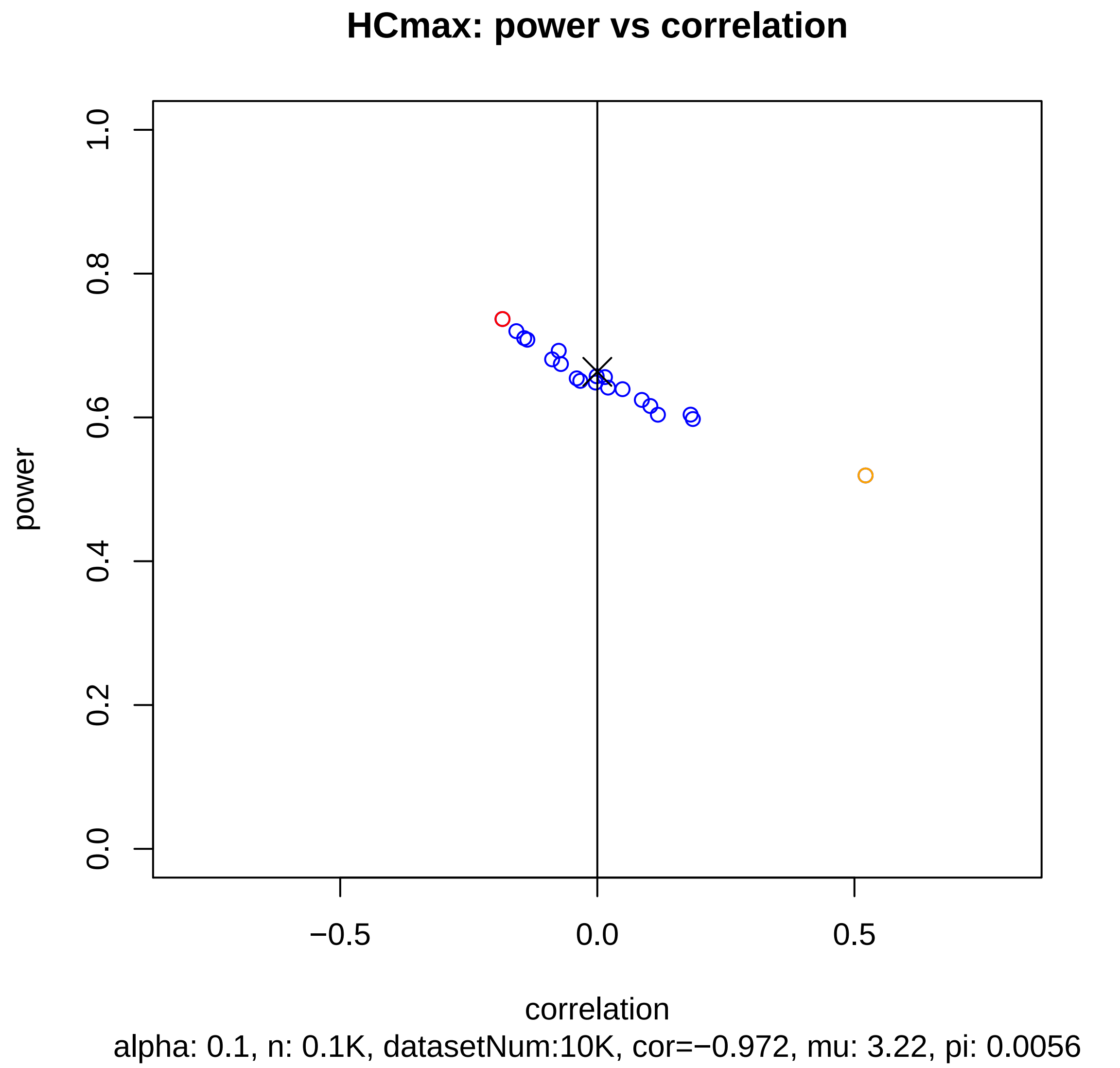
HCmax: power vs correlation
power
0.0
0.2
0.4
0.6
0.8
1.0
−0.5
0.0
0.5
correlation
alpha: 0.1, n: 0.1K, datasetNum:10K, cor=−0.972, mu: 3.22, pi: 0.0056

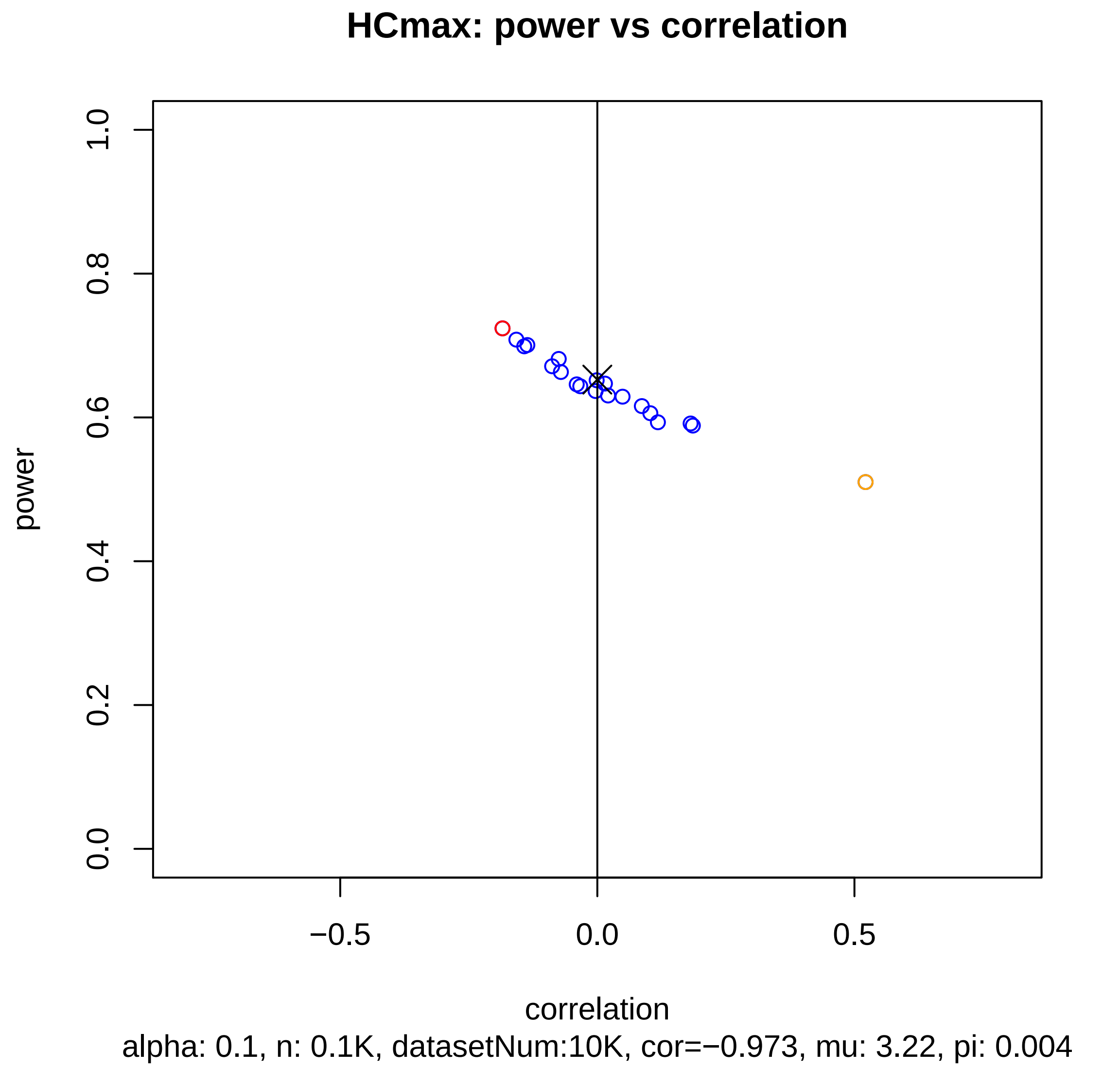

HCmax: power vs correlation
power
0.0
0.2
0.4
0.6
0.8
1.0
−0.5
0.0
0.5
correlation
alpha: 0.1, n: 0.1K, datasetNum:10K, cor=−0.973, mu: 3.22, pi: 0.004

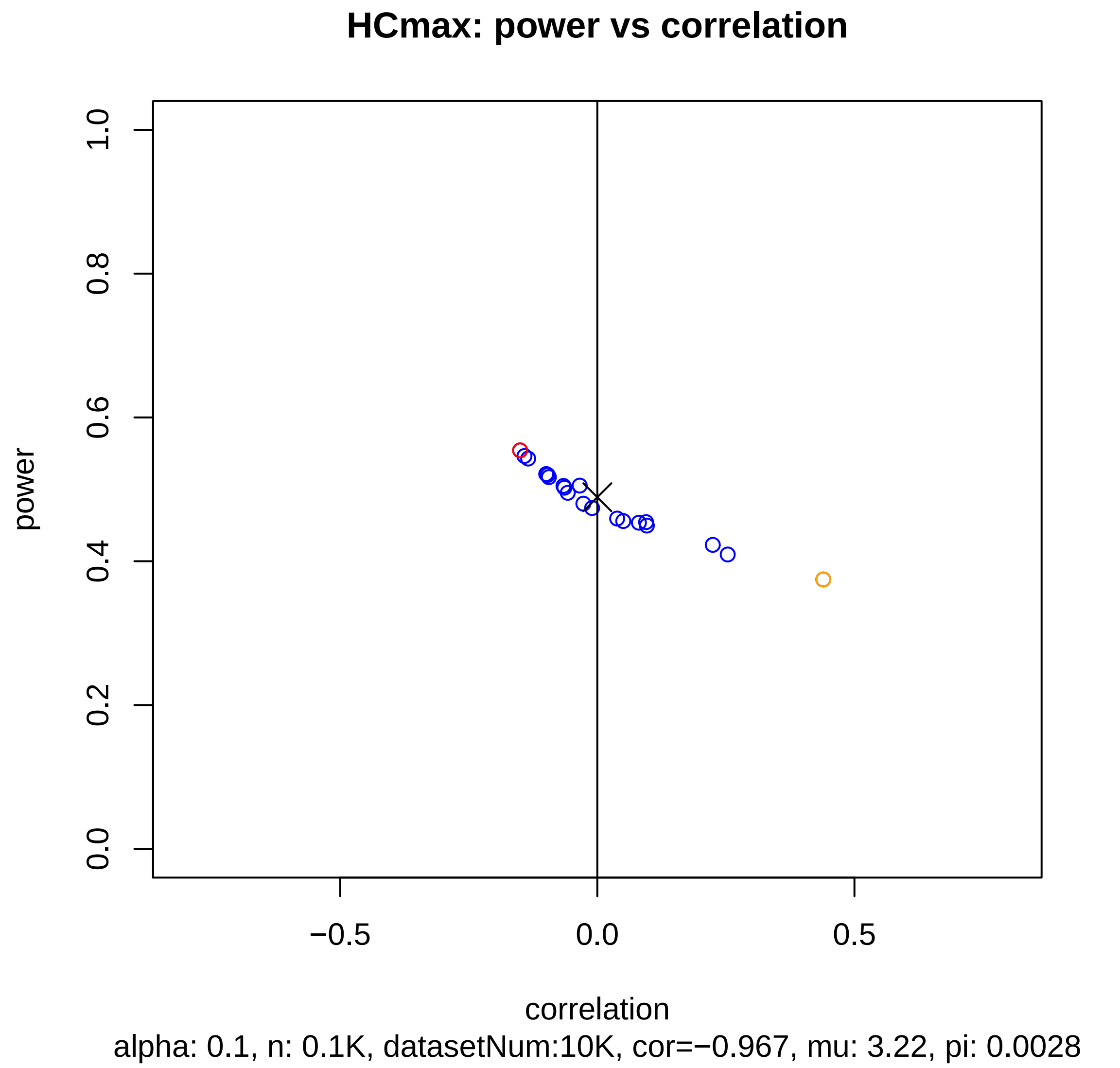

HCmax: power vs correlation
power
correlation
1.0
0.8
0.6
0.4
0.2
0.0
−0.5
0.0
0.5
alpha: 0.1, n: 0.1K, datasetNum:10K, cor=−0.967, mu: 3.22, pi: 0.0028

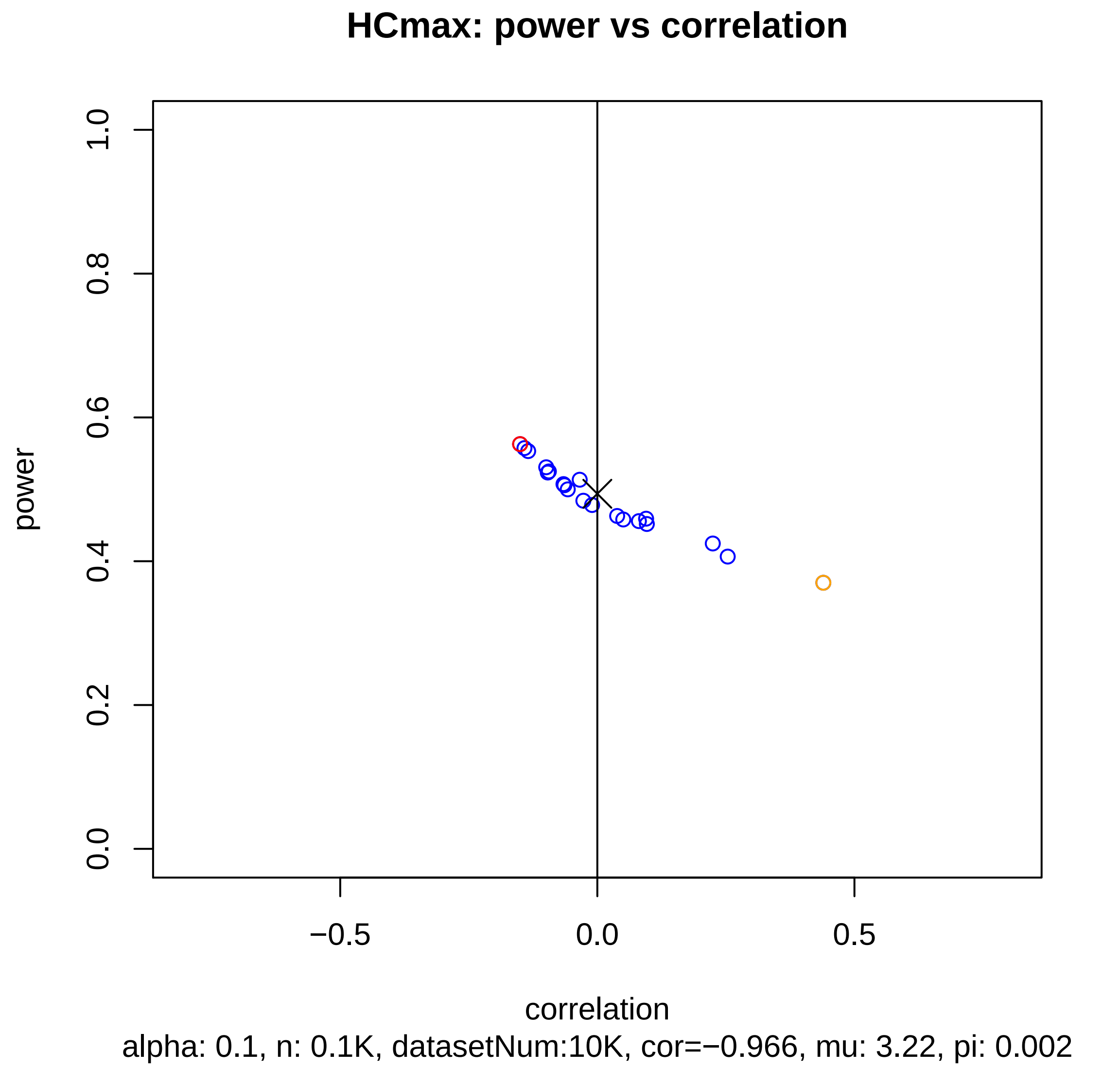
HCmax: power vs correlation
power
1.0
0.8
0.6
0.4
0.2
0.0
−0.5
0.0
0.5
correlation
alpha: 0.1, n: 0.1K, datasetNum:10K, cor=−0.966, mu: 3.22, pi: 0.002

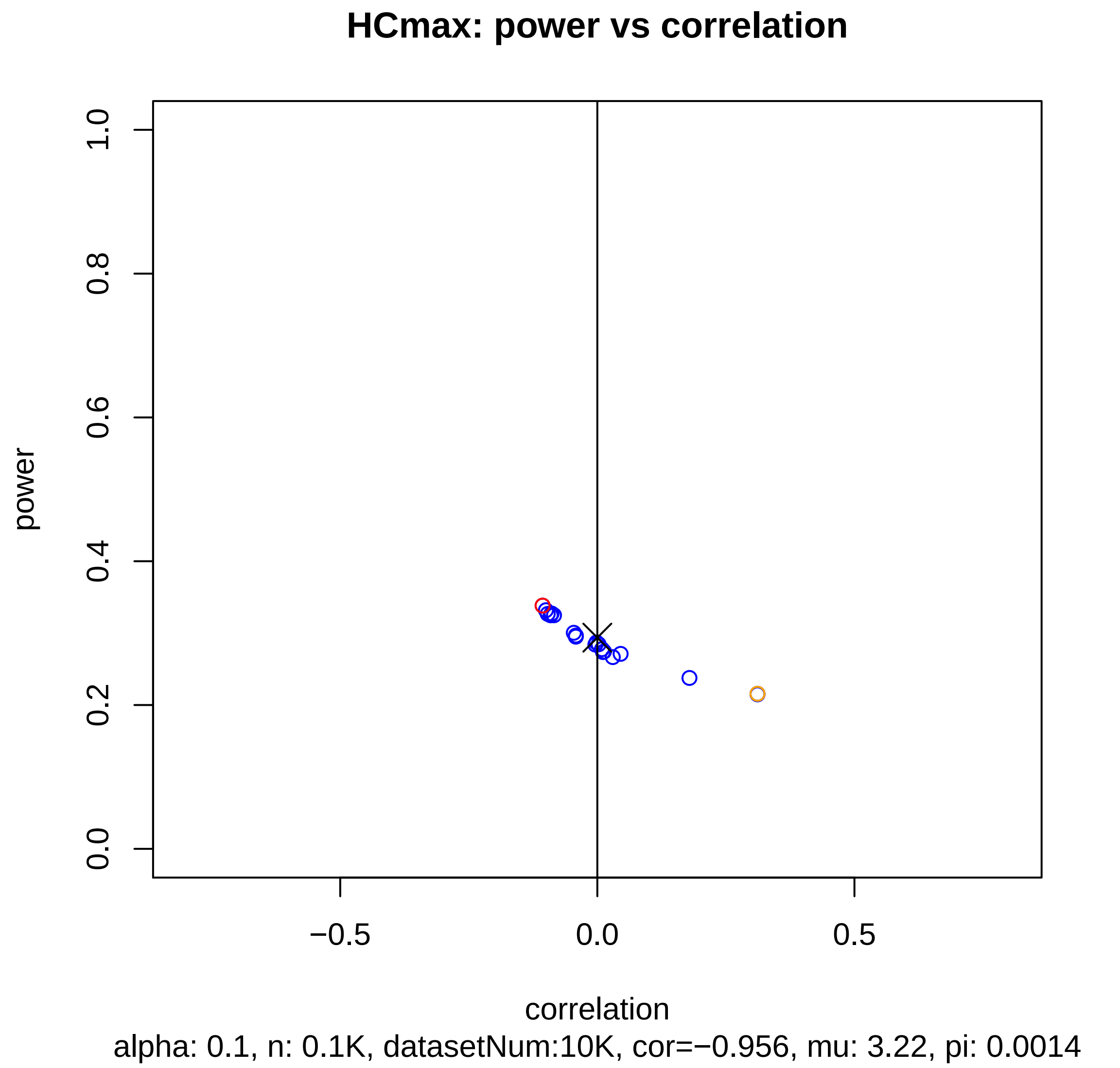

HCmax: power vs correlation
power
1.0
0.8
0.6
0.4
0.2
0.0
−0.5
0.0
0.5
correlation
alpha: 0.1, n: 0.1K, datasetNum:10K, cor=−0.956, mu: 3.22, pi: 0.0014

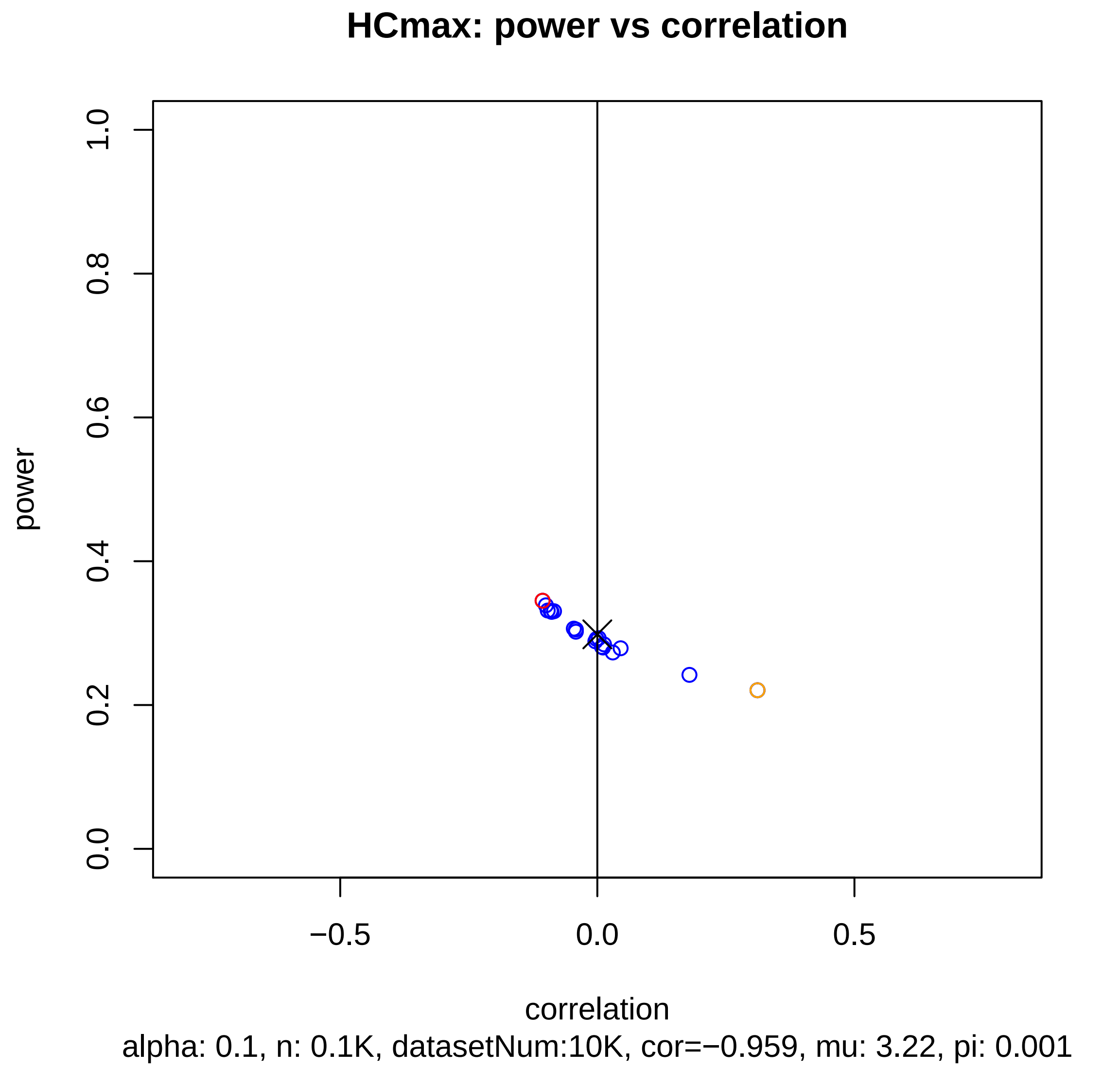

HCmax: power vs correlation
power
1.0
0.8
0.6
0.4
0.2
0.0
−0.5
0.0
0.5
correlation
alpha: 0.1, n: 0.1K, datasetNum:10K, cor=−0.959, mu: 3.22, pi: 0.001

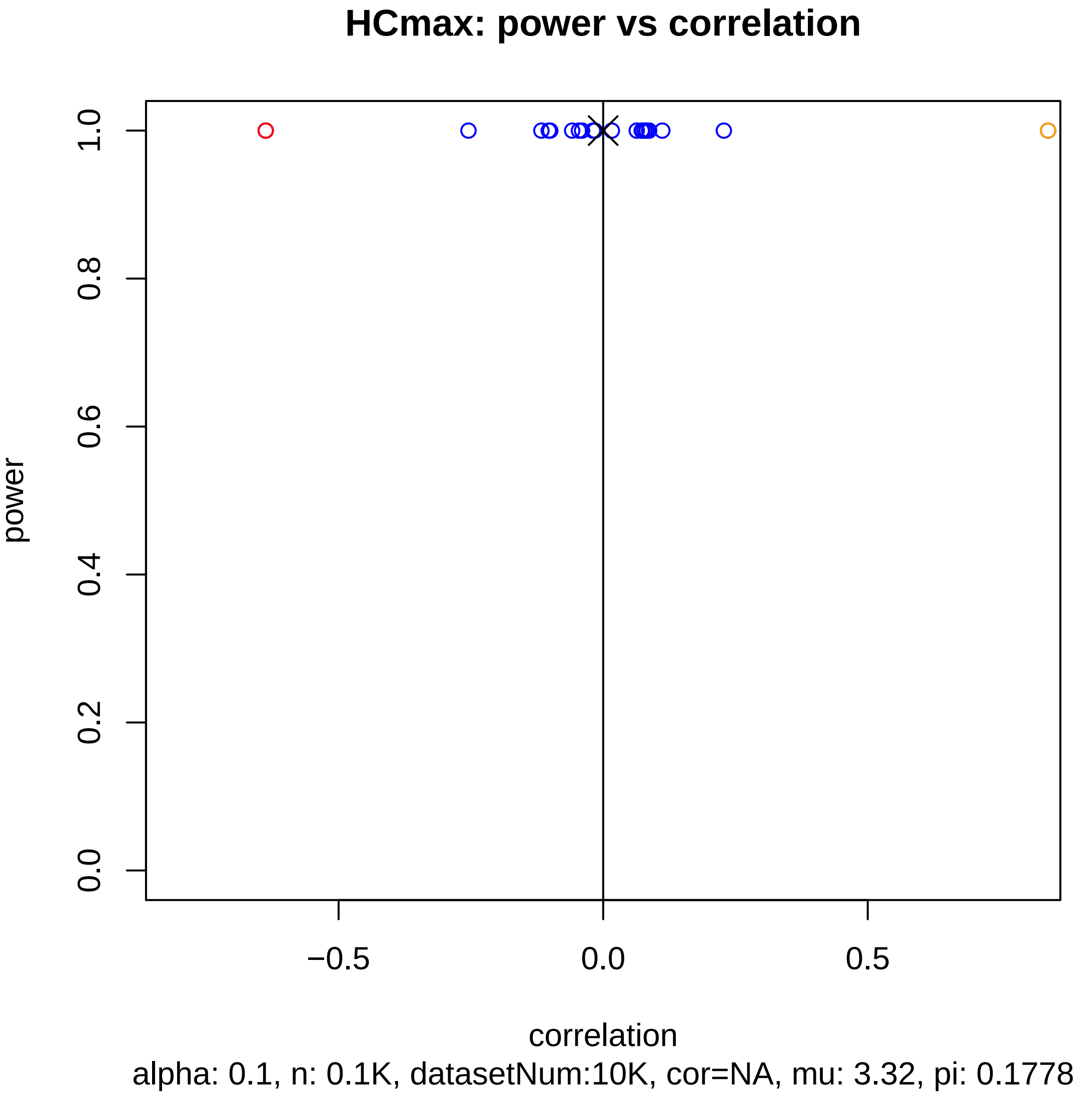
HCmax: power vs correlation
power
1.0
0.8
0.6
0.4
0.2
0.0
−0.5
0.0
0.5
correlation
alpha: 0.1, n: 0.1K, datasetNum:10K, cor=NA, mu: 3.32, pi: 0.1778

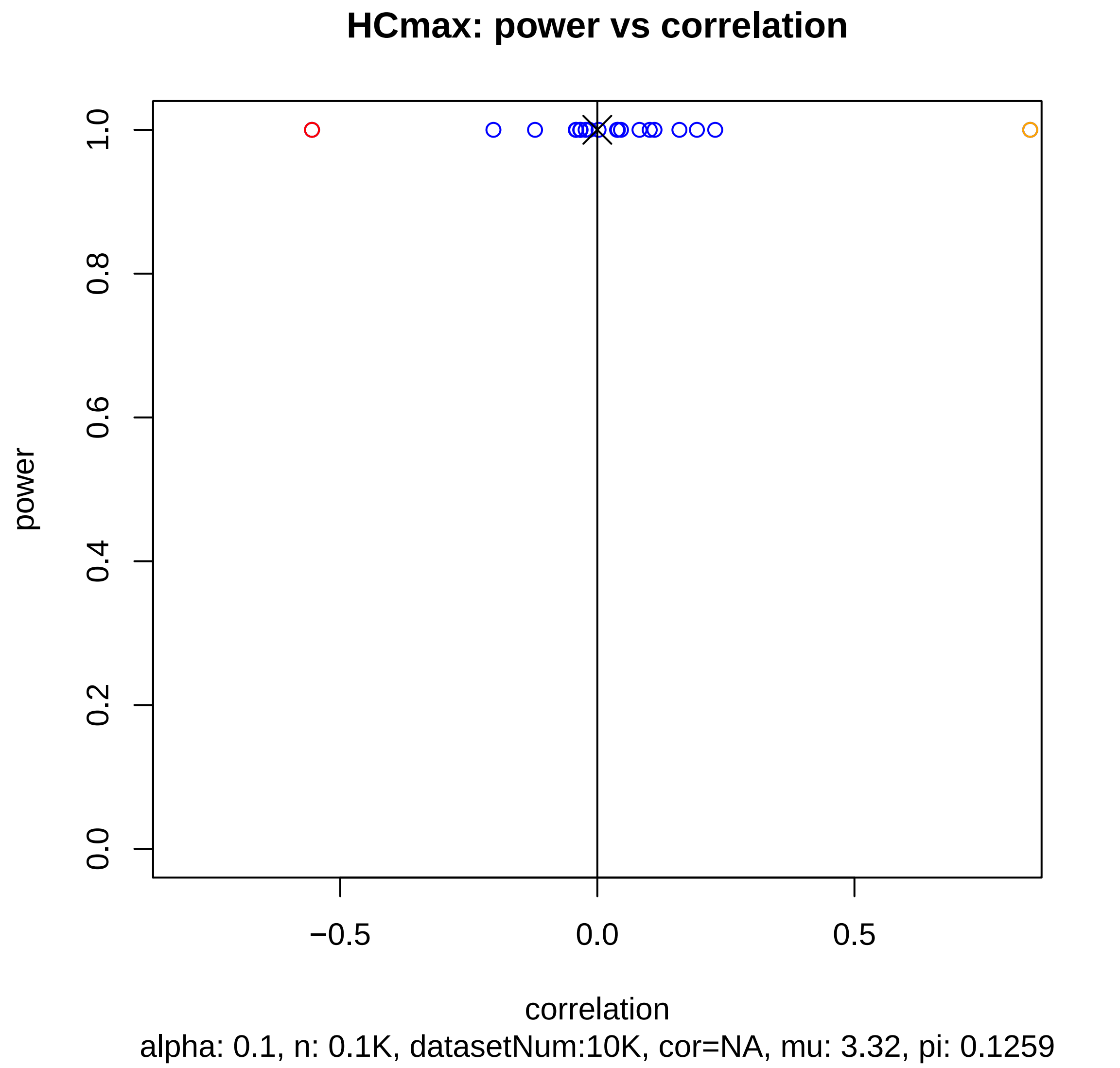
HCmax: power vs correlation
power
1.0
0.8
0.6
0.4
0.2
0.0
−0.5
0.0
0.5
correlation
alpha: 0.1, n: 0.1K, datasetNum:10K, cor=NA, mu: 3.32, pi: 0.1259

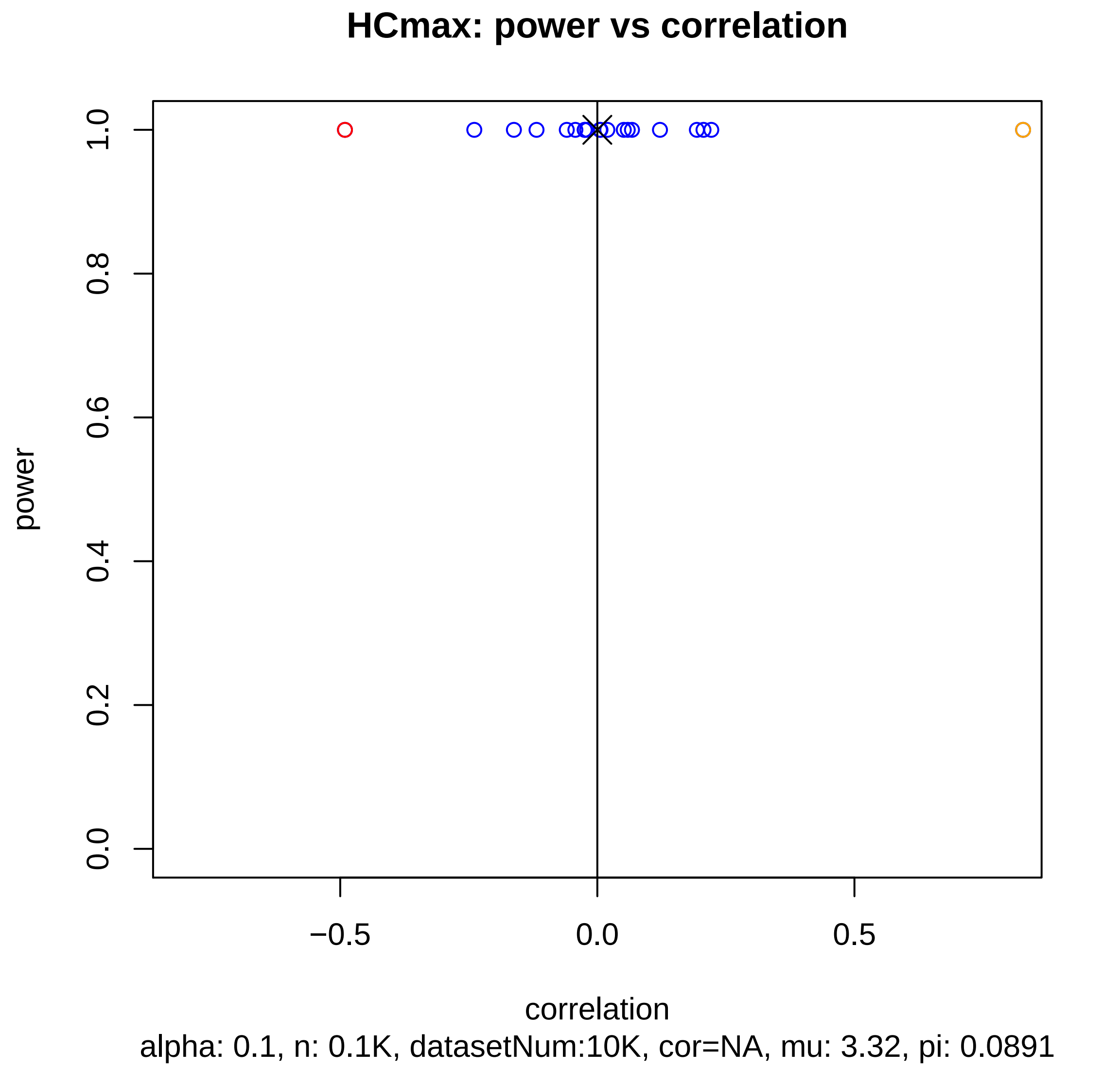
HCmax: power vs correlation
power
1.0
0.8
0.6
0.4
0.2
0.0
−0.5
0.0
0.5
correlation
alpha: 0.1, n: 0.1K, datasetNum:10K, cor=NA, mu: 3.32, pi: 0.0891

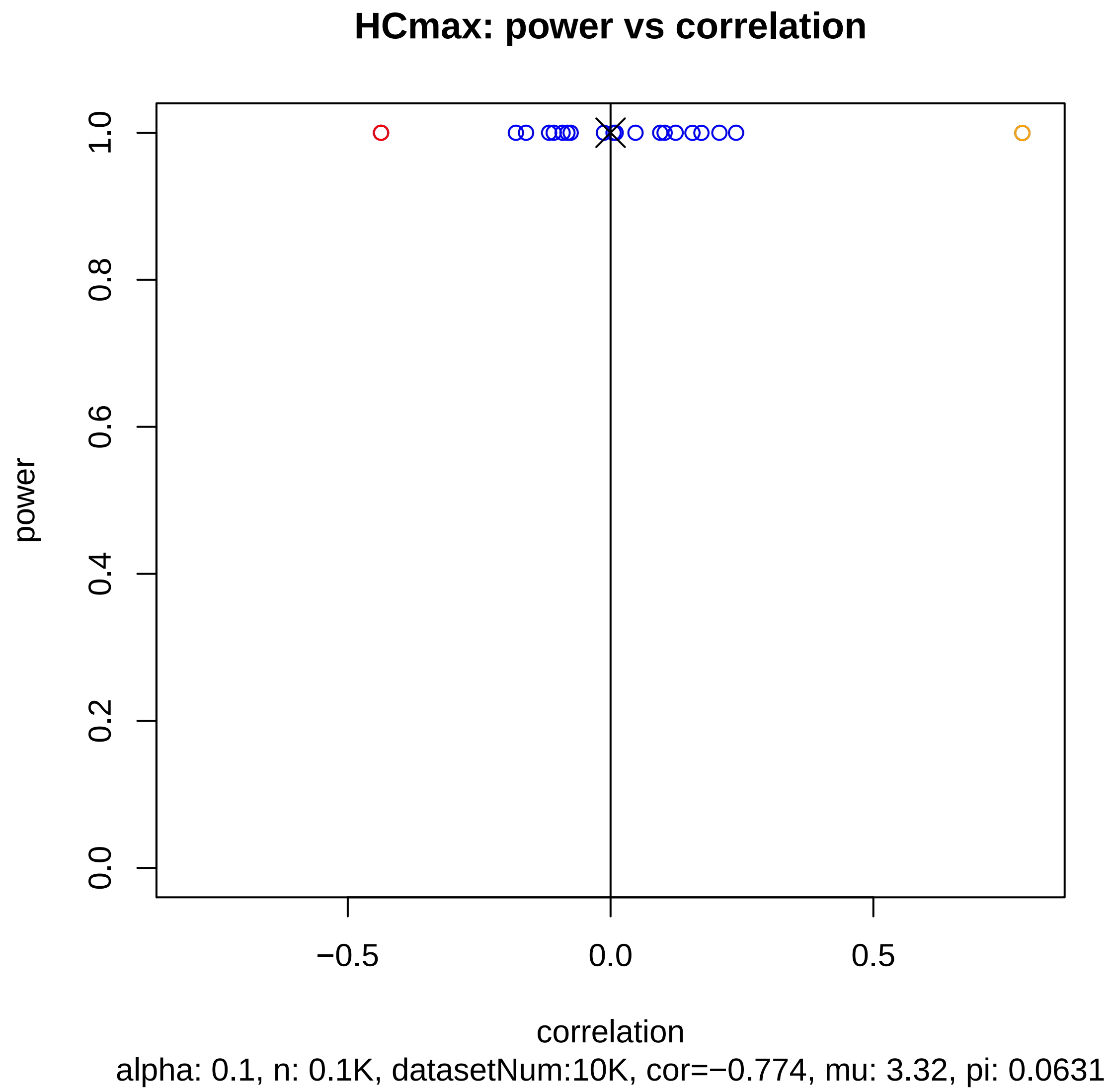
HCmax: power vs correlation
power
0.0
0.2
0.4
0.6
0.8
1.0
−0.5
0.0
0.5
correlation
alpha: 0.1, n: 0.1K, datasetNum:10K, cor=−0.774, mu: 3.32, pi: 0.0631

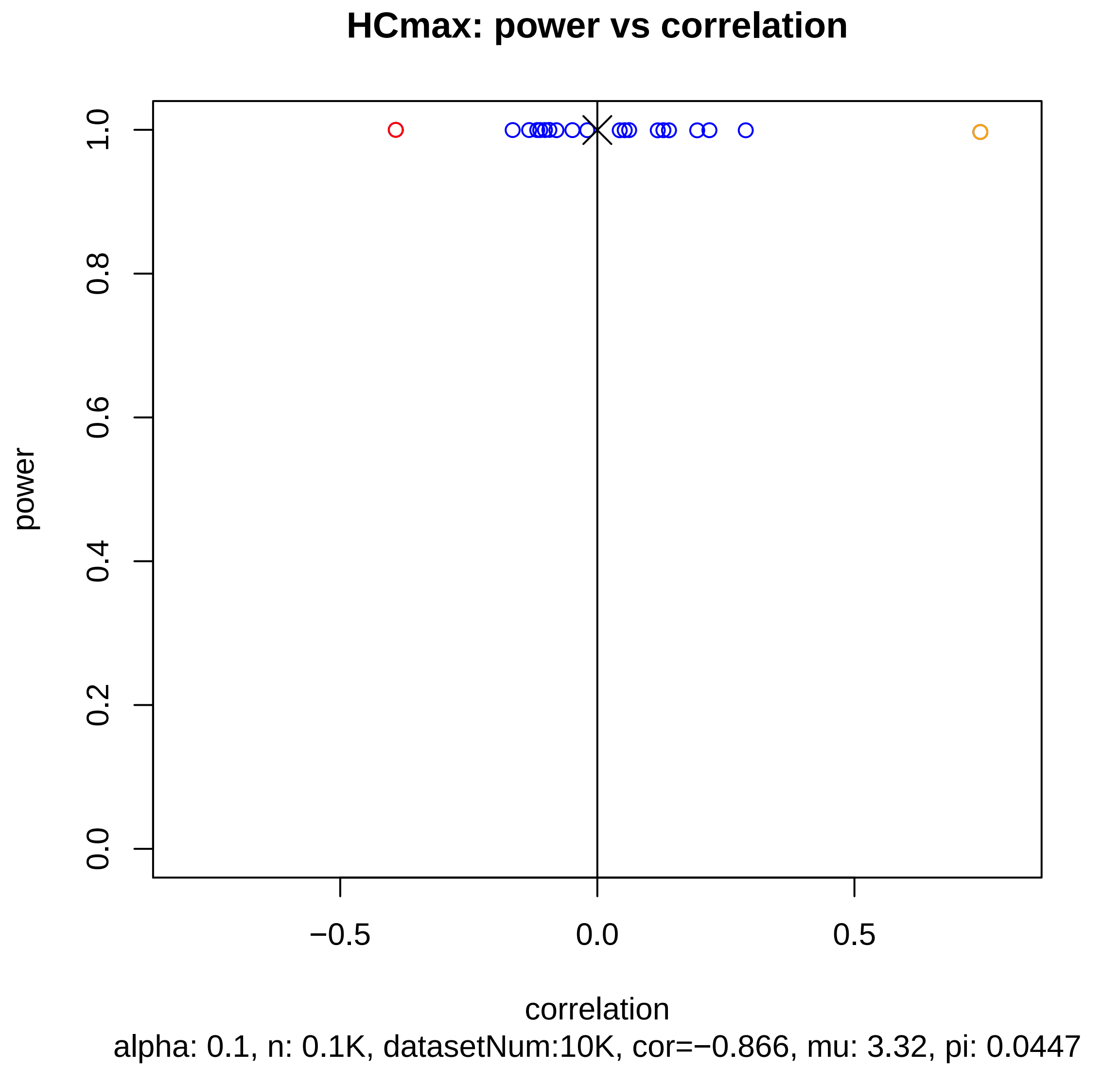
HCmax: power vs correlation
power
0.0
0.2
0.4
0.6
0.8
1.0
−0.5
0.0
0.5
correlation
alpha: 0.1, n: 0.1K, datasetNum:10K, cor=−0.866, mu: 3.32, pi: 0.0447

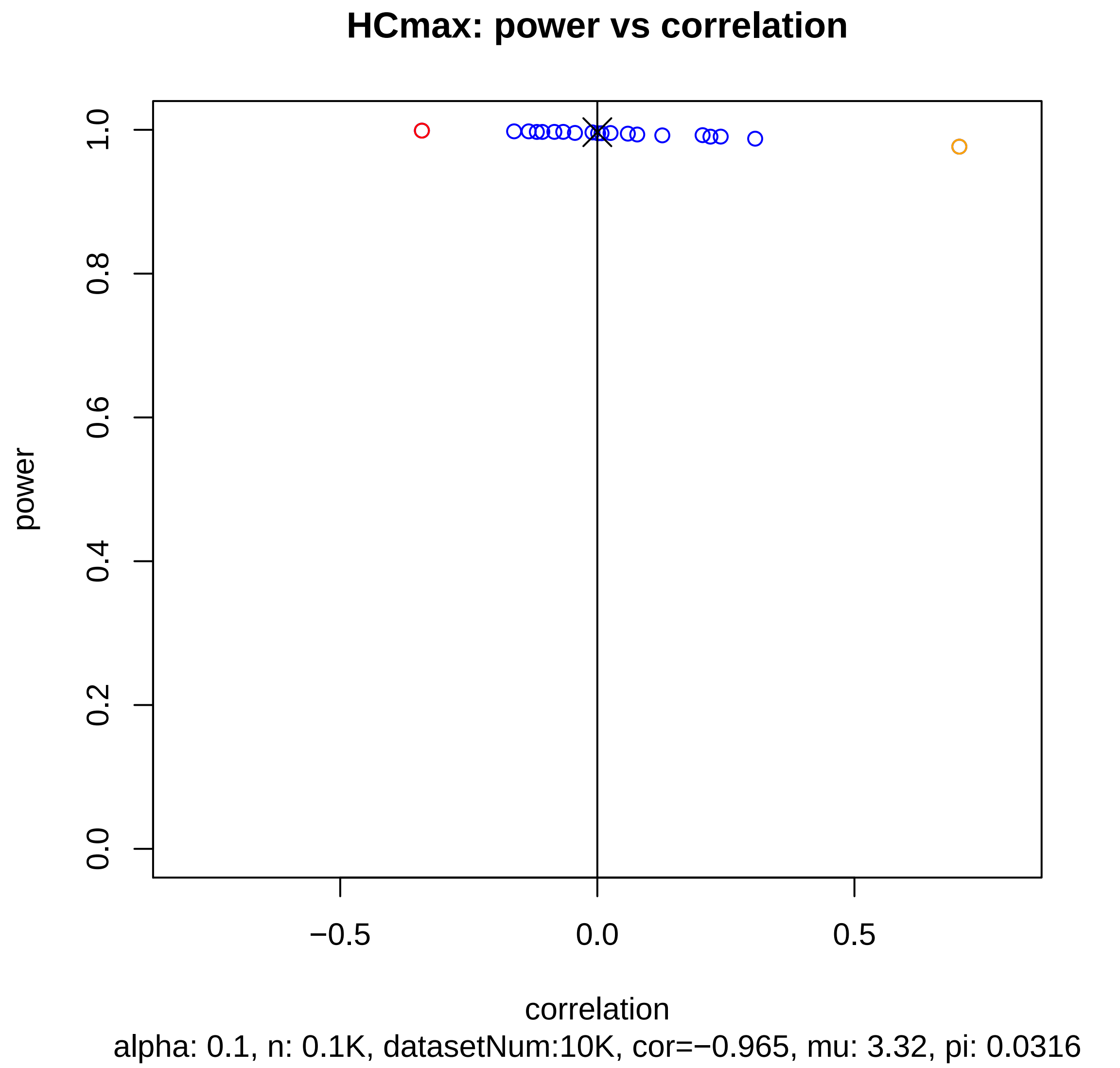
HCmax: power vs correlation
power
0.0
0.2
0.4
0.6
0.8
1.0
−0.5
0.0
0.5
correlation
alpha: 0.1, n: 0.1K, datasetNum:10K, cor=−0.965, mu: 3.32, pi: 0.0316

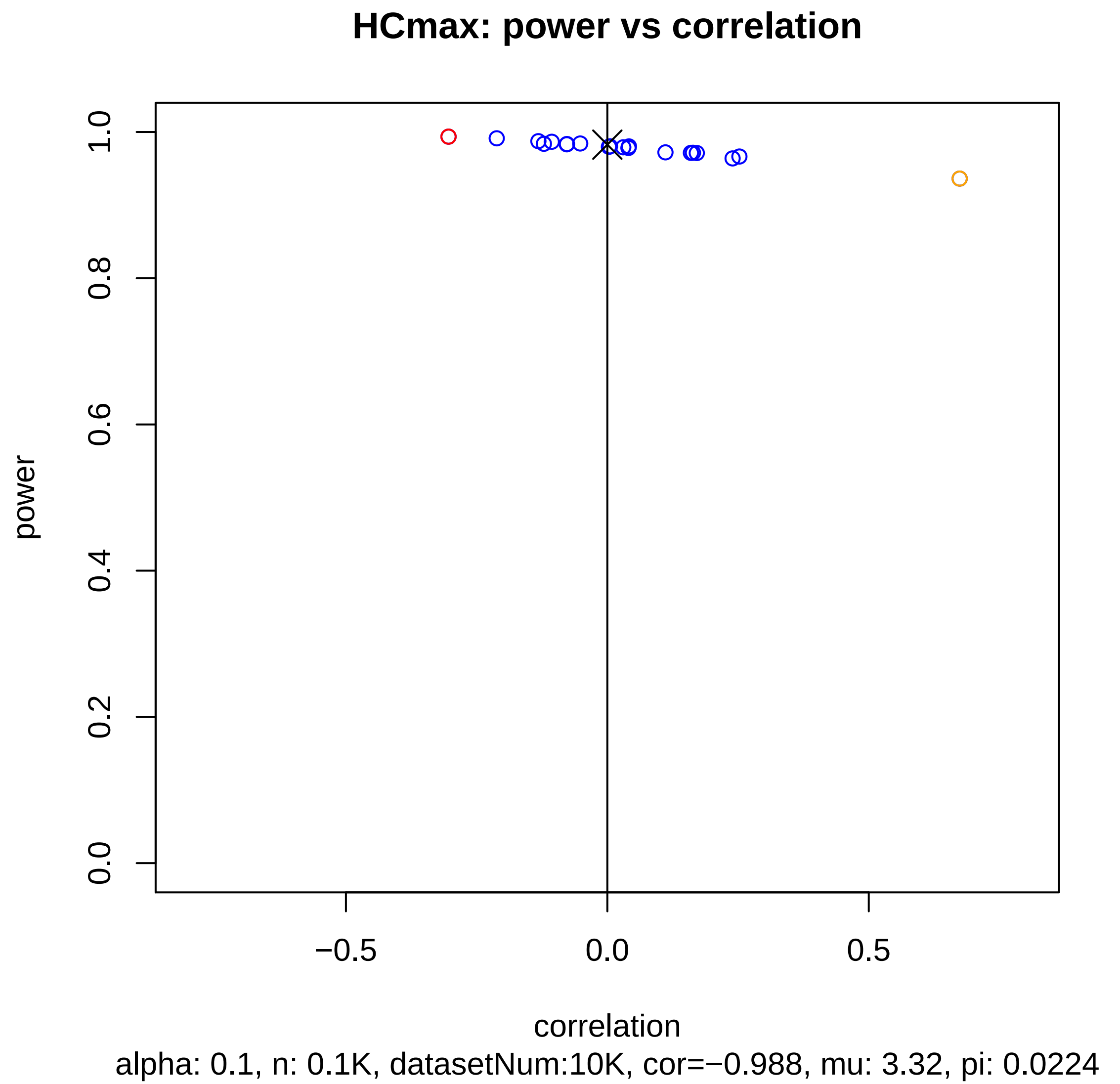
HCmax: power vs correlation
power
1.0
0.8
0.6
0.4
0.2
0.0
−0.5
0.0
0.5
correlation
alpha: 0.1, n: 0.1K, datasetNum:10K, cor=−0.988, mu: 3.32, pi: 0.0224

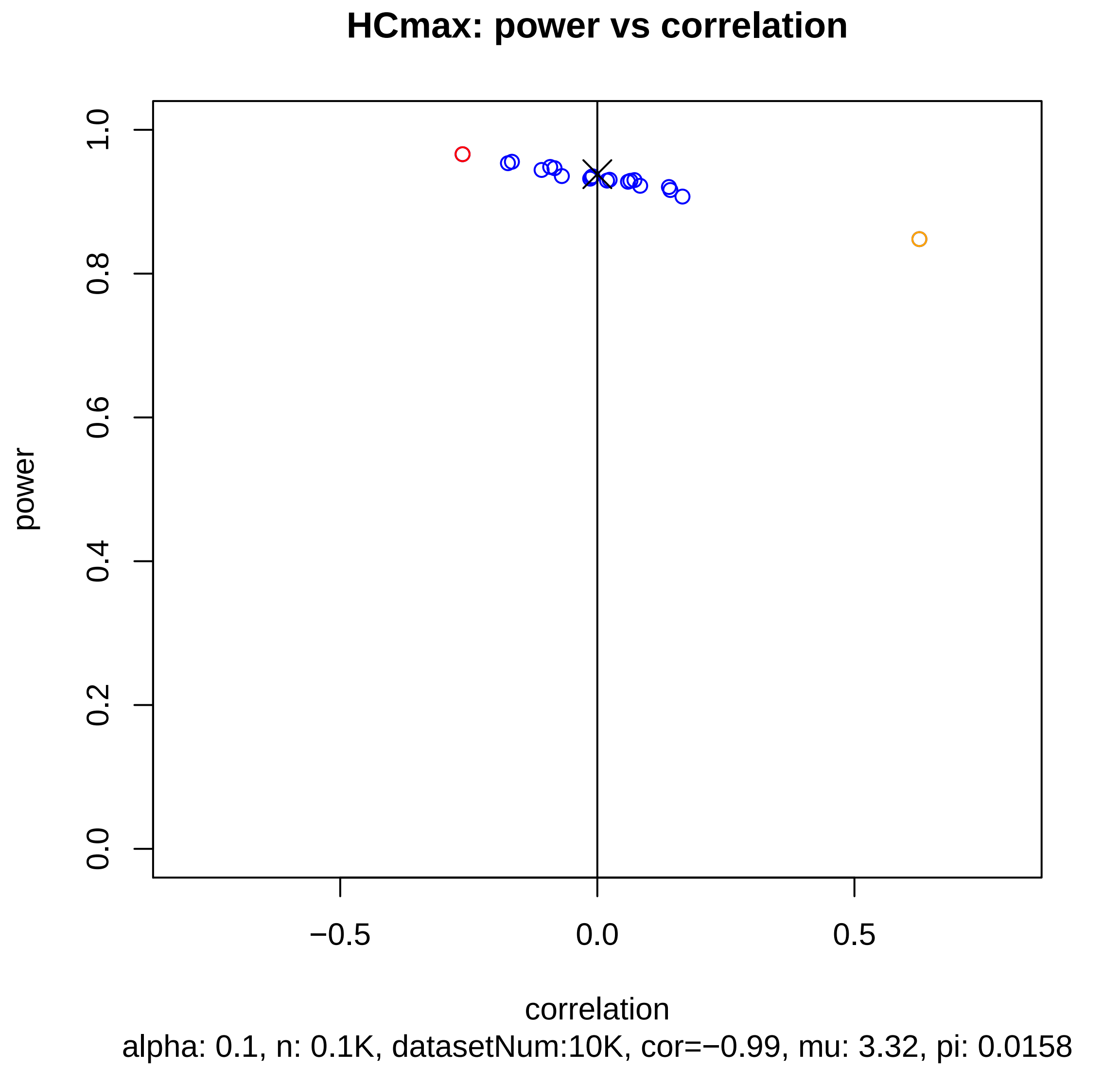
HCmax: power vs correlation
power
1.0
0.8
0.6
0.4
0.2
0.0
−0.5
0.0
0.5
correlation
alpha: 0.1, n: 0.1K, datasetNum:10K, cor=−0.99, mu: 3.32, pi: 0.0158

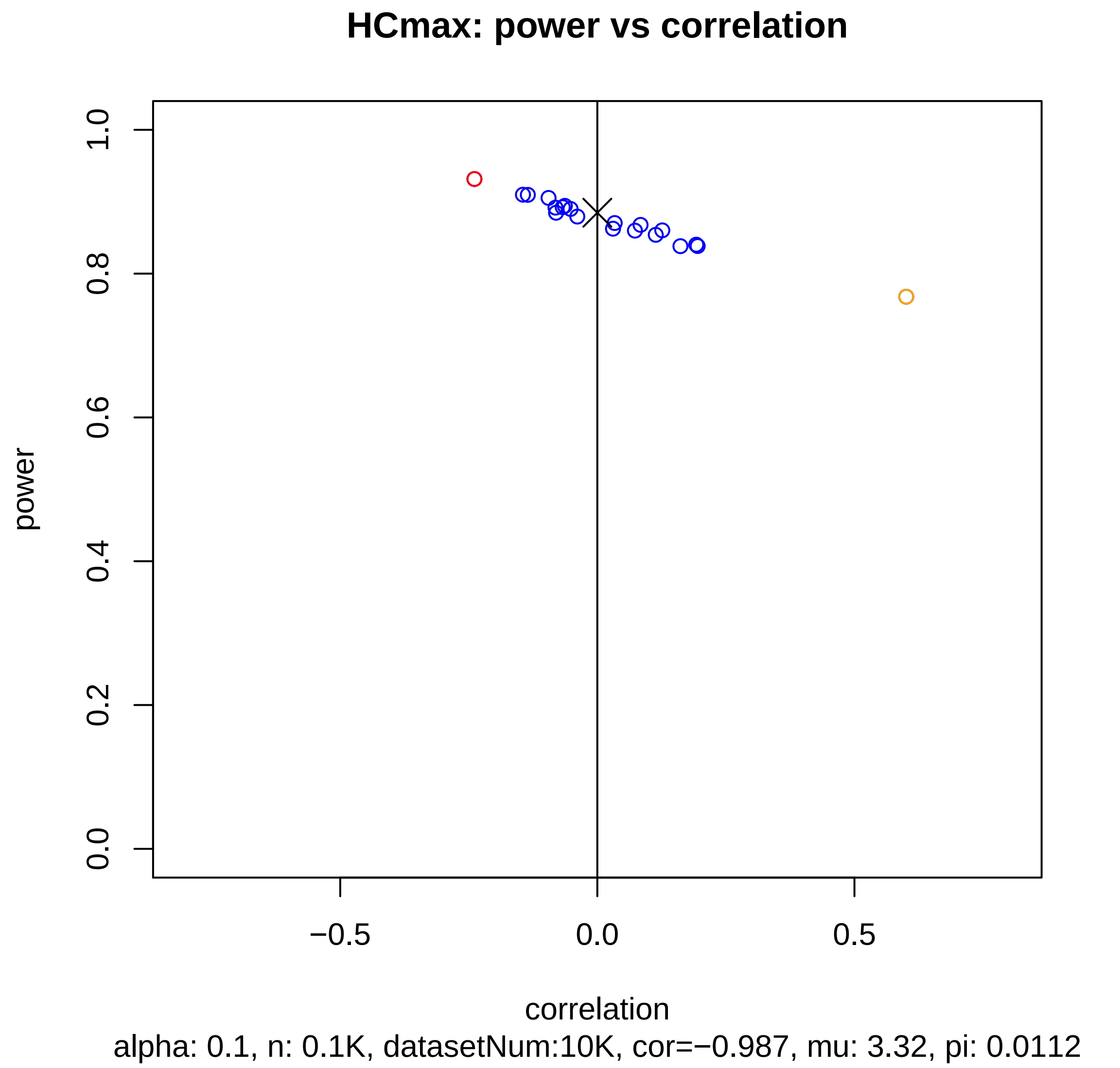
HCmax: power vs correlation
power
1.0
0.8
0.6
0.4
0.2
0.0
−0.5
0.0
0.5
correlation
alpha: 0.1, n: 0.1K, datasetNum:10K, cor=−0.987, mu: 3.32, pi: 0.0112

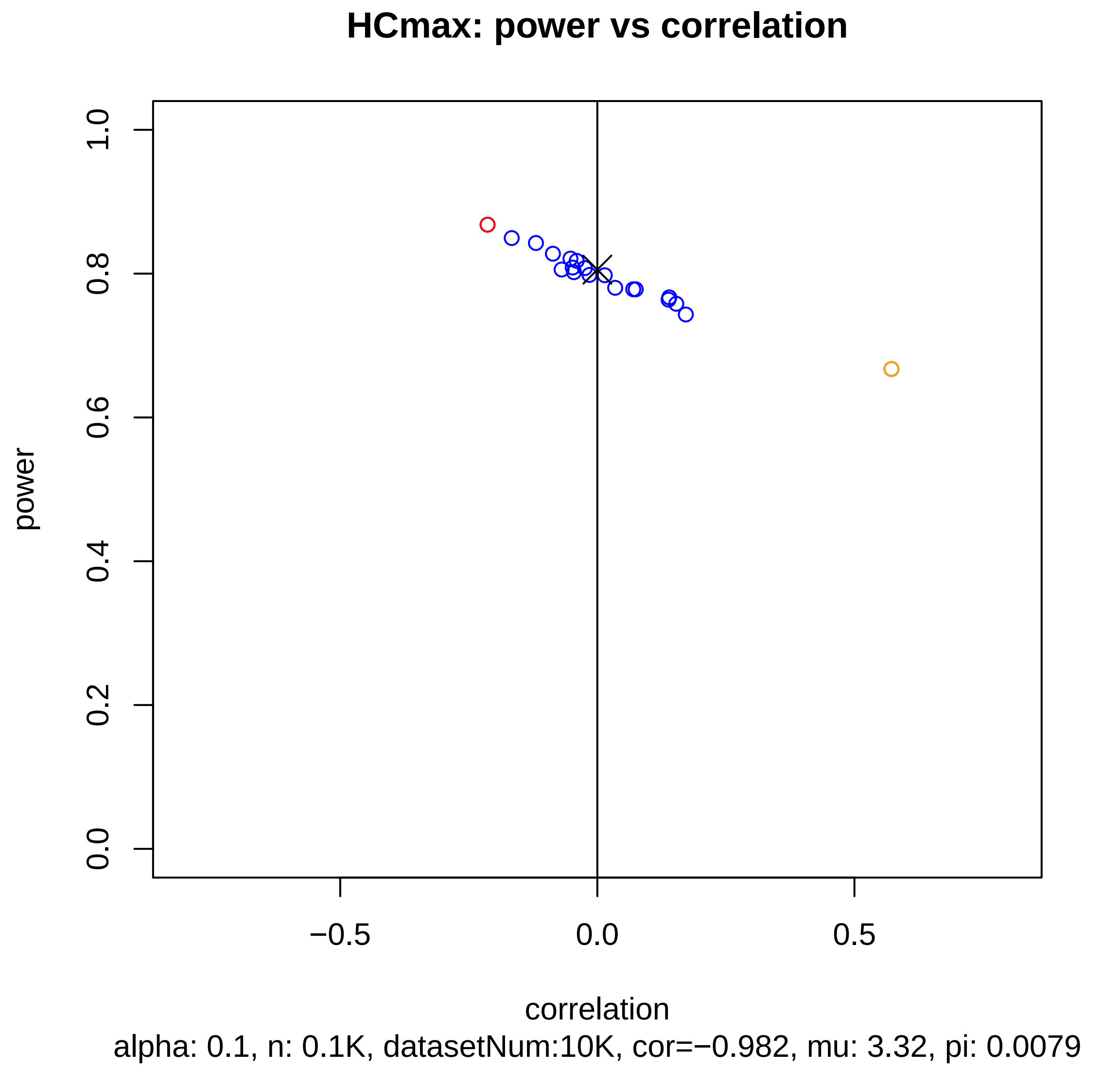
HCmax: power vs correlation
power
1.0
0.8
0.6
0.4
0.2
0.0
−0.5
0.0
0.5
correlation
alpha: 0.1, n: 0.1K, datasetNum:10K, cor=−0.982, mu: 3.32, pi: 0.0079

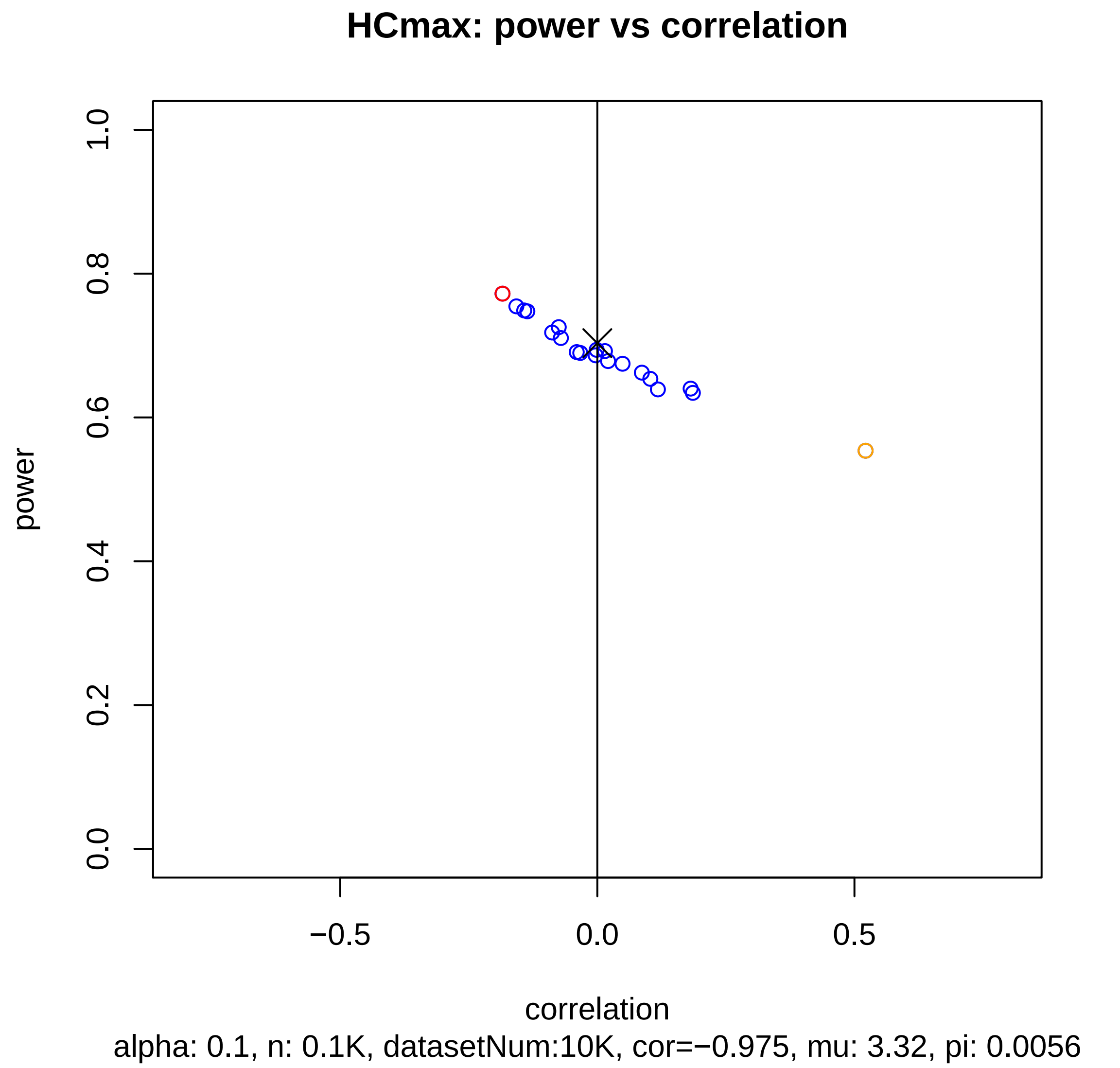
HCmax: power vs correlation
power
correlation
1.0
0.8
0.6
0.4
0.2
0.0
−0.5
0.0
0.5
alpha: 0.1, n: 0.1K, datasetNum:10K, cor=−0.975, mu: 3.32, pi: 0.0056

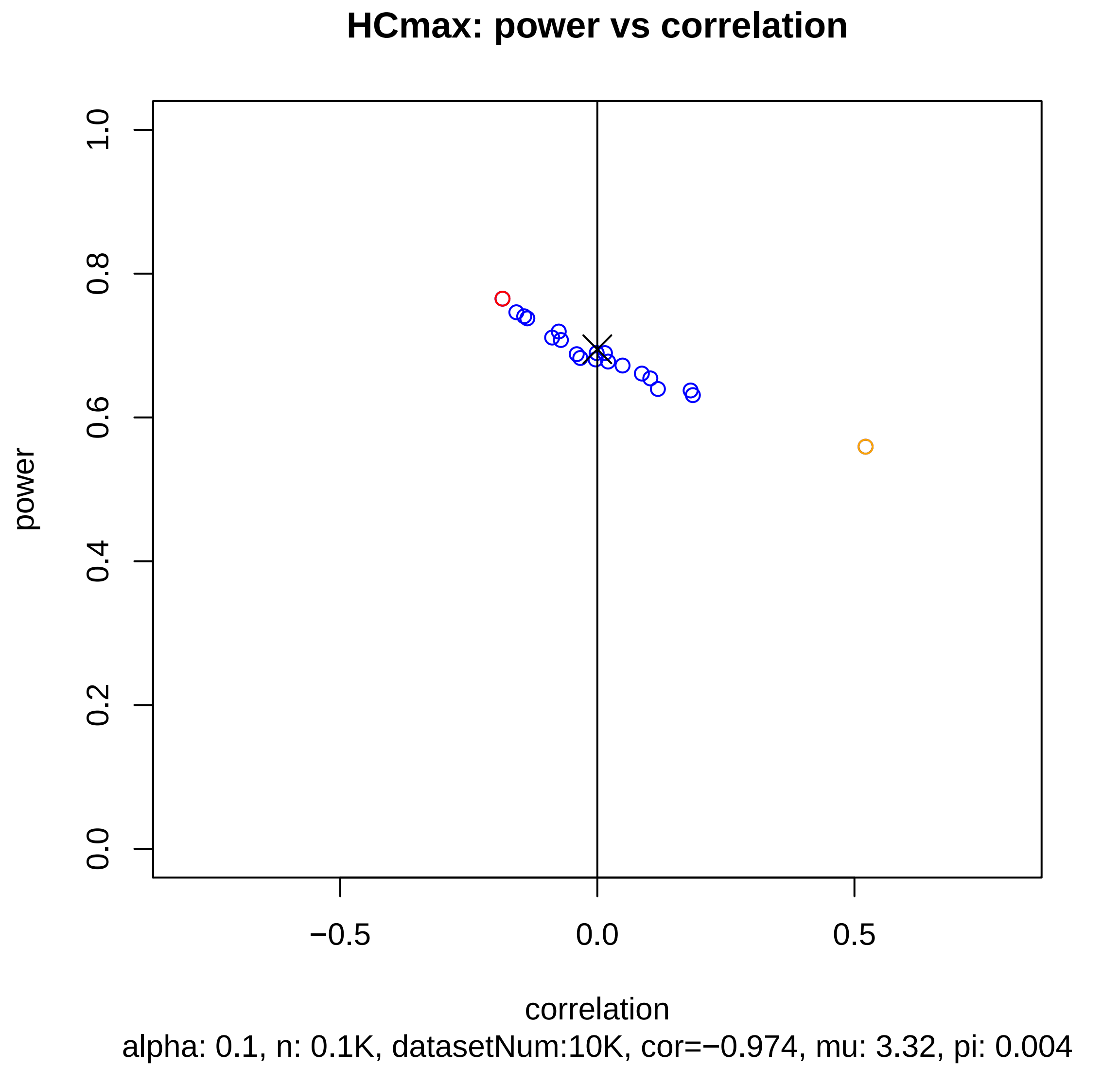

HCmax: power vs correlation
power
0.0
0.2
0.4
0.6
0.8
1.0
−0.5
0.0
0.5
correlation
alpha: 0.1, n: 0.1K, datasetNum:10K, cor=−0.974, mu: 3.32, pi: 0.004

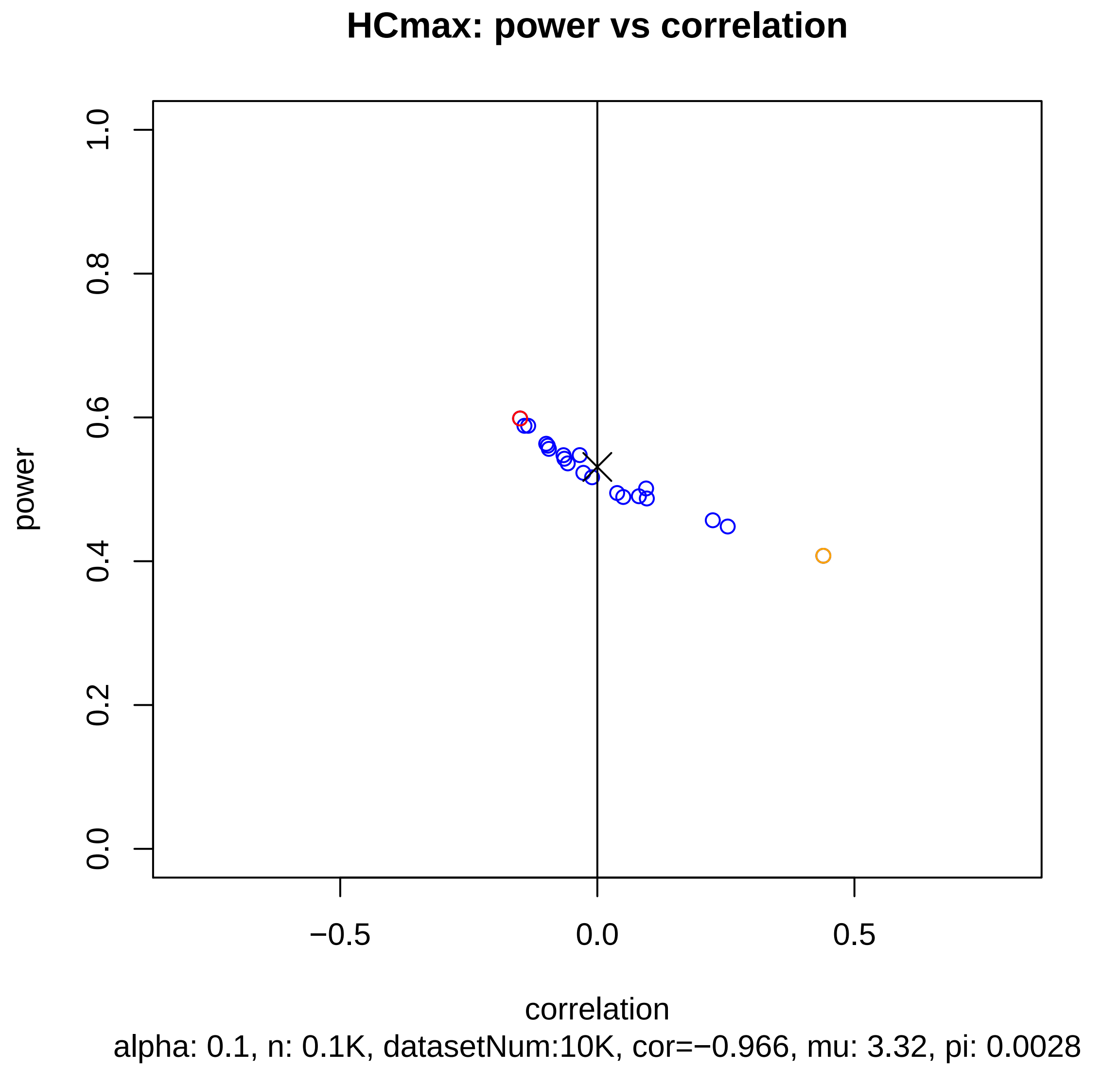
HCmax: power vs correlation
power
1.0
0.8
0.6
0.4
0.2
0.0
−0.5
0.0
0.5
correlation
alpha: 0.1, n: 0.1K, datasetNum:10K, cor=−0.966, mu: 3.32, pi: 0.0028

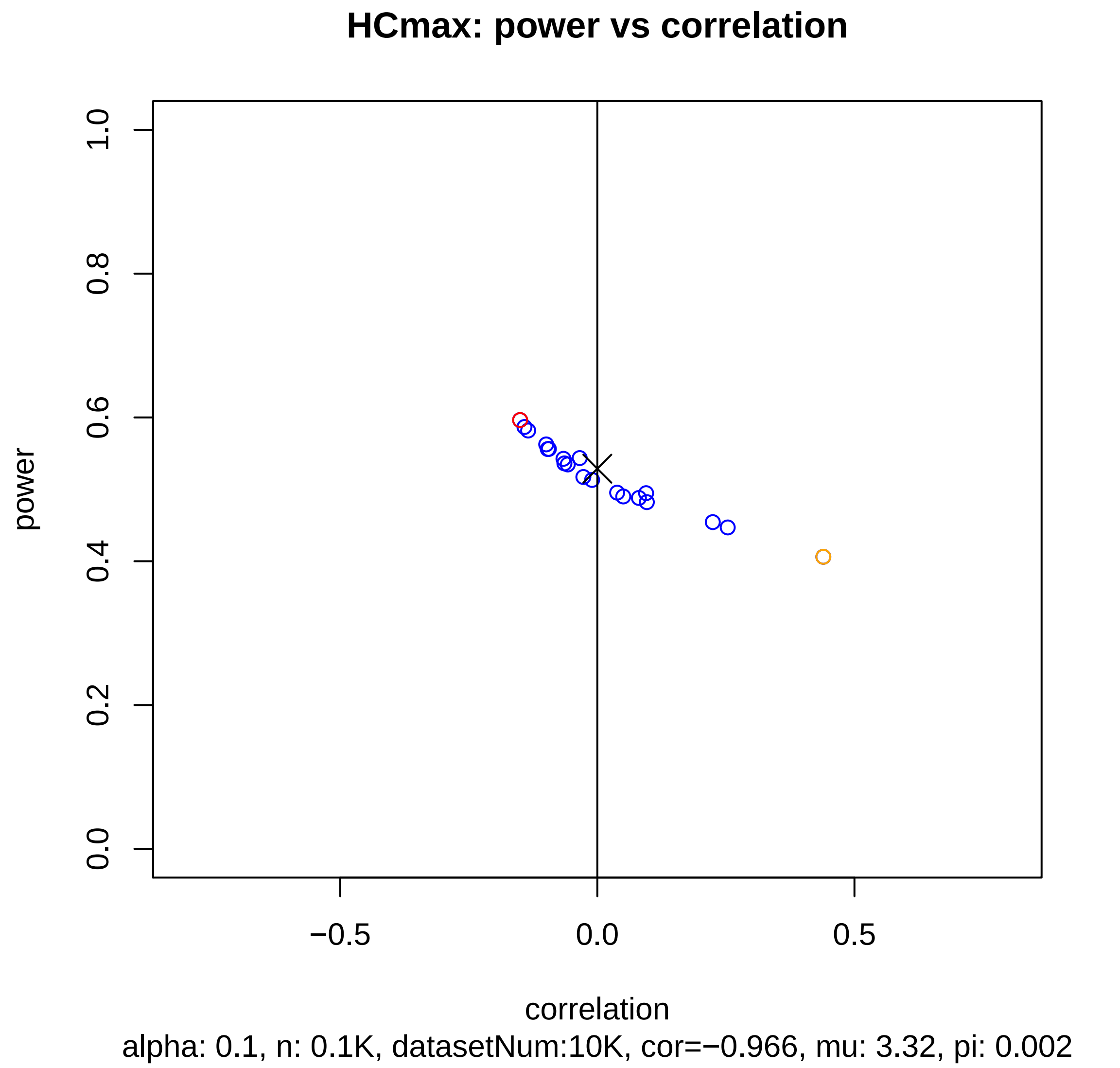
HCmax: power vs correlation
power
1.0
0.8
0.6
0.4
0.2
0.0
−0.5
0.0
0.5
correlation
alpha: 0.1, n: 0.1K, datasetNum:10K, cor=−0.966, mu: 3.32, pi: 0.002

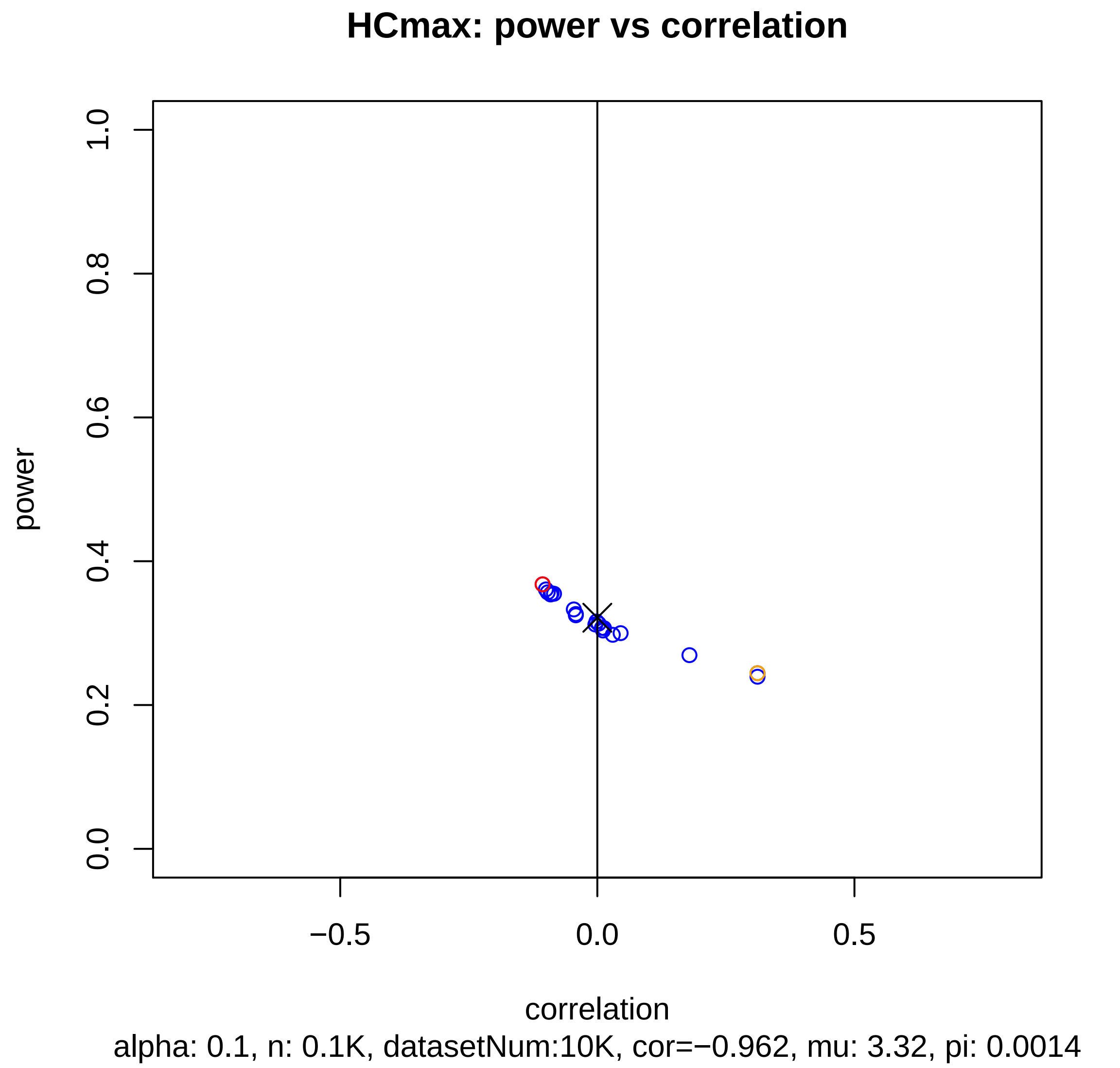

HCmax: power vs correlation
power
0.0
0.2
0.4
0.6
0.8
1.0
−0.5
0.0
0.5
correlation
alpha: 0.1, n: 0.1K, datasetNum:10K, cor=−0.962, mu: 3.32, pi: 0.0014

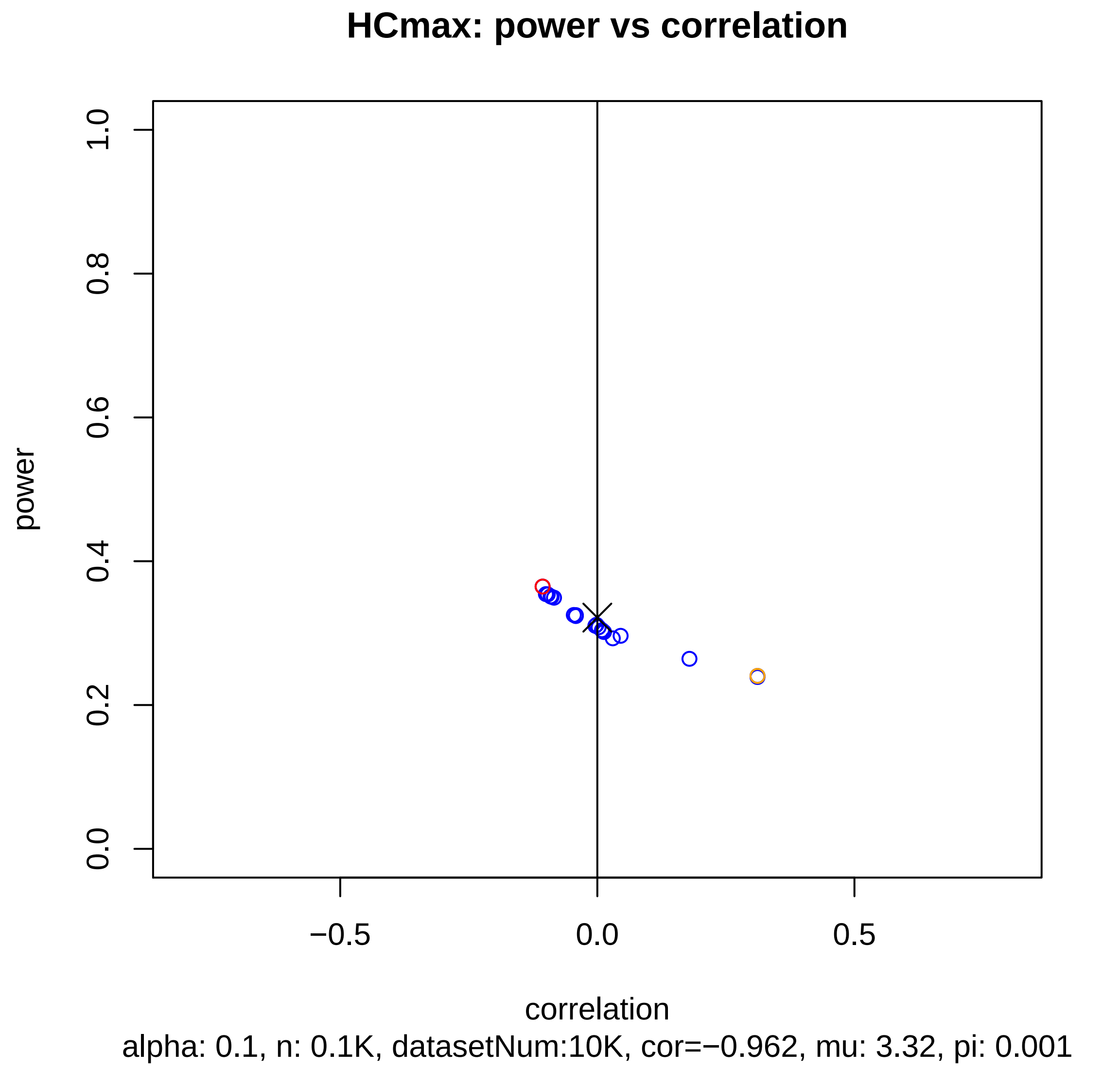
HCmax: power vs correlation
power
0.0
0.2
0.4
0.6
0.8
1.0
−0.5
0.0
0.5
correlation
alpha: 0.1, n: 0.1K, datasetNum:10K, cor=−0.962, mu: 3.32, pi: 0.001

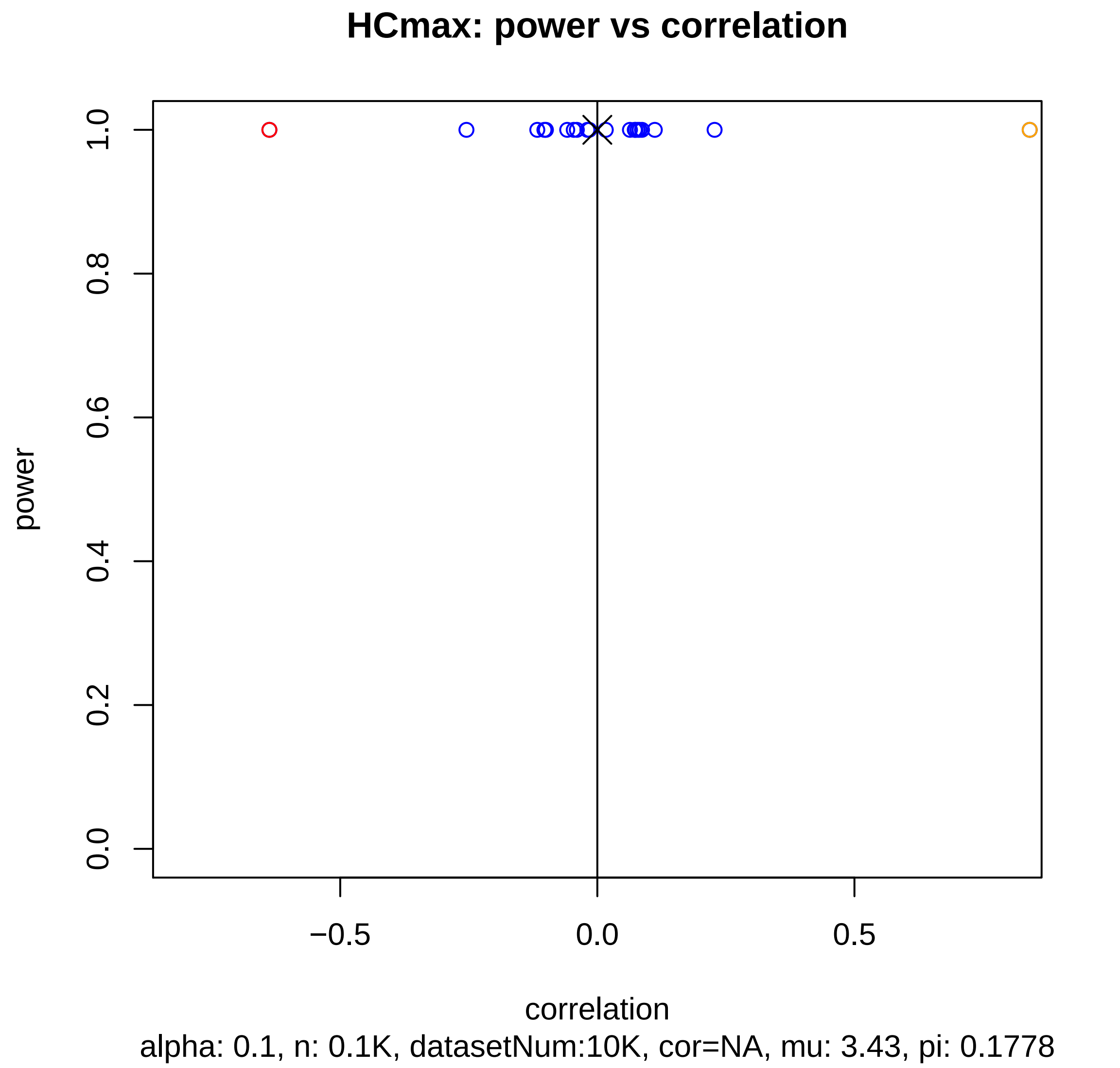
HCmax: power vs correlation
1.0
0.8
0.6
0.4
0.2
0.0
power
−0.5
0.0
0.5
correlation
alpha: 0.1, n: 0.1K, datasetNum:10K, cor=NA, mu: 3.43, pi: 0.1778

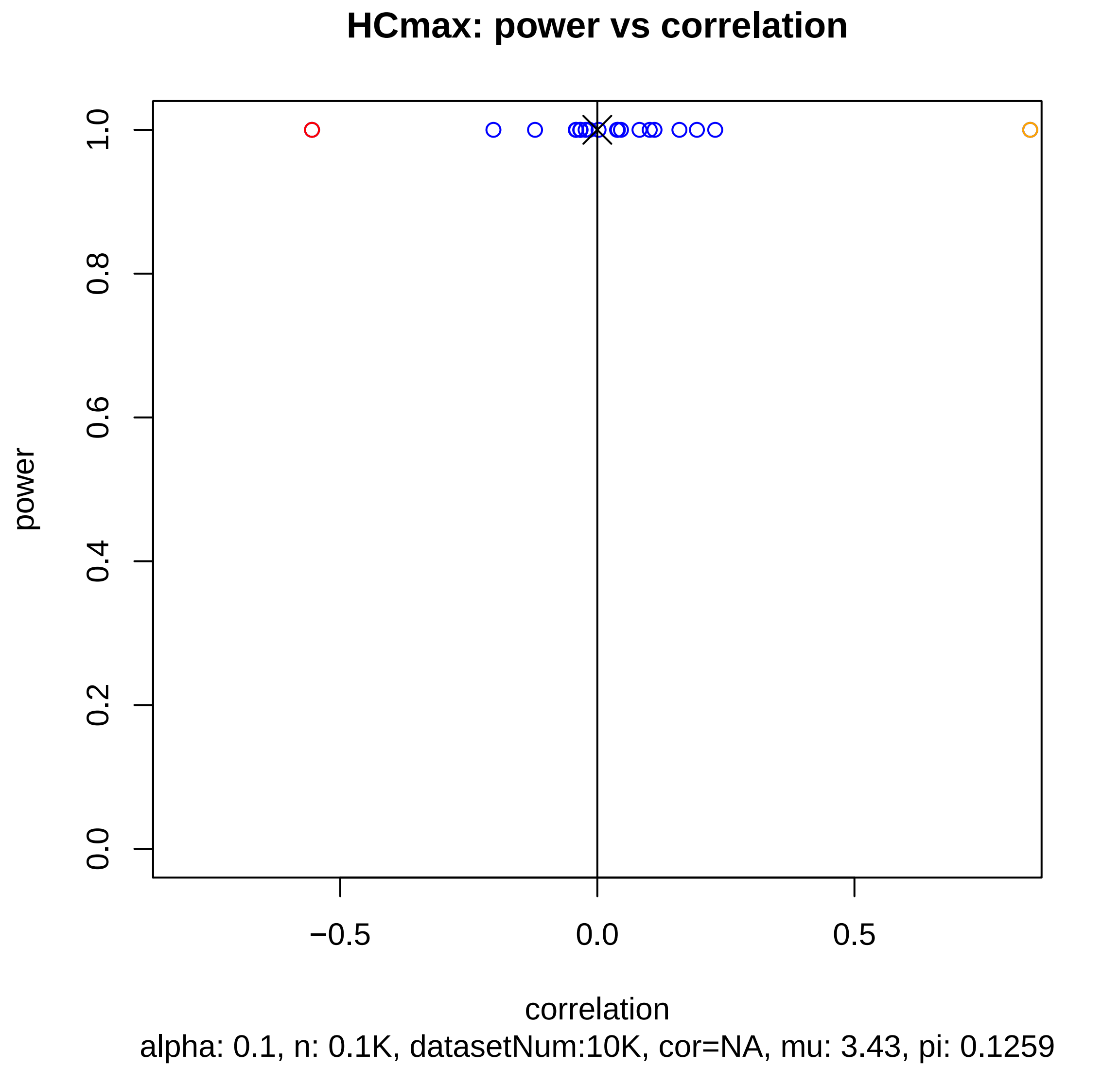

HCmax: power vs correlation
power
1.0
0.8
0.6
0.4
0.2
0.0
−0.5
0.0
0.5
correlation
alpha: 0.1, n: 0.1K, datasetNum:10K, cor=NA, mu: 3.43, pi: 0.1259

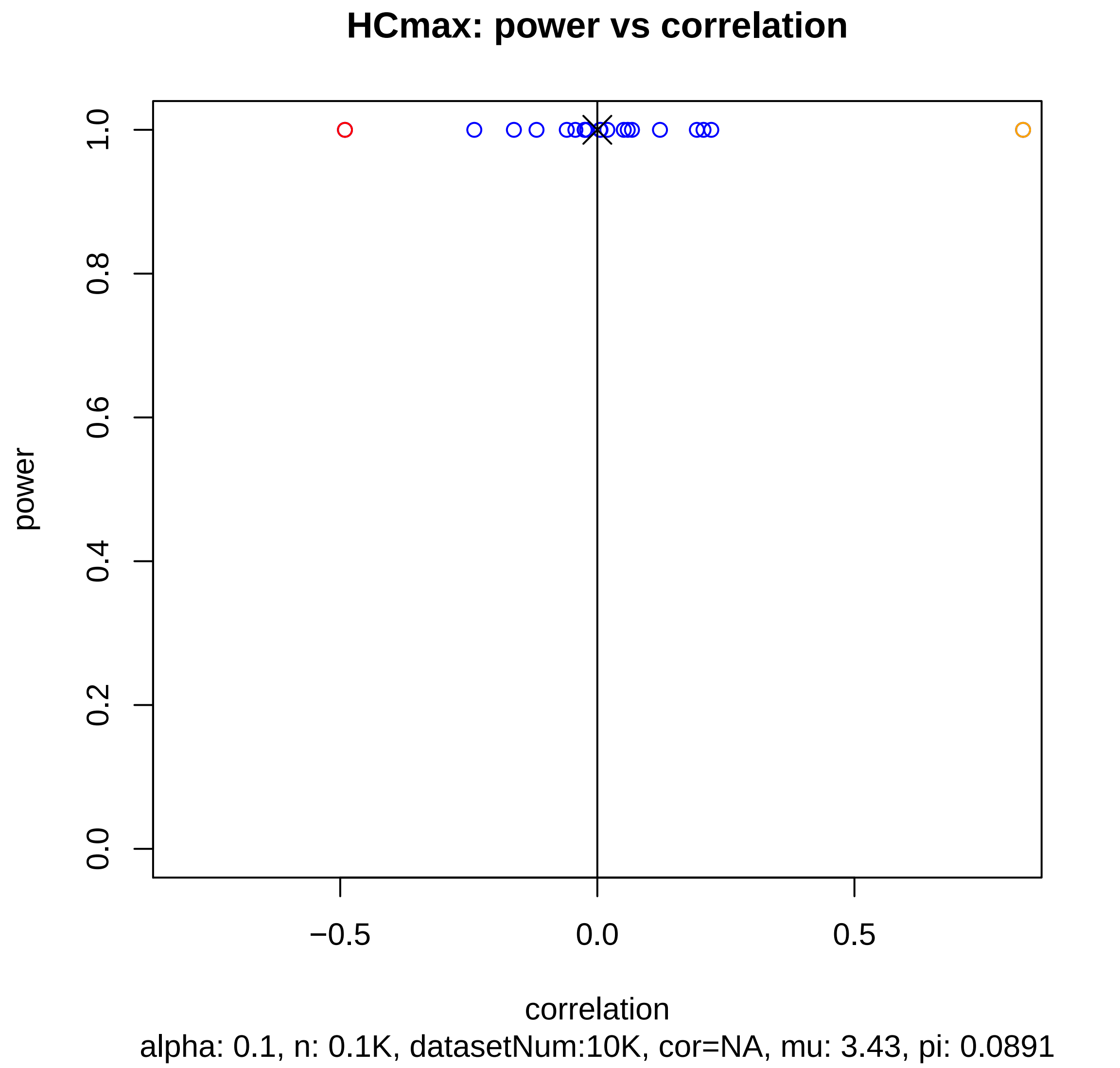
HCmax: power vs correlation
power
1.0
0.8
0.6
0.4
0.2
0.0
−0.5
0.0
0.5
correlation
alpha: 0.1, n: 0.1K, datasetNum:10K, cor=NA, mu: 3.43, pi: 0.0891

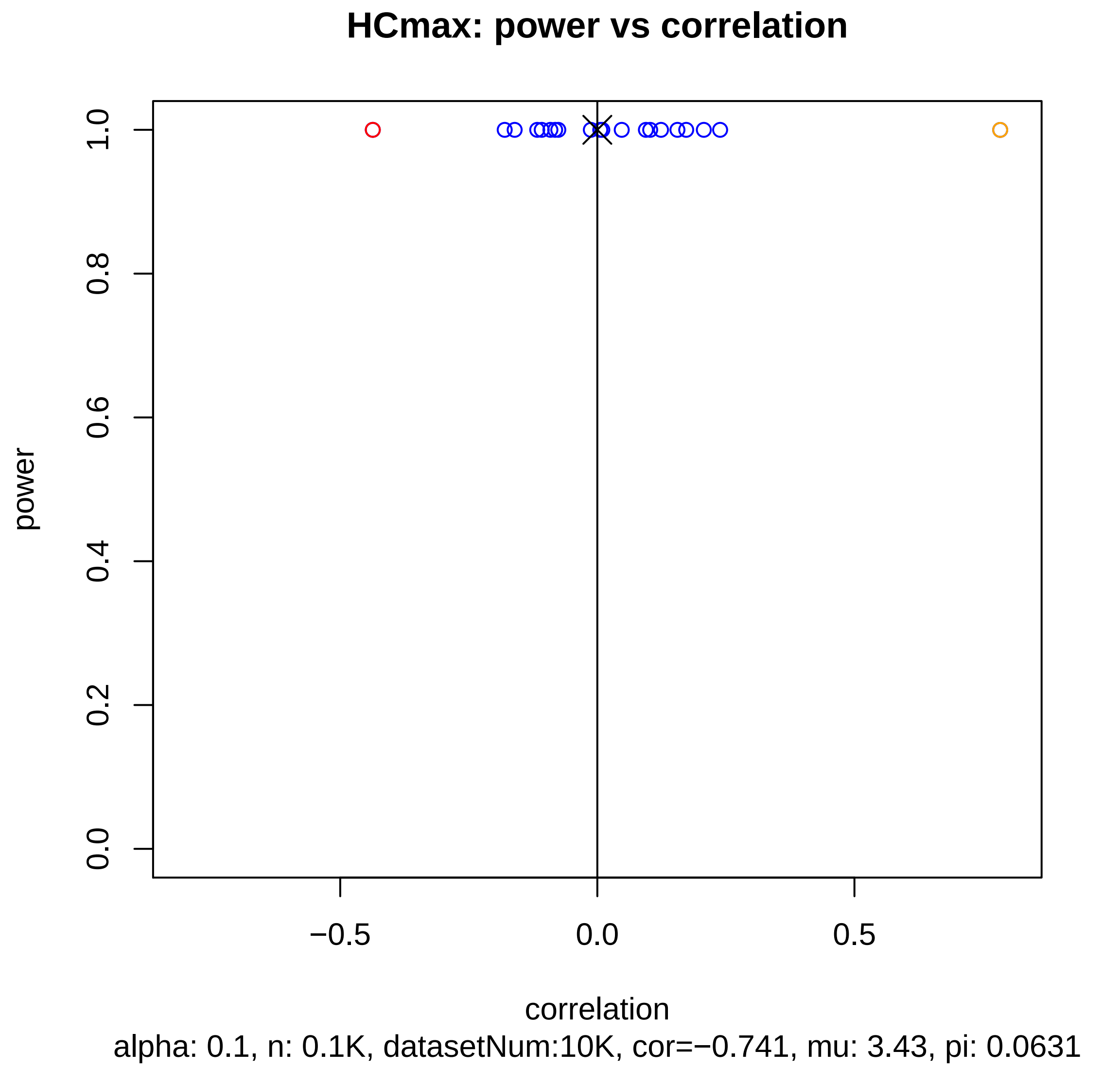
HCmax: power vs correlation
power
1.0
0.8
0.6
0.4
0.2
0.0
−0.5
0.0
0.5
correlation
alpha: 0.1, n: 0.1K, datasetNum:10K, cor=−0.741, mu: 3.43, pi: 0.0631

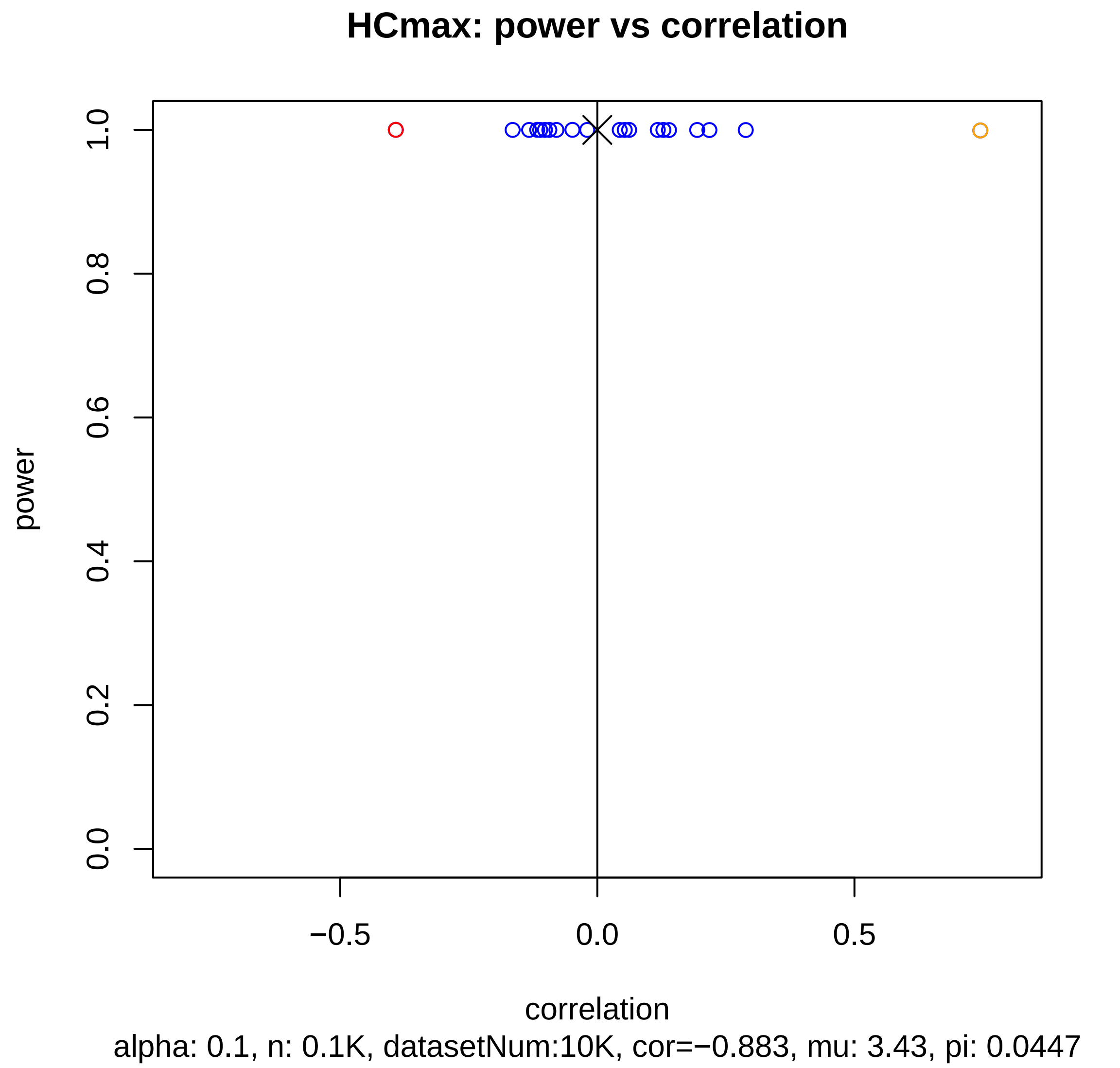
HCmax: power vs correlation
power
1.0
0.8
0.6
0.4
0.2
0.0
−0.5
0.0
0.5
correlation
alpha: 0.1, n: 0.1K, datasetNum:10K, cor=−0.883, mu: 3.43, pi: 0.0447

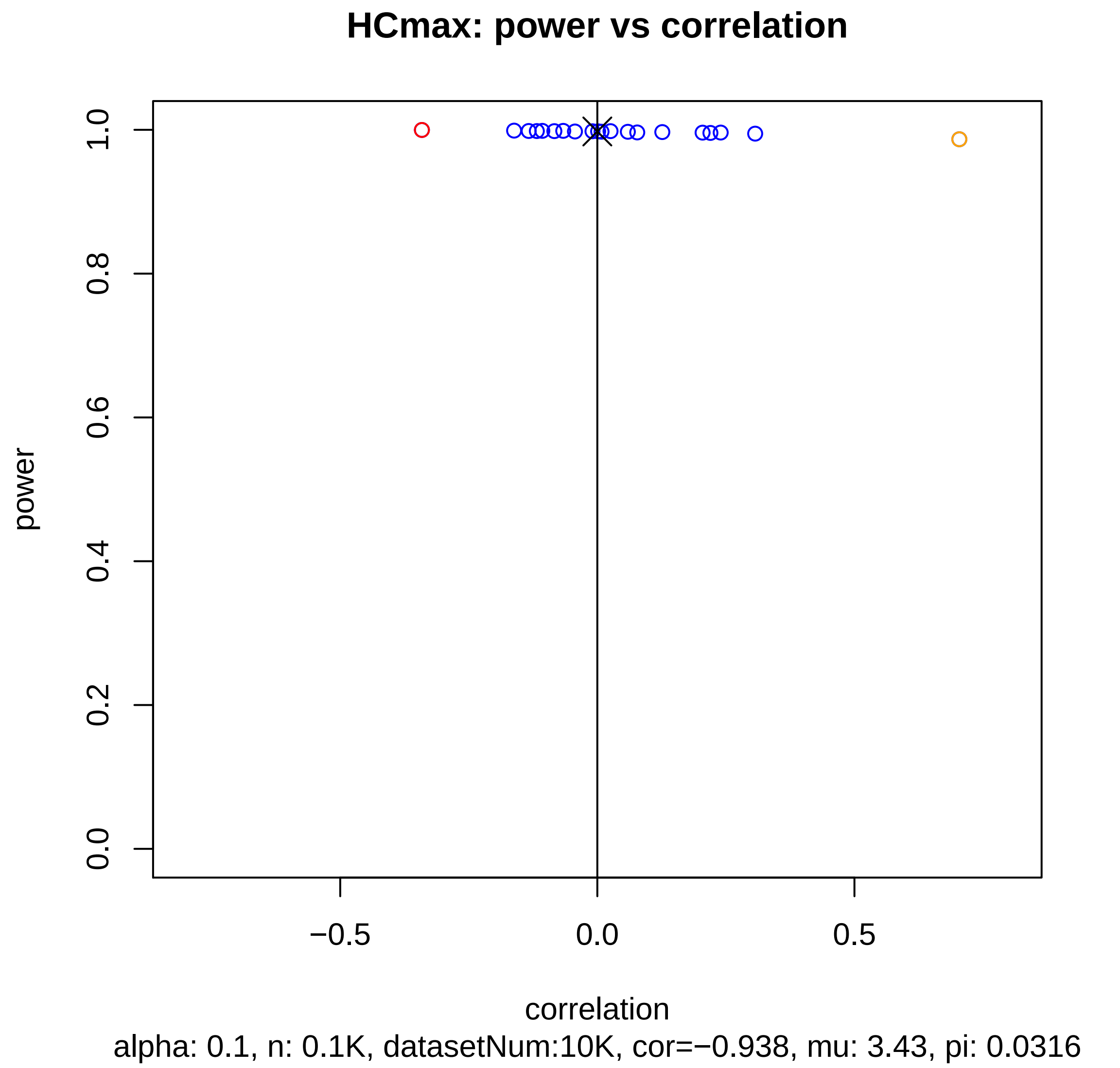
HCmax: power vs correlation
1.0
0.8
0.6
0.4
0.2
0.0
power
−0.5
0.0
0.5
correlation
alpha: 0.1, n: 0.1K, datasetNum:10K, cor=−0.938, mu: 3.43, pi: 0.0316

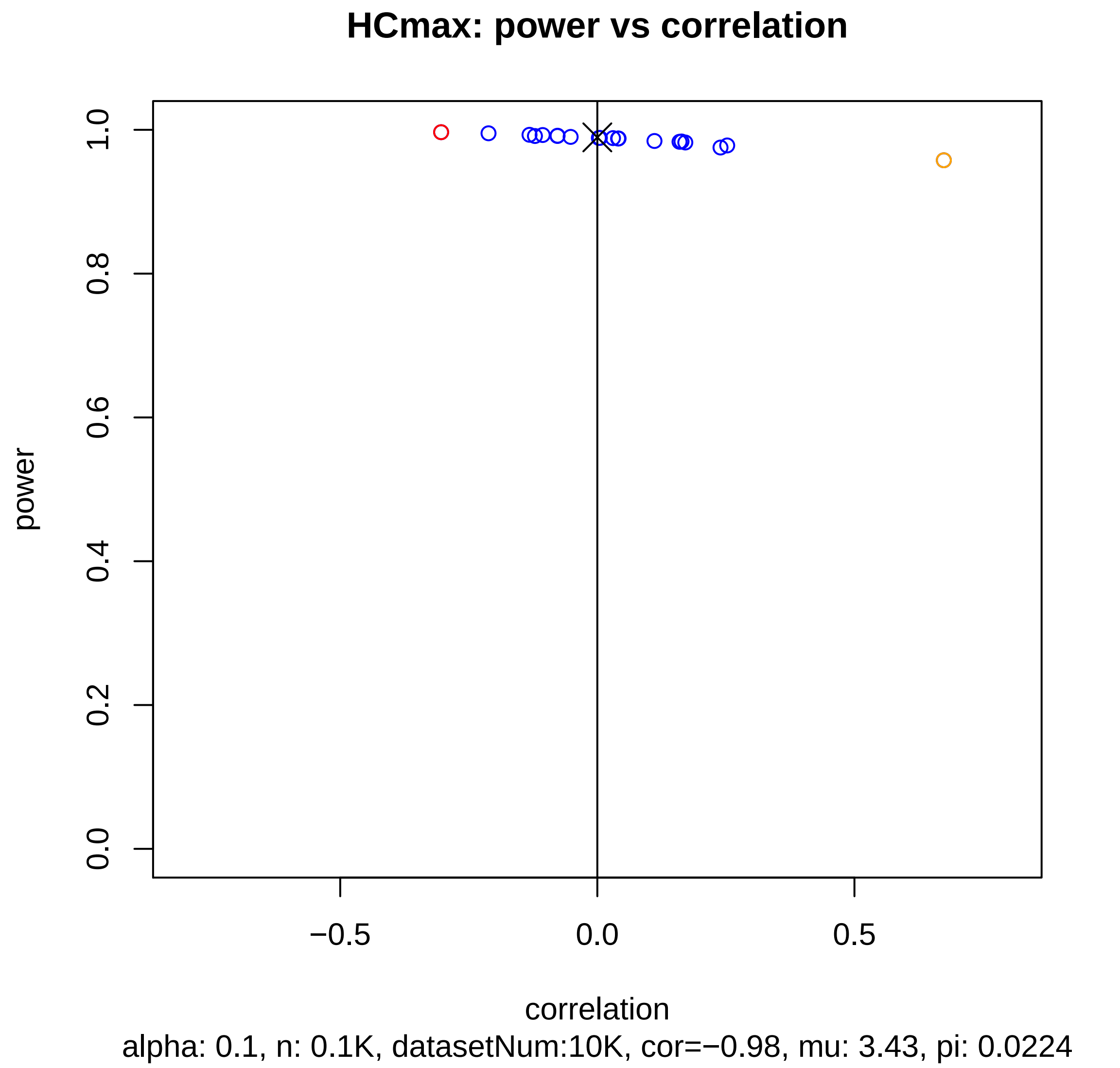
HCmax: power vs correlation
power
0.0
0.2
0.4
0.6
0.8
1.0
−0.5
0.0
0.5
correlation
alpha: 0.1, n: 0.1K, datasetNum:10K, cor=−0.98, mu: 3.43, pi: 0.0224

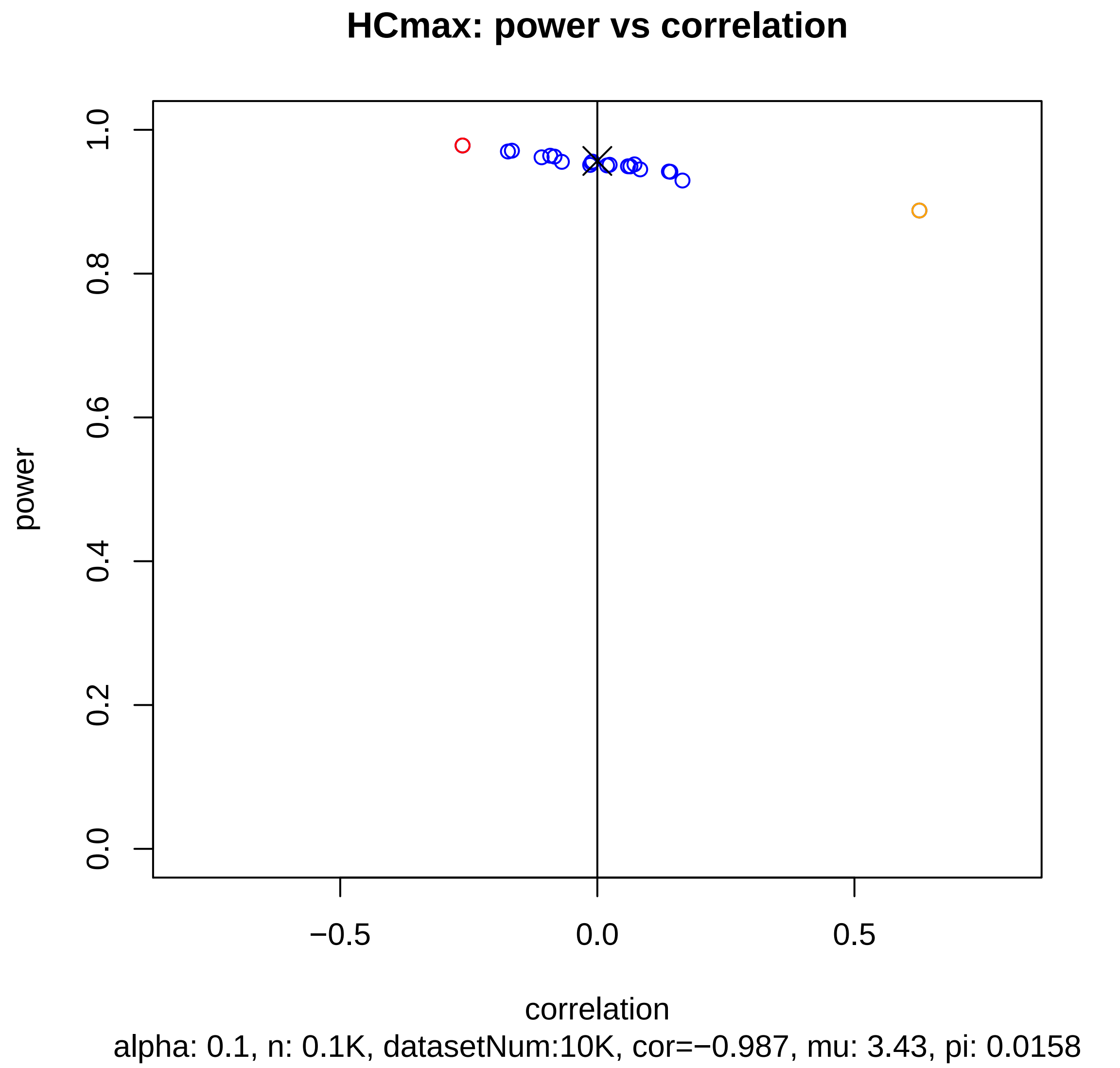
HCmax: power vs correlation
power
1.0
0.8
0.6
0.4
0.2
0.0
−0.5
0.0
0.5
correlation
alpha: 0.1, n: 0.1K, datasetNum:10K, cor=−0.987, mu: 3.43, pi: 0.0158

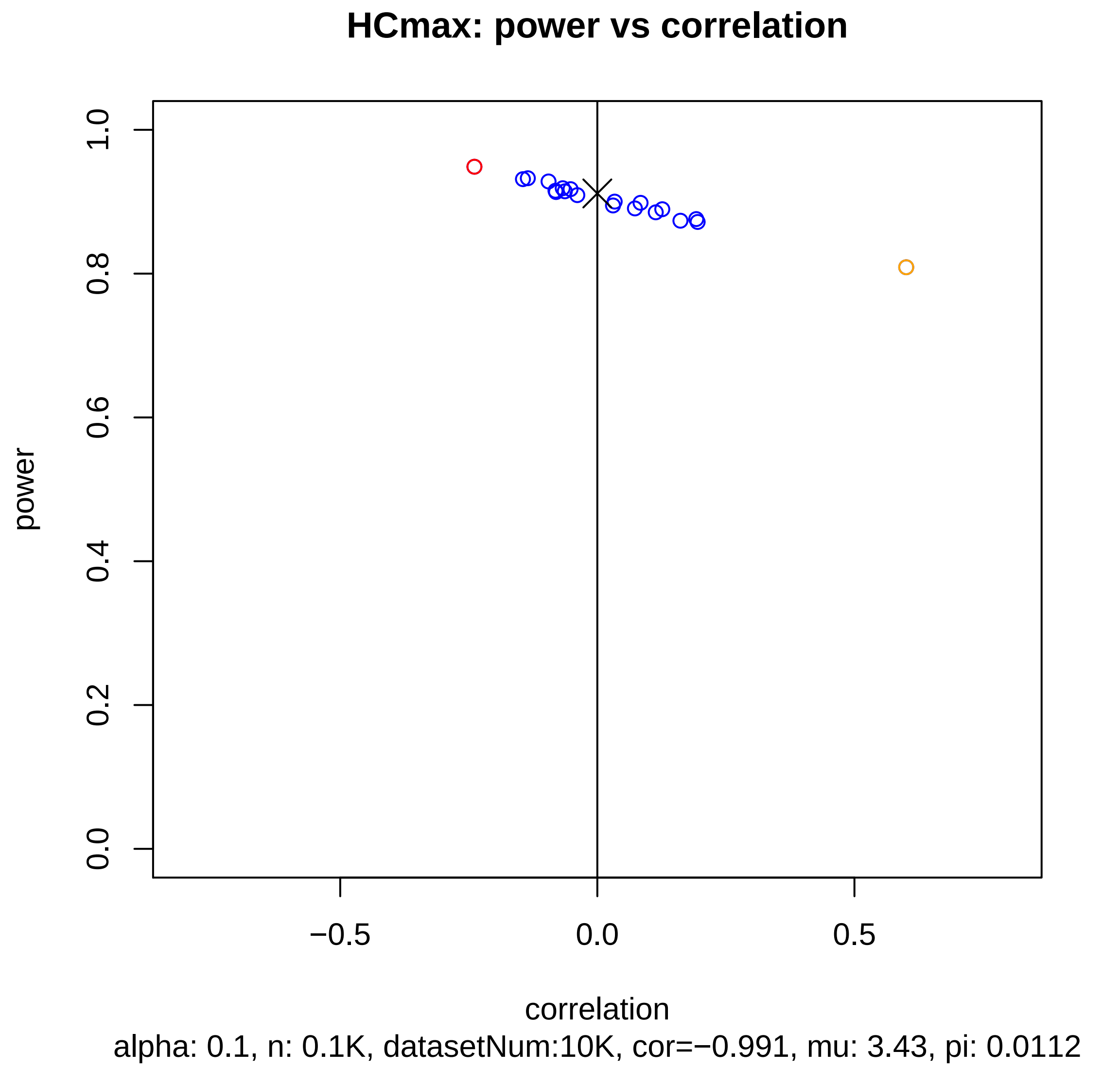
HCmax: power vs correlation
power
0.0
0.2
0.4
0.6
0.8
1.0
−0.5
0.0
0.5
correlation
alpha: 0.1, n: 0.1K, datasetNum:10K, cor=−0.991, mu: 3.43, pi: 0.0112

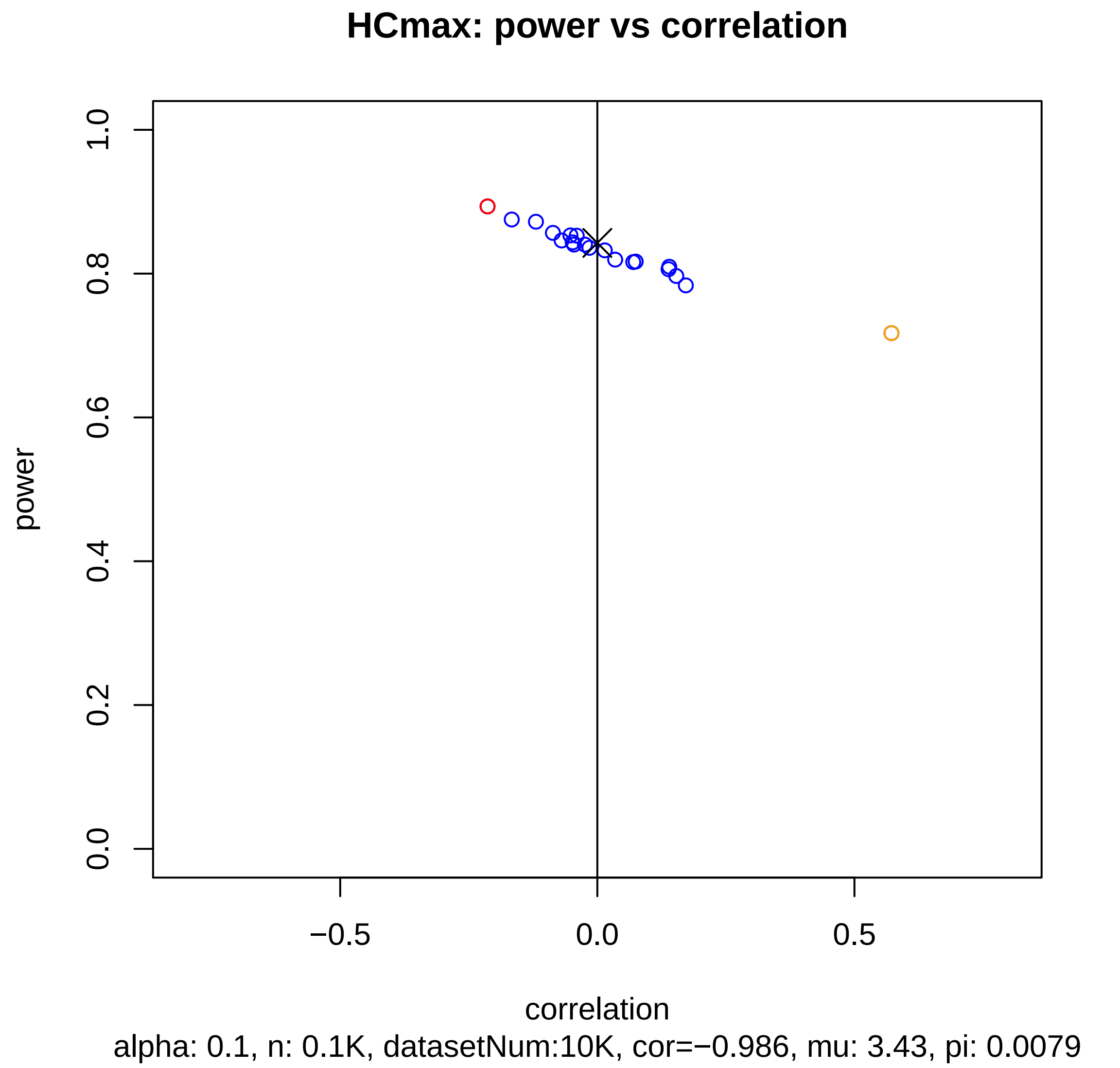
HCmax: power vs correlation
power
1.0
0.8
0.6
0.4
0.2
0.0
−0.5
0.0
0.5
correlation
alpha: 0.1, n: 0.1K, datasetNum:10K, cor=−0.986, mu: 3.43, pi: 0.0079

### HCmax: power vs correlation

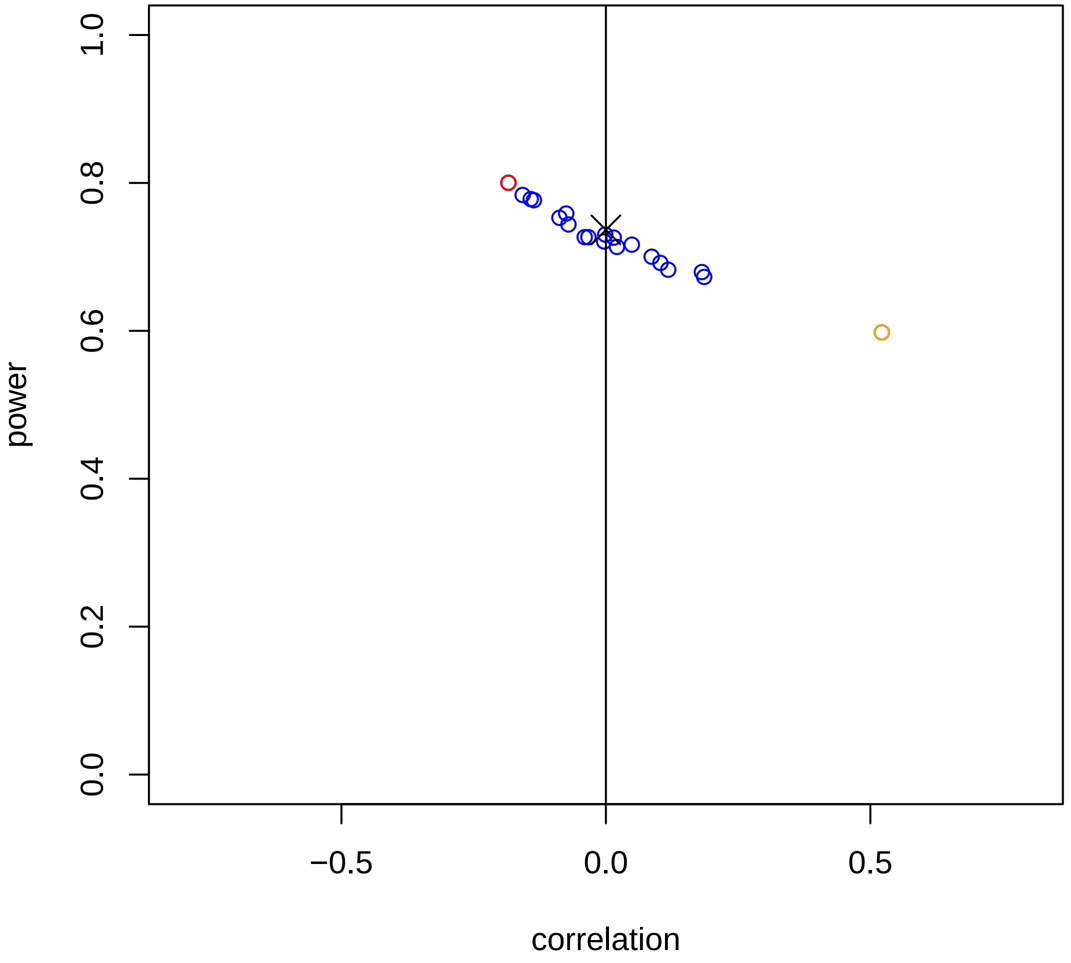


alpha: 0.1, n: 0.1K, datasetNum:10K, cor=−0.978, mu: 3.43, pi: 0.0056

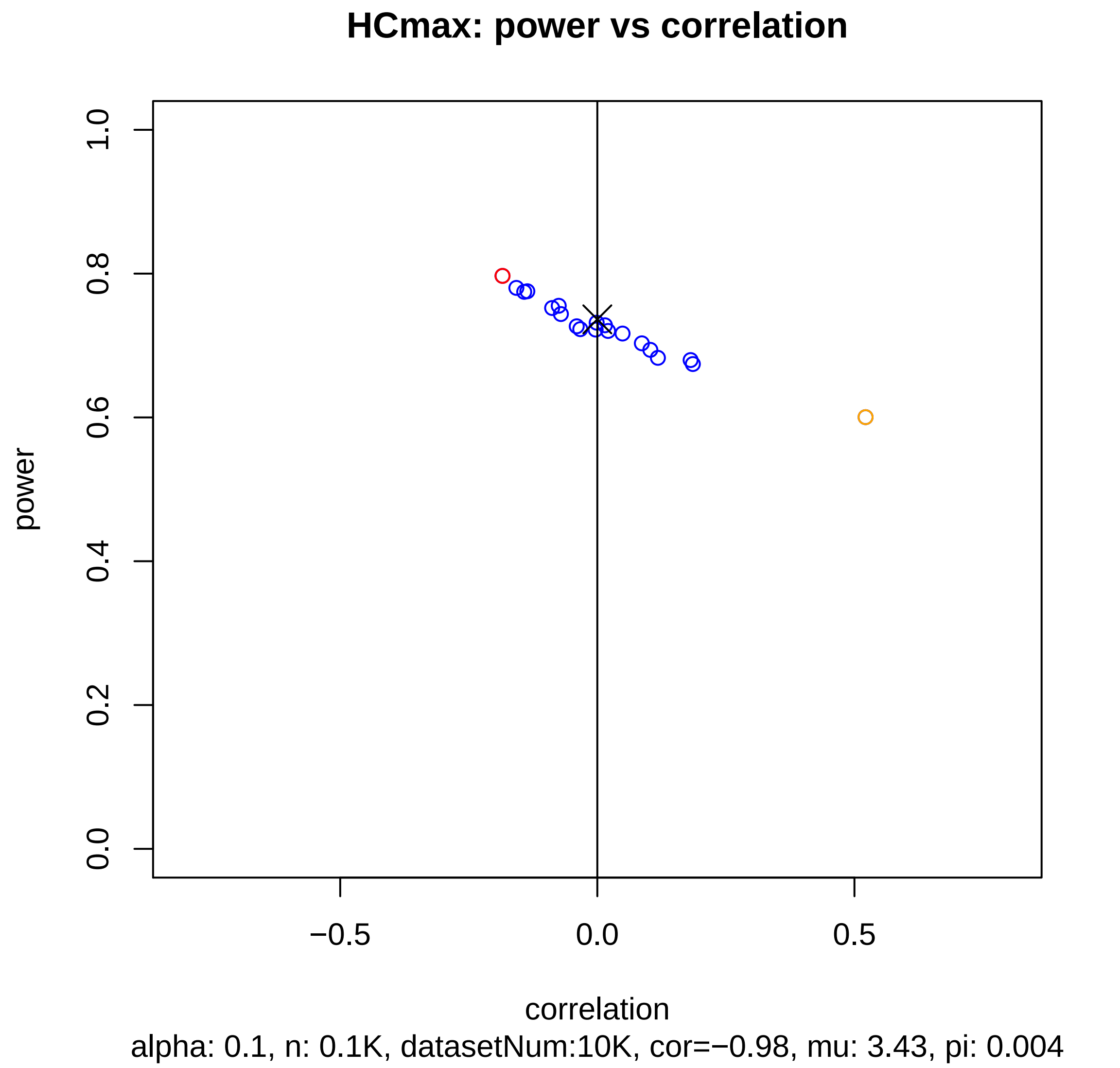
HCmax: power vs correlation
power
0.0
0.2
0.4
0.6
0.8
1.0
−0.5
0.0
0.5
correlation
alpha: 0.1, n: 0.1K, datasetNum:10K, cor=−0.98, mu: 3.43, pi: 0.004

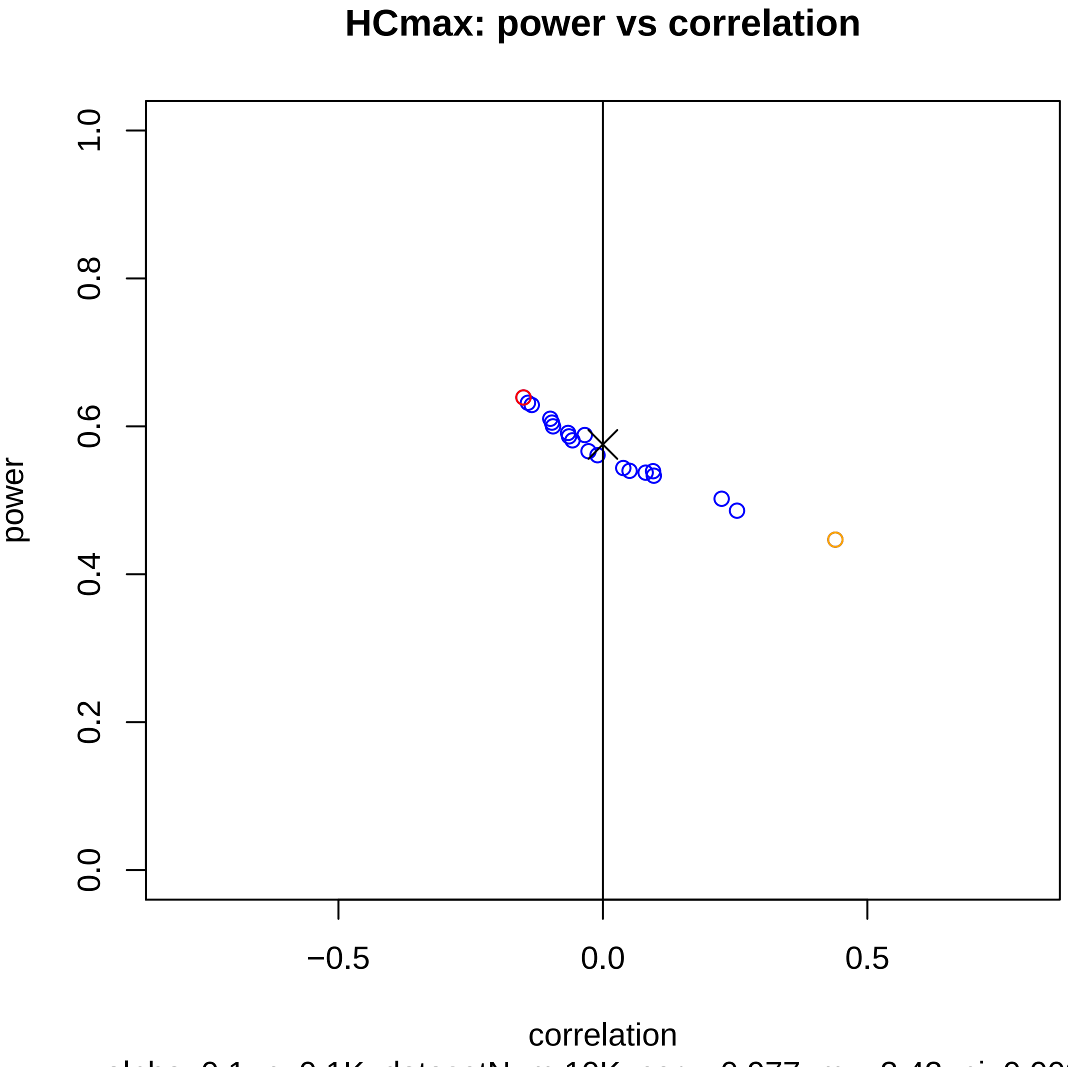

alpha: 0.1, n: 0.1K, datasetNum:10K, cor=−0.977, mu: 3.43, pi: 0.0028

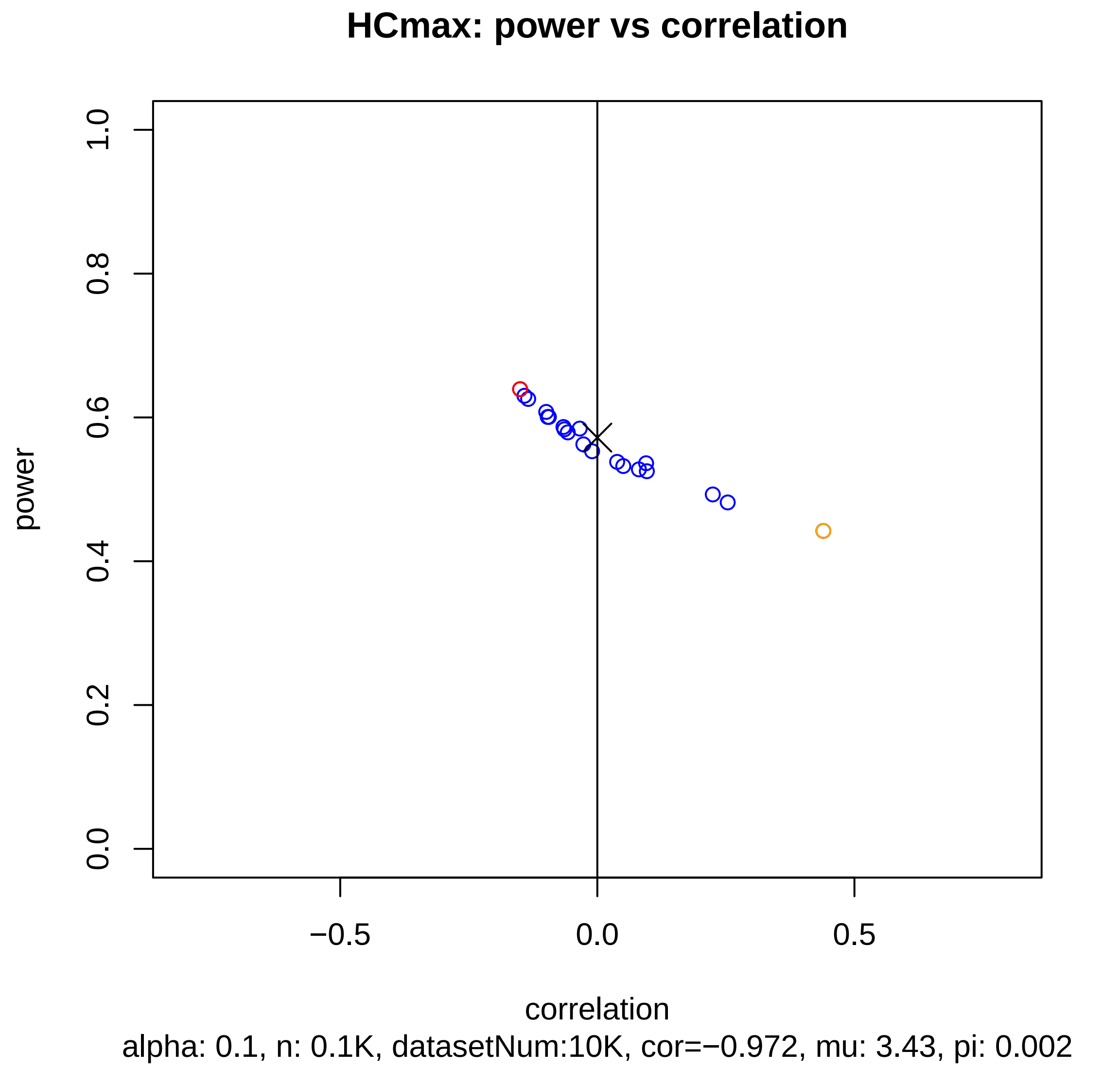
HCmax: power vs correlation
power
1.0
0.8
0.6
0.4
0.2
0.0
−0.5
0.0
0.5
correlation
alpha: 0.1, n: 0.1K, datasetNum:10K, cor=−0.972, mu: 3.43, pi: 0.002

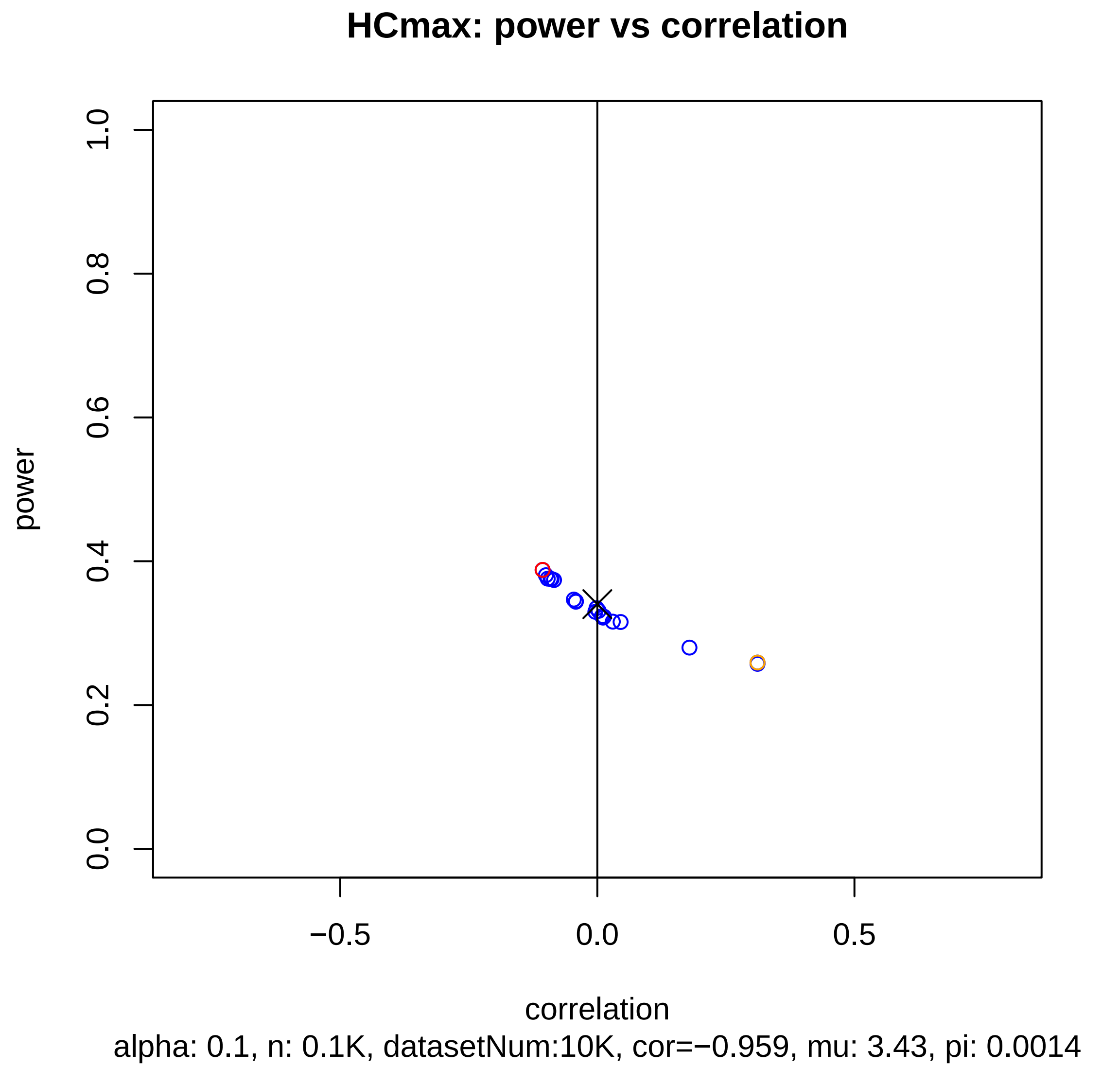
HCmax: power vs correlation
power
1.0
0.8
0.6
0.4
0.2
0.0
−0.5
0.0
0.5
correlation
alpha: 0.1, n: 0.1K, datasetNum:10K, cor=−0.959, mu: 3.43, pi: 0.0014

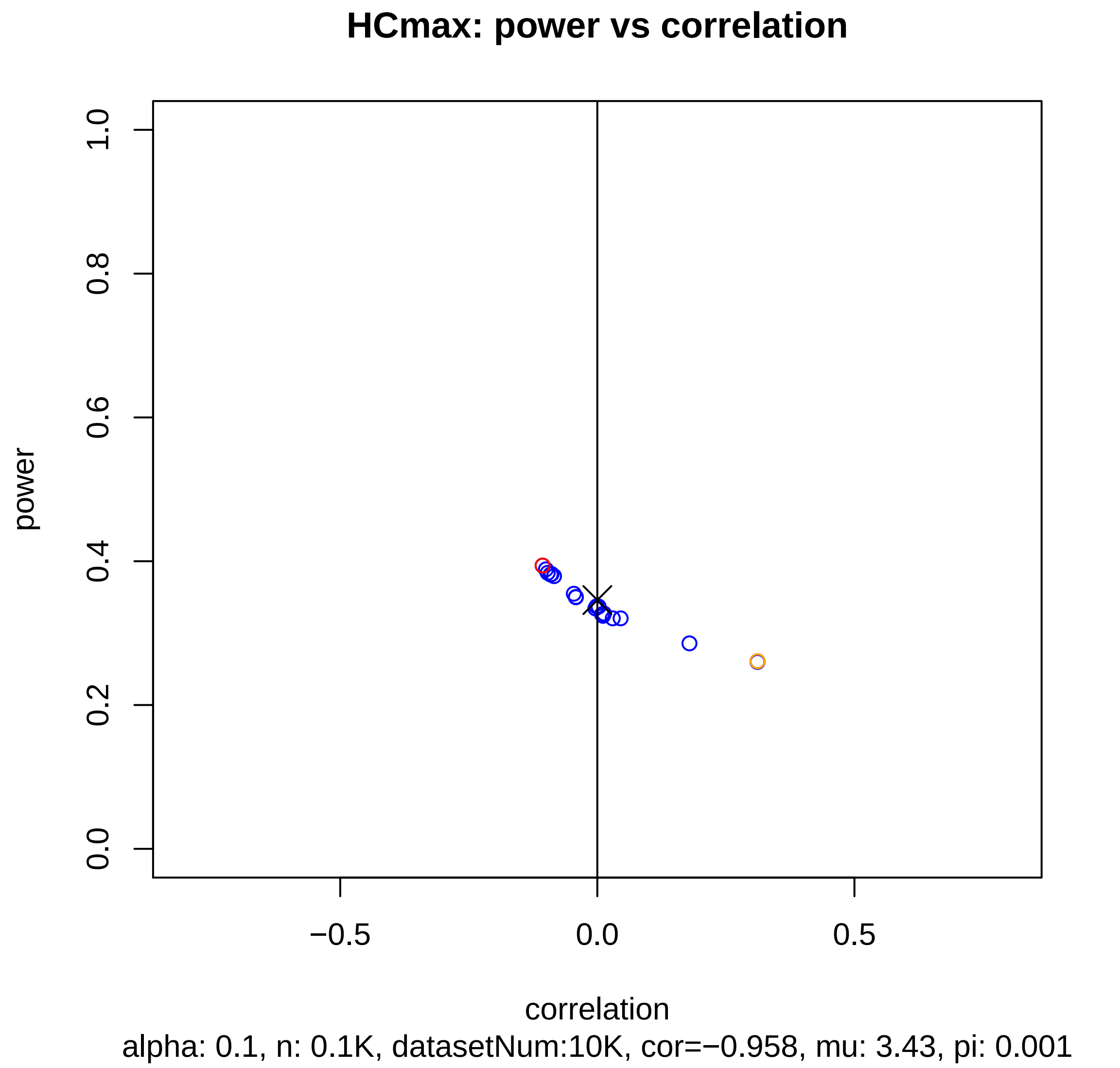
HCmax: power vs correlation
power
1.0
0.8
0.6
0.4
0.2
0.0
−0.5
0.0
0.5
correlation
alpha: 0.1, n: 0.1K, datasetNum:10K, cor=−0.958, mu: 3.43, pi: 0.001

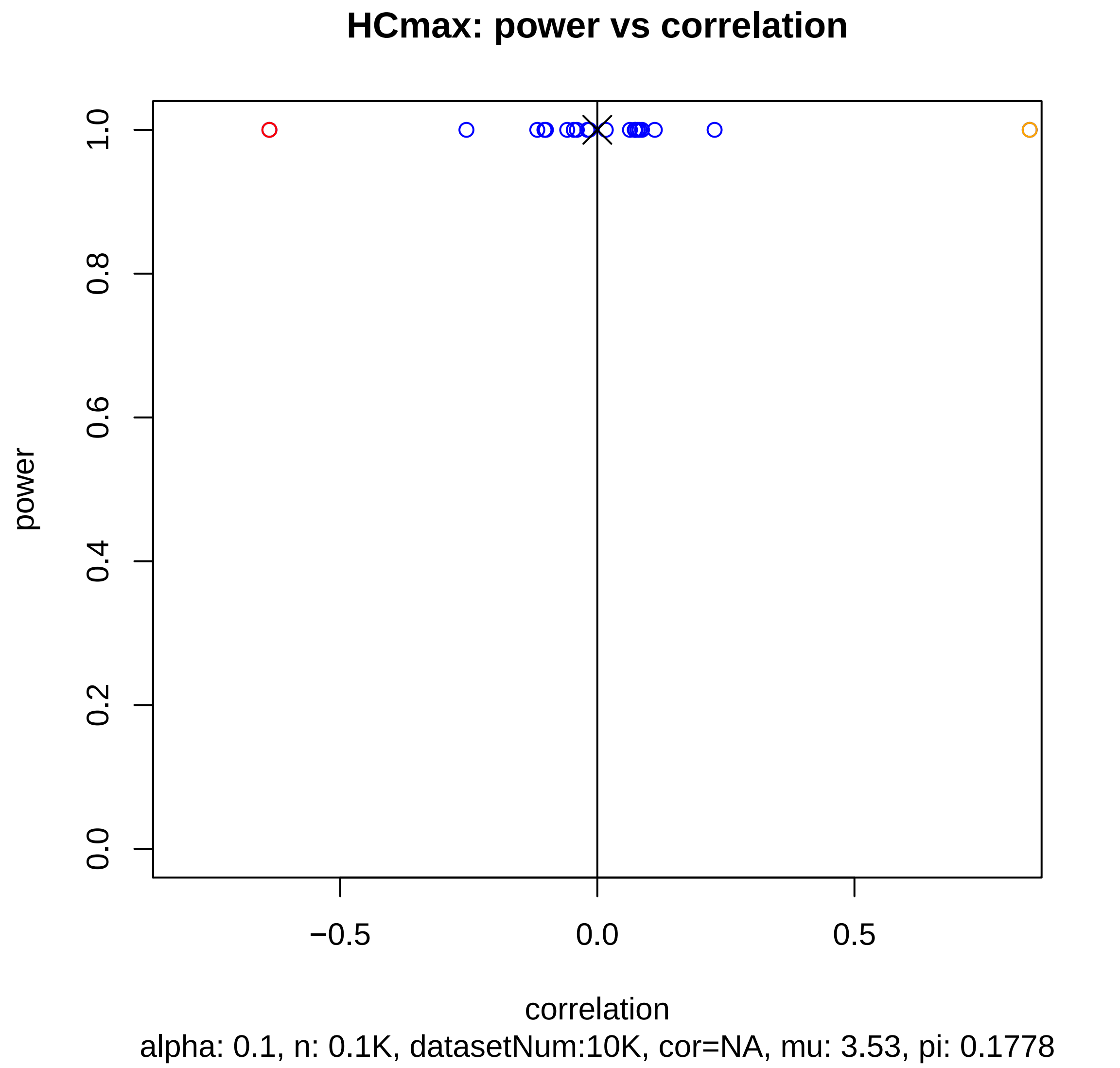
HCmax: power vs correlation
power
1.0
0.8
0.6
0.4
0.2
0.0
−0.5
0.0
0.5
correlation
alpha: 0.1, n: 0.1K, datasetNum:10K, cor=NA, mu: 3.53, pi: 0.1778

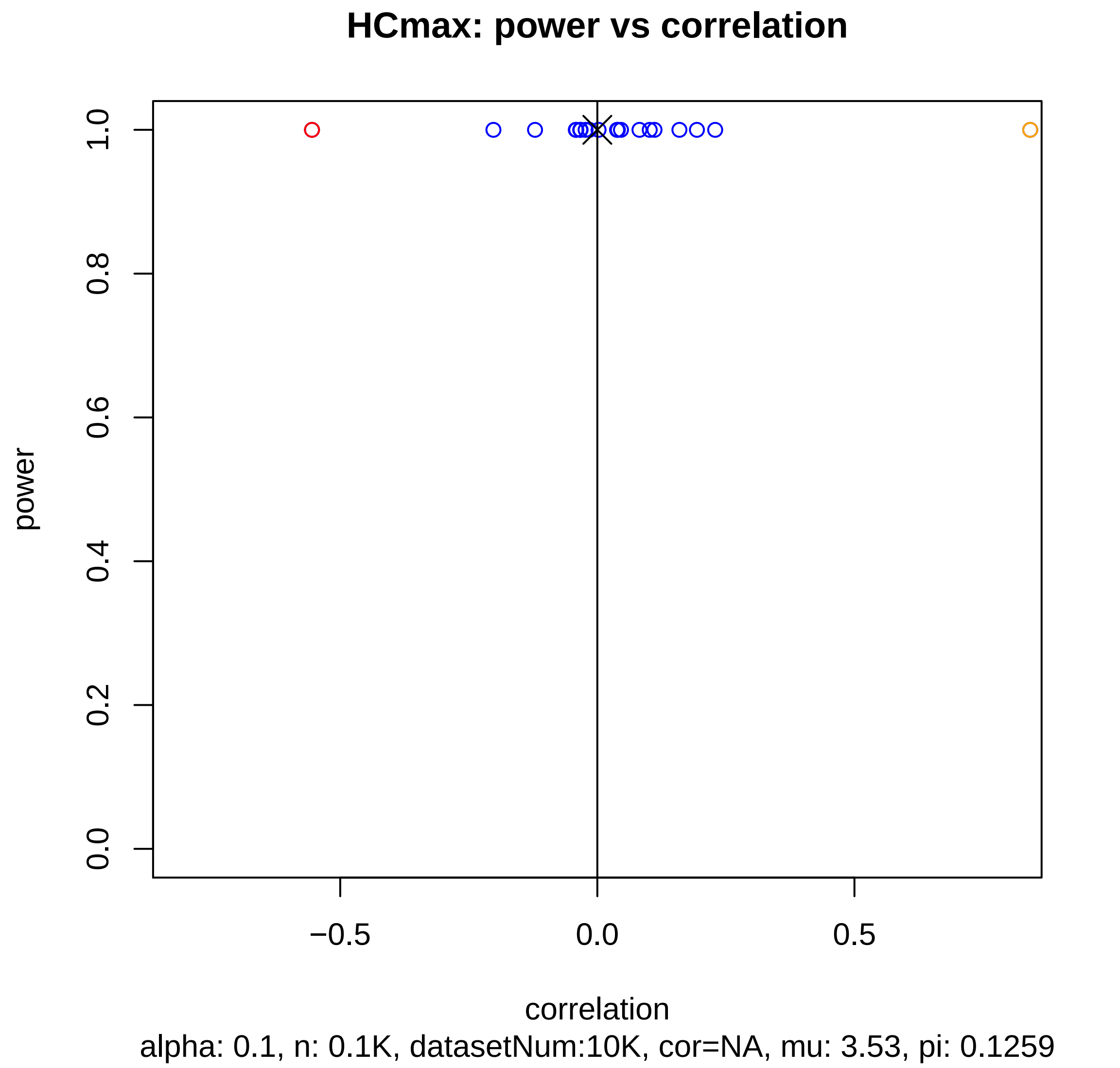
HCmax: power vs correlation
power
1.0
0.8
0.6
0.4
0.2
0.0
−0.5
0.0
0.5
correlation
alpha: 0.1, n: 0.1K, datasetNum:10K, cor=NA, mu: 3.53, pi: 0.1259

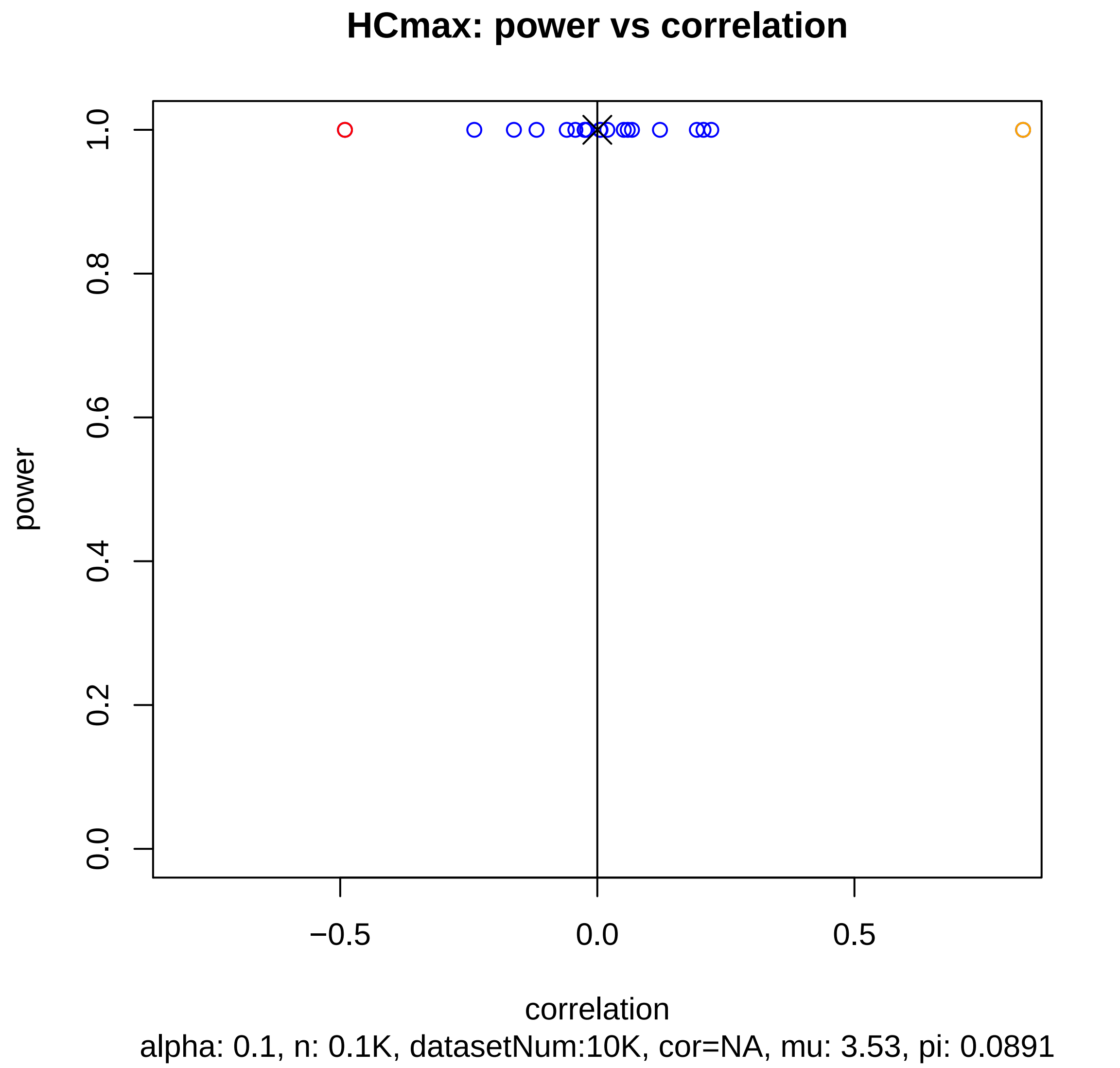

HCmax: power vs correlation
power
1.0
0.8
0.6
0.4
0.2
0.0
−0.5
0.0
0.5
correlation
alpha: 0.1, n: 0.1K, datasetNum:10K, cor=NA, mu: 3.53, pi: 0.0891

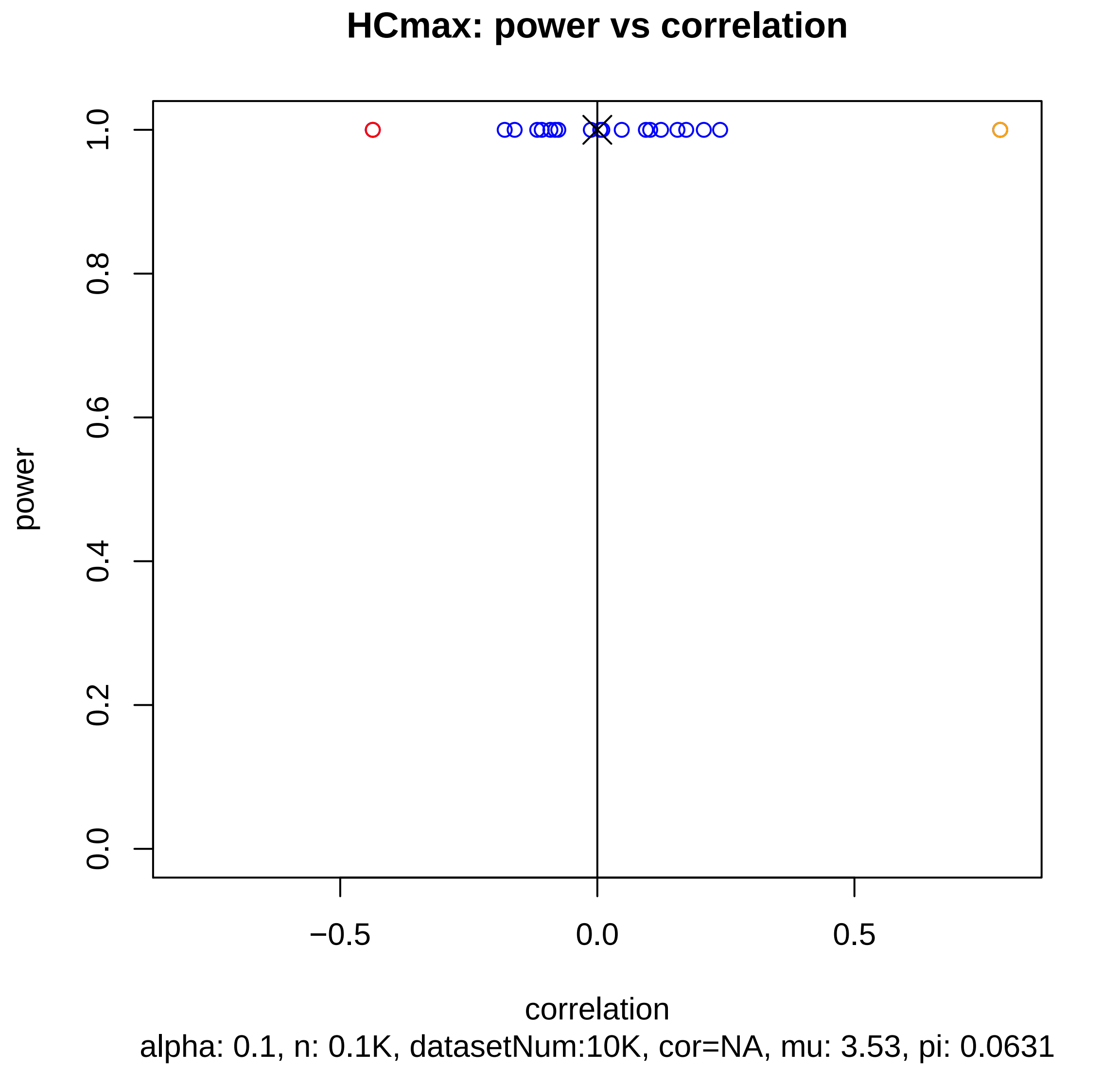
HCmax: power vs correlation
power
1.0
0.8
0.6
0.4
0.2
0.0
−0.5
0.0
0.5
correlation
alpha: 0.1, n: 0.1K, datasetNum:10K, cor=NA, mu: 3.53, pi: 0.0631

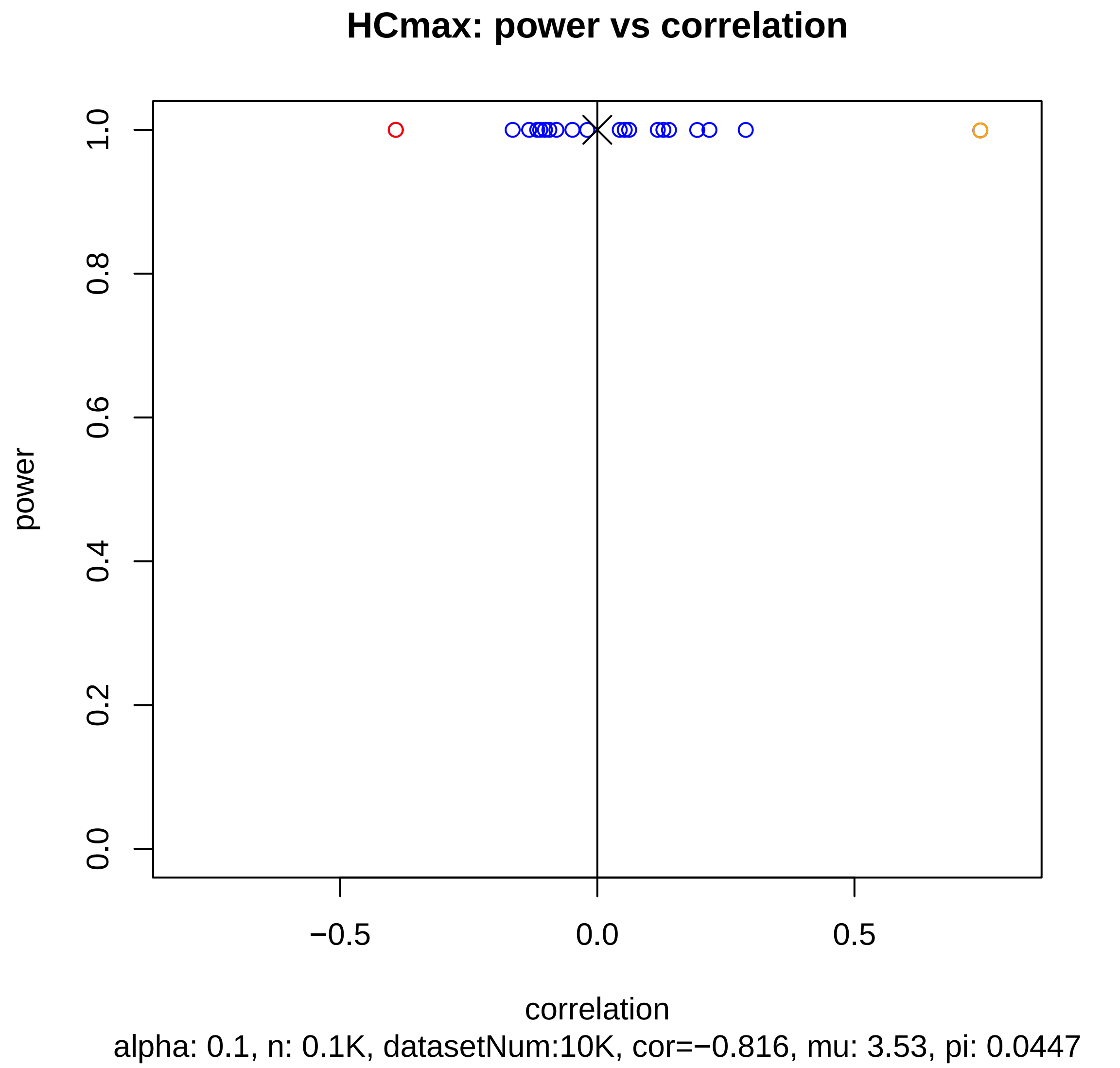
HCmax: power vs correlation
power
1.0
0.8
0.6
0.4
0.2
0.0
−0.5
0.0
0.5
correlation
alpha: 0.1, n: 0.1K, datasetNum:10K, cor=−0.816, mu: 3.53, pi: 0.0447

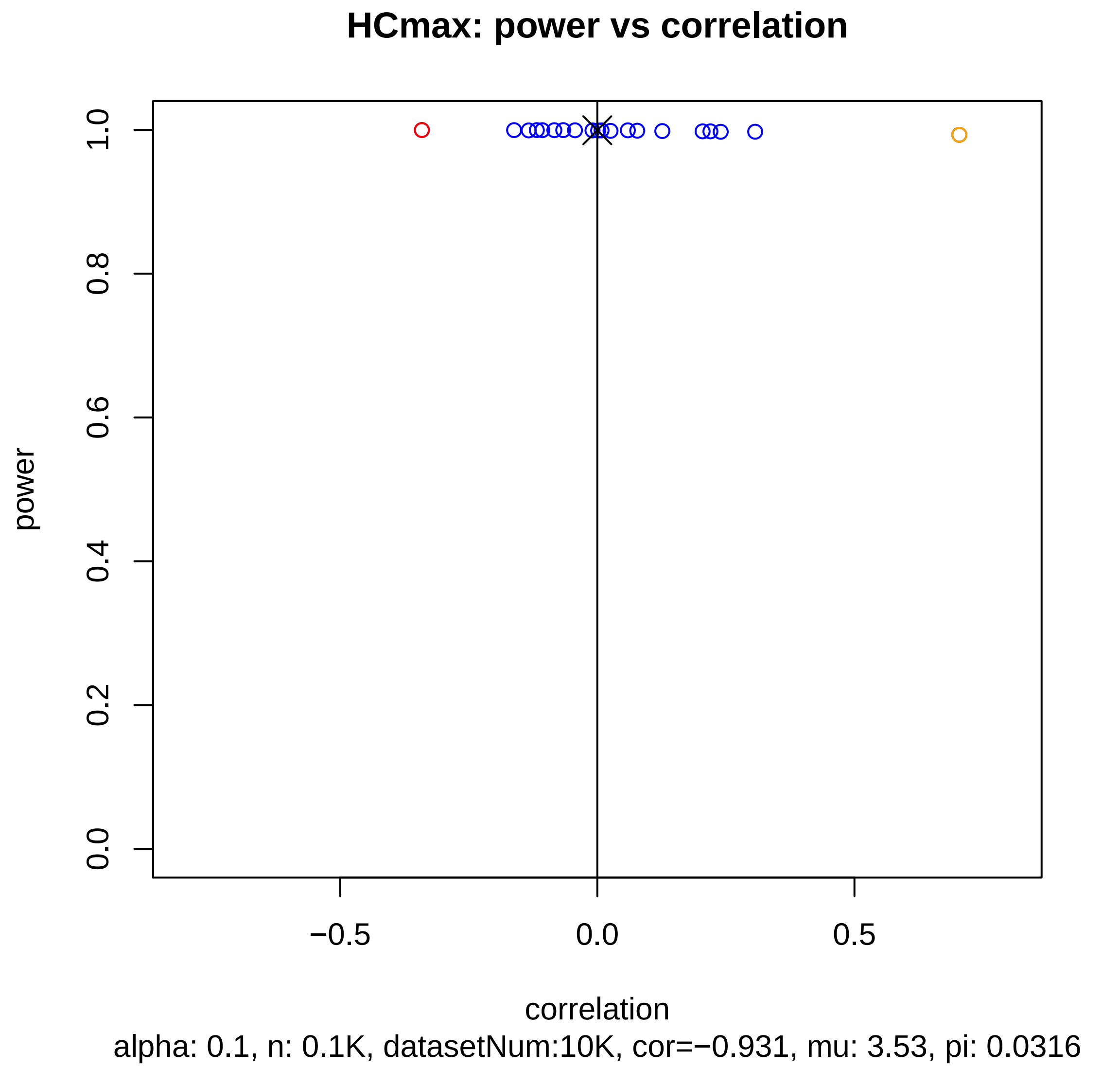
HCmax: power vs correlation
power
1.0
0.8
0.6
0.4
0.2
0.0
−0.5
0.0
0.5
correlation
alpha: 0.1, n: 0.1K, datasetNum:10K, cor=−0.931, mu: 3.53, pi: 0.0316

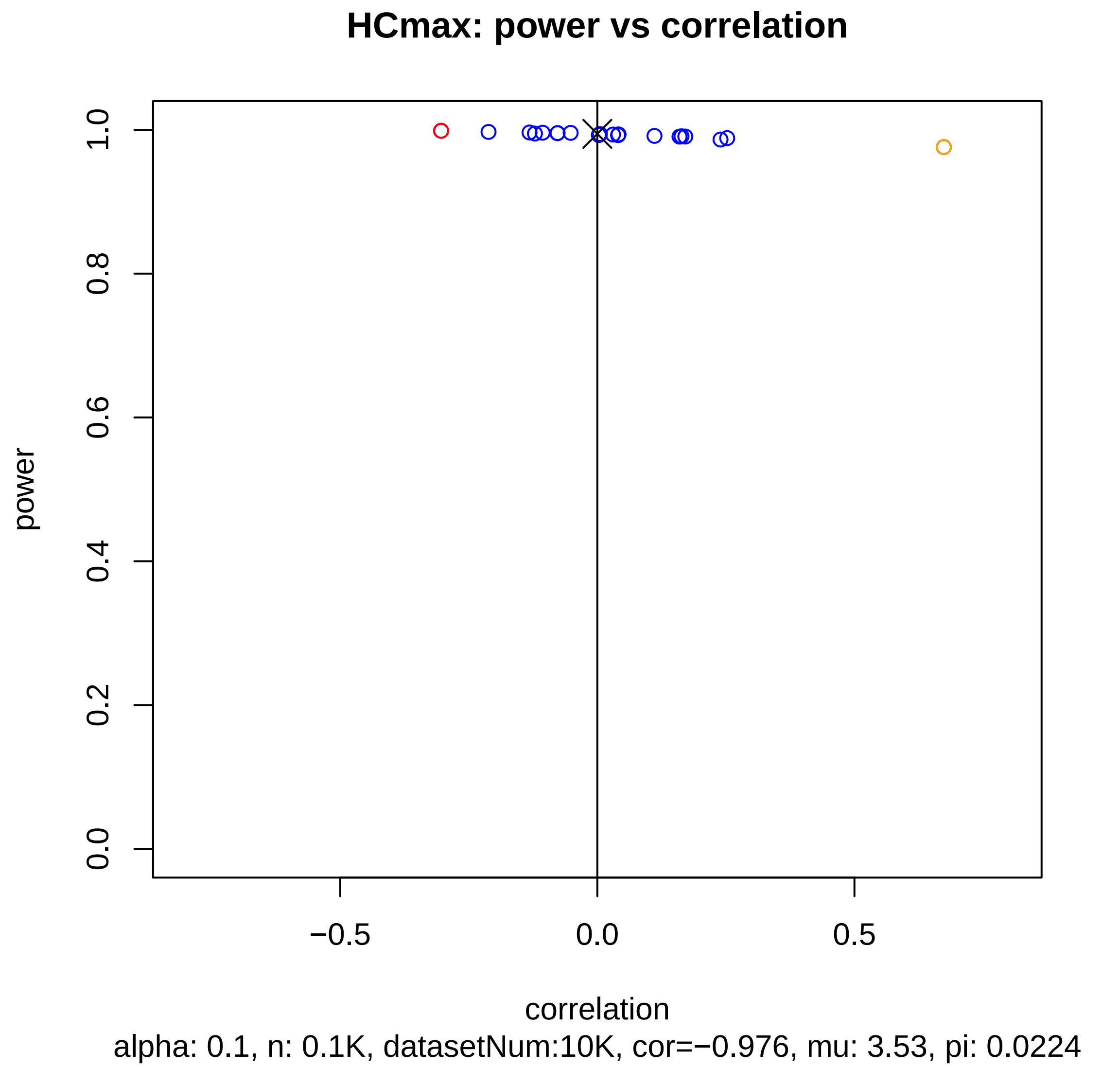

HCmax: power vs correlation
power
1.0
0.8
0.6
0.4
0.2
0.0
−0.5
0.0
0.5
correlation
alpha: 0.1, n: 0.1K, datasetNum:10K, cor=−0.976, mu: 3.53, pi: 0.0224

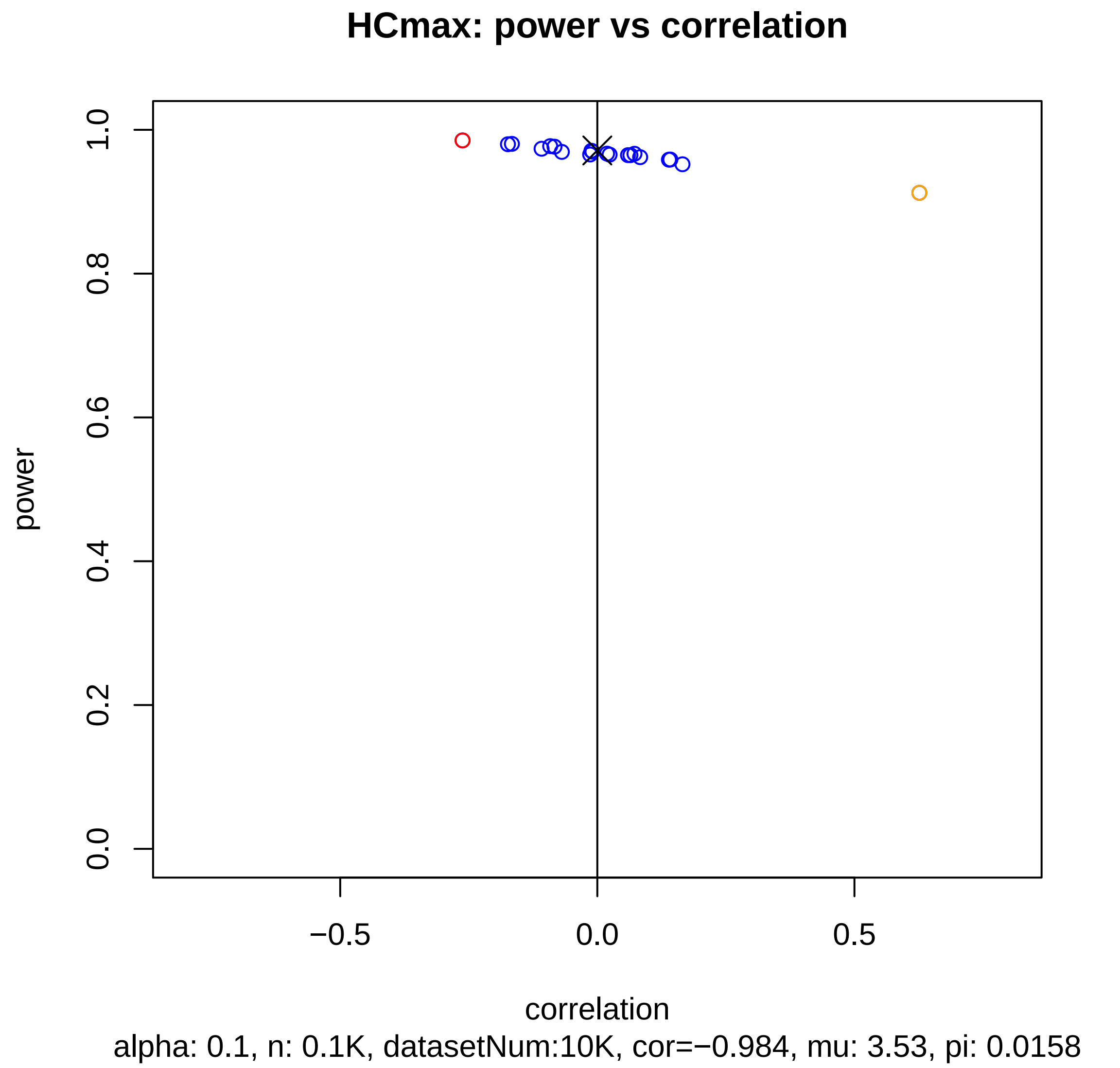
HCmax: power vs correlation
power
1.0
0.8
0.6
0.4
0.2
0.0
−0.5
0.0
0.5
correlation
alpha: 0.1, n: 0.1K, datasetNum:10K, cor=−0.984, mu: 3.53, pi: 0.0158

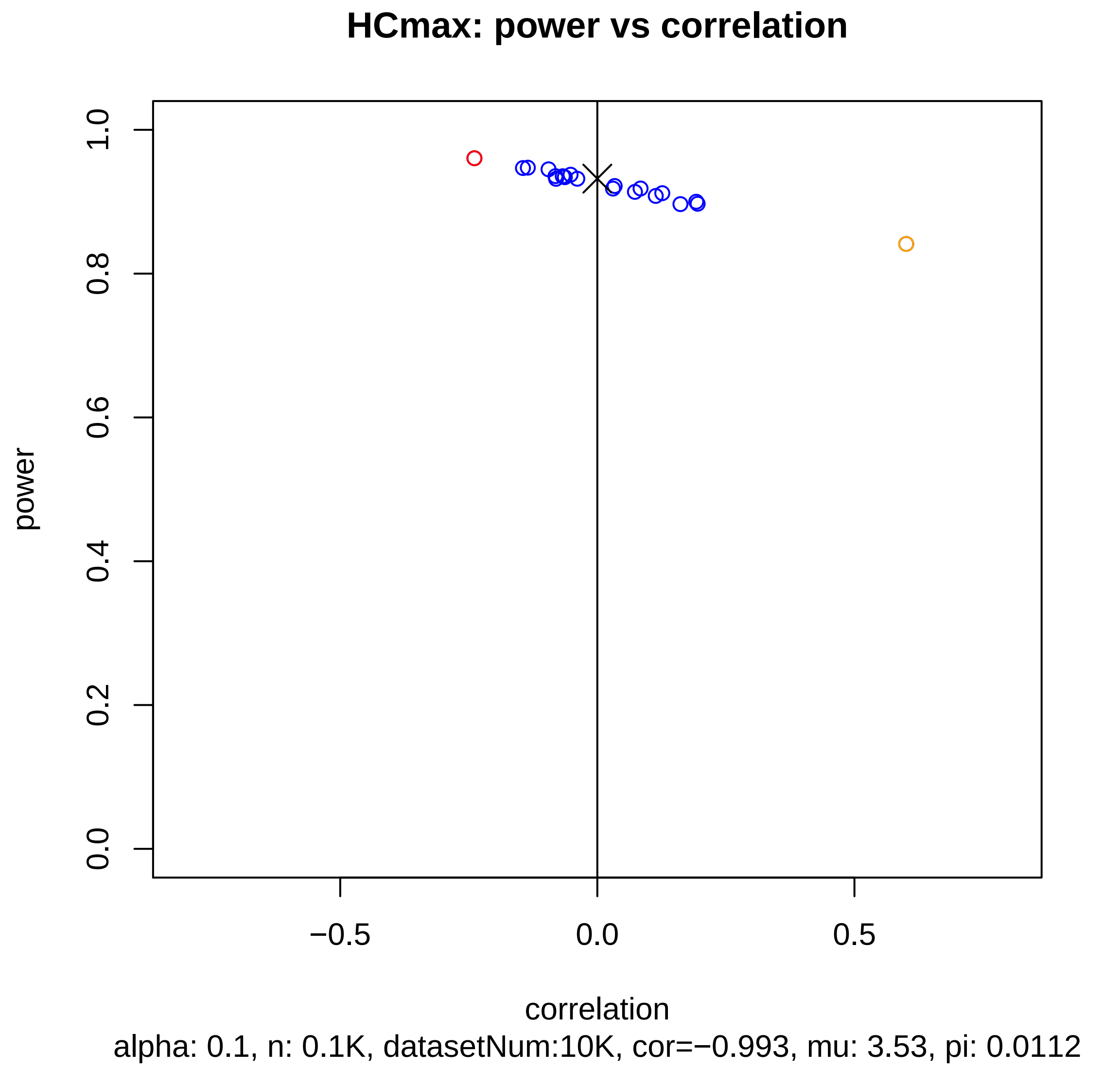

HCmax: power vs correlation
power
0.0
0.2
0.4
0.6
0.8
1.0
−0.5
0.0
0.5
correlation
alpha: 0.1, n: 0.1K, datasetNum:10K, cor=−0.993, mu: 3.53, pi: 0.0112

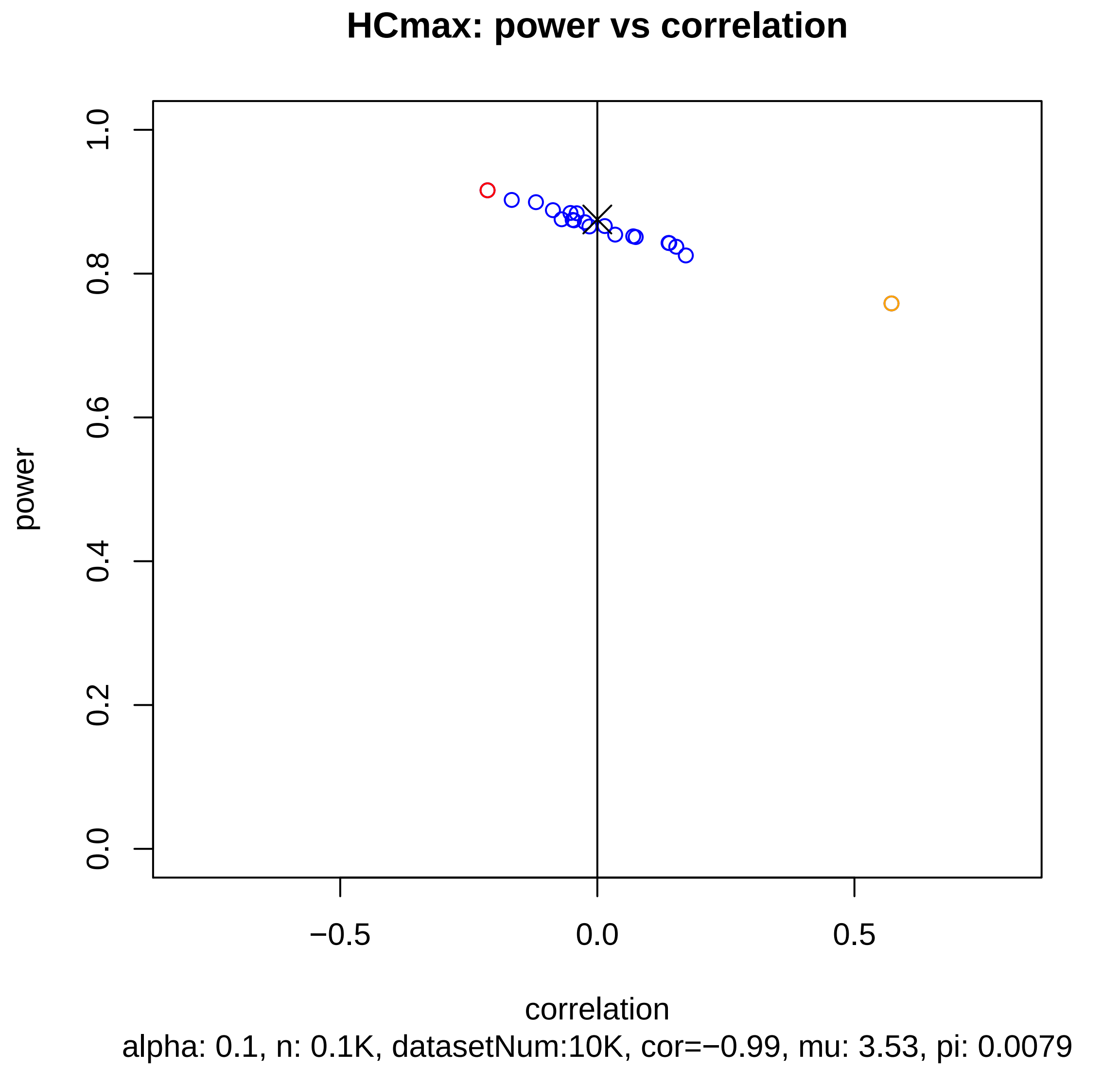
HCmax: power vs correlation
power
0.0
0.2
0.4
0.6
0.8
1.0
−0.5
0.0
0.5
correlation
alpha: 0.1, n: 0.1K, datasetNum:10K, cor=−0.99, mu: 3.53, pi: 0.0079

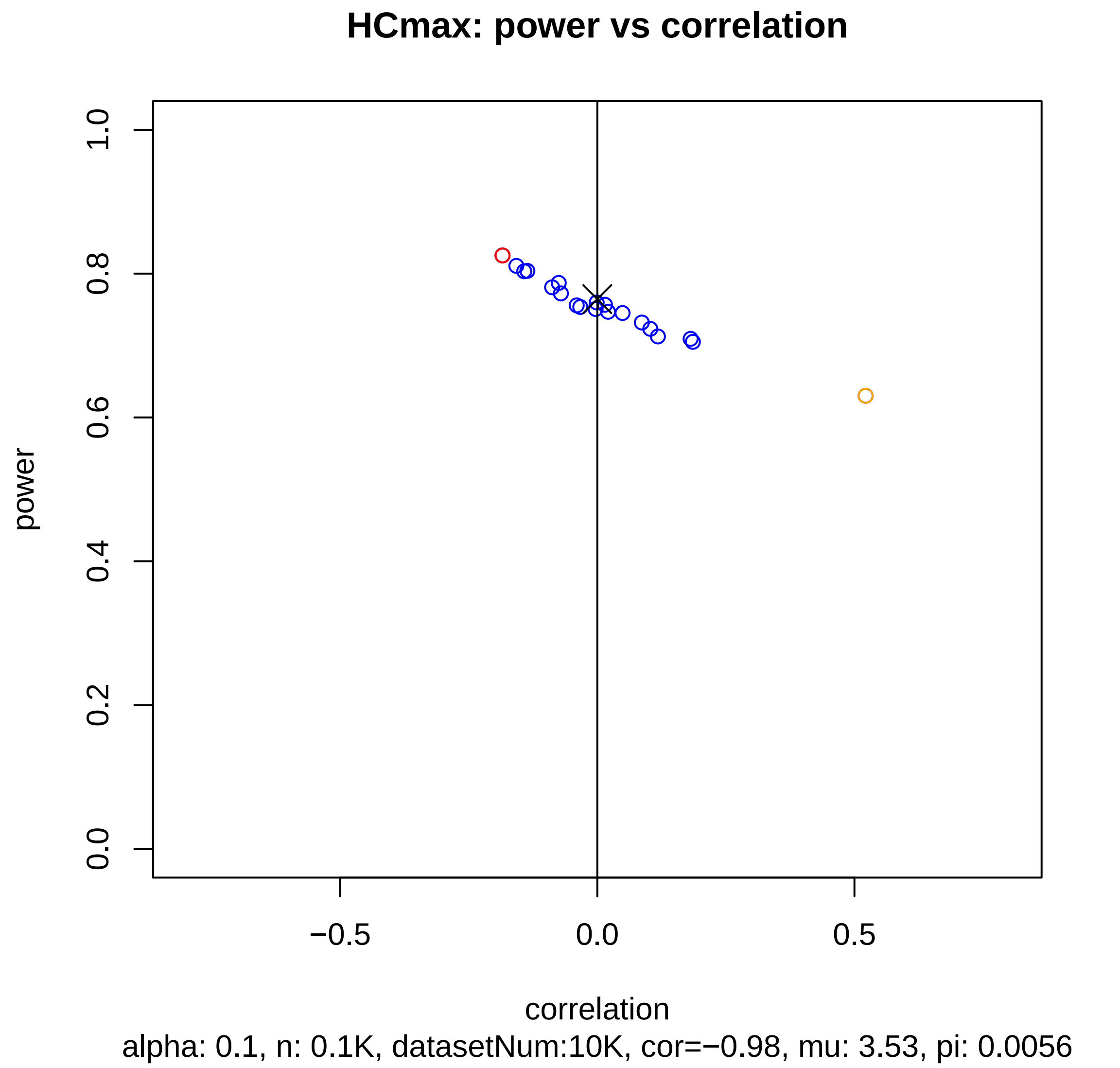

HCmax: power vs correlation
power
1.0
0.8
0.6
0.4
0.2
0.0
−0.5
0.0
0.5
correlation
alpha: 0.1, n: 0.1K, datasetNum:10K, cor=−0.98, mu: 3.53, pi: 0.0056

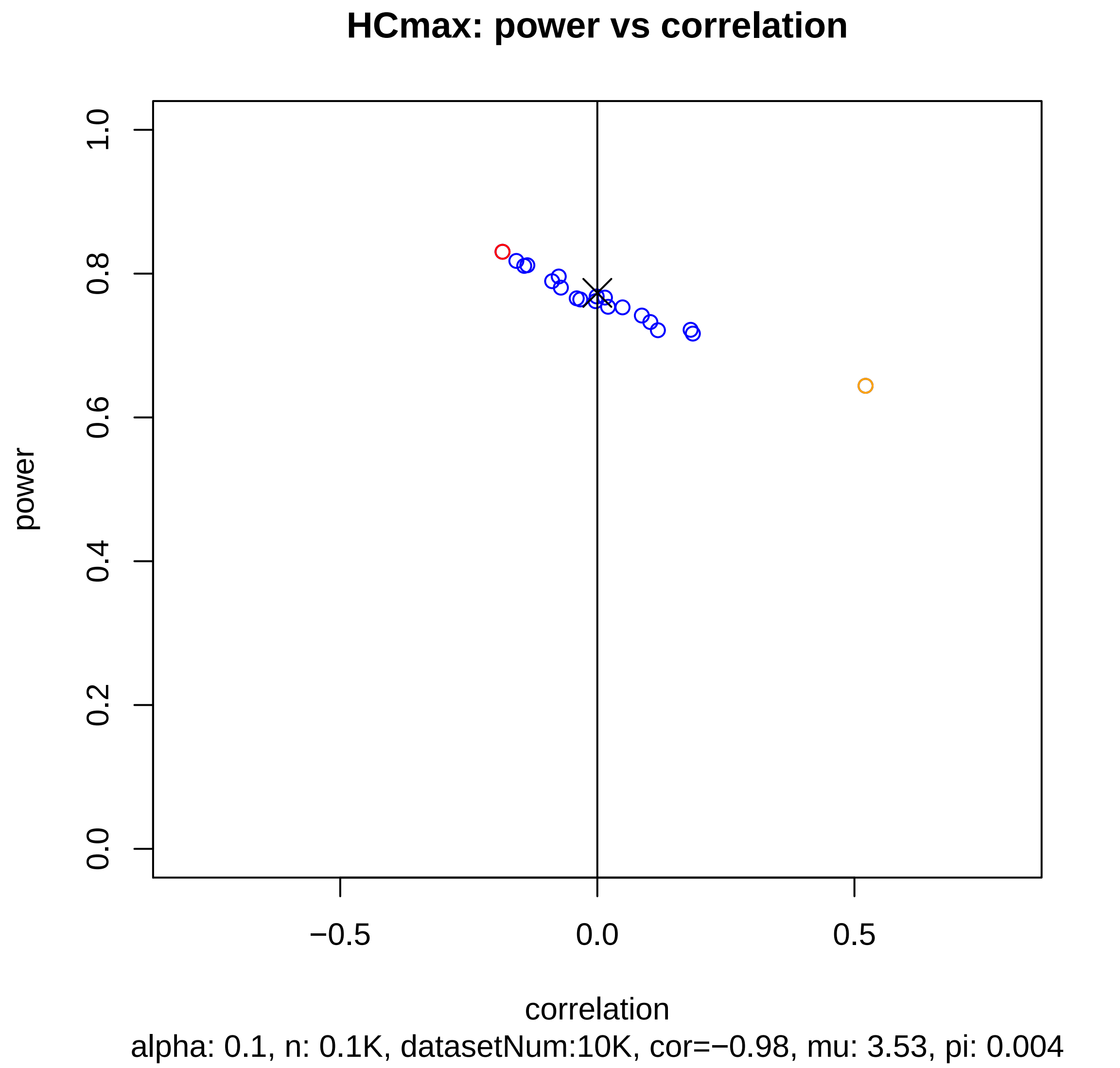
HCmax: power vs correlation
power
1.0
0.8
0.6
0.4
0.2
0.0
−0.5
0.0
0.5
correlation
alpha: 0.1, n: 0.1K, datasetNum:10K, cor=−0.98, mu: 3.53, pi: 0.004

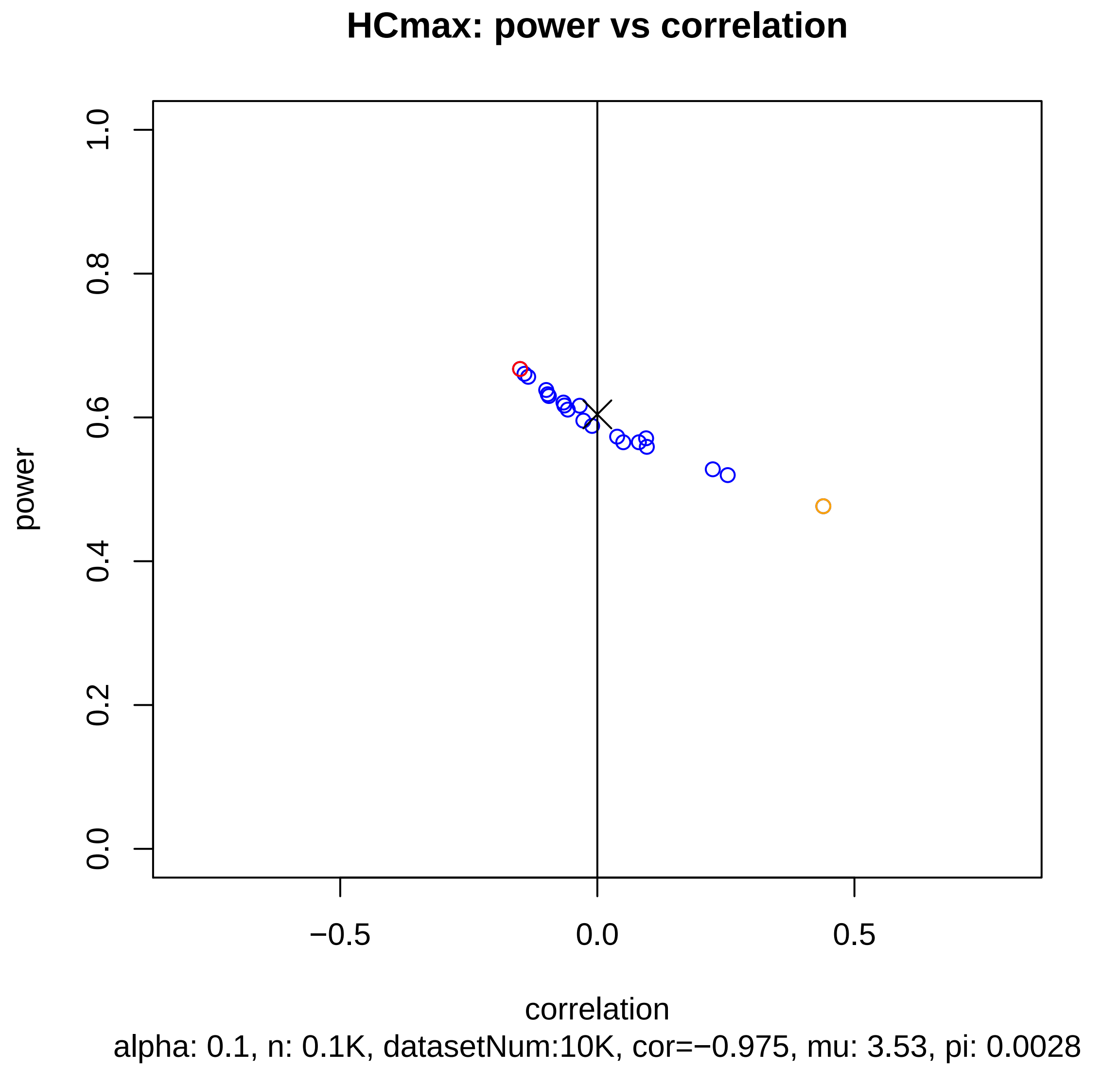
HCmax: power vs correlation
power
1.0
0.8
0.6
0.4
0.2
0.0
−0.5
0.0
0.5
correlation
alpha: 0.1, n: 0.1K, datasetNum:10K, cor=−0.975, mu: 3.53, pi: 0.0028

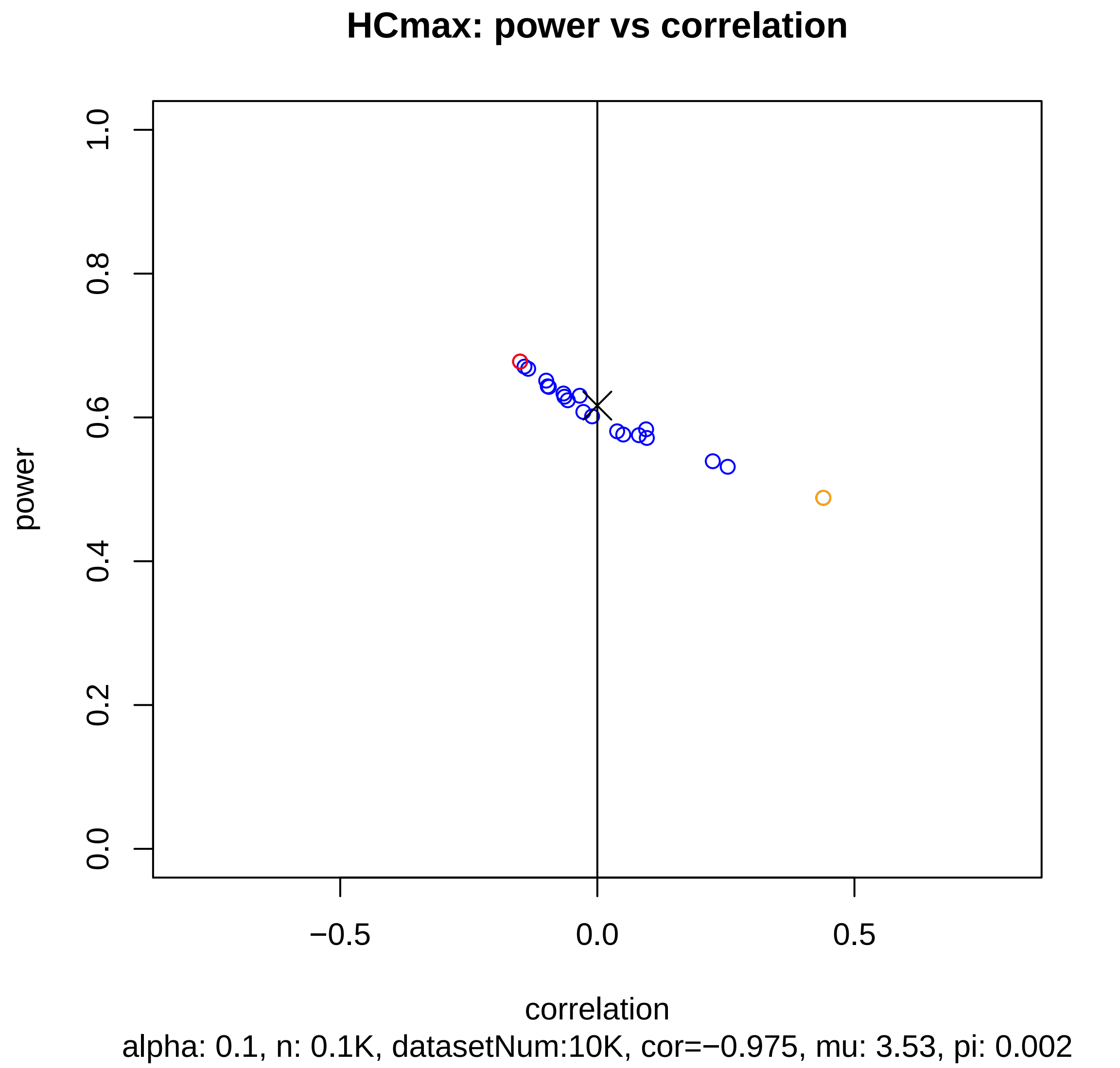
HCmax: power vs correlation
1.0
0.8
0.6
0.4
0.2
0.0
power
−0.5
0.0
0.5
correlation
alpha: 0.1, n: 0.1K, datasetNum:10K, cor=−0.975, mu: 3.53, pi: 0.002

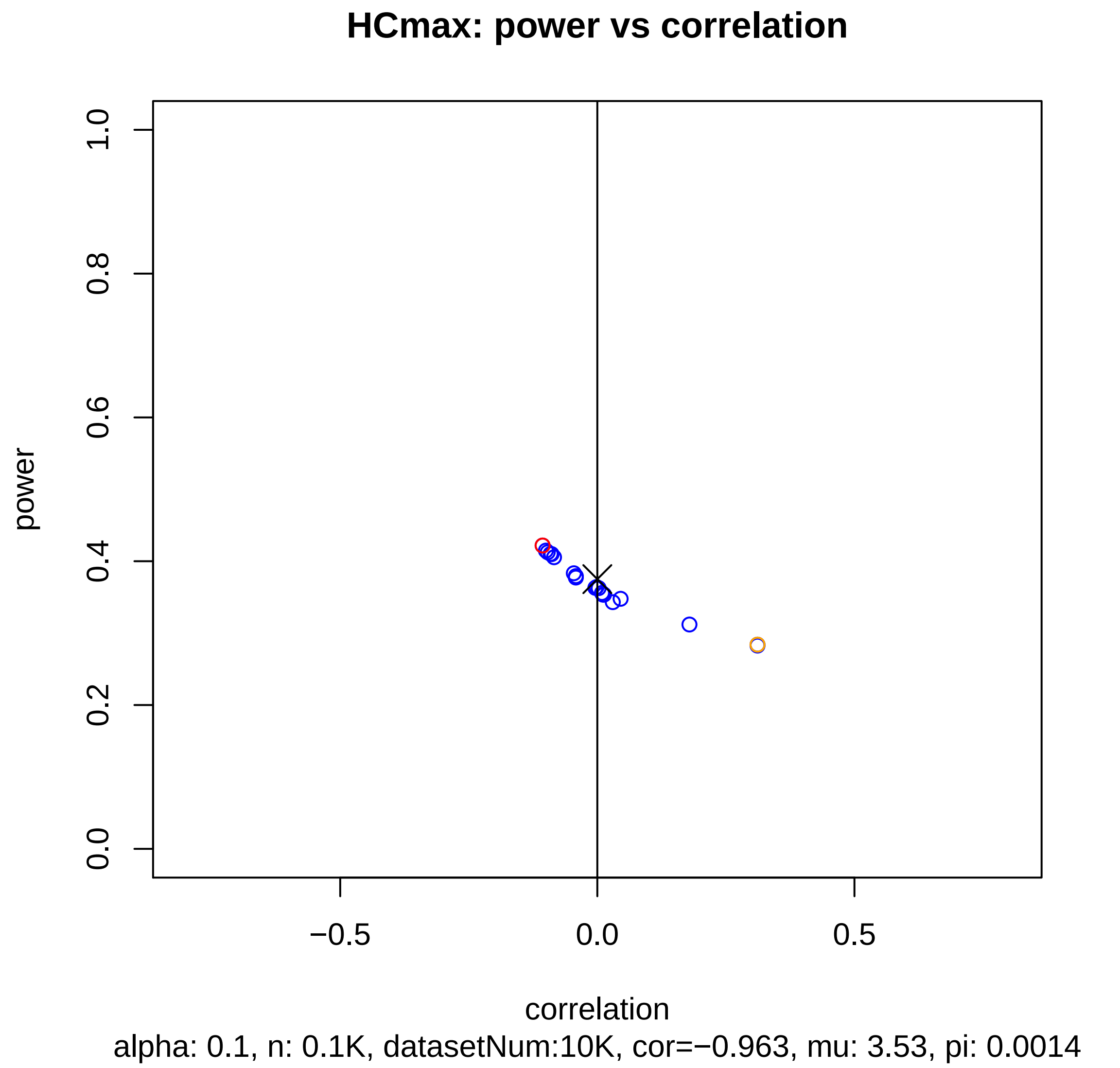
HCmax: power vs correlation
power
1.0
0.8
0.6
0.4
0.2
0.0
−0.5
0.0
0.5
correlation
alpha: 0.1, n: 0.1K, datasetNum:10K, cor=−0.963, mu: 3.53, pi: 0.0014

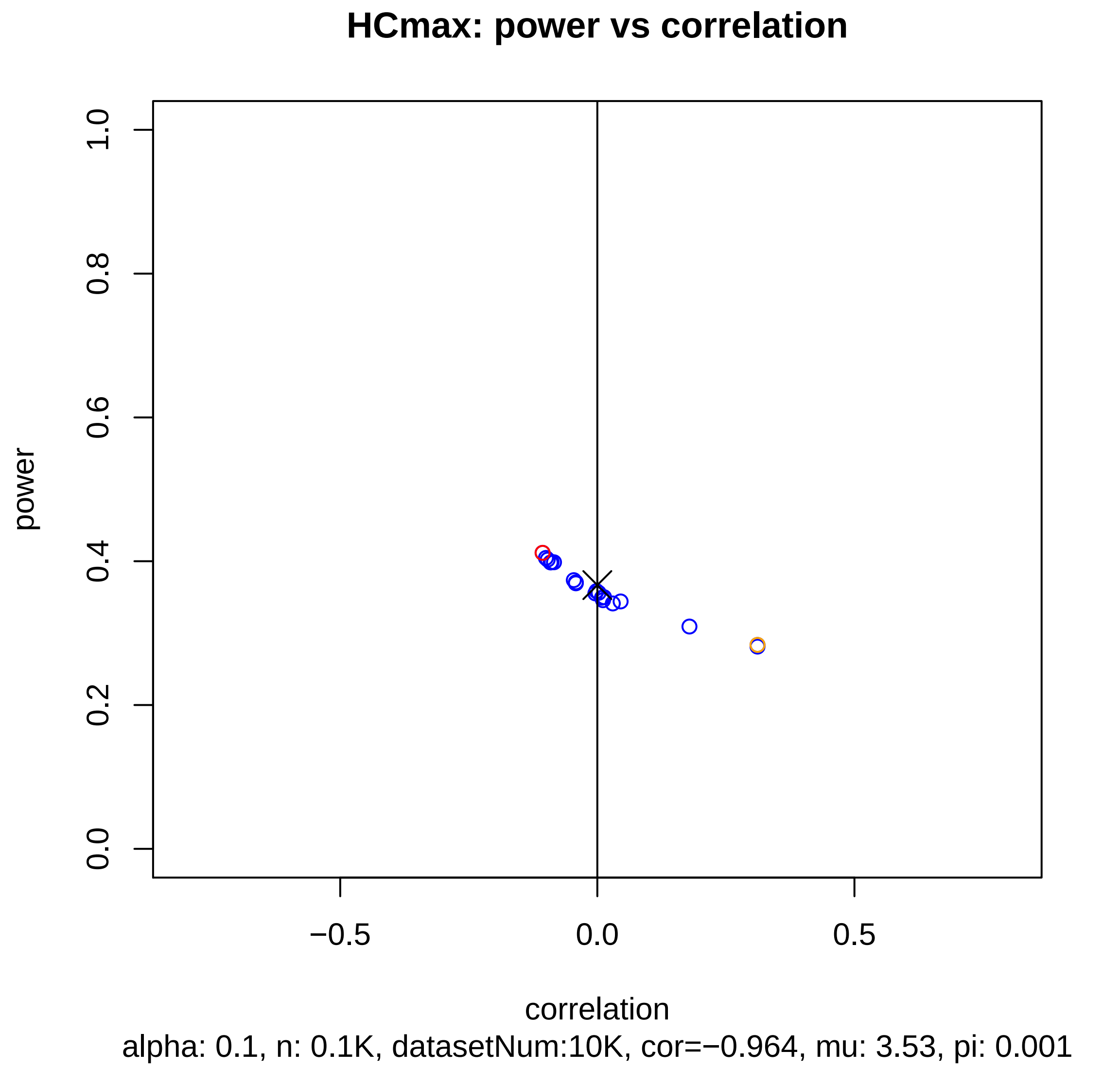
HCmax: power vs correlation
power
1.0
0.8
0.6
0.4
0.2
0.0
−0.5
0.0
0.5
correlation
alpha: 0.1, n: 0.1K, datasetNum:10K, cor=−0.964, mu: 3.53, pi: 0.001

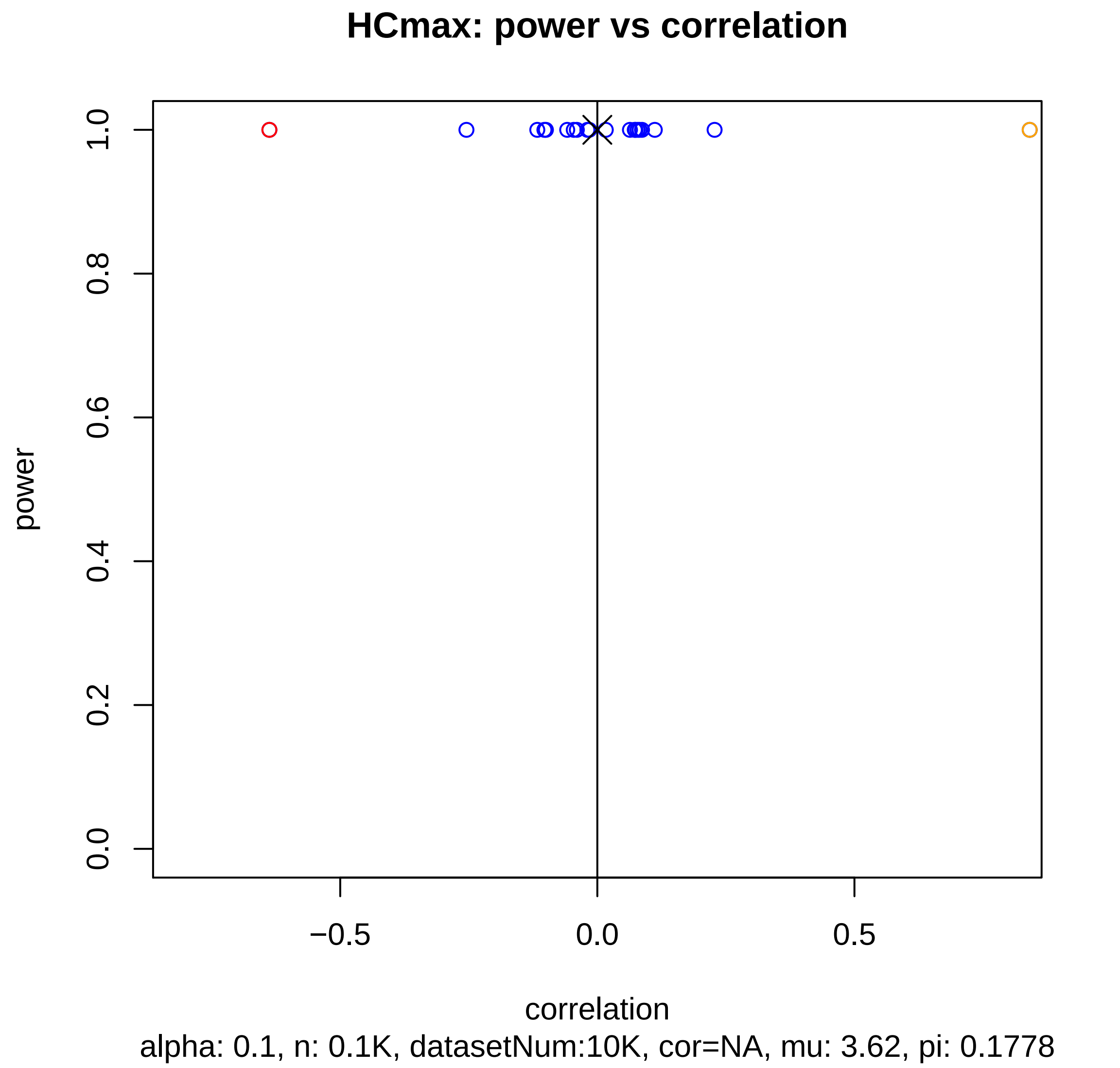
HCmax: power vs correlation
power
1.0
0.8
0.6
0.4
0.2
0.0
−0.5
0.0
0.5
correlation
alpha: 0.1, n: 0.1K, datasetNum:10K, cor=NA, mu: 3.62, pi: 0.1778

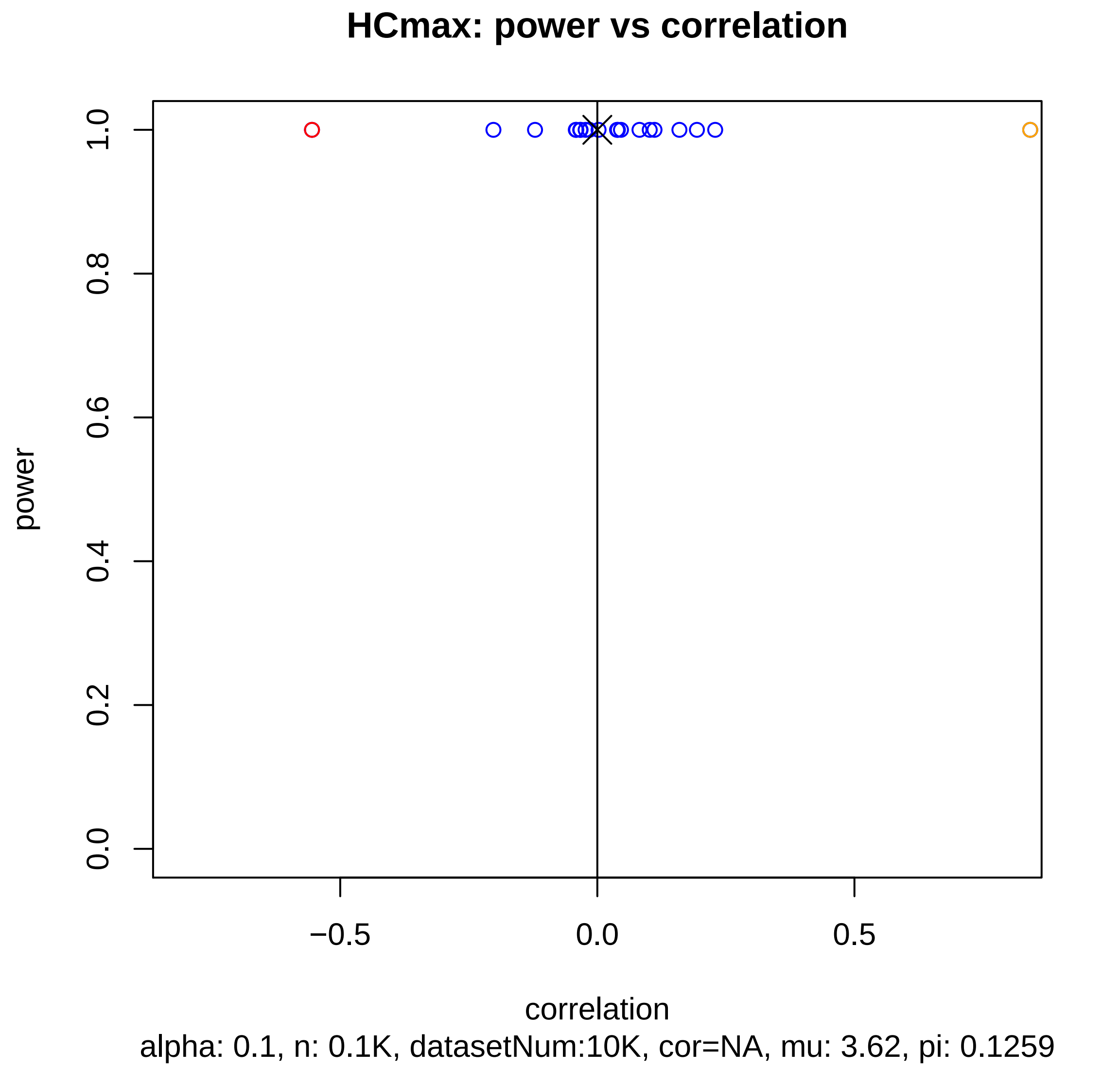
HCmax: power vs correlation
power
0.0
0.2
0.4
0.6
0.8
1.0
−0.5
0.0
0.5
correlation
alpha: 0.1, n: 0.1K, datasetNum:10K, cor=NA, mu: 3.62, pi: 0.1259

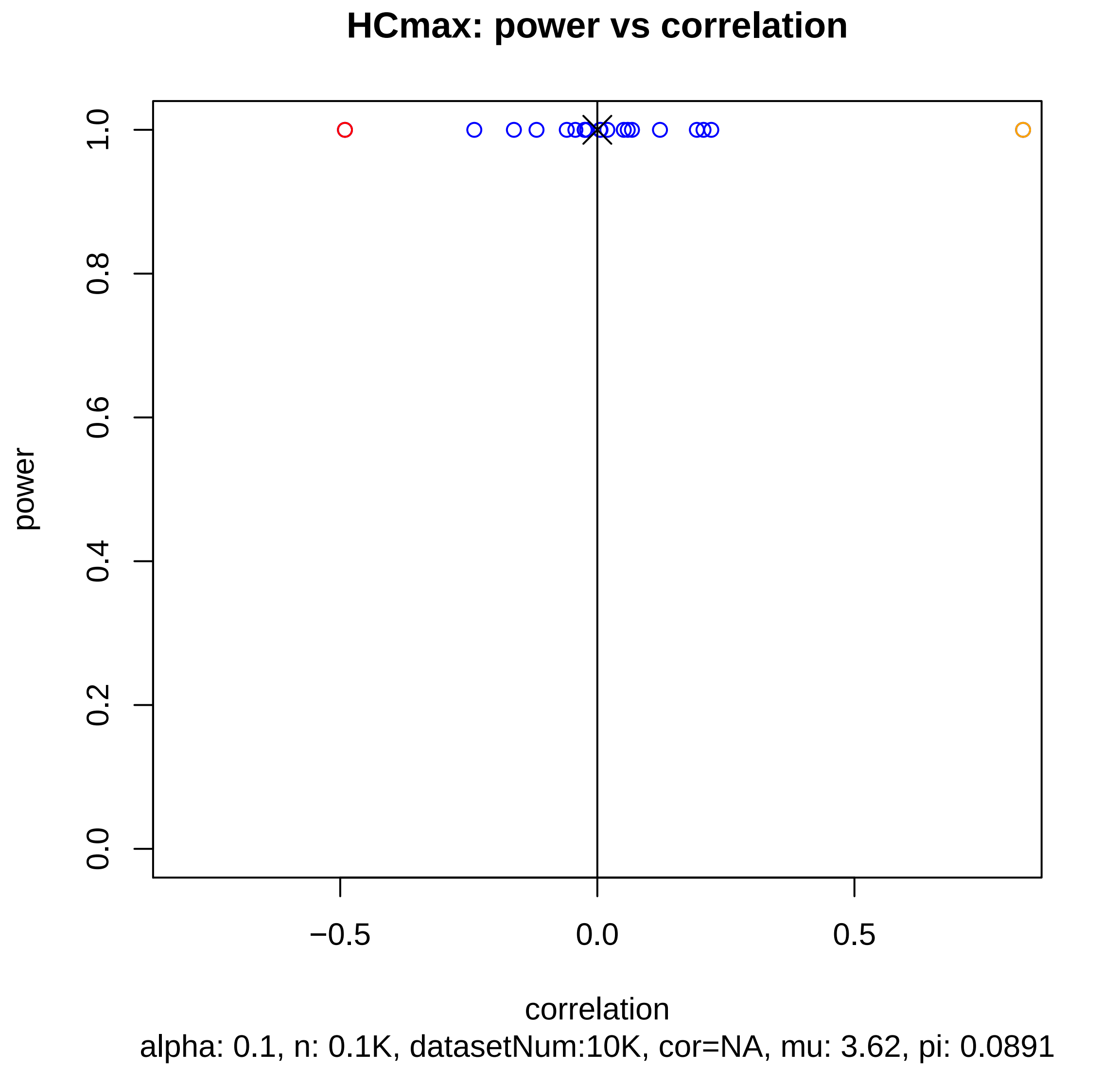
HCmax: power vs correlation
power
correlation
0.0
0.2
0.4
0.6
0.8
1.0
−0.5
0.0
0.5
alpha: 0.1, n: 0.1K, datasetNum:10K, cor=NA, mu: 3.62, pi: 0.0891

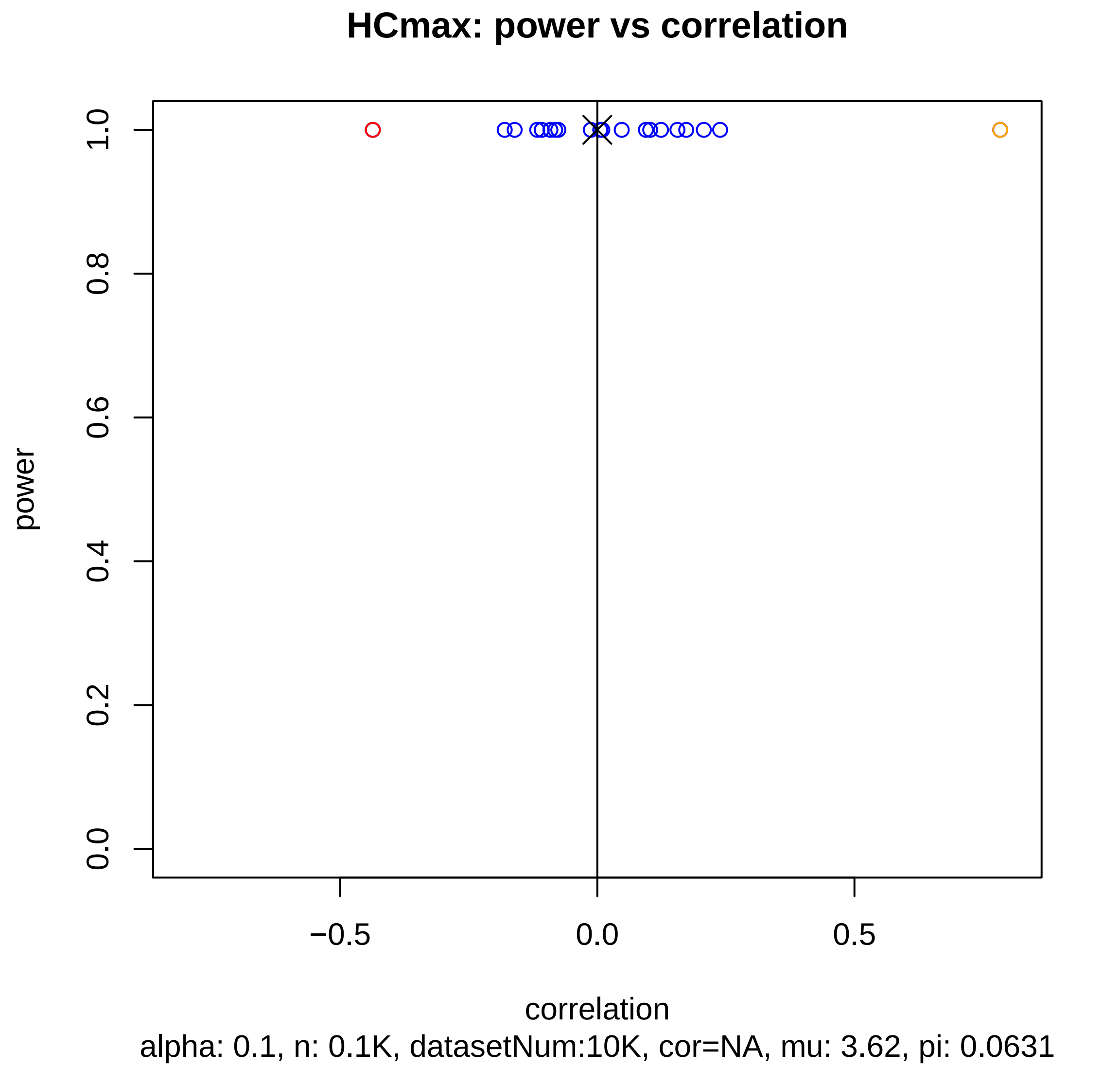
HCmax: power vs correlation
power
1.0
0.8
0.6
0.4
0.2
0.0
−0.5
0.0
0.5
correlation
alpha: 0.1, n: 0.1K, datasetNum:10K, cor=NA, mu: 3.62, pi: 0.0631

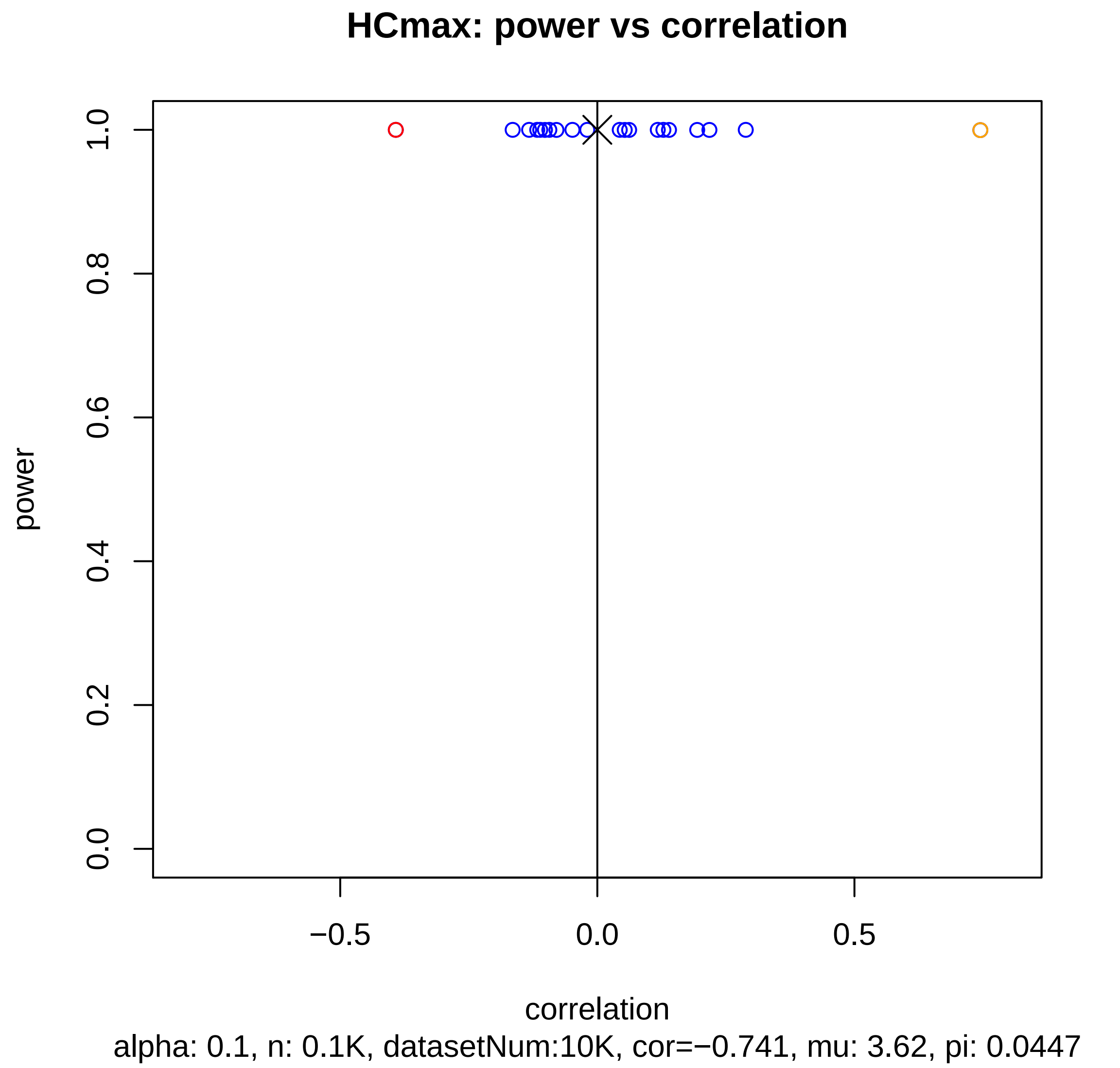
HCmax: power vs correlation
power
0.0
0.2
0.4
0.6
0.8
1.0
−0.5
0.0
0.5
correlation
alpha: 0.1, n: 0.1K, datasetNum:10K, cor=−0.741, mu: 3.62, pi: 0.0447

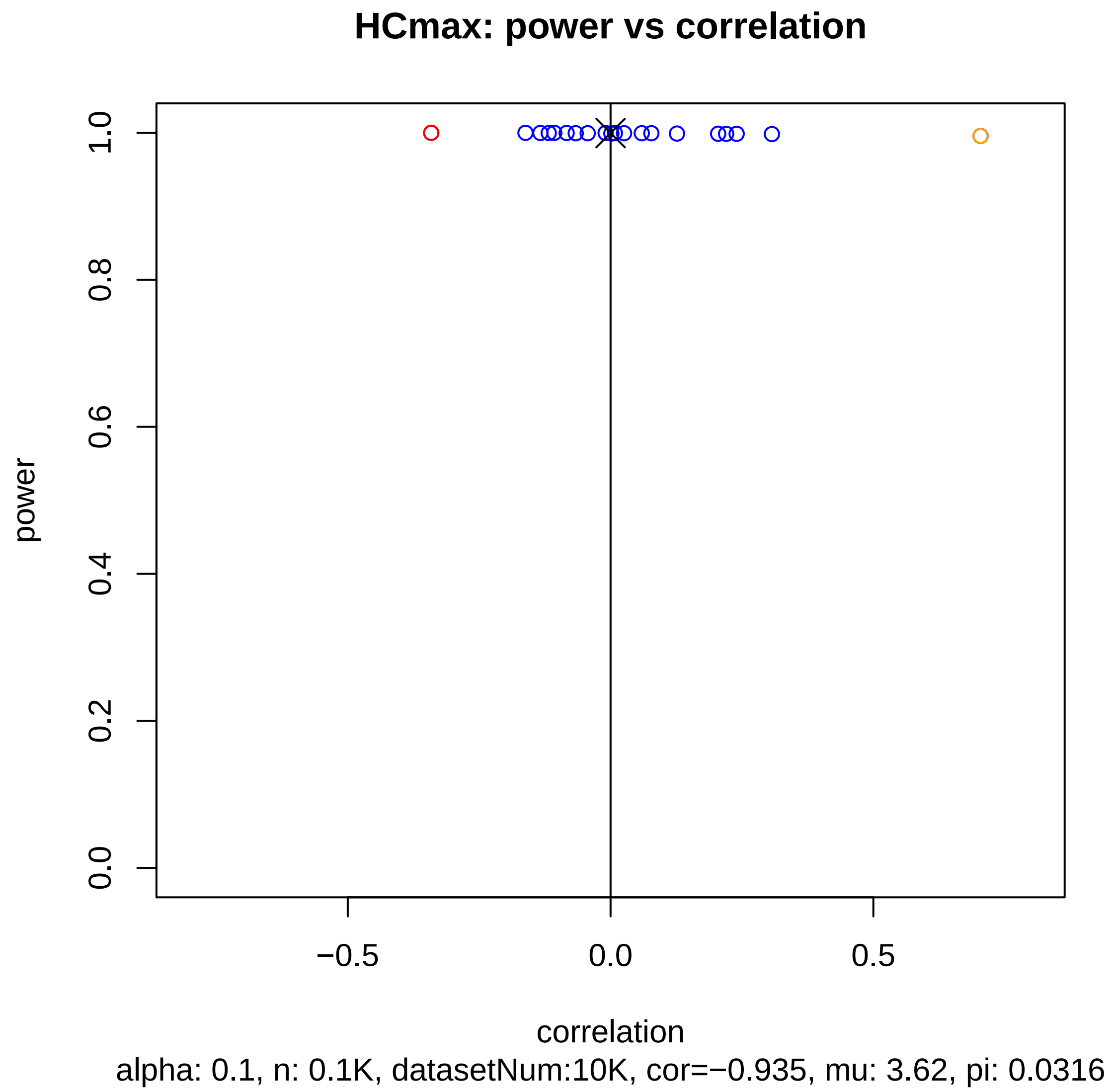
HCmax: power vs correlation
power
1.0
0.8
0.6
0.4
0.2
0.0
−0.5
0.0
0.5
correlation
alpha: 0.1, n: 0.1K, datasetNum:10K, cor=−0.935, mu: 3.62, pi: 0.0316

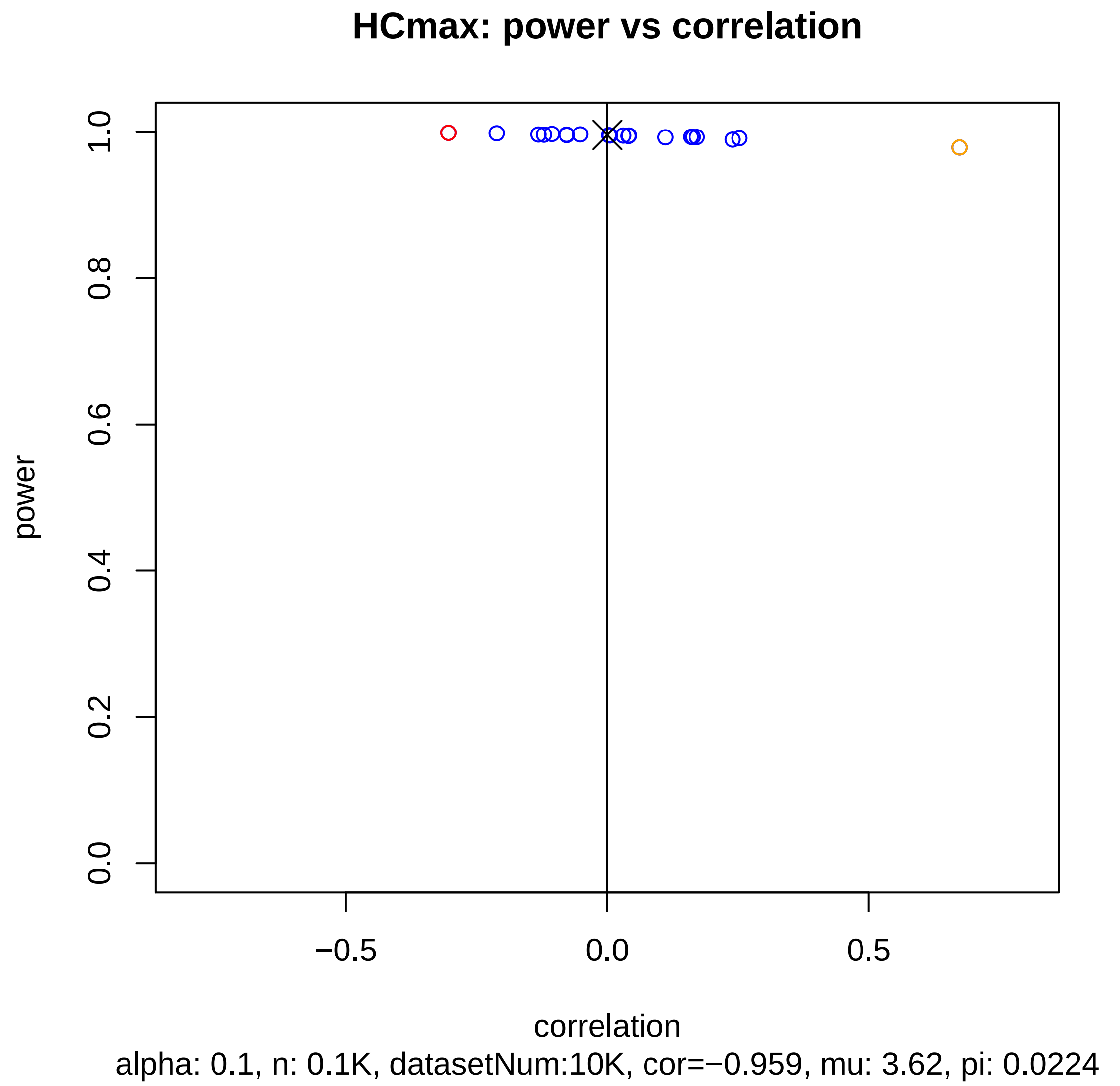
HCmax: power vs correlation
1.0
0.8
0.6
0.4
0.2
0.0
power
−0.5
0.0
0.5
correlation
alpha: 0.1, n: 0.1K, datasetNum:10K, cor=−0.959, mu: 3.62, pi: 0.0224

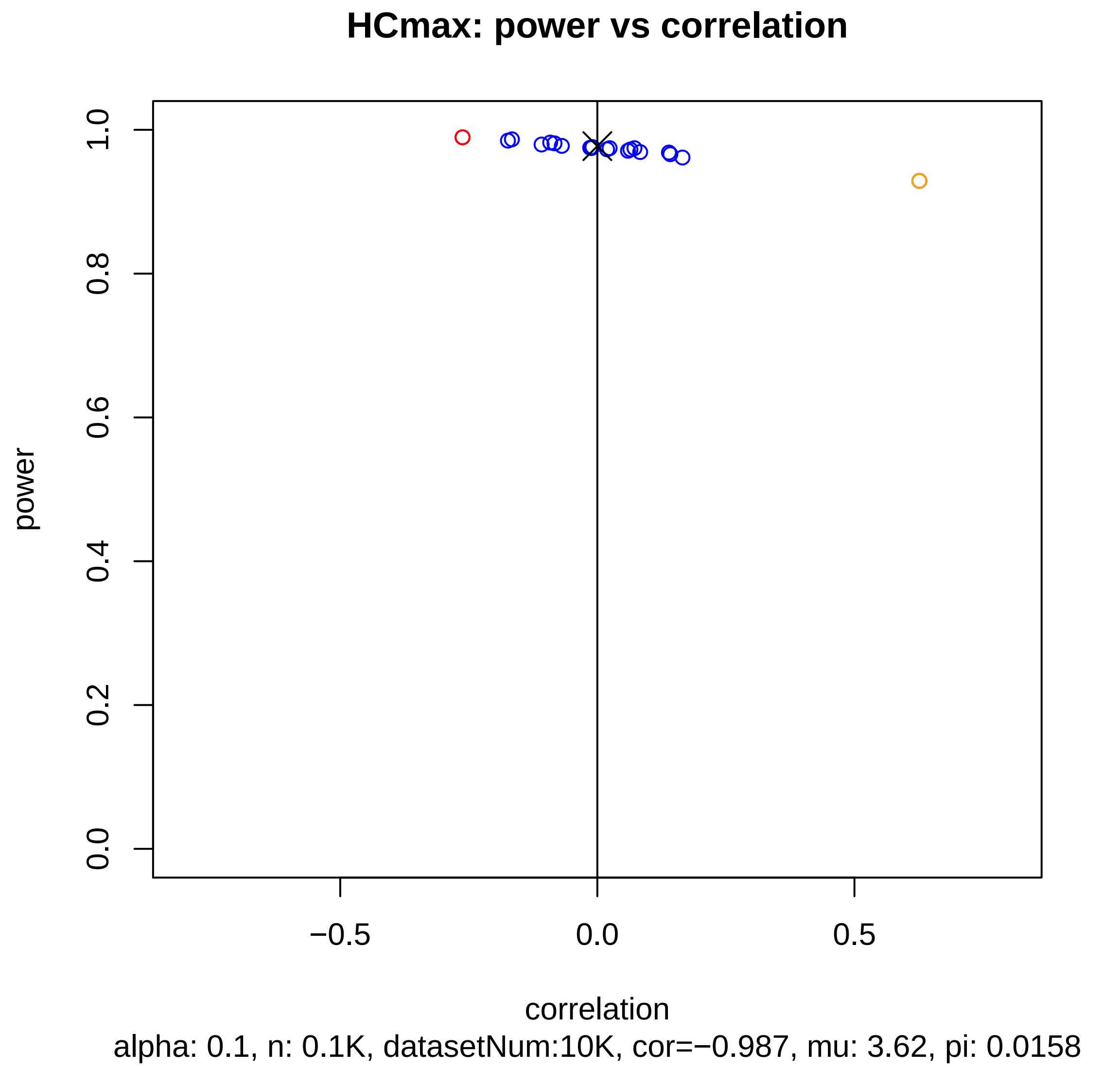
HCmax: power vs correlation
power
0.0
0.2
0.4
0.6
0.8
1.0
−0.5
0.0
0.5
correlation
alpha: 0.1, n: 0.1K, datasetNum:10K, cor=−0.987, mu: 3.62, pi: 0.0158

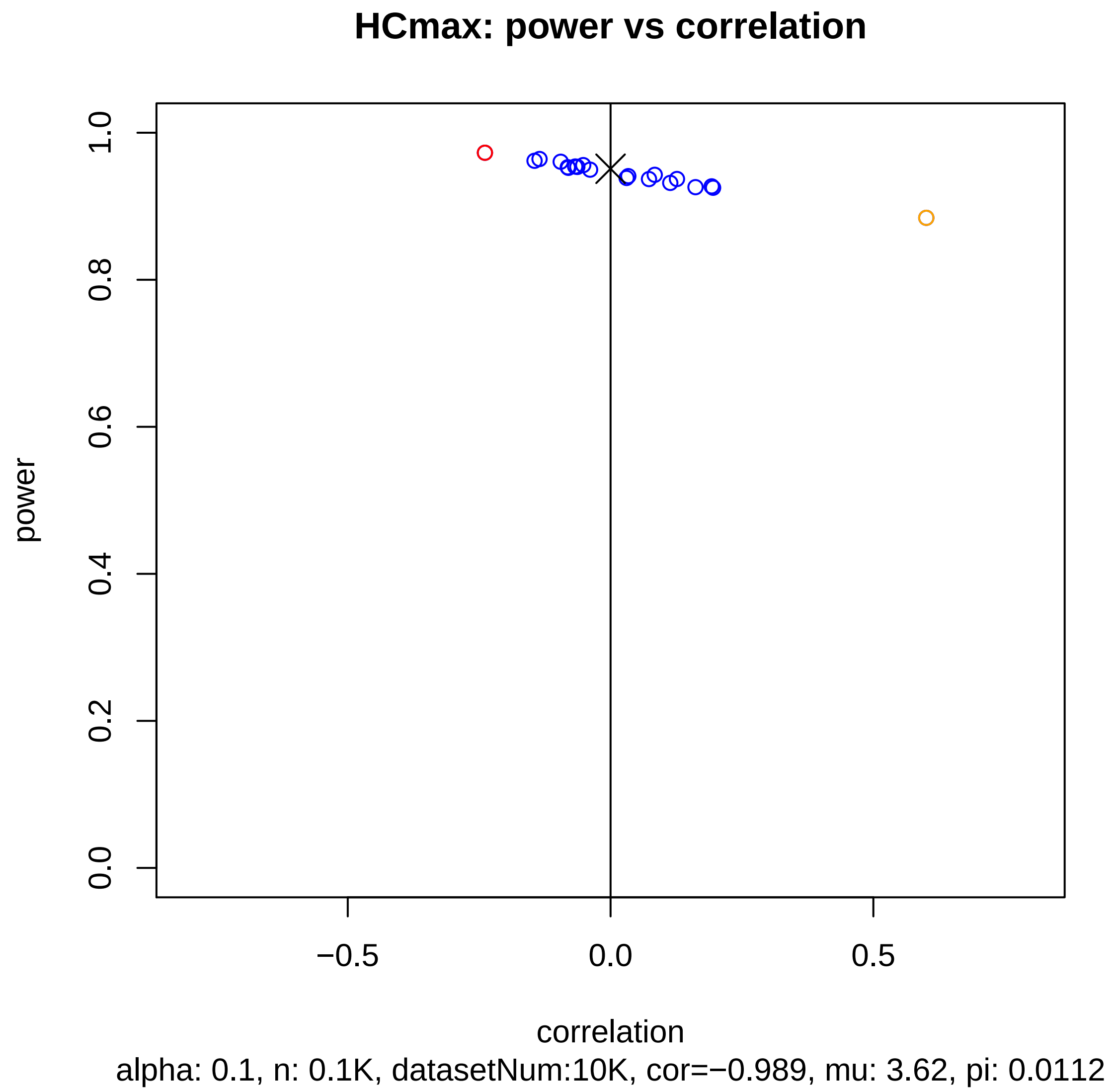

HCmax: power vs correlation
power
1.0
0.8
0.6
0.4
0.2
0.0
−0.5
0.0
0.5
correlation
alpha: 0.1, n: 0.1K, datasetNum:10K, cor=−0.989, mu: 3.62, pi: 0.0112

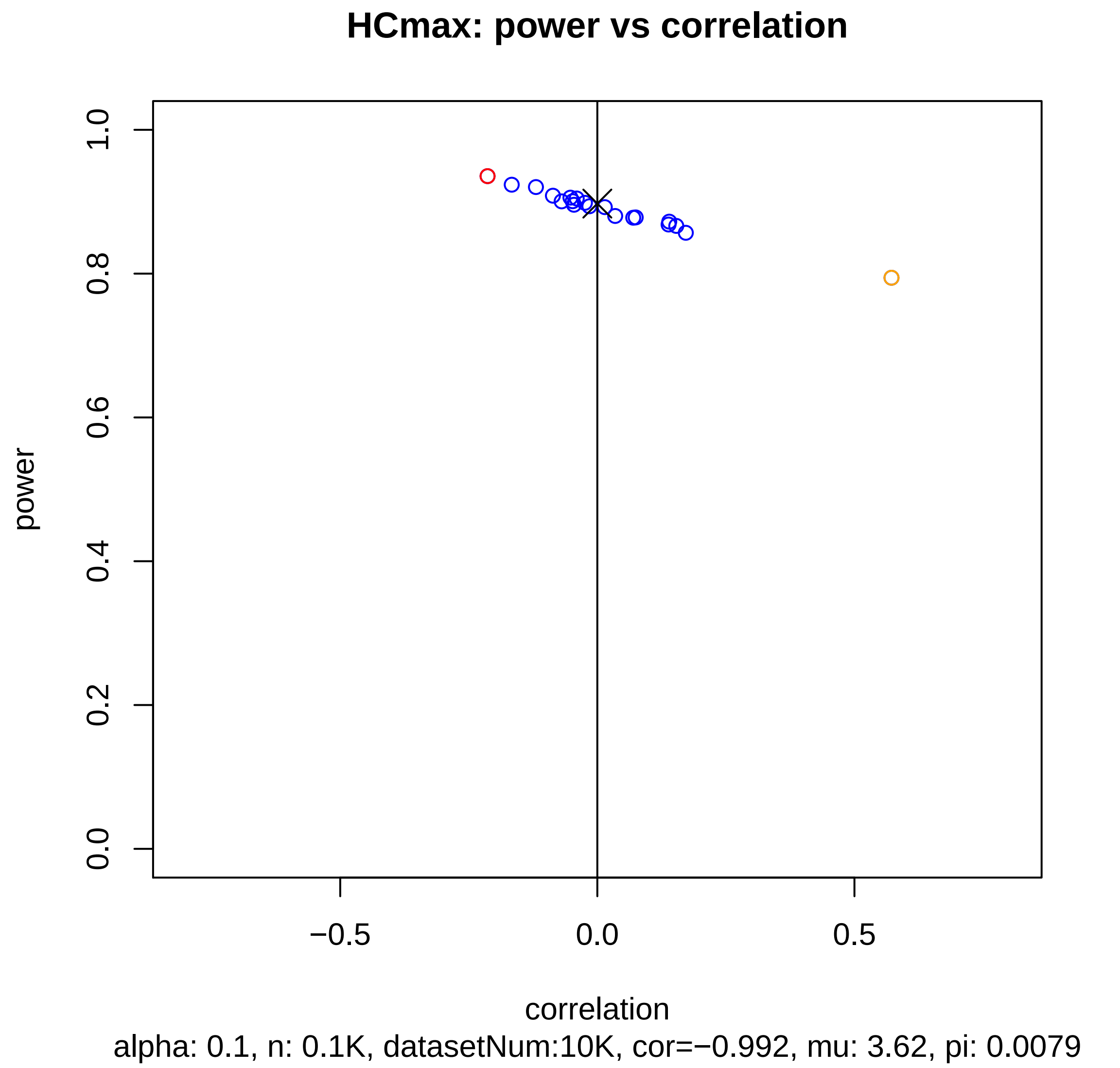
HCmax: power vs correlation
power
0.0
0.2
0.4
0.6
0.8
1.0
−0.5
0.0
0.5
correlation
alpha: 0.1, n: 0.1K, datasetNum:10K, cor=−0.992, mu: 3.62, pi: 0.0079

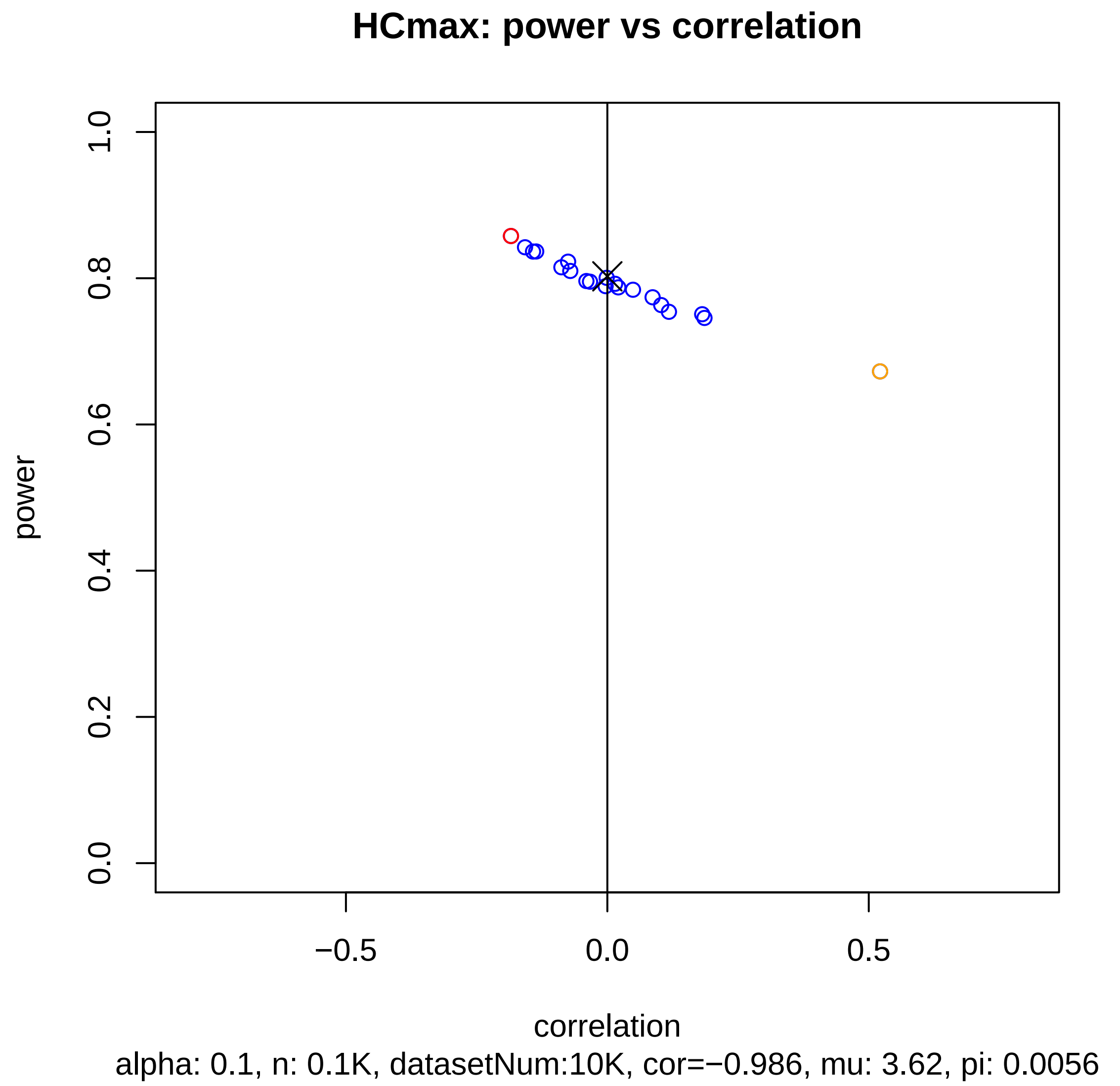
HCmax: power vs correlation
power
0.0
0.2
0.4
0.6
0.8
1.0
−0.5
0.0
0.5
correlation
alpha: 0.1, n: 0.1K, datasetNum:10K, cor=−0.986, mu: 3.62, pi: 0.0056

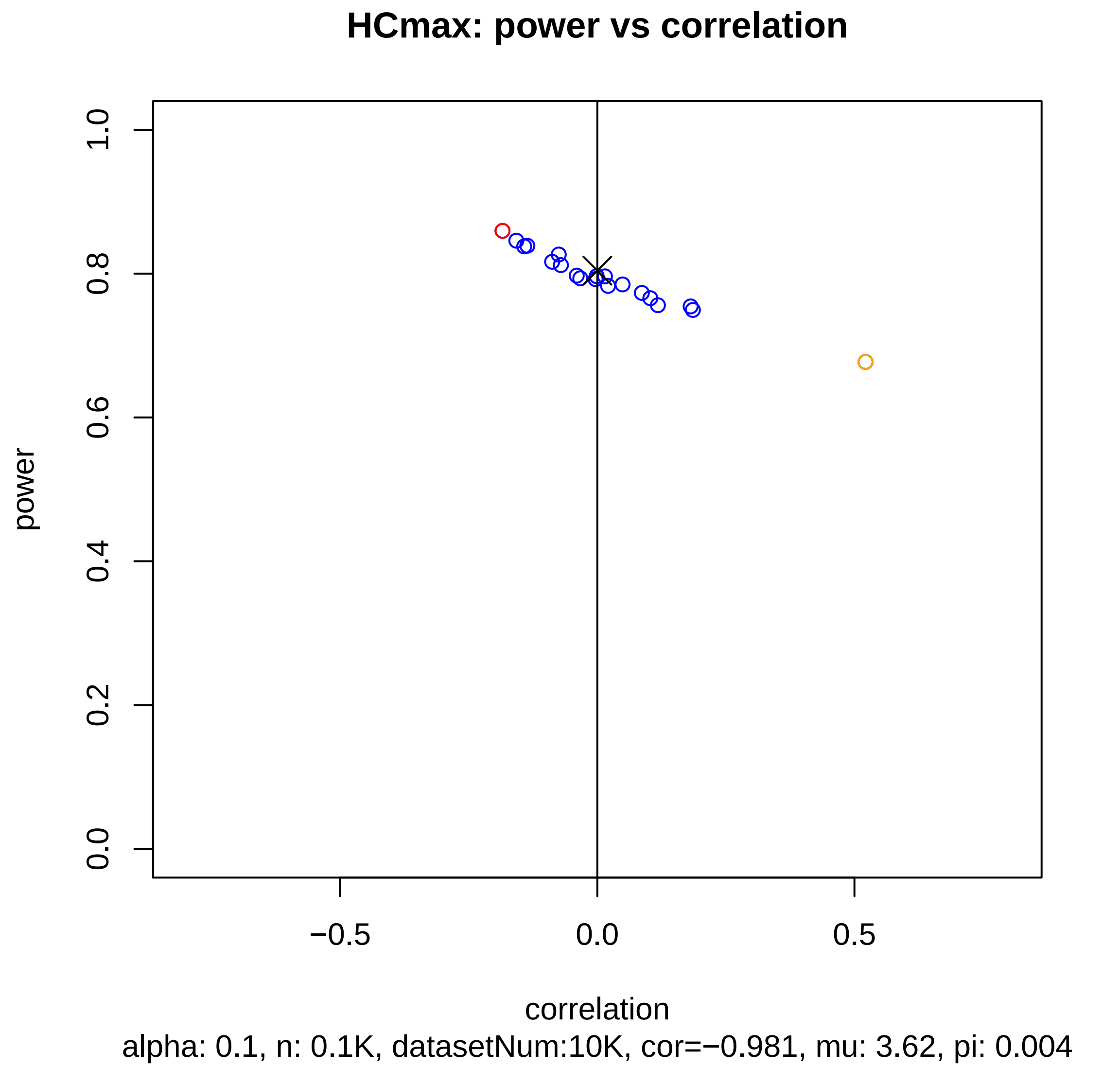
HCmax: power vs correlation
power
0.0
0.2
0.4
0.6
0.8
1.0
−0.5
0.0
0.5
correlation
alpha: 0.1, n: 0.1K, datasetNum:10K, cor=−0.981, mu: 3.62, pi: 0.004

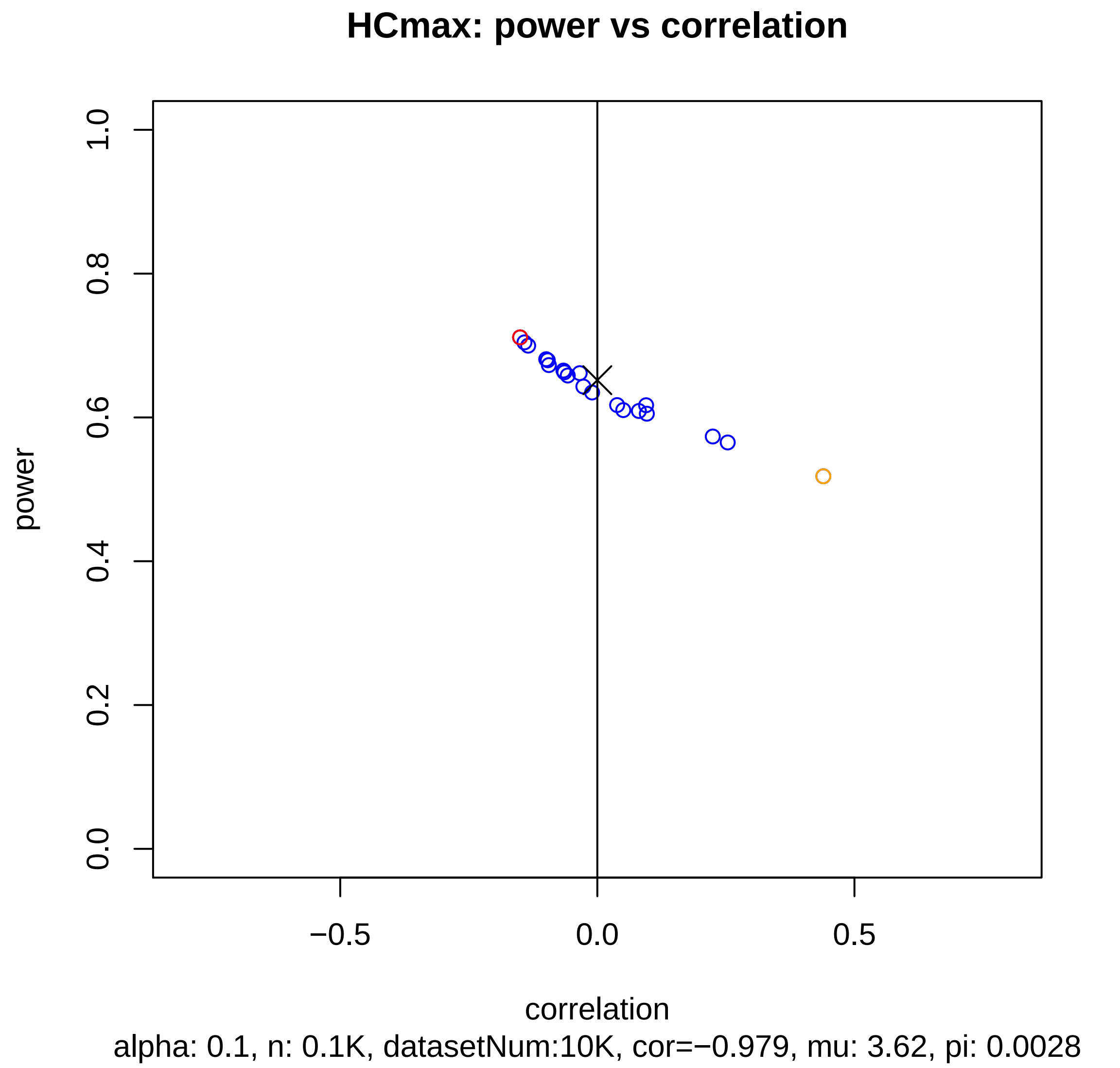
HCmax: power vs correlation
1.0
0.8
0.6
0.4
0.2
0.0
power
−0.5
0.0
0.5
correlation
alpha: 0.1, n: 0.1K, datasetNum:10K, cor=−0.979, mu: 3.62, pi: 0.0028

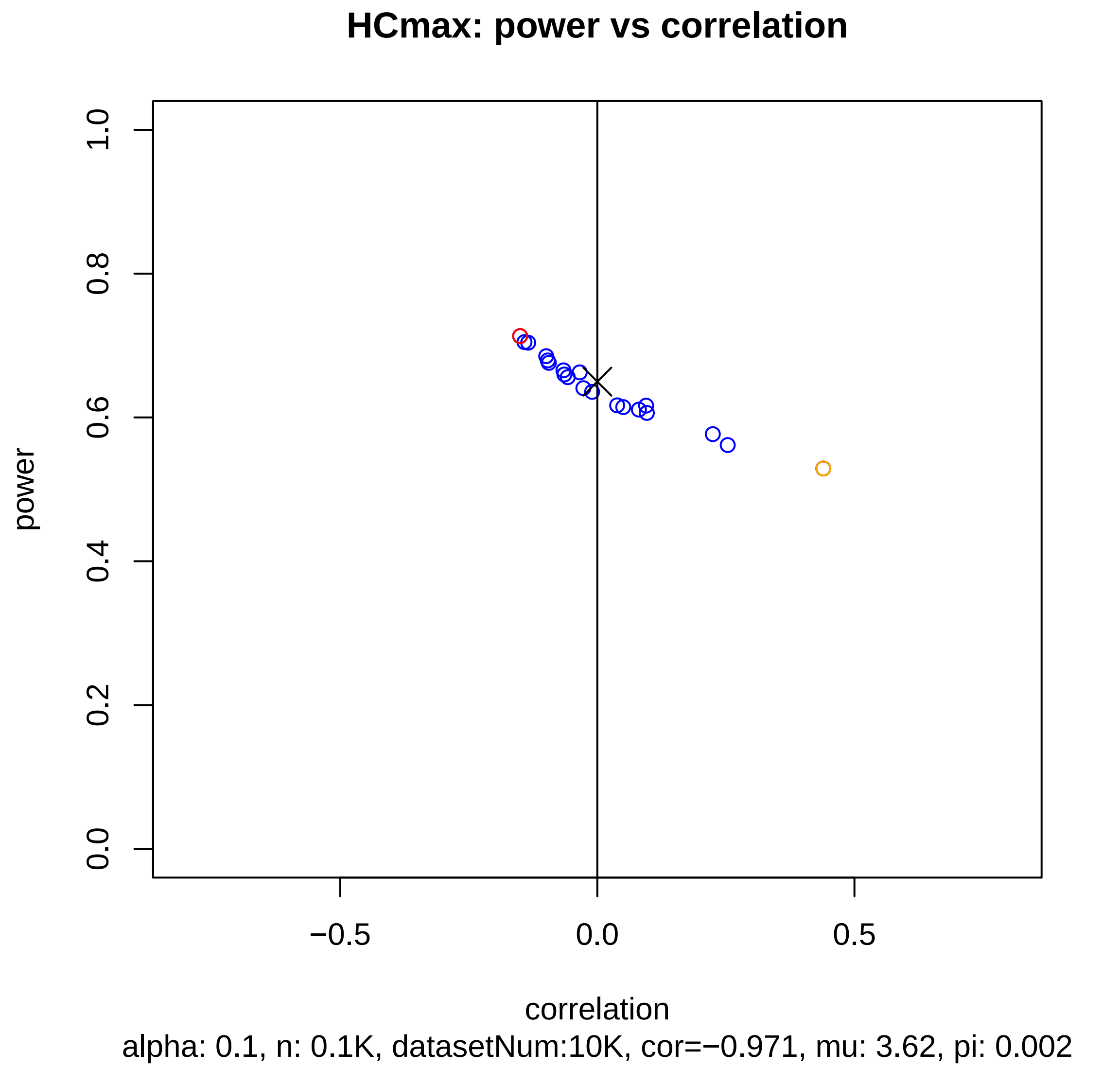
HCmax: power vs correlation
power
0.0
0.2
0.4
0.6
0.8
1.0
−0.5
0.0
0.5
correlation
alpha: 0.1, n: 0.1K, datasetNum:10K, cor=−0.971, mu: 3.62, pi: 0.002

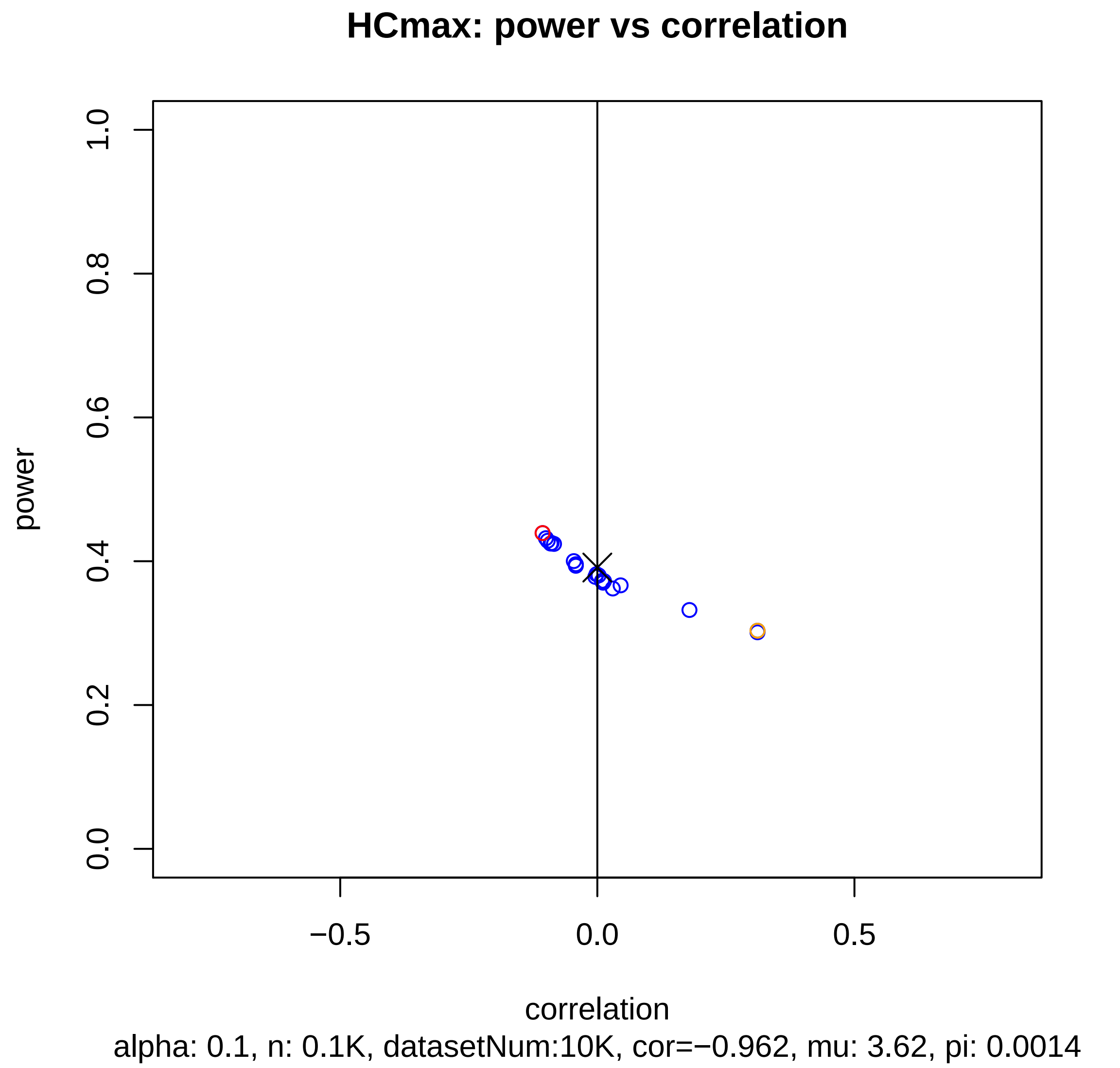
HCmax: power vs correlation
power
1.0
0.8
0.6
0.4
0.2
0.0
−0.5
0.0
0.5
correlation
alpha: 0.1, n: 0.1K, datasetNum:10K, cor=−0.962, mu: 3.62, pi: 0.0014

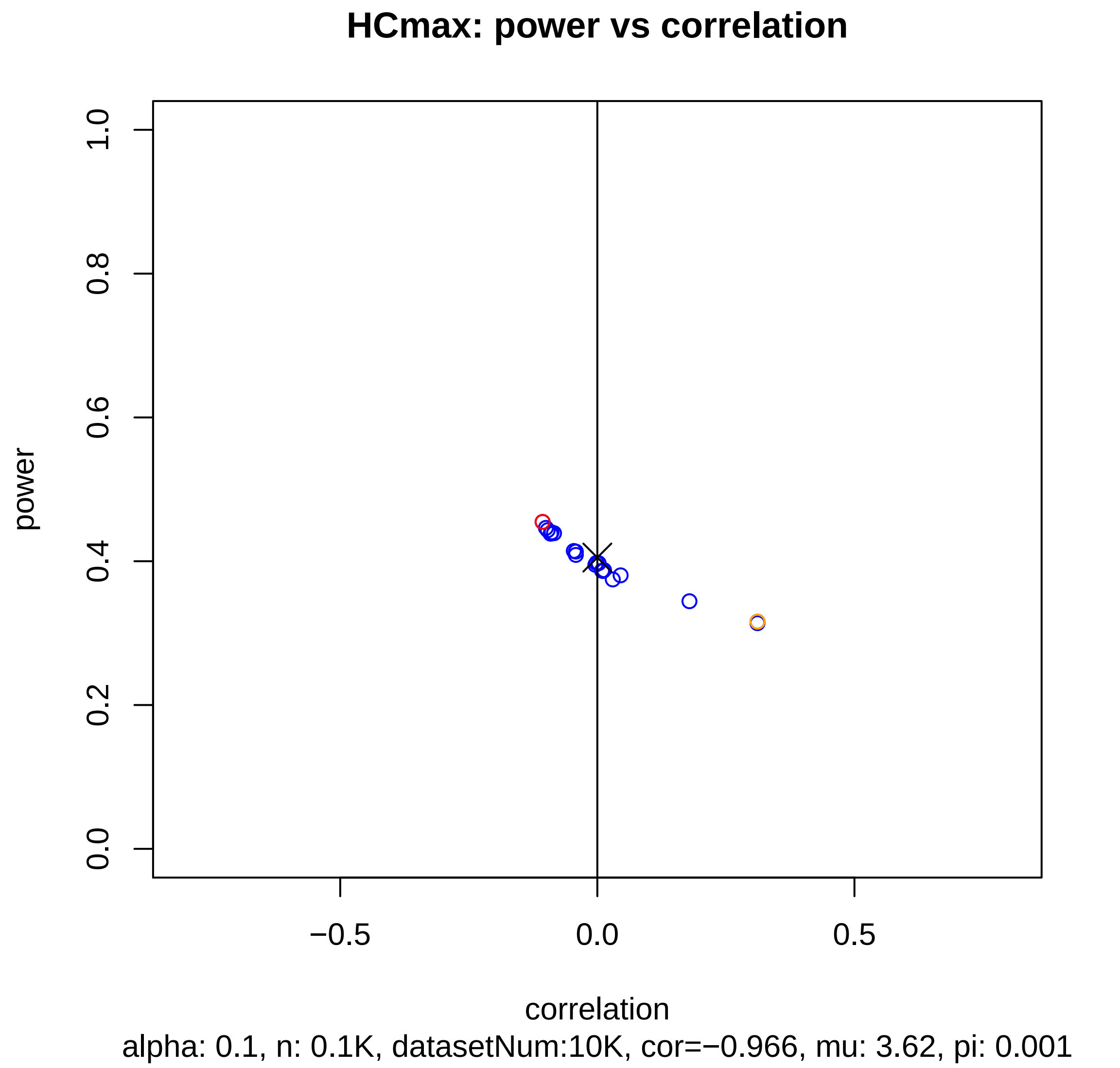
HCmax: power vs correlation
power
0.0
0.2
0.4
0.6
0.8
1.0
−0.5
0.0
0.5
correlation
alpha: 0.1, n: 0.1K, datasetNum:10K, cor=−0.966, mu: 3.62, pi: 0.001

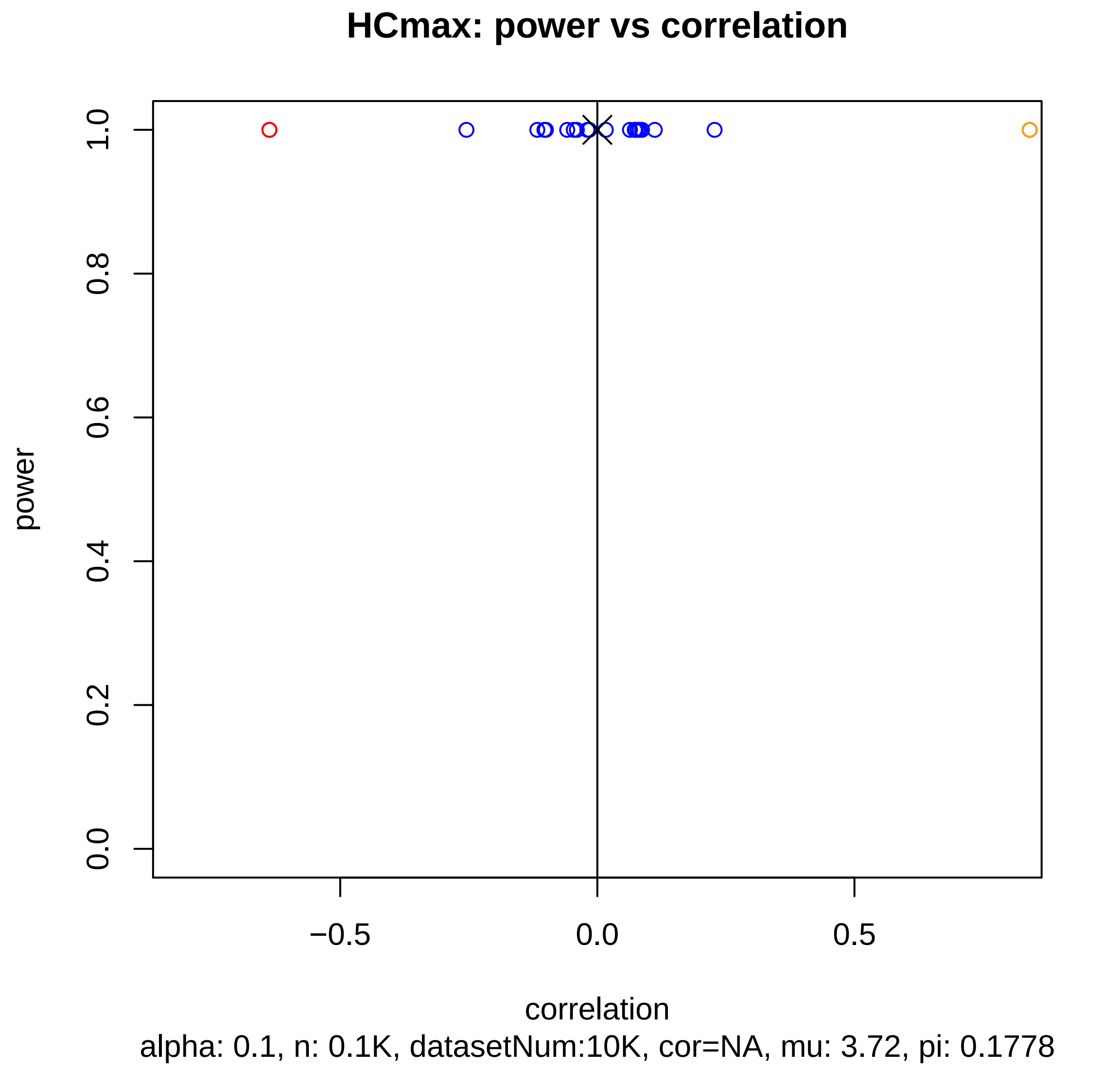
HCmax: power vs correlation
power
1.0
0.8
0.6
0.4
0.2
0.0
−0.5
0.0
0.5
correlation
alpha: 0.1, n: 0.1K, datasetNum:10K, cor=NA, mu: 3.72, pi: 0.1778

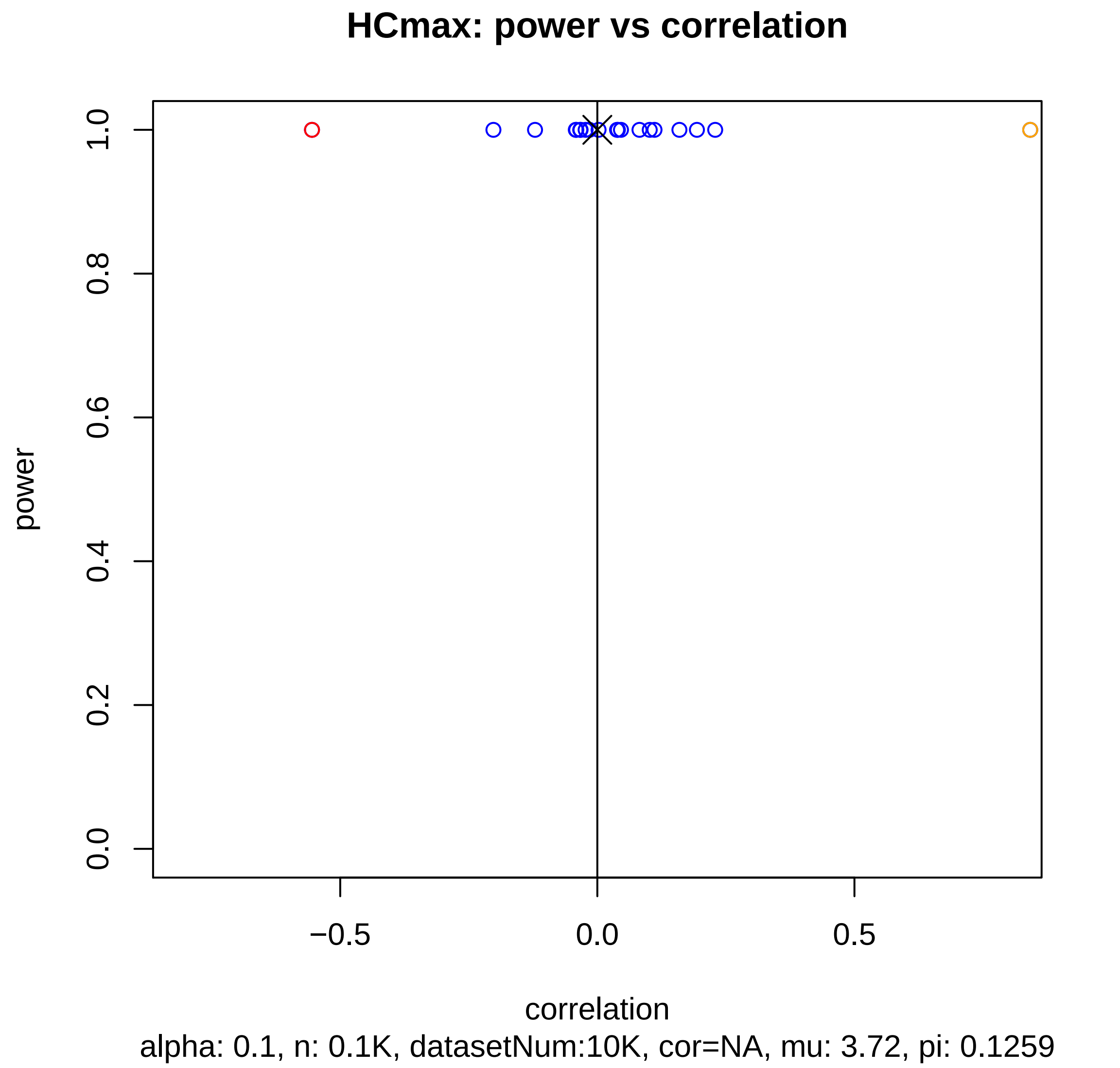

HCmax: power vs correlation
power
1.0
0.8
0.6
0.4
0.2
0.0
−0.5
0.0
0.5
correlation
alpha: 0.1, n: 0.1K, datasetNum:10K, cor=NA, mu: 3.72, pi: 0.1259

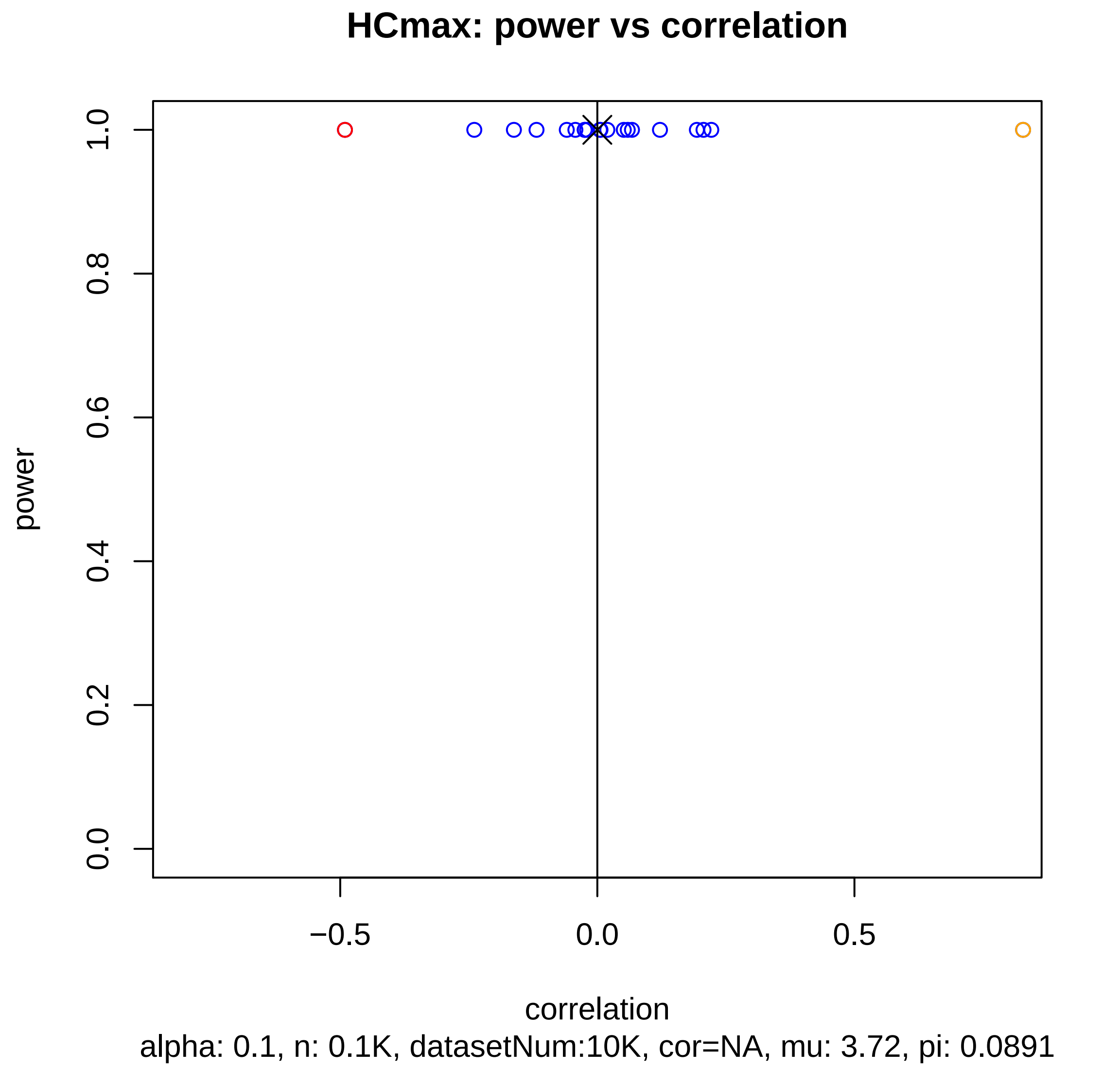
HCmax: power vs correlation
power
1.0
0.8
0.6
0.4
0.2
0.0
−0.5
0.0
0.5
correlation
alpha: 0.1, n: 0.1K, datasetNum:10K, cor=NA, mu: 3.72, pi: 0.0891

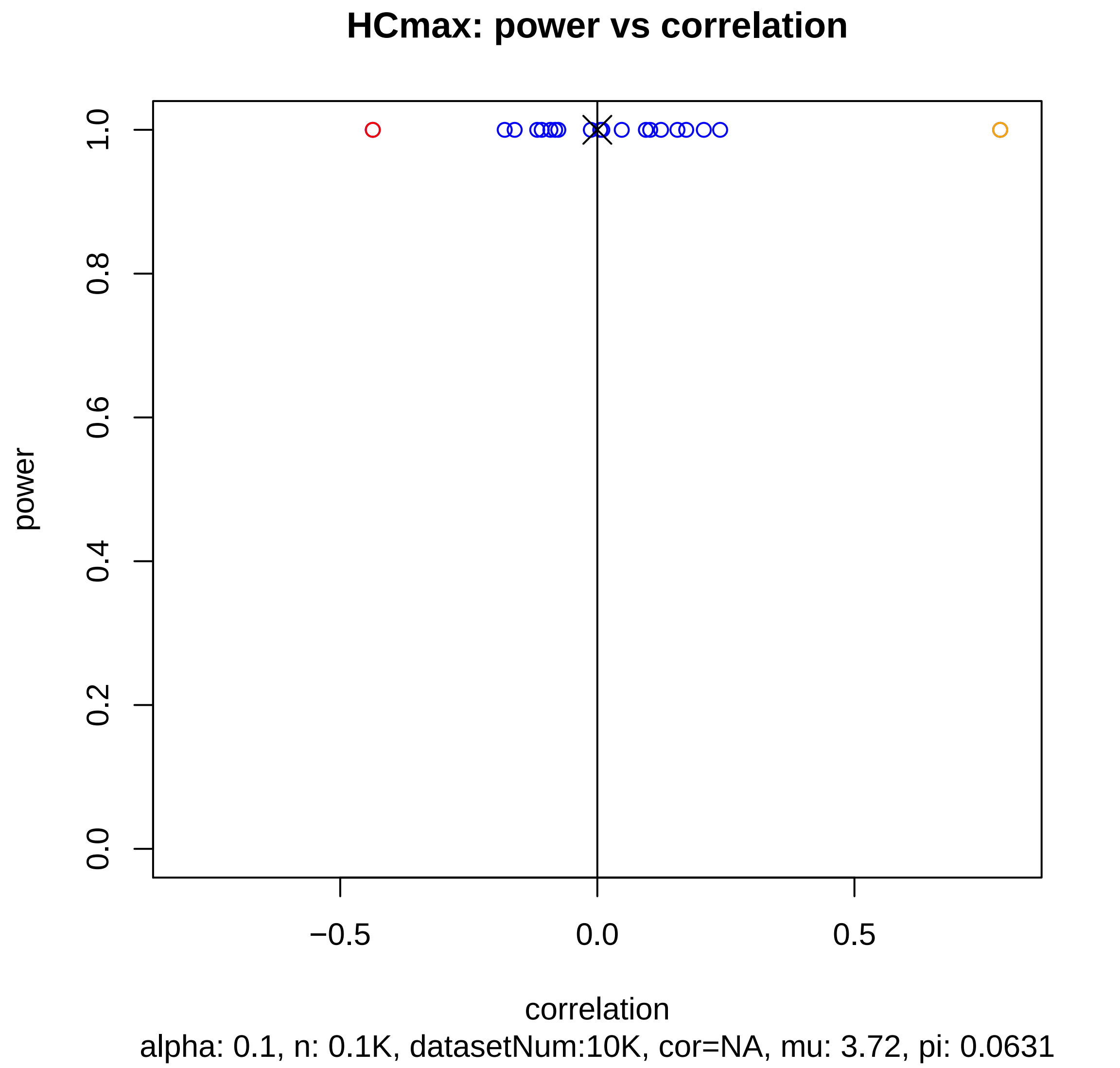
HCmax: power vs correlation
power
0.0
0.2
0.4
0.6
0.8
1.0
−0.5
0.0
0.5
correlation
alpha: 0.1, n: 0.1K, datasetNum:10K, cor=NA, mu: 3.72, pi: 0.0631

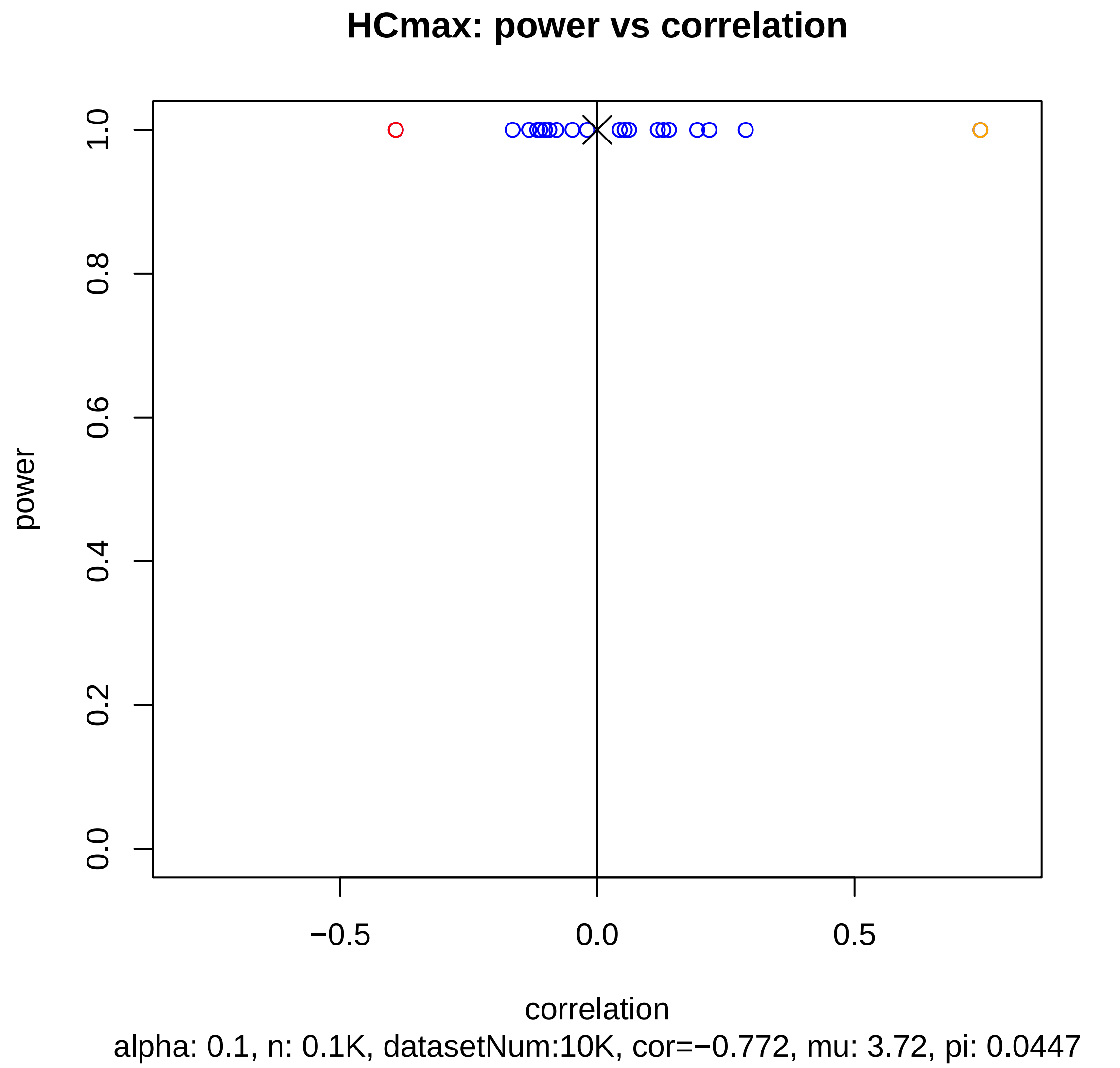
HCmax: power vs correlation
power
0.0
0.2
0.4
0.6
0.8
1.0
−0.5
0.0
0.5
correlation
alpha: 0.1, n: 0.1K, datasetNum:10K, cor=−0.772, mu: 3.72, pi: 0.0447

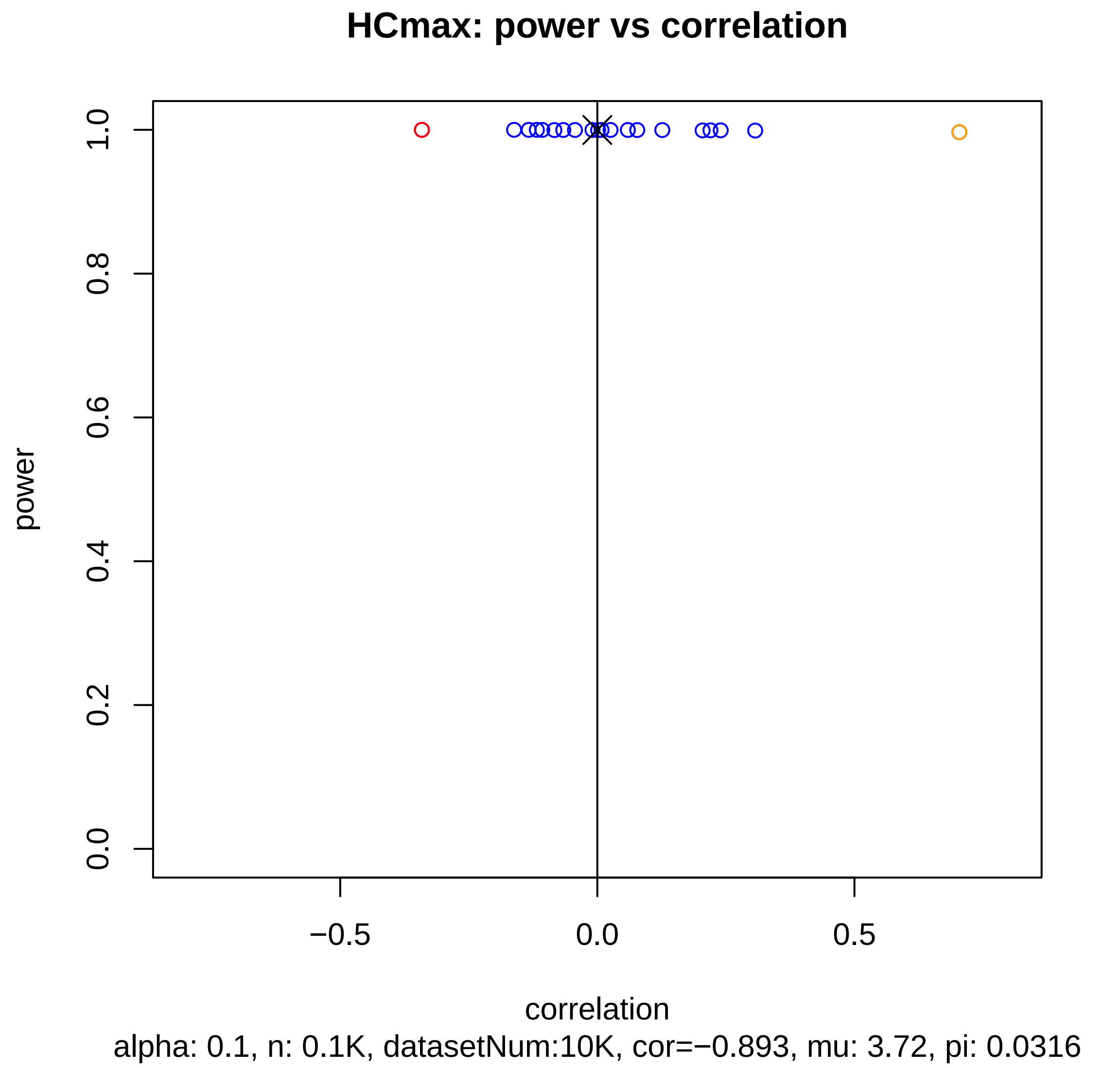
HCmax: power vs correlation
power
1.0
0.8
0.6
0.4
0.2
0.0
−0.5
0.0
0.5
correlation
alpha: 0.1, n: 0.1K, datasetNum:10K, cor=−0.893, mu: 3.72, pi: 0.0316

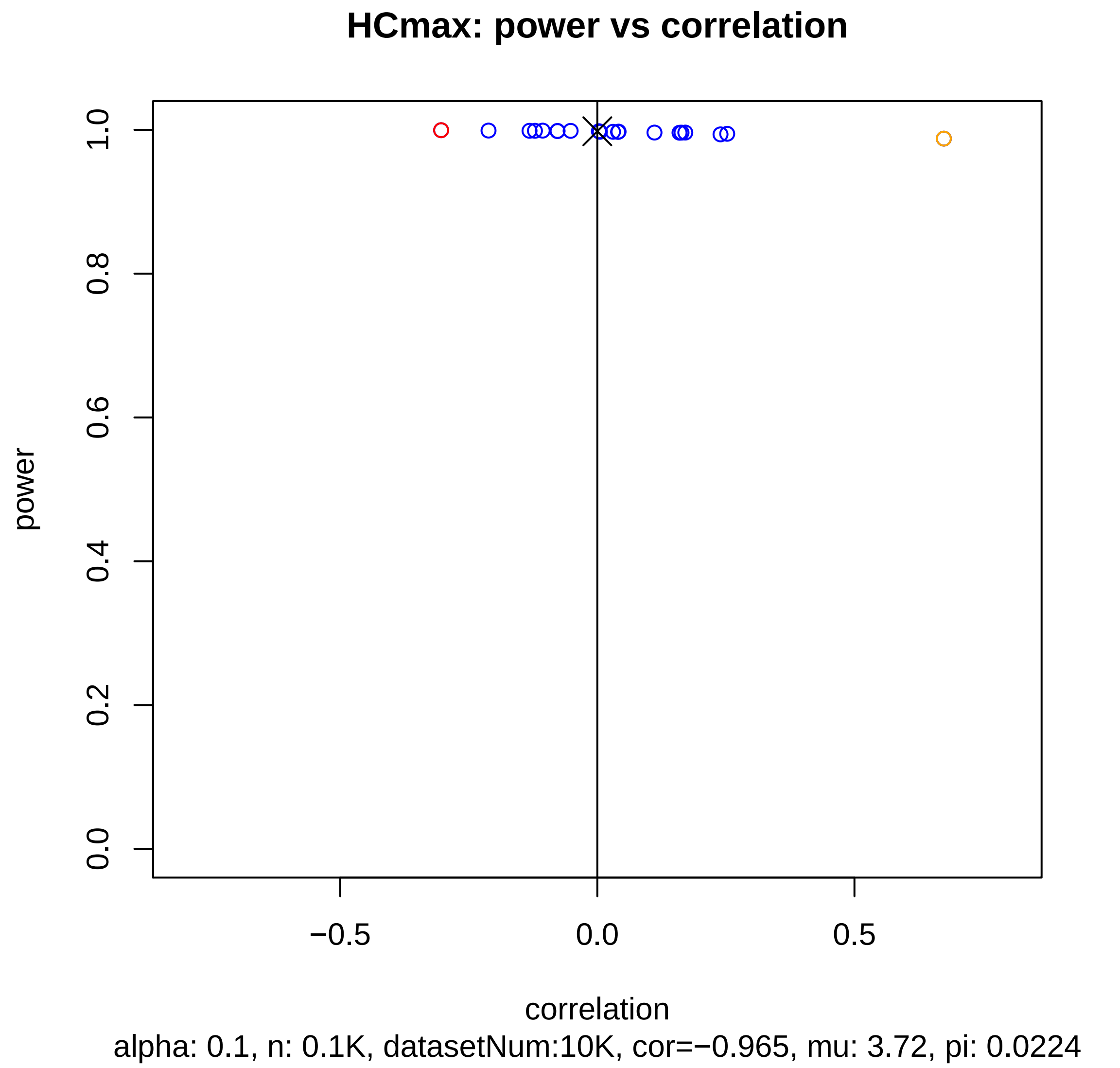

HCmax: power vs correlation
power
1.0
0.8
0.6
0.4
0.2
0.0
−0.5
0.0
0.5
correlation
alpha: 0.1, n: 0.1K, datasetNum:10K, cor=−0.965, mu: 3.72, pi: 0.0224

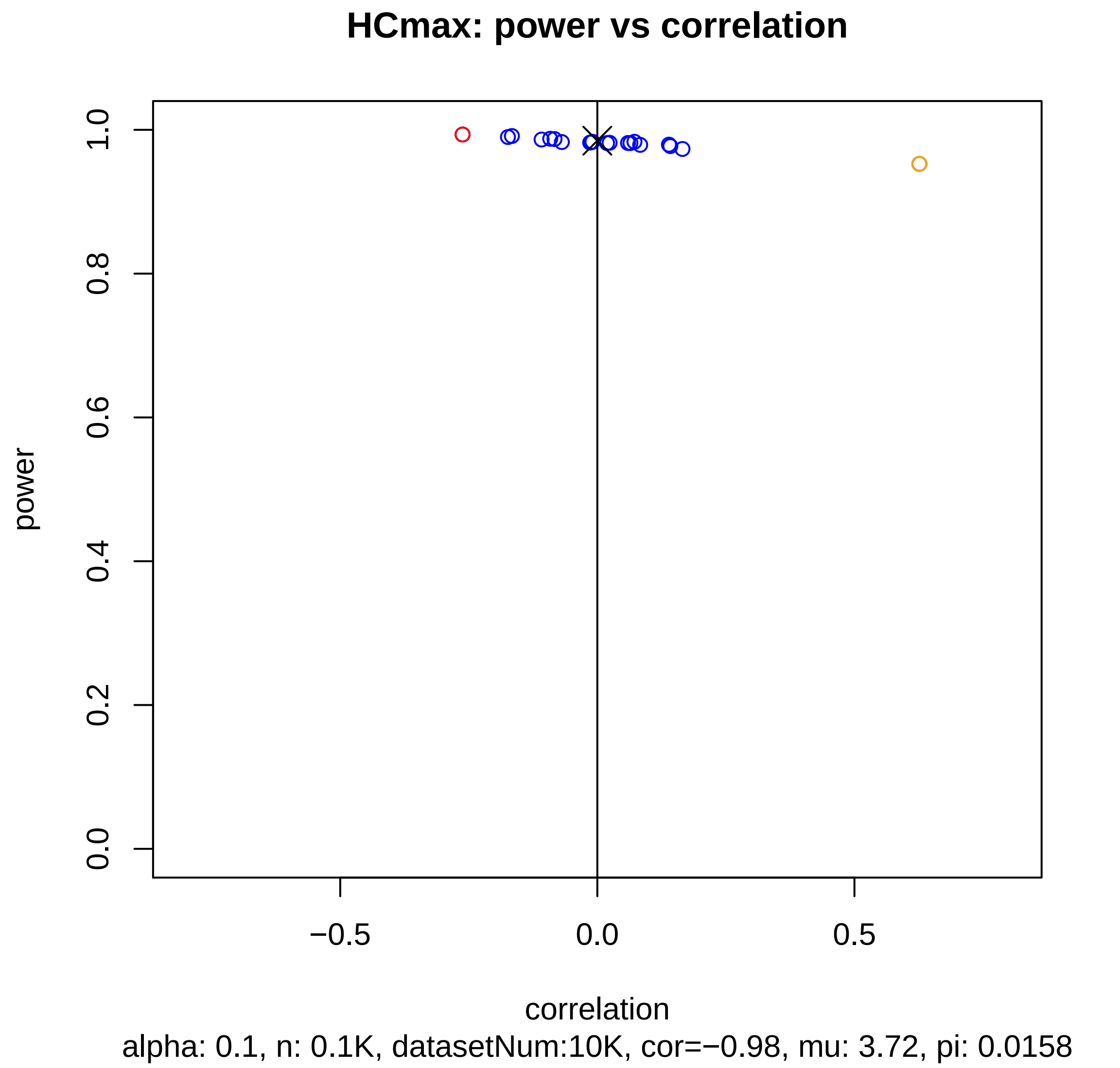

HCmax: power vs correlation
power
1.0
0.8
0.6
0.4
0.2
0.0
−0.5
0.0
0.5
correlation
alpha: 0.1, n: 0.1K, datasetNum:10K, cor=−0.98, mu: 3.72, pi: 0.0158

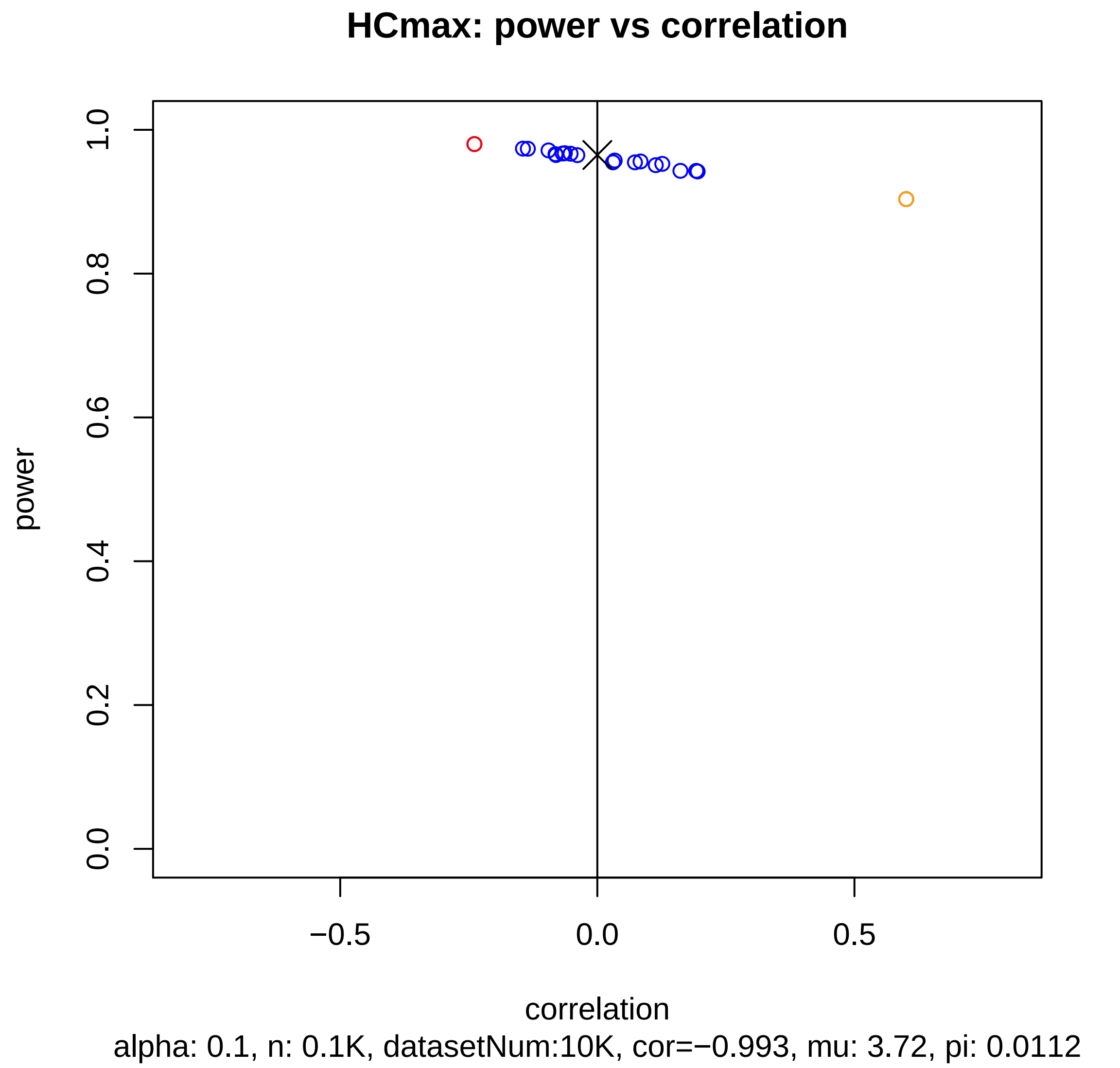
HCmax: power vs correlation
power
0.0
0.2
0.4
0.6
0.8
1.0
correlation
−0.5
0.0
0.5
alpha: 0.1, n: 0.1K, datasetNum:10K, cor=−0.993, mu: 3.72, pi: 0.0112

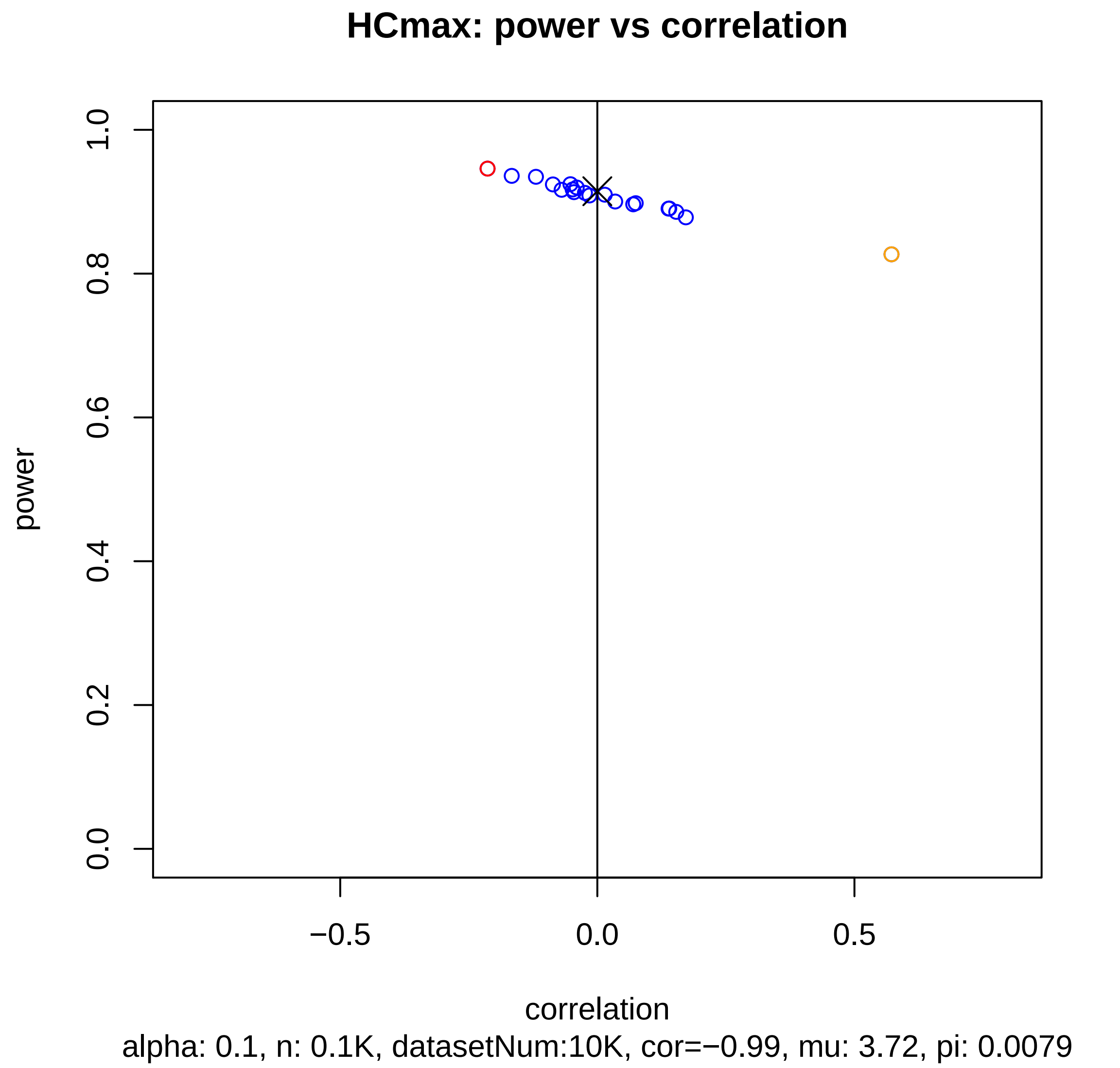
HCmax: power vs correlation
power
1.0
0.8
0.6
0.4
0.2
0.0
−0.5
0.0
0.5
correlation
alpha: 0.1, n: 0.1K, datasetNum:10K, cor=−0.99, mu: 3.72, pi: 0.0079

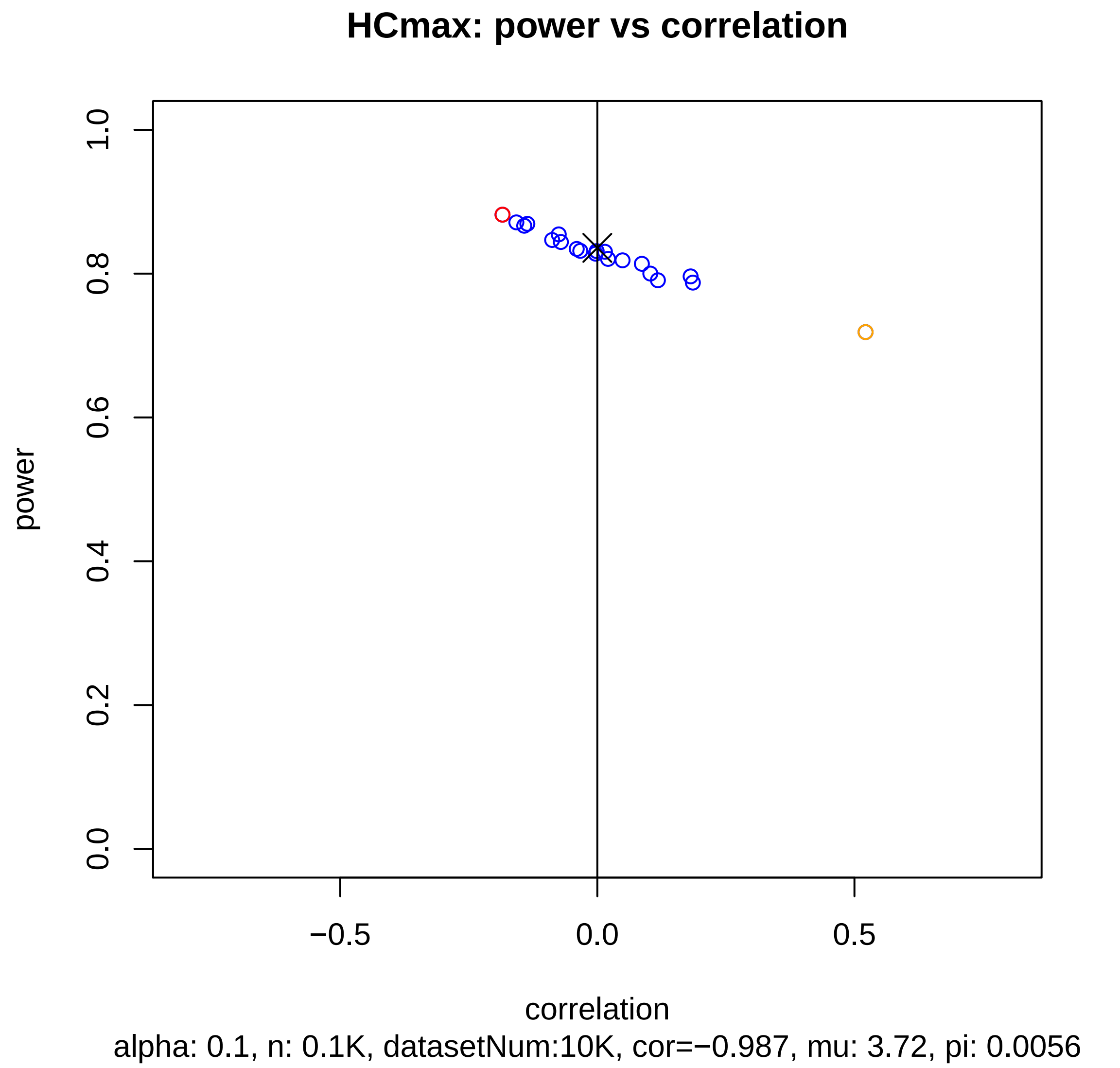
HCmax: power vs correlation
power
1.0
0.8
0.6
0.4
0.2
0.0
−0.5
0.0
0.5
correlation
alpha: 0.1, n: 0.1K, datasetNum:10K, cor=−0.987, mu: 3.72, pi: 0.0056

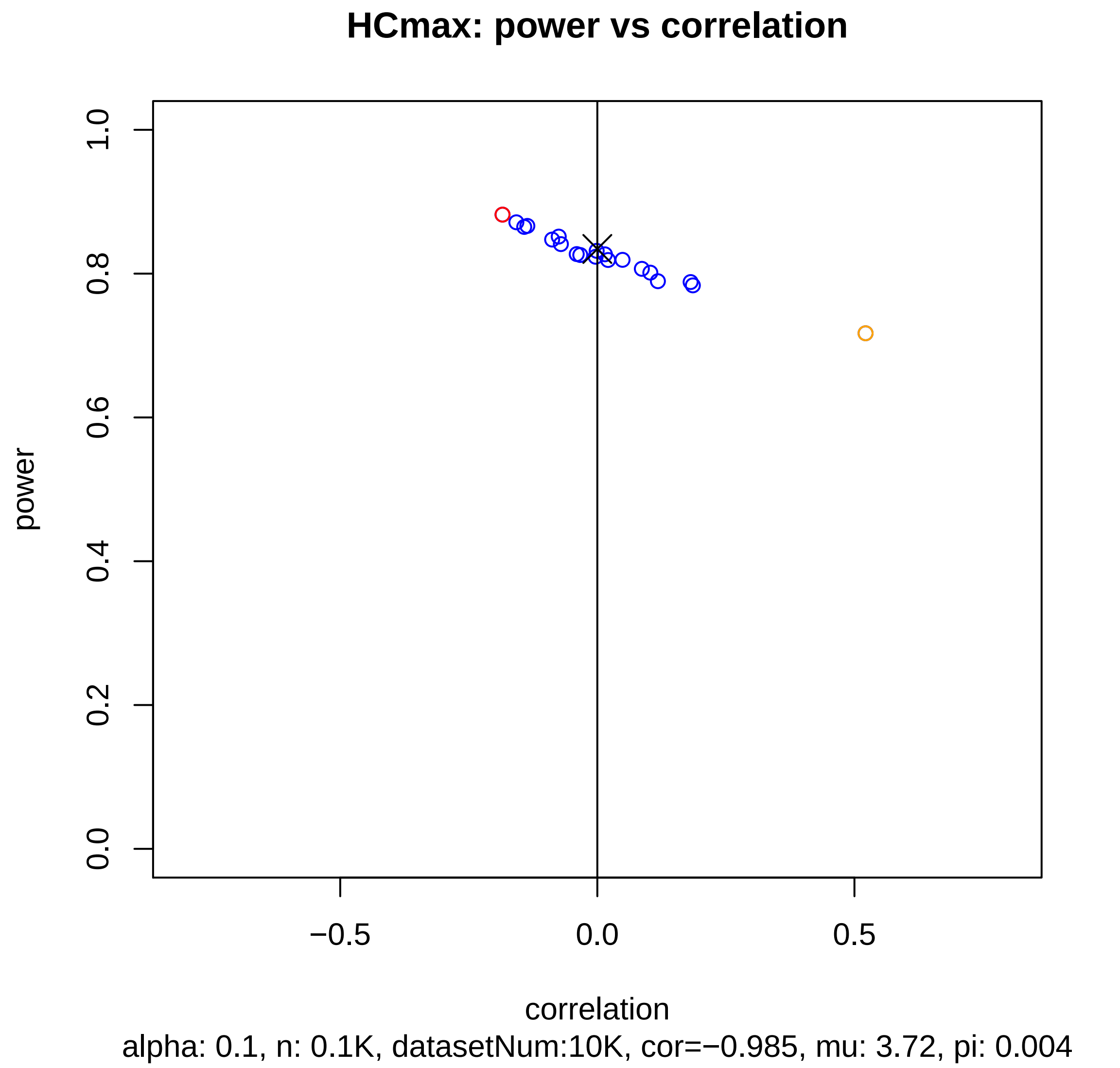
HCmax: power vs correlation
power
0.0
0.2
0.4
0.6
0.8
1.0
−0.5
0.0
0.5
correlation
alpha: 0.1, n: 0.1K, datasetNum:10K, cor=−0.985, mu: 3.72, pi: 0.004

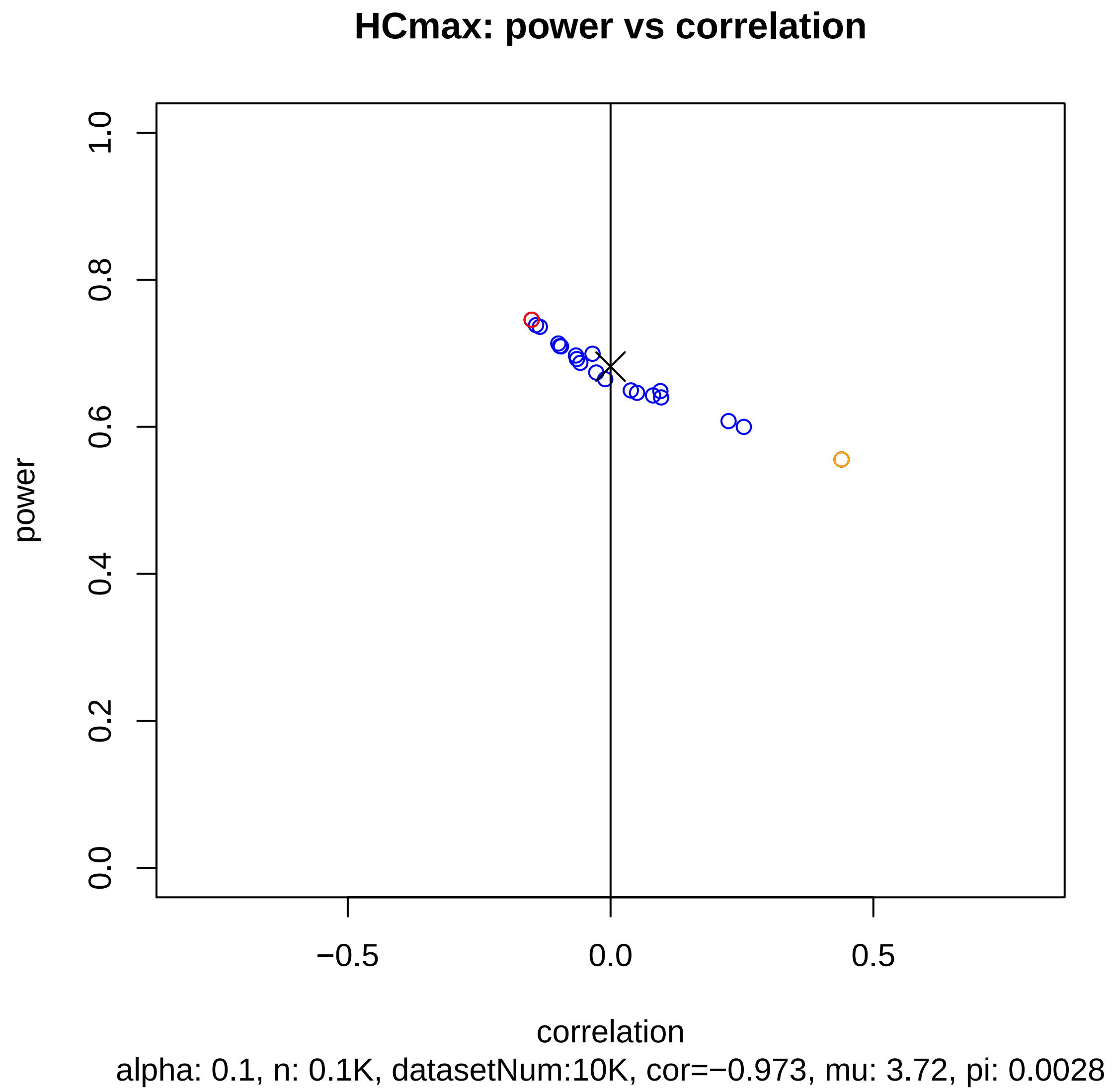
HCmax: power vs correlation
power
0.0
0.2
0.4
0.6
0.8
1.0
−0.5
0.0
0.5
correlation
alpha: 0.1, n: 0.1K, datasetNum:10K, cor=−0.973, mu: 3.72, pi: 0.0028

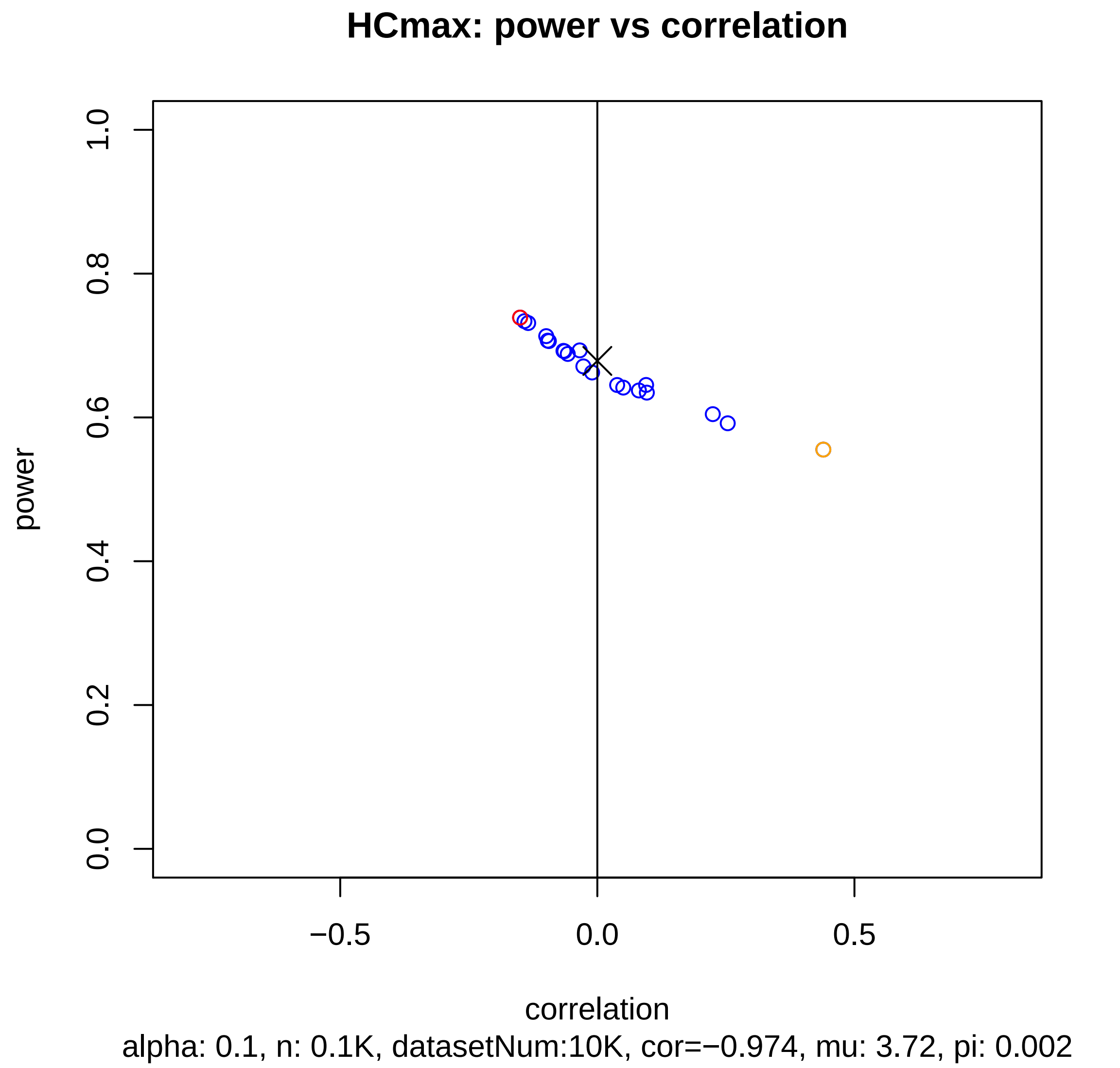
HCmax: power vs correlation
power
0.0
0.2
0.4
0.6
0.8
1.0
−0.5
0.0
0.5
correlation
alpha: 0.1, n: 0.1K, datasetNum:10K, cor=−0.974, mu: 3.72, pi: 0.002

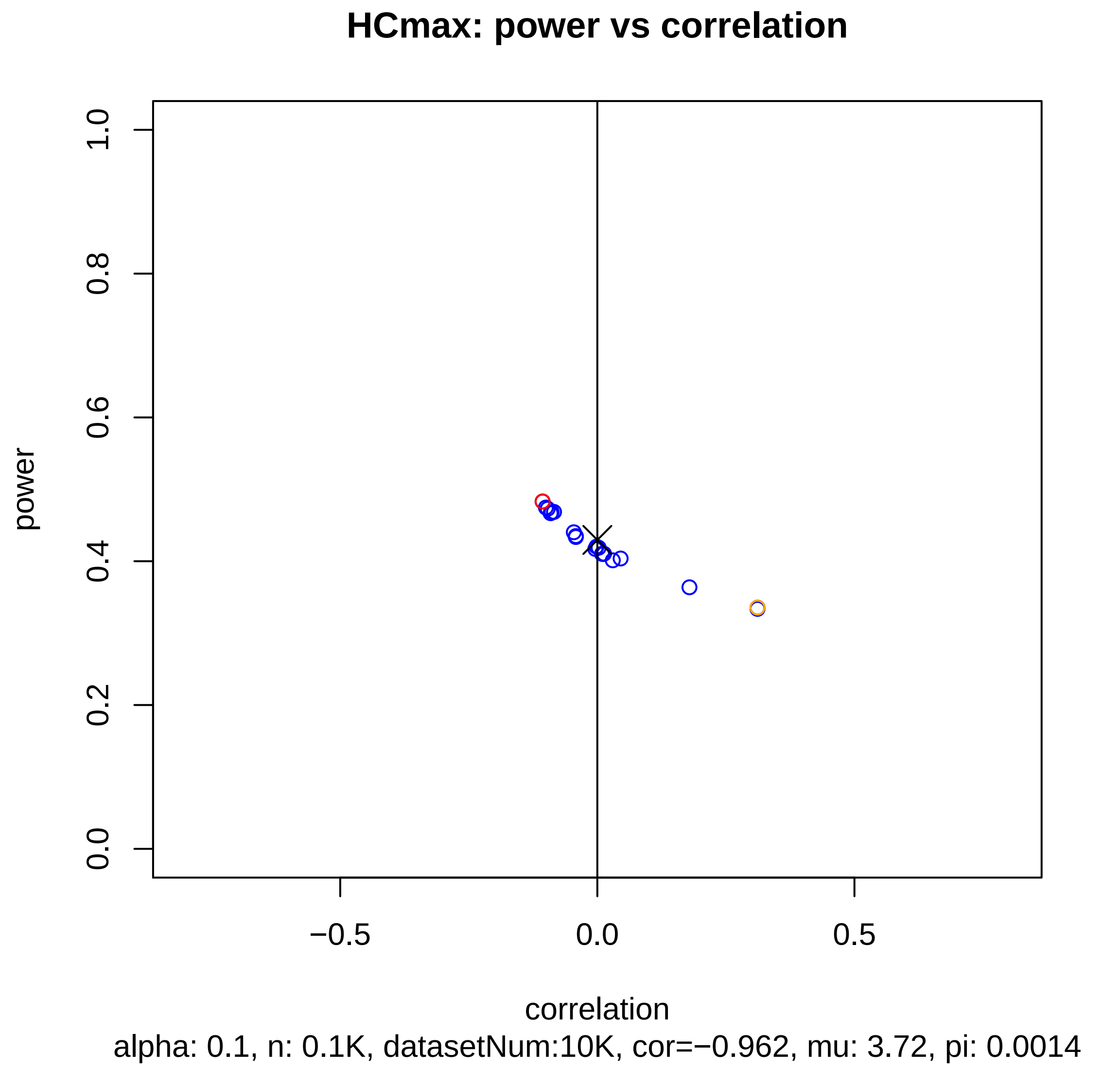
HCmax: power vs correlation
power
1.0
0.8
0.6
0.4
0.2
0.0
−0.5
0.0
0.5
correlation
alpha: 0.1, n: 0.1K, datasetNum:10K, cor=−0.962, mu: 3.72, pi: 0.0014

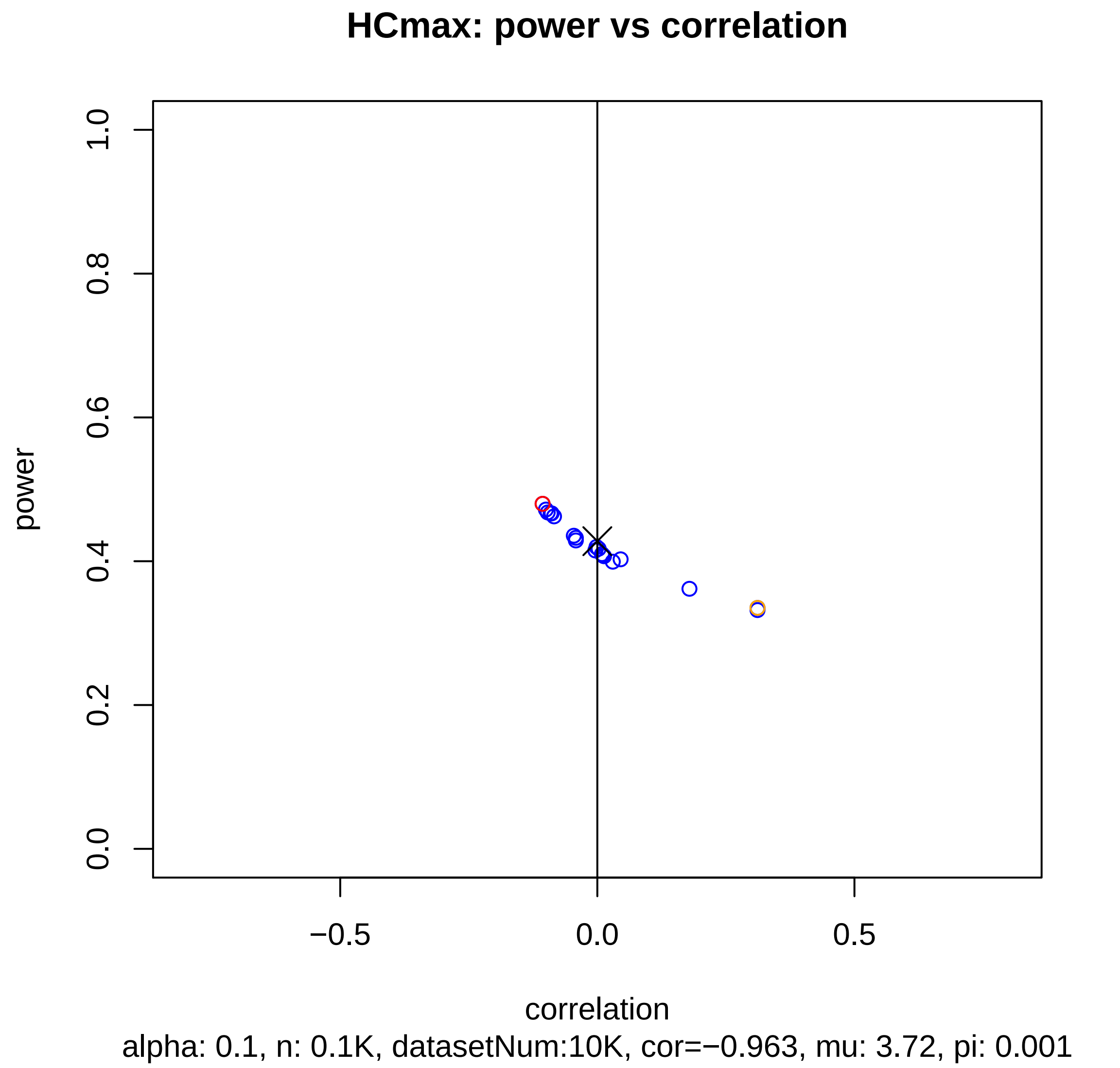
HCmax: power vs correlation
power
1.0
0.8
0.6
0.4
0.2
0.0
−0.5
0.0
0.5
correlation
alpha: 0.1, n: 0.1K, datasetNum:10K, cor=−0.963, mu: 3.72, pi: 0.001
