## supplementary_figure_series_of_power_curve_alpha=0.1_n=1000 for "Incorporating prior information into signal-detection analyses across biologically informed gene-sets"

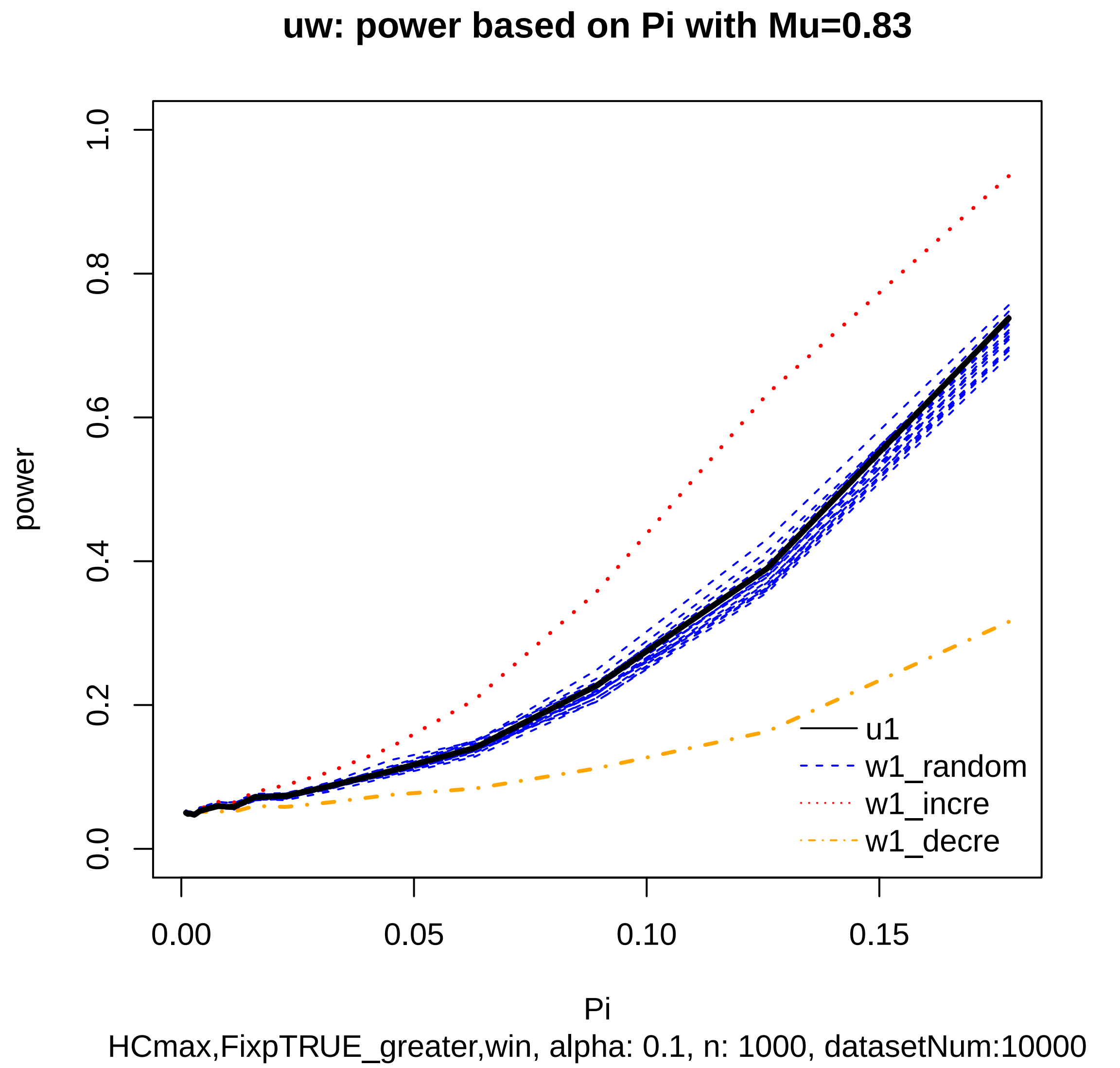
uw: power based on Pi with Mu=0.83
power
1.0
0.8
0.6
0.4
0.2
0.0
0.00
0.05
0.10
0.15
Pi
u1
w1_random
w1_incre
w1_decre
HCmax,FixpTRUE_greater,win, alpha: 0.1, n: 1000, datasetNum:10000

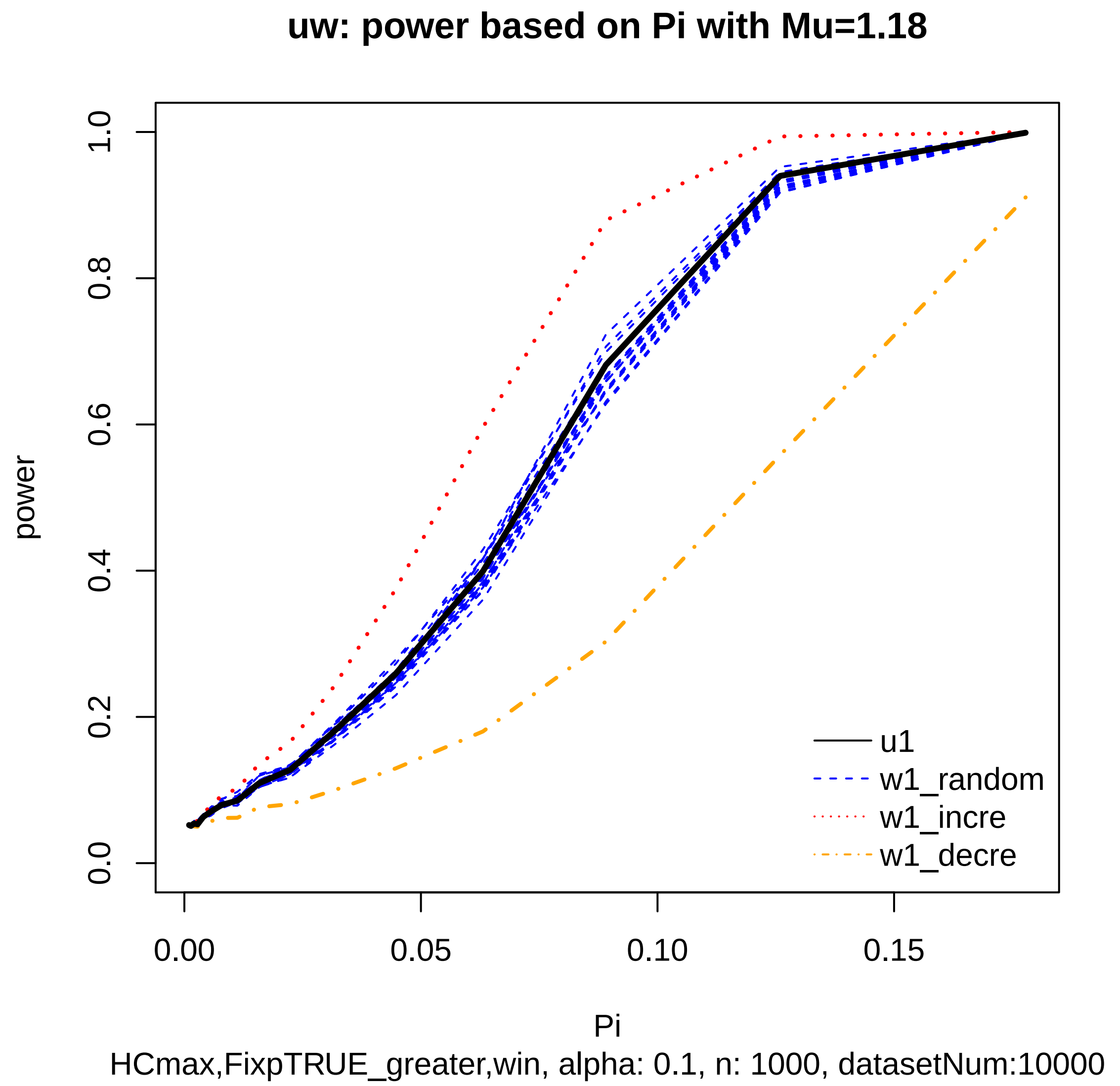
uw: power based on Pi with Mu=1.18
power
1.0
0.8
0.6
0.4
0.2
0.0
0.00
0.05
0.10
0.15
Pi
u1
w1_random
w1_incre
w1_decre
HCmax,FixpTRUE_greater,win, alpha: 0.1, n: 1000, datasetNum:10000

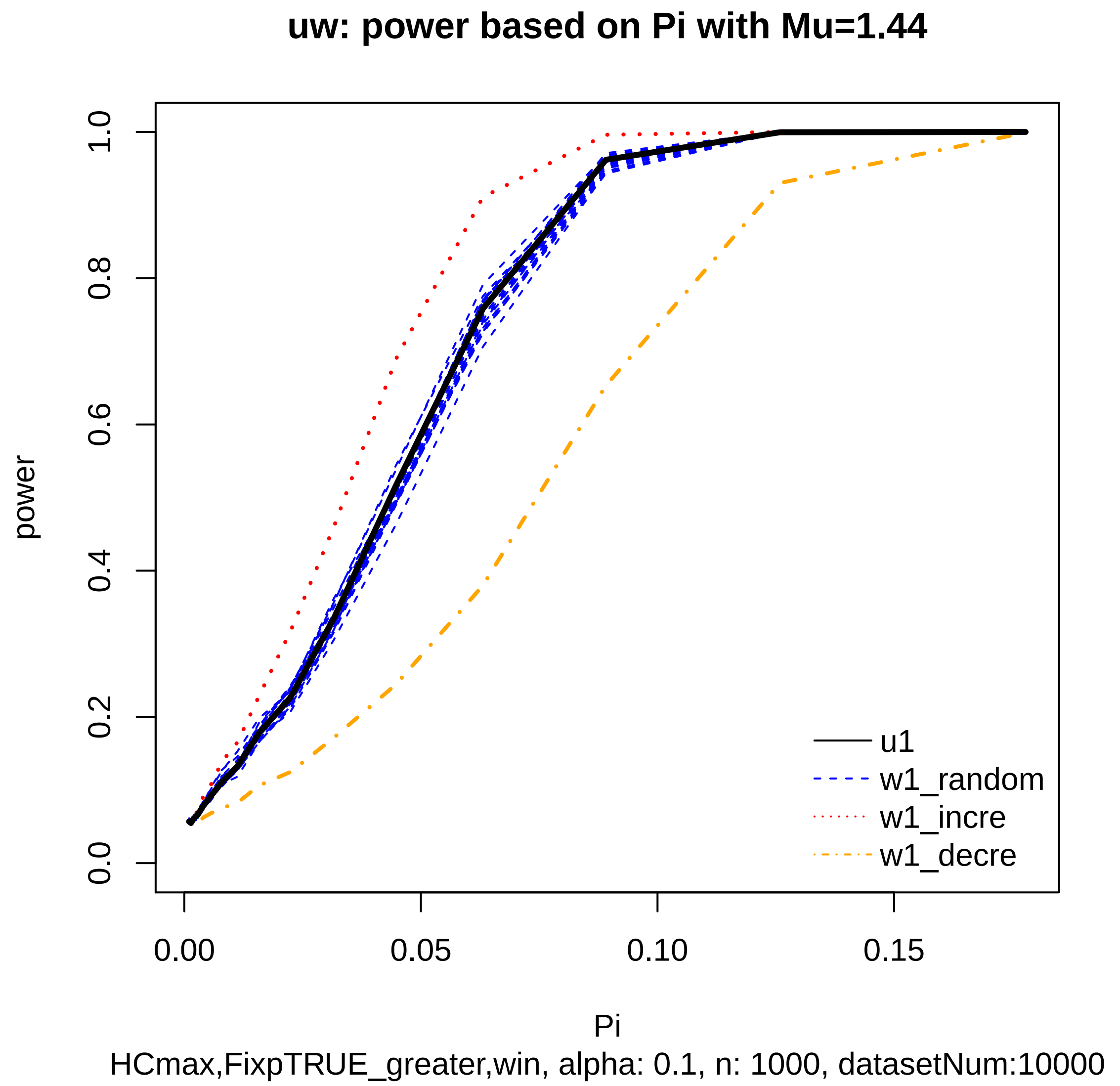
uw: power based on Pi with Mu=1.44
power
0.0
0.2
0.4
0.6
0.8
1.0
0.00
0.05
0.10
0.15
Pi
u1
w1_random
w1_incre
w1_decre
HCmax,FixpTRUE_greater,win, alpha: 0.1, n: 1000, datasetNum:10000

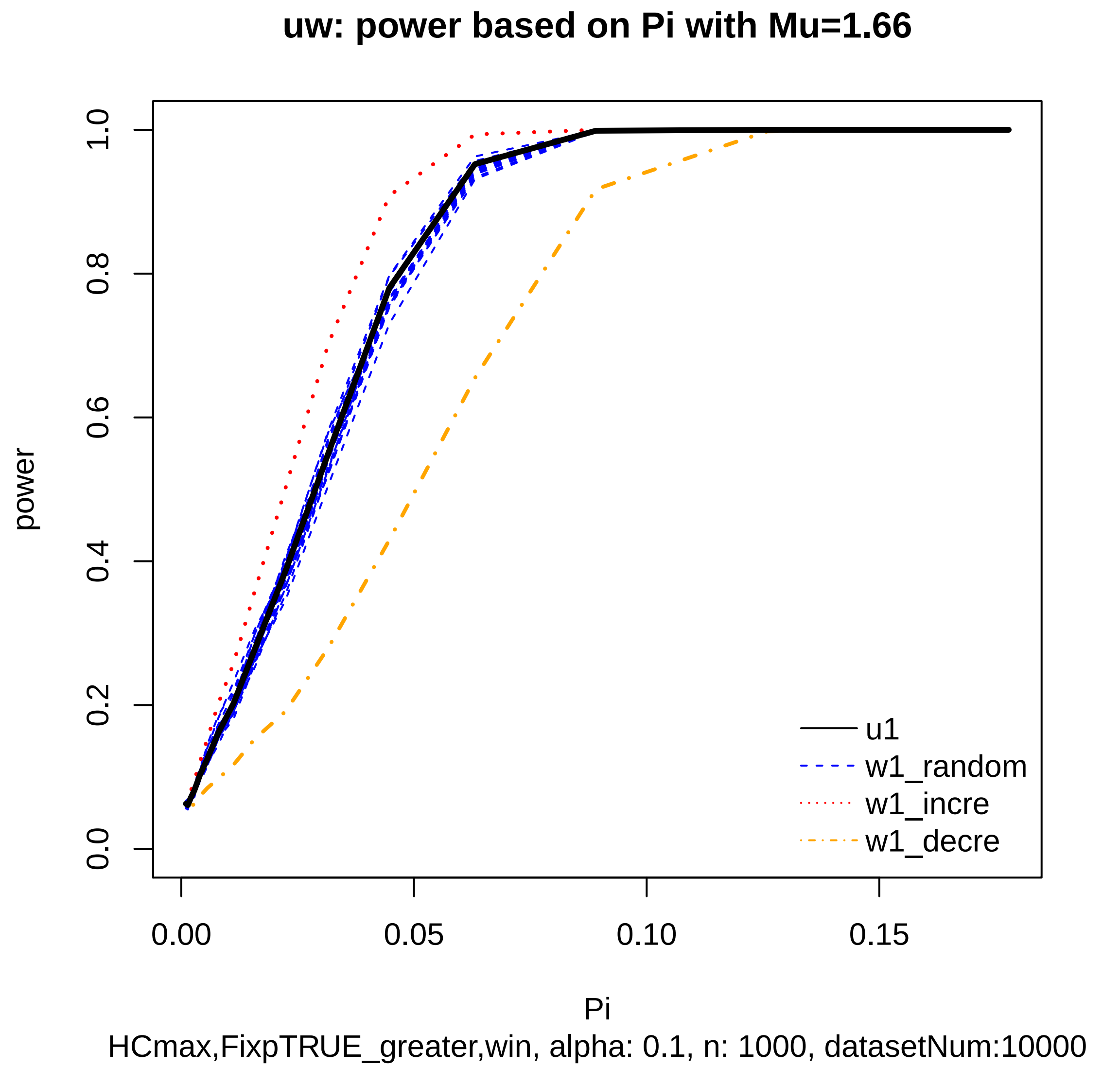

uw: power based on Pi with Mu=1.66
power
0.0
0.2
0.4
0.6
0.8
1.0
0.00
0.05
0.10
0.15
Pi
u1
w1_random
w1_incre
w1_decre
HCmax,FixpTRUE_greater,win, alpha: 0.1, n: 1000, datasetNum:10000

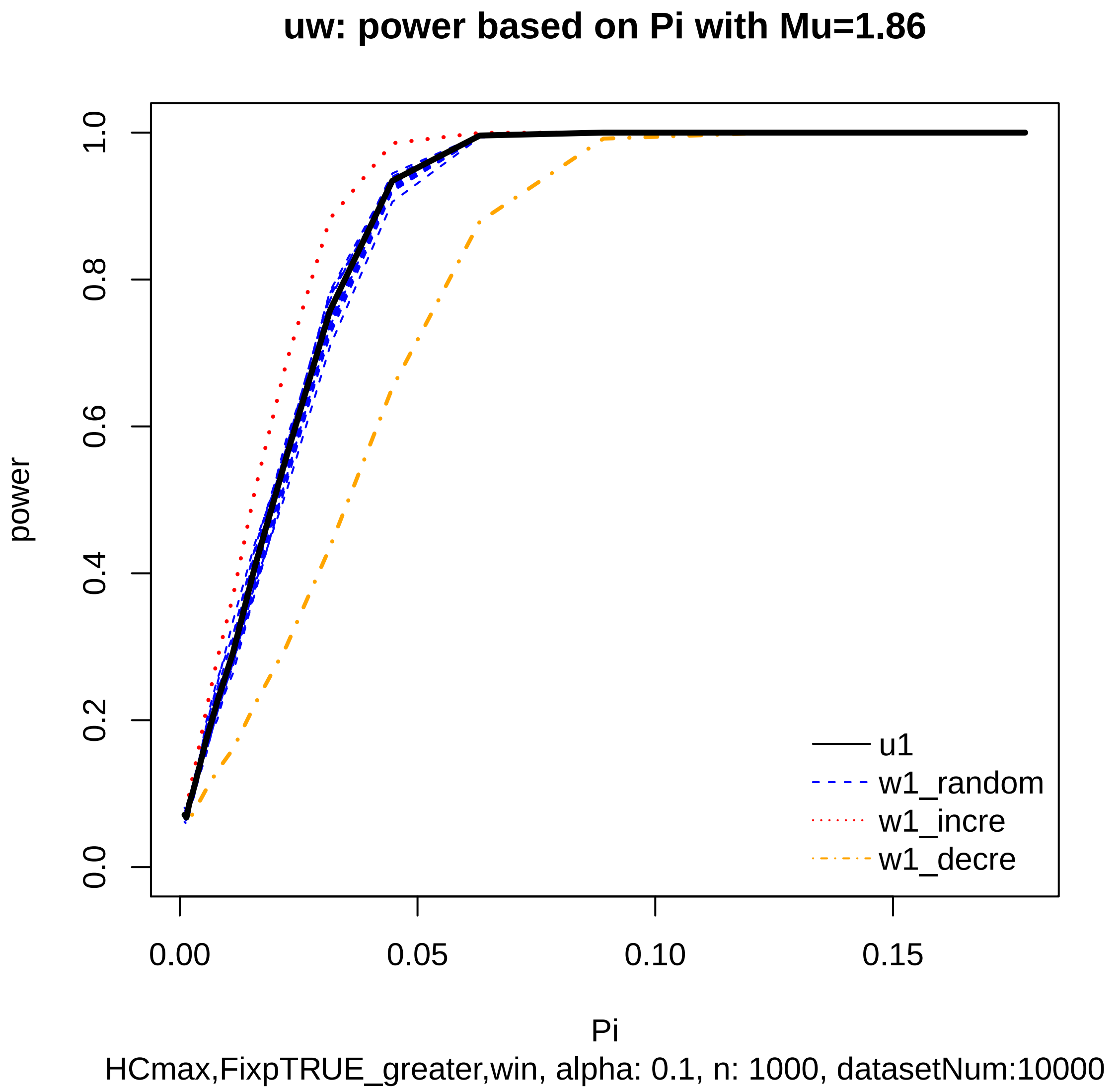
uw: power based on Pi with Mu=1.86
power
1.0
0.8
0.6
0.4
0.2
0.0
0.00
0.05
0.10
0.15
Pi
u1
w1_random
w1_incre
w1_decre
HCmax,FixpTRUE_greater,win, alpha: 0.1, n: 1000, datasetNum:10000

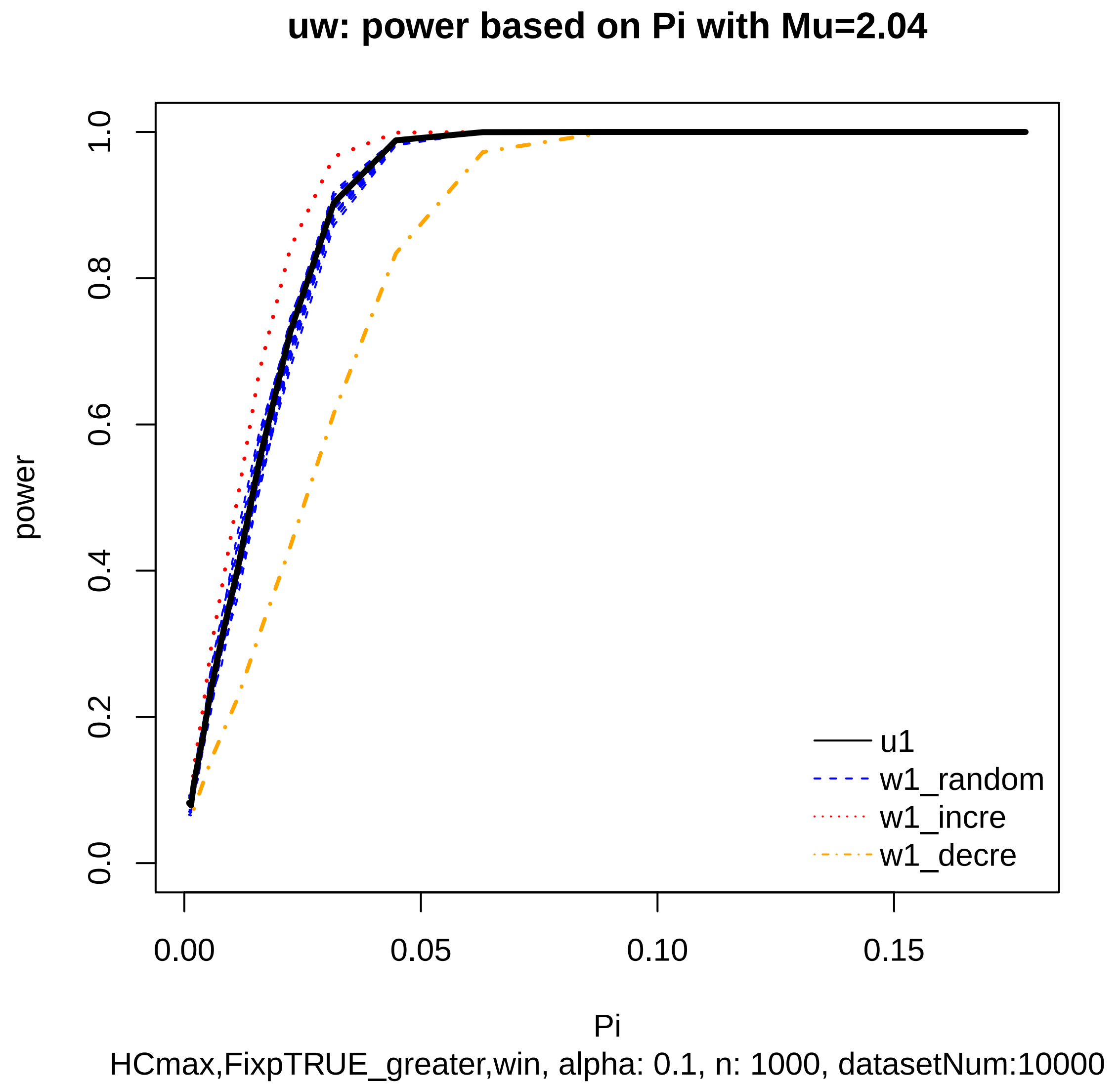
uw: power based on Pi with Mu=2.04
power
0.0
0.2
0.4
0.6
0.8
1.0
0.00
0.05
0.10
0.15
Pi
u1
w1_random
w1_incre
w1_decre
HCmax,FixpTRUE_greater,win, alpha: 0.1, n: 1000, datasetNum:10000

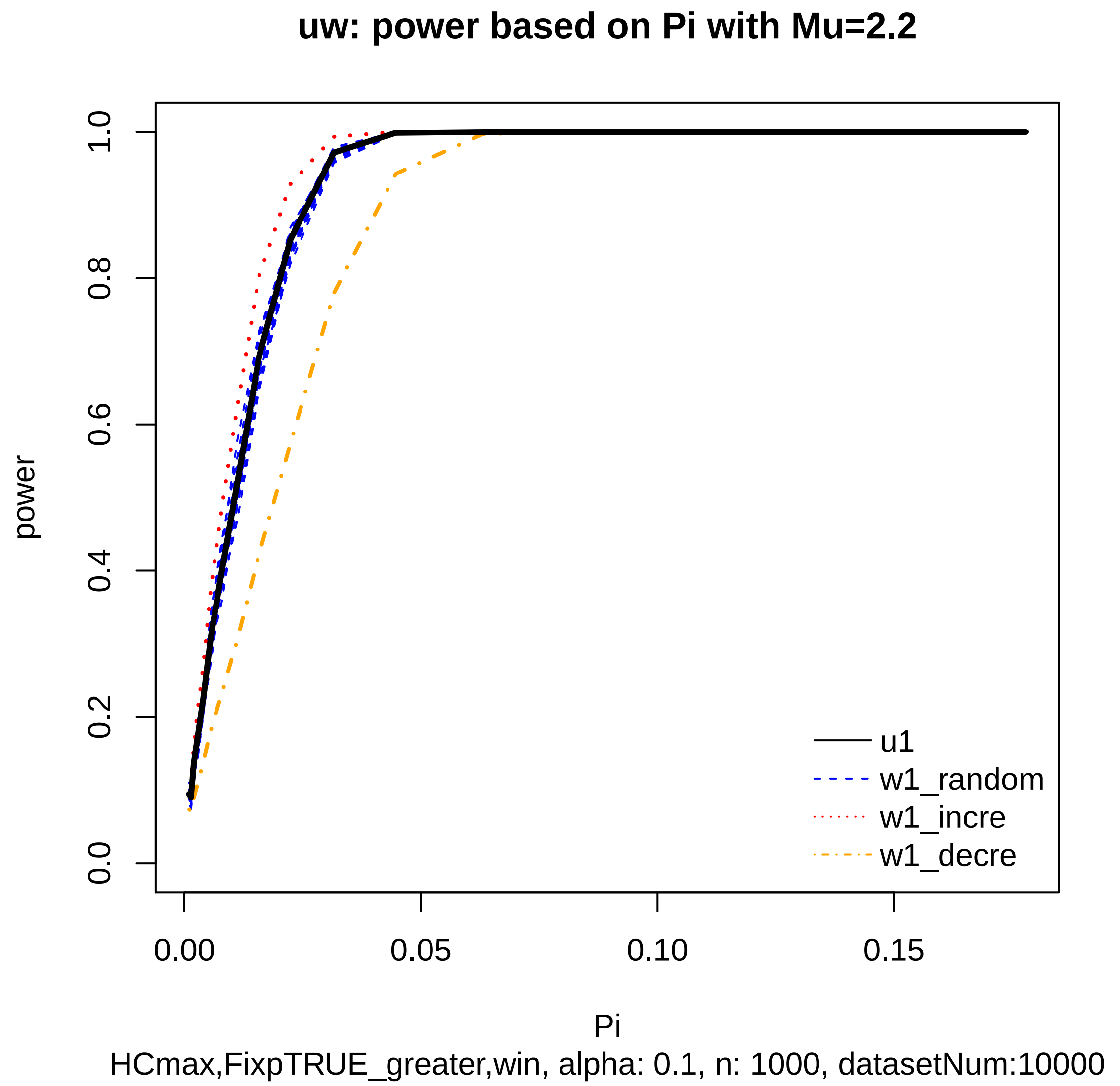
uw: power based on Pi with Mu=2.2
power
0.0
0.2
0.4
0.6
0.8
1.0
0.00
0.05
0.10
0.15
Pi
u1
w1_random
w1_incre
w1_decre
HCmax,FixpTRUE_greater,win, alpha: 0.1, n: 1000, datasetNum:10000

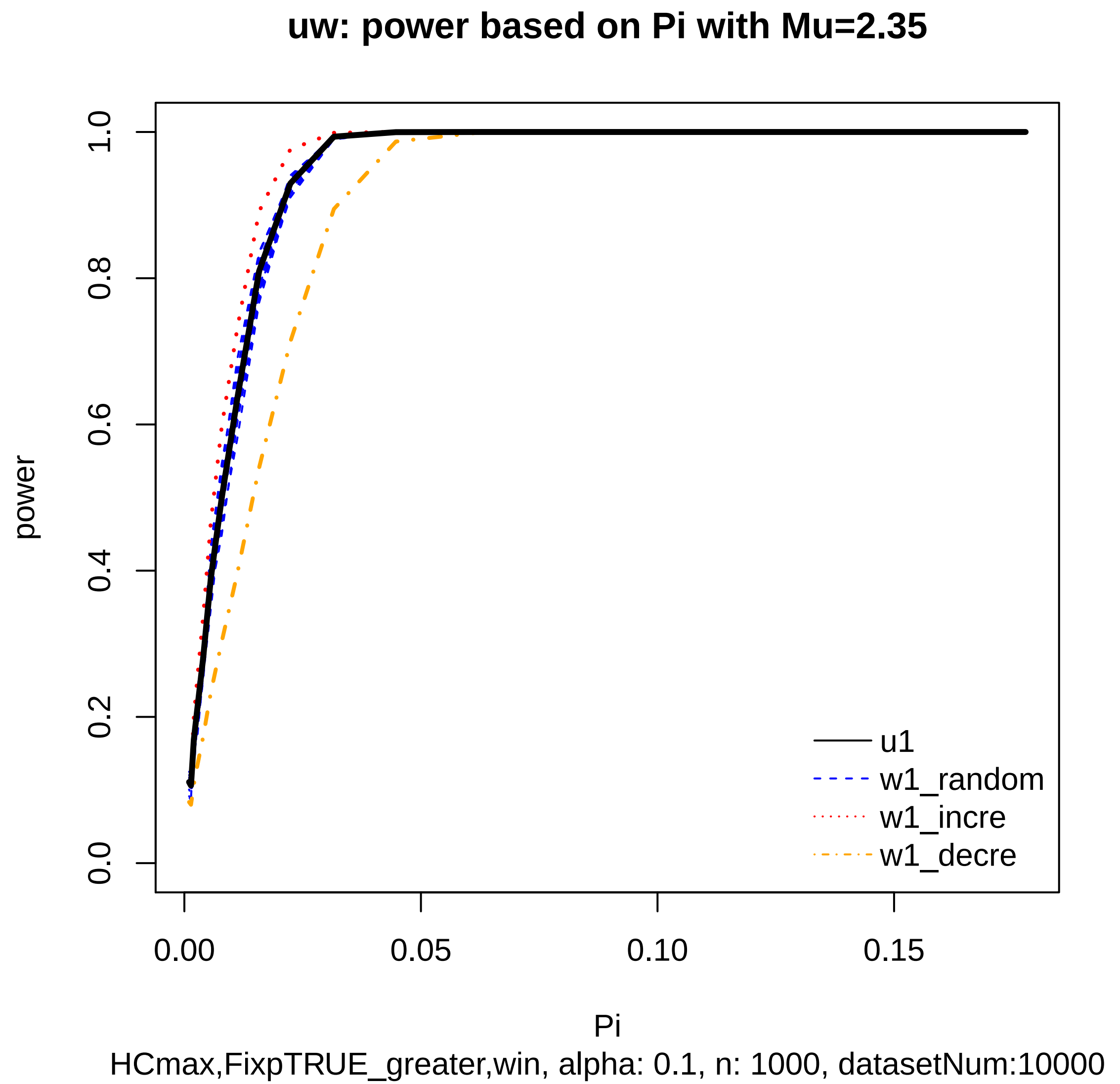
uw: power based on Pi with Mu=2.35
power
0.0
0.2
0.4
0.6
0.8
1.0
0.00
0.05
0.10
0.15
Pi
u1
w1_random
w1_incre
w1_decre
HCmax,FixpTRUE_greater,win, alpha: 0.1, n: 1000, datasetNum:10000

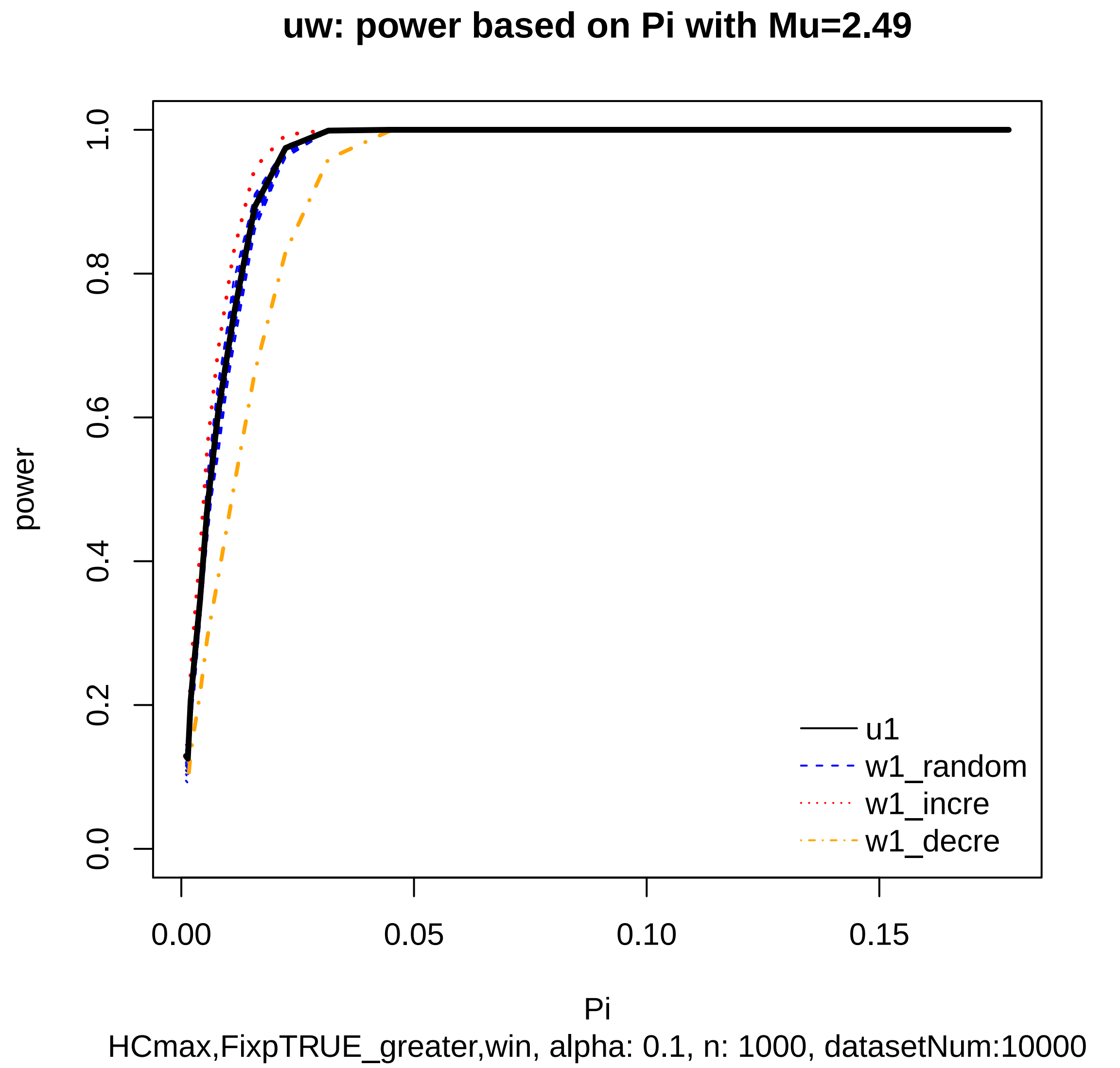

uw: power based on Pi with Mu=2.49
power
0.0
0.2
0.4
0.6
0.8
1.0
0.00
0.05
0.10
0.15
Pi
u1
w1_random
w1_incre
w1_decre
HCmax,FixpTRUE_greater,win, alpha: 0.1, n: 1000, datasetNum:10000

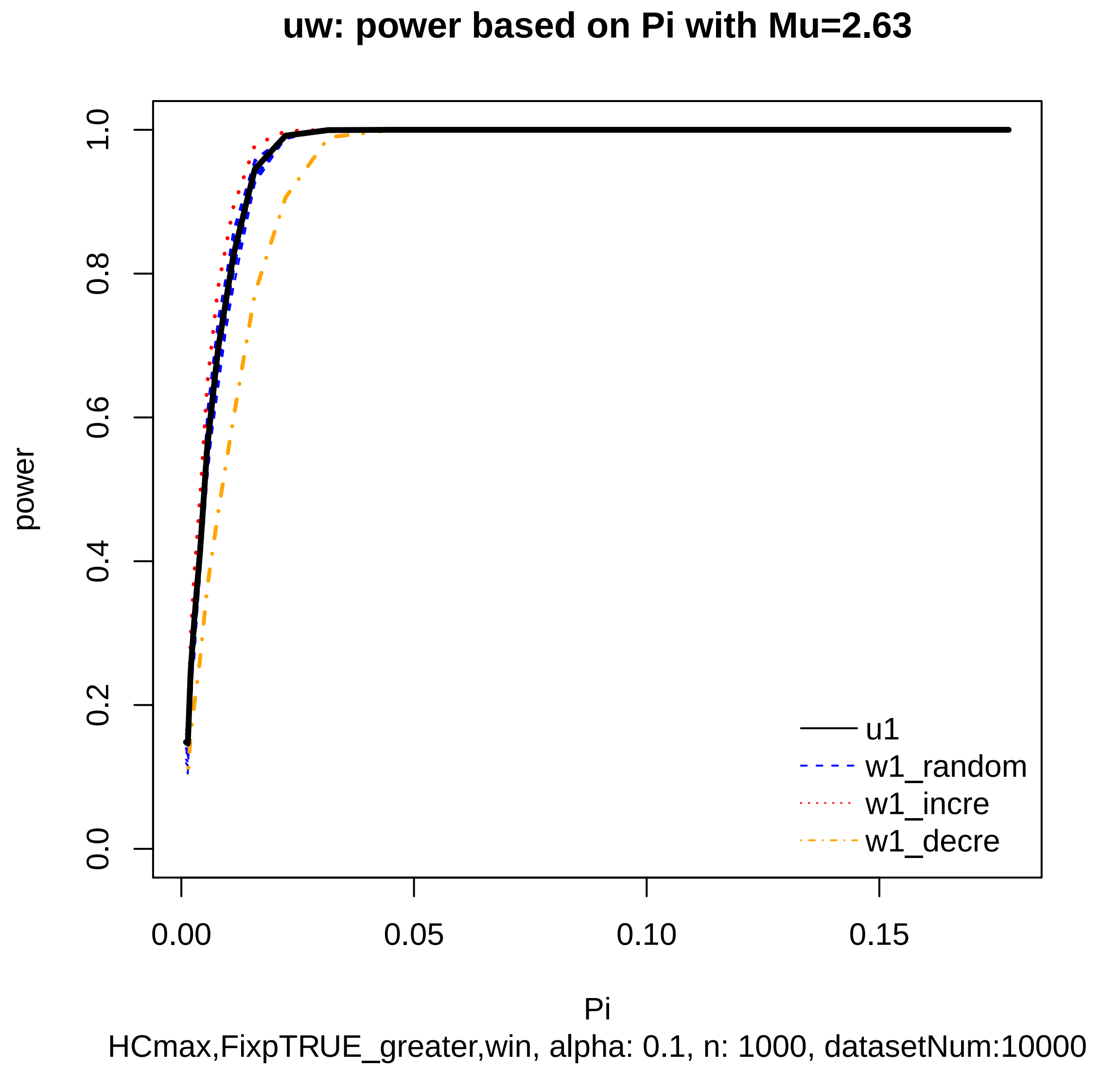
uw: power based on Pi with Mu=2.63
power
0.0
0.2
0.4
0.6
0.8
1.0
0.00
0.05
0.10
0.15
Pi
u1
w1_random
w1_incre
w1_decre
HCmax,FixpTRUE_greater,win, alpha: 0.1, n: 1000, datasetNum:10000

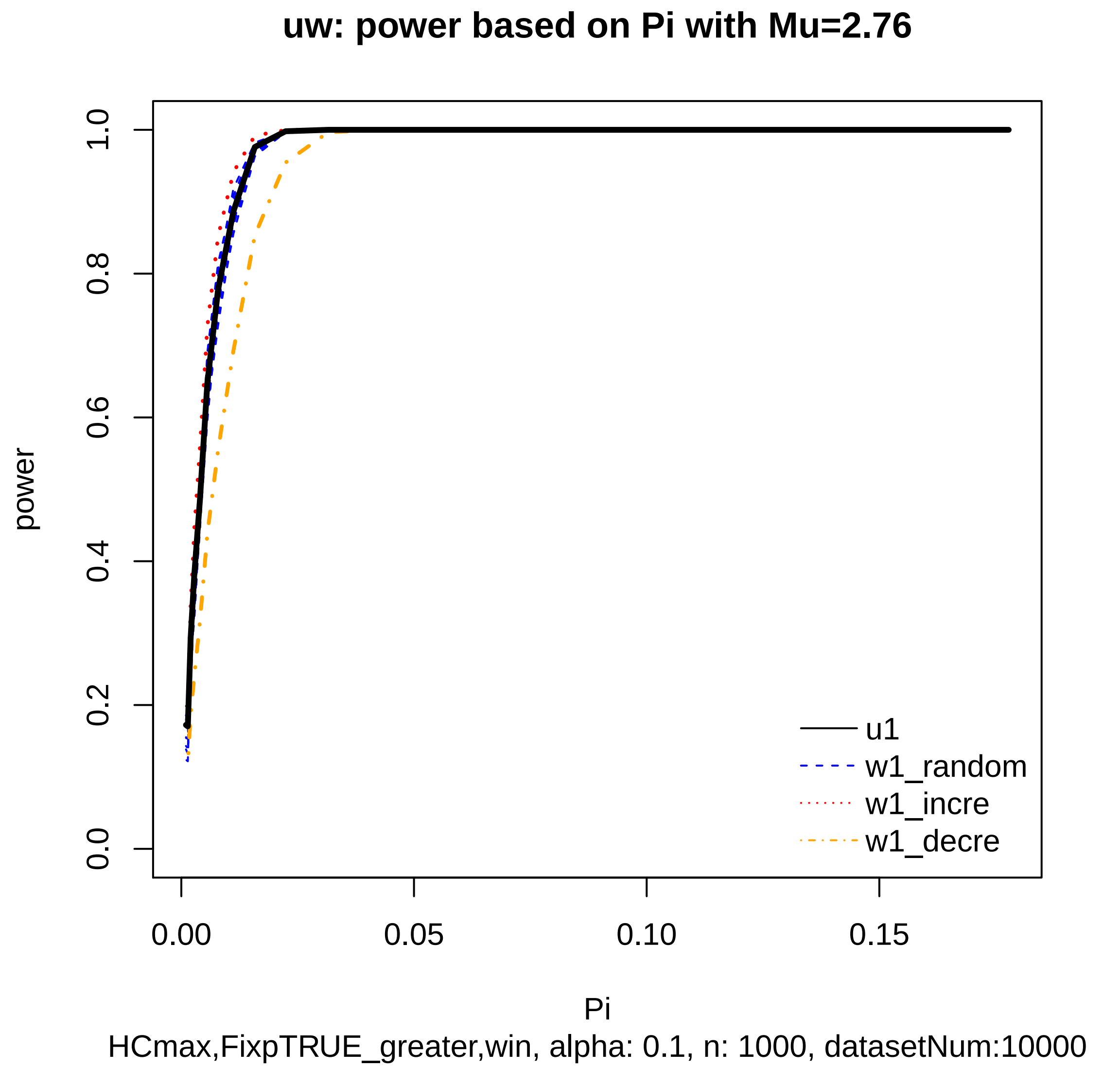
uw: power based on Pi with Mu=2.76
power
1.0
0.8
0.6
0.4
0.2
0.0
0.00
0.05
0.10
0.15
Pi
u1
w1_random
w1_incre
w1_decre
HCmax,FixpTRUE_greater,win, alpha: 0.1, n: 1000, datasetNum:10000

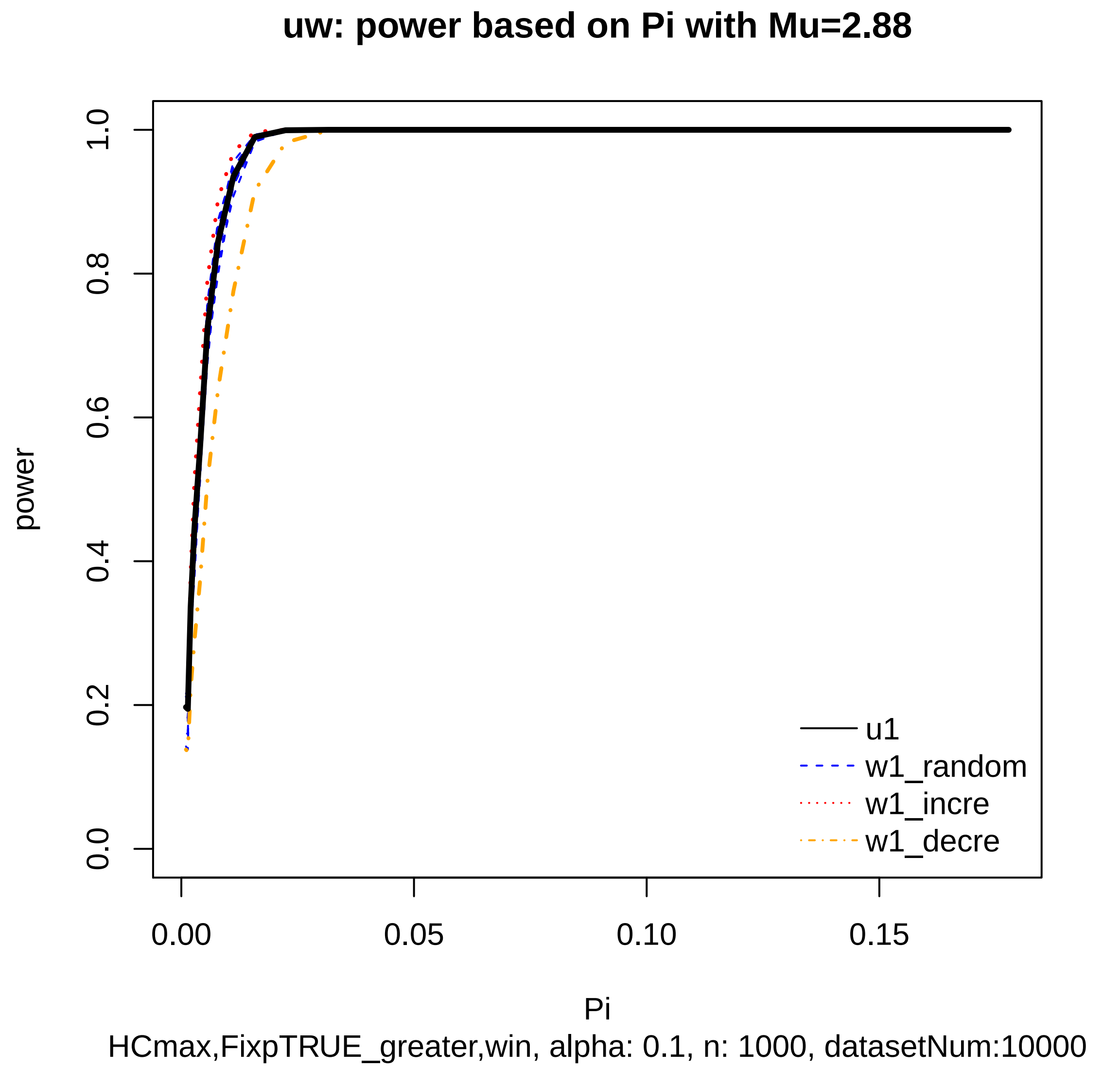
uw: power based on Pi with Mu=2.88
power
1.0
0.8
0.6
0.4
0.2
0.0
0.00
0.05
0.10
0.15
Pi
u1
w1_random
w1_incre
w1_decre
HCmax,FixpTRUE_greater,win, alpha: 0.1, n: 1000, datasetNum:10000

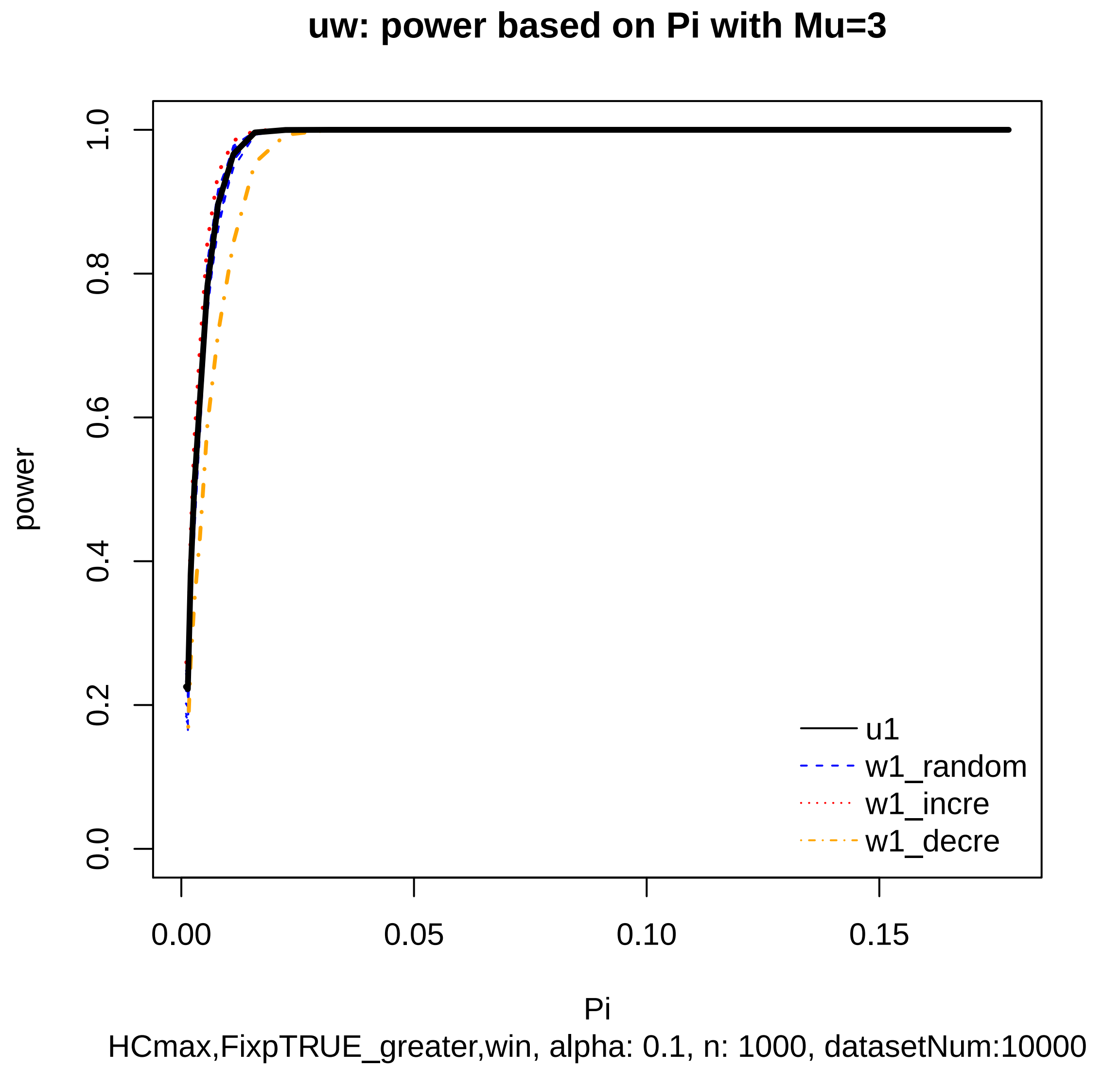
uw: power based on Pi with Mu=3
power
1.0
0.8
0.6
0.4
0.2
0.0
0.00
0.05
0.10
0.15
Pi
u1
w1_random
w1_incre
w1_decre
HCmax,FixpTRUE_greater,win, alpha: 0.1, n: 1000, datasetNum:10000

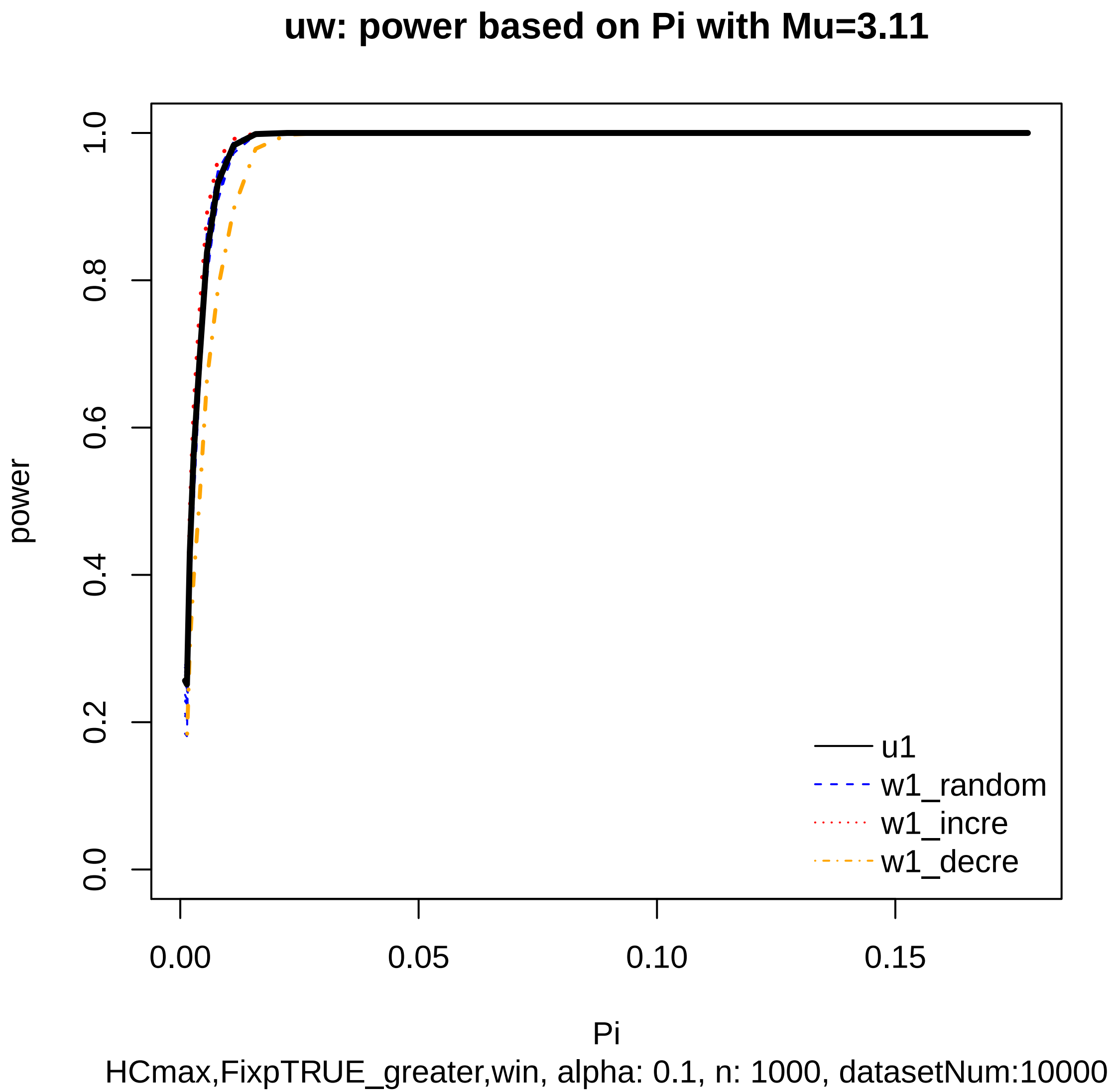
uw: power based on Pi with Mu=3.11
power
0.0
0.2
0.4
0.6
0.8
1.0
0.00
0.05
0.10
0.15
Pi
u1
w1_random
w1_incre
w1_decre
HCmax,FixpTRUE_greater,win, alpha: 0.1, n: 1000, datasetNum:10000

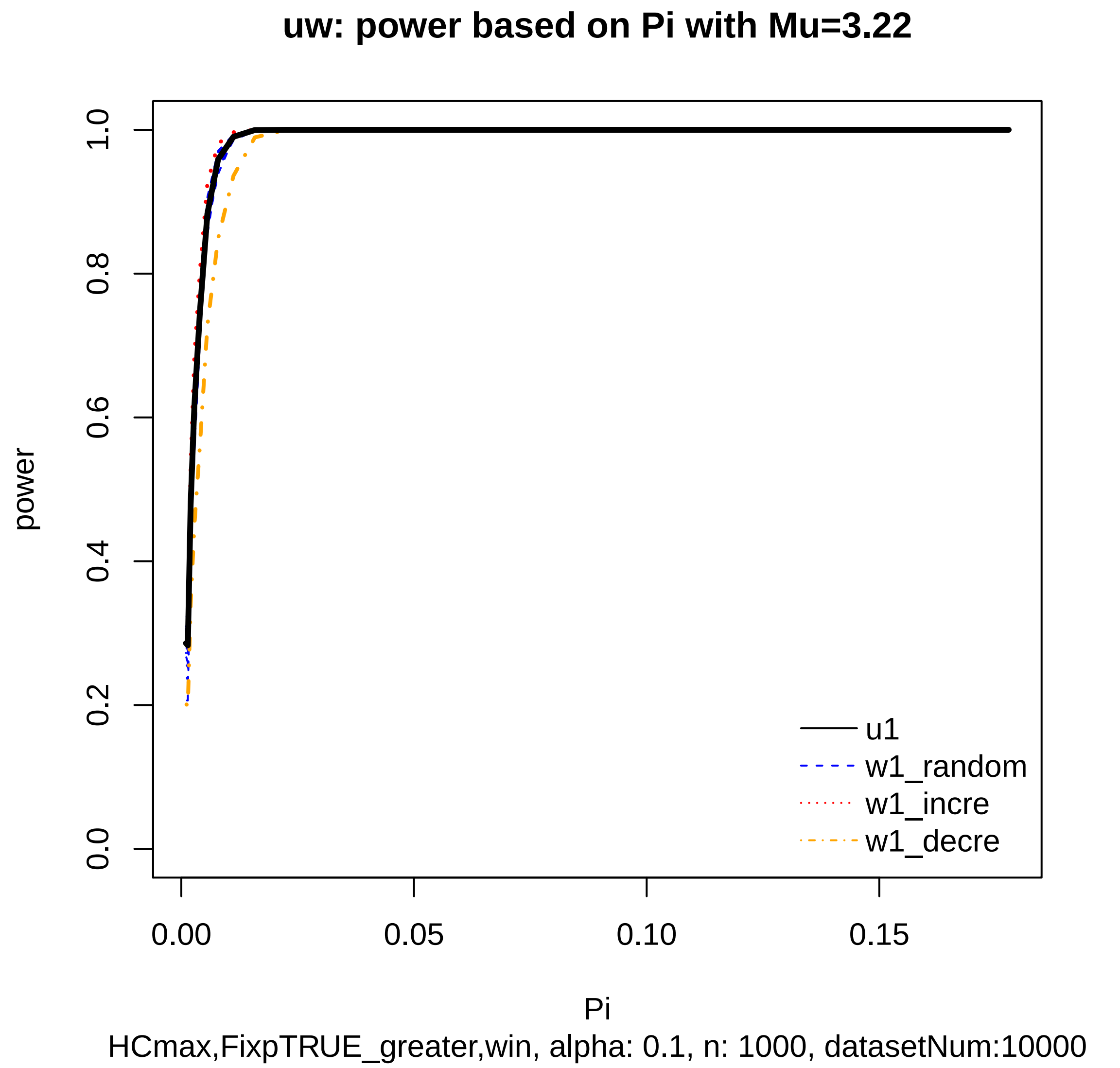
uw: power based on Pi with Mu=3.22
power
0.0
0.2
0.4
0.6
0.8
1.0
0.00
0.05
0.10
0.15
Pi
u1
w1_random
w1_incre
w1_decre
HCmax,FixpTRUE_greater,win, alpha: 0.1, n: 1000, datasetNum:10000

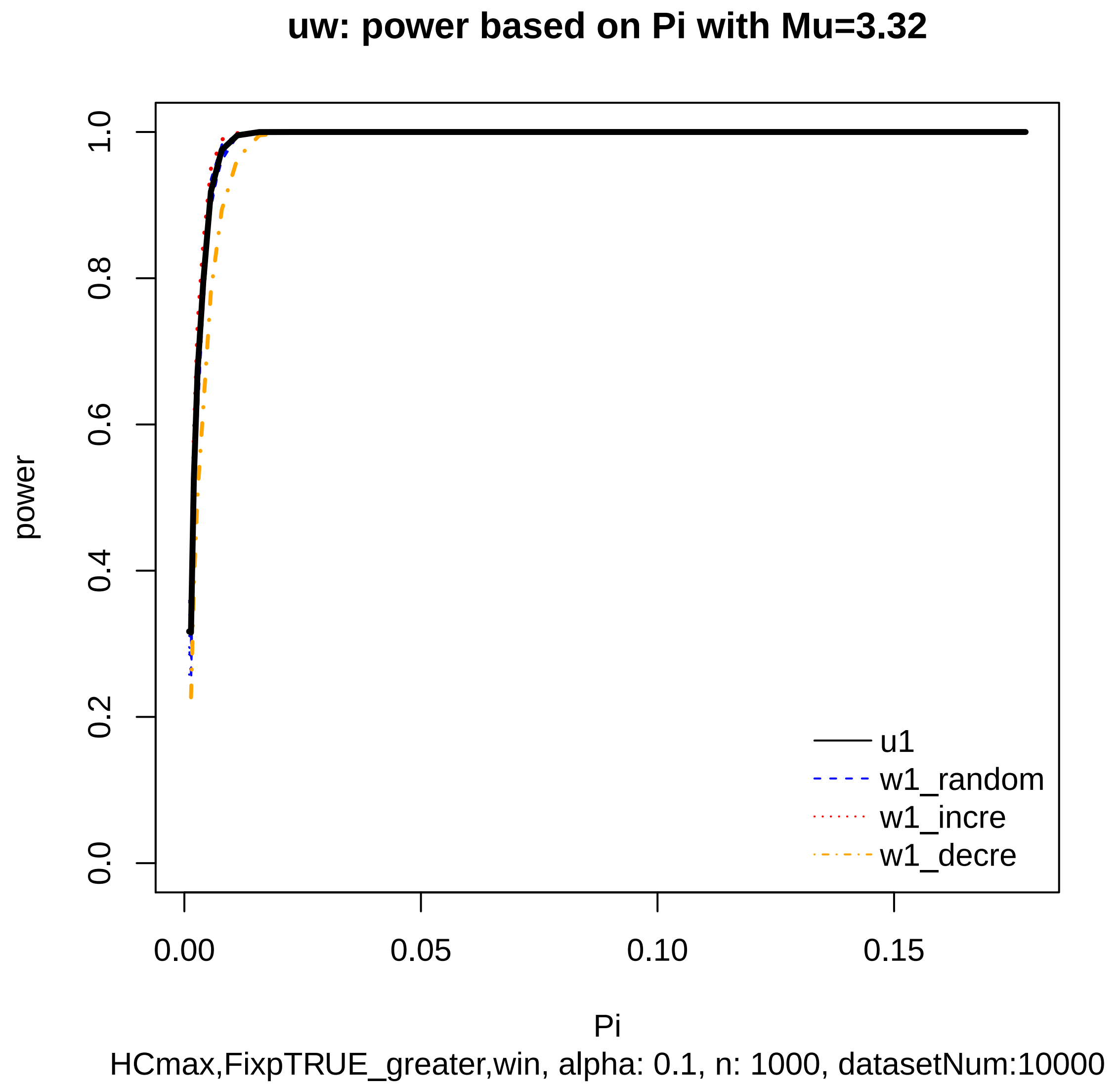
uw: power based on Pi with Mu=3.32
power
1.0
0.8
0.6
0.4
0.2
0.0
0.00
0.05
0.10
0.15
Pi
u1
w1_random
w1_incre
w1_decre
HCmax,FixpTRUE_greater,win, alpha: 0.1, n: 1000, datasetNum:10000

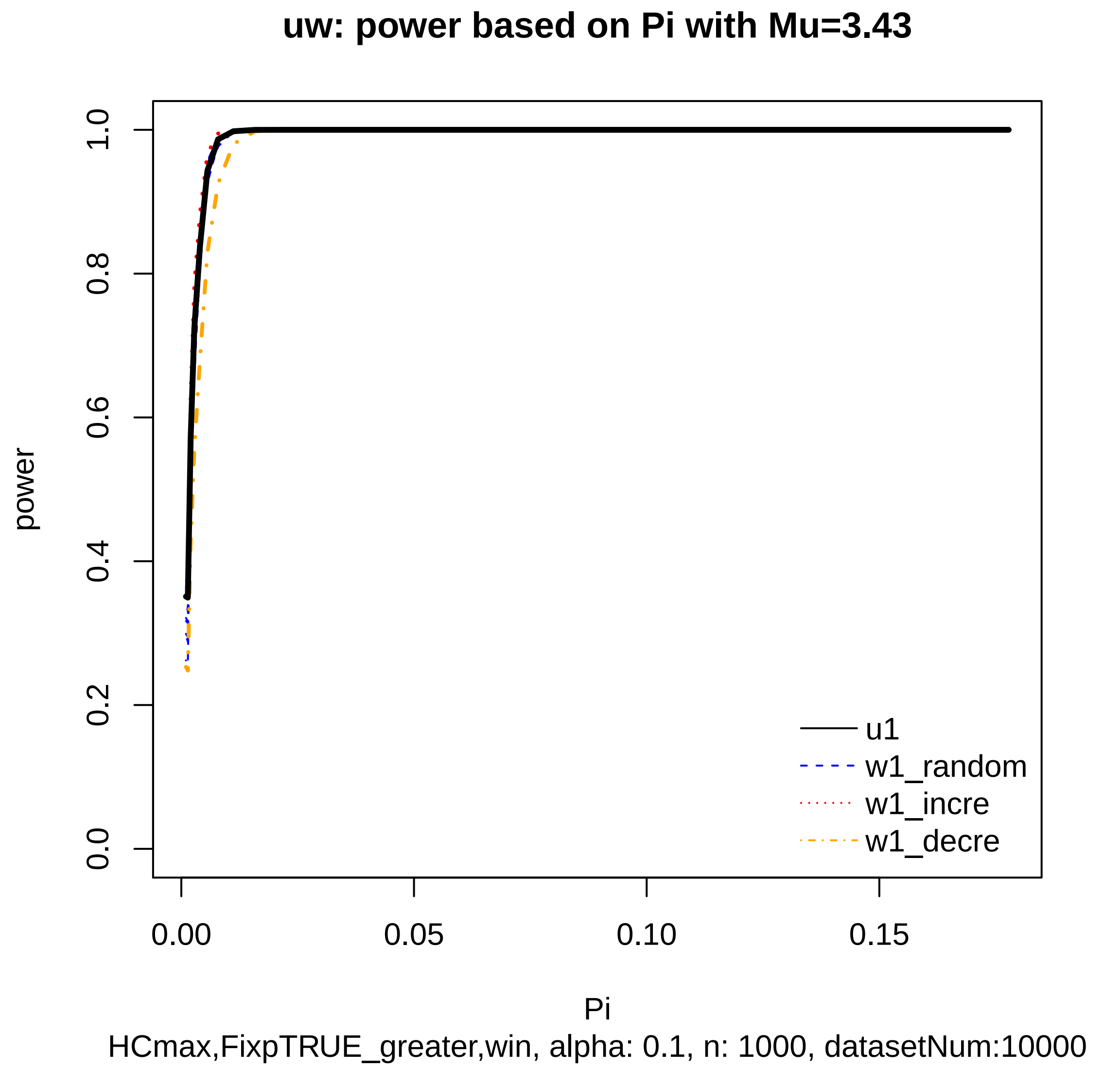
uw: power based on Pi with Mu=3.43
power
1.0
0.8
0.6
0.4
0.2
0.0
0.00
0.05
0.10
0.15
u1
w1_random
w1_incre
w1_decre
Pi
HCmax,FixpTRUE_greater,win, alpha: 0.1, n: 1000, datasetNum:10000

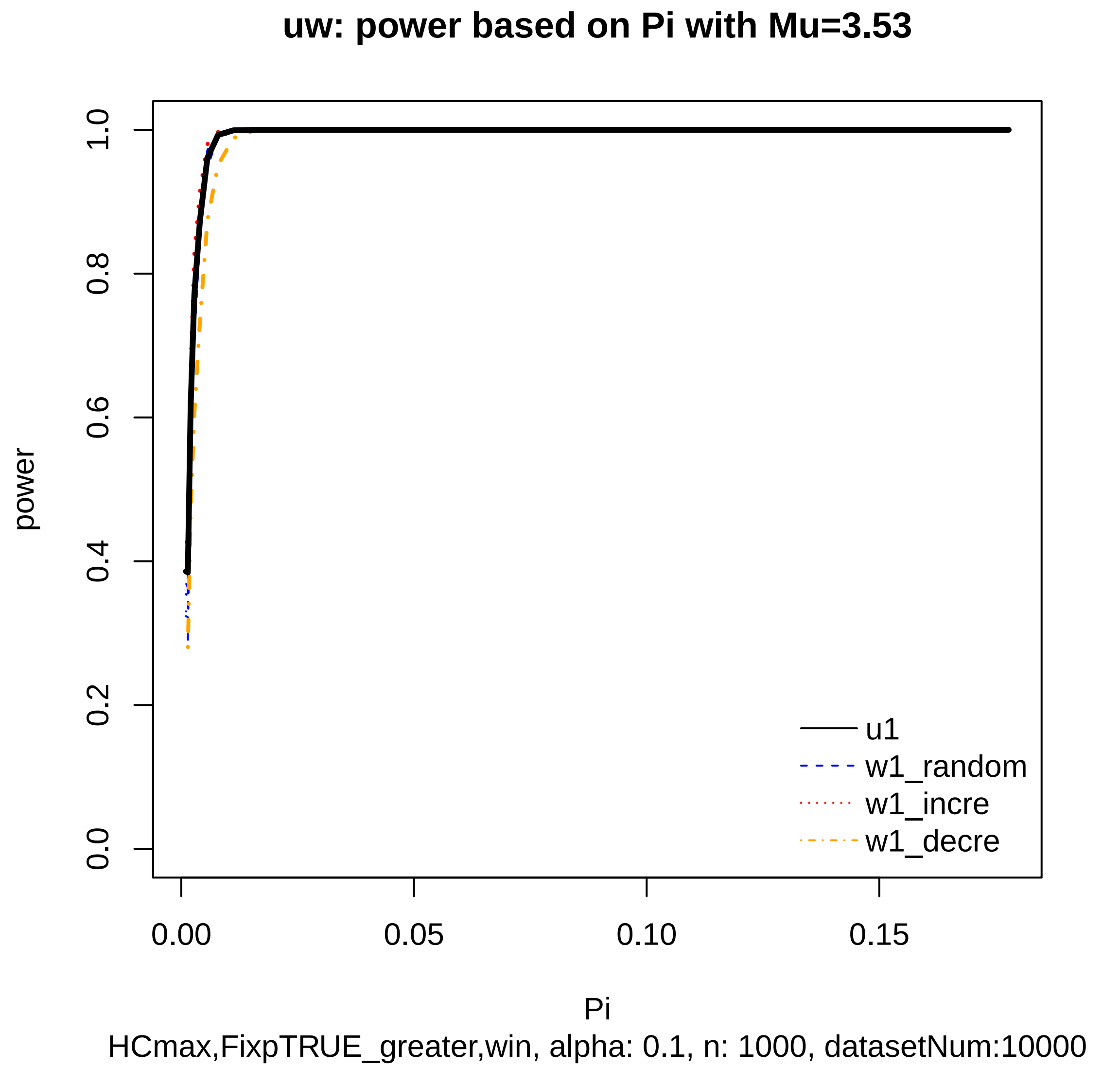
uw: power based on Pi with Mu=3.53
power
1.0
0.8
0.6
0.4
0.2
0.0
0.00
0.05
0.10
0.15
Pi
u1
w1_random
w1_incre
w1_decre
HCmax,FixpTRUE_greater,win, alpha: 0.1, n: 1000, datasetNum:10000

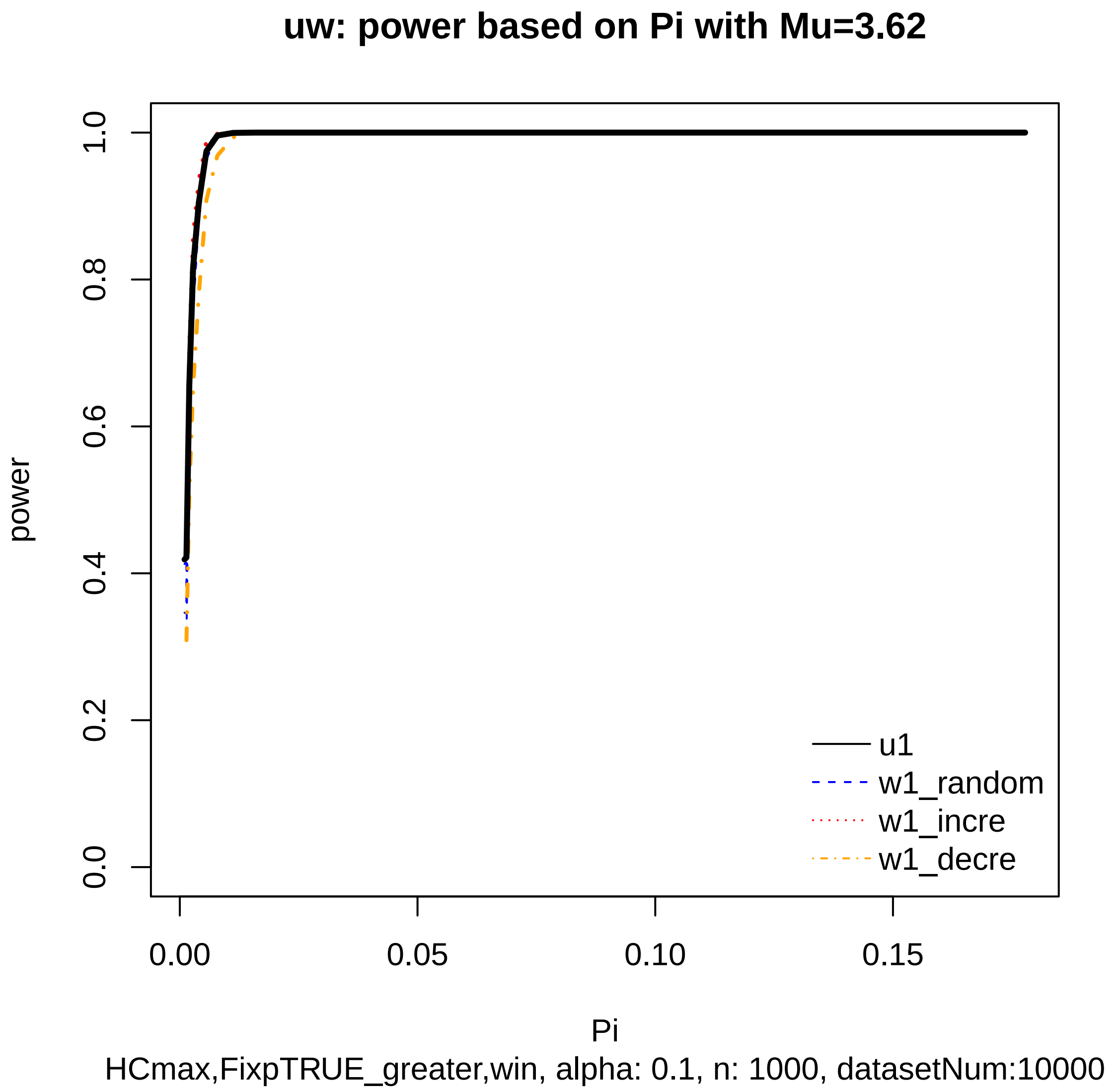

uw: power based on Pi with Mu=3.62
power
1.0
0.8
0.6
0.4
0.2
0.0
0.00
0.05
0.10
0.15
Pi
u1
w1_random
w1_incre
w1_decre
HCmax,FixpTRUE_greater,win, alpha: 0.1, n: 1000, datasetNum:10000

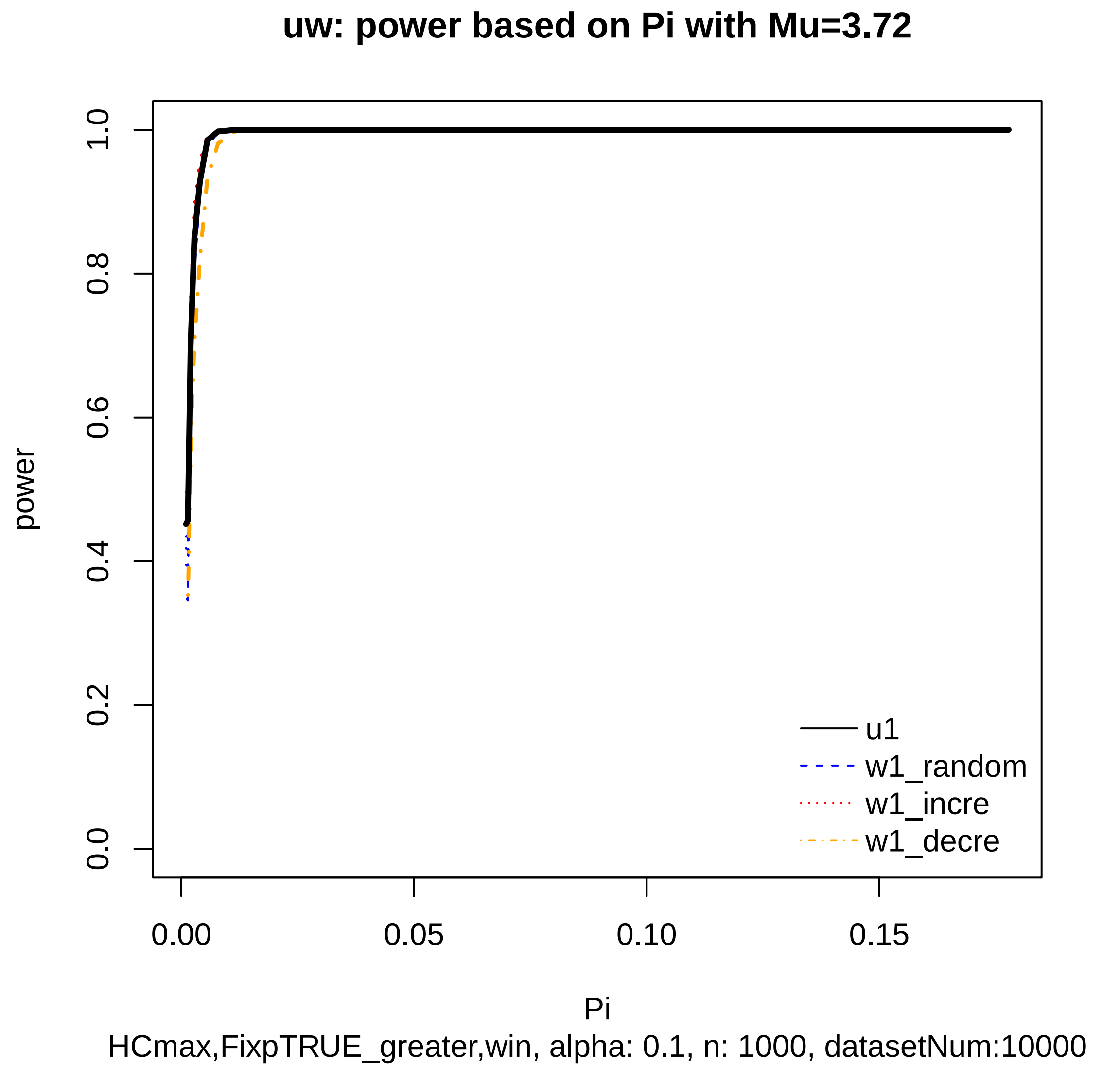
uw: power based on Pi with Mu=3.72
power
0.0
0.2
0.4
0.6
0.8
1.0
0.00
0.05
0.10
0.15
Pi
u1
w1_random
w1_incre
w1_decre
HCmax,FixpTRUE_greater,win, alpha: 0.1, n: 1000, datasetNum:10000

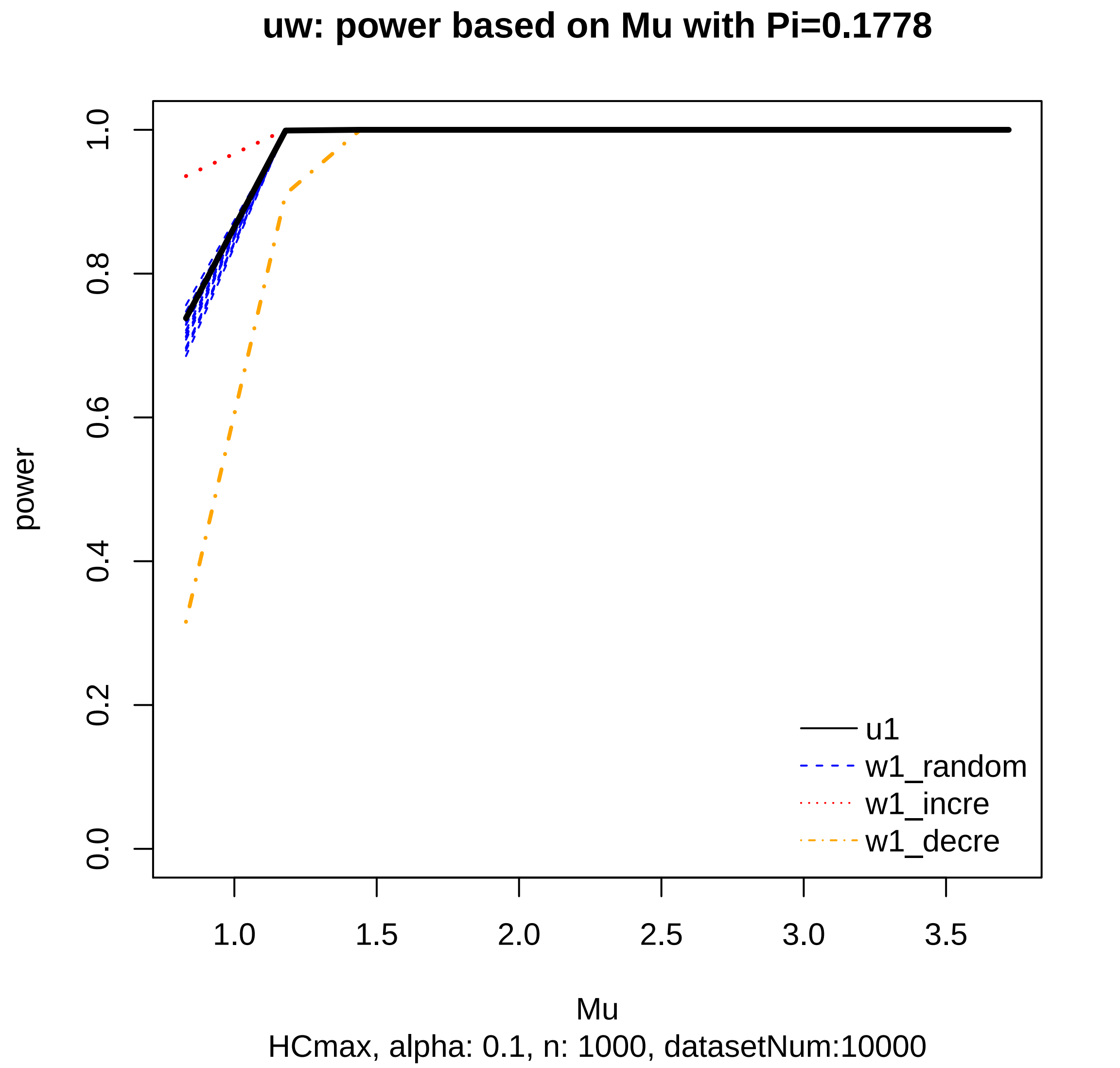
uw: power based on Mu with Pi=0.1778
power
0.0
0.2
0.4
0.6
0.8
1.0
1.0
1.5
2.0
2.5
3.0
3.5
Mu
HCmax, alpha: 0.1, n: 1000, datasetNum:10000
u1
w1_random
w1_incre
w1_decre

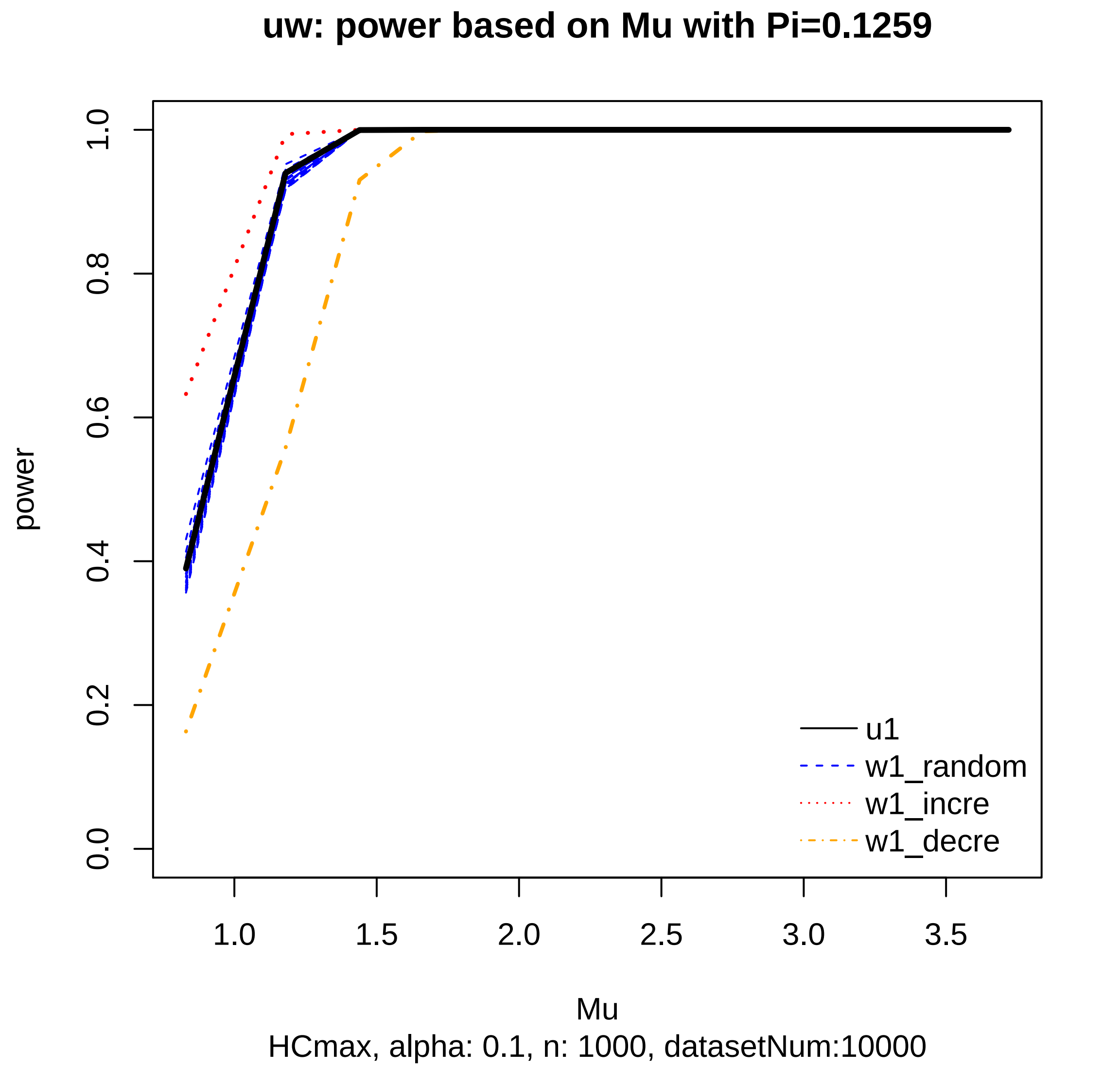

uw: power based on Mu with Pi=0.1259
power
0.0
0.2
0.4
0.6
0.8
1.0
1.0
1.5
2.0
2.5
3.0
3.5
Mu
HCmax, alpha: 0.1, n: 1000, datasetNum:10000
u1
w1_random
w1_incre
w1_decre

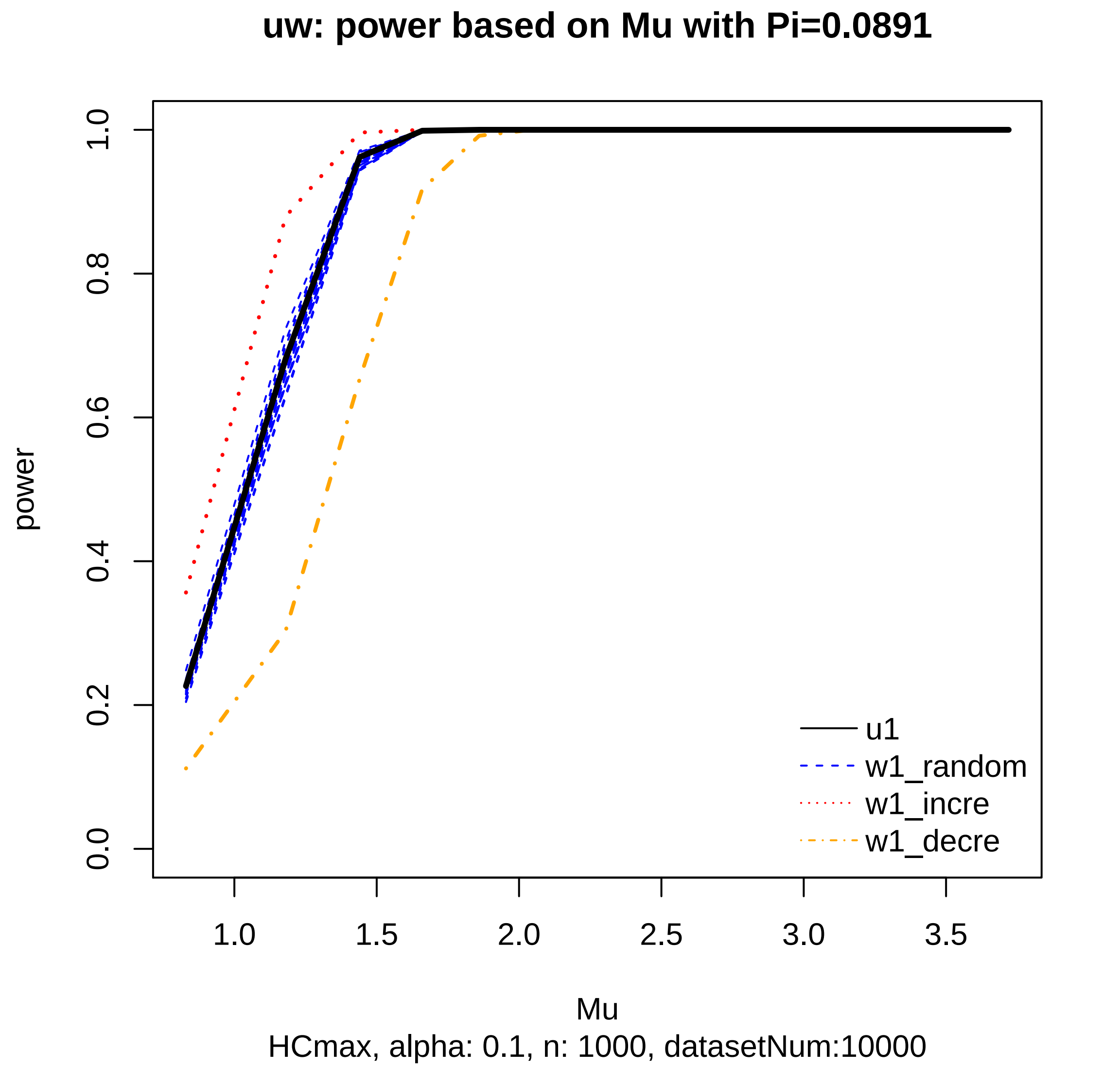
uw: power based on Mu with Pi=0.0891
power
0.0
0.2
0.4
0.6
0.8
1.0
1.0
1.5
2.0
2.5
3.0
3.5
Mu
u1
w1_random
w1_incre
w1_decre
HCmax, alpha: 0.1, n: 1000, datasetNum:10000

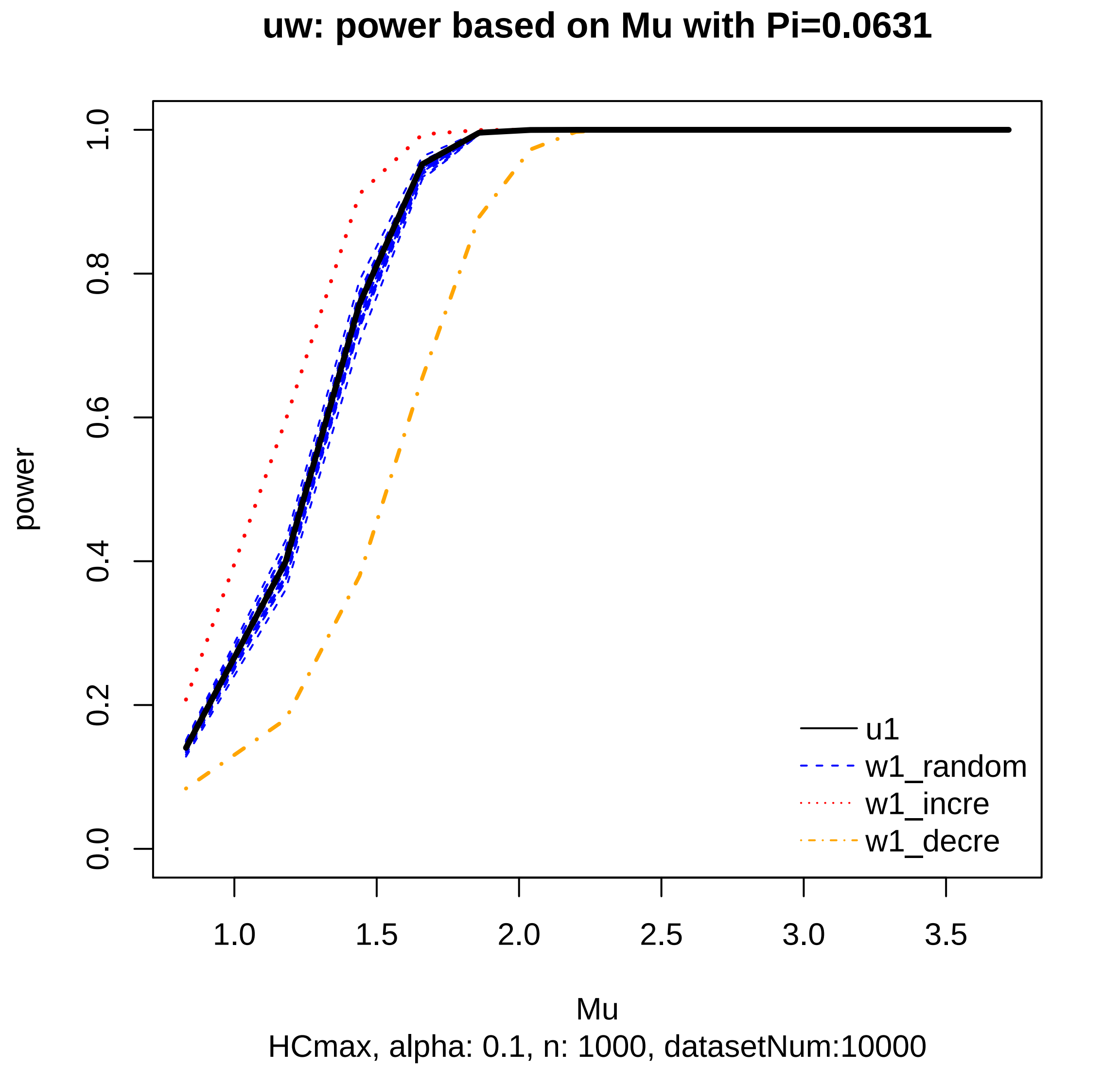
uw: power based on Mu with Pi=0.0631
power
0.0
0.2
0.4
0.6
0.8
1.0
1.0
1.5
2.0
2.5
3.0
3.5
Mu
HCmax, alpha: 0.1, n: 1000, datasetNum:10000
u1
w1_random
w1_incre
w1_decre

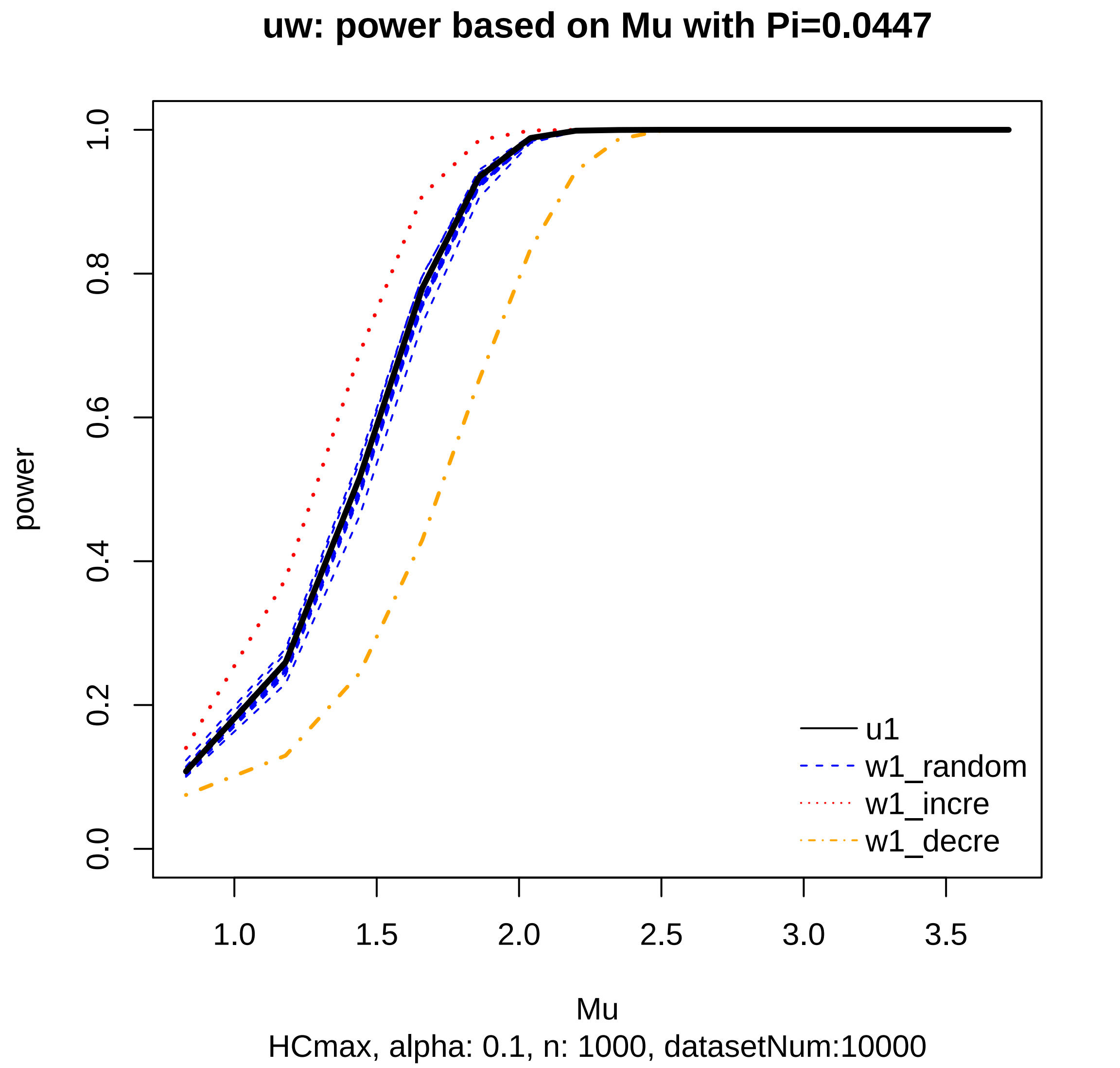
uw: power based on Mu with Pi=0.0447
power
0.0
0.2
0.4
0.6
0.8
1.0
1.0
1.5
2.0
2.5
3.0
3.5
Mu
HCmax, alpha: 0.1, n: 1000, datasetNum:10000
u1
w1_random
w1_incre
w1_decre

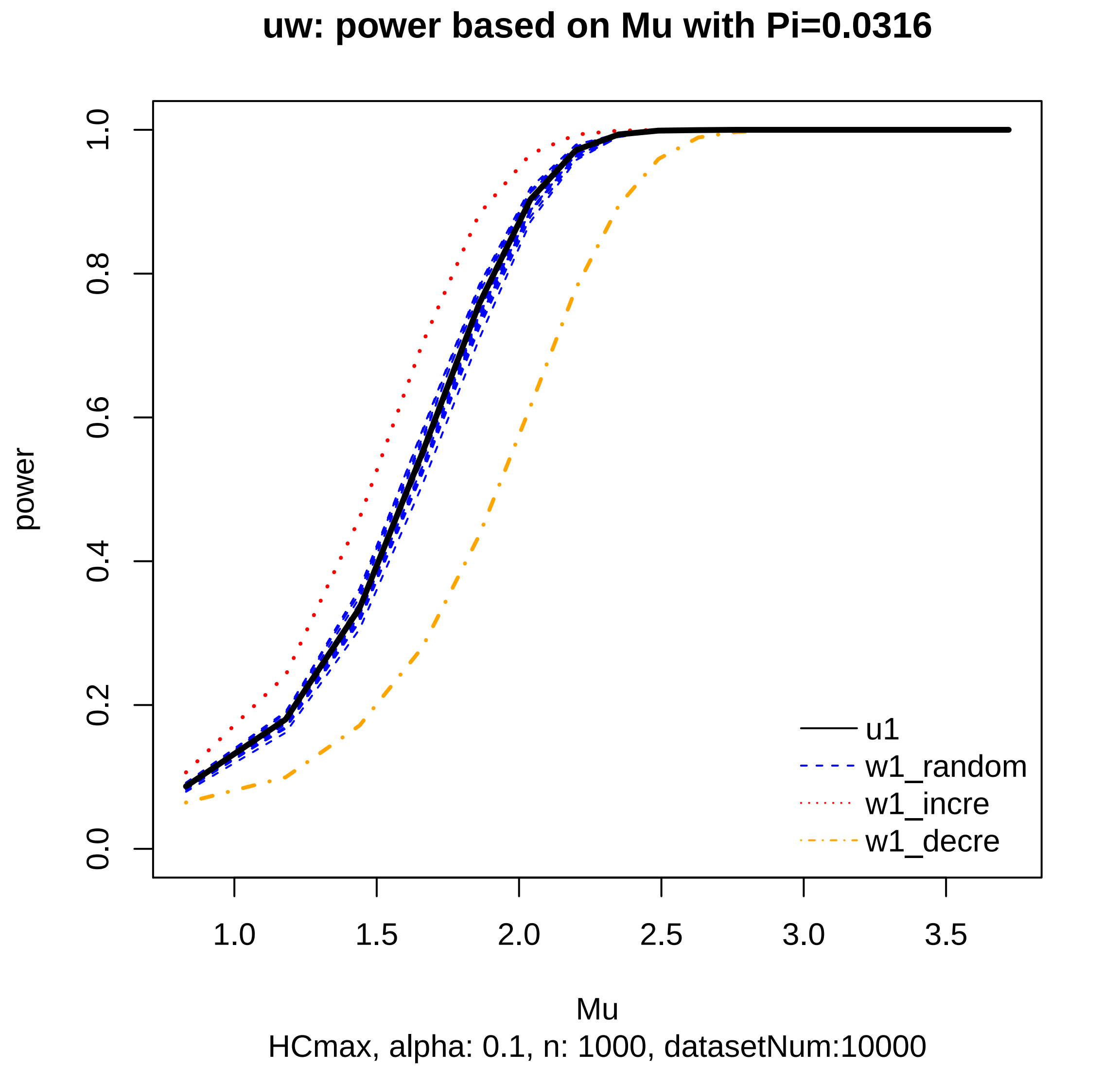
uw: power based on Mu with Pi=0.0316
power
0.0
0.2
0.4
0.6
0.8
1.0
1.0
1.5
2.0
2.5
3.0
3.5
Mu
HCmax, alpha: 0.1, n: 1000, datasetNum:10000
u1
w1_random
w1_incre
w1_decre

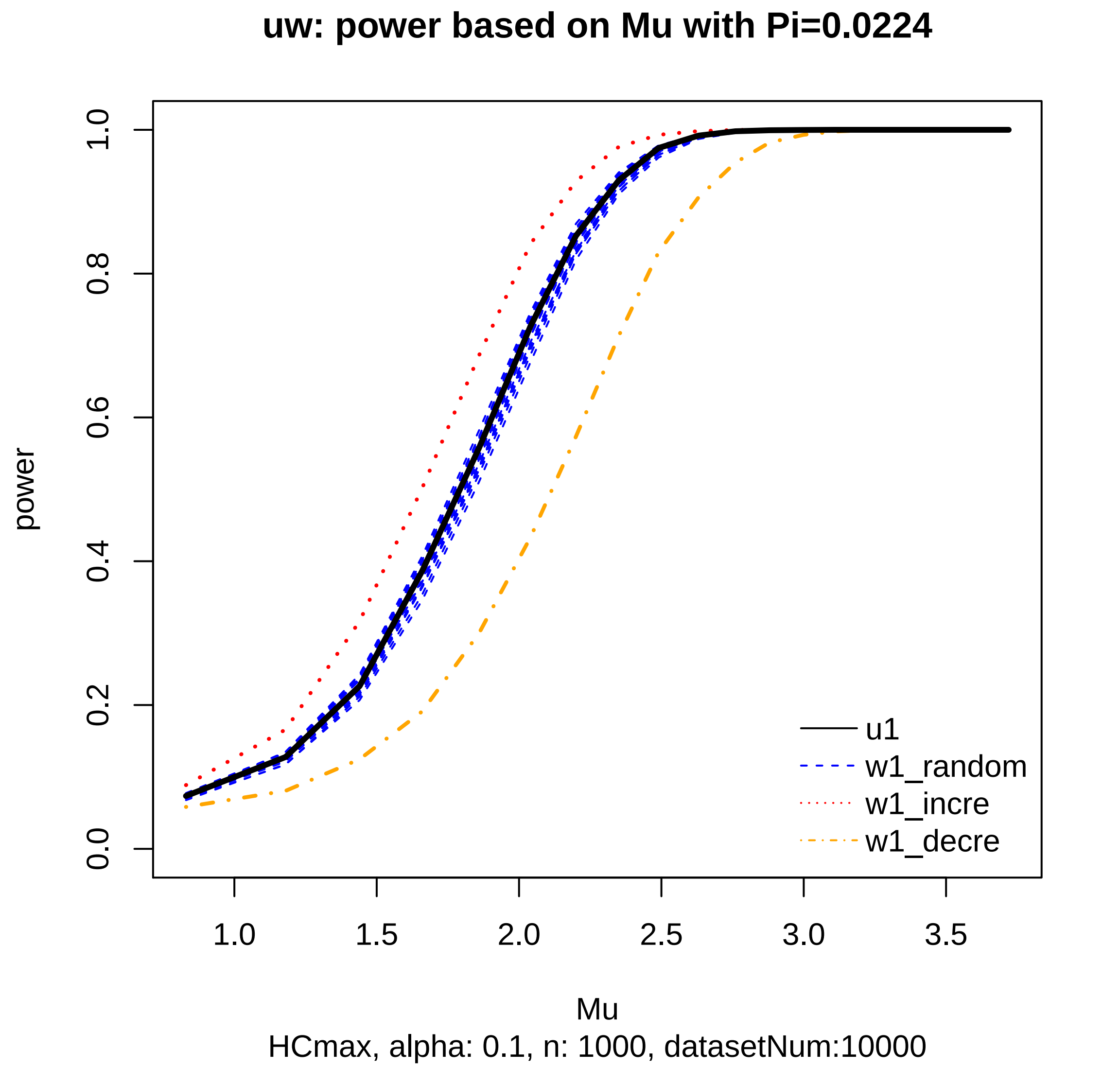
uw: power based on Mu with Pi=0.0224
power
0.0
0.2
0.4
0.6
0.8
1.0
1.0
1.5
2.0
2.5
3.0
3.5
Mu
HCmax, alpha: 0.1, n: 1000, datasetNum:10000
u1
w1_random
w1_incre
w1_decre

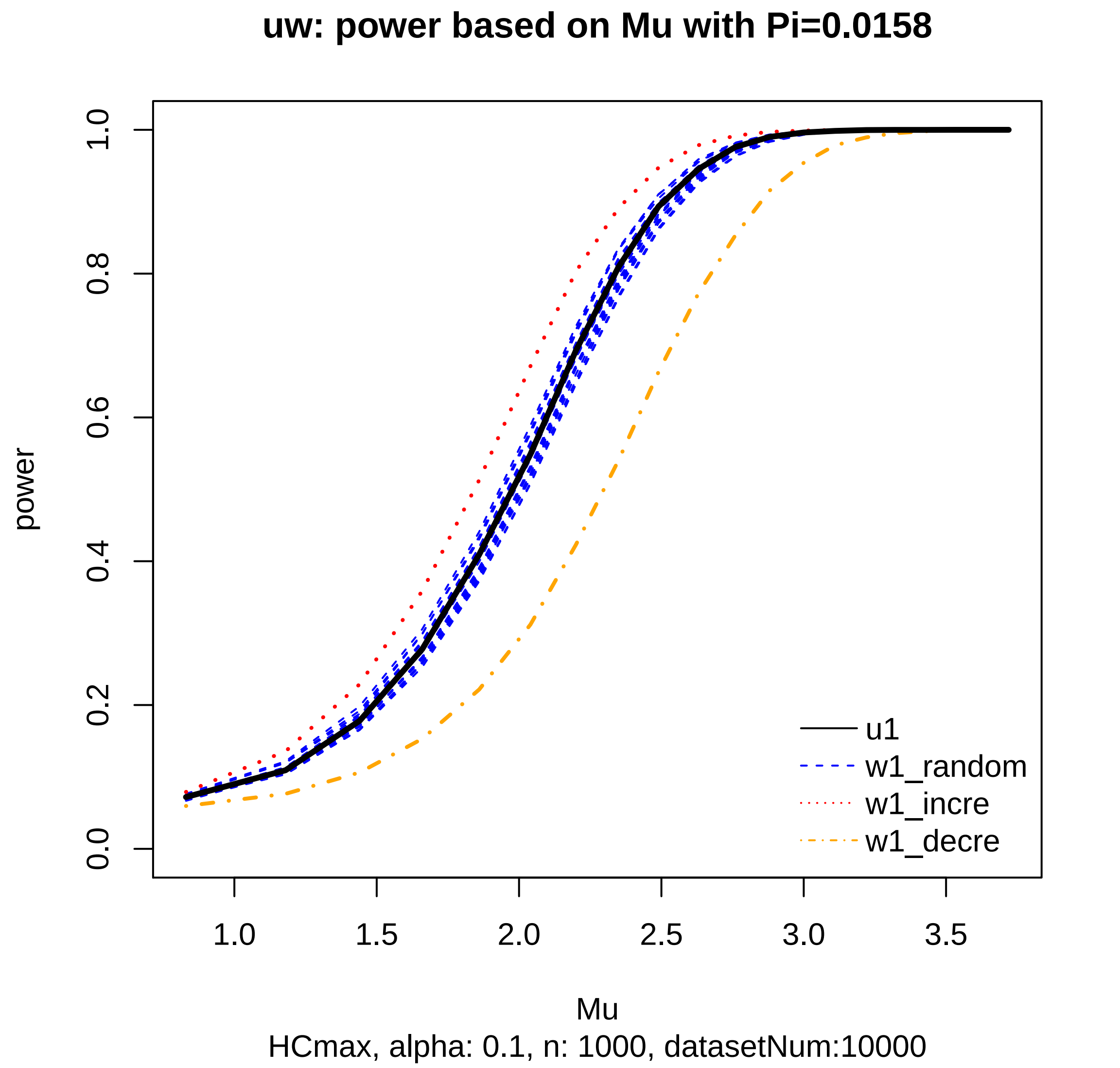
uw: power based on Mu with Pi=0.0158
power
0.0
0.2
0.4
0.6
0.8
1.0
1.0
1.5
2.0
2.5
3.0
3.5
Mu
u1
w1_random
w1_incre
w1_decre
HCmax, alpha: 0.1, n: 1000, datasetNum:10000

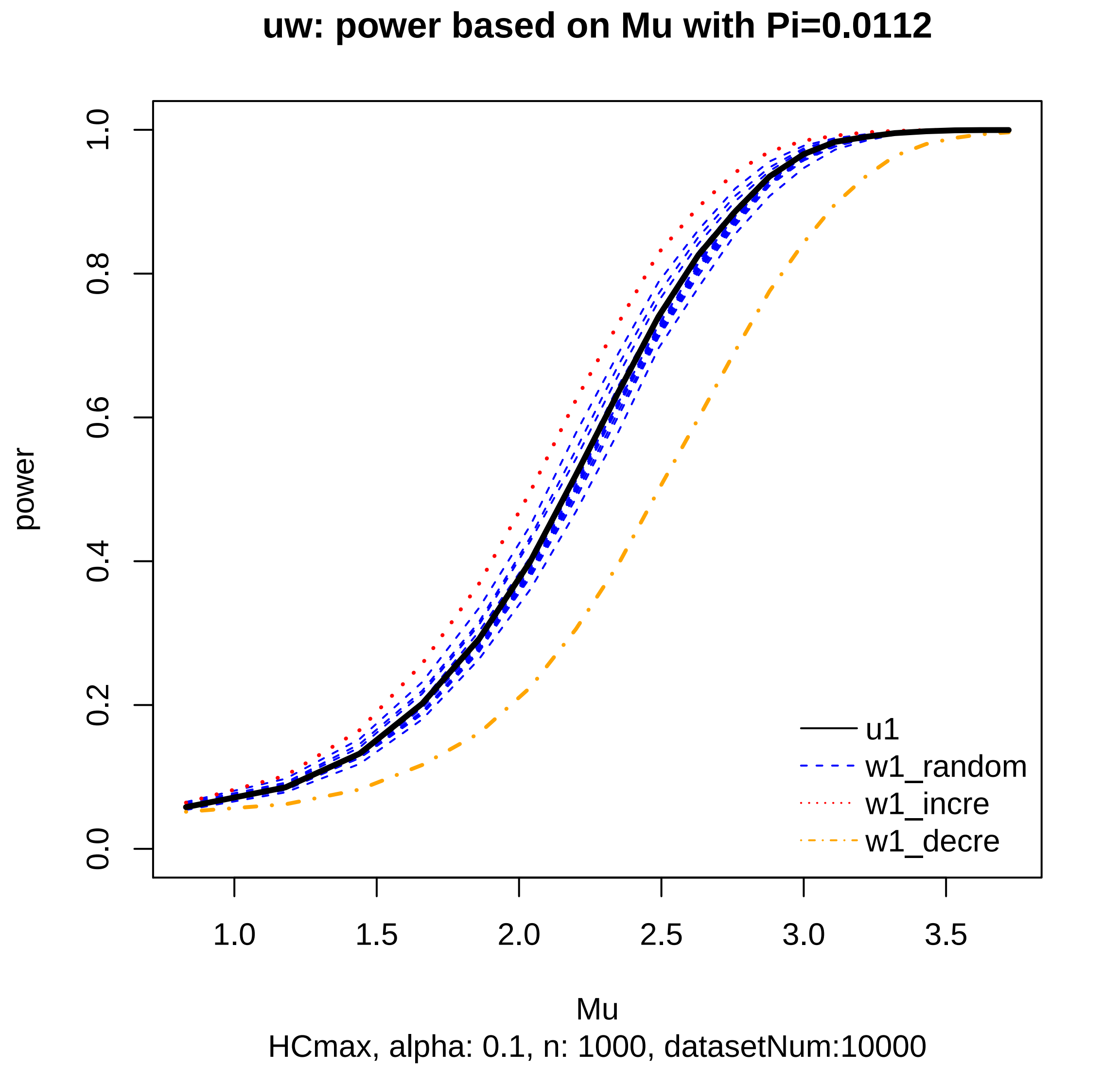

uw: power based on Mu with Pi=0.0112
power
0.0
0.2
0.4
0.6
0.8
1.0
1.0
1.5
2.0
2.5
3.0
3.5
Mu
HCmax, alpha: 0.1, n: 1000, datasetNum:10000
u1
w1_random
w1_incre
w1_decre

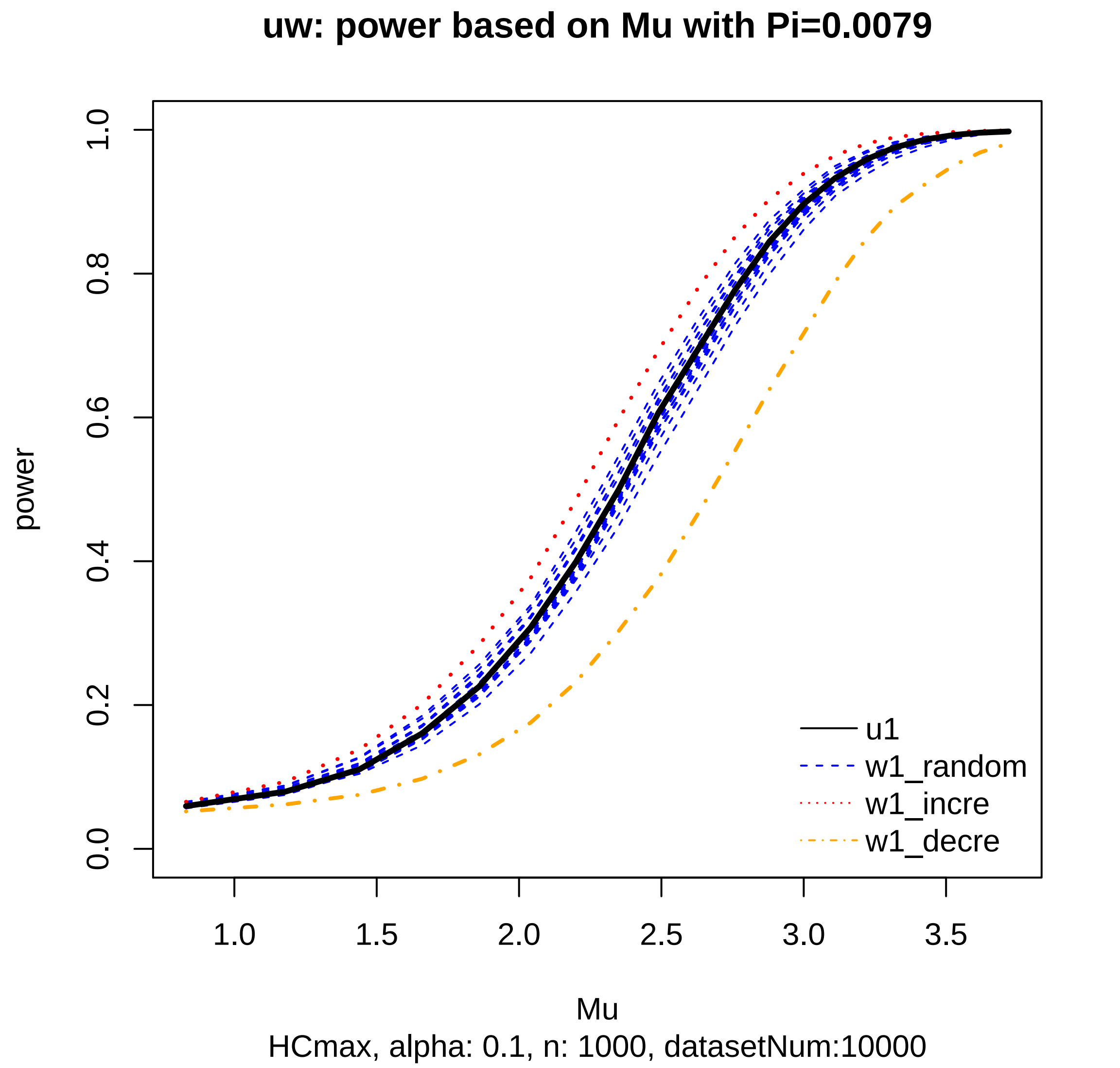
uw: power based on Mu with Pi=0.0079
power
0.0
0.2
0.4
0.6
0.8
1.0
1.0
1.5
2.0
2.5
3.0
3.5
Mu
u1
w1_random
w1_incre
w1_decre
HCmax, alpha: 0.1, n: 1000, datasetNum:10000

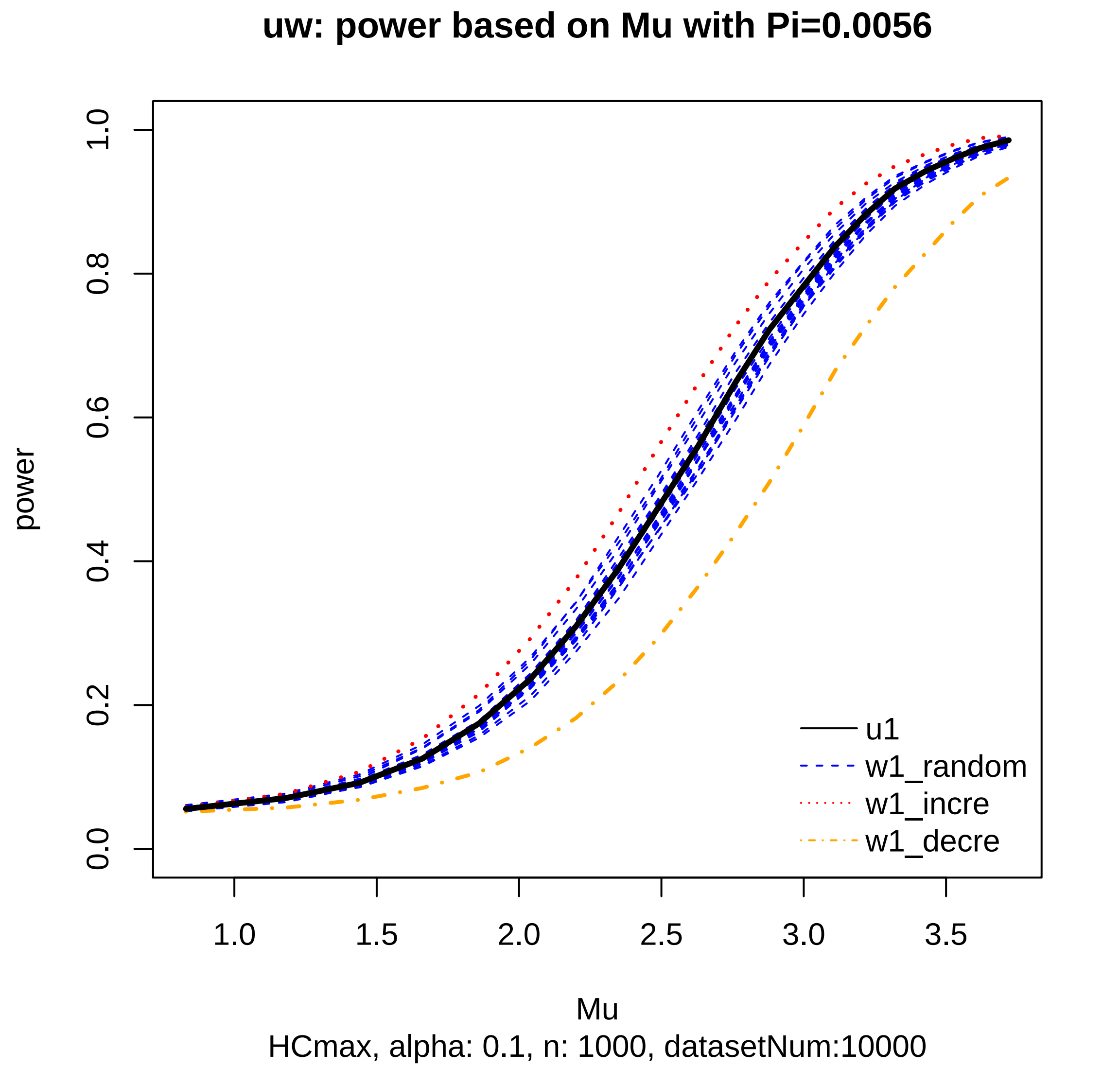
uw: power based on Mu with Pi=0.0056
power
0.0
0.2
0.4
0.6
0.8
1.0
1.0
1.5
2.0
2.5
3.0
3.5
Mu
HCmax, alpha: 0.1, n: 1000, datasetNum:10000
u1
w1_random
w1_incre
w1_decre

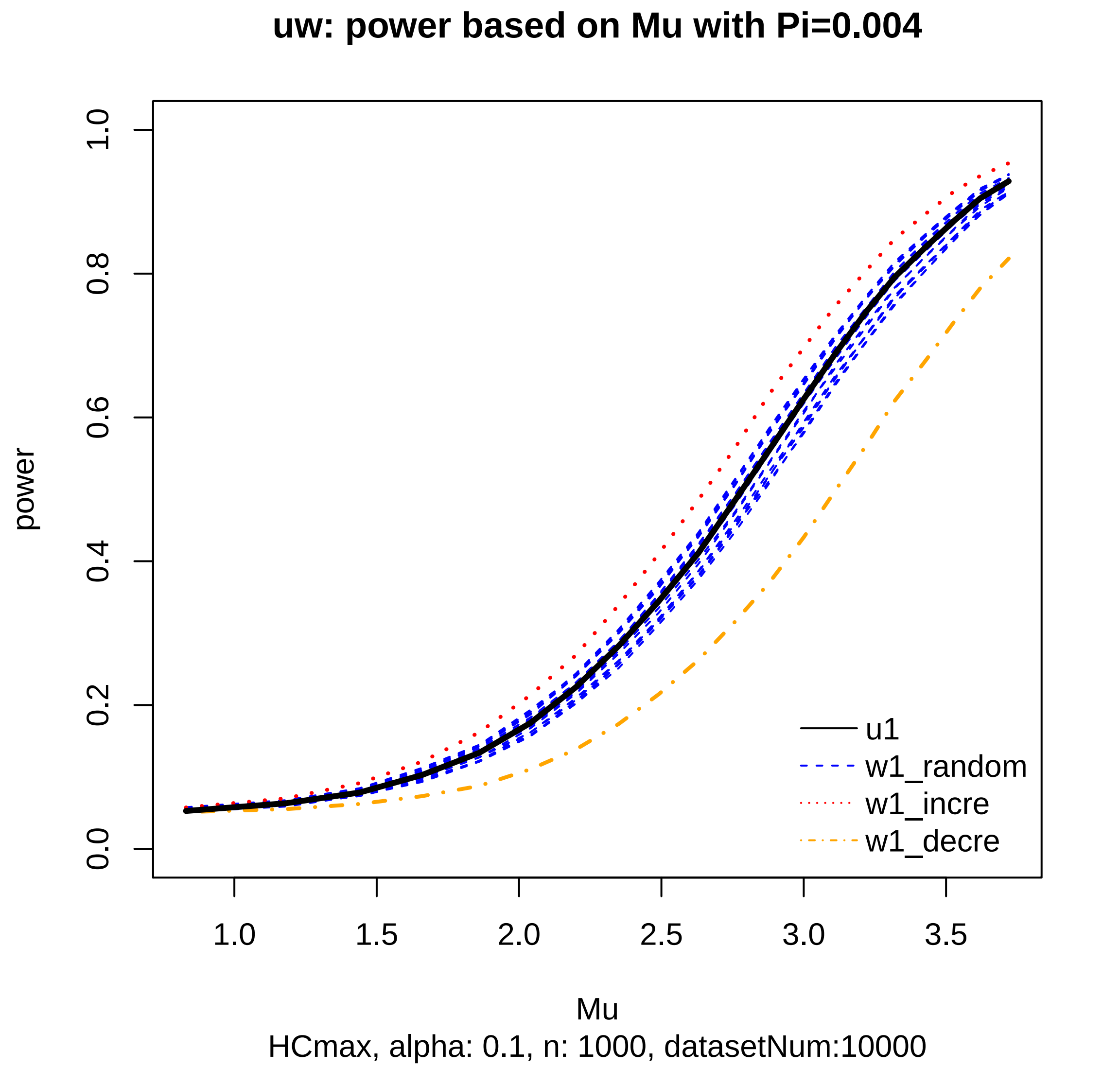
uw: power based on Mu with Pi=0.004
power
0.0
0.2
0.4
0.6
0.8
1.0
1.0
1.5
2.0
2.5
3.0
3.5
Mu
HCmax, alpha: 0.1, n: 1000, datasetNum:10000
u1
w1_random
w1_incre
w1_decre

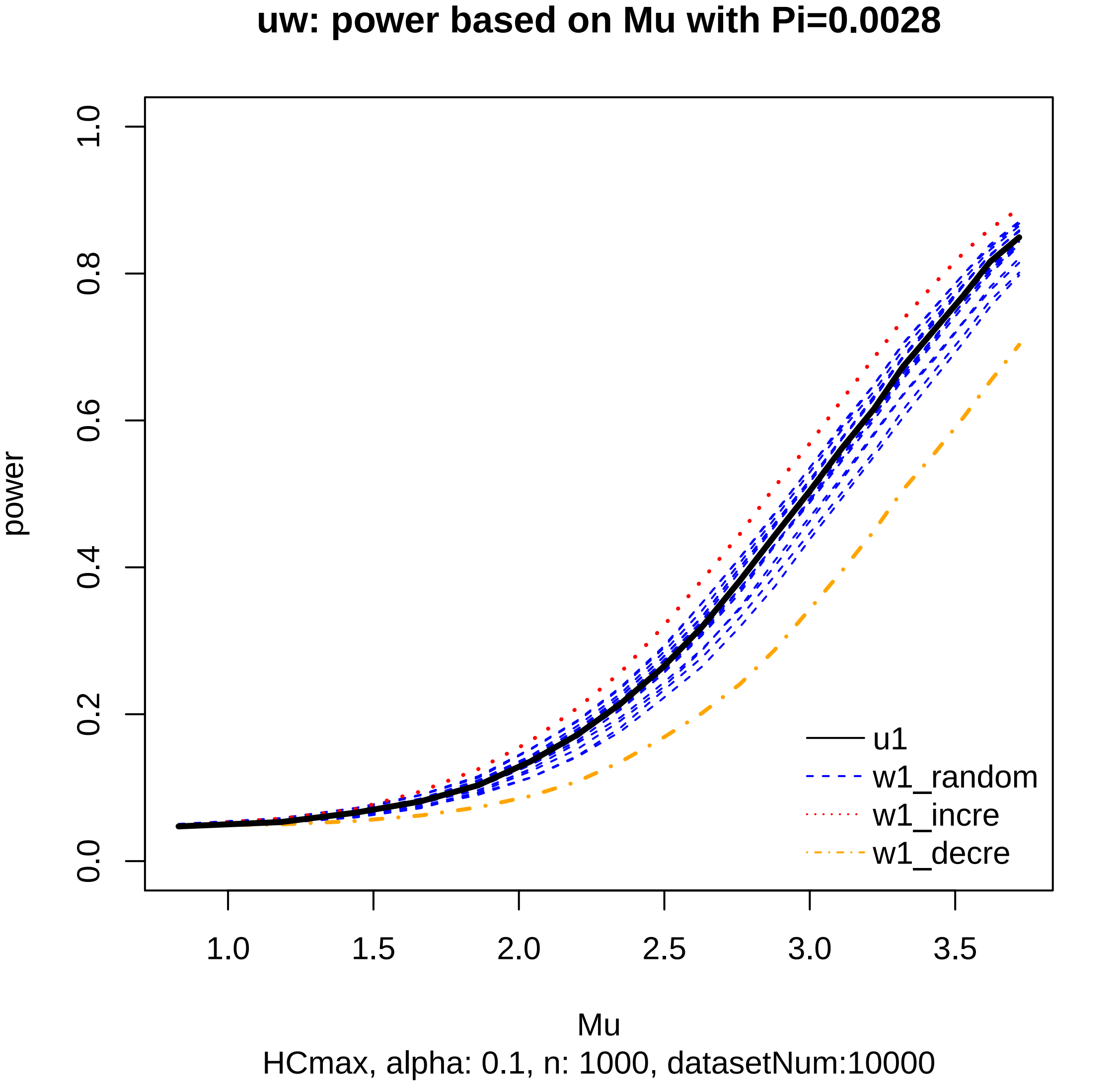

uw: power based on Mu with Pi=0.0028
power
0.0
0.2
0.4
0.6
0.8
1.0
1.0
1.5
2.0
2.5
3.0
3.5
Mu
HCmax, alpha: 0.1, n: 1000, datasetNum:10000
u1
w1_random
w1_incre
w1_decre

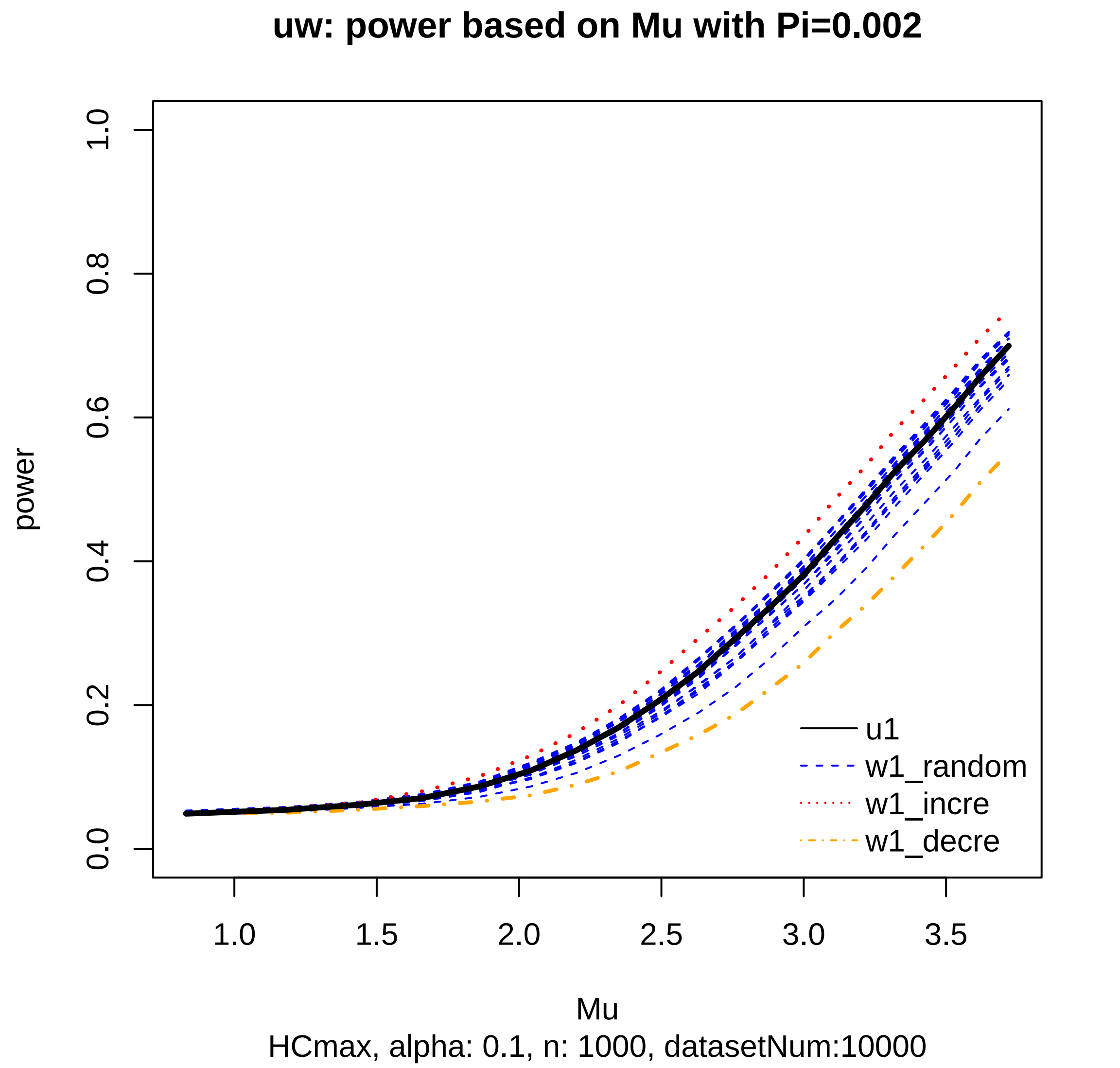
uw: power based on Mu with Pi=0.002
power
1.0
0.8
0.6
0.4
0.2
0.0
1.0
1.5
2.0
2.5
3.0
3.5
Mu
u1
w1_random
w1_incre
w1_decre
HCmax, alpha: 0.1, n: 1000, datasetNum:10000

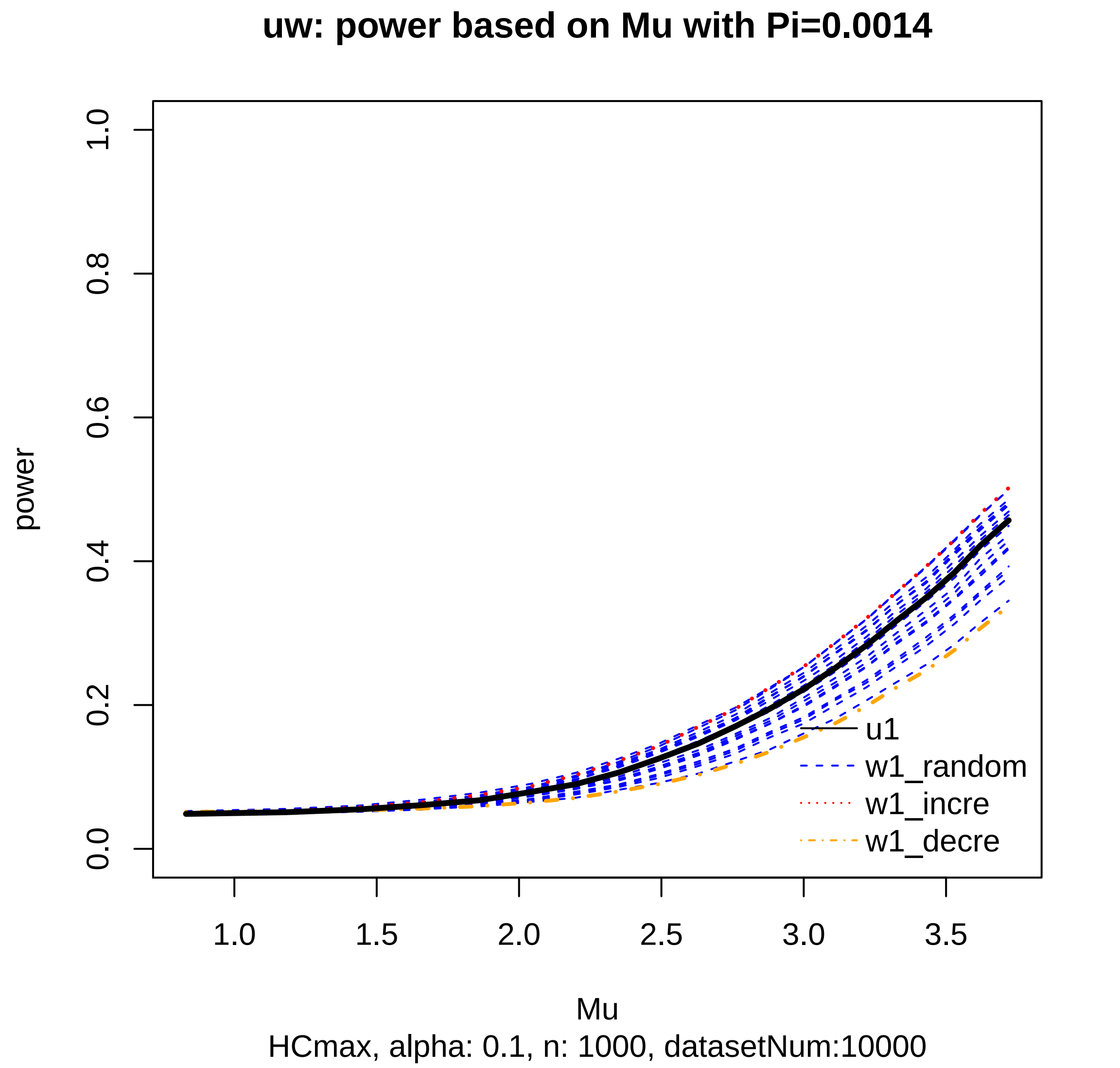
uw: power based on Mu with Pi=0.0014
power
0.0
0.2
0.4
0.6
0.8
1.0
1.0
1.5
2.0
2.5
3.0
3.5
Mu
HCmax, alpha: 0.1, n: 1000, datasetNum:10000
u1
w1_random
w1_incre
w1_decre

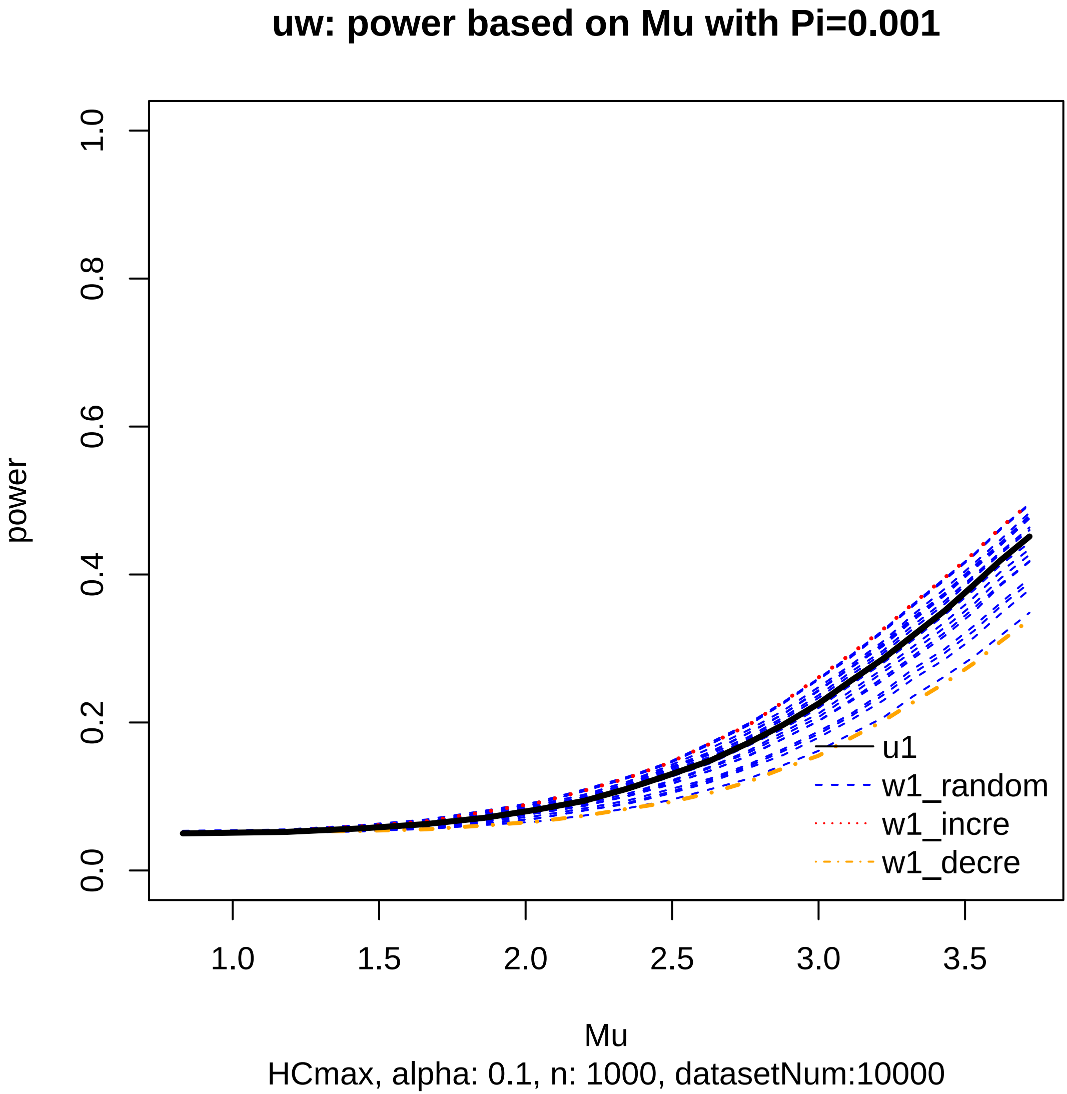

uw: power based on Mu with Pi=0.001
power
0.0
0.2
0.4
0.6
0.8
1.0
1.0
1.5
2.0
2.5
3.0
3.5
Mu
u1
w1_random
w1_incre
w1_decre
HCmax, alpha: 0.1, n: 1000, datasetNum:10000
